# Integrating suboptimal secondary structures, AI-assisted genomic synteny, and evolutionary conservation to identify bacterial ncRNA homologs beyond sequence similarity

**DOI:** 10.64898/2026.07.10.737749

**Authors:** Josef Pánek

## Abstract

A bioinformatic approach for genome-wide identification of homologs of bacterial non-coding RNAs (ncRNAs) integrating structural similarity, genomic synteny, and evolutionary conservation is presented.

The structural similarity is detected using an algorithm for genome-wide identification of loci in genomic intergenic regions (IGRs) containing sequences capable of adopting secondary structures similar to that of the query ncRNA. The algorithm scans IGR sequences using a sliding window with a predefined step. For each window, suboptimal secondary structures are predicted and compared with the template structure to compute structural similarity scores. These scores are evaluated statistically on a genome-wide scale to infer homology of the RNAs represented by the predicted structures.

Loci encoding statistically significant structures are further filtered using genomic synteny of the query ncRNAs inferred from genomic annotations. ChatGPT was used to assist in identifying literature-supported biological relationships between genes with distinct functional annotations. Syntenic loci with the structures are then examined for homologs in related species, as evolutionary conservation among related species is a common feature of ncRNAs

Using this approach, we predicted novel homologs of the spot42 RNA-encoding spf gene in *Glaciecola* and *Pseudoalteromonas* genomes, and ms1 RNA genes in *Frankia* and *Bifidobacterium* genomes, where previous homology searches had failed.

**GRAPHICAL ABSTRACT:** 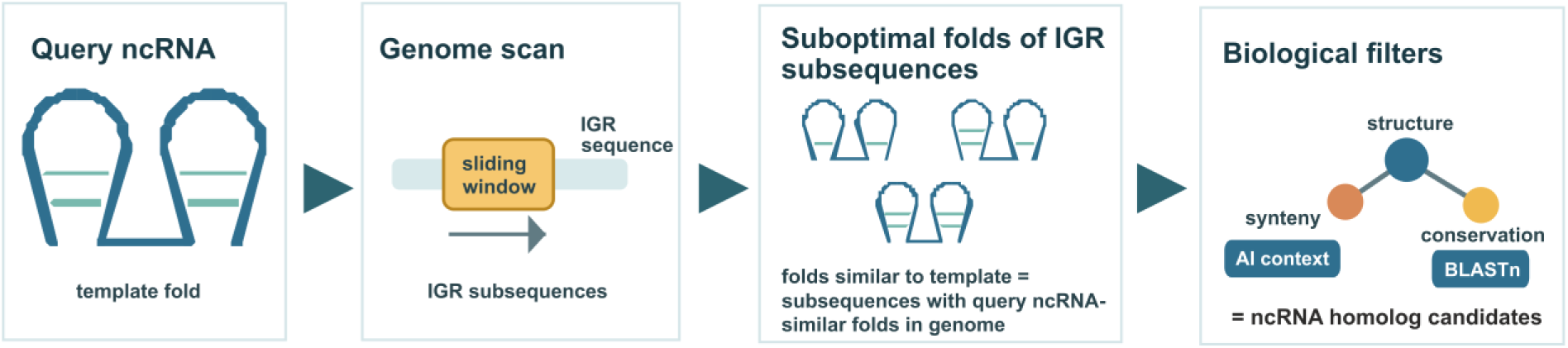

## INTRODUCTION

Bacterial genomes encode a wide variety of non-coding RNAs (ncRNAs) that participate in many cellular processes, ranging from gene expression to cell wall biogenesis and maintenance. Although some ncRNAs act through base pairing with target RNAs, their functions are often determined not only by primary sequence but also by conserved structural features (1). As a result, conventional sequence-based homology searches are often insufficient, particularly when ncRNAs are short, rapidly evolving, conserved mainly at the structural level, or when homologs are sought in evolutionarily distant organisms, where sequence similarity may no longer be detectable. This limitation has motivated the development of computational approaches that incorporate RNA secondary structure, comparative conservation, and probabilistic models of RNA families (2-4).

RNA secondary structure is a key feature that can be conserved among homologous ncRNAs even when their nucleotide sequences have diverged. As a result, secondary structures of ncRNAs can be used as a tool for identification of ncRNA homologs rather than primary sequences. Still the use of secondary structures in homology searches is limited by the inaccuracy of the RNA structure prediction and difficulties in experimental identification of the RNA structures.

The prediction mostly relies on minimum free energy (MFE) structures as a proxy for the native structure. However, a single MFE structure may not adequately describe the structural behavior of an RNA molecule, because RNA sequences can populate ensembles of alternative conformations and energetically close structures (5); (6). Suboptimal structures—those with higher free energies (FEs) than the MFE structure—can therefore provide useful information about structurally plausible states, structural diversity, and alternative conformations, particularly for RNAs that undergo functional structural rearrangements or populate multiple folds (7) (8). The reason is that native conformations of RNA molecules are formed under conditions in the cells that correspond to higher FEs (9,10). Recent reviews of RNA structure and RNA–RNA interaction prediction also emphasize the importance of ensemble-based and suboptimal-structure approaches rather than relying exclusively on a single optimal fold (11).

Recent studies have emphasized the importance of incorporating sets of suboptimal structures to improve the prediction and interpretation of RNA conformations (12) (13) (14) (15). Rather than relying on a single predicted structure, analyzing a broader structural ensemble allows for a more biologically reliably prediction of RNA folding and function. This ensemble-based perspective can be particularly valuable when experimental data suggest a structure that deviates significantly from the MFE prediction. The use of suboptimal secondary structures still remains limited due to the computational and evaluation challenges related to them (16) (17) (18) (19) (8) (20).

The structure-based genome-wide searches can produce false-positive candidates, because even low false-positive rates can generate many spurious predictions when large genomic search spaces are scanned (21). Intergenic regions may contain local sequences that form structures resembling a query RNA by chance. For this reason, structural evidence should be interpreted together with independent biological signals. Genomic synteny provides one such signal based on the conservation of the genomic neighborhood surrounding ncRNA genes. In bacterial ncRNAs discovery, conserved flanking genes, gene synteny, and genomic backbone retention have been used to identify candidate ncRNA homologs across related genomes (22). More generally, bacterial gene order is not random, and conserved synteny reflects functional and evolutionary constraints on genome organization (23). Thus, syntenic context can help distinguish true homologous loci from unrelated regions producing structurally similar folds.

Evolutionary conservation among closely related species provides a further criterion for ncRNA gene identification. Functional ncRNAs are expected to be maintained over evolutionary time, particularly among related genomes in which homologous loci can often still be recognized. The presence of structurally similar candidates in corresponding genomic contexts across multiple related species supports the interpretation that these loci encode conserved ncRNA homologs rather than isolated structural artifacts. Thus, the integration of structural similarity, genomic synteny, and evolutionary conservation offers a more reliable framework than any single criterion alone.

In this work, we present an integrative bioinformatics approach for genome-wide identification of bacterial ncRNA homologs. The method searches genomic intergenic regions using a sliding-window procedure and evaluates whether each window can adopt secondary structures similar to a query ncRNA. For each genomic window, suboptimal secondary structures are predicted and compared with the template structure of the query ncRNA, allowing the method to capture structural similarity beyond a single optimal folding solution. Candidate loci with statistically significant structural similarity are then filtered using genomic synteny inferred from genome annotations in genomic contexts of the query ncRNA. Finally, syntenic candidates are examined across closely related species to assess evolutionary conservation.

By combining these complementary sources of evidence, the proposed approach is designed to improve the detection of ncRNA homologs that are missed by conventional sequence-based searches. The integration of suboptimal structure analysis with genomic context and comparative conservation provides a framework for identifying structurally conserved ncRNA genes in bacterial genomes, including cases where sequence similarity is weak or absent. We demonstrate the utility of this strategy by predicting novel homologs of the spot42 RNA-encoding *spf* gene in *Glaciecola* and *Pseudoalteromonas* genomes, as well as homologs of ms1 RNA genes in *Frankia* and *Bifidobacterium* genomes, where previous homology searches had failed.

## MATERIALS AND METHODS

### Genome–wide structure search algorithm

The algorithm for genome–wide structure search picks up the loci in subject sequences with structure similarity to structure templates. The subject sequences can be any nucleotide sequence. The templates are RNA secondary structures. The approach is demonstrated here with identification of homologs of bacterial regulatory non-coding RNAs (ncRNAs), therefore the templates are experimentally determined secondary structures of selected ncRNAs whose homologs were searched for. These structures were obtained from the Protein Data Bank (PDB) and from published scientific literature. The subject sequences were genomic intergenic regions (IGRs) where ncRNA genes reside. The genomes were obtained from public databases, primarily NCBI GenBank. IGRs in which homologs of the templates are searched are extracted from genomes using coordinates of the genes annotated in the genomes. Templates and genomes used for testing the algorithm are listed in Supplementary Tables S1.

The algorithm is based on a scanning approach via a sliding window using templates. The sliding window moves along IGR sequences in fixed steps of a predefined number of nucleotides, generating subsequences of length equal to the window size. The step size was set to 5 nucleotides as a compromise between the speed and accuracy of the algorithm. It was a maximal possible step for not to miss subsequences in IGRs with the global free energy (FE) minimum. The window length corresponds to the length of the query ncRNA, but also may be set manually when the expected length of the identified homolog differs from that of the template.

For each subsequence, suboptimal secondary structures is predicted. For this purpose, UNAfold [29] was used. The prediction generates ensembles of suboptimal secondary structures that sample the folding space of each subsequence. The ensembles enabled analysis of folding trends rather than relying on a single structure, such as the commonly used minimum free energy (MFE) structure, which provides only limited structural information.

The similarity between predicted suboptimal secondary structures and the template were computed. All suboptimal secondary structures of individual subsequences were compared to the template. For this comparison, RNAdistance [30] was used to calculate tree edit distances (TEDs), where lower values indicated higher similarity. TEDs were then used to compute scores for all subsequences across IGRs in the searched genomes. Subsequences in IGRs with the best scores were selected as candidate loci most likely to encode the sought homologs.

Multiple suboptimal secondary structures predicted for single subsequences made it possible to define multiple scores that grasped different aspects of structural similarity. The scores were tested using the test searches. In this project, five score definitions were proposed as follows:

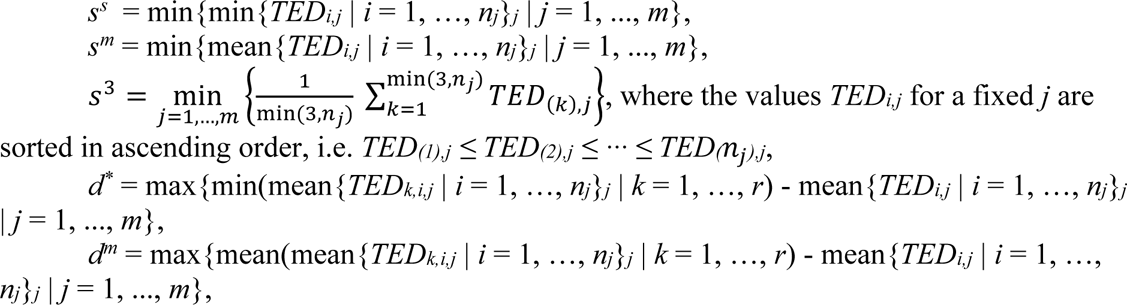

where *n_j_*_=1,…,*m*_, are numbers of suboptimal secondary structures for *j*th of the total *m* subsequences of a subject sequence, *TED_i,j_* is a tree edit distance between the *i*th suboptimal secondary structure of the *j*th subsequence and the template, and *TED_k,i,j_* is a tree edit distance between the *i*th suboptimal secondary structure of the *k*th version with randomly shuffled dinucleotides of the *j*th subsequence and the template. *m* is given by the size of the sliding window, the size of the sliding step, and the length the subject sequence.

Candidates to homologs in IGRs in an inspected genome were obtained, with one candidate produced per IGR, i.e. 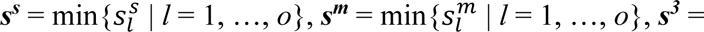 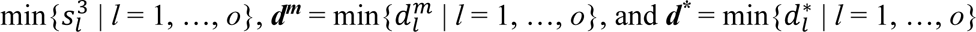, where *o* is a number of subject sequences and ***d^m^***, ***d\****, ***s^s^***, ***s^m^***, ***s^3^*** are vectors.

The biologically reliable candidates were then selected from these sets. To achieve this, the individual scores of candidates were compared using a two-sample t-test against the scores of all other candidates identified within the same genome. This testing produced two sets of p-values corresponding to the scores, as follows:

*p_l_* = t-test(*sc_l_*, *sc_1…l-1,l+1…o_*), where *p_l_* was p-value of *l*th subject sequence, *l* = 1,…,*o*, where *o* is the number of subject sequences, and *sc_l_* was one of the score vectors defined above. Note that *o* was also the number of IGRs in searched genomes, as one IGR had one sequence with candidate with secondary structure most similar to the template. The probability level of the test was set to α = 0.05.

The scores were tested for their efficiency using the test searches as described in the Results section.

### Test searches

Test searches were proceeded using known bacterial non-coding RNAs (ncRNAs) with experimentally determined secondary structures. Representative ncRNAs were selected from Rfam 15.1 (24) and from the literature (Supplementary Table S1, sequences and structures are provided in Supplementary File S1.fa). Secondary structures were obtained from PDB entries (25) or from published sources. Riboswitches were excluded because they are parts of larger structural complexes and are not functional as standalone RNAs.

The presented genome–wide structure search algorithm searched for known homologs of the query ncRNAs. Multiple homologs were used for a single query ncRNA from differently evolutionarily distant bacterial species to test the evolutionary range the algorithm is able to grasp. Also homologs with no detectable sequence similarity were included to find out if the search is able to make such an evolutionary divergence (Supplementary Table S2).

Fifteen ncRNA families were collected, each containing at least one experimentally defined structure and multiple known homologs (Supplementary Table S2). From these, 48 predictions were assembled (Table 1), each searching for homologs of a query ncRNA in diverse bacterial genomes. Both small and large RNAs were included to assess size dependence of the search.

**Table 1.**
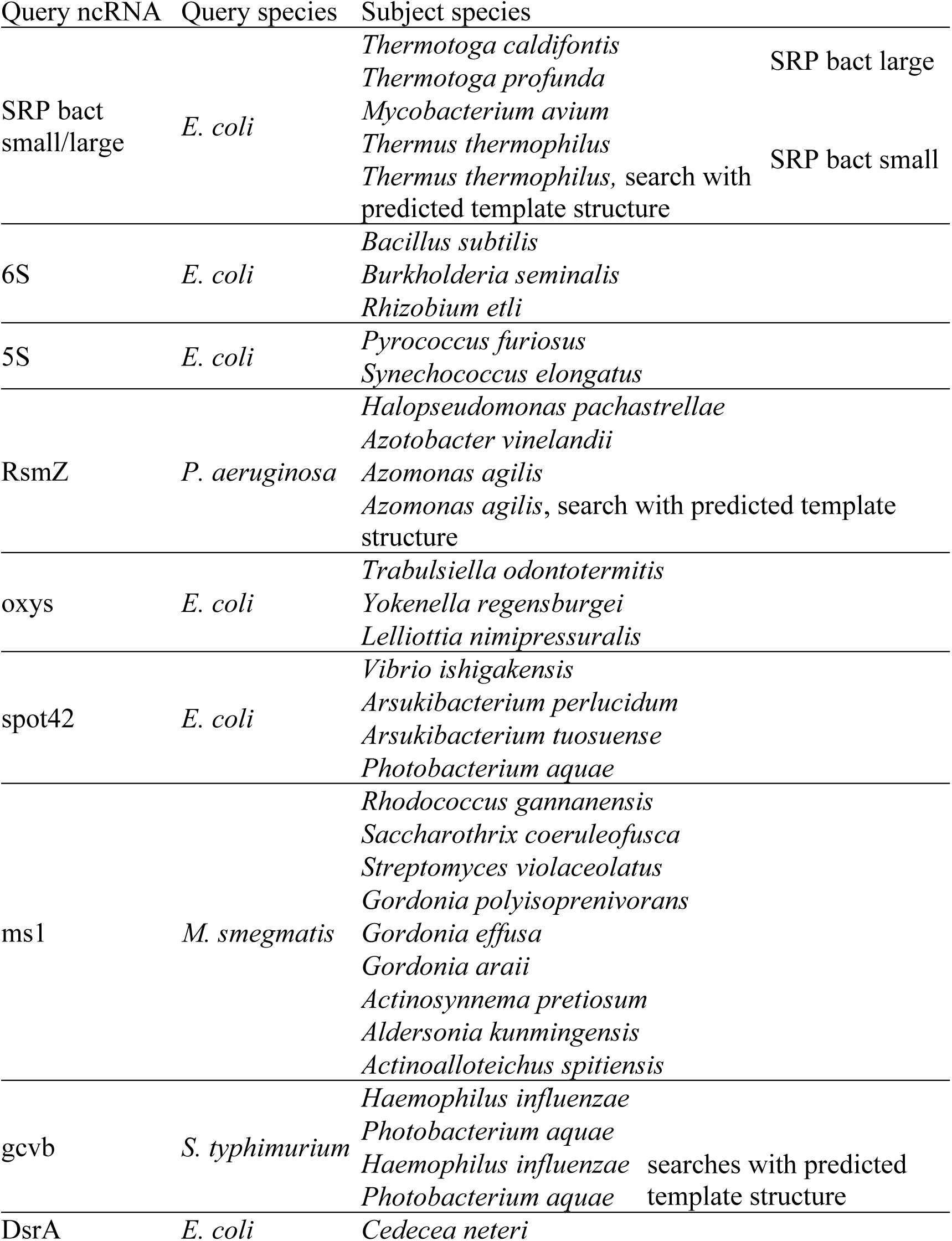
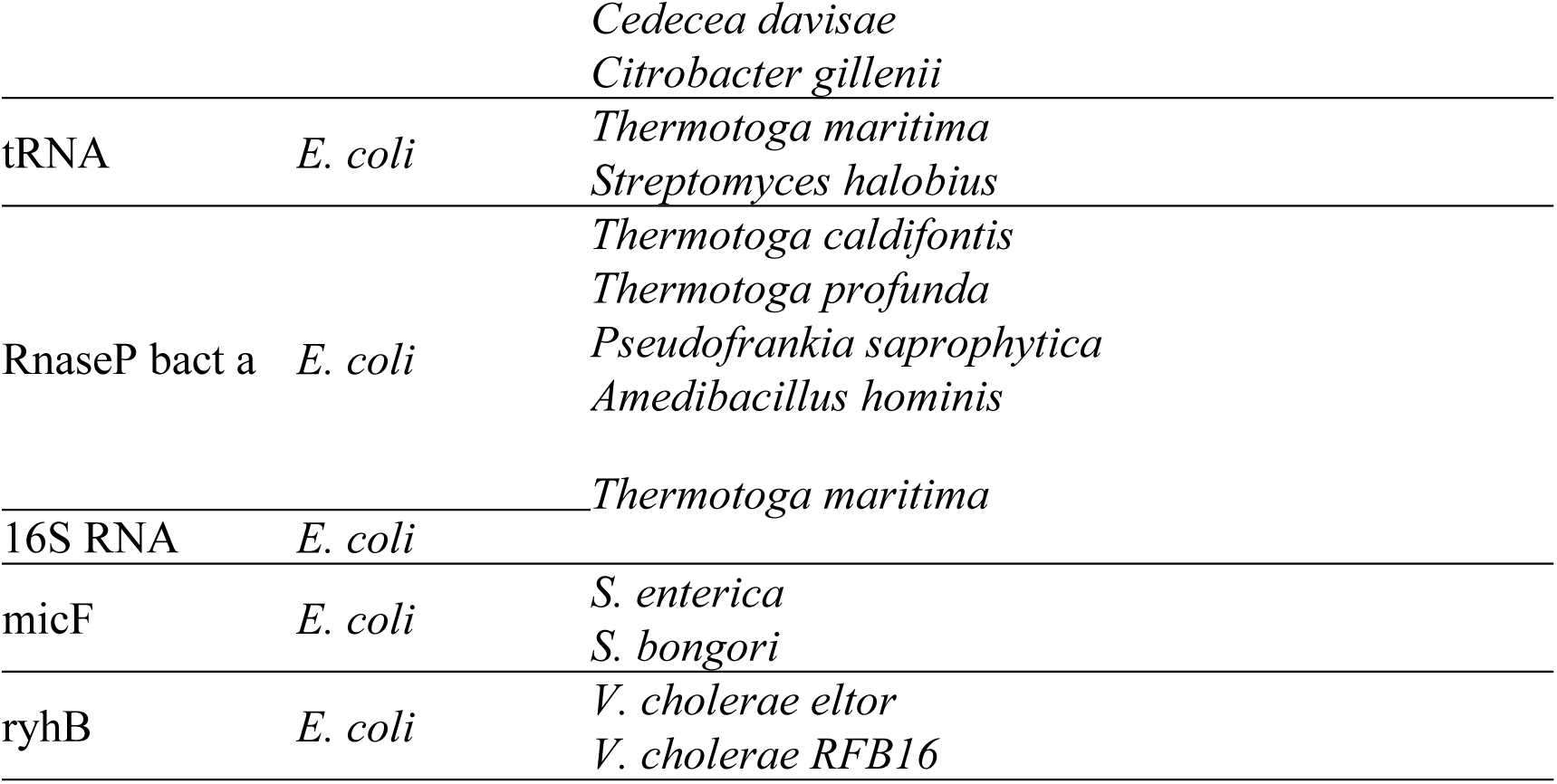
Summary of test searches. For details see Supplementary Table S2.

### Synteny-based filtering

The aim of the synteny-based filtering was to identify the genomic contexts of the structure candidates biologically/functionally consistent with the synteny of the query ncRNAs. The contexts were successions of genomic annotations of up to 7 genes located upstream and downstream of the IGRs containing the candidates. The annotations were obtained from NCBI feature table files. The filtering was accomplished using synteny phrases inferred from the genomic contexts of known homologs of the query ncRNAs were inferred and identified in genomic contexts of structure candidates.

Synteny phrases were text phrases consisting of most repeating words in genomic annotation of genes flanking multiple homologs of query ncRNAs. The phrases did not contain nonspecific words that were typically these: ‘hypothetical’, ‘protein’, ‘type’, ‘family’, ‘domain’, ‘putative’, ‘precursor’, ‘component’, ‘subunit’, ‘subfamily’, ‘conserved’, ‘chain’, ‘or’, ‘superfamily’, ‘unknown’, ‘ribosomal’, ‘function’, ‘of’, ‘conserved’, ‘like’, ‘containing’, ‘II’, ‘and’, ‘short’, ‘dependent’, ‘probable’, ‘associated’, ‘to’, ‘that’, ‘the’, ‘predicted’, ‘uncharacterized’, ‘production’, ‘fold’, ‘in’, ‘by’, ‘universal’, ‘pathway’, ‘involved’, ‘related’, ‘binding’, ‘general’, ‘group’, ‘sequence’, ‘class’, ‘cluster’, ‘accepting’, ‘determining’.

These repeating words were also specific, i.e. they characterized specific biological aspects of gene function. E.g. the ‘ribosome biogenesis GTP-binding protein YihA/YsxC’ annotation contained two nonspecific words, ‘binding’ and ‘protein’ and specific words ‘ribosome’, ‘biogenesis’, ‘GTP’, ‘YihA’ and ‘YsxC’.

Synteny phrases were generated from the most frequent specific words according to their co-occurrence in the annotations from which they were extracted and which defined their semantic binding. For example, the specific words shown above formed the following phrases: ‘ribosome biogenesis GTP’, ‘YihA’ and ‘YsxC’. The phrases were typically two, for downstream and upstream parts of genomic contexts. Both the construction and the use of synteny phrases are demonstrated in the Use cases subsection in the Results section.

Synteny phrases were identified among seven consecutive genes located both downstream and upstream of the IGRs containing statistically significant structural candidates. Seven genes were analyzed to cover a sufficiently long genomic range, allowing the identification of genes that were part of the original syntenic region of the subject ncRNAs, since insertions may have occurred between these genes. The presence of all words from any substring of a synteny phrase, in any order, within the annotations of a genomic context was considered evidence of synteny for the query ncRNA. If either the downstream or upstream synteny phrase was detected on one side of a genomic context, the corresponding phrase was then searched for on the opposite side, accounting for the two possible genomic orientations. In some cases, only one part of the synteny could be detected, as the syntenic region was only partially kept during evolution.

The procedures required for the presented approach were coded in Matlab computational environment. The Matlab scripts are included in Supplementary File S11.

## RESULTS

A bioinformatic approach for genome-wide identification of homologs of non-coding RNAs (ncRNAs) that integrates structure similarity, genomic synteny, and evolutionary conservation is presented. It consists of three steps: (i) genome–wide structure search, (ii) synteny-based filtering, and (iii) evolutionarily conservation-based filtering. The overall scheme of the approach is shown in Figure 1. For further details, see the Materials and Methods section.

**Figure 1.**
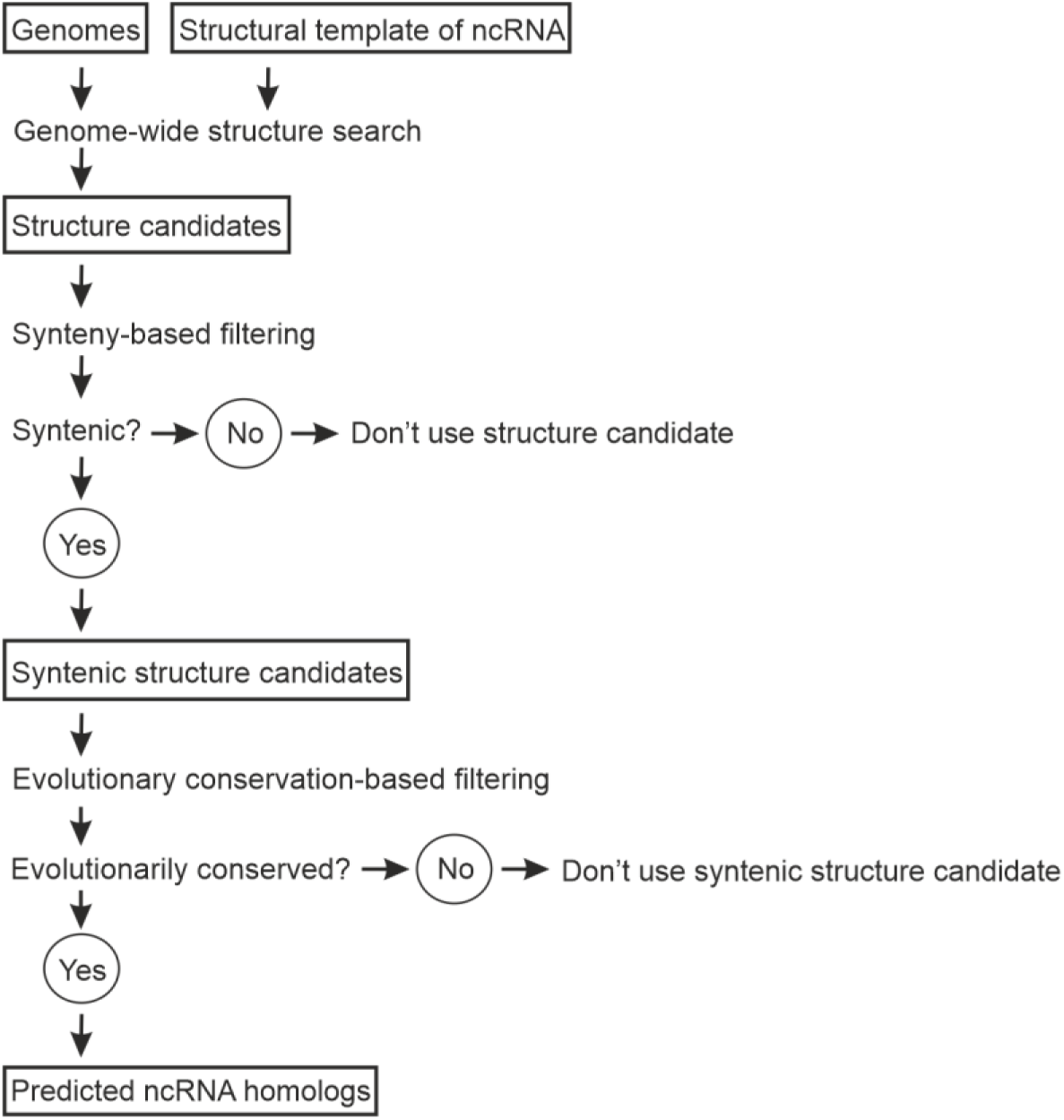
Scheme of workflow of the presented approach.

### (i) Genome-wide structure search

The search is implemented by an algorithm for identifying loci in biological genomes that exhibit structure homology of biological molecules. The structure homology is indicative of the presence of genes of evolutionarily related molecules with the same function in genomes of different biological species called homologs. Such molecules are kept during the evolution because their function is necessary for the species. To get biological relevance to encode for the homologs, the statistical test for the identified loci was developed.

The molecules used in this project for both testing and demonstration of the presented approach, were bacterial regulatory non-coding RNAs (ncRNAs). Two homologs of each ncRNA described in literature and/or included in RNA databases were used, one as a query and one as a target identified using the query. The genomes in which the target homologs were identified were obtained from public databases, primarily NCBI GenBank. The algorithm is described in details in the Genome–wide structure search algorithm subsection of the Materials and Methods section.

First, the algorithm was tested and validated using 48 test searches (Table 1, Supplementary Table S1, S2). Details on the test set are in the Test searches subsection of the Materials and Methods section. Of the 48 identified homologs, 43, 42, 41, 40 and 34 were identified as statistically significant by *s^m^*, *s^3^*, *s^s^*, *d^m^* and *d\** scores, respectively. The effectiveness of the scores for the test searches are shown in Supplementary Figure S1.

Where it was applicable, predicted secondary structures were used instead of experimentally identified structures (Supplementary Table S2) and the test showed no difference between searches using either secondary structures, showing an applicability of the algorithm also with predicted secondary structures as templates. This part of the test was limited as only two ncRNAs with known homologs with substantially differing lengths were found.

The algorithm was also tested with different lengths of sliding window. This was possible for two query ncRNAs, 6S and ms1 RNAs. For these RNAs, experimentally identified homologs with their lengths differing substantially from those of the query were available. The lengths of query RNAs were 184 and 304 nucleotides for 6S and ms1 RNAs. The sequence lengths of their homologs in *Rhizobium etli* and several *Actinomycetes* identified in the test were ∼160 and 227 nucleotides for 6S and ms1 RNAs. The test is summarized in Supplementary Figure S2 and demonstrated that the algorithm is highly length-responsive. This property adds an informative diagnostic capabilities to its uses making it applicable to identification of boundaries and size of the predicted RNAs.

Besides the sought homologs, that were true positives (TPs), that were single in the testing searches, also other candidates were identified as statistically significant that were considered false positives (FPs). Across 48 test searches on the genomes, the average number of false positives (FPs) and their proportion relative to IGRs for the five scores were 75 (3.8%), 56 (2.8%), 56 (2.9%), 84 (4.1%), and 84 (4.1%), respectively. Different percentages for the same FP numbers were caused by different numbers of IGRs from different genomes, because FPs may have been identified in different genomes using different scores.

The test showed that the numbers of FPs were still too high for unambiguous homolog identification. Still the search narrowed an occurrence of the test homologs in test genomes from typically thousands of IGRs per genome to tens, representing thus significant increase in precision. For identification TPs among these statistically significant candidates a synteny based specification was used.

### (ii) Synteny-based filtering

Synteny of ncRNAs was used to identify TPs among statistically significant structure candidates. TPs were homologous candidates that were expected to share at least partly syntenic genomic contexts. The syntenic agreement was identified by searching for characteristic phrases, so called synteny phrases, derived from the genomic contexts of the query ncRNA and their homologs. Synteny phrase construction is described in the Synteny-based filtering subsection of the Materials and Methods section.

Using synteny, the strategy was to identify TPs within the sets of statistically significant structure candidates, each consisting of multiple FPs and a single TP, as determined by individual structure search scores. The sets were examined sequentially, considering only genomes in which no TP had been identified in previous sets (except for the first set). The sets were ordered by decreasing performance of the scores in test searches, i.e. *s^m^*, *s^3^*, *s^s^*, *d^m^* and *d\**. This was done in the following manner: select score *i;* select searched genome *j*; if there is not a syntenic candidate in genome *j* found by previous scores; identified by score *i;* genome *j* by score *i*, if any; select statistically significant structure candidates in genome *j* do the synteny-based filtering of the selected candidates; save syntenic candidates among the selected candidates for next genome j + 1; next score i + 1.

The syntenic candidates were further inspected for evolutionary conservation as described in the following section.

### (iii) Evolutionary conservation-based filtering

The syntenic candidates were checked for their evolutionary conservation. They were considered evolutionarily conserved when there existed similar sequences to them in evolutionarily closely related species in IGRs with syntenic genomic contexts.

Similar sequences were identified by BLAST of the sequences of the candidates in NCBÍs nt_prok and refseq_genomes databases with sensitive setting for cross-species exploration (with parameters -r 1 -q 1 -G 1 -E 2 -W 7 (26) with BLAST E-value threshold set to 0.05).

IGRs with similar sequences were checked for synteny by searching the genomic annotations of their surrounding regions for synteny-related phrases, using the same approach as for the structural candidates. These IGRs contained putative homologs of the query ncRNAs, inferred from the statistically significant sequence similarity to syntenic structural candidates that were also statistically significant. Therefore, these IGRs were also considered to contain putative homologs of the query ncRNAs.

### Use cases

The presented approach is demonstrated using case studies focused on identifying new ncRNA homologs. These represent challenging ncRNA homolog identification problems in which standard sequence/structure- and synteny-based methods had failed or yielded uncertain results. The cases include searches for genes of spot 42 RNA homologs in *Glaciecola* and *Pseudoalteromonas*, and genes of ms1 RNA homologs in *Frankia* and *Bifidobacterium* (27) Vankova Hausnerova, 2022 #683}. None of these homologs were included in 15.1 (24).

The newly predicted homologs were found to be located in intergenic regions with conserved but modified syntenic contexts, likely shaped by evolutionary rearrangements. In *Glaciecola* and *Pseudoalteromonas*, the predicted homologous *spf* genes occurred near the original *spf*-associated genes, polA and yihA/ysxC, although additional conserved genes were inserted between them. In *Frankia* and *Bifidobacterium*, genes of potential ms1 RNA homologs were identified by expanding the synteny criteria to include functionally related septum-site, conjugation, secretion, and pilus-associated genes.

Overall, these use cases show that the approach can detect divergent ncRNA homologs that are difficult to identify by sequence similarity alone. A detailed description of each use case follows.

### spot 42 RNA genes in *Glaciecola*

spot 42 RNA, encoded by the *spf* gene, was first characterized in *Escherichia coli* and later identified in several γ-proteobacterial orders based on sequence similarity (28). In *Glaciecola* and *Pseudoalteromonas* genera, the existence of homologous *spf* genes was questionable. The gene was reported in one species in each, but not in other species of these genera (27).

Here, to identify the *spf* gene in *Glaciecola*, the first step of the presented approach, the genome-wide structure search was applied to 7 *Glaciecola* genomes downloaded from NCBI GenBank (Supplementary Table S3). The template was the *E. coli* spot 42 secondary structure and the sliding window was formed by 119 nucleotides, which was the length of the template sequence.

As ea result, the algorithm identified in average 86 statistically significant structure candidates per genome using the *s^m^* score, which was in average 3.8% of all IGRs in the searched genomes.

The candidates were subjected to the synteny-based filtering, done with the following synteny phrases:

I. {’ribosome biogenesis GTP’ OR ‘ribosome GTPase’ OR ‘YihA’ OR ‘YsxC’ OR ‘EngB’};
II. {’DNA polymerase I’ OR ‘polA’}.

The synteny phrases were derived from the annotations of the genes adjacent to IGRs containing known spf genes. In these cases, the spf-containing IGR was flanked by a gene annotated as “ribosome biogenesis GTP-binding protein YihA/YsxC” or “EngB” on one side and a gene annotated as “DNA polymerase I” on the other. These flanking genes were strongly conserved across species with known spf genes included in the Rfam database (data not shown), and this conserved syntenic context was also used in Baekkedal et al. 2015.

The genomic context of a statistically significant structure candidate was considered syntenic if any string of one phrase occurred in either the upstream or downstream part of the context, while any string the other phrase occurred in the opposite part.

The synteny-based filtering identified syntenic structure candidates in *G. sp. HTCC2999*, G*. sp. MH2013*, *G. sp. 2405UD65-10* and *G. sp. MF2-115*.

In the remaining 3 species, the *s^3^* score was applied to identify statistically significant structure candidates, producing in average ∼102 statistically significant candidates per genome which was in average 4% of all IGRs in the genomes. Among the candidates, the syntenic ones were identified in *G. sp. KUL10* and *G. sp. 33A*.

In the remaining species, *G. siphonariae*, the syntenic candidate was identified among 113 statistically significant structure candidates making it 4% of all IGRs in *G. siphonariae* genome produced by the score *d^m^*. The sequences and genomic data of the candidates can be found in Supplementary File S2 and their secondary structures in Supplementary File S3.

Next, the syntenic candidates were tested for their evolutionary conservation. The sequences of the candidates were BLASTed against the refseq_genomes database in the *Alteromonadales* order to limit the search to related species. In the species in which IGRs with sequence similarity were detected by BLAST, the *spf* synteny was checked using the same synteny-bases filtering as for the candidates described above.

The *G. sp. HTCC2999*, *G. sp. MF2-115, G. sp. KUL10, G. sp. 33A* and *G. siphonariae* candidates had statistically significant sequence similarity to 43, 3, 2, 9 and 4 IGRs in various related *Alteromonadales* species that also exhibited the *spf* synteny, respectively. The species are listed in Supplementary File S3. Among these species, another 4 *Glaciecola* species were identified, namely *Glaciecola sp. SC05*, *Glaciecola sp. G7-5*, *Glaciecola sp. 1036* and *Glaciecola sp. XM2 124*. Their IGRs with the sequence similarity to the syntenic candidates could be considered as containing another *spf* candidates, as they had *spf*-syntenic genomic contexts and statistically significant sequence similarity to the candidates.

No sequence similarity-homologs were found for *G. sp. MH2013* and *G. sp. 2405UD65-10*, but the latter was found among the species with sequence similarity to the candidates, validating its prediction. Thus, except for the *G. sp. MH2013* candidate, the candidates identified in the remaining 6 *Glaciecola* species were proposed as *Glaciecola spf* homologs, as they were *spf*-syntenic and evolutionarily conserved, and exhibited sequence similarity to *spf*-syntenic IGRs in mutiple evolutionarily related species.

The scheme showing the genomic contexts of the proposed *Glaciecola spf* homologs identified by the presented approach is shown in Figure 2. The broader genomic contexts are shown in Supplementary Figure S4. As shown in Figure 2, four candidates, *G. sp. KUL10*, *G. sp. 33A*, *G. siphonariae* and *G. sp. HTCC2999*, retained the original *spf* gene context known from *γ-Proteobacteria* that is DNA polymerase I (polA) and ribosome biogenesis GTP-binding protein YihA/YsxC. The remaining two proposed *spf* homologs in *G. sp. 2405UD65* and *G. sp. MF2*, and the candidate in *G. sp. MH2013* with unverified evolutionary conservation, showed partial synteny: the yihA/ysxC gene was preserved, but polA was separated from the *spf* genes by two to five intervening genes, most likely as a result of local genomic rearrangements during the evolution retaining immediate partial genomic context of yihA/ysxC gene.

**Figure 2.**
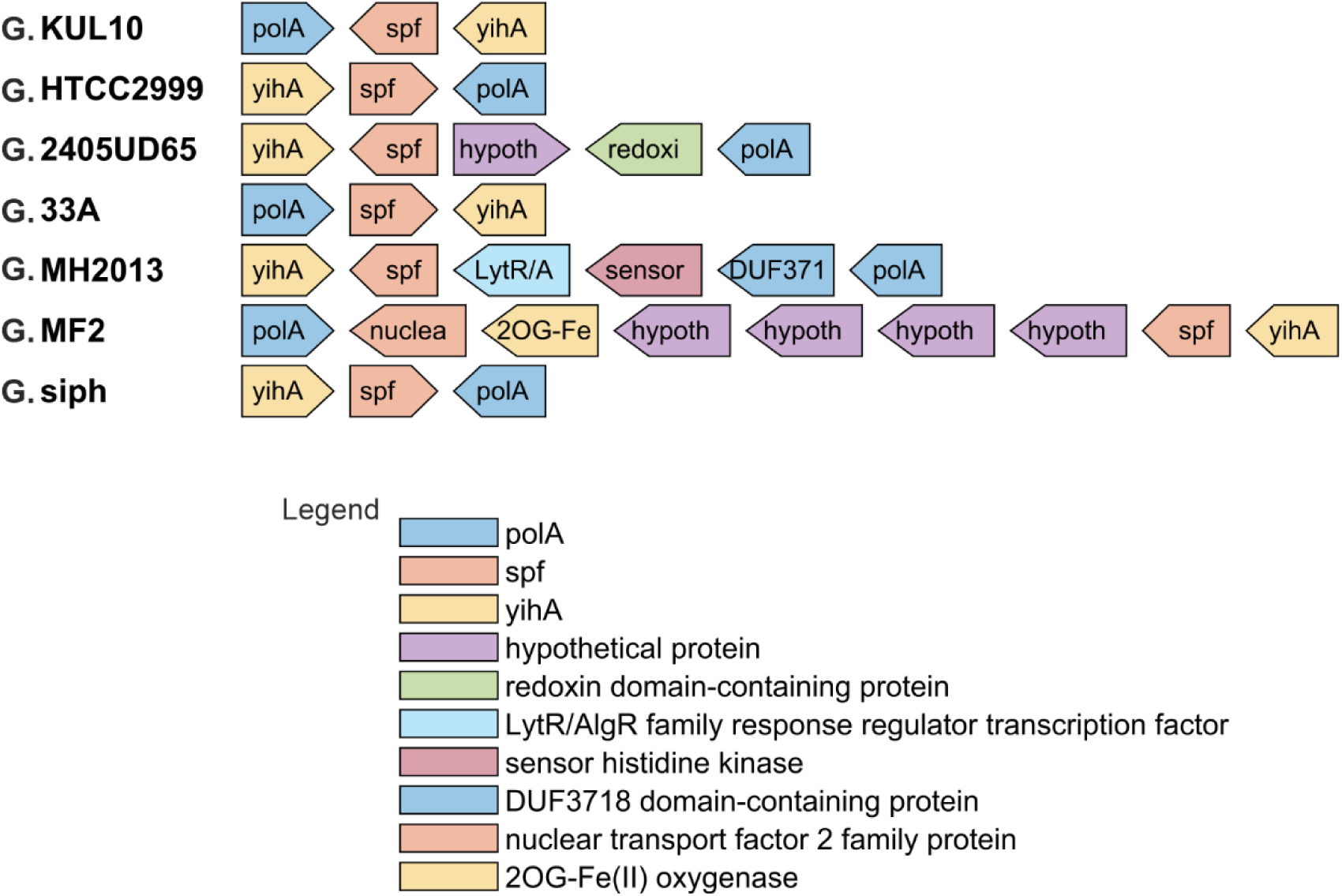
Genomic contexts of *Glaciecola spf* homologs predicted using the presented approach. The contexts are organized in rows of arrow shapes beneath each other, each for one *Glaciecola* species. The species are indicated by their names on the left. The shapes depict genes, the spaces between them are IGRs. The *spf* homologs are indicated by the *spf* labels. Lengths of the both shapes and spaces correspond to lengths of genes and IGRs. The shapes point in the direction of genomic orientation of the genes. The shapes are labeled by shortcuts of gene annotations, shown in full in the legend. The *Glaciecola* species are marked by species names on the left. They are: *G. sp. KUL10* (G. KUL10), *G. sp. HTCC2999* (G. HTCC2999), *G. sp. 2405UD65-10* (G. 2405UD65), *G. sp. 33A* (G. 33A), *G. sp. MH2013* (G. MH2013), *G. sp. MF2-115* (G. MF2) and *G. siphonariae* (G. siph).

### spf homologs in Pseudoalteromonas

In this use case, six *Pseudoalteromonas* genomes from NCBI GenBank (Supplementary Table S3) were searched for *spf* homologs. The structure search was done with the same template and the sliding windows length as for the *Glaciecola* use case.

Using the *s^m^*score, the structure search identified in average 109 statistically significant candidates per genome that was in average 4.3% of all IGRs in the genomes.

Among them, syntenic candidates were identified in four species, *P. atlantica*, *P. maricaloris*, *P. marina* and *P. haloplanktis*, using the same synteny phrases as for the previous use case. Their genomic contexts contained both genes of the original *spf* synteny, polA and yihA/ysxC, but another genes were inserted between them (Figure 3). The inserted genes were repeating among the four species, indicating their evolutionary conservation, unlike in *Glaciecola*, where the inserted genes differed among species.

**Figure 3.**
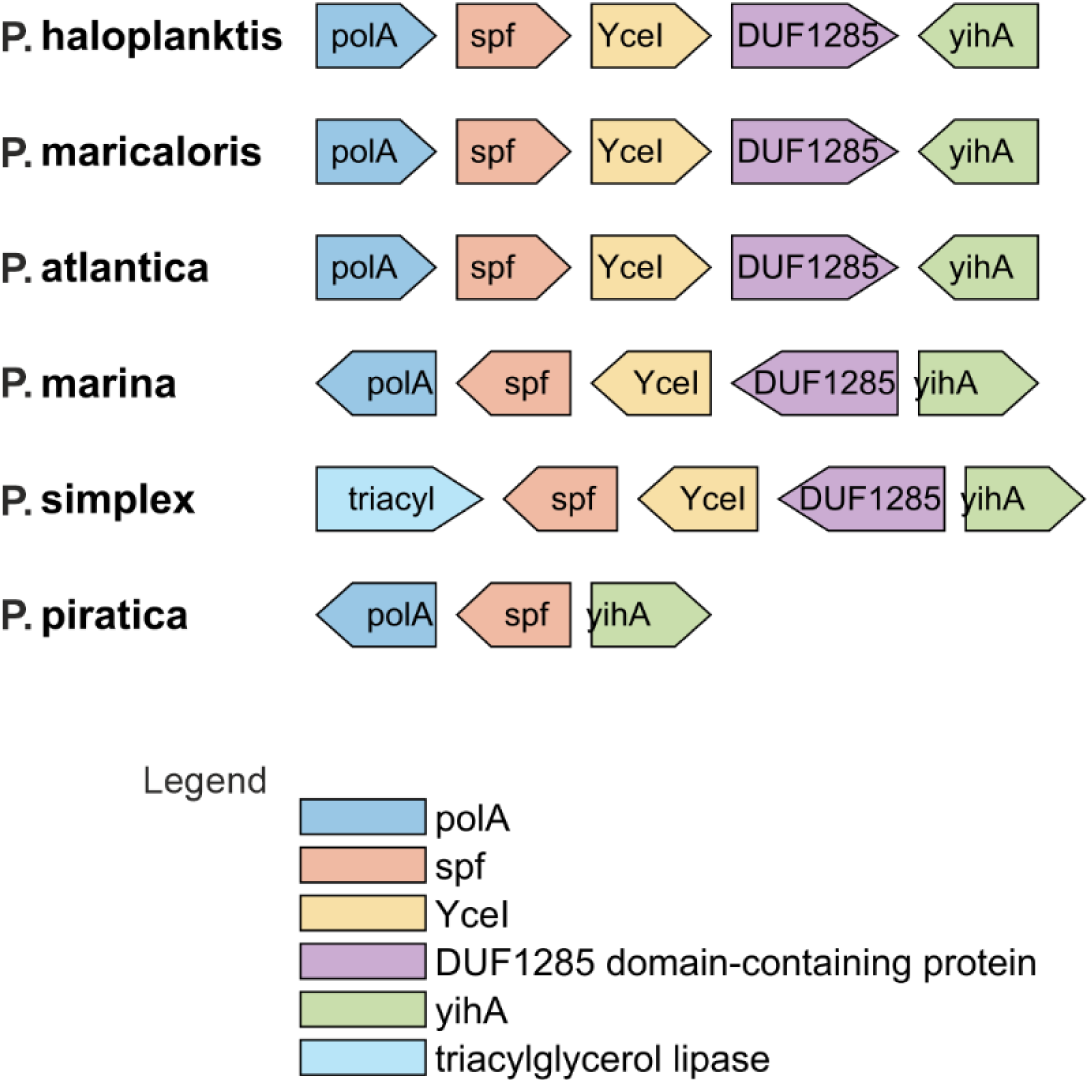
Genomic contexts of *Pseudoalteromonas spf* homologs predicted using the presented approach. The contexts are organized in rows of arrow shapes beneath each other, each for one *Pseudoalteromonas* species. The species are indicated by their names on the left. The shapes depict genes, the spaces between them are IGRs. The *spf* homologs are indicated by the *spf* labels. Lengths of the both shapes and spaces correspond to lengths of genes and IGRs. The shapes point in the direction of genomic orientation of the genes. The shapes are labeled by shortcuts of gene annotations, shown in full in the legend. The *Pseudoalteromonas* species are marked by species names on the left. They are: *P. haloplanktis* (P. haloplanktis), *P. maricaloris* (P. maricaloris), *P. atlantica* (P. atlantica), *P. marina* (P. marina), *P. simplex* (P. simplex) and *P. piratica* (P. piratica).

As the inserted genes appeared to be conserved, a new synteny Ia was designed:

Ia. {[’ribosome biogenesis GTP’ OR ‘ribosome GTPase’ OR ‘YihA’ OR ‘YsxC’ OR ‘EngB’] AND ‘YceI’}.

Using the phrase Ia, a genomic context was considered syntenic if any string of both the sub-phrases of the phrase I occurred in its either upstream or downstream part, while the the phrase II was not used.

This synteny-based filtering identified another syntenic candidate in *P. simplex* with genomic context with YceI and yihA/ysxC genes, whereas polA was missing. Manual search for polA in *P. simplex* genomic annotations showed polA located far in the genome from the predicted *spf* gene, thus also from YceI and yihA/ysxC genes.

A syntenic candidate in the last of the six searched species, *P. piratica*, was identified among the statistically significant candidates obtained by the *d^m^*score. The score identified 132 statistically significant candidates in the genome that was 5% of all IGRs in the genome. The candidate preserved the original polA/yihA synteny. The sequences and secondary structures of the candidates were included in Supplementary File S4 and Supplementary Figure S6. The broader genomic contexts of the candidates are shown in Supplementary Figure S5.

Next, the syntenic candidates were tested for their evolutionary conservation. The candidate sequences were BLASTed against the refseq_genomes database in *Alteromonadales* order to limit the search to the related species.

The six candidates had statistically significant sequence similarity to 41, 51, 49, 47, 36 and 43 IGRs with *spf* synteny in related species for *P. atlantica*, *P. maricaloris*, *P. marina*, *P. haloplanktis*, *P. simplex* and *P. piratica*, respectively. The synteny of the IGRs with the sequence similarity was identified using the synteny-based filtering with the synteny phrases Ia and II and their filtering rules described above. The species in which the syntenic IGRs with sequence similarity was found are listed in Supplementary file S5.

The species with sequence similarity to the syntenic candidates appeared to be only from the *Pseudoalteromonadales* genus. One exception was *P. piratica* whose syntenic candidate had sequence similarity also in other than *Pseudoalteromonadales* genus, in *Flocullibacter*. The BLAST of the *Flocullibacter* IGR sequence in *Alteromonadales* identified statistically significant sequence similarity in IGRs in 29 species of two *Alteromonadales* genera other than *Pseudoalteromonadales*, in *Psychrosphaera* and *Alishewanella* (Supplementary File S6) that were *spf*-syntenic (Supplementary File S7), thus further confirming the validity of the search. The broader genomic contexts of these IGRs are shown in Supplementary Figure S7.

### ms1 RNA genes in *Frankia*

In this use case, the presented approach was applied to the identification of ms1 RNA genes in *Frankia*. The first step of this approach, the structure search, was applied to 25 *Frankia* genomes available in NCBI GenBank (Supplementary Table S3). The template was the experimentally determined secondary structure of *S. coelicolor* ms1 RNA (29).

Using the *s^m^*, *s^3^*, *s^s^*, *d^m^* and *d\**scores, the structure search identified in average 128, 63, 24, 35 and 37 statistically significant candidates per genome that represented 3.9% 2% 0.9% 1% and 1% of all IGRs in the genomes in average.

Among these candidates, using synteny-based filtering, those with the ms1-like syntenic genomic contexts were identified. The synteny phrase for ms1 RNA was derived from conserved genes in the genomic contexts of known ms1 RNA homologs in *Actinomycetes* (29). The contexts of seven flanking genes in each genomic direction from the central ms1 RNA-containing IGR were analyzed. Among them the most conserved were the 4^th^, 3^rd^, 2^nd^ and 1^st^ genes on one side of the IGR and the 1^st^ gene on the other side, where the 1^st^ means the closest gene to the IGR. The genes were in the order 4^th^, 3^rd^, 2^nd^, 1^st^//1^st^, where ‘//’ marks position of IGRs with ms1 RNA: ‘type II secretion system F’, ‘TadA family conjugal transfer-associated ATPase’, ‘septum site-determining protein Ssd’, ‘HAD-IB hydrolase’ or ‘HAD-IB phosphatase’//‘oxidoreductase’ or ‘Fic Filamentation induced by cAMP family protein’ or ‘glycoside hydrolase’.

Exceptions from this scheme occurred in a few *Actinomycetes* genera, but repeated among the species within those genera. These exceptions included the absence of the 1^st^ gene, “HAD-IB hydrolase/phosphatase”’ as observed, for example, in *Actinomadura*; the absence of the 4^th^ gene, “type II secretion system F,” as observed, for example, in *Glycomyces* and *Micromonospora*; and replacement of either the 3^rd^ or 2^nd^ gene by “Flp pilus assembly protein ATPase CpaE/FlpE/TadZ,” as observed, for example, in *Actinoalloteichus*.

The synteny phrases derived from the annotations were as follows:

III. ‘TadA’ AND ‘Ssd’ AND {‘HAD-IB’ AND [‘hydrolase’ OR ‘phosphatase’]};
IV. {‘oxidoreductase’ OR ‘Fic’} OR {‘glycoside’ AND ‘hydrolase’}.

A genomic context of a statistically significant structure candidate was considered syntenic if strings of one phrase, separated in the phrases by the highest level AND, occurred in either the upstream or downstream part of the context at the appropriate positions of flanking genes in the matched context, while strings of the other phrase occurred at the appropriate positions of flanking genes in the matched context in the opposite part.

Filtering the genomic contexts of statistically significant *Frankia* structure candidates using the full synteny phrase did not identify any candidate. Therefore, only the “HAD-IB” AND [“hydrolase” OR “phosphatase”] part of the phrase III, representing genes immediately adjacent to IGRs with structure candidates, was used instead of the full phrase. Filtering using this part identified structure candidates in IGRs between “HAD-IB phosphatase” and “5′–3′ exonuclease” genes in 6 of the 25 searched *Frankia* species (Supplementary Figure S8). Exonucleases are distantly functionally related to glycoside hydrolases, suggesting that this genomic context may be functionally similar to the “HAD-IB hydrolase/phosphatase”//“glycoside hydrolase” context.

Nevertheless, only 12 of the 25 searched *Frankia* species possessed IGRs between “HAD-IB phosphatase” and “5′–3′ exonuclease” genes that were long enough to accommodate ms1 RNA genes. Note, that the length of *S. coelicolor* ms1 RNA is 227 nucleotides, thus the IGRs with its gene should have at least 250 nucleotides. In the 13 remaining species, ms1 RNA could not exist in these IGRs and also could not be detected. The occurrence of these suitable IGRs in only 12/25 *Frankia* species did not indicate evolutionary conservation. The lack of this sequence conservation agreed with the lack of statistically significant structure candidates that were identified in only 6 out of the 12 species with these long enough IGRs. Moreover, the broader contexts of these IGRs did not match the ms1 RNA synteny described above.

It was therefore relevant to determine whether IGRs with statistically significant structure candidates occurred in genomic contexts that, as observed in e.g. *Actinomadura*, lacked HAD-IB hydrolase/phosphatase genes but still contained the other conserved genes of the ms1 RNA synteny.

The contexts of the structure candidates therefore were searched for occurrence of the keywords “Ssd” and “TadA”, which represent unambiguous signatures of two of the most conserved genes in the ms1 RNA synteny: “septum site-determining protein Ssd” and “TadA family conjugal transfer-associated ATPase”. Indeed, candidates in IGRs flanking with the succession of these two genes accompanied by “type II secretion system F family” were found. Such contexts matched the ms1 RNA synteny. But the structure candidates were identified in only 3 such IGRs (Supplementary Figure S9), although IGRs with this context longer than 250 nucleotides, and therefore able to accommodate ms1 RNA genes, were present in 15 of the 25 searched *Frankia* species. Again, the structure candidates were too few to indicate evolutionarily conserved occurrence of homologous ms1 RNA structures in these IGRs.

Therefore, a broader search was performed for the genomic contexts of IGRs with structure candidates that were functionally or biologically related to the ms1 RNA synteny. This search did not rely on highly specific gene names as previously, but instead used keywords from annotations of genes within the ms1 RNA synteny that reflected the associated biological processes. Two such keywords were “septum” and “conjugal”. The contexts that would be identified using “septum” keyword were already identified in the search above using the “Ssd” gene name. The “conjugal” keyword, when not accompanied by “TadA”, occurred in the “conjugal transfer protein TraC” annotations found in 14 contexts in 14 *Frankia* species. Using ChatGPT 5.5, these proteins were identified as functionally related members of distinct ATPase families (Supplementary File S12), suggesting that genes annotated as “conjugal transfer protein TraC” may have occured in genomic contexts functionally related to ms1 RNA synteny. The sources included in the ChatGPT reports in supplementary files, as well as references retrieved by ChatGPT and used in this work, were manually validated.

Some of the contexts containing the genes of TraC proteins, contained “MinD/ParA family ATP-binding protein”, that was adjacent to the IGRs with structure candidates. These proteins were found to be functional analogs of Ssd proteins and belong to the ParA–MinD ATPase superfamily, which includes septum-site-regulating proteins based on ChatGPT 5.5 functional comparison (Supplementary File S13) (30) (31). This classification further supported the evidence that the contexts containing the genes of TraC proteins and were functionally related to the *ms1* synteny.

These MinD-containing contexts, extending up to 6 genes in both directions from the “MinD” gene, consisted of the following annotations (the context from *Frankia casuarinae* used as an example): “conjugal transfer protein TraC,” “SCO6880 family protein,” “hypothetical protein,” “hypothetical protein,” “DUF6112 family protein,” “MinD/ParA family ATP-binding protein,” “helix-turn-helix domain-containing protein,” “hypothetical protein,” “NlpC/P60 family protein,” “hypothetical protein,” and “FAD-binding oxidoreductase.” Functional and biological characterization of the genes in this context, based on both ChatGPT 5.5 functional analysis and sequence similarity of proteins produced by these genes to annotated genes in species in other *Actinomycetes* genera identified by Protein BLAST searches in NCBI, indicated that these contexts were related to septum-site-, conjugation-, and secretion-related cellular processes (Table 2) i.e. to the same cellular processes as the genes in the ms1 RNA synteny.

**Table 2.**
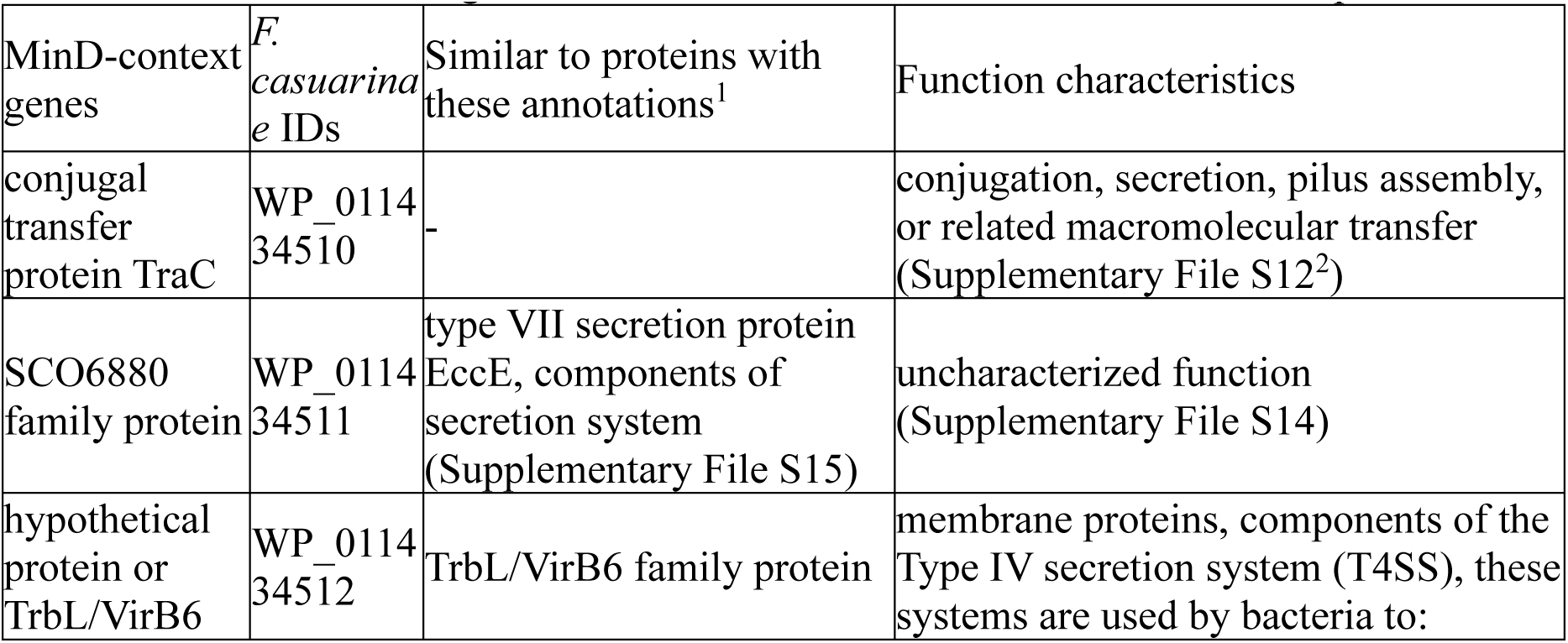
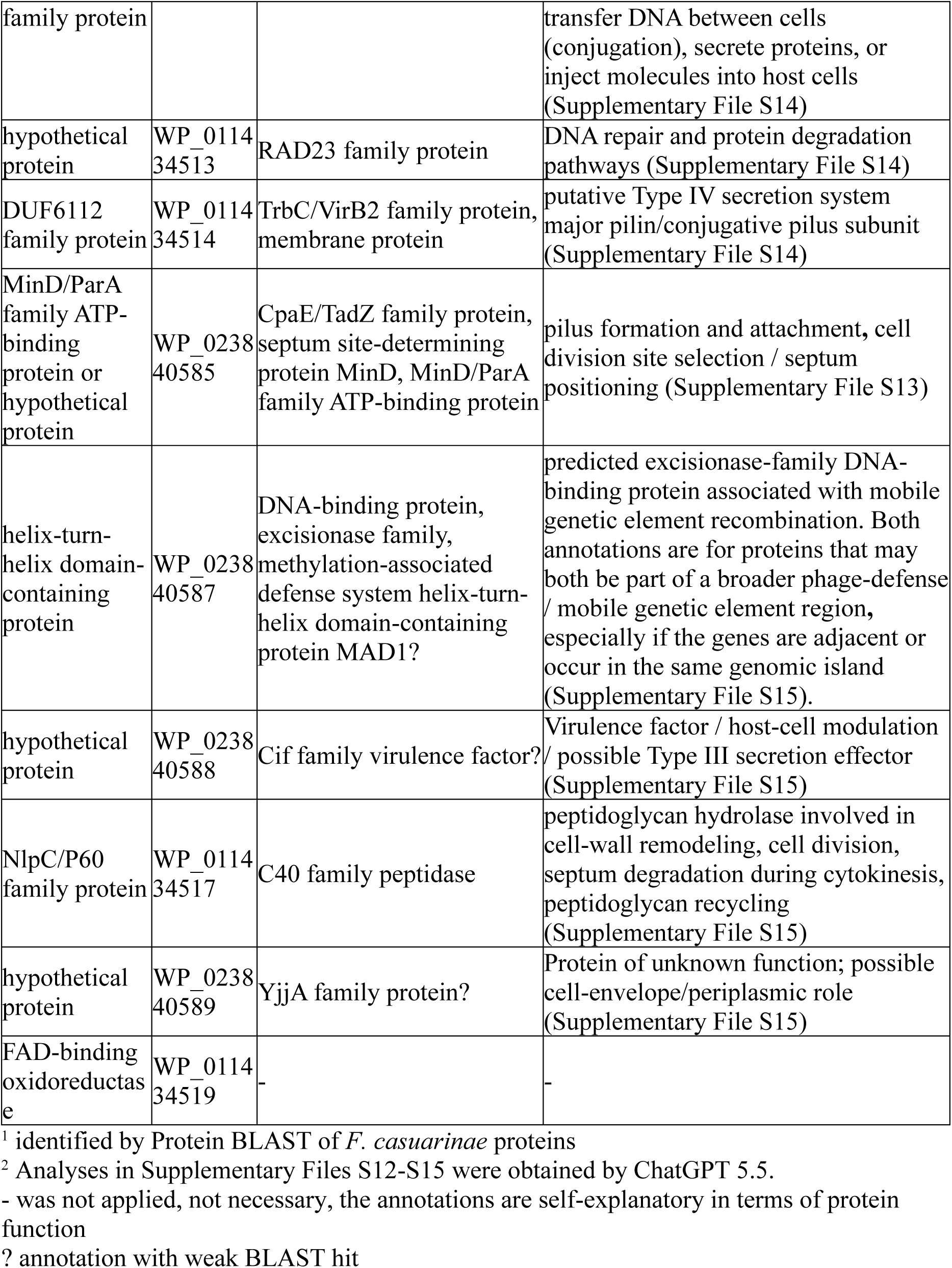
Search for annotated homologs for genes in the MinD-containing contexts using similarity-based analysis by BLAST in *Actinomycetes*. The search is demonstrated here with *Frankia casuarinae*. Some genes could have different annotations in different species.

The synteny phrases were constructed for identification of these contexts among genomic contexts of structure candidates in the analyzed *Frankia* species:

V. ‘NlpC/P60’ OR ‘helix-turn-helix’ OR [‘FAD AND ‘oxidoreductase’];
VI. [‘ MinD’ OR ‘ParA’] OR ‘SCO6880’ OR [‘TraC’ OR ‘conjugal’].

The filtering rule was the same as for the phrases III and IV: at least one string of each phrase had to appear in either the upstream or downstream part from the IGR with a structure candidate in the matched context, while they have to appear in opposite parts.

In 7 of the 25 species, namely *Frankia gtarii*, *tisae*, *alpine*, *sp. KB5*, *EI5c*, *Hr75*, *RB7*, the IGRs with contexts corresponding the synteny phrases V an VI were split between contigs and therefore could not be searched for the ms1 RNA structure. In one *Frankia* species, *Cj5*, this IGR was not long enough. Therefore only 17 analyzed *Frankia* species had IGRs corresponding the synteny phrases V an VI long enough to theoretically accommodate putative homologs of the 227-nucleotide-long *S. coelicolor* ms1 RNA. Of these 17 *Frankia* species, 14 had statistically significant structure candidates in IGRs corresponding the synteny phrases V an VI (Figure 4).

**Figure 4.**
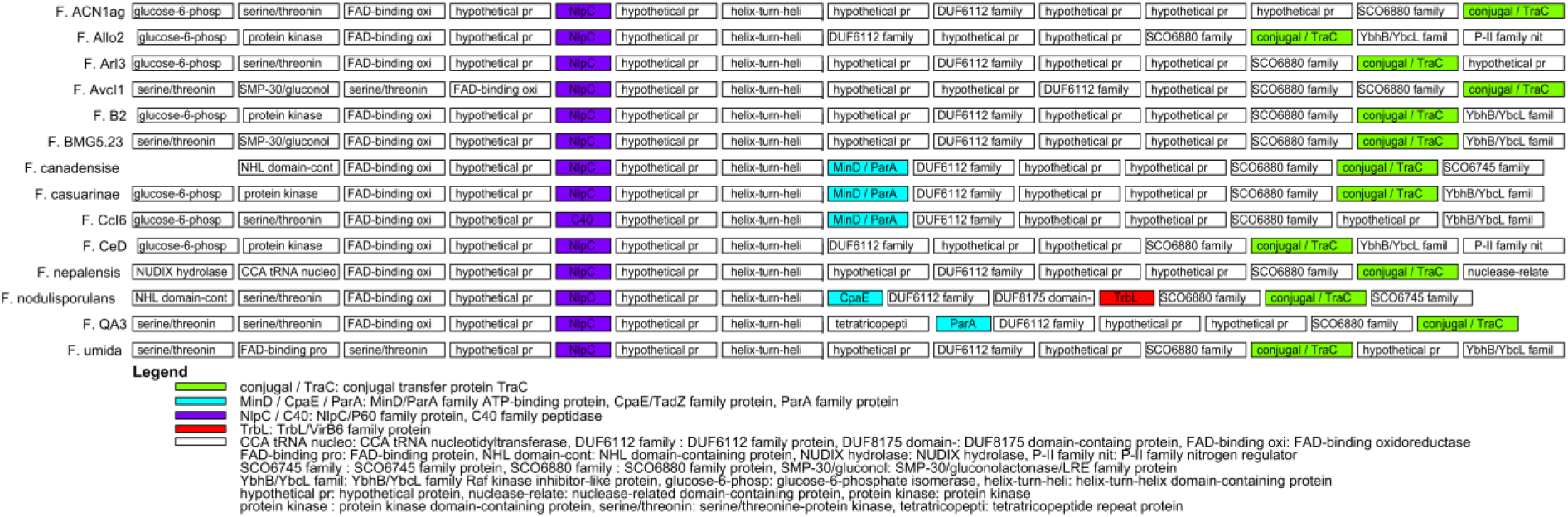
Broader genomic contexts of genes of *Frankia* ms1 RNA homologs predicted using the presented approach. The contexts are organized in rows of boxes beneath each other. The boxes mark genes, the spaces between them IGRs. The lengths of both do not correspond to the lengths of genes and IGRs. The boxes contain shortcuts of gene annotations, shown in full in the legend. The contexts consist of 14 genes, 7 genes in each of the genomic directions from the central IGRs with predicted homologs. If the genes were not available in annotations, the appropriate positions in the contexts are empty. Colored boxes show genes complying with ms1 RNA synteny. The gene annotations corresponding to the colors are shown in the legend.

Structure candidates were positioned in these contexts between “helix-turn-helix domain-containing protein” genes and one of these: “MinD/ParA family ATP-binding protein”, “ParA family protein”, “CpaE/TadZ family protein” and “hypothetical protein’. The hypothetical proteins in these contexts exhibited statistically significant sequence similarity to the MinD/ParA family protein-annotated genes in other *Actinomycetes* species, when examined with Protein BLAST in NCBI (not shown). The predicted secondary structures of these candidates were highly similar to the *S. coelicolor* ms1 RNA template and were also the best-scoring structures in all 14 *Frankia* species (their plots were included in Supplementary Figure S10).

Sequence similarity of the syntenic candidates in related species was examined using NCBI BLASTn Web. Statistically significant hits were obtained in multiple species for each of the candidates, confirming their evolutionary conservation. The species in which the candidates had statistically significant sequence similarity are shown in Supplementary File S8. The sequences and secondary structures in dot-bracket format of the predicted homologs are provided in Supplementary File S7.

Interestingly, when “MinD” keyword was searched in *Frankia* genomic annotations, another “MinD” gene-containing genomic context was found that matched the biological and functional characteristics of the ms1 RNA synteny. This context consisted of the following annotations: “TadE family protein,” “hypothetical protein,” “type II secretion system F family protein,” “type II secretion system F family protein,” “CpaF family protein,” “MinD/ParA family ATP-binding protein,” “SAF domain-containing protein,” “ATP/GTP-binding protein,” “hypothetical protein,” “hypothetical protein,” and “2′-5′ RNA ligase family protein.” It was directly related to bacterial pilus assembly through the CpaF family proteins and indirectly related to conjugation through the TadE family proteins. Nevertheless, these contexts could not harbor homologous ms1 RNA genes in the IGRs flanking the “MinD/ParA family ATP-binding protein” gene, because they were not long enough in any of the 25 analyzed *Frankia* species.

Note that the keywords used to search the annotations were highly specific. They occurred on average only once or twice per genome, unlike terms such as “oxidoreductase” or other annotations that are ubiquitous throughout genomes. The keywords “TadA” and “Ssd,” which represented unambiguous signatures of two of the most conserved genes in the ms1 RNA synteny, “TadA family conjugal transfer-associated ATPase” and “septum site-determining protein Ssd”, occurred in 23 and 20 of the 25 searched *Frankia* species, respectively, and were adjacent to each other in 19 species. The “septum” keyword occurred 66 times in genomic annotations of the 25 *Frankia* species, 20 times in the “septum site-determining protein Ssd”. The 46 remaining occurrences were found in contexts clearly unrelated to the ms1 RNA synteny (not shown). The “conjugal” keyword, when not in “TadA family conjugal transfer-associated ATPase”, occurred in 23 of the 25 species. In 22 of these 23 occurrences, it appeared in the annotation “conjugal transfer protein TraC.”

The conclusion stemming from this brief search in *Frankia* genomic annotations was that these septum-site-, conjugation-, and secretion-related contexts are specific in *Frankia* genomes and that the occurrence of the statistically significant structure candidates in them was not random.

### ms1 RNA genes in *Bifidobacterium*

In this use case, the presented approach was applied to the identification of ms1 RNA genes in *Bifidobacterium*. The structure search was performed on genomes of 22 *Bifidobacterium* species available in NCBI GenBank (Supplementary Table S3). The template was the experimentally determined secondary structure of *S. coelicolor* ms1 RNA (29).

Using the *s^m^* score, structure candidates in 20 of the 22 inspected species were identified. The search identified in average 33 statistically significant structure candidates per searched genome, which was in average 4% of all IGRs in the searched genomes.

The candidates were subjected to the synteny-based filtering. It followed the filtering procedure applied in *Frankia*. First, the original synteny phrases III and IV were used. As ‘HAD-IB hydrolase’ did not occur in *Bifidobacterium*, ‘HAD hydrolase’ was used instead. No candidate was found using either full phrases or the “HAD-IB” AND [“hydrolase” OR “phosphatase”] part of the phrase III.

Therefore as in *Frankia*, the synteny filtering using the rest of the original ms1 RNA synteny was applied. The septum site-determining Ssd proteins did not occur in the searched *Bifidobacterium* genomes. Instead there was ‘CpaE/TadZ family protein’ followed by ‘TadA family conjugal transfer-associated ATPase’ and one or two copies of ‘type II secretion system F family protein’, which as in *Frankia* represented the most conserved part of the original ms1 RNA synteny shown above. In some *Bifidobacterium* species, instead of CpaE/TadZ family proteins, there was either the P-loop NTPase or AAA family ATPase annotations that are broader, less specific annotations that may encompass or overlap with CpaE/TadZ-family proteins (Supplementary File S18). The CpaE/TadZ family proteins are related to the Ssd/MinD/ParA-like septum site-determining ATPases (32) (Supplementary File S19). Also, TadA-family conjugal transfer-associated ATPases were annotated in some *Bifidobacterium* species as CpaE family proteins, as their more general classification is CpaF/TadA-like pilus assembly ATPases (KEGG entry F6J84_03795). Then, the following phrase was constructed and used in synteny filtering:

VII. [’CpaE’ OR ‘P-loop NTPase’ OR ‘AAA family ATPase’] AND [’CpaF’ OR ‘TadA’] AND ‘type II secretion system F’.

The filtering rule required that OR operator-separated substrings from all three AND-separated strings has to be found in the annotations of the corresponding genes in the genomic contexts of IGRs with structure candidates, with the gene matching the “CpaE”-containing string flanking the IGR.

This phrase was identified in genomic contexts of 14 of the 22 analyzed *Bifidobacterium* species, in which the candidates were in the IGRs flanking the CpaE/TadZ family proteins (Figure 5). Of the 22 analyzed *Bifidobacterium* species, 20 had these CpaE/TadZ family proteins–flanking IGRs long enough to theoretically accommodate putative homologs of the 227-nucleotide-long *S. coelicolor* ms1 RNA.

**Figure 5.**
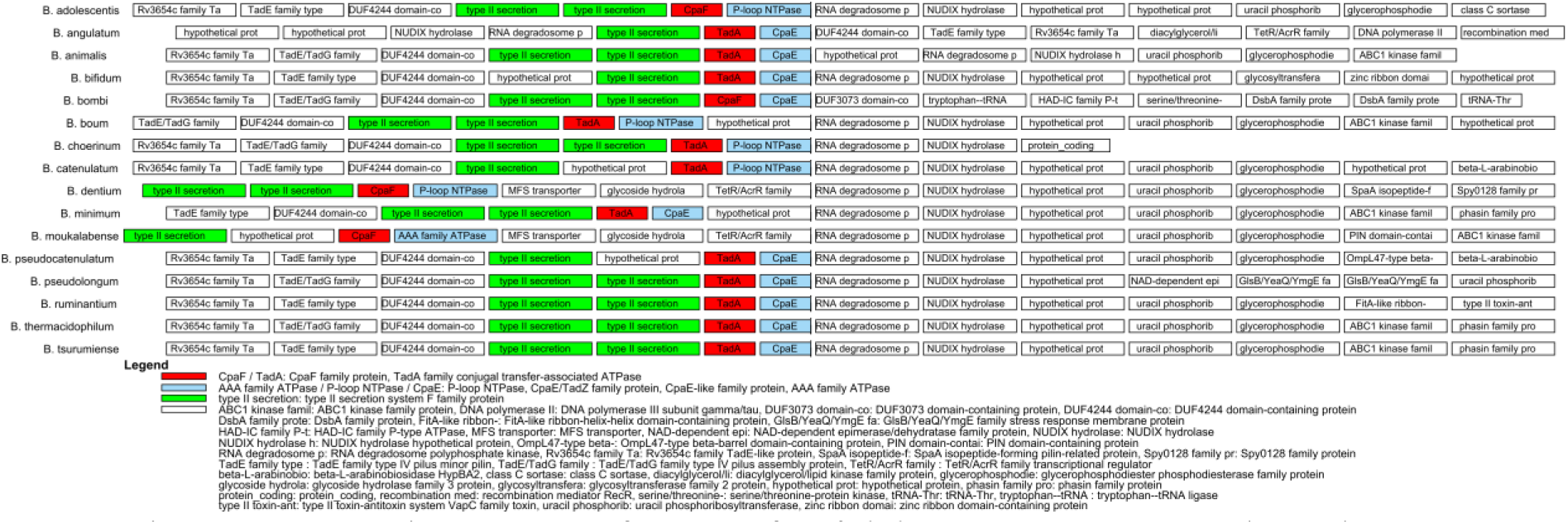
Genomic contexts of genes of *Bifidobacterium* ms1 RNA homologs predicted using the presented approach. The contexts are organized in rows of boxes beneath each other. The boxes mark genes, the spaces between them IGRs. The lengths of both do not correspond to the lengths of genes and IGRs. The boxes contain shortcuts of gene annotations, shown in full in the legend. The contexts consist of 14 genes, 7 genes in each of the genomic directions from the central IGRs with predicted homologs. If the genes were not available in annotations, the appropriate positions in the contexts are empty. Colored boxes show genes complying with ms1 RNA synteny. The gene annotations corresponding to the colors are shown in the legend.

Using the *s^s^* score, statistically significant structure candidates were identified in another 2 species, *B. animalis* and *B. boum*, with the precision of 24 statistically significant structure candidates per searched genome in average, which was 3% of all IGRs in average in the searched genomes. Both the candidates were found syntenic using the above described synteny filtering procedure.

In the four *Bifidobacterium* species without statistically significant structure candidates, no candidates were identified using the remaining scores.

Sequence similarity of the syntenic candidates in related species was examined using NCBI BLASTn Web. Statistically significant hits were obtained in multiple species for each of the candidates, confirming their evolutionary conservation. The species in which the candidates had statistically significant sequence similarity are shown in Supplementary File S10.

The sequences and secondary structures in dot-bracket format of the predicted homologs are provided in Supplementary File S9 and secondary structure plots are in Supplementary Figure S11.

## DISCUSSION

The identification of homologous bacterial ncRNAs remains challenging when sequence similarity is weak or absent, particularly for short regulatory RNAs whose functional constraints may be imposed more strongly on secondary structure, genomic context, or evolutionary conservation than on primary sequence. In this study, we present an integrative bioinformatic approach that combines genome-wide structural similarity searches based on suboptimal RNA secondary structures with synteny-based filtering and comparative conservation analysis. The results show that these complementary criteria can identify plausible ncRNA homologs in bacterial genomes in cases where conventional sequence-based approaches had previously failed.

A central feature of the presented approach is the use of suboptimal secondary structures rather than reliance on a single minimum free energy structure. This is important because bacterial ncRNAs may populate multiple structurally plausible conformations, and experimentally observed structures are not always identical to the predicted MFE fold. By comparing ensembles of suboptimal structures with a template structure, the algorithm evaluates whether a genomic sequence has the capacity to adopt a fold compatible with the query ncRNA. This ensemble-based strategy is especially relevant for ncRNAs whose function depends on conserved structural elements despite extensive sequence divergence. The successful identification of known homologs in the validation searches supports the usefulness of this principle and indicates that structural information can extend the detectable evolutionary range of ncRNA homology searches beyond sequence similarity alone.

The validation searches demonstrated that structural similarity alone can substantially reduce the genomic search space. Across the test searches, the best-performing scores identified most known homologs as statistically significant candidates, while reducing thousands of intergenic regions to a smaller set of candidate loci. Among the tested scores, s^m^ and s^3^ showed the highest recovery of known homologs, suggesting that scores based on the behavior of multiple suboptimal structures are particularly informative. This result supports the assumption that ncRNA homologs may be better represented by structural tendencies across an ensemble than by a single best-matching predicted structure. At the same time, the number of false-positive candidates generated by structure search alone remained too high for confident homolog identification. This outcome is expected in genome-wide RNA structure searches because local sequences may form structures resembling a query RNA by chance. Thus, structural similarity is a powerful enrichment step, but it is not sufficient as a stand-alone criterion for final annotation.

The addition of synteny-based filtering addressed this limitation by incorporating independent biological information. In bacteria, conserved genomic neighborhoods often reflect shared evolutionary history, functional linkage, or constraints on local genome organization. Therefore, a structurally similar candidate located in a genomic context related to that of known homologs is more likely to represent a true homolog than a structurally similar candidate located in an unrelated region. The phrase-based synteny strategy used here offers a flexible way to exploit genome annotations without requiring strict conservation of gene order or exact annotation wording. This flexibility is important because bacterial genomes frequently undergo local rearrangements, gene insertions, deletions, and annotation changes. By allowing partial synteny and alternative annotation terms, the method can retain candidates in evolutionarily modified but biologically related genomic contexts.

Evolutionary conservation provided a third level of support. Candidate loci that are structurally similar to the query ncRNA, occur in syntenic genomic contexts, and are conserved among related species are unlikely to represent isolated structural artifacts. This was particularly evident in the use cases. Predicted *spf* homologs in *Glaciecola* and *Pseudoalteromonas* were supported not only by structural similarity to the *Escherichia coli* Spot 42 RNA template, but also by conservation of genomic neighborhoods involving genes such as *polA* and *yihA/ysxC*. In several cases, the original syntenic arrangement was retained, whereas in others only partial synteny remained because of intervening genes or local rearrangements. The ability of the approach to accommodate these rearrangements was essential for detecting candidates that would otherwise have been missed by a stricter synteny requirement.

The *Pseudoalteromonas* use case further illustrates the value of adaptive synteny interpretation. In several species, conserved inserted genes occurred between the canonical *spf*-associated flanking genes, indicating that the local genomic context had evolved in a genus-specific manner. By modifying the synteny phrase to include this conserved derived context, additional candidates could be identified. This finding highlights an important practical aspect of bacterial ncRNA homology searches: synteny is often conserved at the level of a genomic neighborhood rather than as a perfectly preserved pair of immediately adjacent genes. The detection of related syntenic loci in additional *Alteromonadales* species also suggests that the predicted *spf* homologs are part of a broader conserved evolutionary pattern rather than isolated predictions.

The prediction of genes of ms1 RNA homologs in *Frankia* and *Bifidobacterium* represents a more challenging case and demonstrates both the potential and the interpretative complexity of the method. In these genera, previous sequence- and synteny-based searches had not identified *ms1* homologs (24,29). The presented approach detected structurally significant candidates in genomic contexts that were not identical to the canonical *ms1* synteny known from other *Actinomycetes*, but that were functionally and biologically related to it. In *Frankia*, candidates were found near genes associated with septum-site regulation, conjugation, secretion-related processes, and oxidoreductase functions. Although these contexts differed from the original *ms1* neighborhood, they contained genes with functional relationships to conserved components of the known *ms1* synteny. The recurrence of these contexts across multiple species and the presence of highly similar predicted secondary structures support the interpretation that these loci may encode divergent ms1 RNA homologs.

Similarly, in *Bifidobacterium*, predicted *ms1* homologs were associated with genomic neighborhoods containing CpaE/TadZ-family proteins, TadA- or CpaF-related proteins, and type II secretion system F-family proteins. These genes are related to cellular processes represented in the broader *ms1* syntenic context, even though the exact gene composition differed from that observed in other *Actinomycetes*. This result suggests that homologous ncRNAs may be preserved within functionally equivalent or evolutionarily remodeled genomic contexts. For divergent bacterial lineages, such functional-context conservation may be more informative than strict conservation of individual gene names or gene order.

The *Frankia* and *Bifidobacterium* use cases illustrate both the practical value and the necessary limitations of AI-assisted comparative genomics. Conventional structure- and synteny-based filtering initially failed to recover a convincing set of *ms1* gene candidates in *Frankia*, largely because the genomic contexts differed from the canonical ms1 RNA synteny described in other *Actinomycetes*. In this situation, ChatGPT 5.5 was not used as a stand-alone prediction tool, but rather as a hypothesis-generating aid to recognize functional relationships among heterogeneous gene annotations. By comparing the biological processes represented in the known ms1 RNA synteny with the genomic contexts of structural candidates in *Frankia*, the AI analysis highlighted contexts associated with septum-site regulation, conjugation, secretion, and pilus assembly, even when the specific gene names differed from those in the original synteny model. These suggestions guided subsequent conventional searches using biologically meaningful keywords such as “conjugal,” “TraC,” “MinD,” and “ParA,” which led to the identification of a conserved MinD/ParA-flanked context containing the best-scoring ms1-like structural candidates in most *Frankia* species with suitable IGRs. Importantly, the AI-derived hints were validated by independent keyword-based annotation searches, BLAST analyses, structural scoring, and sequence conservation among related species. Thus, the use of AI did not replace standard bioinformatic validation, but expanded the search space by helping to interpret functionally equivalent yet annotation-divergent genomic contexts. This demonstrates that large language models can be useful in RNA gene discovery when applied cautiously as tools for biological reasoning and annotation synthesis, provided that all AI-generated hypotheses are subjected to reproducible, conventional computational verification.

However, when ChatGPT 5.5 was initially used to identify complete genomic contexts consisting of 14 gene annotations, with the predicted structural candidates positioned centrally, it failed to recover contexts biologically or functionally related to the *ms1* synteny. Instead of evaluating the functional characterization of each context as a whole, it rather identified related contexts based on matches to individual annotations, often driven by shared wording rather than biological relevance (see Supplementary Files S16 and S17 for examples). In this task, ChatGPT largely behaved like the keyword-based text search described here. This suggests that reliable genomic context matching would require foundation LLMs specifically trained or adapted for annotation-based context comparison with explicitly specified relevant genomic data sources. While this result highlights a limitation of commercial general-purpose LLMs in genomic context-matching tasks, ChatGPT 5.5 was nevertheless highly useful for inferring biological and functional relationships between proteins from genomic annotations, saving days or weeks of detailed manual work.

The use cases also show that the method can generate biologically testable hypotheses. The predicted homologous *spf* and *ms1* genes are supported by convergent computational evidence, but experimental validation remains necessary. Northern blotting, RT-PCR, RNA-seq-based expression analysis, or targeted transcript mapping could confirm whether the predicted loci are transcribed. Structural probing could evaluate whether the predicted RNAs adopt the proposed folds in vitro or in vivo. Functional studies, such as deletion or overexpression experiments, would be required to determine whether the predicted homologs perform roles comparable to known Spot 42 or ms1 RNAs. Thus, the presented approach should be viewed as a prioritization framework for candidate discovery rather than a replacement for experimental characterization.

Several limitations should be considered. First, the performance of the structure search depends on the quality of the template structure. Experimentally determined structures are preferable, but they are available only for a limited number of bacterial ncRNAs. The validation results suggest that predicted structures can also be useful as templates, but this conclusion was based on a limited number of cases and should be examined further. Second, the approach depends on genome annotation quality for synteny filtering. Inconsistent, incomplete, or overly general annotations may obscure true syntenic relationships or produce misleading phrase matches. Third, although phrase-based synteny filtering is flexible, it requires careful biological interpretation, especially when moving from strict gene-name conservation to broader functional-context conservation. This is particularly relevant for the *ms1* predictions in *Frankia* and *Bifidobacterium*, where the inferred synteny is based on functional relatedness rather than exact conservation of the canonical context.

Another limitation is that the statistical evaluation of structural candidates is performed within individual genomes. This enables genome-specific ranking of candidate intergenic regions, but it does not by itself establish that a candidate is homologous to the query ncRNA. The false-positive rates observed in the validation searches show that structurally significant candidates can occur by chance. The integration of synteny and conservation reduces this problem, but it does not eliminate it entirely. Future improvements could include more formal models that combine structural score, syntenic evidence, conservation across taxa, sequence similarity among candidates, and genomic distance relationships into a single probabilistic framework.

The algorithm was also shown to be responsive to sliding-window length, which can be both a strength and a limitation. Length responsiveness may help estimate the likely boundaries of candidate ncRNAs and distinguish windows that best accommodate the conserved fold. However, it also means that searches for homologs with substantially different lengths from the query require careful parameter selection. Future versions of the method could implement adaptive window sizes or multi-scale scanning to better detect homologs with insertions, deletions, or lineage-specific expansions.

Despite these limitations, the presented approach provides a useful framework for the discovery of structurally conserved bacterial ncRNAs. Its main advantage lies in the integration of three independent but complementary signals. Structural similarity identifies candidate loci with the potential to adopt a query-like fold. Synteny filtering evaluates whether those loci occur in biologically plausible genomic contexts. Evolutionary conservation then tests whether candidates are preserved across related species. Each layer reduces uncertainty introduced by the others, resulting in a more robust strategy than searches based on sequence, structure, synteny, or conservation alone.

In conclusion, the results demonstrate that suboptimal secondary structure analysis, when combined with genomic synteny and evolutionary conservation, can identify bacterial ncRNA homologs beyond the reach of conventional sequence or sequence-structure similarity searches. The predicted *spf* homologs in *Glaciecola* and *Pseudoalteromonas* and the predicted *ms1* homologs in *Frankia* and *Bifidobacterium* expand the candidate distribution of these ncRNA families and provide targets for future experimental validation. More broadly, this work supports the use of integrative, context-aware approaches for ncRNA discovery in bacterial genomes, especially for divergent homologs whose evolutionary conservation is retained primarily at the levels of structure and genomic organization.

## DATA AVAILABILITY

All data are incorporated into the article and its online supplementary material.

## SUPPLEMENTARY DATA

Supplementary Data are available at *NAR* Online

## ACKNOWLEDGEMENTS

The author thanks Jiří Vohradský for reading the manuscript.

## Authors contribution

JP is the only author of the presented work.

