## Supplementary figures for "Integrating suboptimal secondary structures, AI-assisted genomic synteny, and evolutionary conservation to identify bacterial ncRNA homologs beyond sequence similarity"

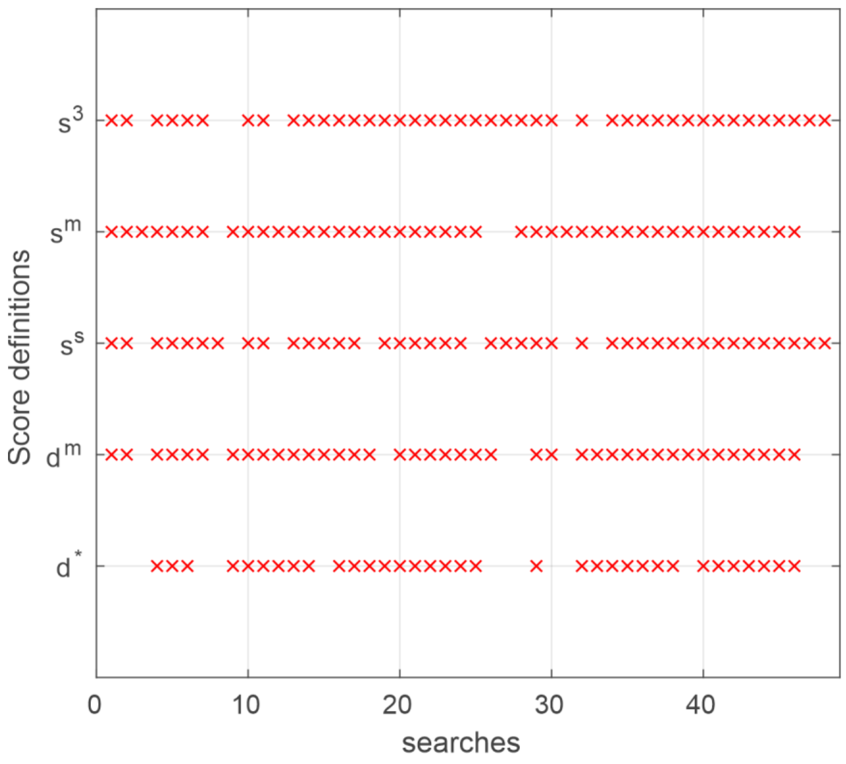


Supplementary Figure S1. Performance of the score definitions in test searches. Y-axis shows the score definitions. X-axis shows searches. Crosses show in which test searches the score definitions identified TP.

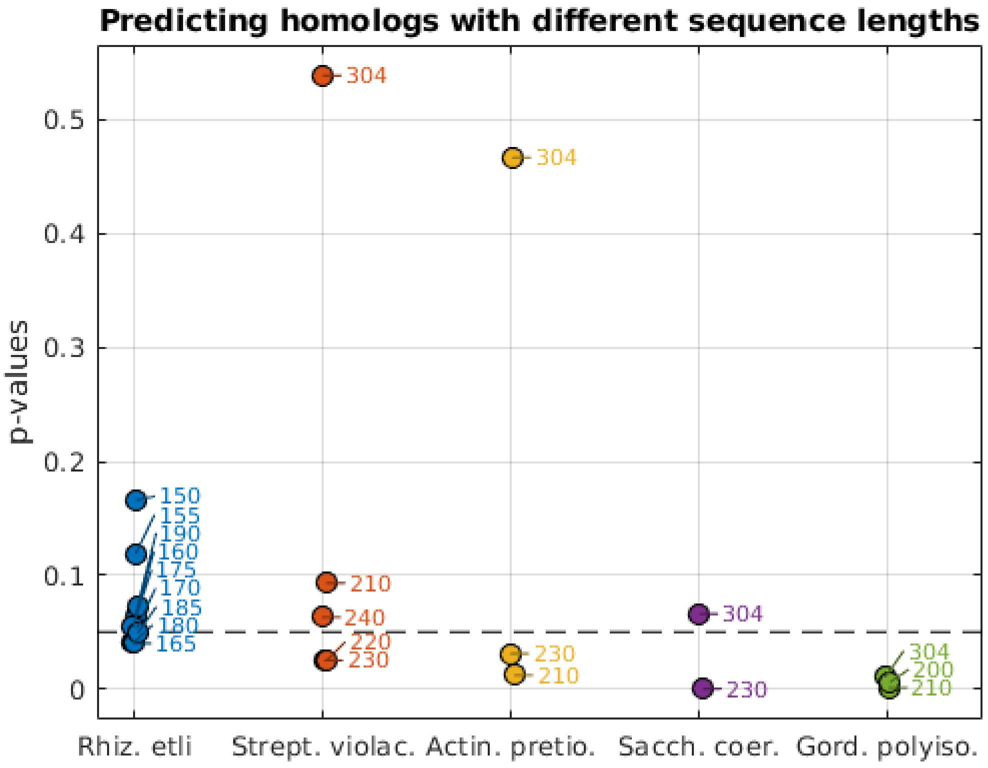


Supplementary Figure S2. Test identification of homologs with different sequence lengths. The test was proceeded with two ncRNAs from the test set for which query RNAs and their experimentally identified homologs had substantially different sequence lengths.

The query RNAs were *E. coli* 6S RNA, 184 nucleotides long, and *M. smegmatis* ms1 RNA with 304 nucleotides.

The homolog of *E. coli* 6S RNA was *Rhizobium etli* 6S RNA with ~160 nucleotides. The homologs of *M. smegmatis* ms1 RNA were ms1 RNAs in *Streptomyces violaceum*, *Actinosynnema pretiousum*, *Saccharothrix coeruleofusca*, *Actinoalloteichus spitiensis* and *Gordonia polyisoprenivorans* with ~230 nucleotides.

In the figure, x-axis shows labels of species in which the test homologs of query ncRNAs were identified. The labels are: Rhiz. etli: *Rhizobium etli*, Strept. violac.*: Streptomyces violaceum*, Actin. pretio.: *Actinosynnema pretiousum,* Sacch. coer.: *Saccharothrix coeruleofusca*, and Gord. polyiso.: *Gordonia polyisoprenivorans*.

The y-axis shows the p-values representing the statistical significance of the identified homologs. Each point represents the p-value obtained from a particular structure-based search. The number next to each point indicates the sliding window length used in that search, which corresponds to the length of the sequence segment analyzed, in nucleotides. The horizontal line indicates the significance threshold of $p = 0.05$.

The figure shows that the choice of sliding window length is important for correctly identifying homologs. Statistically significant p-values, shown as points below the horizontal $p = 0.05$ line, were obtained when the sliding window length corresponded closely to the actual sequence length of the identified homologs.

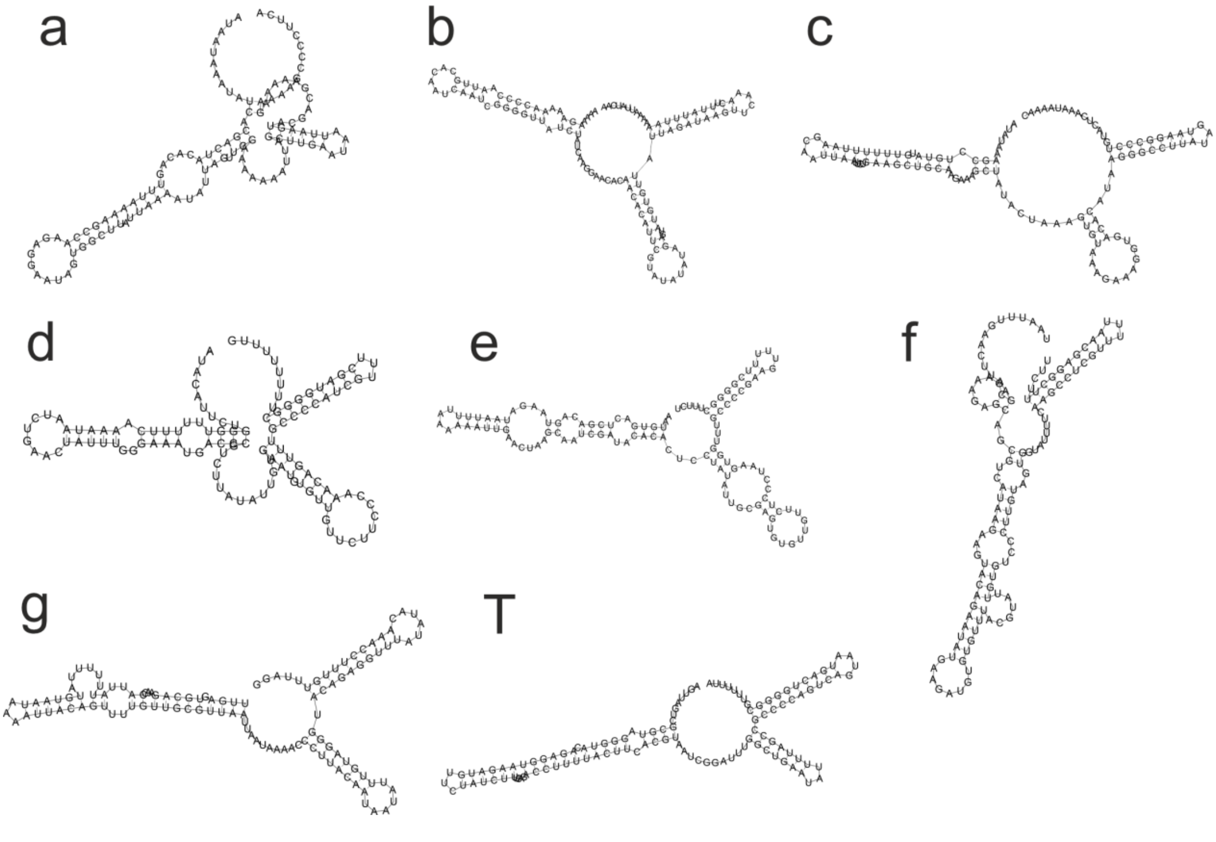


Supplementary Figure S3. Secondary structures of best structure representatives of predicted spot 42 RNA homologs in *Glaciecola*. Species are: *G. sp. MF2* (a), *G. sp. KUL10* (b), *G. sp. 2405UD65-10* (c), *G. sp. 33A* (d), *G. siphonariae* (e), *G. sp. MH2031* (f), *G. sp. HTCC2999* (g). In (T), the *E. coli* spot 42 RNA used as the template in the search.

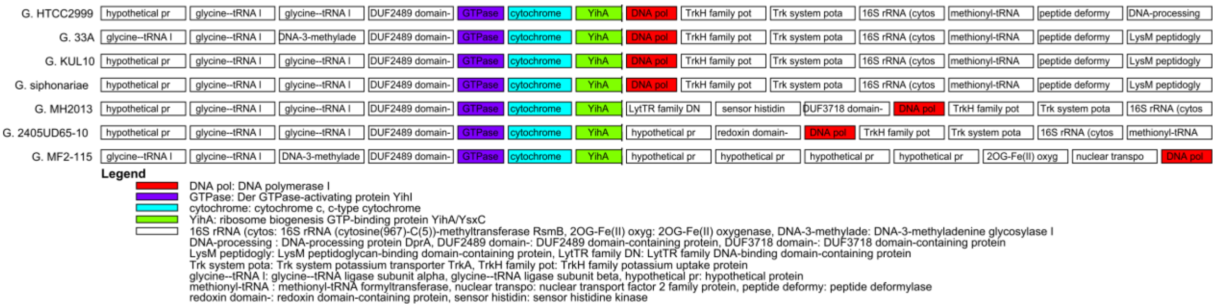


Supplementary Figure S4. Broader genomic contexts of *Glaciecola* spot 42 homologs predicted using the presented approach. The contexts are organized in rows of boxes beneath each other. The boxes mark genes, the spaces between them IGRs. The lengths of both do not correspond to the lengths of genes and IGRs. The boxes contain shortcuts of gene annotations, shown in full in the legend. The contexts consist of 14 genes, 7 genes in each of the genomic directions from the central IGRs with predicted homologs. Colored boxes show genes complying with spot 42 synteny. The genes corresponding to the colors are shown in the legend.

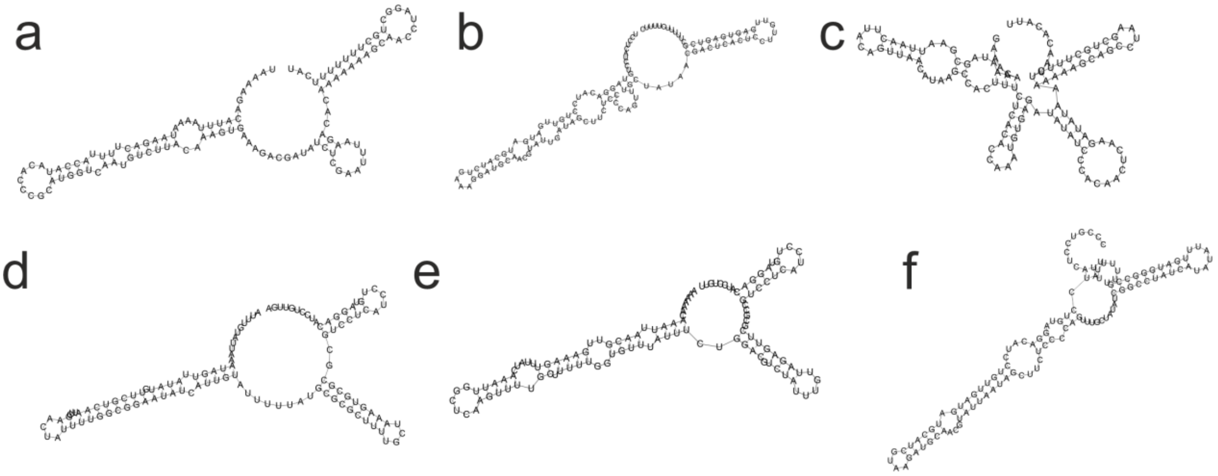


Supplementary Figure S5. Secondary structures of best structure representatives of predicted spot 42 RNA homologs in *Pseudoalteromonas*. Species are: *P. sp. marina* (a), *P. sp. simplex* (b), *P. sp. piratica* (c), *P. sp. atlantica* (d), *P. maricaloris* (e), and *P. sp. haloplanktis* (f). The template used in the search is shown in (T) in Supplementary Figure S3.

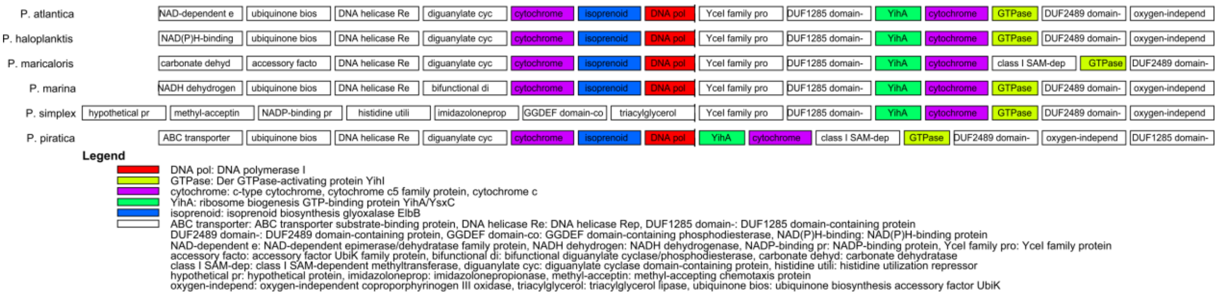


Supplementary Figure S6. Broader genomic contexts of *Pseudoalteromonas* spot 42 homologs predicted using the presented approach. The contexts are organized in rows of boxes beneath each other. The boxes mark genes, the spaces between them IGRs. The lengths of both do not correspond to the lengths of genes and IGRs. The boxes contain shortcuts of gene annotations, shown in full in the legend. The contexts consist of 14 genes, 7 genes in each of the genomic directions from the central IGRs with predicted homologs. Colored boxes show genes complying with spot 42 synteny. The genes corresponding to the colors are shown in the legend.

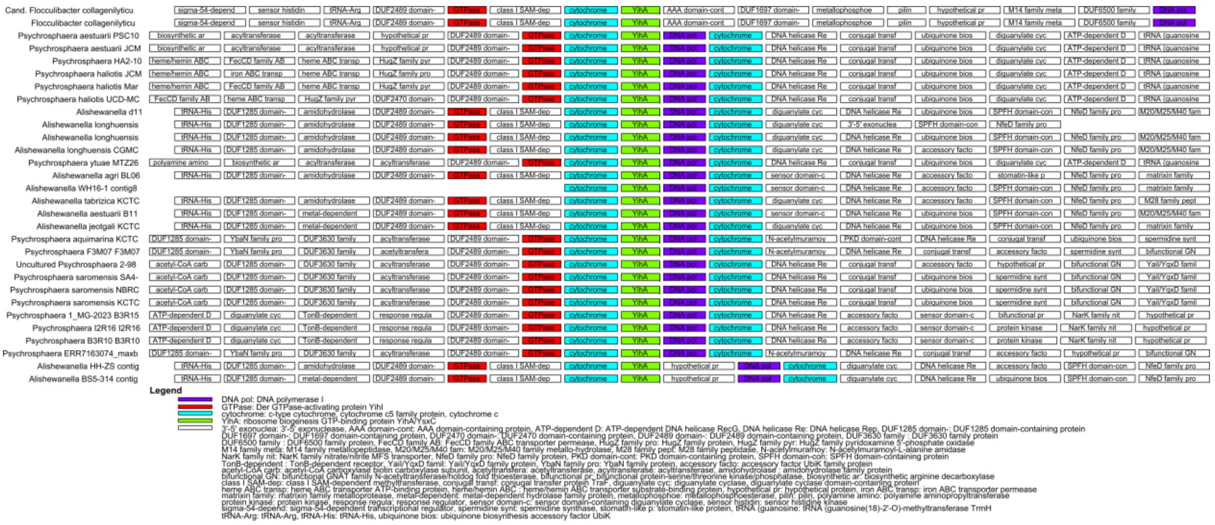

Supplementary Figure S7. spf homologs identified by cascade sequence searches using homologs of *spf* in *Pseudalteromonas* predicted by the presented approach as queries.

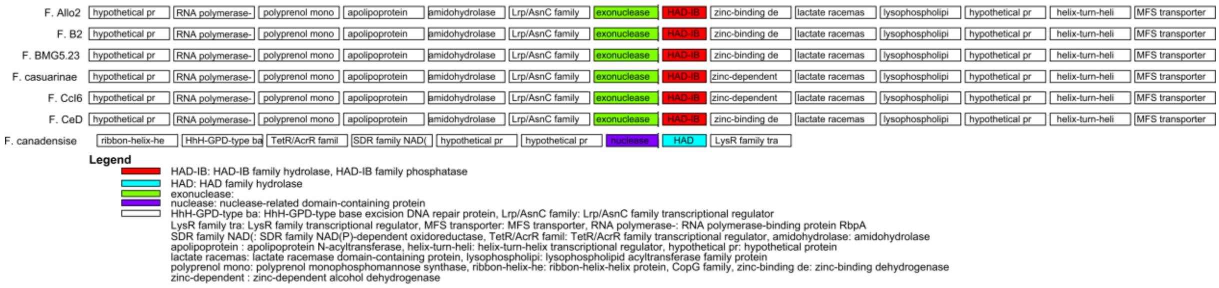


Supplementary Figure S8. *Frankia* HAD-IB-genomic contexts with statistically significant structural candidates to ms1 RNA homologs predicted using the presented approach. The contexts are organized in rows of arrow shapes beneath each other, each for one *Frankia* species indicated above the contexts. The shapes depict genes, the spaces between them are IGRs. The contexts consist of 10 genes, 5 genes in each of the genomic directions from the central IGRs with the candidates, with an exception of the last context formed with 7 genes, 3 and 4 in each of the genomic directions from the central IGRs with the structure candidate. Lengths of the both shapes and spaces correspond to lengths of genes and IGRs. The shapes point in the direction of genomic orientation of the genes. The shapes are labeled by protein IDs and genomic annotations.

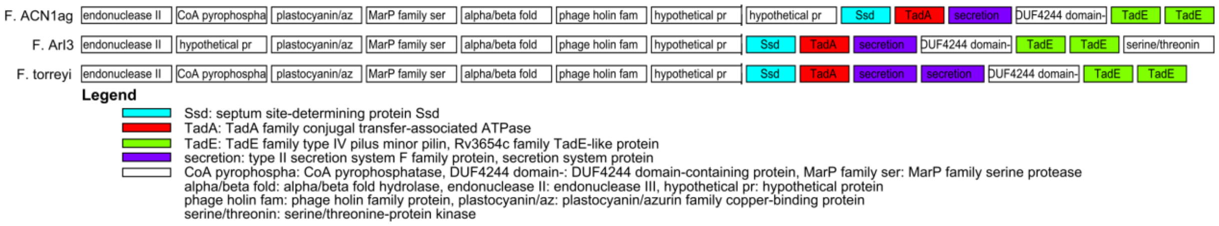


Supplementary Figure S9. *Frankia* septum-genomic contexts with statistically significant structural candidates to ms1 RNA homologs predicted using the presented approach.

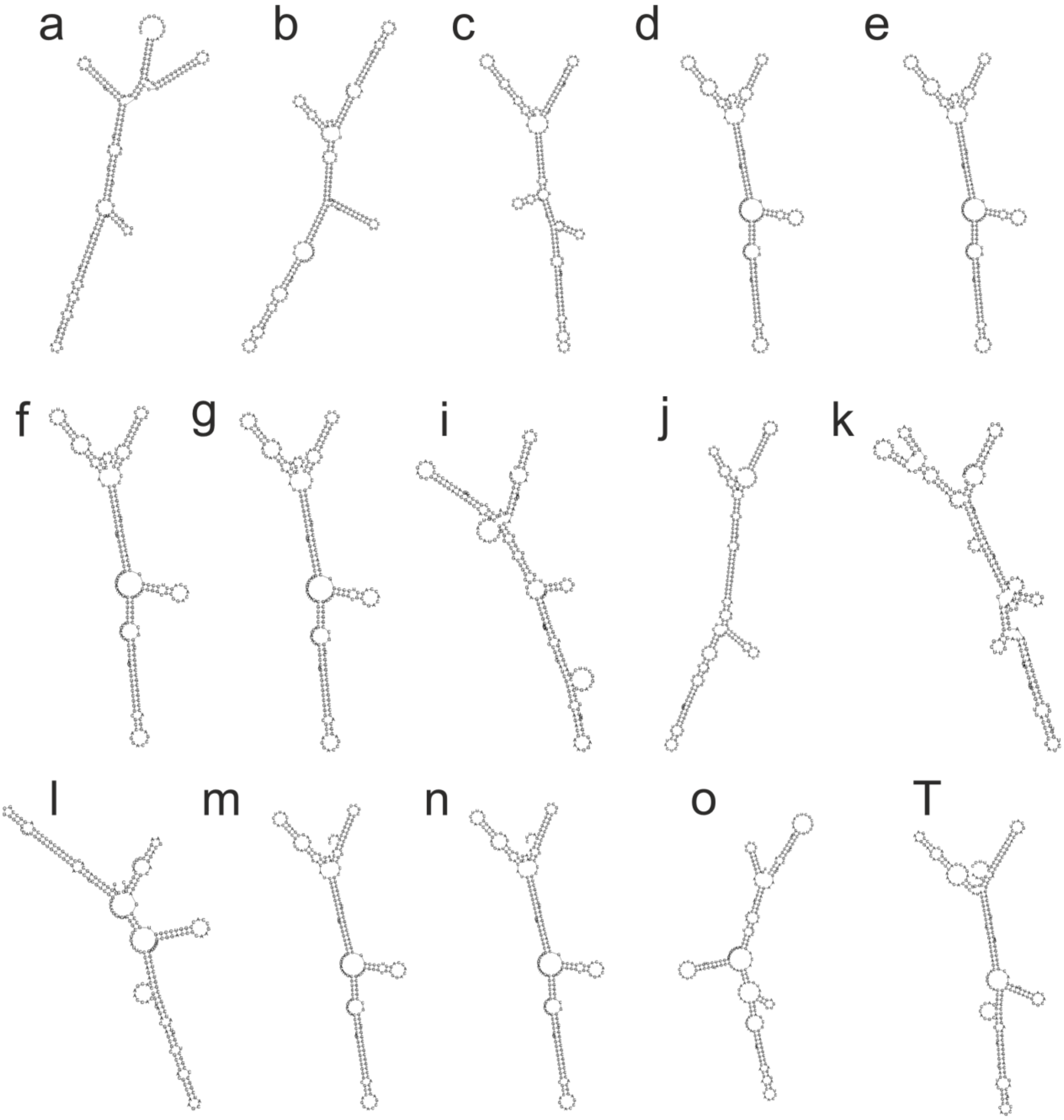


Supplementary Figure S10. Best structure representatives of predicted ms1 RNA homologs in *Frankia*. Species are: *F. nodulisporulans* (a), *F. sp. ArI3* (b), *F. sp. QA3* (c), *F. sp. BMG5.23* (d), *F. casuarinae* (e), *F. sp. Allo2* (f), *F. sp. CcI6* (g), *F. canadensis* (i), *F. sp. ACN1ag* (j), *F. nepalensis* (k), *F. umida* (l), *F. sp. B2* (m), *F. sp. CeD* (n), *F. sp. AvcI1* (o). In (T), the *S. coelicolor* ms1 RNA used as the template in the search.

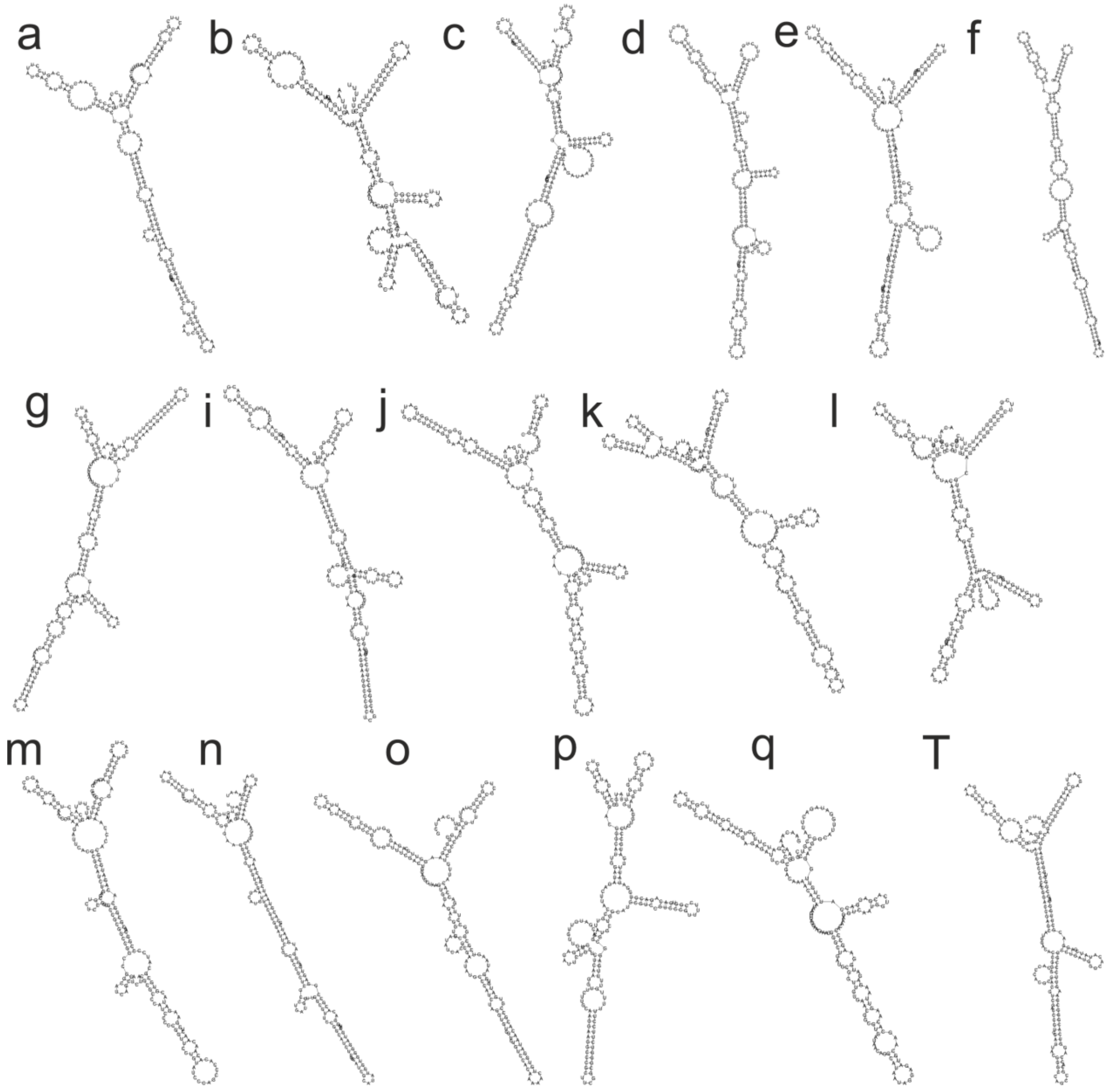


Supplementary Figure S11. Best structure representatives of predicted ms1 RNA homologs in *Bifidobacterium*. Species are: *B. angulatum* (a), *B. pseudolongum* (b), *B. dentium* (c), *B. bombi.23* (d), *B. choerinum* (e), *B. boum* (f), *B. animalis* (g), *B. pseudocatenulatum* (i), *B. tsurumiense* (j), *B. adolescentis* (k), *B. ruminantium* (l), *B. thermacidophilum* (m), *B. moukalabense* (n), *B. bifidum* (o), *B. catenulatum* (p), *B. minimum* (q). In (T), the *S. coelicolor* ms1 RNA used as the template in the search.
