## Supplementary tables for "Integrating suboptimal secondary structures, AI-assisted genomic synteny, and evolutionary conservation to identify bacterial ncRNA homologs beyond sequence similarity"

Supplementary Table S1. Test RNAs. Family names, species, sources of structures used as templates in the test searches, and sequence lengths are shown.

| ncRNA family | Species | Source of template structure | Sequence length (nts) |
|---|---|---|---|
| SRP bact large | *E. coli* | SRPDB {Andersen, 2006 #210} | 114 |
| SRP bact small | *E. coli* | RF00169 {Kalvari, 2021 #690} | 98 |
| 6S | *E. coli* | {Barrick, 2005 #218} | 184 |
| 5S | *E. coli* | PDB ID 1C2X | 120 |
| RsmZ | *P. aueruginosa* | PDB ID 7YR7 | |
| oxys | *E. coli* | {Stih, 2023 #680} | 109 |
| spot42 | *E. coli* | {Baekkedal, 2015 #679} | 119 |
| ms1 | *M. smegmatis* | {Panek, 2011 #603} | 304 |
| gcvb | *S. typhimurium* | {Sharma, 2007 #681} | 206 |
| DsrA | *E. coli* | {Wu, 2017 #682} | 87 |
| tRNA Thr | *E. coli* | PDB ID 1QF6 | 76 |
| RNaseP bact a | *E. coli, T. tencongensis* | PDB IDs 7UO1, 3q1r | 373, 347 |
| 16S RNA | *E. coli* | multiple PDB IDs, RNAfold prediction | 1542 |
| micF | *E. coli* | {Delihas, 2001 #686;Schmidt, 1995 #687} | 95 |
| ryhB | *E. coli* | {Chen, 2021 #685} | 90 |

Supplementary Table S2. Test searches in details.

| ncRNA | Species | Template [1] | Template length (nts) | Sliding window length (nts) [2] | Subject species | Sequence similarity to template [3] | Rfam [4] | Subject genome ID |
|---|---|---|---|---|---|---|---|---|
| SRP bact small/large | *E. coli* | SRP_bact_small_esccol3d-97-11-17-stretched.pdb | 114 | | *Mycobacterium avium* (SRP bact small) | - | + | NZ_CP018019.2 |
| | | | | | *Thermotoga profunda* (SRP bact large) | 59%/4e-10 | + | NZ_AP014510.1 |
| | | | | | *Thermotoga caldifontis* (SRP bact large) | 40%/1e-05 | + | NZ_AP014509.1 |
| | | | | | *Thermus thermophilus (SRP bact small)* | 25%/5e-04 | | NC_006461.1 |

| | | | | | | | | |
|---|---|---|---|---|---|---|---|---|
| | | SRP_bact_small_Eco_RF00169.rf | 98 | | | | ✓ | |
| 6S | *E. coli* | 6S_Ec | 184 | | *Bacillus subtilis* | - | ✓ | NC_000964.3 |
| | | | | | *Burkholderia seminalis* | - | ✓ | NZ_CP013398.1 |
| | | | | 165 | *Rhizobium etli* | | ✓ | |
| 5S | *E. coli* | 5S_Ec_1C2X | 120 | | *Pyrococcus furiosus* | - | ✓ | NZ_CP023154.1 |
| | | | | | *Synechococcus elongatus* | - | + | NZ_CP085785.1 |
| RsmZ | *P. aeruginosa* | RsmZ_Pae_7YR7 | 118 | | *Halopseudomonas pachastrellae* | - | ✓ | NZ_FOUD01000001.1 |
| | | | | | *Azotobacter vinelandii* | 33%/0.024 | ✓ | NC_012560.1 |
| | | RsmZ_Pae_7YR7.rf | | | *Azomonas agilis* | - | ✓ | NZ_VLKG01000004.1 |
| oxys | *E. coli* | oxys_Ec | 109 | | *Trabulsiella odontotermitis* | 87%/9e-24 | ✓ | NZ_CP125781.1 |
| | | | | | *Lelliottia nimipressuralis* | 70%/2e-14 | - | NZ_CP167172.1 |
| | | | | | *Yokenella regensburgei* | 87%/9e-18 | + | NZ_JH417859.1 |
| spot42 | *E. coli* | spot42_Eco | 119 | | *Arsukibacterium perlucidum* | 29%/5e-09 | - | NZ_KB907701.1 |
| | | | | | *Arsukibacterium tuosuense* | 35%/2e-10 | - | NZ_OBEB01000016.1 |
| | | | | | *Vibrio ishigakensis* | 98%/6e-18 | + | NZ_AP024881.1 |
| | | | | | *Photobacterium aquae* | 76%/2e-23 | + | NZ_LDOT01000004.1 |
| ms1 | *M. smegmatis* | Msmest_n_7 | 304 | 230 | *Actinosynnema pretiosum* | 25%/3e-05 | - | CP029607.1 |
| | | | | 230 | *Saccharothrix coeruleofusca* | 24%/0.019 | - | NZ_JAGINV010000001.1 |
| | | | | 230 | *Streptomyces violaceolatus* | - | - | NZ_BAAASK010000002.1 |
| | | | | 230 | *Actinoalloteichus spitiensis* | - | - | NZ_AGVX02000789.1 |
| | | | | | *Gordonia effusa* | 75%/3e-12 | - | NZ_BAEH01000067.1 |
| | | | | | *Gordonia araii* | 61%/4e-16 | - | NZ_JABELY010000001.1 |
| | | | | | *Rhodococcus gannanensis* | 42%/9e-06 | + | NZ_JBHUFB010000022.1 |
| | | | | | *Aldersonia kunmingensis* | 41%/6e-08 | | NZ_LRRB01000288.1 |
| | | | | | *Gordonia polyisoprenivorans* | 26%/0.017 | - | NC_016906.1 |
| gcvb | *S. typhim* | gcvb_St | 206 | | *Haemophilus influenzae* | - | ✓ | NZ_CP085952.1 |
| | | | | | *Photobacterium aquae* | 66%/0.042 | + | NZ_LDOT01000051.1 |

| | | | | | | | | |
|---|---|---|---|---|---|---|---|---|
| | *urium* | gcvb_St.rf | | | *Haemophilus influenzae* | - | ✓ | NZ_CP085952.1 |
| | | | | | *Photobacterium aquae* | 66%/0.042 | + | NZ_LDOT01000051.1 |
| DsrA | *E. coli* | DsrA_Eco_paper | 87 | | *Cedecea neteri* | 83%/3e-19 | + | NZ_CP023525.1 |
| | | | | | *Cedecea davisae* | 83%/7e-13 | + | NZ_CP172054.1 |
| | | | | | *Citrobacter gillenii* | 90%/1e-22 | + | NZ_SUQN01000005.1 |
| tRNA | *E. coli* | tRNA_Thr_Eco_1QF 6 | 76 | | *Thermotoga maritima* | 84%/4e-09 | ✓ | NC_023151.1 |
| | | | | | *Streptomyces halobius* | 72%/2e-06 | + | NZ_CP086322.1 |
| Rnase P bact a | *E. coli* | RNaseP_bact_a_Ec_ 7UO1.rf | 373 | | *Thermotoga profunda* | 68% 4e-09 | + | NZ_AP014510.1 |
| | | | | | *Thermotoga caldifontis* | 47% 6e-07 | + | NZ_AP014509.1 |
| | | | | | *Pseudofrankia saprophytica* | 53% 1e-27 | - | NZ_KI912266.1 |
| | | | | | *Amedibacillus hominis* | 81% 3e-61 | - | NZ_JAKVPQ010000045.1 |
| 16S RNA | *E. coli* | 16S_Ec.rf | 1542 | | *Thermotoga maritima* | 100%/0 | ✓ | NC_023151.1 |
| micF | *E. coli* | micF_Ec | 95 | | *S. enterica* | 100%/7e-39 | ✓ | NC_003197.2 |
| | | | | | *S. bongori* | 98%/ 2e-40 | + | NC_015761.1 |
| ryhB | *E. coli* | ryhB_Ec | 90 | | *V. cholerae eltor* | 30%/2e-05 | ✓ | NC_002506.1 |
| | | | | | *V. cholerae RFB16* | 30%/1e-04 | + | NZ_CP043554.1 |

[1] Template structures are included in Supplementary File S1.fa

[2] Shown only if differs from the length of template ncRNAs shown in Table 1

[3] BLAST Coverage(%)/Expect of hits of templates in inspected IGRs with the sought RNA homologs . '-' for no significant similarity found. The BLAST parameters were: program: blastn, Word size: 11, Expect value: 0.05, Hitlist size: 100, Match/Mismatch scores: 2,-3, Gapcosts: 5,2, Low Complexity Filter: Yes, Filter string: L;m

[4] '✓' for the identified homolog is in Rfam, '+' it is not in Rfam, but other species of the genus, '-' it is not in Rfam, but in literature cited in the Table S1

Supplementary Table S3. Species in use cases.

| Query ncRNA | Subject species | Genomic ID |
|---|---|---|

| | | |
|---|---|---|
| *E. coli* spot 42 | *Glaciecola siphonariae*<br>*Glaciecola sp. KUL10*<br>*Glaciecola sp. HTCC2999*<br>*Glaciecola sp. 33A*<br>*Glaciecola sp. MH2013*<br>*Glaciecola sp. 2405UD65-10*<br>*Glaciecola sp. MF2-115* | NZ_JBHSGU010000001.1<br>NZ_BGNG01000001.1<br>NZ_DS989817.1<br>NZ_PJAR01000044.1<br>NZ_JADNRN010000001.1<br>NZ_JBMEUF010000001.1<br>NZ_JBLBVM010000001.1 |
| | *Pseudoalteromonas haloplanktis*<br>*Pseudoalteromonas maricaloris*<br>*Pseudoalteromonas simplex*<br>*Pseudoalteromonas atlantica*<br>*Pseudoalteromonas marina*<br>*Pseudoalteromonas piratica* | NZ_CAMAPB010000001.1<br>NZ_CP024625.1<br>NZ_JADILN010000001.1<br>NZ_CP023464.1<br>NZ_CP023558.1<br>NZ_CP009888.1 |
| *S. coelicolor* ms1 | *Frankia sp. RB7*<br>*Frankia sp. Hr75.2*<br>*Frankia casuarinae*<br>*Frankia alni ACN14a*<br>*Frankia sp. QA3*<br>*Frankia sp. CcI6*<br>*Frankia sp. BMG5.23*<br>*Frankia sp. CeD*<br>*Frankia sp. Allo2*<br>*Frankia torreyi*<br>*Frankia sp. R43*<br>*Frankia sp. ACN1ag*<br>*Frankia sp. AvcI1*<br>*Frankia sp. EI5c*<br>*Frankia sp. KB5*<br>*Frankia sp. B2*<br>*Frankia nepalensis* | JAUEPE010000058.1<br>CASGGV010000001.1<br>NC_007777.1<br>NC_008278.1<br>NZ_CM001489.1<br>NZ_AYTZ01000001.1<br>NZ_JDWE01000001.1<br>NZ_JPGU01000001.1<br>NZ_JPHT01000001.1<br>NZ_JYFN01000001.1<br>NZ_LFCW01000001.1<br>NZ_LJPA01000001.1<br>NZ_LJFZ01000001.1<br>NZ_LRTK01000001.1<br>NZ_MRUJ01000001.1<br>NZ_SOPN01000100.1<br>NZ_JADWYU010000127.1 |

| | | |
|---|---|---|
| | *Frankia sp. ArI3* | NZ_CP079862.1 |
| | *Frankia umida* | NZ_JALKFT010000100.1 |
| | *Frankia gtarii* | NZ_JAMPTM010000100.1 |
| | *Frankia tisae* | NZ_JANEZT010000100.1 |
| | *Frankia canadensis* | NZ_FZMO01000568.1 |
| | *Candidatus Frankia alpina* | NZ_CADCWT010000001. |
| | *Candidatus Frankia nod-* | 1 |
| | *ulisporulans* | NZ_CADDZU010000001.1 |
| | *Frankia sp. Cj5* | NZ_CAKJTO010000001.1 |
| | *Bifidobacterium longum* | NZ_GG666849 |
| | *Bifidobacterium adolescentis* | NC_008618 |
| | *Bifidobacterium animalis* | NC_011835 |
| | *Bifidobacterium dentium* | NC_013714 |
| | *Bifidobacterium angulatum* | NZ_GG663540 |
| | *Bifidobacterium bifidum* | NC_014616 |
| | *Bifidobacterium gallicum* | NZ_ABXB03000027 |
| | *Bifidobacterium catenulatum* | NZ_ABXY01000031 |
| | *Bifidobacterium thermophilum* | NC_020546 |
| | *Bifidobacterium magnum* | NZ_ATVE01000001 |
| | *Bifidobacterium* | NZ_ATWW01000001 |
| | *pseudolongum* | NZ_ATXM01000001 |
| | *Bifidobacterium minimum* | NZ_AUFH01000001 |
| | *Bifidobacterium subtile* | NZ_AUFI01000006 |
| | *Bifidobacterium* | NZ_AUCL01000001 |
| | *thermacidophilum* | NZ_AXVO01000001 |
| | *Bifidobacterium tsurumiense* | NZ_AWUO01000001 |
| | *Bifidobacterium choerinum* | NZ_AZMV01000001 |
| | *Bifidobacterium sp* | NZ_JHWO01000001 |
| | *Bifidobacterium moukalabense* | NZ_JHWQ01000001 |
| | *Bifidobacterium boum* | NZ_ATLK01000001 |
| | *Bifidobacterium ruminantium* | NZ_JGZF01000001 |
| | *Bifidobacterium bombi* | |

| | |
|---|---|
| | *Bifidobacterium pseudocatenulatum* |
