## Supplementary files for "Integrating suboptimal secondary structures, AI-assisted genomic synteny, and evolutionary conservation to identify bacterial ncRNA homologs beyond sequence similarity"

Supplementary File S1. Sequences of the test query RNAs. Their secondary structures were used as templates and their homologs were identified in test searches.

```
>SRP_bact_small_esccol3d-97-11-17-stretched.pdb
gGGGGCUCUGUUGGUUCUCCCGCAACGCUACUCUGUUUACCAGGUCAGGUCCGGAAGGAAGCAGCCAAGGCAG
AUGACGCGUGUGCCGGGAUGUAGCUGGCAGGGCCCCCACCC
((((((((((((((((.(((((..((((...(((((......(((....(((....)))....)))...))))
)....))))....)))))...))))))))))))))))....
>SRP_bact_small_Eco_RF00169.rf
UGUUGGUUCUCCCGCAACGCUACUCUGUUUACCAGGUCAGGUCCGGAAGGAAGCAGCCAAGGCAGAUGACGCG
UGUGCCGGGAUGUAGCUGGCAGGGC
((((((((((((((((((((..((((((((....(((....(((....)))....))).))))))).)..)))
).)).))))).).))))))))....
>6S_Ec
AUUUCUCUGAGAUGUUCGCAAGCGGGCCAGUCCCCUGAGCCGAUAUUUCAUACCACAAGAAUGUGGCGCUCCG
CGGUUGGUGAGCAUGCUCGGUCCGUCCGAGAAGCCUUAAAACUGCGACGACACAUUCACCUUGAACCAAGGGU
UCAAGGGUUACAGCCUGCGGCGGCAUCUCGGAGAUUCC
...(((((((((((.((((..((((((.....(((((((((.................(((((((.((...((
(((((..((((....(((((.....)))))....)))).))))))).)).)))))))..((((...)))))))
))).))).....)))))).)))))))))))))))....
>5S_Ec_1C2X
UGCCUGGCGGCCGUAGCGCGGUGGUCCCACCUGACCCCAUGCCGAACUCAGAAGUGAAACGCCGUAGCGCCGA
UGGUAGUGUGGGGUCUCCCCAUGCGAGAGUAGGGAACUGCCAGGCAU
((((((((.(.....((.(((.(.......((((.............))))........).)))..)).((..
.....((.((((.....)))).)).......))...).)))))))).
>RsmZ_Pae_7YR7
GCGUACAGGGAACACGCAACCCCGAAGGAUCGGGGAAGGGACGUCGCCAGGGAGGCGAUUCCAUCAGGAUGAU
GACGAGGGACUGAAGAGUGGGCGGGGUAAUACCCCGCCCCUUUUU
((((.........))))..(((((......)))))........(((.(......))))..(.((.......))
)...........(((((.((((((((.....))))))))))))).
>RsmZ_Pae_7YR7.rf
GCGUACAGGGAACACGCAACCCCGAAGGAUCGGGGAAGGGACGUCGCCAGGGAGGCGAUUCCAUCAGGAUGAU
GACGAGGGACUGAAGAGUGGGCGGGGUAAUACCCCGCCCCUUUUU
((((.........))))..((((((....))))))...(((.((((((.....))))))))).((((......
.........))))((((.(((((((((...))))))))))))).. 
>oxys_Ec
GAAACGGAGCGGCACCUCUUUUAACCCUUGAAGUCACUGCCCGUUUCGAGAGUUUCUCAACUCGAAUAACUAA
AGCCAACGUGAACUUUUGCGGAUCUCCAGGAUCCGC
(((((((.((((.((.((...........)).))..)))))))))))..((((......))))..........
.................(((((((.....)))))))
>spot42_Eco
AGUUAGUCGCGUAGGGUACAGAGGUAAGAUGUUCUAUCUUUCAGACCUUUUACUUCACGUAAUCGGAUUUGGC
UGAAUAUUUUAGCCGCCCCAGUCAGUAAUGACUGGGGCGUUUUUUA
........((((.((((.(((((((((((((...))))))....))))))))))).))))..........(((
((((...)))))))((((((((((....))))))))))........
>Msmest_n_7
ACGGAAGCUUGGCGAGGCCAAGACCCAGCCGGAAGAGAAGGCUAGAUCUCCCGACCCAAGCUCCUAGCACGGA
UACCGAGCACCCACGCGGAGCACAUGCCGCGGAAUAGGCAAAAGUGUUGCGGACCUGCGUAGUUUCGAAAAGC
GGACGGCCACGACGGCCCUUUGGGUGGGGUUGCAGCCGUAGCGCAUCGCAAAGACGCCGAGGUCACCCACGCA
ACCCACAUCGCACGCUUGGUCACUCGGGGUCCGUGCUAGCGGGCGGCGAACCGGACAUUGUCCGGGACGCCGC
CCGAUUUCUUUU
..(((.((((((...((...(((...((((.........))))...))).))...)))))))))(((((((((
..(((((...(((.(((..((...((((........))))...((((....(((((((((..((.(((...((
(.(((((..((((.((((...))))...))))..)))))..))).))).))..))))..)))))....)))).
.........)).))).)))...)))))..))))))))).(((((((((..((((((...))))))..))))))
))).........
>gcvb_St
```

ACUUCCUGAGCCGGAACGAAAAGUUUUAUCGGAAUGCGUGUUCUGAUGGGCUUUUGGCUUACGGUUGUGAUGU
UGUGUUGUUGUGUUUGCAAUUGGUCUGCGAUUCAGACCAUGGUAGCGAGACUACCUUUUUCACUUCCUGUAC
AUUUACCCUGUCUGUCCAUAGUGAUUAAUGUAGCACCGCCAUAUUGCGGUGCUUUUUUUU
....(((((((((((......(((((.(((((((......))))))))))))))))))))).)).........
....................(((((((.....))))))).(((((.....)))))).................(
(((.......................)))).((((((((......)))))))).......
>gcvb_St.rf
ACUUCCUGAGCCGGAACGAAAAGUUUUAUCGGAAUGCGUGUUCUGAUGGGCUUUUGGCUUACGGUUGUGAUGU
UGUGUUGUUGUGUUUGCAAUUGGUCUGCGAUUCAGACCAUGGUAGCGAGACUACCUUUUUCACUUCCUGUAC
AUUUACCCUGUCUGUCCAUAGUGAUUAAUGUAGCACCGCCAUAUUGCGGUGCUUUUUUUU
......(((((((((......(((((.((((((((....)))))))))))))))))))))).((..((((...
...(((((.......)))))(((((((.....))))))).(((((.....))))).....))))..))..(((
(((...(((((......)))).)...))))))(((((((......)))))))........
>DsrA_Eco_paper
AACACAUCAGAUUUCCUGGUGUAACGAAUUUUUAAGUGCUUCUUGCUUAAGCAAGUUUCAUCCCGACCCCCU
CAGGGUCGGGAUUU
...(((((((.....)))))))............(((..((((((....))).)))))).((((((((((...
..))))))))))..
>tRNA_Thr_Eco_1QF6
gCCGAUAUAGCUCAGuuGGuAGAGCAGCGCAUUCGUaAUGCGAAGgUCGUAGGuuCGACUCCUAUUAUCGGCA
CCA
(((((((..((((........))))..((((.......))))......(((((.......)))))))))))).
...
>RNaseP_bact_a_Ec_7UO1.rf
gAAGCUGACCAGACAGUCGCCGCUUCGUCGUCCUCUUCGGGGGAGACGGGCGGAGGGGAGGAAAGUCCGG
GCUCCAUAGGGcAGGGUGCCAGGUAACGCCUGGGGGGGAAACCCACGACCAGUGCAACAGAGAGCAAACCGCC
GAUGGCCCGCGCAAGCGGGAUCAGGUAAgGGUGAAAGGGUGCGGUAAGAGCGCACCGCGCGGCUGGUAACAGU
CCGUGGCACGGUAAACUCCACCCGGAGCAAGGCCAAAUAGGGGUUCAUAAgGUACGGCCCGUACUGAACCCGG
GUAGGCUGCUUGAGCCAGUGAGCGAUUGCUGGCCUAGAUGAAUGACUGUCCACGACAGAACCCGGCUUAUCGG
UCAGUUUC
..(.((((((.........((.((((((((((.(((.(....).))))))).)))))).)).....((.((((
((..(........((((((.(((.....))).).((......))..(..........................
.......(((....)).)...................((.((.........)).)).....(...(...)...
.)...............))))))).)...((((.........(........(((.....)))....).....
...))))....(.(((((.((....)).))))))..........................)))))))....))
)))).)..
>16S_Ec.rf
AAAUUGAAGAGUUUGAUCAUGGCUCAGAUUGAACGCUGGCGGCAGGCCUAACACAUGCAAGUCGAACGGUAAC
AGGAAACAGCUUGCUGUUUCGCUGACGAGUGGCGGACGGGUGAGUAAUGUCUGGGAAACUGCCUGAUGGAGGG
GGAUAACUACUGGAAACGGUAGCUAAUACCGCAUAACGUCGCAAGACCAAAGAGGGGGACCUUCGGGCCUCUU
GCCAUCAGAUGUGCCCAGAUGGGAUUAGCUAGUAGGUGGGGUAACGGCUCACCUAGGCGACGAUCCCUAGCUG
GUCUGAGAGGAUGACCAGCCACACUGGAACUGAGACACGGUCCAGACUCCUACGGGAGGCAGCAGUGGGGAAU
AUUGCACAAUGGGCGCAAGCCUGAUGCAGCCAUGCCGCGUGUAUGAAGAAGGCCUUCGGGUUGUAAAGUACUU
UCAGCGGGGAGGAAGGGAGUAAAGUUAAUACCUUUGCUCAUUGACGUUACCCGCAGAAGAAGCACCGGCUAAC
UCCGUGCCAGCAGCCGCGGUAAUACGGAGGGUGCAAGCGUUAAUCGGAAUUACUGGGCGUAAAGCGCACGCAG
GCGGUUUGUUAAGUCAGAUGUGAAAUCCCCGGGCUCAACCUGGGAACUGCAUCUGAUACUGGCAAGCUUGAGU
CUCGUAGAGGGGGGUAGAAUUCCAGGUGUAGCGGUGAAAUGCGUAGAGAUCUGGAGGAAUACCGGUGGCGAAG
GCGGCCCCCUGGACGAAGACUGACGCUCAGGUGCGAAAGCGUGGGGAGCAAACAGGAUUAGAUACCCUGGUAG
UCCACGCCGUAAACGAUGUCGACUUGGAGGUUGUGCCCUUGAGGCGUGGCUUCCGGAGCUAACGCGUUAAGUC
GACCGCCUGGGGAGUACGGCCGCAAGGUUAAAACUCAAAUGAAUUGACGGGGGCCCGCACAAGCGGUGGAGCA
UGUGGUUUAAUUCGAUGCAACGCGAAGAACCUUACCUGGUCUUGACAUCCACGGAAGUUUUCAGAGAUGAGAA
UGUGCCUUCGGGAACCGUGAGACAGGUGCUGCAUGGCUGUCGUCAGCUCGUGUUGUGAAAUGUUGGGUUAAGU
CCCGCAACGAGCGCAACCCUUAUCCUUUGUUGCCAGCGGUCCGGCCGGGAACUCAAAGGAGACUGCCAGUGAU
AAACUGGAGGAAGGUGGGGAUGACGUCAAGUCAUCAUGGCCCUUACGACCAGGGCUACACACGUGCUACAAUG
GCGCAUACAAAGAGAAGCGACCUCGCGAGAGCAAGCGGACCUCAUAAAGUGCGUCGUAGUCCGGAUUGGAGUC
UGCAACUCGACUCCAUGAAGUCGGAAUCGCUAGUAAUCGUGGAUCAGAAUGCCACGGUGAAUACGUUCCCGGG
CCUUGUACACACCGCCCGUCACACCAUGGGAGUGGGUUGCAAAAGAAGUAGGUAGCUUAACCUUCGGGAGGGC

GCUUACCACUUUGUGAUUCAUGACUGGGGUGAAGUCGUAACAAGGUAACCGUAGGGGAACCUGCGGUUGGAUC
ACCUCCUUA
........((((((...(.(((((..(((....(((((((((((((.((..(((.((..(((.((((((((((
........).))))))))).)))..)).)))..))..(((((((....(((((((..((...(((((((.(.(
(..((.((((((....)))))).))...)).).....(((....))).....((((((.((....))))))))
.)))))))..)).)))))))((((((.(((..((((((((.......)))))))))))...))))))..((((
(((.........)))))))....(((((.(((.....)))))))).((((....))))...((((((.....)
)))))).....((((....))))(((((((((((((.((..........((((.(((((.((((((..((..((
((.(((((..(..((.((((((((.......))))))))).))..)....))))).))))..)).((((.(((.
(((((((..(((.((.((......))..)).)))..))).....)))).)))))))((((...)))).(.(((
(((((((((((.(((((((((....((((.((......)).))))...)))))))))..)))))(((.(.(((
(((((..((((.((((...((((((((.(.(((........)))..).))))))))...))))(((.((....
)).)))))))..))).))))).).)))....((((...(((((((..((...((((.........))))))..
))))))))))).(((((.((......)).)))))...(((((.(((((((((...))))).)))).)))))..
))))))))).)((((..((((....))))...))))....((((((((.(.(((((((.......)))).)).
).).).)))))))..)))))).)))))..))))(((((.(((((...(((.((((.(((((((....))))).
...)).)))))))....))))))))))))).))))))))).)))).((((((.........(((.(((......
))))))))))))......))))))))))))))))))))))).)))))))))))).((((((.((.(((((...
..)))))))...(((.(.((((((.....))))))).).))).(((((((..(((((((..(((((((.....(
((((............))).))((((........)))).........))))))).))))))).((((((((((
........)))))))....)))((((.((.((...(((((((.........)))))))...)))))))))(((
(..((....))..)))))((((.((..((..((((((((((((.(..((((((.((((..((...))..))))
))))))..)))))))))))))..)).))))))).))))))).....((((((((((....))))))))))....
..))))))).
>micF_Ec
CGCUAUCAUCAUUAACUUUAUUUAUUACCGUCAUUCAUUUCUGAAUGUCUGUUUACCCCUAUUUCAACCGGAU
GCCUCGCAUUCGGUUUUUUUUA
............................((.((((((....))))))..))..............((((((((
((...))))))))))).......
>ryhB_Ec
GCGAUCAGGAAGACCCUCGCGGAGAACCUGAAAGCACGACAUUGCUCACAUUGCUUCCAGUAUUACUUAGCCA
GCCGGGUGCUGGCUUUU
((((...((.....)))))).......(((.(((((...............))))).)))........(((((
((.....)))))))...

Supplementary File S2. Sequences and secondary structures in dot-bracket format of predicted spot 42 RNA homologs in Glaciecola.

>38_3053771_1_3064242_Glaciecola_sp._HTCC2999_scf_1100007007377,_igr489.f
a_1_l=119_46
UUGAGUGCAGAACCAUUAUUUUUUAUGUAAUAAAAUUACAGUUUUGUUGCGUUAAUUAAUAAAACCCUUACAA
UAAUAUUUGUAGGGUACAGAGGUUUAUAUACAAACCUUUGUUUAGG
((((.(((((...((..((......((((((...))))))))..)))))))))))..........((((((((
......)))))))).((((((((((......))))))))))).....
>32_51040_1_61985_Glaciecola_sp._MH2013_contig1,_igr2065.fa_1_l=119_6
UAAUUUGAACUUAAGCGAAAGAGCAGCGUCAUAAGAAGUACAGAAUAUGAAGAUGUGUGUUUACGUAUUGUGU
CCCUUGUAGUGGUAUUUUCAAGCCUCGUUUUUAACGAGGCUUUCUU
..............((......))..((..((((...(((((((((((........))))))......)))).
..))))))..)).......((((((((((...))))))))))....
>8_6858_1_17951_Glaciecola_sp._2405UD65-10_2405UD65-
10_1,_igr1593.fa_1_l=119_121
AUAUAAAGCCUGUAUGUUUUUUAAGCAAUUAAUUCUGAAGCUGCAAGAAAGCUAUACUAAAGUGUAAAGAAAG
GUGACACAUAAGGGCCUUAUAGUAAGGCCCUGUACUCAAAUAAAAC
......(((.((((.(((((((((....))))....))))))))).....)))........((((........
...))))...(((((((((...)))))))))...............
>2_20316_1_40560_Glaciecola_sp._MF2-115_contig_1,_igr367.fa_1_l=119_16
AUAAUAAAUAUCACGACUACACAGUUUAAAAGCCAAGAGGAAUAGUGGCUUAUUAAAAUAUUAGUUCGAAAAA
AAUUAGCUUGAAUAAUUAAGUACGACGAAAAAGAAAAGCCCCUUCA

..........((.((((((.....(((((((((((..........)))))).)))))....))).))).....
.....((((((....))))))...........))............
>4_47110_1_61287_Glaciecola_sp._KUL10_contig01,_igr953.fa_1_l=119_36
AAAAAGAAAAAACCCCAAUUGCACAAUCAAUCGGGGUUAUCUAUCAAGGAACACAACACAUUCGUAUAUAGA
GUAUGUGUUAUUAGAUAAGUUCAAACUUUAUUUAAAAAAUUAUCAA
....(((..((((((.((((......)))).)))))).)))............(((((((((.........))
..))))))).((((((((((....)).))))))))...........
>24_73801_1_86777_Glaciecola_sp._33A_contig1,_rc_igr2350.fa_1_l=119_96
AUACAUUCUGGGUUUUUUCAAAAUAAUCUGAACUAUUUGGGAAAUGACCCGUCUUAUAUUGUGAAUGUGUUGU
UCUUCCCAAACAGUUUGCCCCAUCGUUUCGAUGGGGCUUUUUUUUG
.........((((..((((.(((((........)))))..))))..))))............((((.((((..
........))))))))(((((((((...))))))))).........
>48_16840_1_925_Glaciecola_siphonariae_strain_CCUG_igr178.fa_1_l=119_13
AAUGUGACUCGACAGUAAGAUAAUUUUAAAAAAUUGAACUAAGCAAUCGAUACACACUCCUAUAUUGCGAGUG
UGUUGUUCUCCCUAAGUGGUUUGCCCCGAAGUUUUUCGGGGCUUUCU
..((((..((((..((....((((((....))))))......))..))))..))))...((((.....(((..
.......))).....))))...((((((((....))))))))).....

Supplementary File S3. Evolutionary conservation of predicted Glaciecola spot 42 homologs in related species. The lines contains names of species of the spot 42 candidates, introduced by “Cand.”, lists of species in which statistically significant BLAST hits of the candidate sequences in spot 42-syntenic IGRs were obtained follow the arrows.

Cand. Glaciecola HTCC2999 scf_110000 -> Opacimonas viscosa LMIT007, Aliiglaciecola M165 scaffold11, Paraglaciecola _36 , Paraglaciecola metabat_bins.17, Paraglaciecola chathamensis S1, Plancto marinum K7, Paraglaciecola polaris LMG, Opacimonas immobilis LMIT016, Paraglaciecola chathamensis KC, Paraglaciecola chathamensis IB, Alteromonas M12, Paraglaciecola Bin_40470_ID136, Paraglaciecola ERR599341_binne, Paraglaciecola polaris ERR2762, Aliiglaciecola NS0011-25_25, Plancto marinum KCTC, Paraglaciecola chathamensis KC, Plancto marinum ena-yuan-GCF, Paraglaciecola chathamensis UB, Paraglaciecola mesophila GPM4, Plancto R6 contig41, Paraglaciecola chathamensis A3, Aliiglaciecola lipolytica D2R0, Paraglaciecola chathamensis E2, Paraglaciecola chathamensis A1, Aliiglaciecola LC.122.F.C4_512, Alteromonas LMIT006, Aliiglaciecola SL4 16, Paraglaciecola agarilytica NO2, Plancto marinum Dej080120, Aliiglaciecola 1_MG-2023 B3R02, Aliiglaciecola 2_MG-2023 B3R02, Bowmanella yangjiangensis Y26, Alteromonas facilis P0213, Paraglaciecola mesophila KMM, Alishewanella maricola 16-MA, Bowmanella yangjiangensis Y57, Paraglaciecola G1-13, Alishewanella HL-SH06, Alishewanella HL-SH05, Paraglaciecola mesophila AS12, Alishewanella SMS8 NG, Paraglaciecola T6c, Aliiglaciecola LCG003.
Cand. Glaciecola MF2-115 contig_35 -> Brumicola nitratireducens FR10, Glaciecola 33A contig6, Brumicola pallidula DSM, Brumicola pallidula DSM.
Cand. Glaciecola KUL10 contig17 -> Agaribacter flavus KCTC, Glaciecola SC05 scaffold15, Glaciecola G7-5.
Cand. Glaciecola 33A contig6 -> Brumicola pallidula DSM, Brumicola pallidula DSM, Brumicola nitratireducens FR10, Alteromonas W364 14, Brumicola blandensis W409, Glaciecola MF2-115 contig, Glaciecola SC05 scaffold15, Glaciecola G7-5, Glaciecola 2405UD65-10 2405UD6, Glaciecola siphonariae CCUG.
Cand. Glaciecola siphonariae CCUG 60 -> Glaciecola SC05 scaffold15, Glaciecola 1036 NODE, Glaciecola XM2 124, Agaribacter marinus NBRC, Agaribacter flavus KCTC.

Supplementary File S4. Sequences and secondary structures in dot-bracket format of predicted spot 42 RNA homologs in Pseudalteromonas.

>28_14081_1_30893_Pseudoalteromonas_haloplanktis_strain_CIP_rc_igr446.fa_1_l=119_211
CCCGUCCUCAUCCUGUAGGACAUCCUGUUGAUGAUGCAUCGUAAGAUGCAACGUAUUAAUAGCUUCUCCCCAG
UUUGCUAUAUCGGCCUAUCAUAUAUUUGAUGGGCCGUUUUUUUAUU
.........((.(((..(((....(((((((((.((((((....))))))...)))))))))....))).)))
..........((((((((((......))))))))))......)).
>41_276997_1_289917_Pseudoalteromonas_maricaloris_strain_SCSIO_rc_igr184.fa_1_l=119_128
AAAAAACAAAUUAACGUUGAAAGUUUAUCAAAUUGGCUCAAGUUUUGCUUUUUGGUGUUUAUUUCUGGACGUC
UAUUUGUUAGAGUUCGCGCCGUCCUCAUCCUGUAGGACAUCCUGUU
.......((((.((((..((((((.....(((((......))))).)).))))..)))).))))..((((.((
((.....)))))))).....((((((.....).)))))........
>14_81742_1_91213_Pseudoalteromonas_atlantica_strain_ECSMB14104_rc_igr202.fa_1_l=119_141
AUUUGUAUUAAAUAGUUAUAUGUUCGUCAAUUCGAACUAUUUUGGCGGAAUAUCAUUGUAUUUUUAUGCGCGC
UUUUGCUAAAGUGCGCGCGUCCUCAUCCUGUAGGACAUCCUGUUGA
...........(((((.((((.((((((((...((....)))))))))))))).))))).........((((((
(((....)))))))))..((((((.....).))))).........
>11_26343_1_16610_Pseudoalteromonas_piratica_strain_OCN003_rc_igr246.fa_1_l=119_441
GAAACAAAAUAGCGAAUUAACUUACAGUUAACAUAAGCCACUUUCUCACACCAAAUGUGAGAUAUAUCCCACA
ACUCAAGAUAUAAAAAAAGCAGCCUAAGCUGCUUUUUCAACACAUU
.....(((...((...(((((.....)))))).....))...)))((((((.....)))))).((((((.....
......)))))).((((((((((....))))))))))........
>7_1923325_1_1939262_Pseudoalteromonas_marina_strain_ECSMB14103_rc_igr1964.fa_1_l=119_581
UAAAAGACAUUUAAAAUAAGACUUUUACCAUACACCCGCAUGGUCAAUGUCUUACAAAGUGAAAGACGAUAUC
UCGAAUUUAAGACACAAAAAAAGCAACCUAGGCUGCUUUUUUUCAU
.......(((((....((((((.((.(((((........))))).)).))))))..)))))..........((
(........)))...((((((((((........))))))))))...
>9_9590_1_21477_Pseudoalteromonas_simplex_strain_A520_igr1493.fa_1_l=119_226
UCCUCAUCCUGUAGGACAUCCUGUUGAUGAUGCAUCUGAAAGGAUGCAACGUAUUGAUAGCUUCUCCCCAGUU
UGCUAUAACGACUCACUCCUUGUUGAGUGAGUCGCUUUUUCUUUUC
..........((((((....((((..(((.(((((((....)))))))...)))..))))....)))......
))).....((((((((((......))))))))))............

Supplementary File S5. Evolutionary conservation of predicted Pseudalteromonas spot 42 homologs in related species. The lines contains names of species of the spot 42 candidates, introduced by “Cand.”, lists of species in which statistically significant BLAST hits of the candidate sequences in spot 42-syntenic IGRs were obtained follow the arrows.

Cand. Pseudoalteromonas piratica OCN -> Pseudoalteromonas Mix_sand22.2, Pseudoalteromonas G4 NODE, Pseudoalteromonas SSM20 contig, Pseudoalteromonas P1-9 AN214, Pseudoalteromonas T1lg24 NODE, Pseudoalteromonas spongiae S36, Pseudoalteromonas MMG024 , Pseudoalteromonas spongiae HQB, Pseudoalteromonas ABV4_021 con, Pseudoalteromonas XMcav1-K NOD, Pseudoalteromonas spongiae CE1, Pseudoalteromonas spongiae SAO, Pseudoalteromonas spongiae UST, Pseudoalteromonas tunicata D2, Pseudoalteromonas tunicata SMS, Pseudoalteromonas tunicata , Pseudoalteromonas tunicata D2, Pseudoalteromonas tunicata D2, Pseudoalteromonas xiamenensis , Pseudoalteromonas ulvae TC14, Pseudoalteromonas ulvae UL12, Pseudoalteromonas fenneropenae, Pseudoalteromonas peptidolytic, Pseudoalteromonas peptidolytic, Pseudoalteromonas J010 scaffol, Pseudoalteromonas peptidolytic, Pseudoalteromonas UG3-1 tig000,

Pseudoalteromonas SCSIO 43201, Pseudoalteromonas SMS1 scaffol, Pseudoalteromonas luteoviolace, Pseudoalteromonas piscicida 36, Pseudoalteromonas T1lg10 NODE, Pseudoalteromonas piscicida US, Pseudoalteromonas SSDWG2 conti, Pseudoalteromonas pernae YIC-4, Pseudoalteromonas qingdaonensi, Pseudoalteromonas galatheae S4, Pseudoalteromonas piscicida JC, Pseudoalteromonas HM-SA03 scaf, Pseudoalteromonas flavipulchra, Pseudoalteromonas piscicida AT, Pseudoalteromonas piscicida S1, Pseudoalteromonas piscicida S2, Pseudoalteromonas piscicida S2, Pseudoalteromonas Of11M-6 scaf, Pseudoalteromonas ABE4_541 Con, Pseudoalteromonas GCY 2, Pseudoalteromonas galatheae pB, Pseudoalteromonas piscicida WJ, Pseudoalteromonas piscicida Y9, Pseudoalteromonas GCY Scaffold, Pseudoalteromonas OANN1 NODE, Pseudoalteromonas NJ631 Scaffo, Pseudoalteromonas maricaloris .

Cand. Pseudoalteromonas piratica OCN -> Pseudoalteromonas G4 NODE, Pseudoalteromonas Mix_sand22.2, Pseudoalteromonas SSM20 contig, Pseudoalteromonas spongiae UST, Pseudoalteromonas P1-9 AN214, Pseudoalteromonas T1lg24 NODE, Pseudoalteromonas spongiae S36, Pseudoalteromonas MMG024 , Pseudoalteromonas spongiae HQB, Pseudoalteromonas ABV4_021 con, Pseudoalteromonas XMcav1-K NOD, Pseudoalteromonas spongiae CE1, Pseudoalteromonas spongiae SAO, Pseudoalteromonas tunicata D2, Pseudoalteromonas tunicata SMS, Pseudoalteromonas tunicata , Pseudoalteromonas tunicata D2, Pseudoalteromonas tunicata D2, Pseudoalteromonas ulvae TC14, Pseudoalteromonas ulvae UL12, Pseudoalteromonas xiamenensis , Pseudoalteromonas fenneropenae, Pseudoalteromonas SMS1 scaffol, Pseudoalteromonas luteoviolace, Pseudoalteromonas peptidolytic, Pseudoalteromonas peptidolytic, Pseudoalteromonas luteoviolace, Pseudoalteromonas J010 scaffol, Pseudoalteromonas peptidolytic, Pseudoalteromonas UG3-1 tig000, Pseudoalteromonas SCSIO 43201, Pseudoalteromonas ABE4_541 Con, Pseudoalteromonas luteoviolace, Pseudoalteromonas luteoviolace, Pseudoalteromonas T1lg10 NODE, Pseudoalteromonas luteoviolace, Pseudoalteromonas luteoviolace, Pseudoalteromonas SSDWG2 conti, Pseudoalteromonas luteoviolace, Pseudoalteromonas luteoviolace, Pseudoalteromonas luteoviolace, Pseudoalteromonas luteoviolace, Pseudoalteromonas luteoviolace, Pseudoalteromonas luteoviolace, Pseudoalteromonas luteoviolace, Pseudoalteromonas piscicida 36, Pseudoalteromonas piscicida US, Pseudoalteromonas pernae YIC-4, Pseudoalteromonas qingdaonensi, Pseudoalteromonas galatheae S4, Pseudoalteromonas maricaloris , Pseudoalteromonas XMcav11-Q NO, Pseudoalteromonas JC3 , Pseudoalteromonas flavipulchra.

Cand. Pseudoalteromonas haloplanktis -> Pseudoalteromonas MQS005 Scaff, Pseudoalteromonas G5_S1_54_S4 , Pseudoalteromonas S3260 S3260, Pseudoalteromonas tetraodonis , Pseudoalteromonas tetraodonis , Pseudoalteromonas tetraodonis , Pseudoalteromonas SiA1 SiA1-1, Pseudoalteromonas SM9913 I, Pseudoalteromonas SD03 1, Pseudoalteromonas TB43-MNA-CIB, Pseudoalteromonas issachenkoni, Pseudoalteromonas tetraodonis , Pseudoalteromonas TB41 ctg84, Pseudoalteromonas 23_GOM-1509m, Pseudoalteromonas 6BO_GOM-1096, Pseudoalteromonas S201 S201, Pseudoalteromonas A2 contig16, Pseudoalteromonas APC 3213, Pseudoalteromonas issachenkoni, Pseudoalteromonas tetraodonis , Pseudoalteromonas M58, Pseudoalteromonas G24-MNA-CIBA, Pseudoalteromonas issachenkoni, Pseudoalteromonas tetraodonis , Pseudoalteromonas APC 3495, Pseudoalteromonas undina DSM, Pseudoalteromonas tetraodonis , Pseudoalteromonas S1608 S1608, Pseudoalteromonas S1610 S1610, Pseudoalteromonas ERR2094167_b, Pseudoalteromonas US3C1004 1, Pseudoalteromonas US3C1013 1, Pseudoalteromonas SCSIO_11900 ,

Pseudoalteromonas tetraodonis , Pseudoalteromonas 3J6 Pseudoal, Pseudoalteromonas Scap25 1, Pseudoalteromonas Scap03 1, Pseudoalteromonas Scap06 1, Pseudoalteromonas DL-6 1, Pseudoalteromonas RW-H-Ap-1 NZ, Pseudoalteromonas KJ71-7 1, Pseudoalteromonas Mix_sand18.2, Pseudoalteromonas SK18 4, Pseudoalteromonas SCSIO 43095, Pseudoalteromonas QJM3NY_12, Pseudoalteromonas nigrifaciens, Pseudoalteromonas nigrifaciens, Pseudoalteromonas CAL260-MNA-C.
Cand. Pseudoalteromonas haloplanktis -> Pseudoalteromonas issachenkoni, Pseudoalteromonas tetraodonis , Pseudoalteromonas TB41 ctg84, Pseudoalteromonas MQS005 Scaff, Pseudoalteromonas 23_GOM-1509m, Pseudoalteromonas 6BO_GOM-1096, Pseudoalteromonas G5_S1_54_S4 , Pseudoalteromonas S201 S201, Pseudoalteromonas S3260 S3260, Pseudoalteromonas A2 contig16, Pseudoalteromonas tetraodonis , Pseudoalteromonas tetraodonis , Pseudoalteromonas tetraodonis , Pseudoalteromonas APC 3213, Pseudoalteromonas issachenkoni, Pseudoalteromonas SiA1 SiA1-1, Pseudoalteromonas SM9913 I, Pseudoalteromonas SD03 1, Pseudoalteromonas tetraodonis , Pseudoalteromonas TB43-MNA-CIB, Pseudoalteromonas M58, Pseudoalteromonas G24-MNA-CIBA, Pseudoalteromonas issachenkoni, Pseudoalteromonas tetraodonis , Pseudoalteromonas APC 3495, Pseudoalteromonas undina DSM, Pseudoalteromonas S1608 S1608, Pseudoalteromonas S1610 S1610, Pseudoalteromonas US3C1004 1, Pseudoalteromonas US3C1013 1, Pseudoalteromonas 3J6 Pseudoal, Pseudoalteromonas Scap25 1, Pseudoalteromonas Scap03 1, Pseudoalteromonas Scap06 1, Pseudoalteromonas DL-6 1, Pseudoalteromonas RW-H-Ap-1 NZ, Pseudoalteromonas KJ71-7 1, Pseudoalteromonas Mix_sand18.2, Pseudoalteromonas SK18 4, Pseudoalteromonas tetraodonis , Pseudoalteromonas ERR2094167_b, Pseudoalteromonas SCSIO_11900 , Pseudoalteromonas QJM3NY_12, Pseudoalteromonas tetraodonis , Pseudoalteromonas SCSIO 43095, Pseudoalteromonas nigrifaciens, Pseudoalteromonas nigrifaciens, Pseudoalteromonas CAL260-MNA-C.
Cand. Pseudoalteromonas maricaloris -> Pseudoalteromonas flavipulchra, Pseudoalteromonas CO342X NODE, Pseudoalteromonas piscicida S2, Pseudoalteromonas A22 2, Pseudoalteromonas maricaloris , Pseudoalteromonas maricaloris , Pseudoalteromonas XMcav11-Q NO, Pseudoalteromonas JC3 , Pseudoalteromonas NC201, Pseudoalteromonas L23 L23, Pseudoalteromonas L7 L7, Pseudoalteromonas piscicida M1, Pseudoalteromonas M8 2, Pseudoalteromonas JC3 1, Pseudoalteromonas flavipulchra, Pseudoalteromonas Of11M-6 scaf, Pseudoalteromonas GCY 2, Pseudoalteromonas piscicida DE, Pseudoalteromonas GCY Scaffold, Pseudoalteromonas piscicida DE, Pseudoalteromonas piscicida DE, Pseudoalteromonas HM-SA03 scaf, Pseudoalteromonas piscicida S1, Pseudoalteromonas piscicida S2, Pseudoalteromonas piscicida S2, Pseudoalteromonas piscicida Y9, Pseudoalteromonas OANN1 NODE, Pseudoalteromonas NJ631 Scaffo, Pseudoalteromonas galatheae S4, Pseudoalteromonas galatheae pB, Pseudoalteromonas piscicida JC, Pseudoalteromonas piscicida AT, Pseudoalteromonas CO348 NODE, Pseudoalteromonas OF7H-1 NODE, Pseudoalteromonas piscicida WC, Pseudoalteromonas piscicida WJ, Pseudoalteromonas maricaloris , Pseudoalteromonas maricaloris , Pseudoalteromonas piscicida 36, Pseudoalteromonas piscicida US, Pseudoalteromonas UG3-1 tig000, Pseudoalteromonas peptidolytic, Pseudoalteromonas peptidolytic, Pseudoalteromonas J010 scaffol, Pseudoalteromonas peptidolytic, Pseudoalteromonas SCSIO 43201, Pseudoalteromonas luteoviolace, Pseudoalteromonas luteoviolace, Pseudoalteromonas luteoviolace, Pseudoalteromonas ABE4_541 Con, Pseudoalteromonas SMS1 scaffol, Pseudoalteromonas luteoviolace.

Cand. Pseudoalteromonas maricaloris  -> Pseudoalteromonas flavipulchra, Pseudoalteromonas CO342X NODE, Pseudoalteromonas piscicida S2, Pseudoalteromonas A22 2, Pseudoalteromonas maricaloris , Pseudoalteromonas maricaloris , Pseudoalteromonas XMcav11-Q NO, Pseudoalteromonas JC3 , Pseudoalteromonas NC201, Pseudoalteromonas L23 L23, Pseudoalteromonas L7 L7, Pseudoalteromonas piscicida M1, Pseudoalteromonas M8 2, Pseudoalteromonas JC3 1, Pseudoalteromonas flavipulchra, Pseudoalteromonas Of11M-6 scaf, Pseudoalteromonas GCY 2, Pseudoalteromonas GCY Scaffold, Pseudoalteromonas HM-SA03 scaf, Pseudoalteromonas piscicida DE, Pseudoalteromonas piscicida DE, Pseudoalteromonas piscicida DE, Pseudoalteromonas galatheae S4, Pseudoalteromonas CO348 NODE, Pseudoalteromonas OF7H-1 NODE, Pseudoalteromonas galatheae pB, Pseudoalteromonas piscicida WC, Pseudoalteromonas maricaloris , Pseudoalteromonas maricaloris , Pseudoalteromonas piscicida S1, Pseudoalteromonas piscicida S2, Pseudoalteromonas piscicida S2, Pseudoalteromonas piscicida Y9, Pseudoalteromonas OANN1 NODE, Pseudoalteromonas NJ631 Scaffo, Pseudoalteromonas piscicida JC, Pseudoalteromonas piscicida AT, Pseudoalteromonas piscicida WJ, Pseudoalteromonas piscicida 36, Pseudoalteromonas piscicida US, Pseudoalteromonas UG3-1 tig000, Pseudoalteromonas peptidolytic, Pseudoalteromonas peptidolytic, Pseudoalteromonas J010 scaffol, Pseudoalteromonas peptidolytic, Pseudoalteromonas SCSIO 43201, Pseudoalteromonas luteoviolace, Pseudoalteromonas luteoviolace, Pseudoalteromonas luteoviolace, Pseudoalteromonas ABE4_541 Con, Pseudoalteromonas SMS1 scaffol, Pseudoalteromonas luteoviolace.
Cand. Pseudoalteromonas atlantica NB -> Pseudoalteromonas agarivorans , Pseudoalteromonas Xi13 1, Pseudoalteromonas UBA6610 UBA6, Pseudoalteromonas Angola-9 Ang, Pseudoalteromonas Angola-30 An, Pseudoalteromonas CST3 CST3, Pseudoalteromonas Angola-18 An, Pseudoalteromonas 2CM37A NODE, Pseudoalteromonas ERR599005_bi, Pseudoalteromonas S979 S979, Pseudoalteromonas S981 S981, Pseudoalteromonas S1731 S1731, Pseudoalteromonas S983 S983, Pseudoalteromonas S2893 S2893, Pseudoalteromonas S326 S326, Pseudoalteromonas S327 S327, Pseudoalteromonas atlantica CI, Pseudoalteromonas agarivorans , Pseudoalteromonas agarivorans , Pseudoalteromonas ERR599005_bi, Pseudoalteromonas agarivorans , Pseudoalteromonas atlantica EC, Pseudoalteromonas agarivorans , Pseudoalteromonas atlantica Ma, Pseudoalteromonas atlantica Ma, Pseudoalteromonas UBA2102 UBA2, Pseudoalteromonas CuT4-3, Pseudoalteromonas espejiana DS, Pseudoalteromonas P1-13-1a AN3, Pseudoalteromonas C12FD-1 Cont, Pseudoalteromonas MM1 1, Pseudoalteromonas carrageenovo, Pseudoalteromonas carrageenovo, Pseudoalteromonas agarivorans , Pseudoalteromonas carrageenovo, Pseudoalteromonas RW-H-Ap-1 1, Pseudoalteromonas MMG006 , Pseudoalteromonas MMG007 , Pseudoalteromonas carrageenovo, Pseudoalteromonas carrageenovo, Pseudoalteromonas carrageenovo.
Cand. Pseudoalteromonas atlantica NB -> Pseudoalteromonas agarivorans , Pseudoalteromonas UBA6610 UBA6, Pseudoalteromonas Angola-9 Ang, Pseudoalteromonas Angola-30 An, Pseudoalteromonas CST3 CST3, Pseudoalteromonas Angola-18 An, Pseudoalteromonas ERR599005_bi, Pseudoalteromonas S983 S983, Pseudoalteromonas S326 S326, Pseudoalteromonas S327 S327, Pseudoalteromonas atlantica CI, Pseudoalteromonas agarivorans , Pseudoalteromonas ERR599005_bi, Pseudoalteromonas agarivorans , Pseudoalteromonas atlantica EC, Pseudoalteromonas agarivorans , Pseudoalteromonas UBA2102 UBA2, Pseudoalteromonas Xi13 1, Pseudoalteromonas 2CM37A NODE, Pseudoalteromonas S979 S979, Pseudoalteromonas S981 S981,

Pseudoalteromonas S1731 S1731, Pseudoalteromonas S2893 S2893, Pseudoalteromonas agarivorans , Pseudoalteromonas C12FD-1 Cont, Pseudoalteromonas atlantica Ma, Pseudoalteromonas atlantica Ma, Pseudoalteromonas MM1 1, Pseudoalteromonas CuT4-3, Pseudoalteromonas P1-13-1a AN3, Pseudoalteromonas espejiana DS, Pseudoalteromonas carrageenovo, Pseudoalteromonas carrageenovo, Pseudoalteromonas agarivorans , Pseudoalteromonas carrageenovo, Pseudoalteromonas MMG006 , Pseudoalteromonas carrageenovo, Pseudoalteromonas carrageenovo, Pseudoalteromonas carrageenovo, Pseudoalteromonas MMG007 , Pseudoalteromonas distincta CI, Pseudoalteromonas distincta Ma.

Cand. Pseudoalteromonas marina REMEI -> Pseudoalteromonas marina -21, Pseudoalteromonas SRR3933286_b, Pseudoalteromonas S554 S554, Pseudoalteromonas SIMBA_162 NO, Pseudoalteromonas PS1M3 1, Pseudoalteromonas marina DSM, Pseudoalteromonas ECSMB14103 c, Pseudoalteromonas marina 4G09, Pseudoalteromonas ESRF-bin5 co, Pseudoalteromonas marina AF, Pseudoalteromonas marina 13-15, Pseudoalteromonas 13-15 1, Pseudoalteromonas CR1 NODE, Pseudoalteromonas APM04 APM04, Pseudoalteromonas 10-33 1033, Pseudoalteromonas S186 S186, Pseudoalteromonas S407 S407, Pseudoalteromonas S408 S408, Pseudoalteromonas S410 S410, Pseudoalteromonas PSi26 9, Pseudoalteromonas aliena EH1, Pseudoalteromonas SG44-4 NODE, Pseudoalteromonas aliena 4-110, Pseudoalteromonas FUC4 scaffol, Pseudoalteromonas Z1A8 Scaffol, Pseudoalteromonas SG45-1 NODE, Pseudoalteromonas SWN29 WN-29, Pseudoalteromonas SWN166 WN-16, Pseudoalteromonas distincta CI, Pseudoalteromonas SG43-1 NODE, Pseudoalteromonas SR44-2 NODE, Pseudoalteromonas SG43-4 NODE, Pseudoalteromonas SR41-7 NODE, Pseudoalteromonas SG43-3 NODE, Pseudoalteromonas SG45-6 NODE, Pseudoalteromonas SG45-2 NODE, Pseudoalteromonas arctica 2-12, Pseudoalteromonas B62 contig, Pseudoalteromonas B131b contig, Pseudoalteromonas distincta KM, Pseudoalteromonas distincta Ma, Pseudoalteromonas ERR2762177_b, Pseudoalteromonas distincta MK, Pseudoalteromonas arctica AS26, Pseudoalteromonas AS71 , Pseudoalteromonas LC2018020214, Pseudoalteromonas RB2-MNA-CIBA, Pseudoalteromonas CAL107-MNA-C, Pseudoalteromonas distincta 8N, Pseudoalteromonas HL-AS1.

Cand. Pseudoalteromonas marina ECSMB -> Pseudoalteromonas ECSMB14103 c, Pseudoalteromonas marina 13-15, Pseudoalteromonas marina REMEI, Pseudoalteromonas S554 S554, Pseudoalteromonas 13-15 1, Pseudoalteromonas PS1M3 1, Pseudoalteromonas marina DSM, Pseudoalteromonas APM04 APM04, Pseudoalteromonas marina AF, Pseudoalteromonas marina -21, Pseudoalteromonas SRR3933286_b, Pseudoalteromonas SIMBA_162 NO, Pseudoalteromonas marina 4G09, Pseudoalteromonas ESRF-bin5 co, Pseudoalteromonas S186 S186, Pseudoalteromonas S407 S407, Pseudoalteromonas S408 S408, Pseudoalteromonas S410 S410, Pseudoalteromonas CR1 NODE, Pseudoalteromonas PSi26 9, Pseudoalteromonas 10-33 1033, Pseudoalteromonas P1-13-1a AN3, Pseudoalteromonas C12FD-1 Cont, Pseudoalteromonas MM1 1, Pseudoalteromonas agarivorans , Pseudoalteromonas UBA6610 UBA6, Pseudoalteromonas Angola-9 Ang, Pseudoalteromonas Angola-30 An, Pseudoalteromonas CST3 CST3, Pseudoalteromonas Angola-18 An, Pseudoalteromonas ERR599005_bi, Pseudoalteromonas S326 S326, Pseudoalteromonas S327 S327, Pseudoalteromonas agarivorans , Pseudoalteromonas ERR599005_bi, Pseudoalteromonas agarivorans , Pseudoalteromonas atlantica EC, Pseudoalteromonas agarivorans , Pseudoalteromonas UBA2102 UBA2, Pseudoalteromonas atlantica NB, Pseudoalteromonas 2CM37A NODE, Pseudoalteromonas S979 S979, Pseudoalteromonas S981 S981, Pseudoalteromonas S1731 S1731, Pseudoalteromonas atlantica Ma, Pseudoalteromonas atlantica Ma, Pseudoalteromonas CuT4-3.

Cand. Pseudoalteromonas simplex A520 -> Pseudoalteromonas apostichopi , Pseudoalteromonas P1-26 AN394, Pseudoalteromonas Gpa_10 GpaG, Pseudoalteromonas Isolate3 M13, Pseudoalteromonas shioyasakien, Pseudoalteromonas HF66 NODE, Pseudoalteromonas D15MCD-2 CON, Pseudoalteromonas SCSIO 43101, Pseudoalteromonas gelatinilyti, Pseudoalteromonas CH_XMU1449-3, Pseudoalteromonas CS_790 CS, Pseudoalteromonas CO133X NODE, Pseudoalteromonas CO302Y NODE, Pseudoalteromonas G1.bin.8.str, Pseudoalteromonas T1lg122 NODE, Pseudoalteromonas S4491 S4491, Pseudoalteromonas S4492 S4492, Pseudoalteromonas CJ01B_5.24 k, Pseudoalteromonas 68 DY56-GL68, Pseudoalteromonas CO109Y NODE, Pseudoalteromonas shioyasakien, Pseudoalteromonas CJ01B_5.24 k, Pseudoalteromonas arabiensis J, Pseudoalteromonas BZA4 NODE, Pseudoalteromonas T1lg21 NODE, Pseudoalteromonas shioyasakien, Pseudoalteromonas 2102 NODE, Pseudoalteromonas gelatinilyti, Pseudoalteromonas CnMc7-37 NOD, Pseudoalteromonas shioyasakien, Pseudoalteromonas shioyasakien, Pseudoalteromonas lipolytica M, Pseudoalteromonas lipolytica M, Pseudoalteromonas lipolytica M, Pseudoalteromonas ERR594412_bi, Pseudoalteromonas lipolytica G, Pseudoalteromonas lipolytica M.

Supplementary File S6. Sequences of spf homologs identified by cascade sequence searches using homologs of spf in Pseudalteromonas predicted by the presented approach as queries.

Cand. Flocculibacter collagenilyticu -> Flocculibacter collagenilyticu, Psychrosphaera aestuarii PSC10, Psychrosphaera aestuarii JCM, Psychrosphaera HA2-10, Psychrosphaera haliotis JCM, Psychrosphaera haliotis Mar, Psychrosphaera haliotis UCD-MC, Alishewanella d11, Alishewanella longhuensis #55, Alishewanella longhuensis #56, Alishewanella longhuensis CGMC, Psychrosphaera ytuae MTZ26, Alishewanella agri BL06, Alishewanella WH16-1 contig8, Alishewanella tabrizica KCTC, Alishewanella aestuarii B11, Alishewanella jeotgali KCTC, Alishewanella HH-ZS contig, Alishewanella BS5-314 contig, Psychrosphaera aquimarina KCTC, Psychrosphaera F3M07 F3M07, Uncultured Psychrosphaera 2-98, Psychrosphaera saromensis SA4-, Psychrosphaera saromensis NBRC, Psychrosphaera saromensis KCTC, Psychrosphaera 1_MG-2023 B3R15, Psychrosphaera I2R16 I2R16, Psychrosphaera B3R10 B3R10, Psychrosphaera ERR7163074_maxb.

Supplementary File S7. Sequences and secondary structures in dot-bracket format of predicted ms1 RNA homologs in Frankia.

```
>1_55949_1_75499_Candidatus_Frankia_nodulisporulans_strain_igr1065.fa_1_l
=227_621
AGCGGGGCGGGGCCGACGCGCGCGGUGGACGGCUGCGUCGCGGGGUGCUGCCAGUGCCGGUGCCCGAGGUCGA
UCUGUCCUGCCCGGAGCGUGCGGACGCAUCGGUGCCGGACGGUCGGGGAGGGCGGUCGGCAUCCGGCUGGGCG
CGGGGGGCUACCGCGGAGGGGCGCCCGGCCCGUCGUCCCCGGCCACUCCGGGGGCGGCCACCCUCCGUUACUU
CUGGUGUC
(((((((.((((((..((((((((((.....)))))).))))(((((((.((..(((.(((((((...(((((
.(((((((.(((.((.(((.(((.(((....)))))).))).)).)))))))))))))))..(((((...)).
))).)))).)))))..))))))))))))))).(((((((((((.....)))))))))))..)))))))))....
........
>9_969536_1_973066_Frankia_sp._ArI3_chromosome,_rc_igr1318.fa_1_l=227_124
6
CGAAGCAGUCGCCGCAGGUCCCGGGCCUACCGCGAUCCCCCCGCACGCCUCCGGGCCGCUGCCCGUCCCUGAU
CCCCCCGCCGCCGCGACGCCCCCGCGCAGCCGGCCGCCCAGGGCAUCGGCUCCACCCCGGGCGGCGGCCACCC
GGCCGGUCCCGGCCGGGGGACCCCGGGGGCCCCGGUGCGGCUGCCACGGAGCGGGGCGACCGGCCCGUCACCG
CCGUUCCG
```

....((((((((....((((.(((.....))).))))...(((...(((((((((((((((((((...(((((
(((...((.(((((..((......))..).)))).))...))).)))))........))))))))))((.(((
(((((....))))))))))..)))))))))..))).))))))))..((((((((((.(((......))).)).
))))))))
>12_2514651_1_2523346_Frankia_sp._QA3_chromosome,_igr2721.fa_1_l=227_701
UCCGGCCCGCUCCGCCGCCGAUGCCGCCGGCGCCCGGAACGCGCGUAGGGAGGUGACUCGGCGUCCUCGGUGG
UCGAGGUCUCGUCGCGUCCGUACUGGUCGUCUCGGUCCGGCAGCCGGUUGGAACGGGAGGCCGUCGUGGGAU
GCGGGCUCCUAUGAGCGAAACGGGCCGCGCGUCGCCUUCCUCACCGUACGGUGCCUGUGGCACAUCGCUCACG
CGCCGUCG
....((.((.((((.(((((.......)))))..)))).)).))..(((((((((((.((.((.((.....))
.)).((((.((((((((((..(((((((.(((((.(((..(.....)..))).))))))))))..))..))))
))).((((....))))...)))))).)).))))))))))))...((.(((((((..(((((.....)).))))
))))))))
>16_13934_1_11236_Frankia_sp._BMG5.23_BMG523Draft_scaffold_0.1,_rc_igr183
4.fa_1_l=227_816
ACGAAGCGUGGCAGAGCGAGGCAUUCCACCUCGAUGAUGCGACAUCGGCGGCCCGUUCGGGUGCGGACCGUAC
AAACACGUCCGACGACACACCGCCAUCGGUGCCGGGCGCCAGGACCGAGGCACCCGGCGCCGGGCCGGGCCGG
GCGACCACGGGCGCGGACACCAGCUCCCGGCGCACCCGCGGCGCCGCUGCCCGGCGCGCCGGCCGCCCGUCGG
CGAUUCCG
.......((.(((...(((((.......)))))....))).))..((((((((((..((((((((........
.....((((((........(((((..((((((((((.(((........))).))))))))))))).))..)))
)))....((((.((........)).)))).))))))))))).))))))))..(((..(((((((.....)))))
))...)))
>16_22790_1_18093_Frankia_casuarinae,_complete_sequence_igr21.fa_1_l=227_
576
ACGAAGCGUGGCAGAGCGAGGCAUUCCACCUCGAUGAUGCGACAUCGGCGGCCCGUUCGGGUGCGGACCGUAC
AAACACGUCCGACGACACACCGCCAUCGGUGCCGGGCGCCAGGACCGAGGCACCCGGCGCCGGGCCGGGCCGG
GCGACCACGGGCGCGGACACCAGCUCCCGGCGCACCCGCGGCGCCGCUGCCCGGCGCGCCGGCCGCCCGUCGG
CGAUUCCG
.......((.(((...(((((.......)))))....))).))..((((((((((..((((((((........
.....((((((........(((((..((((((((((.(((........))).))))))))))))).))..)))
)))....((((.((........)).)))).))))))))))).))))))))..(((..(((((((.....)))))
))...)))
>16_56182_1_75199_Frankia_sp._CcI6_CcI6DRAFT_scaffold_0.1,_igr359.fa_1_l=
227_576
ACGAAGCGUGGCAGAGCGAGGCAUUCCACCUCGAUGAUGCGACAUCGGCGGCCCGUUCGGGUGCGGACCGUAC
AAACACGUCCGACGACACACCGCCAUCGGUGCCGGGCGCCAGGACCGAGGCACCCGGCGCCGGGCCGGGCCGG
GCGACCACGGGCGCGGACACCAGCUCCCGGCGCACCCGCGGCGCCGCUGCCCGGCGCGCCGGCCGCCCGUCGG
CGAUUCCG
.......((.(((...(((((.......)))))....))).))..((((((((((..((((((((........
.....((((((........(((((..((((((((((.(((........))).))))))))))))).))..)))
)))....((((.((........)).)))).))))))))))).))))))))..(((..(((((((.....)))))
))...)))
>16_3504573_1_3523545_Frankia_sp._Allo2_ALLO2DRAFT_scaffold_0.1,_rc_igr54
.fa_1_l=227_576
ACGAAGCGUGGCAGAGCGAGGCAUUCCACCUCGAUGAUGCGACAUCGGCGGCCCGUUCGGGUGCGGACCGUAC
AAACACGUCCGACGACACACCGCCAUCGGUGCCGGGCGCCAGGACCGAGGCACCCGGCGCCGGGCCGGGCCGG
GCGACCACGGGCGCGGACACCAGCUCCCGGCGCACCCGCGGCGCCGCUGCCCGGCGCGCCGGCCGCCCGUCGG
CGAUUCCG
.......((.(((...(((((.......)))))....))).))..((((((((((..((((((((........
.....((((((........(((((..((((((((((.(((........))).))))))))))))).))..)))
)))....((((.((........)).)))).))))))))))).))))))))..(((..(((((((.....)))))
))...)))
>20_59319_1_85678_Frankia_sp._AvcI1_Ga0078773_101,_rc_igr1048.fa_1_l=227_
146
CGACCCAUCCGUUGUGGGUAGCCUCAACCAGAGCUUGUCAUACUCGAAUGGCUCGUGUUACAUCGAUCCGACC
GCAGCCGAAGGCAAGUAACCGUAAGUAACCUUAACUGGCACUAGCUAACCUGUAGAGAGCCGGGAGGGCGACG

```
GCGCGUGAGCGAUGUGCCACAGGCACCGUACGGUGAGGAAGGCGACGCGCGGCCCGUUUCGCUCAUAGGAGCC
CGCAUCCC
..((((((.....))))))..((((.(((...((.((((..........((((((.(((..........))))
).))))...((((.....((((......((((..(((((.((.((......))..)).)))))))))...)))
)(((....)))...))))...))))..))..)))))))..((.((.(((.(((((............)).)))
))).))))
>24_20438_1_3948_Frankia_canadensis_isolate_FRACA_ARgP5,_igr791.fa_1_l=22
7_716
ACCGGACCUGGAGGCGCGGCCCGGAUCACCCCGACGAGCGGGUGAUCGGCCACGCGUGACCAUGCCGGGGCCG
GCGGGGGGUAGUCGGGAUCAGCGGCGCGCAUCGUGCGGCAGACGAAAGGAGCGUCGGUGGCGCAGGACGGCUU
CUCGAUGGCGACGCCCCGAUCAUCCGGUCGCCCCGGUCGCCCGGCGGGCCCACAGGGAUCCACGCAGGGUCGU
GCCCUGCC
...(((((((...(((.((((..((((((((........))))))))))))).)))...........(((.(((
.((((.(...((((((....(((.(((.(((((((((.((((((.......))))..)).)))).........
..)))))))).))))))))...((((.....)))).).))))).))).))).))))..)))..((((((....
.)))))).
>26_17014_1_11590_Frankia_sp._ACN1ag_Ga0074159_101,_rc_igr120.fa_1_l=227_
231
GACCUGCGGCGACUGCUUCGGCGGGGGCGGCUGCGGGGCCGUGAUCGGCCCCGGGACCGGAGCCCGCGGUCUG
UGCGGUCGUGCCUGUGCCGGUGCCUACUCCGGUGCCGGCGGCAUGGUCGGAGCCGUCCGCUCGGGACGGGACC
AUCGCGUCCCGCGACGAUCCCGGGGCCGAGGCGUCGGCGCCGCCGGGCGGCGCCGGGCGGCGGGAAGGGUCUG
CCUGGUUG
..((.((.((..((((....))))..)).)).((((.((((((.((((((((((((.((..((..((((...(
(.((..((((((...(((((..((......))..)))))))))))..))..))...))))..((((((.(...
..).))))))))..)).))))))))))..)).))))).))))).)).....((((((((((.......).)))
))))))..
>26_1951092_1_1969440_Frankia_nepalensis_strain_CN7_rc_igr4320.fa_1_l=227
_461
CCGCGGUGCCUGCCCGGGGGACACCCCAUACCCCAGACGCGGUGACGGACGGUACACCACCCUUGGUAGCCAU
GGUUAGGCUCUAUCGGAGCUUGUCAUACUUCAGCGUCGCGGGUUACCCCGAUCUGGUGGUGGCCCGGCGCAAG
UAACGGUAAGUAAUCCCAAGUGGGACCAACAAGCCUGAACCAGCCAGGAGGGCGACGGCACGUGAGCAAUGUG
CCACCGAC
.(((.(((((..((.((((....)))).((((........))))..))..)))))....((((((((.....(
((((((((.(((((((.....)).(((((..((((..((((..(((((.....)).)))..)))))))))))
)).))))).....((((....))))......))))))).))))))))).)))))..((((((.....)))))
))......
>30_13846_1_22850_Frankia_sp._CeD_CEDDRAFT_scaffold_0.1,_rc_igr1634.fa_1_
l=227_556
CGACGAAGCGUGGCAGAGCGAGGCAUUCCACCUCGAUGAUGCGACAUCGGCGGCCCGUUCGGGUGCGGACCGU
ACAAACACGUCCGACGACACACCGCCAUCGGUGCCGGGCGCCAGGACCGAGGCACCCGGCGCCGGGCCGGGCC
GGGCGACCACGGGCGCGGACACCAGCUCCCGGCGCACCCGCGGCGCCGCUGCCCGGCGCGCCGGCCGCCCGUC
GGCGAUUC
((.((....((.(((...(((((.......)))))....))).))..((((((((((..((((((((......
.......((((((........(((((..((((((((((.(((.........))).))))))))))).)))..)
)))))....((((.((........)).)))).))))))))))).)))))))..)).))(((((((.....)))
))))....
>30_106682_1_3339_Frankia_sp._B2_Contig_100_length_3772_cov_81.341289,_ig
r2001.fa_1_l=227_556
CGACGAAGCGUGGCAGAGCGAGGCAUUCCACCUCGAUGAUGCGACAUCGGCGGCCCGUUCGGGUGCGGACCGU
ACAAACACGUCCGACGACACACCGCCAUCGGUGCCGGGCGCCAGGACCGAGGCACCCGGCGCCGGGCCGGGCC
GGGCGACCACGGGCGCGGACACCAGCUCCCGGCGCACCCGCGGCGCCGCUGCCCGGCGCGCCGGCCGCCCGUC
GGCGAUUC
((.((....((.(((...(((((.......)))))....))).))..((((((((((..((((((((......
.......((((((........(((((..((((((((((.(((.........))).))))))))))).)))..)
)))))....((((.((........)).)))).))))))))))).)))))))..)).))(((((((.....)))
))))....
>34_27130_1_5917_Frankia_umida_strain_Ag45_Mut15_igr2717.fa_1_l=227_1361
```

GAGCCGGCGAGCACCGUGGCGCCCGAGGCCGCUGGCCCGAUGGGUGCCGCGAGCUCCGUCGGCCCGCACACG
GCGUGAUUCCCGAGACCACGAACCACGGUGAGACGAGCGCGCCGACGCCGAAGCAUCGGCCCGGCUCGCGUCA
UCUGCCGGUCUCGGAGCCGUCCGGACCGCAAGCCACCGGUCCGGUGCCGGAGCCGCAGGACGAAGUCCCGGUA
CGGCUGUC
...(((((((((..(((((((((((.((((...))))...))))))))))).))).))))))).........((
((.......((((((((.........(((.(((.(((((.((((..(((((....))))).)))).))).)).
))))))))))))))......((((((((........)))))))).))))).(((((..((((...)))).....
)))))...

Supplementary File S8. Evolutionary conservation of predicted Frankia ms1 RNA homologs in related species. The lines contains names of species of the spot 42 candidates, introduced by “Cand.”, lists of species in which statistically significant BLAST hits of the candidate sequences in spot 42-syntenic IGRs were obtained follow the arrows.

Cand. Frankia nodulisporulans AgUmAS -> Frankia umida Ag45/Mut15, Frankia AgPm24 NODE.
Cand. Frankia ArI3 -> Frankia ACN1ag Ga0074159.
Cand. Frankia QA3 -> Frankia AvcI1 NODE, Frankia AvcI1 Ga0078773, Frankia alni ACN14a, Frankia ACN10a NODE, Frankia torreyi CpI1, Frankia CpI1-P FF86, Frankia ACN1ag Ga0074159, Frankia ArI3, Frankia canadensis FRACA, Frankia canadensis FRACA1, Frankia CgMI4 contig, Frankia casuarinae Cg70.9, Frankia CeD CEDDRAFT, Frankia BMG5.23 BMG523Draft, Frankia CeCAU2 Frankia, Frankia B2 Contig, Frankia AgKG'84/4 ena-SAMPLE-T, Frankia CoBUL1 Frankia, Frankia CoWAU3 Frankia, Frankia AiPs1 ena-SAMPLE-TAB-1, Frankia AiPa1 ena-SAMPLE-TAB-0, Frankia AiPa1 NODE, Frankia CcCS1 Frankia, Frankia CeTa3Moorea Frankia, Frankia CeRuru1 Frankia, Frankia casuarinae, Frankia CgS1 contig, Frankia casuarinae Thr, Frankia CcI6 CcI6DRAFT, Frankia Allo2 ALLO2DRAFT, Frankia umida Ag45/Mut15, Frankia nodulisporulans AgUmAS, Frankia AgPm24 NODE, Parafrankia discariae BCU11050, Parafrankia soli NRRL, Parafrankia BMG5.11 contig, Parafrankia soli FMc6, Frankia Mgl5 NODE, Parafrankia irregularis DSM, Frankia R43 contig006, Parafrankia CH37 Contig1, Frankia CcI49 contig, Parafrankia EAN1pec, Pseudofrankia BMG5.37 Scaffold, Pseudofrankia asymbiotica NRRL, Pseudofrankia saprophytica CN3, Pseudofrankia EUN1h contig, Frankia nepalensis CN6, Frankia nepalensis CN7, Pseudofrankia inefficax.
Cand. Frankia BMG5.23 BMG523Draft_sc -> Frankia casuarinae, Frankia CgMI4 contig, Frankia CgS1 contig, Frankia casuarinae Cg70.9, Frankia casuarinae Thr, Frankia CeD CEDDRAFT, Frankia Allo2 ALLO2DRAFT, Frankia CcI6 CcI6DRAFT, Frankia B2 Contig, Frankia CeCAU2 Frankia, Frankia CeTa3Moorea Frankia, Frankia CeRuru1 Frankia, Frankia CoBUL1 Frankia, Frankia CoWAU3 Frankia, Frankia CcCS1 Frankia, Frankia QA3, Frankia alni ACN14a, Frankia AvcI1 Ga0078773, Frankia ACN10a NODE, Frankia AvcI1 NODE, Frankia torreyi CpI1, Frankia CpI1-P FF86, Frankia ACN1ag Ga0074159, Frankia ArI3, Frankia AiPs1 ena-SAMPLE-TAB-1, Frankia AiPa1 ena-SAMPLE-TAB-0, Frankia AiPa1 NODE, Frankia canadensis FRACA, Frankia canadensis FRACA1, Frankia AgKG'84/4 ena-SAMPLE-T, Frankia umida Ag45/Mut15, Frankia nodulisporulans AgUmAS, Frankia AgPm24 NODE.
Cand. Frankia casuarinae -> Frankia BMG5.23 BMG523Draft, Frankia CgMI4 contig, Frankia CgS1 contig, Frankia casuarinae Cg70.9, Frankia casuarinae Thr, Frankia CeD CEDDRAFT, Frankia Allo2 ALLO2DRAFT, Frankia CcI6 CcI6DRAFT, Frankia B2 Contig, Frankia CeCAU2 Frankia, Frankia CeTa3Moorea Frankia, Frankia CeRuru1 Frankia, Frankia CoBUL1 Frankia, Frankia CoWAU3 Frankia, Frankia CcCS1 Frankia, Frankia QA3, Frankia alni ACN14a, Frankia AvcI1 Ga0078773, Frankia ACN10a NODE, Frankia AvcI1 NODE, Frankia torreyi CpI1, Frankia CpI1-P FF86, Frankia ACN1ag Ga0074159, Frankia ArI3, Frankia AiPs1 ena-SAMPLE-TAB-1, Frankia AiPa1 ena-SAMPLE-TAB-0, Frankia AiPa1 NODE, Frankia canadensis FRACA, Frankia canadensis

FRACA1, Frankia AgKG'84/4 ena-SAMPLE-T, Frankia umida Ag45/Mut15, Frankia nodulisporulans AgUmAS, Frankia AgPm24 NODE.
Cand. Frankia CcI6 CcI6DRAFT_scaffol -> Frankia casuarinae, Frankia BMG5.23 BMG523Draft, Frankia CgMI4 contig, Frankia CgS1 contig, Frankia casuarinae Cg70.9, Frankia casuarinae Thr, Frankia CeD CEDDRAFT, Frankia Allo2 ALLO2DRAFT, Frankia B2 Contig, Frankia CeCAU2 Frankia, Frankia CeTa3Moorea Frankia, Frankia CeRuru1 Frankia, Frankia CoBUL1 Frankia, Frankia CoWAU3 Frankia, Frankia CcCS1 Frankia, Frankia QA3, Frankia alni ACN14a, Frankia AvcI1 Ga0078773, Frankia ACN10a NODE, Frankia AvcI1 NODE, Frankia torreyi CpI1, Frankia CpI1-P FF86, Frankia ACN1ag Ga0074159, Frankia ArI3, Frankia AiPs1 ena-SAMPLE-TAB-1, Frankia AiPa1 ena-SAMPLE-TAB-0, Frankia AiPa1 NODE, Frankia canadensis FRACA, Frankia canadensis FRACA1, Frankia AgKG'84/4 ena-SAMPLE-T, Frankia umida Ag45/Mut15, Frankia nodulisporulans AgUmAS, Frankia AgPm24 NODE.
Cand. Frankia Allo2 ALLO2DRAFT_scaff -> Frankia casuarinae, Frankia BMG5.23 BMG523Draft, Frankia CgMI4 contig, Frankia CgS1 contig, Frankia casuarinae Cg70.9, Frankia casuarinae Thr, Frankia CeD CEDDRAFT, Frankia CcI6 CcI6DRAFT, Frankia B2 Contig, Frankia CeCAU2 Frankia, Frankia CeTa3Moorea Frankia, Frankia CeRuru1 Frankia, Frankia CoBUL1 Frankia, Frankia CoWAU3 Frankia, Frankia CcCS1 Frankia, Frankia QA3, Frankia alni ACN14a, Frankia AvcI1 Ga0078773, Frankia ACN10a NODE, Frankia AvcI1 NODE, Frankia torreyi CpI1, Frankia CpI1-P FF86, Frankia ACN1ag Ga0074159, Frankia ArI3, Frankia AiPs1 ena-SAMPLE-TAB-1, Frankia AiPa1 ena-SAMPLE-TAB-0, Frankia AiPa1 NODE, Frankia canadensis FRACA, Frankia canadensis FRACA1, Frankia AgKG'84/4 ena-SAMPLE-T, Frankia umida Ag45/Mut15, Frankia nodulisporulans AgUmAS, Frankia AgPm24 NODE.
Cand. Frankia AvcI1 Ga0078773_115 -> Frankia alni ACN14a, Frankia ACN10a NODE, Frankia AvcI1 NODE, Frankia torreyi CpI1, Frankia CpI1-P FF86, Frankia ACN1ag Ga0074159, Frankia ArI3, Frankia QA3, Frankia canadensis FRACA, Frankia canadensis FRACA1, Frankia casuarinae, Frankia BMG5.23 BMG523Draft, Frankia CgMI4 contig, Frankia CgS1 contig, Frankia casuarinae Cg70.9, Frankia CeCAU2 Frankia, Frankia casuarinae Thr, Frankia CeD CEDDRAFT, Frankia CcI6 CcI6DRAFT, Frankia B2 Contig, Frankia Allo2 ALLO2DRAFT, Frankia CoBUL1 Frankia, Frankia CoWAU3 Frankia, Frankia AgKG'84/4 ena-SAMPLE-T, Frankia CeTa3Moorea Frankia, Frankia CeRuru1 Frankia, Frankia CcCS1 Frankia, Frankia AiPs1 ena-SAMPLE-TAB-1, Frankia AiPa1 ena-SAMPLE-TAB-0, Frankia AiPa1 NODE, Frankia umida Ag45/Mut15, Frankia nodulisporulans AgUmAS, Frankia AgPm24 NODE, Parafrankia discariae BCU11050, Parafrankia soli NRRL, Parafrankia BMG5.11 contig, Parafrankia soli FMc6, Frankia Mgl5 NODE, Parafrankia EAN1pec, Frankia R43 contig006, Frankia CcI49 contig, Parafrankia irregularis DSM, Parafrankia CH37 Contig1, Pseudofrankia asymbiotica NRRL, Pseudofrankia BMG5.37 Scaffold, Pseudofrankia saprophytica CN3, Pseudofrankia EUN1h contig, Frankia nepalensis CN6, Frankia nepalensis CN7.
Cand. Frankia canadensis FRACA_ARgP5 -> Frankia canadensis FRACA1, Frankia QA3, Frankia torreyi CpI1, Frankia CpI1-P FF86, Frankia ACN1ag Ga0074159, Frankia ArI3, Frankia alni ACN14a, Frankia ACN10a NODE, Frankia AvcI1 NODE, Frankia BMG5.23 BMG523Draft, Frankia CgS1 contig.
Cand. Frankia ACN1ag Ga0074159_101 -> Frankia ArI3, Frankia QA3.
Cand. Frankia nepalensis CN4 NODE_39 -> Frankia nepalensis CN6, Frankia nepalensis CN7, Pseudofrankia inefficax, Pseudofrankia saprophytica CN3, Pseudofrankia EUN1h contig, Pseudofrankia asymbiotica NRRL, Pseudofrankia BMG5.37 Scaffold, Frankia QA3, Frankia AgKG'84/4 ena-SAMPLE-T, Frankia torreyi CpI1, Frankia alni ACN14a, Frankia CpI1-P FF86, Frankia AvcI1 Ga0078773, Frankia ACN1ag Ga0074159, Frankia ACN10a NODE, Frankia AvcI1 NODE, Frankia ArI3, Frankia canadensis FRACA, Frankia canadensis FRACA1, Frankia CoBUL1 Frankia, Frankia CoWAU3 Frankia, Parafrankia BMG5.11 contig, Parafrankia soli FMc6, Frankia Mgl5 NODE, Frankia AiPs1 ena-

SAMPLE-TAB-1, Frankia AiPa1 ena-SAMPLE-TAB-0, Frankia AiPa1 NODE, Frankia Allo2 ALLO2DRAFT, Frankia casuarinae, Frankia BMG5.23 BMG523Draft, Frankia CgMI4 contig, Frankia CgS1 contig, Frankia casuarinae Cg70.9, Frankia CeCAU2 Frankia, Frankia CcCS1 Frankia, Frankia casuarinae Thr, Frankia CeD CEDDRAFT, Frankia CcI6 CcI6DRAFT, Frankia B2 Contig, Parafrankia discariae BCU11050, Parafrankia soli NRRL, Parafrankia EAN1pec, Frankia CeTa3Moorea Frankia, Frankia CeRuru1 Frankia, Frankia umida Ag45/Mut15, Frankia nodulisporulans AgUmAS, Frankia AgPm24 NODE, Parafrankia irregularis DSM, Frankia R43 contig006, Parafrankia CH37 Contig1, Frankia CcI49 contig.
Cand. Frankia CeD CEDDRAFT_scaffold_ -> Frankia casuarinae, Frankia BMG5.23 BMG523Draft, Frankia CgMI4 contig, Frankia CgS1 contig, Frankia casuarinae Cg70.9, Frankia casuarinae Thr, Frankia Allo2 ALLO2DRAFT, Frankia CcI6 CcI6DRAFT, Frankia B2 Contig, Frankia CeCAU2 Frankia, Frankia CeTa3Moorea Frankia, Frankia CeRuru1 Frankia, Frankia CoBUL1 Frankia, Frankia CoWAU3 Frankia, Frankia CcCS1 Frankia, Frankia QA3, Frankia alni ACN14a, Frankia AvcI1 Ga0078773, Frankia ACN10a NODE, Frankia AvcI1 NODE, Frankia torreyi CpI1, Frankia CpI1-P FF86, Frankia ACN1ag Ga0074159, Frankia ArI3, Frankia AiPs1 ena-SAMPLE-TAB-1, Frankia AiPa1 ena-SAMPLE-TAB-0, Frankia AiPa1 NODE, Frankia canadensis FRACA, Frankia canadensis FRACA1, Frankia AgKG'84/4 ena-SAMPLE-T, Frankia umida Ag45/Mut15, Frankia nodulisporulans AgUmAS, Frankia AgPm24 NODE.
Cand. Frankia B2 Contig_36_length_50 -> Frankia casuarinae, Frankia BMG5.23 BMG523Draft, Frankia CgMI4 contig, Frankia CgS1 contig, Frankia casuarinae Cg70.9, Frankia casuarinae Thr, Frankia CeD CEDDRAFT, Frankia Allo2 ALLO2DRAFT, Frankia CcI6 CcI6DRAFT, Frankia CeCAU2 Frankia, Frankia CeTa3Moorea Frankia, Frankia CeRuru1 Frankia, Frankia CoBUL1 Frankia, Frankia CoWAU3 Frankia, Frankia CcCS1 Frankia, Frankia QA3, Frankia alni ACN14a, Frankia AvcI1 Ga0078773, Frankia ACN10a NODE, Frankia AvcI1 NODE, Frankia torreyi CpI1, Frankia CpI1-P FF86, Frankia ACN1ag Ga0074159, Frankia ArI3, Frankia AiPs1 ena-SAMPLE-TAB-1, Frankia AiPa1 ena-SAMPLE-TAB-0, Frankia AiPa1 NODE, Frankia canadensis FRACA, Frankia canadensis FRACA1, Frankia AgKG'84/4 ena-SAMPLE-T, Frankia umida Ag45/Mut15, Frankia nodulisporulans AgUmAS, Frankia AgPm24 NODE.

Supplementary File S9. Sequences and secondary structures in dot-bracket format of predicted ms1 RNA homologs in Bifidobacterium.

```
>20_31487_1_7507_Bifidobacterium_animalis_subsp._lactis_rc_igr126.fa_1_l=
227_231
GACGCUUAGGCGAUUCCGACUUGCGUAAUCCUAAAUUCCGAGCAUAAUUGCAAGUACAAGGACAACCGAAGAG
CAGACCCACAAAAGUAGAUGAUCAGCAAGAUUAUUACGUUGGGCGGGCAAUGGAAAGACUUACCGUUACAACG
AUGAGGUUGUGCAGAGAUUCUGCCUCGUUGGACUUGGGAAAGAGACCCACCUCGUGUGGGUCUCUUUUUUAUU
UCAUGAAA
.((((..((.((....)).)).))))............((((((.((((....((((((......(((....(
(...((((.....((.(((((((.....)))))))))..))))...))..))).....((((.(((....)))
.)))).))))))...)))).))).)))...((..(..(((((((((((((.....)))))))))))))..)..
))...... (-62.3)
>16_14774_1_35725_Bifidobacterium_boum_DSM_20432_rc_igr218.fa_1_l=227_106
CGUGGCGUGCAGUGAUCUGGCUGUGCGUUCUGGUGCGGGUUUGUGCGGGUGUGGUUCGUUUGGGGCGAGGGGG
GCGCACCGCCGCGACCGGCGUGUCUUGAGGUGUGCACACAACCGCACAUAUGAACCGGUUCGAUGUGCACAAU
AAUCAGCAGGUUGGCGGCUUCGUGAUGUGUUGUUGAACACUUGCGCAAAGAAGCGAACAAUCCGCCUGCAUGG
GGCGGAUG
((..(((..((((......))))..)))..)).(((...(((((((..((((....((......((((((..(
(((...)))).((.((..(.(((..(((..((((((((.(((((((....))...)))))...))))))))..
..)))))).)..)))).)))))).......))....))))..)))))))...)))....((((((((.....)
))))))). (-86.5)
```

>26_13939_1_25127_Bifidobacterium_pseudocatenulatum_DSM_20438_rc_igr272.f
a_1_l=227_306
AAGAGUGUAUGGCCAAGAUAGGUCGUUACGUUGAACGGGCAUUGGAUAGACUUACCGUUACGACGAUGUUGUU
GUGCAGAGAUUCUGCCCUUAAGGUUCUUGCAAGGAAGGCCCGCUCGGCGGGCCUUCCUUUUUGUUUGCCUGUU
CUCAGGGAGUUUUUCGGGAAAACGACGGACGCCGAUUAUCGUCGUUGUGUGGAUAACGGUGCAUAACACGCCG
GGGUUGUG
..((((.(((..((((......(((((......)))))...))))))).)))).....(((((((((..(((
(.((..........((((..((((....((((((((((((((...)))))))))))))...)))..)))))...
...)))).(((..(((......)))..)))))))))...))))))))))..(((..(((((.......)))))
..)))... (-76.6)
>31_103073_1_2145_Bifidobacterium_tsurumiense_DSM_17777_igr207.fa_1_l=227
_106
CUGCGGUGAAUGGGCUGCGGUGAAUGGGCUGCGGUGAUUGGGCUGCAGUGAAUGAGCUGCGGUGAAUGGAUAU
GGUUGGCGGGAAUUAGGUGUUUGAUGUUUGGAUAGCGCUGUUGUGAAUCGGCGAGGUCUUAAGAUAUGCAUUU
GGCUUUGUGAGAGGACUGGAAGUCCUUUGAGAAUACCCGUGGACGCAGGGCUCAGCACGGUACGUAUGCGAUU
GGCUAUGC
.(((.........((((((((..((..((((((((......))))))))..))..))))))))....((...(
(((..(((((...((..(((...((.((..(((..(((((........)))))..)))..)).))..)))..)
)........(((((((.....)))))))........)))))..)).))...))..))).....((((.((....
.))))))). (-70)
>3_1522873_1_1538129_Bifidobacterium_pseudolongum_AGR2145_G627DRAFT_scaff
old00001.1_C,_igr349.fa_1_l=227_406
AAUUAGUGAACAUUAGCUGAAUCAAGUUGCGGAAUGGCAAAUAGCGAGCAUAAUUACAAGUACAAGAACAACU
GAAUAGCAGACCCACAAAAAGUAUAUGAUCAGCAAGAUUAUUACGUUGAGCGGGUAUUGGAAAGACUUACCGU
UGCAACGAUGAGGUUGUGCAGAGAUUCUGCGCGGUUGGACUUGUAGGGAAGGCCCACAAUAGCGUGGGUCUUC
CCUUUUUU
.....((((...((((((........((((......))))......))).)))))))...((((((..((((.
........((((((..........((((((.....))))))..((((((((((.....((.....))..))))
).))))).)).))))((((((.....)))))).))))..)))))))((((((((((((......))))))))))
))...... (-68.4)
>3_8570_1_36392_Bifidobacterium_angulatum_DSM_20098_igr111.fa_1_l=227_401
CGGUAAGUCUAUCCAAUGCCCGUCCAACGUACCAGUCCUUAUGACCAUUACUUCUUUGUGGGCUCUGCUCUGU
UCUGUUGUCCUUGUACCUCACAAUUUAGGCGCGUUCGAGAGCCUGCGCAACCUGGUCGGAAACCAUGUUCGCU
UUCGUGCUCAAAUGGUGCAAAUCGACAGGAAAACACAGUGAUUUGAAAUGGAAUGUGCACUUCGCUCCAUCCA
UAUGUUUC
...((((((.........(((((......(((...(((.....)))...)))........))))).(((.....(
(((((((...(((((((.....((((..((((....((((((..(((.....((((.....)))))))..)))
))))))..)))))))))))..)))))))).....))).)))))))..(((((.((.((.....))..))))))
)....... (-55.4)
>39_7710_1_16811_Bifidobacterium_adolescentis_ATCC_15703,_rc_igr82.fa_1_l
=227_36
UUGCAAAACUCAGGAAAAUUAAGCAGGAGUUAACAUUCGUUCGCCGAGUUGGUGAAAUAUGCUUCCGACACGC
CGUUUUUUCGGGGCAAAUGAACGUGAAUGAACAAGCAAGCGUUGCGAUUCGCACAGCAUGGAGCAUUUUUGCG
AUUCUUGCGAUCGGAAAGCAACAGUUGCGAUAAAUACGGAACAUCGUCCCGUUUCGGUUUGCACUUGCCGCAA
UGGCGAGC
.(((((......(((......((((.((((....)))).((((((.....))))))...)))))))).....((
((......((((((........((...(((....(((((..((((((...((.(.....)..))....)))))
)..))))))..))).....))....(((.((.......)).)))...))))))...)))))))))((((((....
.))))))). (-58.9)
>40_84709_1_103671_Bifidobacterium_ruminantium_DSM_6489_igr549.fa_1_l=227
_221
ACAUUGCCCGGUUCCGGUUUGCAUCCAUGGAAAUGGCGAGCAUAAUUACGAAGUACAAGGACAACUGAAGAGC
AGAGCCCCCACAAGGUGUGUAUGACCGAGAAUGGUUGUUACGUUGAACGGGCAUUGGAUAGACUUACCGUUAC
GACGAUGUUGUUGUGCAGCGAUUCUGCUCCGAGGAUUCUUGCAGGAAGGCCCGCUUCGGCGGGUCUUCCUUUU
UGUUGCGC
.....((.......((...(((..((((....))))...))).......)).....((((((...((...((((
(((((((...(((.(((...(..(((......)))..)))).)))...))))..............((((((

((((....)))))))..)))..)))))))..))..)))))).((((((((((((....))))))))))))...
....))..  (-74)
>4_17892_1_32585_Bifidobacterium_dentium_Bd1,_complete_rc_igr126.fa_1_l=2
27_136
AUUGCGACGUGUCAGGUGGACUUCGGCGUCAUUCGCCUGAUGGUUGGCGUUCCGGCGGUUCGACGUCGCCGUA
UGGUGGGUUGCGUACGACACGCCGGAGUACGGCAUGAAGCGCACGCAAUCGCACAAUCCUCAGGUGAGAAUCG
CACAAGAUAAUGCAAAUCUCCGAUUAUGGCGAUCGGAAGGGAACAGUUUGGCAAAUAACGCAACAUAGGCCCG
UUUCGGUU
.((((....((((((((((......)))...)))))))).((..(((((((.((((((....(((((...
..((((((((((.((.((.((((.....)))).))...)).))))))))).)))........)))))))))))
)................(((((((.....)))))))..))))).).)..))......)))))).....(((..
....))).  (-78.8)
>44_1630971_1_1640630_Bifidobacterium_moukalabense_DSM_27321_rc_igr518.fa
_1_l=227_71
UUAGUUUGAUGAAUUUGCGAAGAGUGCAUCGAUGGAUGUUCGGUUCUCGGCGGGGAAGGGGGUUGUCGAAGCA
UGUUGCGUGGAUUCCGGCUCCGCUCGCCUGAUGGUUGGCGUUCCGGCAUCUCGGCAUCGCAUAUGGUGGGGAU
GCGUACGACACGCCGGAGUGAGGGAUGAAUCGCACGUAAUCGCACAAACCUCAGGGAAGAAACGCACAGGAUG
AUGCAAAU
...((((......(..((..((((.(((((....)))))....))))..))..)...((((.(((((((....
..((((((((((((...((((((((.((....))..(((((..(((((((((..(((((....))))))))))
))...))..))))).))))).)))..))))).)))))))))).)))).))))......))))((((.....)
.))))....  (-67.9)
>46_14845_1_29181_Bifidobacterium_bifidum_S17,_complete_igr829.fa_1_l=227
_1081
CGGGCGAAGGUUGCAGAGAGGUUGGCUCGGAUUCCUCCCGACUCCGCUGCCGGGUGUUUUCGACUGGAUGCUA
AGUGGAGACGGUUCCAUAUCACGCGAAUAUGCCCAUCUAGGUGUCGUCUGUCCACAAAAACUGUGGAUAACAC
GCCGUGAUGUGGAUAAGUCGCGGGCCUUCGACCACAUUGUCGCCUCACGUUGCGCCGCUGUGGCGCUAUUCGU
UUCAAUCG
.((((((.(((((.((((....((((.((((.((.....)).))))..))))....)))))))))........
.((((.((.((..(........((((.....((((((..(((((.((.((((((((.....)))))))))))))
)))..))))).)).....))))))..)).))..))))....)))))).(((..(((((.....))))).....))))
........  (-79.8)
>46_3934301_1_3952661_Bifidobacterium_catenulatum_DSM_16992_igr11.fa_1_l=
227_311
UAUGGCCAAGAUAGGUCGUUACGUUGGACGGGCAUUGGAUAGACUUACCGUUACGACGAUGUUGUUGUGCAGA
GAUUCUGCUCUCAAGGUUCUUGCAAGGAAGGCCCGCUUCGGCGGGUCUUCCUUUUUUGUUUUGAGGCGUGCUG
GCAUGCGUCCGUCCCGUUUUCCGGUGACAGACGUGGAUAUUUGAUGCCCUGUGGAUAGCUGUGGAUGACACGC
CGUGGUUG
.....((((..(((....)))..))))..((((((..((((......(((..(((.............((((.
....)))).(((((..(......(((((((((((((....)))))))))))))....)..))))).))))..))
)...(((((.((((((.....))).))).)))))...)))))..)))))).......((((..(((........)
))..))))  (-84.5)
>54_13024_1_13199_Bifidobacterium_minimum_DSM_20102_rc_igr430.fa_1_l=227_
151
GAACAAAAAGGCCGAUCUUCGUGUUCGGAUUCGCUCAACGGGUCGGUUCAAACGAGAGAUUCGCGACAAAACA
GAUAAGUUAUGAACAAACGGGCUGAUGCGGUUGUGAACGGCGUGAUAACAGGCAUAAUAGGAGAUGCAAGCAC
AAGUUAAGAGAUGGAAGCCCACAAAGGUCAAUGGUCACGAGGACCAUCUGUUGGACGGGCAUUGGAUAGACUU
ACCGUCGC
..........((..(((((..((((.((((..(((.....)))..)))).)))).)))))..)).....((((
((................(((((....((.((...(((...(((.......((((.........))))...)))
..)))..))..))..)))))).....((((..((....))..)))).)))))).(((((...............
.))))))..  (-51.4)
>9_22110_1_19169_Bifidobacterium_bombi_DSM_19703_rc_igr864.fa_1_l=227_176
AUGGUAAUCCGCGGCUUCGUGGCUUUGGGAGUUGCGGUUCUUGCACUCGUUGUCAGGUGUCGGUCUGGUUUGC
CUUUUCUGCUGCUUUUUGUGAUGUGAACAUUUGGUUCUCAGUGCCAUCCGUGUGCGUUUUCAAGCGUUCGUUC
UCCUCUGAUGGGCGAACCUCAACAAUGCGAUUGUGACAUUGGACAGGUUUUCGAGCGUCCGUCCUUUUCUGAU
GGACGGAU

...((((.((((..((((.........))))..))))...)))).(((((((((..(((((.....(((((((
((.......((((...((.(((((..((...((((.......))))...))..)))))..)))))).....((
......)).)))))))))...(((((...))))))))))..)))))).....)))).((((((((.........
)))))))) (-66.7)
>
46_42441_1_60427_Bifidobacterium_choerinum_DSM_20434_rc_igr680.fa_1_l=227
_551
CAAUACCCGCCCAACGUAAUAAUCUUGCUGAUCAUAUACUUUUUGUGGGUCUGCUAUUCAGUUGUUCUUGUAC
UUGAACAUUAUGCUCGAUUCAUCGGUCUUCGCAAGUCGACGCGGCCGCCGAUCGGGCGAUGUUCGCAUUUUUU
CCAUUUUGUGCGCCGCACCCCCGGACGCAACAUGAAAACGAGCCAUAUUGUCCGUUACGGACGACCGUAUGUU
CAUUCGAA
....((((((..((.(((((.(((.....))).)).))).))..))))))......(((((((((((((((..
..(((((((..((((((....(((((..((((........))))..))))))))))))))))))((((.....
.......))))...))).....))).))))).))))...((((.((((((((((...))))))...)))))))
)....... (-57.6)
>43_2665120_1_2668098_Bifidobacterium_thermacidophilum_subsp._thermacidop
hilum_igr207.fa_1_l=227_176
GCGGCGUUAUCUUUACGUUCCUGUACGCGAGCCGACGCAACAACCCGAAGAUUCACGCGCAUCUGAGCGUCUC
CGGCGCCGCGGCCGGCCGGCAUCAGCCAUCUACGCGCCCAUCCCCCGGUUCCUGUAUCCCAGCCCACACAACA
ACUCGAAGAUACGUGCGCAUCCCCGGCAACGCGGCACAGAUGCGCACCCAAACGCGAUGUCCUGCACUGGAGG
GCAGACAG
...(((((.........(((.(((..(((......))).)))....))).......(((((((((..((....
))..(((((((((((..(((....))).....(((((..(((...(((((..(((............)))..)
)).))..)))..))))).....)))))..)))))).))))))))).....)))))..((((((.......)))
))).... (-68.6)

Supplementary File S10. Evolutionary conservation of predicted Bifidobacterium ms1 RNA homologs in related species. The lines contains names of species of the spot 42 candidates, introduced by “Cand.”, lists of species in which statistically significant BLAST hits of the candidate sequences in spot 42-syntenic IGRs were obtained follow the arrows.

Cand. Bifidobacterium animalis -> Bifidobacterium KRGSERBCFTRI, Bifidobacterium italicum Rab10, Bifidobacterium anseris Goo31D, Bifidobacterium anseris LH, Bifidobacterium AGR2158 G606DR, Bifidobacterium canis SY61, Bifidobacterium castoris 2020B, Bifidobacterium canis GSD1FS, Bifidobacterium choerinum FMB-, Bifidobacterium choerinum DSM, Bifidobacterium choerinum LMG, Bifidobacterium choerinum LH, Bifidobacterium choerinum LH, Bifidobacterium choerinum LH, Bifidobacterium choerinum Sow4, Bifidobacterium choerinum LH, Bifidobacterium choerinum DSM, Bifidobacterium RUG13504, Bifidobacterium criceti Ham19E, Bifidobacterium pseudolongum g, Bifidobacterium fecal__452 fec, Bifidobacterium pseudolongum L, Bifidobacterium pseudolongum p, Bifidobacterium pseudolongum p, Bifidobacterium pseudolongum 4, Bifidobacterium pseudolongum 4, Bifidobacterium pseudolongum 4, Bifidobacterium pseudolongum p, Bifidobacterium pseudolongum p, Bifidobacterium pseudolongum S, Bifidobacterium pseudolongum S, Bifidobacterium pseudolongum L, Bifidobacterium pseudolongum 4, Bifidobacterium pseudolongum p, Bifidobacterium pseudolongum p, Bifidobacterium pseudolongum p, Bifidobacterium pseudolongum p, Bifidobacterium pseudolongum p, Bifidobacterium pseudolongum p, Bifidobacterium pseudolongum g, Bifidobacterium pseudolongum A, Bifidobacterium pseudolongum g, Bifidobacterium pseudolongum g, Bifidobacterium pseudolongum g, Bifidobacterium pseudolongum M, Bifidobacterium pseudolongum g, Bifidobacterium pseudolongum g, Bifidobacterium pseudolongum g, Bifidobacterium pseudolongum P, Bifidobacterium pseudolongum g, Bifidobacterium pseudolongum g, Bifidobacterium

pseudolongum g, Bifidobacterium pseudolongum g, Bifidobacterium pseudolongum g, Bifidobacterium pseudolongum 2, Bifidobacterium pseudolongum N, Bifidobacterium pseudolongum Y, Bifidobacterium pseudolongum A, Bifidobacterium pseudolongum g, Bifidobacterium pseudolongum g, Bifidobacterium pseudolongum g, Bifidobacterium pseudolongum g, Bifidobacterium pseudolongum g, Bifidobacterium pseudolongum g, Bifidobacterium pseudolongum g, Bifidobacterium pseudolongum g, Bifidobacterium pseudolongum g, Bifidobacterium pseudolongum g, Bifidobacterium pseudolongum g, Bifidobacterium pseudolongum g, Bifidobacterium pseudolongum 1, Bifidobacterium pseudolongum g, Bifidobacterium pseudolongum g, Bifidobacterium pseudolongum g, Bifidobacterium S0468_NS37_NS8, Bifidobacterium fecal__148 fec, Bifidobacterium pseudolongum 9, Bifidobacterium pseudolongum 2, Bifidobacterium DOME__3616 361, Bifidobacterium RUG11560, Bifidobacterium RUG082, Bifidobacterium pseudolongum g, Bifidobacterium pseudolongum g, Bifidobacterium pseudolongum 2.
Cand. Bifidobacterium boum -> Bifidobacterium RUG11914, Bifidobacterium RUG13422, Bifidobacterium RUG10593, Bifidobacterium FUKU07_bin.013, Bifidobacterium thermophilum J, Bifidobacterium thermacidophil, Bifidobacterium CO.B43.metabat, Bifidobacterium thermophilum M, Bifidobacterium thermophilum R, Bifidobacterium thermophilum L, Bifidobacterium thermophilum L, Bifidobacterium thermophilum L, Bifidobacterium thermophilum L, Bifidobacterium thermophilum L, Bifidobacterium thermophilum M, Bifidobacterium thermophilum D, Bifidobacterium thermophilum J, Bifidobacterium thermophilum 1, Bifidobacterium thermophilum 1, Bifidobacterium thermophilum L, Bifidobacterium thermophilum B, Bifidobacterium LCP19S3_E9 con, Bifidobacterium LCP19S3_H1 con, Bifidobacterium thermacidophil, Bifidobacterium thermacidophil, Bifidobacterium porcinum JCM70, Bifidobacterium thermacidophil, Bifidobacterium thermacidophil, Bifidobacterium porcinum M521A, Bifidobacterium SUG2225 k141, Bifidobacterium thermacidophil, Bifidobacterium apri DSM, Bifidobacterium apri CCM, Bifidobacterium apri M106A, Bifidobacterium apri M114A, Bifidobacterium apri 115B, Bifidobacterium apri M63A, Bifidobacterium apri M64A.
Cand. Bifidobacterium pseudocatenula -> Bifidobacterium D5_SB_87 k141, Bifidobacterium ERR10149223_bi, Bifidobacterium D4_SB_77 k141, Bifidobacterium MSK.13.1 NODE, Bifidobacterium MSK.13.7 NODE, Bifidobacterium D6_SB_62 k141, Bifidobacterium catenulatum ka, Bifidobacterium D2_SB_85 k141, Bifidobacterium REFINED_METABA, Bifidobacterium catenulatum LC, Bifidobacterium catenulatum LC, Bifidobacterium catenulatum LC, Bifidobacterium catenulatum pu, Bifidobacterium catenulatum RT, Bifidobacterium catenulatum LC, Bifidobacterium catenulatum LC, Bifidobacterium catenulatum LC, Bifidobacterium catenulatum LC, Bifidobacterium catenulatum LC, Bifidobacterium catenulatum LC, Bifidobacterium catenulatum LC, Bifidobacterium catenulatum LC, Bifidobacterium catenulatum LC, Bifidobacterium catenulatum LC, Bifidobacterium catenulatum LC, Bifidobacterium catenulatum LC, Bifidobacterium catenulatum LC, Bifidobacterium catenulatum LC, Bifidobacterium catenulatum LC, Bifidobacterium catenulatum LC, Bifidobacterium catenulatum LC, Bifidobacterium catenulatum LC, Bifidobacterium catenulatum LC, Bifidobacterium catenulatum LC, Bifidobacterium catenulatum LC, Bifidobacterium catenulatum LC, Bifidobacterium catenulatum LC, Bifidobacterium catenulatum LC, Bifidobacterium catenulatum LC, Bifidobacterium catenulatum LC, Bifidobacterium P040-20 NODE, Bifidobacterium catenulatum RT, Bifidobacterium catenulatum RT, Bifidobacterium catenulatum RT, Bifidobacterium catenulatum RT,

Bifidobacterium catenulatum RT, Bifidobacterium catenulatum RT,
Bifidobacterium catenulatum RT, Bifidobacterium catenulatum RT,
Bifidobacterium catenulatum RT, Bifidobacterium catenulatum RT,
Bifidobacterium REFINED_METABA, Bifidobacterium catenulatum BI,
Bifidobacterium catenulatum BI, Bifidobacterium catenulatum 18,
Bifidobacterium catenulatum CL, Bifidobacterium catenulatum 22,
Bifidobacterium catenulatum ca, Bifidobacterium SRR17798043_bi,
Bifidobacterium N5G01 N5G01, Bifidobacterium gSpreadC_ID_24,
Bifidobacterium catenulatum pu, Bifidobacterium hominis Bg155.,
Bifidobacterium Map_89_012 con, Bifidobacterium hominis CLA-AA,
Bifidobacterium catenulatum RT, Bifidobacterium catenulatum RT,
Bifidobacterium catenulatum RT, Bifidobacterium catenulatum RT,
Bifidobacterium catenulatum RT, Bifidobacterium catenulatum MG,
Bifidobacterium hominis YIT, Bifidobacterium hominis YIT, Bifidobacterium catenulatum ca, Bifidobacterium hominis M264, Bifidobacterium hominis Bg4172, Bifidobacterium catenulatum pu, Bifidobacterium catenulatum ka,
Bifidobacterium MGYG-HGUT-0075, Bifidobacterium catenulatum 22,
Bifidobacterium catenulatum BS, Bifidobacterium catenulatum 22,
Bifidobacterium catenulatum 22, Bifidobacterium catenulatum BS,
Bifidobacterium catenulatum RT, Bifidobacterium catenulatum RT,
Bifidobacterium catenulatum RT, Bifidobacterium catenulatum 22,
Bifidobacterium N5_245_000G1_d, Bifidobacterium N4G05 N4G05,
Bifidobacterium catenulatum RC, Bifidobacterium catenulatum RC,
Bifidobacterium catenulatum ka, Bifidobacterium M3-R-103 conti,
Bifidobacterium catenulatum 22, Bifidobacterium DRR171859_bin_,
Bifidobacterium ERR10149218_bi, Bifidobacterium ERR10960878_bi.
Cand. Bifidobacterium pseudolongum -> Bifidobacterium S0468_NS37_NS8,
Bifidobacterium fecal__148 fec, Bifidobacterium DOME__3616 361,
Bifidobacterium RUG11560, Bifidobacterium RUG082, Bifidobacterium fecal__452 fec, Bifidobacterium anseris Goo31D, Bifidobacterium anseris LH, Bifidobacterium AGR2158 G606DR, Bifidobacterium choerinum FMB-,
Bifidobacterium choerinum DSM, Bifidobacterium choerinum LMG,
Bifidobacterium choerinum LH, Bifidobacterium choerinum LH,
Bifidobacterium choerinum LH, Bifidobacterium choerinum Sow4,
Bifidobacterium choerinum LH, Bifidobacterium choerinum DSM,
Bifidobacterium italicum Rab10, Bifidobacterium criceti Ham19E,
Bifidobacterium RUG13504, Bifidobacterium castoris 2020B, Bifidobacterium canis SY61, Bifidobacterium canis GSD1FS, Bifidobacterium animalis anima,
Bifidobacterium animalis YL2, Bifidobacterium animalis anima,
Bifidobacterium animalis anima, Bifidobacterium animalis A82,
Bifidobacterium animalis anima, Bifidobacterium animalis MRD34,
Bifidobacterium animalis anima, Bifidobacterium animalis anima,
Bifidobacterium animalis anima, Bifidobacterium animalis anima,
Bifidobacterium animalis anima, Bifidobacterium animalis anima,
Bifidobacterium animalis anima, Bifidobacterium animalis YL2,
Bifidobacterium animalis LH, Bifidobacterium animalis anima,
Bifidobacterium animalis lacti, Bifidobacterium animalis lacti,
Bifidobacterium animalis lacti, Bifidobacterium animalis lacti,
Bifidobacterium animalis CGMCC, Bifidobacterium animalis lacti,
Bifidobacterium animalis lacti, Bifidobacterium animalis lacti,
Bifidobacterium animalis lacti, Bifidobacterium animalis lacti,
Bifidobacterium animalis lacti, Bifidobacterium animalis LHCA2,
Bifidobacterium animalis LHCA2, Bifidobacterium animalis ATA-B,
Bifidobacterium animalis lacti, Bifidobacterium animalis CM18M,
Bifidobacterium animalis 10012, Bifidobacterium animalis C3,
Bifidobacterium animalis 10012, Bifidobacterium animalis D6,
Bifidobacterium animalis G1, Bifidobacterium animalis 10012,
Bifidobacterium animalis 10012, Bifidobacterium animalis 10012,

Bifidobacterium animalis B11, Bifidobacterium animalis B2, Bifidobacterium animalis 10012, Bifidobacterium animalis 10012, Bifidobacterium animalis C11, Bifidobacterium animalis D1, Bifidobacterium animalis A12, Bifidobacterium animalis lacti, Bifidobacterium animalis lacti, Bifidobacterium animalis 10012, Bifidobacterium animalis 10012, Bifidobacterium animalis 10012, Bifidobacterium animalis lacti, Bifidobacterium animalis lacti, Bifidobacterium animalis lacti, Bifidobacterium animalis lacti, Bifidobacterium animalis lacti, Bifidobacterium animalis lacti, Bifidobacterium animalis lacti, Bifidobacterium animalis lacti, Bifidobacterium animalis RC150, Bifidobacterium animalis lacti, Bifidobacterium animalis lacti, Bifidobacterium animalis ZK-77.
Cand. Bifidobacterium angulatum -> Bifidobacterium merycicum DSM, Bifidobacterium merycicum DSM, Bifidobacterium merycicum LMG, Bifidobacterium merycicum LH, Bifidobacterium merycicum S749, Bifidobacterium merycicum RUG1, Bifidobacterium merycicum DOME, Bifidobacterium merycicum SRR1, Bifidobacterium merycicum SRR1, Bifidobacterium merycicum RUG1, Bifidobacterium pongonis 64T4, Bifidobacterium panos DSM, Bifidobacterium stellenboschen, Bifidobacterium LH_421 LH, Bifidobacterium parmae Uis4E, Bifidobacterium platyrrhinorum, Bifidobacterium callitrichos R, Bifidobacterium callitrichos D, Bifidobacterium callitrichos D, Bifidobacterium callitrichos U, Bifidobacterium callitrichos L, Bifidobacterium callitrichos L, Bifidobacterium callitrichos L, Bifidobacterium UTCIF-38 conti, Bifidobacterium UTCIF-37 conti, Bifidobacterium saguinibicolor, Bifidobacterium CP2 Contig, Bifidobacterium LH_456 LH, Bifidobacterium LH_424 LH, Bifidobacterium LH_426 LH, Bifidobacterium LH_413 LH, Bifidobacterium LH_419 LH, Bifidobacterium aesculapii LH, Bifidobacterium rousetti LH, Bifidobacterium aesculapii DSM, Bifidobacterium rousetti RST9, Bifidobacterium rousetti LH, Bifidobacterium rousetti LH.
Cand. Bifidobacterium adolescentis -> Bifidobacterium KRGSERBCFTRI, Bifidobacterium REFINED_METABA, Bifidobacterium Gw_Prim_bin_23, Bifidobacterium ERR10149223_bi, Bifidobacterium MSK.17.10 NODE, Bifidobacterium REFINED_METABA, Bifidobacterium S06C.meta.bin_, Bifidobacterium D2_MB_49 k141, Bifidobacterium D3_SB_66 k141, Bifidobacterium REFINED_METABA, Bifidobacterium S07C.meta.bin_, Bifidobacterium S07B.meta.bin_, Bifidobacterium D1_SB_76 k141, Bifidobacterium D4_SB_20 k141, Bifidobacterium S08C.meta.bin_, Bifidobacterium S08B.meta.bin_, Bifidobacterium S08A.meta.bin_, Bifidobacterium UBA8278 contig, Bifidobacterium pseudocatenula, Bifidobacterium pseudocatenula, Bifidobacterium pseudocatenula, Bifidobacterium pseudocatenula, Bifidobacterium pseudocatenula, Bifidobacterium pseudocatenula, Bifidobacterium pseudocatenula, Bifidobacterium pseudocatenula, Bifidobacterium pseudocatenula, Bifidobacterium pseudocatenula, Bifidobacterium pseudocatenula, Bifidobacterium pseudocatenula, Bifidobacterium pseudocatenula, Bifidobacterium pseudocatenula, Bifidobacterium pseudocatenula, Bifidobacterium pseudocatenula, Bifidobacterium pseudocatenula, Bifidobacterium pseudocatenula, Bifidobacterium pseudocatenula, Bifidobacterium pseudocatenula, Bifidobacterium pseudocatenula, Bifidobacterium pseudocatenula, Bifidobacterium pseudocatenula, Bifidobacterium pseudocatenula, Bifidobacterium pseudocatenula, Bifidobacterium pseudocatenula, Bifidobacterium pseudocatenula, Bifidobacterium pseudocatenula, Bifidobacterium pseudocatenula, Bifidobacterium pseudocatenula, Bifidobacterium pseudocatenula, Bifidobacterium pseudocatenula, Bifidobacterium pseudocatenula, Bifidobacterium pseudocatenula, Bifidobacterium pseudocatenula, Bifidobacterium pseudocatenula,

Bifidobacterium pseudocatenula, Bifidobacterium pseudocatenula,
Bifidobacterium pseudocatenula, Bifidobacterium D4_SB_77 k141,
Bifidobacterium catenulatum ka, Bifidobacterium pseudocatenula,
Bifidobacterium pseudocatenula, Bifidobacterium pseudocatenula,
Bifidobacterium LH_743 LH, Bifidobacterium pseudocatenula,
Bifidobacterium pseudocatenula, Bifidobacterium pseudocatenula,
Bifidobacterium pseudocatenula, Bifidobacterium pseudocatenula,
Bifidobacterium pseudocatenula, Bifidobacterium pseudocatenula,
Bifidobacterium pseudocatenula, Bifidobacterium pseudocatenula,
Bifidobacterium pseudocatenula, Bifidobacterium pseudocatenula,
Bifidobacterium pseudocatenula, Bifidobacterium pseudocatenula,
Bifidobacterium pseudocatenula, Bifidobacterium D2_SB_85 k141,
Bifidobacterium D6_SB_62 k141, Bifidobacterium D5_SB_87 k141,
Bifidobacterium pseudocatenula, Bifidobacterium pseudocatenula,
Bifidobacterium pseudocatenula, Bifidobacterium pseudocatenula,
Bifidobacterium pseudocatenula, Bifidobacterium pseudocatenula,
Bifidobacterium pseudocatenula, Bifidobacterium pseudocatenula,
Bifidobacterium pseudocatenula, Bifidobacterium pseudocatenula,
Bifidobacterium pseudocatenula, Bifidobacterium pseudocatenula,
Bifidobacterium pseudocatenula, Bifidobacterium pseudocatenula,
Bifidobacterium pseudocatenula, Bifidobacterium pseudocatenula,
Bifidobacterium pseudocatenula, Bifidobacterium pseudocatenula.
Cand. Bifidobacterium ruminantium -> Bifidobacterium MGYG-HGUT-0345,
Bifidobacterium CM_BacBif1601., Bifidobacterium LH_743 LH,
Bifidobacterium adolescentis C, Bifidobacterium UBA8278 contig,
Bifidobacterium adolescentis C, Bifidobacterium adolescentis C,
Bifidobacterium adolescentis M, Bifidobacterium adolescentis M,
Bifidobacterium adolescentis M, Bifidobacterium adolescentis M,
Bifidobacterium adolescentis M, Bifidobacterium adolescentis S,
Bifidobacterium adolescentis S, Bifidobacterium adolescentis S,
Bifidobacterium adolescentis S, Bifidobacterium adolescentis S,
Bifidobacterium adolescentis S, Bifidobacterium adolescentis S,
Bifidobacterium adolescentis S, Bifidobacterium adolescentis S,
Bifidobacterium adolescentis S, Bifidobacterium adolescentis 2,
Bifidobacterium adolescentis 7, Bifidobacterium adolescentis S,
Bifidobacterium adolescentis 8, Bifidobacterium adolescentis 4,
Bifidobacterium adolescentis I, Bifidobacterium adolescentis I,
Bifidobacterium adolescentis 1, Bifidobacterium adolescentis 1,
Bifidobacterium adolescentis B, Bifidobacterium adolescentis D,
Bifidobacterium adolescentis 1, Bifidobacterium REFINED_METABA,
Bifidobacterium adolescentis 1, Bifidobacterium adolescentis B,
Bifidobacterium adolescentis B, Bifidobacterium adolescentis M,
Bifidobacterium adolescentis B, Bifidobacterium adolescentis R,
Bifidobacterium adolescentis D, Bifidobacterium adolescentis B,
Bifidobacterium adolescentis R, Bifidobacterium adolescentis R,
Bifidobacterium adolescentis B, Bifidobacterium adolescentis 1,
Bifidobacterium adolescentis R, Bifidobacterium adolescentis B,
Bifidobacterium adolescentis R, Bifidobacterium adolescentis B,
Bifidobacterium adolescentis R, Bifidobacterium adolescentis 1,
Bifidobacterium adolescentis B, Bifidobacterium adolescentis B,
Bifidobacterium adolescentis B, Bifidobacterium adolescentis C,
Bifidobacterium adolescentis C, Bifidobacterium adolescentis C,
Bifidobacterium adolescentis C, Bifidobacterium adolescentis C,
Bifidobacterium adolescentis C, Bifidobacterium adolescentis C,
Bifidobacterium adolescentis L, Bifidobacterium adolescentis B,
Bifidobacterium adolescentis E, Bifidobacterium adolescentis R,
Bifidobacterium adolescentis R, Bifidobacterium adolescentis R,
Bifidobacterium adolescentis 2, Bifidobacterium adolescentis 2,

Bifidobacterium adolescentis L, Bifidobacterium adolescentis L,
Bifidobacterium adolescentis L, Bifidobacterium adolescentis L,
Bifidobacterium adolescentis L, Bifidobacterium adolescentis E,
Bifidobacterium adolescentis E, Bifidobacterium adolescentis R,
Bifidobacterium D1_SB_76 k141, Bifidobacterium D2_MB_49 k141,
Bifidobacterium D3_SB_66 k141, Bifidobacterium D4_SB_20 k141,
Bifidobacterium adolescentis B, Bifidobacterium KRGSERBCFTRI,
Bifidobacterium adolescentis i, Bifidobacterium adolescentis C,
Bifidobacterium adolescentis L, Bifidobacterium adolescentis i,
Bifidobacterium adolescentis A, Bifidobacterium adolescentis A,
Bifidobacterium adolescentis S, Bifidobacterium adolescentis G,
Bifidobacterium Gw_Prim_bin_23, Bifidobacterium ERR10149223_bi,
Bifidobacterium adolescentis N.
Cand. Bifidobacterium dentium -> Bifidobacterium MSTE12 contig0,
Bifidobacterium N2_071_000G1_d, Bifidobacterium N3_173_000G1_d,
Bifidobacterium moukalabense G, Bifidobacterium moukalabense C,
Bifidobacterium moukalabense E, Bifidobacterium moukalabense D,
Bifidobacterium moukalabense G, Bifidobacterium moukalabense G,
Bifidobacterium moukalabense G, Bifidobacterium moukalabense G,
Bifidobacterium moukalabense G, Bifidobacterium moukalabense G,
Bifidobacterium moukalabense E, Bifidobacterium moukalabense C,
Bifidobacterium adolescentis L, Bifidobacterium adolescentis D,
Bifidobacterium adolescentis E, Bifidobacterium adolescentis L,
Bifidobacterium MSK.17.10 NODE, Bifidobacterium adolescentis M,
Bifidobacterium adolescentis M, Bifidobacterium adolescentis M,
Bifidobacterium adolescentis M, Bifidobacterium adolescentis M,
Bifidobacterium adolescentis M, Bifidobacterium adolescentis S,
Bifidobacterium adolescentis S, Bifidobacterium adolescentis S,
Bifidobacterium adolescentis S, Bifidobacterium adolescentis S,
Bifidobacterium adolescentis S, Bifidobacterium adolescentis 2,
Bifidobacterium adolescentis S, Bifidobacterium adolescentis 8,
Bifidobacterium adolescentis 4, Bifidobacterium adolescentis I,
Bifidobacterium adolescentis M, Bifidobacterium adolescentis D,
Bifidobacterium adolescentis 1, Bifidobacterium REFINED_METABA,
Bifidobacterium adolescentis K, Bifidobacterium adolescentis K,
Bifidobacterium adolescentis B, Bifidobacterium adolescentis D,
Bifidobacterium adolescentis S, Bifidobacterium adolescentis S,
Bifidobacterium adolescentis G, Bifidobacterium adolescentis 2,
Bifidobacterium adolescentis 1, Bifidobacterium adolescentis 1,
Bifidobacterium adolescentis 1, Bifidobacterium adolescentis M,
Bifidobacterium adolescentis B, Bifidobacterium adolescentis 1,
Bifidobacterium adolescentis R, Bifidobacterium adolescentis B,
Bifidobacterium adolescentis D, Bifidobacterium adolescentis 1,
Bifidobacterium adolescentis R, Bifidobacterium adolescentis R,
Bifidobacterium adolescentis R, Bifidobacterium adolescentis 1,
Bifidobacterium adolescentis R, Bifidobacterium adolescentis 1,
Bifidobacterium adolescentis R, Bifidobacterium adolescentis 1,
Bifidobacterium adolescentis 1, Bifidobacterium adolescentis 1,
Bifidobacterium adolescentis 1, Bifidobacterium adolescentis 1,
Bifidobacterium adolescentis C, Bifidobacterium adolescentis C,
Bifidobacterium adolescentis C, Bifidobacterium adolescentis C,
Bifidobacterium adolescentis C, Bifidobacterium adolescentis B,
Bifidobacterium adolescentis E, Bifidobacterium adolescentis R,
Bifidobacterium adolescentis 2, Bifidobacterium adolescentis 2,
Bifidobacterium adolescentis L, Bifidobacterium adolescentis L,
Bifidobacterium adolescentis E, Bifidobacterium D1_SB_76 k141,
Bifidobacterium D2_MB_49 k141, Bifidobacterium D3_SB_66 k141,
Bifidobacterium D4_SB_20 k141, Bifidobacterium adolescentis B,

Bifidobacterium KRGSERBCFTRI, Bifidobacterium adolescentis i, Bifidobacterium adolescentis L, Bifidobacterium adolescentis A, Bifidobacterium adolescentis A, Bifidobacterium adolescentis S, Bifidobacterium Gw_Prim_bin_23, Bifidobacterium ERR10149223_bi, Bifidobacterium adolescentis N.
Cand. Bifidobacterium moukalabense -> Bifidobacterium dentium MB0385, Bifidobacterium dentium MB0477, Bifidobacterium dentium MB0076, Bifidobacterium MSTE12 contig0, Bifidobacterium dentium N8, Bifidobacterium dentium EYE, Bifidobacterium N2_071_000G1_d, Bifidobacterium dentium AF54-1, Bifidobacterium dentium 181B, Bifidobacterium dentium 70T2, Bifidobacterium dentium BRDF, Bifidobacterium dentium JCM, Bifidobacterium dentium DE-29, Bifidobacterium dentium cx, Bifidobacterium dentium Bdenti, Bifidobacterium dentium JCM, Bifidobacterium dentium NCTC11, Bifidobacterium dentium ATCC, Bifidobacterium dentium LFYP24, Bifidobacterium N3_173_000G1_d, Bifidobacterium dentium AM96-0, Bifidobacterium dentium 924B, Bifidobacterium dentium 2124B, Bifidobacterium dentium 898B, Bifidobacterium dentium 2075B, Bifidobacterium dentium VB29, Bifidobacterium dentium JCM, Bifidobacterium dentium BIOML-, Bifidobacterium dentium BIOML-, Bifidobacterium dentium BIOML-, Bifidobacterium dentium Bd1, Bifidobacterium dentium RTP214, Bifidobacterium dentium RTP214, Bifidobacterium dentium RTP214, Bifidobacterium dentium RTP214, Bifidobacterium dentium 78T4, Bifidobacterium dentium 77T2, Bifidobacterium dentium 68T9, Bifidobacterium dentium 74T1, Bifidobacterium dentium MM0176, Bifidobacterium dentium MM0074, Bifidobacterium dentium MB0185, Bifidobacterium dentium MB0148, Bifidobacterium dentium MB0372, Bifidobacterium dentium MB0114, Bifidobacterium dentium 2201st, Bifidobacterium dentium 2201st, Bifidobacterium dentium 2201st, Bifidobacterium dentium UN03-6, Bifidobacterium dentium 2347B, Bifidobacterium dentium JCVIHM, Bifidobacterium dentium E7, Bifidobacterium dentium 1893B, Bifidobacterium dentium BSD278, Bifidobacterium dentium 100127, Bifidobacterium dentium LH, Bifidobacterium dentium UMB001, Bifidobacterium dentium BRDF, Bifidobacterium dentium 100129, Bifidobacterium dentium ATCC, Bifidobacterium dentium OM11-4, Bifidobacterium dentium 679B, Bifidobacterium dentium 793B, Bifidobacterium dentium 848B, Bifidobacterium dentium 831B, Bifidobacterium dentium 841B, Bifidobacterium dentium 2078B, Bifidobacterium dentium LHP009, Bifidobacterium dentium LHP008, Bifidobacterium dentium BRDF, Bifidobacterium dentium UT, Bifidobacterium dentium UT, Bifidobacterium dentium ATCC, Bifidobacterium dentium MB0224, Bifidobacterium dentium 2091B.
Cand. Bifidobacterium bifidum -> Bifidobacterium REFINED_METABA, Bifidobacterium REFINED_METABA, Bifidobacterium SKU, Bifidobacterium TKU, Bifidobacterium hapali DSM, Bifidobacterium saguinibicolor, Bifidobacterium simiiventris 8, Bifidobacterium simiiventris L, Bifidobacterium CP2 Contig, Bifidobacterium rousetti RST9, Bifidobacterium rousetti LH, Bifidobacterium aerophilum LH, Bifidobacterium goeldii 2034B, Bifidobacterium aerophilum TRE, Bifidobacterium amazonense MA1, Bifidobacterium samirii 2033B, Bifidobacterium biavatii DSM, Bifidobacterium biavatii DSM, Bifidobacterium rousetti LH, Bifidobacterium rousetti LH, Bifidobacterium goeldii 2034B, Bifidobacterium biavatii DSM, Bifidobacterium biavatii DSM, Bifidobacterium miconis 82T10, Bifidobacterium LH_730 LH, Bifidobacterium ramosum DSM, Bifidobacterium ramosum LH, Bifidobacterium reuteri LH, Bifidobacterium reuteri LH, Bifidobacterium ramosum TREM, Bifidobacterium pullorum JCM, Bifidobacterium pullorum DSM,

Bifidobacterium pullorum LMG, Bifidobacterium A104_L1_330330, Bifidobacterium D4_SB_67 k141, Bifidobacterium samirii 2033B, Bifidobacterium subtile KCTC, Bifidobacterium reuteri RST19, Bifidobacterium subtile DSM, Bifidobacterium subtile LMG, Bifidobacterium reuteri LH, Bifidobacterium subtile DSM, Bifidobacterium MGYG-HGUT-0338, Bifidobacterium scardovii 981, Bifidobacterium scardovii HROM, Bifidobacterium subtile UW, Bifidobacterium subtile UW, Bifidobacterium scardovii N2, Bifidobacterium scardovii N2, Bifidobacterium scardovii N5, Bifidobacterium scardovii L2, Bifidobacterium scardovii SRR9.
Cand. Bifidobacterium catenulatum -> Bifidobacterium SRR17798043_bi, Bifidobacterium pseudocatenula, Bifidobacterium pseudocatenula, Bifidobacterium pseudocatenula, Bifidobacterium pseudocatenula, Bifidobacterium pseudocatenula, Bifidobacterium pseudocatenula, Bifidobacterium hominis Bg155., Bifidobacterium Map_89_012 con, Bifidobacterium hominis YIT, Bifidobacterium hominis M264, Bifidobacterium MGYG-HGUT-0075, Bifidobacterium P040-20 NODE, Bifidobacterium REFINED_METABA, Bifidobacterium hominis CLA-AA, Bifidobacterium hominis Bg4172, Bifidobacterium N4G05 N4G05, Bifidobacterium ERR10960878_bi, Bifidobacterium N5G01 N5G01, Bifidobacterium gSpreadC_ID_24, Bifidobacterium M3-R-103 conti, Bifidobacterium hominis SS, Bifidobacterium hominis 306, Bifidobacterium hominis YIT, Bifidobacterium N5_245_000G1_d, Bifidobacterium DRR171859_bin_, Bifidobacterium ERR10149218_bi, Bifidobacterium hominis Bg064., Bifidobacterium UMGS603, Bifidobacterium REFINED_METABA, Bifidobacterium pseudocatenula, Bifidobacterium pseudocatenula, Bifidobacterium pseudocatenula, Bifidobacterium pseudocatenula, Bifidobacterium pseudocatenula, Bifidobacterium pseudocatenula, Bifidobacterium pseudocatenula, Bifidobacterium pseudocatenula, Bifidobacterium pseudocatenula, Bifidobacterium pseudocatenula, Bifidobacterium pseudocatenula, Bifidobacterium pseudocatenula, Bifidobacterium pseudocatenula, Bifidobacterium pseudocatenula, Bifidobacterium pseudocatenula, Bifidobacterium pseudocatenula, Bifidobacterium pseudocatenula, Bifidobacterium pseudocatenula, Bifidobacterium pseudocatenula, Bifidobacterium pseudocatenula, Bifidobacterium pseudocatenula, Bifidobacterium pseudocatenula, Bifidobacterium pseudocatenula, Bifidobacterium pseudocatenula, Bifidobacterium pseudocatenula, Bifidobacterium pseudocatenula, Bifidobacterium pseudocatenula, Bifidobacterium pseudocatenula, Bifidobacterium pseudocatenula, Bifidobacterium pseudocatenula, Bifidobacterium pseudocatenula, Bifidobacterium pseudocatenula, Bifidobacterium pseudocatenula, Bifidobacterium pseudocatenula, Bifidobacterium pseudocatenula, Bifidobacterium pseudocatenula, Bifidobacterium pseudocatenula, Bifidobacterium pseudocatenula, Bifidobacterium pseudocatenula, Bifidobacterium pseudocatenula, Bifidobacterium pseudocatenula, Bifidobacterium pseudocatenula, Bifidobacterium pseudocatenula, Bifidobacterium pseudocatenula, Bifidobacterium pseudocatenula, Bifidobacterium pseudocatenula, Bifidobacterium pseudocatenula, Bifidobacterium pseudocatenula, Bifidobacterium pseudocatenula, Bifidobacterium pseudocatenula, Bifidobacterium pseudocatenula, Bifidobacterium pseudocatenula, Bifidobacterium pseudocatenula, Bifidobacterium pseudocatenula, Bifidobacterium pseudocatenula, Bifidobacterium pseudocatenula, Bifidobacterium pseudocatenula, Bifidobacterium pseudocatenula, Bifidobacterium pseudocatenula, Bifidobacterium D6_SB_62 k141, Bifidobacterium pseudocatenula, Bifidobacterium pseudocatenula, Bifidobacterium pseudocatenula, Bifidobacterium pseudocatenula, Bifidobacterium pseudocatenula, Bifidobacterium pseudocatenula, Bifidobacterium pseudocatenula, Bifidobacterium pseudocatenula, Bifidobacterium pseudocatenula, Bifidobacterium pseudocatenula.

Cand. Bifidobacterium minimum -> Bifidobacterium aquikefiricola, Bifidobacterium WK041_bin.7 NO, Bifidobacterium fermentum WK01, Bifidobacterium WK004_bin.7 NO, Bifidobacterium WK013_bin.6 NO, Bifidobacterium WK033_bin.11 N, Bifidobacterium WK044_bin.6 NO, Bifidobacterium WK048_bin.6 NO, Bifidobacterium WK045_bin.8 NO, Bifidobacterium WK025_bin.2 NO, Bifidobacterium fermentum WK04, Bifidobacterium fermentum WK01, Bifidobacterium crudilactis UW, Bifidobacterium WK016_bin.5 NO, Bifidobacterium WK015_bin.5 NO, Bifidobacterium WK021_bin.7 NO, Bifidobacterium crudilactis LM, Bifidobacterium crudilactis AO, Bifidobacterium crudilactis NA, Bifidobacterium crudilactis UW, Bifidobacterium crudilactis UW, Bifidobacterium crudilactis UW, Bifidobacterium crudilactis AS, Bifidobacterium crudilactis 12, Bifidobacterium crudilactis 13, Bifidobacterium crudilactis 11, Bifidobacterium crudilactis 13, Bifidobacterium crudilactis 12, Bifidobacterium crudilactis 10, Bifidobacterium crudilactis 11, Bifidobacterium crudilactis 61, Bifidobacterium crudilactis 86, Bifidobacterium crudilactis 59, Bifidobacterium crudilactis UW, Bifidobacterium crudilactis UW, Bifidobacterium crudilactis UW, Bifidobacterium crudilactis SR, Bifidobacterium aquikefiri LMG, Bifidobacterium aquikefiri WK0, Bifidobacterium aquikefiri WK0, Bifidobacterium aquikefiri WK0, Bifidobacterium aquikefiri WK0, Bifidobacterium aquikefiri WK0, Bifidobacterium aquikefiri WK0, Bifidobacterium aquikefiri WK0, Bifidobacterium aquikefiri WK0, Bifidobacterium aquikefiri WK0, Bifidobacterium aquikefiri WK0, Bifidobacterium aquikefiri WK0, Bifidobacterium aquikefiri WK0, Bifidobacterium aquikefiri WK0, Bifidobacterium aquikefiri WK0, Bifidobacterium aquikefiri WK0, Bifidobacterium aquikefiri WK0, Bifidobacterium aquikefiri WK0, Bifidobacterium aquikefiri WK0, Bifidobacterium aquikefiri WK0, Bifidobacterium aquikefiri WK0, Bifidobacterium aquikefiri WK0, Bifidobacterium aquikefiri ERR, Bifidobacterium mongoliense DS, Bifidobacterium mongoliense DS, Bifidobacterium mongoliense SU, Bifidobacterium mongoliense 22, Bifidobacterium mongoliense 22, Bifidobacterium mongoliense 22, Bifidobacterium mongoliense 22, Bifidobacterium mongoliense 22, Bifidobacterium mongoliense 22, Bifidobacterium mongoliense 22, Bifidobacterium mongoliense 20, Bifidobacterium mongoliense 22, Bifidobacterium mongoliense 22, Bifidobacterium mongoliense BM, Bifidobacterium mongoliense 12, Bifidobacterium mongoliense 59, Bifidobacterium mongoliense 62, Bifidobacterium mongoliense 64, Bifidobacterium mongoliense UW, Bifidobacterium mongoliense UW, Bifidobacterium mongoliense UW, Bifidobacterium mongoliense UW, Bifidobacterium mongoliense UW, Bifidobacterium mongoliense UW, Bifidobacterium mongoliense UW, Bifidobacterium mongoliense UW, Bifidobacterium mongoliense UW, Bifidobacterium mongoliense 30, Bifidobacterium psychraerophil, Bifidobacterium psychraerophil, Bifidobacterium psychraerophil, Bifidobacterium psychraerophil, Bifidobacterium psychraerophil, Bifidobacterium psychraerophil, Bifidobacterium psychraerophil, Bifidobacterium psychraerophil, Bifidobacterium psychraerophil, Bifidobacterium psychraerophil.
Cand. Bifidobacterium choerinum -> Bifidobacterium italicum Rab10, Bifidobacterium castoris 2020B, Bifidobacterium anseris Goo31D, Bifidobacterium anseris LH, Bifidobacterium AGR2158 G606DR, Bifidobacterium criceti Ham19E, Bifidobacterium RUG13504, Bifidobacterium pseudolongum Y, Bifidobacterium pseudolongum A, Bifidobacterium pseudolongum g, Bifidobacterium pseudolongum g, Bifidobacterium pseudolongum g, Bifidobacterium pseudolongum g, Bifidobacterium pseudolongum g, Bifidobacterium pseudolongum g, Bifidobacterium

```
pseudolongum g, Bifidobacterium pseudolongum g, Bifidobacterium
pseudolongum g, Bifidobacterium pseudolongum g, Bifidobacterium
pseudolongum g, Bifidobacterium pseudolongum g, Bifidobacterium
pseudolongum g, Bifidobacterium pseudolongum g, Bifidobacterium
pseudolongum g, Bifidobacterium pseudolongum g, Bifidobacterium
pseudolongum g, Bifidobacterium pseudolongum g, Bifidobacterium
pseudolongum g, Bifidobacterium pseudolongum g, Bifidobacterium
pseudolongum g, Bifidobacterium pseudolongum g, Bifidobacterium
pseudolongum g, Bifidobacterium pseudolongum g, Bifidobacterium
pseudolongum g, Bifidobacterium pseudolongum g, Bifidobacterium
pseudolongum g, Bifidobacterium pseudolongum g, Bifidobacterium
pseudolongum g, Bifidobacterium pseudolongum g, Bifidobacterium
pseudolongum M, Bifidobacterium pseudolongum 1, Bifidobacterium
pseudolongum p, Bifidobacterium pseudolongum g, Bifidobacterium
pseudolongum 4, Bifidobacterium pseudolongum 4, Bifidobacterium
pseudolongum 4, Bifidobacterium pseudolongum g, Bifidobacterium
pseudolongum g, Bifidobacterium pseudolongum g, Bifidobacterium
S0468_NS37_NS8, Bifidobacterium pseudolongum S, Bifidobacterium
pseudolongum S, Bifidobacterium pseudolongum g, Bifidobacterium
fecal__148 fec, Bifidobacterium pseudolongum 9, Bifidobacterium
pseudolongum 2, Bifidobacterium pseudolongum L, Bifidobacterium
pseudolongum L, Bifidobacterium RUG11560, Bifidobacterium RUG082,
Bifidobacterium pseudolongum 4, Bifidobacterium pseudolongum p,
Bifidobacterium pseudolongum g, Bifidobacterium pseudolongum g,
Bifidobacterium pseudolongum g, Bifidobacterium pseudolongum p,
Bifidobacterium pseudolongum p, Bifidobacterium pseudolongum p,
Bifidobacterium pseudolongum g, Bifidobacterium pseudolongum g,
Bifidobacterium pseudolongum g, Bifidobacterium pseudolongum g,
Bifidobacterium pseudolongum g, Bifidobacterium pseudolongum p,
Bifidobacterium pseudolongum 2, Bifidobacterium pseudolongum 2,
Bifidobacterium pseudolongum p, Bifidobacterium pseudolongum g,
Bifidobacterium pseudolongum N.
```

Supplementary File S11. Matlab scripts implementing the presented structure search, synteny and conservation filtering.

```
Structure search

Main scripts

rsol.m:
-> rsol_module.m:
   -> f_sample_sq_by_sopts_to_tm__no_PDF_Unaf.m:
      -> f_reformat_RNAsubopt.m, f_scan_RNAdistance_Xf.m;
-> merge_mats_app.m
   -> f_get_RhoTermPredict_py.m, f_read_tts_from_standard_file.m,
f_read_ARNold_into_standard_file.m;
-> present_results_table_no_stats.m, histo.m, btstrp.m,
f_get_conserved_region.m,
draw_run_f_sample_sq_by_sopts_to_tm_simple_csd_rgn.m.

Bootstrap scripts

btstrp.m:
-> f_bt_strp.m

General functions
```

f_find_str_in_cells.m, cut_string.m, f_remove_substrs_from_str_rsol.m.

Matlab Code

rsol.m

```
function rsol(varargin)
pid__ = num2str(feature('getpid'));
STEP__ = 5;
disp([newline 'RNASubOptLoc - an app for localization of homologous RNA
gene candidates on genomic sequences using suboptimal secondary
structures'])
fnsqs__ = []; fntm__ = []; ttfn__ = [];
WP__ = '';
message__ = ['Usage: ' char(10) ...
'For help:' char(10) ...
'-h(elp) for help.' newline newline...
'For new search:' newline ...
'./rsol.sh -sqs=dir/file.fa -tm=dir/file.br (-tt=dir/file.ext -
SAMPLINGLENGTH -DO_TERMINATORS=[0]|1 -DO_3_END_U=0|[1] -
NO_OF_PROCESSES=<1,[number of cores|number of query sequences]> -
GRAPHICAL_OUTPUT=0|[1] -SAVE_FIGURES=[0]|1 -
SHOW_ONLY_ORIENTATION_MATCHING_TEMPLATE=[0]|1 -TEST=[0]|1  -
NUMBER_OF_BEST_SETS_TO_SHOW=<[0],6> -DO_ALTERNATIVE_CANDIDATES=<[0],6-
NUMBER_OF_BEST_SETS_TO_SHOW> -SHOW_SEQUENCE_SIMILARITY_REGIONS=[0]|1 -
DO_OVERLAP_WITH_SEQUENCE_SIMILARITY_REGIONS=[0]|1 -ALPHA=[0.02]).'
newline newline ...
'For displaying & analyzing saved search: ' newline ...
'./rsol.sh -m(at)=dir/file.mat (-GRAPHICAL_OUTPUT=0|[1] -
SAVE_FIGURES=[0]|1 -SHOW_ONLY_ORIENTATION_MATCHING_TEMPLATE=[0]|1 -
TEST=[0]|1 -NUMBER_OF_BEST_SETS_TO_SHOW=<[0],6> -
DO_ALTERNATIVE_CANDIDATES=<[0],6-NUMBER_OF_BEST_SETS_TO_SHOW> -
HISTOGRAM=[0]|1 -SHOW_SEQUENCE_SIMILARITY_REGIONS=[0]|1 -
DO_OVERLAP_WITH_SEQUENCE_SIMILARITY_REGIONS=[0]|1 -ALPHA[=0.02] -
BOOTSTRAP[=0])' newline newline ...
'Required arguments:' char(10) ...
'-sqs: fasta file with input sequences' char(10) ...
'-tm: file with template structure in dot-bracket format' newline ...
'-m(at): file with saved search (MAT file).' newline newline ...
'Optional arguments:' char(10) ...
'-tt: file with Rho-independent terminators in standard format' char(10)
...
'-SAMPLINGLENGTH: sampling length; default: length of template; max.:
length of input sequences' char(10) ...
'-DO_TERMINATORS=[0]|1: if 1, display Rho-independent terminators, when -
tt set, try to predict using RhoTermPredict.py and display, when -tt
unset; if 0, do nothing' char(10) ...
'-DO_3_END_U=0|[1]: show uracils/thymines in last 10% of input sequences
as clue for presence of Rho-independent terminators' char(10) ...
'-NO_OF_PROCESSES=<1,[number of cores|number of query sequences]>: number
of processor cores user for computing' newline ...
'-GRAPHICAL_OUTPUT=0|[1]: make figures with interactive UI' char(10) ...
'-SAVE_FIGURES=[0]|1: save figures in EPS format' newline ...
'-SHOW_ONLY_ORIENTATION_MATCHING_TEMPLATE=[0]|1: generate output with
candidates with sequence orientation matching template only' char(10) ...
```

```
'-DO_OVERLAP_WITH_SEQUENCE_SIMILARITY_REGIONS=[0]|1: compute regions with
sequence similarity to template in input sequences, if exists, using
BLAST alignment and candidates overlapping it and show them in
interactive figures' char(10) ...
'-TEST=[0]|1: do two-sample t-test of best candidates against other,
worse candidates' newline ...
'-NUMBER_OF_BEST_SETS_TO_SHOW=<[0],6>: no. of best sets to show. Sum with
alternative candidates must be <= 6.' newline ...
'-DO_ALTERNATIVE_CANDIDATES=<[0],6-NUMBER_OF_BEST_SETS_TO_SHOW>: compute
gene candidates predicted using best single suboptimal structures along
those predicted using sets of suboptimal structures' newline ...
'-HISTOGRAM=[0]|1: show a diagram comparing scores (TEDs or TED-based
scores) of best candidates to scores of the other candidates for
individual input sequences' newline ...
'-SHOW_SEQUENCE_SIMILARITY_REGIONS=[0]|1: display regions with sequence
similarity to template' newline ...
'-ALPHA [0.02]: significance level for two-sample t-test' newline ...
'-DIST_LETT=[f]|h|w|c|F|H|W|C|P: distance representation in RNAdistance'
newline ...
'-BOOTSTRAP [0]: if = 0, no bootstrap; if = 1, use already computed
bootstrap, if > 9, compute new bootstrap (min. bootstrap rounds = 10)'
newline ...
'-MAXSTRSPERSQ' newline ...
'-WINDOWSIZE' newline ...
'-PROCENTFE' newline];

if isempty(varargin) | strcmp(lower(varargin{1}), '-h') |
strcmp(lower(varargin{1}), '-help')
      disp(message__);
      return
end

[v v] = unix('which RNAplot');
if contains(v, 'no RNAplot in')
      disp([newline 'No RNA Vienna package installed. Stop.']), return
else
      if strfind(lower(varargin{1}), '-m')
            [v v] = unix('RNAplot -V');
            v(length(v)) = [];
            s = strsplit(v, '.');
            v = strsplit(v, ' ');
            if str2num(s{2}) < 5
                  disp([newline 'Too old version of RNA Vienna package ('
v{2} ') installed. No structure figures will be available due to RNAplot
flaws. Ignoring.' newline])
                  TOO_OLD_RNA_VIENNA = 1;
            end
      end
end
[v v] = unix('which blastn');
[x x] = unix('find ~/ -name "blastn"');
if isempty(strfind(x, 'bin/blastn')) & ~isempty(strfind(v, 'no blastn
in'))
    cnsd_rgn = 'blastn not installed/configured. Install it or put on
PATH. Ignoring.';
      BLASTNPATH = 'No BLAST.';
else
      if ~contains(v, 'no blastn in')
```

```
            BLASTNPATH = '';
      else
            x=strsplit(x,newline);
            x = x{find(contains(x, 'bin/blastn'))};
            i = find(x == '/');
            BLASTNPATH = x(1 : i(length(i)));
      end
end

[CT2B_DIR CT2B_DIR] = unix('echo $DIR');
if CT2B_DIR(length(CT2B_DIR)) == newline, CT2B_DIR(length(CT2B_DIR)) =
[]; end
if CT2B_DIR(length(CT2B_DIR)) == '/', CT2B_DIR = [CT2B_DIR
'rsol_module/application/']; else, CT2B_DIR = [CT2B_DIR
'/rsol_module/application/']; end

if strfind(lower(varargin{1}), '-m')
      fnload_ = strtrim(varargin{1}(find(varargin{1} == '=') + 1 :
length(varargin{1})));
      if ~exist(fnload_, 'file')
            disp([newline fnload_ ' not exist. Stop.']); return
      end
      i__ = find(fnload_ == '.');
      disp([char(10) 'Doing ' fnload_ char(10)])
      load(fnload_)
      i = find(fnload_ == '/');
      if ~isempty(i)
            WP__ = fnload_(1 : i(length(i)));
      end
      for i__ = 2 : length(varargin)
            if ~isnumeric(varargin{i__}) & any(varargin{i__} == '=') &
any(varargin{i__} == '-')
                  name__ = strtrim(varargin{i__}(1 : find(varargin{i__} ==
'=') - 1));
                  switch name__
                        case '-SAVE_FIGURES'
                              SAVE_FIGURES =
str2num(varargin{i__}(find(varargin{i__} == '=') + 1 :
length(varargin{i__})));
                              name__ = [];
                        case '-TEST'
                              TEST =
str2num(varargin{i__}(find(varargin{i__} == '=') + 1 :
length(varargin{i__})));
                              name__ = [];
                        case '-BOOTSTRAP'
                              BOOTSTRAP =
str2num(varargin{i__}(find(varargin{i__} == '=') + 1 :
length(varargin{i__})));
                              name__ = [];
                        case '-GRAPHICAL_OUTPUT'
                              GRAPHICAL_OUTPUT =
str2num(varargin{i__}(find(varargin{i__} == '=') + 1 :
length(varargin{i__})));
                              name__ = [];
                        case '-KEEP_DBS'
```

```
                        KEEP_DBS =
str2num(varargin{i__}(find(varargin{i__} == '=') + 1 :
length(varargin{i__})));
                        name__ = [];
                    case '-SHOW_ONLY_ORIENTATION_MATCHING_TEMPLATE'
                        SHOW_ONLY_ORIENTATION_MATCHING_TEMPLATE =
str2num(varargin{i__}(find(varargin{i__} == '=') + 1 :
length(varargin{i__})));
                        name__ = [];
                    case '-DO_ALTERNATIVE_CANDIDATES'
                        DO_ALTERNATIVE_CANDIDATES =
str2num(varargin{i__}(find(varargin{i__} == '=') + 1 :
length(varargin{i__})));
                        name__ = [];
                    case '-HISTOGRAM'
                        HISTOGRAM =
str2num(varargin{i__}(find(varargin{i__} == '=') + 1 :
length(varargin{i__})));
                        name__ = [];
                    case '-SHOW_SEQUENCE_SIMILARITY_REGIONS'
                        SHOW_SEQUENCE_SIMILARITY_REGIONS =
str2num(varargin{i__}(find(varargin{i__} == '=') + 1 :
length(varargin{i__})));
                        name__ = [];
                    case '-
DO_OVERLAP_WITH_SEQUENCE_SIMILARITY_REGIONS'
                        DO_OVERLAP_WITH_SEQUENCE_SIMILARITY_REGIONS =
str2num(varargin{i__}(find(varargin{i__} == '=') + 1 :
length(varargin{i__})));
                        name__ = [];
                    case '-ALPHA'
                        ALPHA__ =
str2num(varargin{i__}(find(varargin{i__} == '=') + 1 :
length(varargin{i__})));
                        name__ = [];
                    case '-NUMBER_OF_BEST_SETS_TO_SHOW'
                        NUMBER_OF_BEST_SETS_TO_SHOW =
str2num(varargin{i__}(find(varargin{i__} == '=') + 1 :
length(varargin{i__})));
                        name__ = [];
                    case '-DIST_LETT'
                        DIST_LETT =
strtrim(varargin{i__}(find(varargin{i__} == '=') + 1 :
length(varargin{i__})));
                        name__ = [];
                    case '-MAXSTRSPERSQ'
                        MAXSTRSPERSQ =
str2num(varargin{i__}(find(varargin{i__} == '=') + 1 :
length(varargin{i__})));
                        name__ = [];
                    case '-PROCENTFE'
                        PROCENTFE =
str2num(varargin{i__}(find(varargin{i__} == '=') + 1 :
length(varargin{i__})));
                        name__ = [];
                    case '-WINDOWSIZE'
```

```
                            WINDOWSIZE =
str2num(varargin{i__}(find(varargin{i__} == '=') + 1 :
length(varargin{i__})));
                            name__ = [];
                end
                if ~isempty(name__)
                      disp(['Unknown argument ' num2str(i__) '. Ignoring
it.'])
                      return
                end
          else
                disp(['Invalid argument ' num2str(i__) '. Stop.'])
                return
          end
     end

     if ~exist('TEST'), TEST = 0; end
     if ~exist('BOOTSTRAP'), BOOTSTRAP = 0; end
     if ~exist('GRAPHICAL_OUTPUT'), GRAPHICAL_OUTPUT = 1; end
     if ~exist('KEEP_DBS'), KEEP_DBS = 1; end
     if ~exist('SAVE_FIGURES'), SAVE_FIGURES = 0; end
     if ~exist('SHOW_ONLY_ORIENTATION_MATCHING_TEMPLATE'),
SHOW_ONLY_ORIENTATION_MATCHING_TEMPLATE = 0; end
     if ~exist('DO_ALTERNATIVE_CANDIDATES'), DO_ALTERNATIVE_CANDIDATES =
0; end
     if ~exist('HISTOGRAM', 'var'), HISTOGRAM = 0; end
     if ~exist('SHOW_SEQUENCE_SIMILARITY_REGIONS'),
SHOW_SEQUENCE_SIMILARITY_REGIONS = 0; end
     if ~exist('DO_OVERLAP_WITH_SEQUENCE_SIMILARITY_REGIONS'),
DO_OVERLAP_WITH_SEQUENCE_SIMILARITY_REGIONS=0; end
     if ~exist('ALPHA__'), ALPHA__=0.02; end
     if ~exist('NUMBER_OF_BEST_SETS_TO_SHOW'),
NUMBER_OF_BEST_SETS_TO_SHOW = 1; end
     if ~exist('DIST_LETT'), DIST_LETT = 'f'; end
     if ~exist('MAXSTRSPERSQ'), MAXSTRSPERSQ = 60; end
     if ~exist('PROCENTFE'), PROCENTFE = 2000; end
     if ~exist('WINDOWSIZE'), WINDOWSIZE = 2; end



     if ~GRAPHICAL_OUTPUT, SAVE_FIGURES = 0; end
else
     for i__ = 1 : length(varargin)
          if ~isnumeric(varargin{i__}) & any(varargin{i__} == '=') &
any(varargin{i__} == '-')
                name__ = strtrim(varargin{i__}(1 : find(varargin{i__} ==
'=') - 1));
                switch name__
                      case '-sqs'
                            fnsqs__ =
strtrim(varargin{i__}(find(varargin{i__} == '=') + 1 :
length(varargin{i__})));
                            name__ = [];
                      case '-tm'
                            fntm__ =
strtrim(varargin{i__}(find(varargin{i__} == '=') + 1 :
length(varargin{i__})));
                            name__ = [];
```

```
                        case '-tt'
                               ttfn__ =
strtrim(varargin{i__}(find(varargin{i__} == '=') + 1 :
length(varargin{i__})));
                               name__ = [];
            %           case '-SCORINGSCHEME'
            %                 SCORINGSCHEME =
str2num(varargin{i__}(find(varargin{i__} == '=') + 1 :
length(varargin{i__})));
                        case '-SAMPLINGLENGTH'
                              SAMPLINGLENGTH__ =
str2num(varargin{i__}(find(varargin{i__} == '=') + 1 :
length(varargin{i__})));
                              name__ = [];
                        case '-
DO_OVERLAP_WITH_SEQUENCE_SIMILARITY_REGIONS'
                              DO_OVERLAP_WITH_SEQUENCE_SIMILARITY_REGIONS =
str2num(varargin{i__}(find(varargin{i__} == '=') + 1 :
length(varargin{i__})));
                              name__ = [];
                        case '-DO_TERMINATORS'
                              DO_TERMINATORS =
str2num(varargin{i__}(find(varargin{i__} == '=') + 1 :
length(varargin{i__})));
                              name__ = [];
                        case '-DO_3_END_U'
                              DO_3_END_U =
str2num(varargin{i__}(find(varargin{i__} == '=') + 1 :
length(varargin{i__})));
                              name__ = [];
                        case '-NO_OF_PROCESSES'
                              NO_OF_PROCESSES =
str2num(varargin{i__}(find(varargin{i__} == '=') + 1 :
length(varargin{i__})));
                              name__ = [];
                        case '-SAVE_FIGURES'
                              SAVE_FIGURES =
str2num(varargin{i__}(find(varargin{i__} == '=') + 1 :
length(varargin{i__})));
                              name__ = [];
                        case '-TEST'
                              TEST =
str2num(varargin{i__}(find(varargin{i__} == '=') + 1 :
length(varargin{i__})));
                              name__ = [];
                        case '-BOOTSTRAP'
                              BOOTSTRAP =
str2num(varargin{i__}(find(varargin{i__} == '=') + 1 :
length(varargin{i__})));
                              name__ = [];
                        case '-GRAPHICAL_OUTPUT'
                              GRAPHICAL_OUTPUT =
str2num(varargin{i__}(find(varargin{i__} == '=') + 1 :
length(varargin{i__})));
                              name__ = [];
                        case '-KEEP_DBS'
```

```
                              KEEP_DBS =
str2num(varargin{i__}(find(varargin{i__} == '=') + 1 :
length(varargin{i__})));
                              name__ = [];
                        case '-SHOW_ONLY_ORIENTATION_MATCHING_TEMPLATE'
                              SHOW_ONLY_ORIENTATION_MATCHING_TEMPLATE =
str2num(varargin{i__}(find(varargin{i__} == '=') + 1 :
length(varargin{i__})));
                              name__ = [];
                        case '-DO_ALTERNATIVE_CANDIDATES'
                              DO_ALTERNATIVE_CANDIDATES =
str2num(varargin{i__}(find(varargin{i__} == '=') + 1 :
length(varargin{i__})));
                              name__ = [];
                        case '-HISTOGRAM'
                              HISTOGRAM =
str2num(varargin{i__}(find(varargin{i__} == '=') + 1 :
length(varargin{i__})));
                              name__ = [];
                        case '-SHOW_SEQUENCE_SIMILARITY_REGIONS'
                              SHOW_SEQUENCE_SIMILARITY_REGIONS =
str2num(varargin{i__}(find(varargin{i__} == '=') + 1 :
length(varargin{i__})));
                              name__ = [];
                        case '-ALPHA'
                              ALPHA__ =
str2num(varargin{i__}(find(varargin{i__} == '=') + 1 :
length(varargin{i__})));
                              name__ = [];
                        case '-NUMBER_OF_BEST_SETS_TO_SHOW'
                              NUMBER_OF_BEST_SETS_TO_SHOW =
str2num(varargin{i__}(find(varargin{i__} == '=') + 1 :
length(varargin{i__})));
                              name__ = [];
                        case '-DIST_LETT'
                              DIST_LETT =
strtrim(varargin{i__}(find(varargin{i__} == '=') + 1 :
length(varargin{i__})));
                              name__ = [];
                        case '-MAXSTRSPERSQ'
                              MAXSTRSPERSQ =
str2num(varargin{i__}(find(varargin{i__} == '=') + 1 :
length(varargin{i__})));
                              name__ = [];
                        case '-PROCENTFE'
                              PROCENTFE =
str2num(varargin{i__}(find(varargin{i__} == '=') + 1 :
length(varargin{i__})));
                              name__ = [];
                        case '-WINDOWSIZE'
                              WINDOWSIZE =
str2num(varargin{i__}(find(varargin{i__} == '=') + 1 :
length(varargin{i__})));
                              name__ = [];
                  end
                  if ~isempty(name__)
                        disp(['Unknown argument ' num2str(i__) '. Stop.'])
                        return
```

```
                end
            else
                disp(['Invalid argument ' num2str(i__) '. Stop.'])
                return
            end
        end

        SCORINGSCHEME=0;% if ~exist('SCORINGSCHEME'), SCORINGSCHEME=0; end
        if ~exist('DO_OVERLAP_WITH_SEQUENCE_SIMILARITY_REGIONS'),
DO_OVERLAP_WITH_SEQUENCE_SIMILARITY_REGIONS=0; end
        if ~exist('DO_TERMINATORS'), DO_TERMINATORS=0; end
        if ~exist('DO_3_END_U'), DO_3_END_U=1; end
        if ~exist('NO_OF_PROCESSES'), NO_OF_PROCESSES=feature('numcores');
end
        if ~exist('SAVE_FIGURES'), SAVE_FIGURES = 0; end
        if ~exist('TEST'), TEST = 0; end
        if ~exist('BOOTSTRAP'), BOOTSTRAP = 0; end
        if ~exist('GRAPHICAL_OUTPUT'), GRAPHICAL_OUTPUT = 1; end
        if ~exist('KEEP_DBS'), KEEP_DBS = 1; end
        if ~exist('SHOW_ONLY_ORIENTATION_MATCHING_TEMPLATE'),
SHOW_ONLY_ORIENTATION_MATCHING_TEMPLATE = 0; end
        if ~exist('DO_ALTERNATIVE_CANDIDATES'), DO_ALTERNATIVE_CANDIDATES =
0; end
        if ~exist('HISTOGRAM', 'var'), HISTOGRAM = 0; end
        if ~exist('SHOW_SEQUENCE_SIMILARITY_REGIONS'),
SHOW_SEQUENCE_SIMILARITY_REGIONS = 0; end
        if ~exist('ALPHA__'), ALPHA__ = 0.02; end
        if ~exist('NUMBER_OF_BEST_SETS_TO_SHOW'),
NUMBER_OF_BEST_SETS_TO_SHOW = 1; end
        if ~exist('DIST_LETT'), DIST_LETT = 'f'; end
        if ~exist('MAXSTRSPERSQ'), MAXSTRSPERSQ = 60; end
        if ~exist('PROCENTFE'), PROCENTFE = 2000; end
        if ~exist('WINDOWSIZE'), WINDOWSIZE = 2; end

        if ~GRAPHICAL_OUTPUT, SAVE_FIGURES = 0; end

        if isempty(fnsqs__) | isempty(fntm__) |
~strfind(lower(varargin{1}), '-m')
            disp([newline '-sqs and/or -tm, or -m(at) argument(s) is/are
missing. Stop.' newline message__])
            return
        end
        disp([char(10) 'Input sequences: ' fnsqs__ char(10) 'Template: '
fntm__ char(10)])
        if exist(fntm__, 'file')
% FIND SAME IDS IN INPUT SQS AND ADD A SIGN TO THEM TO DISTINGUISH THEM
            id__ = fopen(fntm__); t = textscan(id__, '%s', 'delimiter',
char(10)); fclose(id__);
            id__ = fopen(fntm__); x = fread(id__); fclose(id__);
            if x(length(x)) ~= char(10)
                disp(['Newline character missing in the end of template
file. Stop.']);
                return
            end
            if ~exist('SAMPLINGLENGTH__'), SAMPLINGLENGTH__ =
length(t{1}{2});end
            if exist(fnsqs__, 'file')
                if ~any(fnsqs__ == '/')
```

```
                        fnsqs__ = [pwd '/' fnsqs__];
                  end
                  if ~any(fntm__ == '/')
                        fntm__ = [pwd '/' fntm__];
                  end
                  sx__ = cut_string(fnsqs__, '/');
                  ssx__ = cut_string(fntm__, '/');
                  fnload_ = ['rsol_' pid__ '_' sx__{length(sx__)} '__'
ssx__{length(ssx__)} '__' num2str(SAMPLINGLENGTH__) '.mat'];
                  unix(['rm rsol_' sx__{length(sx__)} '__'
ssx__{length(ssx__)} '__' num2str(SAMPLINGLENGTH__) '*.mat >> /dev/null
2>&1']);
% CHECK IF THE FASTA FILE IS ALL RIGHT
                  id__ = fopen(fnsqs__); f__ = textscan(id__, '%s',
'delimiter', char(10)); fclose(id__);
                  if isempty(f__{1})
                        disp([fnsqs__ ' is empty. Stop.']), return
                  end
                  if rem(length(f__{1}), 2)
                        disp(['Wrong fasta. ' fnsqs__ ' is not dividable
by 2. Stop.']);
                        return
                  end
                  for if__ = 1 : 2 : length(f__{1})
                        if f__{1}{if__}(1) ~= '>' | f__{1}{if__ + 1}(1) ==
'>'
                              disp(['Wrong fasta. Mixed headers and
sequences in ' fnsqs__ ' at sequence ' num2str(if__) '. Stop.']);
                              return
                        end
                        if ~isempty(strfind(f__{1}{if__}, '  ')) |
~isempty(strfind(f__{1}{if__ + 1}, '  '))
                              disp(['Wrong fasta. Bad data in ' fnsqs__ '
at sequence ' num2str(if__) '. Stop.']);
                              return
                        end
                  end
% TO RENAME THE SAME IDS
% FIRST FIND OUT AND ASK - ALL RUNNING RSOLS WILL STOP HERE BEFORE ONE OF
THEM STARTS WRITING INTO THE fnsqs__ FILE.
                  f__ = fastaread(fnsqs__);
                  for if__ = 1 : length(f__)
                        s__ = cut_string(f__(if__).Header, ' ');
%                       k__ = f_find_str_in_cells({f__.Header}, s__{1},
1);
                        k__ = [];
                        for ixx__ = 1 : length(f__)
                              x__ = cut_string(f__(ixx__).Header, ' ');
                              if strcmp(x__{1}, s__{1});
                                    k__ = [k__ ixx__];
                              end
                        end
                        if length(k__) > 1
%                             reply = input(['Identical ID ('
f__(if__).Header ') for sequences #' num2str(k__) ' in ' fnsqs__ '.
Rename them? [y]/n'], 's');
%                             if reply == 'y'
% THEN RENAME
```

```
                            id__ = fopen(fnsqs__, 'w');
                            while ~isempty(f__)
                                 s__ = cut_string(f__(1).Header, '
');
%                                k__ =
f_find_str_in_cells({f__.Header}, s__{1});
                                 k__ = [];
                                 for ixx__ = 1 : length(f__)
                                      x__ =
cut_string(f__(ixx__).Header, ' ');
                                      if strcmp(x__{1}, s__{1});
                                           k__ = [k__ ixx__];
                                      end
                                 end
                                 if length(k__) > 1
                                      for l__ = 1 : length(k__)
                                           s__ =
cut_string(f__(k__(l__)).Header, ' ');
                                           ss__ =
cut_string(s__{1}, '.');
                                           name__ = [ss__{1} '.'
num2str(l__)];
                                           if length(ss__) > 1
                                                name__ = [name__
'.' ss__{2}];
                                           end
                                           for is__ = 2 :
length(s__)
                                                name__ = [name__
' ' s__{is__}];
                                           end
                                           fwrite(id__, ['>'
name__  char(10) f__(k__(l__)).Sequence char(10)]);
                                      end
                                 else
                                      fwrite(id__, ['>'
f__(1).Header char(10) f__(1).Sequence char(10)]);
                                 end
                                 f__(k__) = [];
                            end
                            fclose(id__);
                            break
%                      else
%                           disp('Rename manually. Stop.')
%                           return
%                      end
                  end
             end
%FIND HOW MANY PROCESSES TO RUN
             sqs__ = fastaread(fnsqs__);
             if NO_OF_PROCESSES > length(sqs__), NO_OF_PROCESSES =
length(sqs__); end
             a__ = ones(1, NO_OF_PROCESSES) .* floor(length(sqs__) /
NO_OF_PROCESSES);
             a__(1 : rem(length(sqs__), NO_OF_PROCESSES)) = a__(1 :
rem(length(sqs__), NO_OF_PROCESSES)) + 1;
             DO_THESE_OF_SQS__ = cell(1, NO_OF_PROCESSES);
             DO_THESE_OF_SQS__{1} = 1 : a__(1);
```

```
                  for i__ = 2 : length(a__)
                        for ii__ = 1 : a__(i__)
                              DO_THESE_OF_SQS__{i__} =
[DO_THESE_OF_SQS__{i__} DO_THESE_OF_SQS__{i__ -
1}(length(DO_THESE_OF_SQS__{i__ - 1})) + ii__];
                        end
                  end
% RUN PROCESSES
                  pause('on')
                  [MR MR] = unix('echo $MR'); if MR(length(MR)) ==
newline, MR(length(MR)) = []; end
                  [x__ x__] = unix('tty');
                  ptts__ = cell(1, length(DO_THESE_OF_SQS__));
                  for iDO_THESE_OF_SQS__ = 1 : length(DO_THESE_OF_SQS__)
                        disp(['Searching input sequence(s) '
num2str(DO_THESE_OF_SQS__{iDO_THESE_OF_SQS__}) ' in process #'
num2str(iDO_THESE_OF_SQS__) '.'])
                        ptts__{iDO_THESE_OF_SQS__} = [pid__ '_'
num2str(iDO_THESE_OF_SQS__)];
                        [w w] = unix([CT2B_DIR 'run_rsol_module.sh ' MR
...
                                        ' -sqs=' fnsqs__ ' -tm=' fntm__ ' -
SAMPLINGLENGTH=' num2str(SAMPLINGLENGTH__) ...
                                           ' -DO_THESE_OF_SQS=''['
num2str(DO_THESE_OF_SQS__{iDO_THESE_OF_SQS__}) ']'' -ptt='
ptts__{iDO_THESE_OF_SQS__} ...
                                           ' -STEP=' num2str(STEP__) ' -
dist_lett=' num2str(DIST_LETT) ...
                                           ' -MAXSTRSPERSQ='
num2str(MAXSTRSPERSQ) ' -WINDOWSIZE=' num2str(WINDOWSIZE) ' -PROCENTFE='
num2str(PROCENTFE) ...
                                           ' > rsol_' sx__{length(sx__)}
'_' ssx__{length(ssx__)} '_' num2str(SAMPLINGLENGTH__) '_'
num2str(iDO_THESE_OF_SQS__) ...
                                           '_' ptts__{iDO_THESE_OF_SQS__}
'.inf 2>&1 &']);
% for testing rsol_module as a MATLAB function script (have to change
rsol_module.m too)
%                       rsol_module(fnsqs__, fntm__, SAMPLINGLENGTH__,
DO_THESE_OF_SQS__{iDO_THESE_OF_SQS__}, ptts__{iDO_THESE_OF_SQS__},
STEP__, DIST_LETT);
                  end
% WAIT FOR PROCESSES HAVE TIME TO GET STARTED
                  pause(2);
                  while 1
                        [x__ x__] = unix('ps -fu $whoami|grep
''rsol_module '' |grep -v ''grep''');
                        finished__ = [];
                        for iptts__ = 1 : length(ptts__)
                              if ~contains(x__, ['-ptt=' ptts__{iptts__}])
                                    disp(['Process -ptt=' ptts__{iptts__} '
finished.'])
                                    finished__ = [finished__ iptts__];
                              end
                        end
                        ptts__(finished__) = [];
                        if ~isempty(ptts__)
                              pause(2)
```

```
                    else
                        break
                    end
                end

                FLAG__ = 1;
                merge_mats_app
                if ~FLAG__
                    return
                end
% % you have to save these names with path if it is not in them
%               i = find(fnsqs__ == '/');
%               y = fnsqs__;
%               if isempty(i)
%                   y = [pwd '/' fnsqs__];
%               end
%               i = find(fntm__ == '/');
%               x = fntm__;
%               if isempty(i)
%                   x = [pwd '/' fntm__];
%               end
%               save(fnload_,'x','y',
'SAMPLINGLENGTH','DO_3_END_U','DO_ALTERNATIVE_CANDIDATES','ALPHA__','-
append');
                save(fnload_,'fntm__','fnsqs__',
'SAMPLINGLENGTH','DO_3_END_U','DO_ALTERNATIVE_CANDIDATES','ALPHA__','-
append');
            else
              disp(['No input sequences ' fnsqs__ '. Stop.']), return
            end
        else
            disp(['No template ' fntm__ '. Stop.']), return
        end
end
if ~exist('cnsd_rgn_') & (SHOW_SEQUENCE_SIMILARITY_REGIONS |
DO_OVERLAP_WITH_SEQUENCE_SIMILARITY_REGIONS)
        cnsd_rgn_ = f_get_conserved_region(fntm__, fnsqs__, BLASTNPATH);
        if ischar(cnsd_rgn_)
            disp(cnsd_rgn_)
            return
        else
            save(fnload_, 'cnsd_rgn_', '-append');
        end
end
if ~exist('isqsor__') & exist('cnsd_rgn_')
% FIND THOSE WITH TEMPLATE-MATCHING ORIENTATION FOR OUTPUTS
        REMOVEFROMHEADERS = {'MAG '; 'MAG: '; 'TPA_asm ' ; 'TPA_asm: ';
'candidatus '; 'sp. '; 'isolate '; ' bacterium '; 'UNVERIFIED_ORG: ';
'Sample ';};
        sqs = fastaread(fnsqs__);
        isqsor__ = [];
        for isqs = 1 : length(sqs)
            s = f_remove_substrs_from_str_rsol(sqs(isqs).Header,
REMOVEFROMHEADERS, 0, 0);
            n = strsplit(s, ' ');
%           cnsd_rgn_i = f_find_str_in_cells(cnsd_rgn_(:,1), n{1});
            cnsd_rgn_i = find(contains(cnsd_rgn_(:,1), n{1}));
            if isempty(cnsd_rgn_i)
```

```
                isqsor__ = [isqsor__ isqs];
            else
                for i = 1 : length(cnsd_rgn_i)
                    if cnsd_rgn_{cnsd_rgn_i(i), 4} == '+'
                        isqsor__ = [isqsor__ isqs];
                    end
                end
            end
        end
        save(fnload_, 'isqsor__', '-append');
end

% inner par: for h, h_or_p_ = 0, for p, h_or_p_ = 1
h_or_p_ = 0;

if TEST & BOOTSTRAP
        disp([newline 'TEST and BOOTSTRAP parameters both set to be done
(both > 0). Impossible to use both. Proceeding with BOOTSTRAP.'])
        TEST = 0;
end
if TEST
        psc_ = zeros(1, length(fn_best_sopts_)); pc_ = psc_; hsc_ = psc_;
hc_ = psc_;
% this is just to put all scores into a single vector, as there is no
conversion from cells to vector, fuck
        x= []; xx = x;
        for isqs = 1 : length(fn_best_sopts_)
            x = [x sc_{isqs}];
            xx = [xx c_{isqs}];
        end
        for isqs = 1 : length(fn_best_sopts_)
% YOU SHOULD NOT TEST AGAINST NON-BESTS OF ALL INPUT SEQUENCES...
% here you test all-best ones against all scores of all other input
sequences.
            [hsc_(isqs) psc_(isqs)] = ttest2(sc_{isqs}(sci_{isqs}(1)), x,
'Alpha', ALPHA__);
            [hc_(isqs) pc_(isqs)] = ttest2(min(c_{isqs}), xx, 'Alpha',
ALPHA__);
% ... BUT ONLY WITHIN INDIVIDUAL INPUT SEQUENCES
% here you test best ones against non-best ones within indiidual input
sequences.
%           [hsc_(isqs) psc_(isqs)] = ttest2(sc_{isqs}(sci_{isqs}(1)),
sc_{isqs});
%          [hc_(isqs) pc_(isqs)] = ttest2(min(c_{isqs}), c_{isqs});
        end
        save(fnload_, 'psc_', 'pc_', '-append');
end
if BOOTSTRAP > 1 & BOOTSTRAP <= 9
        disp(['BOOTSTRAP set to ' num2str(BOOTSTRAP) '. Min. bootstrap
rounds is 10. Proceeded.'])
        BOOTSTRAP = 10;
end
if BOOTSTRAP > 9
        btstrp
        if sum(c_rand_sopt_) == 0 | sum(sc_rand_set_) == 0
            disp('Something wrong in btstrp.m. Stop.');
            return
        end
```

```
      save(fnload_, 'sc_rand_set_', 'c_rand_sopt_', 'scs_rand_set_',
'scs_rand_sopt_', '-append');
end
if BOOTSTRAP == 1 | BOOTSTRAP > 9
      if exist('sc_rand_set_')
% YOU CAN HAVE PSC_ AND PC_ OVERWRITTEN WITH TEST, SO YOU SHOULD COMPUTE
THEM AGAIN from already computed sc_rand_set_ and c_rand_sopt_
      psc_ = zeros(1, length(fn_best_sopts_)); pc_ = psc_; hsc_ = psc_;
hc_ = psc_;
% this is just to put all scores into a single vector
      x = reshape(sc_rand_set_, [size(sc_rand_set_, 2) *
length(fn_best_sopts_) 1]);
      xx = reshape(c_rand_sopt_, [size(c_rand_sopt_, 2) *
length(fn_best_sopts_) 1]);
      for isqs = 1 : length(fn_best_sopts_)
% here you test all-best ones against all scores of all other input
sequences.
            [hsc_(isqs) psc_(isqs)] = ttest2(sc_{isqs}(sci_{isqs}(1)), x,
'Alpha', ALPHA__);
            [hc_(isqs) pc_(isqs)] = ttest2(min(c_{isqs}), xx, 'Alpha',
ALPHA__);
% here you test best ones against non-best ones within indiidual input
sequences.
%           [hsc_(isqs)  psc_(isqs)] = ttest2(sc_{isqs}(sci_{isqs}(1)),
sc_rand_set_(isqs, :));
%           [hc_(isqs) pc_(isqs)] = ttest2(min(c_{isqs}),
c_rand_sopt_(isqs, :));
      end
      save(fnload_, 'psc_', 'pc_', '-append');
      else
            disp('No bootstrap. Compute bootstrap first using -
BOOTSTRAP>=10. Stop.'), return
      end
end
present_results_table_no_stats
if GRAPHICAL_OUTPUT
      draw_run_f_sample_sq_by_sopts_to_tm_simple_csd_rgn
end
if HISTOGRAM
      if TEST | BOOTSTRAP
            if BOOTSTRAP & ~exist('sc_rand_set_')
                  disp('No bootstrap. Compute bootstrap first using -
BOOTSTRAP parameter. Stop.'), return
            else
                  histo
            end
      else
            disp('Say whether t-test (-TEST=1) or bootstrap (-BOOTSTRAP>0)
scores should be histogrammed. Stop.'), return
      end
end
% if exist('sc_rand_set_')
% %   ALPHA=.001;
%     if exist('ALPHA', 'var')
% %         for isqs = 1 : length(fn_best_sopts_)
%                 [h p]=ttest2(sc_{1}(sci_{1}(1)), sc_rand_set_,
'alpha',ALPHA);
%                 disp(['Best set sc: ' num2str([h p]) '.']);
```

```
%               [h p]=ttest2(c_{1}(sci_{1}(1)), c_rand_sopt_,
'alpha',ALPHA);
%               disp(['Best set c: ' num2str([h p]) '.']);
%               [h p]=ttest2(min(c_{1}), c_rand_sopt_, 'alpha',ALPHA);
%               disp(['Best-at-all c: ' num2str([h p]) '.']);
% %         end
%     else
% %         for isqs = 1 : length(fn_best_sopts_)
%               [h p]=ttest2(sc_{1}(sci_{1}(1)), sc_rand_set_);
%               disp(['Best set sc: ' num2str(p) '.']);
%               [h p]=ttest2(c_{1}(sci_{1}(1)), c_rand_sopt_);
%               disp(['Best set c: ' num2str(p) '.']);
%               [h p]=ttest2(min(c_{1}), c_rand_sopt_);
%               disp(['Best-at-all c: ' num2str(p) '.']);
% %   end
%     end
% end
% if ~GRAPHICAL_OUTPUT & ~KEEP_DBS
%     disp(['Deleting *.db...']);
%     for i = 1 : length(fn_best_sopts_)
%         for j = 1 : length(fn_best_sopts_{i})
%               eval(['!rm ' fn_best_sopts_{i}{j}])
%         end
%     end
% end
end
```

rsol_module.m

```
% % The rev. complementary strand has not been kept with the forward
strand. It was last saved on Jul 12th 2024.

function rsol_module(varargin)
varargin{:}

fn_ = strtrim(varargin{1}(find(varargin{1} == '=') + 1 :
length(varargin{1})));
fntm_ = strtrim(varargin{2}(find(varargin{2} == '=') + 1 :
length(varargin{2})));
SAMPLINGLENGTH = str2num(varargin{3}(find(varargin{3} == '=') + 1 :
length(varargin{3})));
DO_THESE_OF_SQS = str2num(varargin{4}(find(varargin{4} == '=') + 1 :
length(varargin{4})));
ptt = strtrim(varargin{5}(find(varargin{5} == '=') + 1 :
length(varargin{5})));
STEP = str2num(varargin{6}(find(varargin{6} == '=') + 1 :
length(varargin{6})));
dist_lett = strtrim(varargin{7}(find(varargin{7} == '=') + 1 :
length(varargin{7})));
MAXSTRSPERSQ = str2num(varargin{8}(find(varargin{8} == '=') + 1 :
length(varargin{8})));
WINDOWSIZE = str2num(varargin{9}(find(varargin{9} == '=') + 1 :
length(varargin{9})));
PROCENTFE = str2num(varargin{10}(find(varargin{10} == '=') + 1 :
length(varargin{10})));

% % FOR RUNNING RSOL_MODULE IN NONCOMPILED STATE
% fn_ = varargin{1};
```

```
% fntm_ = varargin{2};
% ptt = varargin{5};
% SAMPLINGLENGTH = varargin{3};
% DO_THESE_OF_SQS = varargin{4};
% STEP = varargin{6};
% dist_lett = varargin{7};

fn_
fntm_
SAMPLINGLENGTH
DO_THESE_OF_SQS
ptt
STEP
dist_lett
MAXSTRSPERSQ
WINDOWSIZE
PROCENTFE

[CT2B_DIR CT2B_DIR] = unix('echo $DIR');
if CT2B_DIR(length(CT2B_DIR)) == newline, CT2B_DIR(length(CT2B_DIR)) =
[]; end
if CT2B_DIR(length(CT2B_DIR)) == '/', CT2B_DIR = [CT2B_DIR
'rsol_module/application/']; else, CT2B_DIR = [CT2B_DIR
'/rsol_module/application/']; end

% DO_THESE_OF_SQS = 'all';
CONTINUE_COMPUTATION = 0;

%
fn_='spot42_EcoK12_IGR.fa',fntm_='~/hs/spot42_Eco.rf';SAMPLINGLENGTH=119;
DO_THESE_OF_SQS=1;

% spot42

% % 1st example
% fntm_ = '~/hs/spot42_Eco.rf'; SAMPLINGLENGTH = 120;

% should have been for validation but unfortunatelly, I couldn't find the
genomes
% fn_ = 'spot42_Escherichia_in_RF00021.fa';
% % Escherichias not in Rfam got by BLAST of known E. coli K-12 spot42
that is in RF00021: (? Escherichia_coli_K-12_in_G-prot.bo ->
% % Escherichia_coli_K-12_in_G-prot.bo.fa ->) Escherichia_IGRs.bo ->
Escherichia_IGRs.bo.fa -> Escherichia_IGRs.bo.fa.igr - for validation /
paper -
% except for first E. coli, they are not in Rfam
% fn_ = 'Escherichia_IGRs.bo.fa.igr';
% fn_ = 'Escherichia_IGRs.bo.fa_rc.igr';

% fn_ = 'Arsukibacterium_from_manual_BLAST_IGRs.fa.edtd.igr';
% fn_ = 'Arsukibacterium_from_manual_BLAST_IGRs.fa.edtd_rc.igr';

% fn_ = 'Rheinheimera_IGRs.bo.fa.igr';
% fn_ = 'Rheinheimera_IGRs.bo.fa_rc.igr';

% ms1

% tm = '~/hs/Msmest.br';
```

```
% % % tm = '~/hs/Ms1_Sco_published.fa.br';
% SAMPLINGLENGTH = 220;
% % fn_= 'Kineosporia_IGRs.bo.fa.igr';
% % fn_= 'Gordonia.igr';
% fn_= 'Streptomyces.igr';

% Y RNA

% the predicted structures of Ys in Rfam are quite different...? See 'Y
RNA Aves from RF00019.br'.
% fntm_ = '~/cl/Y_Ldi.fa.br'; SAMPLINGLENGTH = 112; %
'>JJRK01039655.1/21487-21376 Leptosomus discolor contig39655, whole
genome shotgun sequence.' in Y_RNA_Aves_in_RF00019.fa
% fntm_ = '~/cl/Y_Hal.fa.br'; SAMPLINGLENGTH = 111; % '>JJRL01013369.1/1-
111 Haliaeetus albicilla contig13369, whole genome shotgun sequence.' in
Y_RNA_Aves_in_RF00019.fa
% fntm_ = '~/cl/Y_Hal_9.fa.br'; SAMPLINGLENGTH = 102; %
'>9_JJRL01040244.1 Haliaeetus albicilla contig40244 2122 2021' in
Y_RNA_Aves_in_RF00019.fa.edtd

% % for validation
% % fn_='~/cl/Y_RNA_Aves_in_RF00019.fa.edtd.igr';
% fn_='~/cl/Y_RNA_Aves_in_RF00019.fa.edtd_rc.igr';

% 3rd example
% fn_= '~/cl/Haliaeetus_albicilla_Y_RNA_in_selected_Aves.bo.fa.igr';
% fn_= '~/cl/Haliaeetus_albicilla_Y_RNA_in_selected_Aves.bo.fa_rc.igr';
% fn_= 'Haliaeetus albicilla Y RNA in Acomys russatus chromosome
10.bo.fa.igr';

% fn_= '~/cl/Y_Hal_in_Acomys_russatus_bad_hits_plus_chrom_22.bo.fa';
% fntm_ = '~/cl/Y_Hal.fa.br';

sqs = fastaread(fn_);
if strcmp(DO_THESE_OF_SQS, 'all'), DO_THESE_OF_SQS = 1 : length(sqs); end
if ~CONTINUE_COMPUTATION
      sc_ = cell(1, length(sqs)); sci_= sc_; c_= sc_; fn_pdf_ = sc_;
fn_pdf_c_ = sc_; fn_best_sopts_= sc_; scs_ = sc_;
end
%
% addpath(pwd);
% cd(workdir_);

for iii = 1 : length(DO_THESE_OF_SQS)
      isqs = DO_THESE_OF_SQS(iii);

      s = cut_string(fn_, '/');
      pid = num2str(feature('getpid'));
      name = [ptt '_' pid '_' s{length(s)} '_' num2str(isqs)]
      disp(['Doing ' name ', i.e. ' num2str(isqs) ' of '
num2str(DO_THESE_OF_SQS) '.'])
      id = fopen(name, 'w');
      fwrite(id, ['>' ptt '_' pid '_' sqs(isqs).Header char(10)
sqs(isqs).Sequence char(10)]);
      fclose(id);
% create a copy of template for a single IGR sq to be able to run more
copies of the f_sample_sq_by_sopts_to_tm.m function with different IGR
sqs in a single .igr file
```

```
      r = strsplit(fntm_, '/');
      fntmlocal = [ptt '_' pid '_' r{length(r)}];
      ss = strsplit(fntmlocal, '.');
      unix(['echo ''>''' ss{1} ' > ' fntmlocal]);
      unix(['sed -n ''2,3p'' < ' fntm_ ' | cat >> ' fntmlocal]);
% the function is for one sequence
%
tmfn,      sqfn, SAMPLINGLENGTH, STEP,
RNASUBOPT,SCORINGSCHEME,DISPLAYPDF,VERBOSE,MAKEPDFS,MAKEALLPDFS,CT2B_DIR,
DO_DBS, dist_let
%     [sc sci fn_pdf c fn_pdf_c fn_best_sopts scs] =
f_sample_sq_by_sopts_to_tm(fntmlocal, name, SAMPLINGLENGTH, STEP,   0,
0,            0,          0,      0,       0,          CT2B_DIR, 1);
%     [sc sci c fn_best_sopts scs] = f_sample_sq_by_sopts_to_tm__no_PDF(
fntmlocal, name, SAMPLINGLENGTH, STEP,   0,        0,
0,                           CT2B_DIR, 1, dist_lett);
      [sc sci c fn_best_sopts scs] =
f_sample_sq_by_sopts_to_tm__no_PDF_Unaf(   fntmlocal, name,
SAMPLINGLENGTH, STEP,   0,        0,                        0,
CT2B_DIR, 1, dist_lett, MAXSTRSPERSQ, WINDOWSIZE, PROCENTFE);
      unix(['rm ' fntmlocal]);
      unix(['rm ' name]);

% sc - evaluated scores acc. to SCORINGSCHEME for sopts sets for each
sampled sq. sci - indexes of sorted scores in sc. sc is not sorted. ii -
indexes of sopts best sim to template in each set of sopts.
% c - scores (i.e. tree edit distances) of sopts most sim to template in
each set of sopts.
      if ~isempty(sc)
            if isnumeric(sc)
      % store all pdfs and indexes, when MAKEALLPDFS == 1, for
eval_run_f_sample_sq_by_sopts_to_tm.m
%                 fn_pdf_{isqs} = fn_pdf;
%                 fn_pdf_c_{isqs} = fn_pdf_c;
                  sc_{isqs} = sc;
                  sci_{isqs} = sci;
                  c_{isqs} = c;
                  fn_best_sopts_{isqs} = fn_best_sopts;
                  scs_{isqs} = scs;
      % get best sopt at all
                  [x xi]=min(c);
%                 if isempty(fn_pdf)
                        disp(['Best set #' num2str(sci(1)) ' with sc='
num2str(sc(sci(1))) ' with sopt with TED=' num2str(c(sci(1))) ...
                              '. Sopt most sim to tm at all is in set #'
num2str(xi) ' with TED=' num2str(x) '.'])
%                 else
%                       disp(['Best set: ' fn_pdf{sci(1)} ' #'
num2str(sci(1)) ' with sc=' num2str(sc(sci(1))) ' with sopt with TED = '
num2str(c(sci(1))) ...
%                             '. Sopt most sim to tm at all is in set #'
num2str(xi) ' ' fn_pdf_c{xi} ' with TED=' num2str(x) '.'])
%                 end
            else
                  sc_{isqs} = sc;
            end
      end
end
```

```
ss = cut_string(fntm_, '/');
ptt = cut_string(ptt, '_');
s = ['rsol_' ptt{1} '_' s{length(s)} '__'  ss{length(ss)} '__'
num2str(SAMPLINGLENGTH)];
for iii = 1 : length(DO_THESE_OF_SQS)
      s = [s '_' num2str(DO_THESE_OF_SQS(iii))];
end
disp(scs_);
% save([s '.mat'],
'sc_','sci_','c_','fn_pdf_','fn_pdf_c_','fn_best_sopts_','scs_',
'fn_','fntm_','SAMPLINGLENGTH');
save([s '.mat'], 'sc_','sci_','c_','fn_best_sopts_','scs_',
'fn_','fntm_','SAMPLINGLENGTH');
% clear sc sci fn_pdf c fn_best_sopts fn_pdf_c fn_best_set_sopt
CONTINUE_COMPUTATION sqs isqs iii id x xi DO_THESE_OF_SQS tag fntmlocal
name r f k l is ss s
end

f_sample_sq_by_sopts_to_tm__no_PDF_Unaf.m

% The rev. complementary strand has not been kept with the forward
strand. It was last saved on Jul 12th 2024.

function [sc_ sci_ c fn_best_sopts scs] =
f_sample_sq_by_sopts_to_tm__no_PDF_Unaf(tmfn, sqfn, SAMPLINGLENGTH, STEP,
RNASUBOPT, SCORINGSCHEME, VERBOSE, CT2B_DIR, DO_DBS, dist_lett,
MAXSTRSPERSQ, WINDOWSIZE, PROCENTFE)

% sc_ - scores for sets of sopts. The sets are evaluated acc. to
SCORINGSCHEME. It is one evaluated score for one set of sopts. One
sampling sq produces one set of sopts.
% sci_ - indexes of sorted sc_.
% c - scores of best sopts in sets of sopts.
% fn_best_sopts - .db (dot bracket) files for best sopts of all sampling
sqs of this input sq.

% WORKS ON A SINGLE SQ ONLY!!!!!! run_f_sample_sq_by_sopts_to_tm.m runs
this function for more sqs.

% STRAND=0: orig. sq., STRAND=1: rev. compl. of the org. sq., STRAND = 2:
both.
% SCORINGSCHEME=0: mean(sc), SCORINGSCHEME=1: min(sc), SCORINGSCHEME=2:
mean(sc(1:3).

sc_ = []; sci_ = []; c = []; fn_best_sopts = {}; scs = {};

if ~exist('STEP'), STEP = 1; end
if ~exist('RNASUBOPT'), RNASUBOPT = 0; end
if ~exist('SCORINGSCHEME'), SCORINGSCHEME = 0; end
if ~exist('VERBOSE'), VERBOSE = 0; end
if ~exist('DO_DBS'), DO_DBS = 1; end
% if ~exist('MAXSTRSPERSQ'), MAXSTRSPERSQ= 60; end
% if ~exist('WINDOWSIZE'), WINDOWSIZE= 2; end
% if ~exist('PROCENTFE'), PROCENTFE= 2000; end

id = fopen(sqfn); t=textscan(id,'%s','delimiter',char(10)); fclose(id);
sq = t{1}{2};
sqfn = [sqfn '_l=' num2str(SAMPLINGLENGTH)];
```

```
if length(sq) >= SAMPLINGLENGTH & exist(tmfn)
      if length(sq) == SAMPLINGLENGTH
            steps = 1;
      else
            steps = 1 : STEP : length(sq) - SAMPLINGLENGTH;
      end
      fn_best_sopts = cell(1, length(steps)); sc_ = zeros(1,
length(steps)); sci_ = sc_; c = sc_;
      for isteps = 1 : length(steps)
            step = steps(isteps);
            if VERBOSE, disp(['Doing ' num2str(step) '/[' num2str(1 : STEP
: length(sq) - SAMPLINGLENGTH) '].']); end
%           id = fopen([sqfn '_' num2str(step) '.fa'], 'w'); fwrite(id,
['>' sqfn '_' num2str(step) char(10) sq(step : step + SAMPLINGLENGTH -
1)]); fclose(id);
            if RNASUBOPT
                  unix(['RNAsubopt < ' sqfn '_' num2str(step) '.fa > '
sqfn '_' num2str(step) '.fa.rs']);
                  f = f_reformat_RNAsubopt([sqfn '_' num2str(step)
'.fa.rs'],'all',1);
            else
%                 f = f_get_sopt_strs__with_ct2b_pl([sqfn '_'
num2str(step) '.fa'], MAXSTRSPERSQ, WINDOWSIZE, PROCENTFE, CT2B_DIR);
%                 f = f_get_sopt_strs__with_ct2b_pl([sqfn '_'
num2str(step) '.fa'], 60, 2, 2000, CT2B_DIR);
%                 f = f_get_sopt_strs__with_ct2b_pl([sqfn '_'
num2str(step) '.fa'], 100, -5, 20, CT2B_DIR);
            fn = [sqfn '_' num2str(step) '.fa.br'];
            eval(['!echo -n > ' fn])
            fndm = [fn '.dm']; fndm(find(fndm == ' ')) = '_';
            id = fopen(fndm, 'w'); fwrite(id, sq(step : step +
SAMPLINGLENGTH - 1)); fclose(id);
            if WINDOWSIZE < 0
                  WINDOWSIZE = round(length(sq(step : step +
SAMPLINGLENGTH - 1)) / abs(WINDOWSIZE));
            end
            eval(['!UNAFold.pl -P ' num2str(PROCENTFE) ' -W '
num2str(WINDOWSIZE) ' -X ' num2str(MAXSTRSPERSQ) ' ' fndm ' >> null
2>&1'])
%           eval(['!UNAFold.pl -P 20 -W ' num2str(round(length(sq(step :
step + SAMPLINGLENGTH - 1)) / 5)) ' -X 100 ' fndm ' >> null 2>&1'])
%           eval(['!UNAFold.pl -P 20 -W ' num2str(round(length(sq(step :
step + SAMPLINGLENGTH - 1)) / 10)) ' -X 60 ' fndm ' >> null 2>&1'])

            [x x] = unix(['ls ' fndm '_*.ct']);
            if isempty(strfind(x, 'ls: cannot access'))
                  x(length(x)) = [];
                  x(find(x == ' ')) = char(10);
                  x=strsplit(x, char(10));
                  for ix =1:length(x)
                        eval(['!echo ''>' num2str(ix) '_' sqfn '_'
num2str(step) ''' >> ' fn])
                        eval(['!' CT2B_DIR 'ct2b.pl ' fndm '_' num2str(ix)
'.ct >> ' fn])
                  end
                  eval(['!rm ' fndm '*.ct'])
                  eval(['!rm ' fndm])
```

```
                  % eval(['!rm ' fndm '.log'])
                  eval(['!rm ' fndm '.ann'])
                  eval(['!rm ' fndm '.det'])
                  eval(['!rm ' fndm '.dG'])
                  eval(['!rm ' fndm '.h-num'])
                  eval(['!rm ' fndm '.rnaml'])
                  eval(['!rm ' fndm '.run'])
                  eval(['!rm ' fndm '.ss-count'])
                  eval(['!rm ' fndm '.plot'])
                  % eval(['!rm ' fndm '*.pdf'])
                  % eval(['!rm ' fndm '*.ps'])
            end
            end
            [x x] = unix(['wc -l < ' fn]);
            if ~str2num(x)
                  sc_ = ['Str. prediction impossible for this subsequence
(most likely for nt composition): ' fn ': ' sq(step : step +
SAMPLINGLENGTH - 1) '. Stop.'];
%                 disp(sc_)
                  return
            else
                  unix(['cat ' tmfn ' > ' sqfn '_' num2str(step) '.br']);
                  unix(['cat ' fn ' >> ' sqfn '_' num2str(step) '.br']);
                  unix(['RNAdistance -Xf -D' dist_lett ' < ' sqfn '_'
num2str(step) '.br > ' sqfn '_' num2str(step) '.br.d']);
                  sc = f_scan_RNAdistance_Xf([sqfn '_' num2str(step)
'.br.d'], dist_lett);
                  scs = [scs; sc];
                  [sci sci] = sort(sc, 'ascend');
      % for each set of sopts, i.e. sopts for each sampled subsq, store
score and index of the sopt best sim to template
                  c(isteps) = sc(sci(1));
      % scoring schemes
                  if SCORINGSCHEME == 0
                        sc_(isteps) = mean(sc);
                  else
                        if SCORINGSCHEME == 1
                              sc_(isteps) = sc(sci(1));
                        else
                              if SCORINGSCHEME == 2
                                    if length(sc) > 2, sc_(isteps) =
mean(sc(sci(1:3))); else, sc_(isteps) = mean(sc); end
                              end
                        end
                  end
                  if DO_DBS
                        fn_best_sopts{isteps} = [sqfn '_'
num2str(num2str(step)) 'nt_' num2str(sci(1)) 'th.db'];
                        eval(['!sed -n ''' num2str((sci(1)-1)*3+1) ','
num2str((sci(1)-1)*3+3) 'p'' < ' fn ' | cat > ' fn_best_sopts{isteps}])
                  end
            end
      end
% Evaluated scores of the sets of sopts of the sampling sqs. Whatever is
the SCORINGSCHEME, you take the least value of them acc. to
SCORINGSCHEME.
      [sci_ sci_] = sort(sc_);
```

```
      unix(['rm ' sqfn '_*.br ' sqfn '_*.br.d ']); if RNASUBOPT,
unix(['rm ' sqfn '_*.rs ' sqfn '_*.rs_r']);  end
else
      sc_ = ['length(sq) (' num2str(length(sq)) ') <= SAMPLINGLENGTH ('
num2str(SAMPLINGLENGTH) '), or tmfn not exist. Stop.'];
%     disp(sc_)
end
end
```

f_reformat_RNAsubopt.m

```
function fn = f_reformat_RNAsubopt(fn, MAXSOPTNOSIMGLESQ, rename)
% function fn = f_reformat_RNAsubopt(fn, MAXSOPTNOSIMGLESQ, rename)

id = fopen(fn); t=textscan(id, '%s', 'delimiter', char(10)); fclose(id);
if rename, fn = [fn '_r']; end
ido = fopen(fn, 'w');

i = [strmatch('>', t{1}); length(t{1}) + 1];
for ii = 1 : length(i) - 1
      if ischar(MAXSOPTNOSIMGLESQ) | i(ii + 1) - i(ii) - 2 <
MAXSOPTNOSIMGLESQ
            soptno = i(ii + 1) - i(ii) - 2;
      else
            soptno = MAXSOPTNOSIMGLESQ;
      end

      header = t{1}{i(ii)}(2 : length(t{1}{i(ii)}));
      if (ischar(MAXSOPTNOSIMGLESQ) & (~isempty(strfind(header,
MAXSOPTNOSIMGLESQ)) | strcmp(MAXSOPTNOSIMGLESQ, 'all'))) |
~ischar(MAXSOPTNOSIMGLESQ)
            if header(1) == ' ', header(1) = []; end
            s = cut_string(header, ' '); header = s{1};

            sq = cut_string(t{1}{i(ii) + 1} , ' '); sq = sq{1};

            for is = i(ii) + 2 : i(ii) + soptno + 1
                  br = cut_string(t{1}{is}, ' ');
                  fwrite(ido, ['>' num2str(is - i(ii) - 1) '_' header
char(10) sq char(10) br{1} char(10)]);
            end
      end
end
fclose(ido);
```

f_scan_RNAdistance_Xf.m

```
function [scs labels]= f_scan_RNAdistance_Xf(fn, dist_let)
% scs = f_scan_RNAdistance_Xf(fn, dist_let)
id = fopen(fn); t = textscan(id, '%s', 'delimiter', char(10));
fclose(id);
scs = ones(1, (length(t{1}) - 1) / 2) .* -1;
labels = {};
if isempty(strmatch([dist_let ': '], t{1}{3})),
disp(['f_scan_RNAdistance_Xf: ' fn ' is not a RNAdistance -Xf file.']),
return, end
```

```
for it = 3 : 2 : length(t{1})
      scs(floor(it / 2)) = str2num(t{1}{it}(3 : length(t{1}{it})));
      labels = [labels t{1}{it - 1}(2 : length(t{1}{it - 1}))];
end
end
```

merge_mats_app.m

```
% The rev. complementary strand has not been kept with the forward
strand. It was last saved on Jul 12th 2024.

i = strfind(fnload_, '.mat');
[x x] = unix(['ls ' fnload_(1 : i - 1) '_*.mat']);
x(find(double(x)==9)) = char(10);
x = strsplit(x, char(10));
if isempty(x{length(x)}), x(length(x)) = []; end

if isempty(strfind(x{1}, 'No such file or directory')) &
exist(x{1},'file')
      load(x{1});
else
      disp([fnload_(1 : i - 1) '*.mat not exist. Stop.'])
      FLAG__ = 0;
      return
end
sc = cell(1, length(sc_)); sci = sc; c = sc; fn_best_sopts = sc; scs =
sc; %fn_pdf = sc;fn_pdf_c = sc;

for ix = 1 : length(x)
      load(x{ix});
      for iI = 1 : length(sc_)
            if ~isempty(sc_{iI})
                  if isnumeric(sc_{iI})
                        sc(iI) = sc_(iI);
                        sci(iI) = sci_(iI);
%                       fn_pdf(iI) = fn_pdf_(iI);
                        c(iI) = c_(iI);
%                       fn_pdf_c(iI) = fn_pdf_c_(iI);
                        fn_best_sopts(iI) = fn_best_sopts_(iI);
                        scs(iI) = scs_(iI);
                  else
                        disp([sc_{iI} ' See ' x{ix} ' *.inf. Stop.'])
                        FLAG__ = 0;
                        return
                  end
            end
      end
end
sc_ = sc; sci_ = sci;c_ = c; fn_best_sopts_ = fn_best_sopts; scs_ = scs;
% fn_pdf_ = fn_pdf; fn_pdf_c_ = fn_pdf_c;
% save(fnload_,'sc_', 'sci_', 'fn_pdf_',
'c_','fn_pdf_c_','fn_best_sopts_', 'scs_');
save(fnload_,'sc_', 'sci_', 'c_','fn_best_sopts_', 'scs_');
unix(['rm ' fnload_(1 : i - 1) '_*.mat']);
if DO_TERMINATORS
      if ~exist('ttfn__')
            ttfn__='';
```

```
      end
      if ~isempty(ttfn__)
            [tt5_ tt3_ ttor_] = f_read_tts_from_standard_file(ttfn__);
            if isempty(tt5_)
                  disp('Invalid standard file with terminators. No
terminators.')
            end
      else
            disp('No -tt argument. Trying RhoTermPredict.py.')
            [tt5_ tt3_ ttor_] = f_get_RhoTermPredict_py(fn_, CT2B_DIR);
            if isempty(tt5_)
                  disp('RhoTermPredict fault. No terminators.')
            else
                  !rm info_about_predictions_*.txt
predictions_coordinates_*.csv
            end
      end
      if ~isempty(tt5_)
            save(fnload_, 'tt5_','tt3_','ttor_','ttfn__', '-append');
      end
end
clear x NO_OF_MATS iI sc sci fn_best_sopts s is ix scs
% clear fn_pdf c fn_pdf_c
```

f_get_RhoTermPredict_py.m

```
function [tt5 tt3 ttor] = f_get_RhoTermPredict(fn, CT2B_DIR)
tt3 = []; tt5 = []; ttor = [];
[x x] = unix(['python3 ' CT2B_DIR '/RhoTermPredict_algorithm_edited.py '
fn]);
if ~isempty(strfind(x, ['RhoTermPredict is working, please wait...'
char(10) 'Work finished, see output files in the current directory']))
      f=fastaread(fn);
      tt5 = cell(1, length(f)); tt3 = tt5; ttor = tt5;
      for fi=1:length(f)
            s=cut_string(f(fi).Header, ' ');
            a = importdata(['predictions_coordinates_' s{1} '.csv']);
            a(f_find_str_in_cells(a, 'Region')) = [];
            tt5{fi} = zeros(1, length(a)); tt3{fi} = tt5{fi}; ttor{fi} =
char(ones(1, length(a)) .* '-');
            for i = 1 : length(a)
                  aa = strsplit(a{i}, char(9));
                  tt5{fi}(i) = str2num(aa{2});
                  tt3{fi}(i) = str2num(aa{3});
                  if strcmp(aa{4}, 'plus')
                        ttor{fi}(i) = '+';
                  end
            end
      end
% else
%     disp('RhoTermPredict_algorithm_edited.py does not work all right.
(From unknown reason(s). No communication from authors.)')
end
```

f_read_tts_from_standard_file.m

```
function [tt5 tt3 ttor] = f_read_tts_from_standard_file(fn)
```

```
id = fopen(fn); t = textscan(id, '%u%u%u%s', 'delimiter', char(9));
fclose(id);
if size(t, 2) ~= 4, tt5 = []; tt3 = [] ;ttor = []; return, end
tt5 = cell(1, t{1}(length(t{1}))); tt3 = tt5; ttor = tt5;
for it = 1 : length(t{1})
      tt5{t{1}(it)} = [tt5{t{1}(it)} t{2}(it)];
      tt3{t{1}(it)} = [tt3{t{1}(it)} t{3}(it)];
      ttor{t{1}(it)} = [ttor{t{1}(it)} t{4}{it}];
end
```

f_read_ARNold_into_standard_file.m

```
function fn = f_read_ARNold_into_standard_file(fn)
if ~exist(fn), return, end
id = fopen(fn); t = textscan(id, '%s', 'delimiter', char(10));
fclose(id);
it = 1;
tti = 0;
id = fopen([fn '.txt'], 'w');
while it <= length(t{1})
      if ~isempty(t{1}{it})
            if t{1}{it}(1) == '>'
                  tti = tti + 1;
            else
                  tt = strsplit(t{1}{it}, ' ');
                  if length(tt) == 5
                        fwrite(id, [num2str(tti) char(9) tt{1} char(9)
num2str(str2num(tt{1}) + length(tt{4}) - 1) char(9) tt{3} char(10)]);
                  end
            end
      end
      it = it + 1;
end
fclose(id);
fn = [fn '.txt'];
```

present_results_table_no_stats.m

```
% load(fnload_)
if SHOW_ONLY_ORIENTATION_MATCHING_TEMPLATE
      isqsor = isqsor__;
else
      isqsor = 1 : length(fn_best_sopts_);
end
disp([char(10) 'RNASubOptLoc results for:' char(10) 'Input sequences: '
fnsqs__ char(10) 'Template: ' fntm__ char(10)])
s = ['Main prediction based on average TED of sets of suboptimal
structures produced by individual candidate sequences:' char(10) 'Input
sequence' char(9) 'Predicted candidate gene position*' char(9) 'Score**'
char(9)];
if TEST | BOOTSTRAP
      if h_or_p_
            s = [s 'p****' char(9)];
      else
            s = [s 't****' char(9)];
      end
end
```

```
s = [s 'Candidate representative structure plot' char(10)];
fprintf(s);
id = fopen([fnload_ '.txt'], 'w');
fwrite(id, s);
for isqsfig = 1 : length(isqsor)
      isqs = isqsor(isqsfig);
      s = strsplit(fn_best_sopts_{isqs}{sci_{isqs}(1)}, '.');
      s = strsplit(s{length(s) - 1}, '_');
      pos = s{4}(~isletter(s{4}));
      ss = strsplit(s{3}, '=');
      len = ss{2};
      todisp = [num2str(isqsor(isqsfig)) char(9) char(9) ...
                    pos ' : ' char(9) num2str(str2num(pos) + str2num(len)
- 1) char(9) char(9) char(9) char(9) ...
                    num2str(round(sc_{isqs}(sci_{isqs}(1))*10)/10)
char(9)];
      towrite = [num2str(isqsor(isqsfig)) char(9) ...
                     pos ' : ' num2str(str2num(pos) + str2num(len) - 1)
char(9) ...
                     num2str(round(sc_{isqs}(sci_{isqs}(1))*10)/10)
char(9)];
      if TEST | BOOTSTRAP
            if h_or_p_
                  todisp = [todisp num2str(round(psc_(isqs)*1000)/1000)
char(9)];
                  towrite = [towrite num2str(round(psc_(isqs)*1000)/1000)
char(9)];
            else
                  todisp = [todisp num2str(round(hsc_(isqs)*1000)/1000)
char(9)];
                  towrite = [towrite num2str(round(hsc_(isqs)*1000)/1000)
char(9)];
            end
      end
      disp([todisp fn_best_sopts_{isqs}{sci_{isqs}(1)}]);
      fwrite(id, [towrite fn_best_sopts_{isqs}{sci_{isqs}(1)} char(10)]);
end
s = [char(10) '* In input sequences.' char(10) '** Average tree edit
distance (TED) to template of the set of suboptimal structures produced
by the candidate gene sequence (the lesser the better).' char(10)];
if DO_ALTERNATIVE_CANDIDATES
      ss = [char(10) 'Alternative prediction based on best, single
suboptimal structures for candidates:' char(10) 'Input sequence' char(9)
'Predicted candidate gene position*' char(9) 'Score***' char(9)];
      if TEST     | BOOTSTRAP
            if h_or_p_
                  ss = [ss 'p****' char(9)];
            else
                  ss = [ss 't****' char(9)];
            end
      end
      ss = [ss 'Candidate representative structure plot' char(10)];
      fprintf(ss);
      fwrite(id, ss);
      for isqsfig = 1 : length(isqsor)
            isqs = isqsor(isqsfig);
            [i i] = min(c_{isqs});
            ss = strsplit(fn_best_sopts_{isqs}{i}, '.');
```

```
            ss = strsplit(ss{length(ss) - 1}, '_');
            pos = str2num(ss{4}(~isletter(ss{4})));
            sss = strsplit(ss{3}, '=');
            len = str2num(sss{2});

            todisp = [num2str(isqsor(isqsfig)) char(9) char(9) ...
                          num2str(pos) ' : ' char(9) num2str(pos + len -
1) char(9) char(9) char(9) char(9) ...
                          num2str(round(c_{isqs}(i)*10)/10) char(9)
char(9)];
            towrite = [num2str(isqsor(isqsfig)) char(9) ...
                           num2str(pos) ' : ' num2str(pos + len - 1)
char(9) ...
                           num2str(round(c_{isqs}(i)*10)/10) char(9) ];
            if TEST | BOOTSTRAP
                  if h_or_p_
                        todisp = [todisp
num2str(round(pc_(isqs)*1000)/1000) char(9)];
                        towrite = [towrite
num2str(round(pc_(isqs)*1000)/1000) char(9)];
                  else
                        todisp = [todisp
num2str(round(hc_(isqs)*1000)/1000) char(9)];
                        towrite = [towrite
num2str(round(hc_(isqs)*1000)/1000) char(9)];
                  end
            end
            disp([todisp fn_best_sopts_{isqs}{i}]);
            fwrite(id, [towrite fn_best_sopts_{isqs}{i} char(10)]);
      end
      s = [s '*** Best, i.e. the least TED of TEDs of all suboptimal
structures of candidate gene sequence.' char(10)];
end
if TEST | BOOTSTRAP
      if h_or_p_
            s = [s '**** p-value of two-sample t-test of best candidates
against either other worse candidates or candidates of randomized
sequences' newline];
      else
            s = [s '**** The decision t of two-sample t-test for best
candidates against either non-best candidates or candidates of randomized
sequences (0-unsignificant, 1-significant at Alpha=' num2str(ALPHA__) ')'
newline newline newline];
      end
end
fprintf(s);
fwrite(id, s);
fclose(id);
clear isqs s ss x i row pos len id sss pc psc todisp towrite isqsfig
isqsor

histo.m

binsno = 10;
if GRAPHICAL_OUTPUT, figure; hold on; end
if BOOTSTRAP
      if GRAPHICAL_OUTPUT, hist(reshape(sc_rand_set_, [size(sc_rand_set_,
2) * length(fn_best_sopts_) 1]), binsno); end
```

```
      [counts centers] = hist(reshape(sc_rand_set_, [size(sc_rand_set_, 
2) * length(fn_best_sopts_) 1]), binsno);
else
      x = [];
      for isqs = 1 : length(fn_best_sopts_)
            x = [x sc_{isqs}];
      end
      [counts centers] = hist(x, binsno);
      if GRAPHICAL_OUTPUT, hist(x, binsno); end
end
factor = max(counts) / 25;
best = [];
for i=1:length(fn_best_sopts_)
      best = [best sc_{i}(sci_{i}(1))];
end
[best i] = sort(best);
% [counts centers] = hist(best, (ceil(max(best)) - floor(min(best))) + 
1);
[counts centers] = hist(best, binsno);
ibest = 1;
for icounts = 1 : length(counts)
      y = 1;
      for j = ibest : ibest + counts(icounts) - 1
%           text(centers(icounts), y, 
num2str(i(j)),'color','r','horizontalalignment','center');
            if GRAPHICAL_OUTPUT, text(best(ibest), y * factor, 
num2str(i(j)),'color','r','horizontalalignment','center'); end
            y = y + 1;
      end
%     line([best(ibest) best(ibest)],[0 counts(icounts)], 'color','r');
      ibest = ibest + counts(icounts);
end

if GRAPHICAL_OUTPUT
      box, grid
      a = axis;
      axis([floor(min(best)) a(2:4)]);
      if BOOTSTRAP
            title([fnsqs__ ',' newline fntm__ ', SAMPLING LENGTH=' 
num2str(SAMPLINGLENGTH) ', BOOTSTRAPS=' num2str(size(sc_rand_set_, 2))], 
'interpreter','none','FontSize', 8);
            ylabel(['Counts of predicted candidates'], 'FontSize', 8)
            xlabel(['sc (TED-based set score) (the lesser, the better). 
Legend: blue - histogram of scs for' newline 'random sequences, red - scs 
for best candidates marked with query #' newline '(position of the 
numbers on y axis is irrelevant).'],'FontSize', 8);
      end
      if TEST
            title([fnsqs__ ',' newline fntm__ ', SAMPLING LENGTH=' 
num2str(SAMPLINGLENGTH)], 'interpreter','none','FontSize', 8);
            xlabel(['sc (TED-based set score) (the lesser, the better). 
Legend: blue - histogram of scs' newline 'for non-best candidates, red - 
scs for best candidates marked with query #' newline '(position of the 
numbers on y axis is irrelevant).'],'FontSize', 8);
            ylabel('Counts of predicted candidates', 'FontSize', 8)
      end
end
```

```
for i = 1 : size(sc_rand_set_, 1)
      disp(['Best sc = ' num2str(round(sc_{i}(sci_{i}(1))*10)/10) ',
sc_rand_set_ (mean/median) = ' num2str(round(mean(sc_rand_set_(i,
:))*10)/10)...
              '/' num2str(round(median(sc_rand_set_(i, :))*10)/10) ', sc
distances (from mean/median) = ' ...
            num2str(round((mean(sc_rand_set_(i, :)) -
sc_{i}(sci_{i}(1)))*10)/10) '/' num2str(round((median(sc_rand_set_(i, :))
- sc_{i}(sci_{i}(1)))*10)/10)]);
% disp(['sc_rand_set_ = ' num2str(round(sc_rand_set_*10)/10) '.']);
end

if DO_ALTERNATIVE_CANDIDATES
      if GRAPHICAL_OUTPUT, figure; hold on;box, grid, end
      if BOOTSTRAP
            if GRAPHICAL_OUTPUT, hist(reshape(c_rand_sopt_,
[size(c_rand_sopt_, 2) * length(fn_best_sopts_) 1]), binsno); end
            [counts centers] = hist(reshape(c_rand_sopt_,
[size(c_rand_sopt_, 2) * length(fn_best_sopts_) 1]), binsno);
      else
            x = [];
            for isqs = 1 : length(fn_best_sopts_)
                  x = [x c_{isqs}];
            end
            [counts centers] = hist(x, binsno);
            if GRAPHICAL_OUTPUT, hist(x, binsno); end
      end
      factor = max(counts) / 25;
      best = [];
      for i=1:length(fn_best_sopts_)
            best = [best min(c_{i})];
      end
      [best i] = sort(best);
      % [counts centers] = hist(best, (ceil(max(best)) -
floor(min(best))) + 1);
      [counts centers] = hist(best, binsno);
      ibest = 1;
      for icounts = 1 : length(counts)
            y = 1;
            for j = ibest : ibest + counts(icounts) - 1
                  if GRAPHICAL_OUTPUT,
            %           text(centers(icounts), y,
num2str(i(j)),'color','r','horizontalalignment','center');
                        text(best(ibest), y * factor,
num2str(i(j)),'color','r','horizontalalignment','center');
                  end
                  y = y + 1;
            end
      %     line([best(ibest) best(ibest)],[0 counts(icounts)],
'color','r');
            ibest = ibest + counts(icounts);
      end
      if GRAPHICAL_OUTPUT,
            a = axis;
            axis([floor(min(best)) a(2:4)]);
            if BOOTSTRAP
```

```
                 title([fnsqs__ ',' newline fntm__ ', SAMPLING LENGTH='
num2str(SAMPLINGLENGTH) ', BOOTSTRAPS=' num2str(size(sc_rand_set_, 2))],
'interpreter','none','FontSize', 8);
                 ylabel(['Counts of predicted candidates'], 'FontSize',
8)
                 xlabel(['TED (the lesser, the better). Legend: blue -
histogram of TEDs for' newline 'random sequences, red - TEDs for best
candidates marked with query # overlapped' newline 'with the histogram
(position of the query numbers on y axis is irrelevant).'],'FontSize',
8);
           end
           if TEST
                 title([fnsqs__ ',' newline fntm__ ', SAMPLING LENGTH='
num2str(SAMPLINGLENGTH)], 'interpreter','none','FontSize', 8);
                 xlabel(['TED (the lesser, the better). Legend: blue -
histogram of TEDs' newline 'for non-best candidates, red - TEDs for best
candidates marked with query #  overlapped' newline 'with the histogram
(position of the query numbers on y axis is irrelevant).'],'FontSize',
8);
                 ylabel('Counts of predicted candidates', 'FontSize', 8)
           end
      end

      for i=1:length(fn_best_sopts_)
           disp(['Best c = ' num2str(round(min(c_{i})*10)/10) ',
c_rand_sopt_ (mean/median) = ' num2str(round(mean(c_rand_sopt_(i,
:))*10)/10)...
                  '/' num2str(round(median(c_rand_sopt_(i, :))*10)/10)
', c distances (from mean/median) = ' ...
                 num2str(round((mean(c_rand_sopt_(i, :)) -
min(c_{i}))*10)/10) '/' num2str(round((median(c_rand_sopt_(i, :)) -
min(c_{i}))*10)/10)]);
%      disp(['c_rand_sopt_ = ' num2str(round(c_rand_sopt_*10)/10) '.']);
      end
end
clear binsno x isqs best i counts centers ibest icounts y a factor

btstrp.m

if ~exist('DO_NEIGHBOURS', 'var'), DO_NEIGHBOURS = 0;end
if BOOTSTRAP > 9
      sc_rand_set_ = zeros(length(fn_best_sopts_), BOOTSTRAP);
                 % sc_rand_sopt_ = sc_rand_set_;
      c_rand_sopt_ = zeros(length(fn_best_sopts_), BOOTSTRAP);
           % c_rand_set_ = sc_rand_set_;
      if DO_NEIGHBOURS
           scs_rand_set_ = cell(length(fn_best_sopts_), 3);
      else
           scs_rand_set_ = cell(length(fn_best_sopts_), 1);
      end
      scs_rand_sopt_ = scs_rand_set_;

      for isqs = 1 : length(fn_best_sopts_)
           disp(['Bootstrapping ' num2str(isqs) '/'
num2str(length(fn_best_sopts_)) ' query sequences.'])
           % here the structure files are used only because their sqs
show position for bootstrapping on input sqs
```

```
                %
SAMPLINGLENGTH, STEP,RNASUBOPT, STRAND, SCORINGSCHEME, RANDSAMPLES
                disp('Doing best sopt of best set.')
                [scrn crn scsrn] = f_bt_strp(fntm__,
fn_best_sopts_{isqs}{sci_{isqs}(1)}, SAMPLINGLENGTH, STEP__, 0, 0,
BOOTSTRAP, CT2B_DIR, pid__, DIST_LETT, MAXSTRSPERSQ, WINDOWSIZE,
PROCENTFE);
                if isempty(scrn)
                     disp(crn)
                     return
                else
                     sc_rand_set_(isqs, :) = scrn;
           %    c_rand_set_(isqs, :) = crn;
                     scs_rand_set_{isqs, 1} = scsrn;
                end

                if DO_NEIGHBOURS
                     disp('Doing best sopt of best set left-
neighbour.')
                     if sci_{isqs}(1) - 1
                          [scrn crn scsrn] = f_bt_strp(fntm__,
fn_best_sopts_{isqs}{sci_{isqs}(1) - 1}, SAMPLINGLENGTH, STEP__,0,
0,BOOTSTRAP, CT2B_DIR, pid__, DIST_LETT, MAXSTRSPERSQ, WINDOWSIZE,
PROCENTFE);
                     if isempty(scrn)
                          disp(crn)
                          return
                     else
                          scs_rand_set_{isqs, 2} = scsrn;
                     end
                     else
                          scs_rand_set_{isqs, 2} = [];
                     end

                     disp('Doing best sopt of best set right-neighbour,
if exists.')
                     if sci_{isqs}(1) + 1 <= length(sci_{isqs})
                          [scrn crn scsrn] = f_bt_strp(fntm__,
fn_best_sopts_{isqs}{sci_{isqs}(1) + 1}, SAMPLINGLENGTH, STEP__,0,  0,
BOOTSTRAP, CT2B_DIR, pid__, DIST_LETT, MAXSTRSPERSQ, WINDOWSIZE,
PROCENTFE);
                          if isempty(scrn)
                               disp(crn)
                               return
                          else
                               scs_rand_set_{isqs, 3} = scsrn;
                          end
                     else
                          scs_rand_set_{isqs, 3} = [];
                     end
                end

                disp('Doing individual best sopt.')
                [i i] = min(c_{isqs});
                [scrn crn scsrn] = f_bt_strp(fntm__,
fn_best_sopts_{isqs}{i}, SAMPLINGLENGTH, STEP__, 0,  0, BOOTSTRAP,
CT2B_DIR, pid__, DIST_LETT, MAXSTRSPERSQ, WINDOWSIZE, PROCENTFE);
                if isempty(scrn)
```

```
                        disp(crn), return
                  else
            %     sc_rand_sopt_(isqs, :) = scrn;
                        c_rand_sopt_(isqs, :) = crn;
                        scs_rand_sopt_{1, isqs} = scsrn;
                  end

                  if DO_NEIGHBOURS
                        disp('Doing individual best sopt left-neighbour,
if exists.')
                        if i - 1
                              [scrn crn scsrn] = f_bt_strp(fntm__,
fn_best_sopts_{isqs}{i - 1}, SAMPLINGLENGTH, STEP__, 0, 0, BOOTSTRAP,
CT2B_DIR, pid__, DIST_LETT, MAXSTRSPERSQ, WINDOWSIZE, PROCENTFE);
                              if isempty(scrn)
                                    disp(crn), return
                              else
                                    scs_rand_sopt_{isqs, 2} = scsrn;
                              end
                        else
                              scs_rand_sopt_{isqs, 2} = [];
                        end

                        disp('Doing individual best sopt right-neighbour,
if exists.')
                        if i + 1 <= length(c_{isqs})
                              [scrn crn scsrn] = f_bt_strp(fntm__,
fn_best_sopts_{isqs}{i + 1}, SAMPLINGLENGTH, STEP__,0, 0, BOOTSTRAP,
CT2B_DIR, pid__, DIST_LETT, MAXSTRSPERSQ, WINDOWSIZE, PROCENTFE);
                              if isempty(scrn)
                                    disp(crn), return
                              else
                                    scs_rand_sopt_{isqs, 3} = scsrn;
                              end
                        else
                              scs_rand_sopt_{isqs, 3} = [];
                        end
                  end
      end
      clear scrn crn isqs i
else
      disp('BOOTSTRAP < 10. No btstrp.m.')
end
```

f_get_conserved_region.m

```
function cnsd_rgn = f_get_conserved_region(tmfn, fn, BLASTNPATH)
% You do local BLAST of template against input sequences
% find out if fn_ is fasta
if isempty(fn), cnsd_rgn = [fn ' is empty.']; return, end
[x x] = unix(['head -n1 ' fn]);
if strfind(x, 'cannot open') | strfind(x, 'No such file or directory') |
x(1) ~= '>'
      cnsd_rgn = [fn ' not exist or not fasta.'];
      return
end
if exist('BLASTNPATH') & strcmp(BLASTNPATH, 'No BLAST.')
```

```
      cnsd_rgn = 'blastn not installed/configured. No conserved regions
computed.'; return
end
if ~exist('BLASTNPATH')
      [v v] = unix('which blastn');
      [x x] = unix('find ~/ -name "blastn"');
      if ~contains(x, 'bin/blastn') | contains(v, 'no blastn in')
            cnsd_rgn = 'blastn not installed/configured. Install it or put
on PATH. No conserved regions computed.'; return
      else
            if ~contains(v, 'no blastn in')
                  BLASTNPATH = '';
            else
                  x=strsplit(x,newline);
                  x = x{find(contains(x, 'bin/blastn'))};
                  i = find(x == '/');
                  BLASTNPATH = x(1 : i(length(i)));
            end
      end
end

tag = strsplit(num2str(rand), '.');
bl = ['BLASTn_' tag{2}];

unix([BLASTNPATH 'makeblastdb -in ' fn ' -dbtype nucl -out ' bl '.db']);%
>> /dev/null 2>&1']);
unix([BLASTNPATH 'blastn -query ' tmfn ' -db ' bl '.db -out ' bl '.bo.txt
-evalue 0.05 -word_size 7 -gapopen 1 -gapextend 2 -penalty -1 -reward 1 -
max_target_seqs 100 -outfmt 6']);%  >> /dev/null 2>&1']);
id=fopen([bl
'.bo.txt']);t=textscan(id,'%s%s%f%d%u%u%u%u%u%u%f%f','delimiter',char(9))
; fclose(id);
cnsd_rgn = cell(length(t{1}), 4);
idel = [];
for it = 1 : length(t{1})
% % you have to use only +/+ BLAST hits, as the template str should be
where its sq is.
%         if t{9}(it) <= t{10}(it)
%             cnsd_rgn(it, 1) = t{2}(it);
%             cnsd_rgn{it, 2} = t{9}(it);
%             cnsd_rgn{it, 3} = t{10}(it);
%         else
%             cnsd_rgn(it, 1) = t{2}(it);
%         end
% here you use both directions
            cnsd_rgn(it, 1) = t{2}(it);
      if t{9}(it) <= t{10}(it)
            cnsd_rgn{it, 2} = t{9}(it);
            cnsd_rgn{it, 3} = t{10}(it);
            cnsd_rgn{it, 4} = '+';
      else
            cnsd_rgn{it, 3} = t{9}(it);
            cnsd_rgn{it, 2} = t{10}(it);
            cnsd_rgn{it, 4} = '-';
      end
end
unix(['rm ' bl '.db.n* >> /dev/null 2>&1']);
```

draw_run_f_sample_sq_by_sopts_to_tm_simple_csd_rgn.m

```
warning('off')
% parameters that can come from rsol.m
if ~exist('DO_3_END_U'), DO_3_END_U = 1; end
if ~exist('SAVE_FIGURES'), SAVE_FIGURES = 0; end
if ~exist('DO_OVERLAP_WITH_SEQUENCE_SIMILARITY_REGIONS'),
DO_OVERLAP_WITH_SEQUENCE_SIMILARITY_REGIONS = 0; end
if ~exist('SHOW_SEQUENCE_SIMILARITY_REGIONS'),
SHOW_SEQUENCE_SIMILARITY_REGIONS = 0; end
if ~exist('DO_ALTERNATIVE_CANDIDATES'), DO_ALTERNATIVE_CANDIDATES = 0;
end
if ~exist('SHOW_ONLY_ORIENTATION_MATCHING_TEMPLATE'),
SHOW_ONLY_ORIENTATION_MATCHING_TEMPLATE = 0; end
if ~exist('TOO_OLD_RNA_VIENNA'), TOO_OLD_RNA_VIENNA = 0; end
% inner parameters
if ~exist('NUMBER_OF_BEST_SETS_TO_SHOW'), NUMBER_OF_BEST_SETS_TO_SHOW =
1; end
if ~exist('TEST'), TEST = 0; end
if ~exist('BOOTSTRAP'), BOOTSTRAP = 0; end
if TEST & BOOTSTRAP, TEST = 0; end
if TEST
      textinfig = 't';
end
if BOOTSTRAP
      textinfig = 'b';
end

SUFFIX = {'th'; 'st'; 'nd'; 'rd';
'th';'th';'th';'th';'th';'th';'th';'th';}';
MAXNOOFSTRS = 6;

sqs = fastaread(fnsqs__);
% hse=figure('MenuBar', 'none','ToolBar', 'none', 'Name',
'','NumberTitle', 'off');
hse=figure;
hold on, grid, box, data.value = zeros(1, 3); data.strs = zeros(1, 1);
guidata(hse, data);
dcm_obj = datacursormode(hse);
set(dcm_obj, 'DisplayStyle', 'window', 'SnapToDataVertex', 'on');
set(dcm_obj, 'UpdateFcn', @customDatatipCallback);
set(gcf, 'Pointer', 'hand');
region = []; %zeros(length(sqs) * 2, 4);
ll = [];
if SHOW_ONLY_ORIENTATION_MATCHING_TEMPLATE
      isqsor = isqsor__;
else
      isqsor = 1 : length(sqs);
end
for isqsfig = 1 : length(isqsor)
      isqs = isqsor(isqsfig);
      if NUMBER_OF_BEST_SETS_TO_SHOW > length(fn_best_sopts_{isqs}),
NUMBER_OF_BEST_SETS_TO_SHOW = length(fn_best_sopts_{isqs}); end
      if NUMBER_OF_BEST_SETS_TO_SHOW > MAXNOOFSTRS,
NUMBER_OF_BEST_SETS_TO_SHOW = MAXNOOFSTRS; end
      if DO_ALTERNATIVE_CANDIDATES > length(fn_best_sopts_{isqs}),
DO_ALTERNATIVE_CANDIDATES = length(fn_best_sopts_{isqs}); end
```

```
      if DO_ALTERNATIVE_CANDIDATES > MAXNOOFSTRS,
DO_ALTERNATIVE_CANDIDATES = MAXNOOFSTRS; end
      if NUMBER_OF_BEST_SETS_TO_SHOW + DO_ALTERNATIVE_CANDIDATES >
MAXNOOFSTRS
            NUMBER_OF_BEST_SETS_TO_SHOW = 3;
            DO_ALTERNATIVE_CANDIDATES = 3;
      end
      ll = [ll length(sqs(isqs).Sequence)];

      n = strsplit(sqs(isqs).Header, ' ');
      k = find(contains(n, 'chromosome'));
      if ~isempty(k) & ~isempty(str2num(n{k + 1})) &
~isletter(str2num(n{k + 1}))
            n = strsplit(f_remove_substrs_from_str_rsol(sqs(isqs).Header,
{'MAG '; 'MAG: '; 'TPA_asm ' ; 'TPA_asm: '; 'candidatus '; 'strain ';'sp.
'; 'isolate '; ' bacterium '; 'UNVERIFIED_ORG: '; 'Sample ';}, 0, 0), '
');
            name = [n{2} ' ' n{3}];
            k = find(contains(n, 'chromosome'));
            name = [name ' chr. ' n{k + 1}];
      else
            n = strsplit(f_remove_substrs_from_str_rsol(sqs(isqs).Header,
{'MAG '; 'MAG: '; 'TPA_asm ' ; 'TPA_asm: '; 'chromosome '; 'candidatus ';
'strain ';'sp. '; 'isolate '; ' bacterium '; 'UNVERIFIED_ORG: '; 'Sample
';}, 0, 0), ' ');
            n1 = 2;
            if sum(isletter(n{1})) > sum(~isletter(n{1}))
                  n1 = 1;
            end
            name = n{n1};
            for in = n1 + 1 : n1 + 2
                  name = [name ' ' n{in}];
            end
      end
      if exist('cnsd_rgn_')
            cnsd_rgn_i = find(contains(cnsd_rgn_(:,1), n{1}));
            if ~isempty(cnsd_rgn_i)
%                 if length(cnsd_rgn_i) > 1, disp('More than 1 queries
with the same ID in cnsd_rgn_. Taking 1st of them.'), end
                  cnsd_rgn_i = cnsd_rgn_i(1);
                  name = [name ' ' cnsd_rgn_{cnsd_rgn_i, 4}];
            else
                  if SHOW_SEQUENCE_SIMILARITY_REGIONS
                        disp(['Note: no sequence similarity to template
detected by BLAST, thus no sequence similarity region for query #'
num2str(isqs) ', ' name '.'])
                  end
            end
      end
      text(0, length(isqsor) - isqsfig + 1, name, 'FontSize', 8,
'VerticalAlignment', 'bottom','interpreter','none');
% draw a line for input sq
      line([0 length(sqs(isqs).Sequence)],[length(isqsor) - isqsfig + 1
length(isqsor) - isqsfig + 1],'color', [173/255 216/255 230/255],
'LineWidth', 2 );
% draw a line for sequence similarity region, if available
      linestep = .1;
```

```
% draw lines for best sopt from best set of sopts followed with score
      look_for_overlap_flag = 1;
      c = .05;
      for j = 1 : NUMBER_OF_BEST_SETS_TO_SHOW
            if exist([WP__ fn_best_sopts_{isqs}{sci_{isqs}(j)}], 'file')
                  id = fopen([WP__ fn_best_sopts_{isqs}{sci_{isqs}(j)}]);
sq = textscan(id,'%s','delimiter',char(10)); fclose(id);
                  s = strsplit(fn_best_sopts_{isqs}{sci_{isqs}(j)}, '.');
                  s = strsplit(s{length(s) - 1}, '_');
                  pos = str2num(s{4}(~isletter(s{4})));
                  ss = strsplit(s{3}, '=');
                  len = str2num(ss{2});
                  if TOO_OLD_RNA_VIENNA
                        disp(['No regular PS figure for ' WP__
fn_best_sopts_{isqs}{sci_{isqs}(j)} '. Most likely too old version of RNA
Vienna package installed.'])
                  else
                        unix(['RNAplot -t1 < ' WP__
fn_best_sopts_{isqs}{sci_{isqs}(j)}]);
                        unix(['gs -sDEVICE=pngalpha -dEPSCrop -o ' WP__
sq{1}{1}(2 : length(sq{1}{1})) '_ss.png ' sq{1}{1}(2 : length(sq{1}{1}))
'_ss.ps >> /dev/null 2>&1']);
                        unix(['rm ' sq{1}{1}(2 : length(sq{1}{1}))
'_ss.ps']);
                  end
                  c = .05;
                  lineHandle = line([pos pos+len-1],[length(isqsor) -
isqsfig + 1 - linestep length(isqsor) - isqsfig + 1 - linestep],'color',
'r');
            %     line([pos pos+len-1],[length(isqsor) - isqsfig + 1 -
linestep length(isqsor) - isqsfig + 1 - linestep],'color',
'r','marker','>','MarkerSize',3);
                  if j > 1
                        s = num2str(j);
                        tt = [name ': ' s SUFFIX{str2num(s(length(s))) +
1} ' best set candidate' newline 'representative str. (TED='
num2str(c_{isqs}(sci_{isqs}(j))) ')'];
                  else
                        tt = [name ': best set candidate' newline
'representative str. (TED=' num2str(c_{isqs}(sci_{isqs}(j))) ')'];
                  end
      % test representative individual structures in the structure
windows. Do it...? It's the sets, what decides here.
      %           if DO_TEST
      %                 if BOOTSTRAP_SOPTS
      %                       [h pc] = ttest2(c_{isqs}(sci_{isqs}(j)),
c_rand_set_(isqs, :));
      %                 else
      %                       [h pc] = ttest2(c_{isqs}(sci_{isqs}(j)),
c_{isqs});
      %                 end
      %                 tt = [tt ', p='
num2str(num2str(round(pc*1000)/1000)) ')'];
      %           else
      %                 tt = [tt ')'];
      %           end
```

```
                if DO_OVERLAP_WITH_SEQUENCE_SIMILARITY_REGIONS &
exist('cnsd_rgn_') & ~isempty(cnsd_rgn_i) &
~isempty(cnsd_rgn_{cnsd_rgn_i, 2})
      % here, find out whether the best set overlaps sequence similarity
region because of the str. window title
      % here you have overlap that must be positive and longer than 25%
of the structure sequence length
      %                 left = pos;
      %                 if cnsd_rgn_{cnsd_rgn_i, 2} >= pos
      %                      left = cnsd_rgn_{cnsd_rgn_i, 2};
      %                 end
      %                 right = pos + len - 1;
      %                 if cnsd_rgn_{cnsd_rgn_i, 3} <= pos + len - 1
      %                      right = cnsd_rgn_{cnsd_rgn_i, 3};
      %                 end
      %                 if right - left > len / 100 * 25
      % here you have the sequence similarity region within the predicted
structure
                     if cnsd_rgn_{cnsd_rgn_i, 2} >= pos &
cnsd_rgn_{cnsd_rgn_i, 3} <= pos + len
                          look_for_overlap_flag = 0;
                          tt = [tt '. Also overlaps sequence similarity
region.'];
                     end
                end
                hPatch = patch('XData', [pos-c pos+len-1+c pos+len-1-c
pos+c], ...
                                                  'YData', [length(isqsor) -
isqsfig + 1 - linestep - c length(isqsor) - isqsfig + 1 - linestep - c
length(isqsor) - isqsfig + 1 - linestep + c length(isqsor) - isqsfig + 1
- linestep + c], ...
                                                  'FaceColor', 'none',
'EdgeColor', 'none', ...
                                                  'HitTest', 'on',
'PickableParts', 'all', ...
                                                  'ButtonDownFcn',@(src, evt)
lineClicked(src, evt, [WP__ sq{1}{1}(2 : length(sq{1}{1})) '_ss.png'],
tt, isqsfig, hse.Position([1, 2])));
                region = [region; [pos-c length(isqsor)-isqsfig+1-
linestep-c (pos+len-1+c)-(pos-c) 2*c]];
      % show data only for sets, as this is analysis based on the set
score
                if TEST | BOOTSTRAP
                     if h_or_p_
      %                      text(pos+len+5, length(isqsor) - isqsfig + 1
- linestep, ['sc=' num2str(round(sc_{isqs}(sci_{isqs}(j))*10)/10) ' (p='
num2str(round(psc_(isqs)*1000)/1000) '), TED='
num2str(round(c_{isqs}(sci_{isqs}(j))*10)/10) ], 'FontSize', 8 );
                          text(pos+len+5, length(isqsor) - isqsfig + 1
- linestep, ['sc=' num2str(round(sc_{isqs}(sci_{isqs}(j))*10)/10) ', p='
num2str(round(psc_(isqs)*1000)/1000) ], 'FontSize', 8 );
                     else
      %                      text(pos+len+5, length(isqsor) - isqsfig + 1
- linestep, ['sc=' num2str(round(sc_{isqs}(sci_{isqs}(j))*10)/10) ' ('
textinfig '=' num2str(hsc_(isqs)) '), TED='
num2str(round(c_{isqs}(sci_{isqs}(j))*10)/10) ], 'FontSize', 8);
```

```
                        text(pos+len+5, length(isqsor) - isqsfig + 1
- linestep, ['sc=' num2str(round(sc_{isqs}(sci_{isqs}(j))*10)/10) ', '
textinfig '=' num2str(hsc_(isqs)) ], 'FontSize', 8);
                  end
            else
      %           text(pos+len+5, length(isqsor) - isqsfig + 1 -
linestep, ['sc=' num2str(round(sc_{isqs}(sci_{isqs}(j))*10)/10) ', TED='
num2str(round(c_{isqs}(sci_{isqs}(j))*10)/10) ], 'FontSize', 8 );
                  text(pos+len+5, length(isqsor) - isqsfig + 1 -
linestep, ['sc=' num2str(round(sc_{isqs}(sci_{isqs}(j))*10)/10)],
'FontSize', 8 );
            end
      % draw dots for Us in 3' tail
            if DO_3_END_U
                  for i = len - 10 : len
                        if sq{1}{2}(i) == 'U' | sq{1}{2}(i) == 'T'
                              plot(pos + i - 1, length(isqsor) -
isqsfig + 1 - linestep - .05, 'k.');
                        end
                  end
            end
            linestep = linestep + .15;
         else
            disp([WP__ fn_best_sopts_{isqs}{sci_{isqs}(j)} ' not
exist. No picture.'])
         end
      end
% find a predicted str that overlap the sequence similarity region and
draw it
      if DO_OVERLAP_WITH_SEQUENCE_SIMILARITY_REGIONS &
look_for_overlap_flag & ~isempty(cnsd_rgn_i) & exist('cnsd_rgn_') &
~isempty(cnsd_rgn_{cnsd_rgn_i, 2})
         for j = NUMBER_OF_BEST_SETS_TO_SHOW + 1 : length(sci_{isqs})
            s = strsplit(fn_best_sopts_{isqs}{sci_{isqs}(j)}, '.');
            s = strsplit(s{length(s) - 1}, '_');
            pos = str2num(s{4}(~isletter(s{4})));
            ss = strsplit(s{3}, '=');
            len = str2num(ss{2});
% here you have the sequence similarity region within the predicted
structure
            if cnsd_rgn_{cnsd_rgn_i, 2} >= pos &
cnsd_rgn_{cnsd_rgn_i, 3} <= pos + len
% % here you have overlap that must be positive and longer tha 25% of the
structure sequence length
%           left = pos; if cnsd_rgn_{cnsd_rgn_i, 2} >= pos, left =
cnsd_rgn_{cnsd_rgn_i, 2}; end
%           right = pos + len - 1; if cnsd_rgn_{cnsd_rgn_i, 3} <=
pos + len - 1, right = cnsd_rgn_{cnsd_rgn_i, 3}; end
%           if right - left > len / 100 * 25
                  id = fopen([WP__
fn_best_sopts_{isqs}{sci_{isqs}(j)}]); sq =
textscan(id,'%s','delimiter',char(10)); fclose(id);
                  if TOO_OLD_RNA_VIENNA
                        disp(['No regular PS figure for ' WP__
fn_best_sopts_{isqs}{sci_{isqs}(j)} '. Most likely too old version of RNA
Vienna package installed.'])
                  else
```

```
                            unix(['RNAplot -t1 < ' WP__
fn_best_sopts_{isqs}{sci_{isqs}(j)}]);
                            unix(['gs -sDEVICE=pngalpha -dEPSCrop -o '
WP__ sq{1}{1}(2 : length(sq{1}{1})) '_ss.png ' sq{1}{1}(2 :
length(sq{1}{1})) '_ss.ps >> /dev/null 2>&1']);
                            unix(['rm ' sq{1}{1}(2 : length(sq{1}{1}))
'_ss.ps']);
                      end
                      lineHandle = line([pos pos+len-1],[length(isqsor)
- isqsfig + 1 - linestep length(isqsor) - isqsfig + 1 -
linestep],'color', 'r','LineStyle','--');
                      s = num2str(j);
%                     if DO_TEST
% % test for individual structures
%                           if BOOTSTRAP_SOPTS
%                                 [h pc] =
ttest2(c_{isqs}(sci_{isqs}(j)), c_rand_set_(isqs, :));
%                           else
%                                 [h pc] =
ttest2(c_{isqs}(sci_{isqs}(j)), c_{isqs});
%                           end
%                           tt = ['Input sequence #' num2str(isqs) ':
best set sequence similarity region-overlapping' char(10) 'candidate (' s
SUFFIX{str2num(s(length(s))) + 1} ' best in total), its representative
str. (TED=' num2str(c_{isqs}(sci_{isqs}(j))) ', p='
num2str(num2str(round(pc*1000)/1000)) ')'];
%                     else
                            tt = [name ': best set sequence similarity'
newline 'region-overlapping candidate (' s SUFFIX{str2num(s(length(s))) +
1} ' best in total), its representative str. (TED='
num2str(c_{isqs}(sci_{isqs}(j))) ')'];
%                     end
                      hPatch = patch('XData', [pos-c pos+len-1+c
pos+len-1-c pos+c], ...
                                                      'YData',
[length(isqsor) - isqsfig + 1 - linestep - c length(isqsor) - isqsfig + 1
- linestep - c length(isqsor) - isqsfig + 1 - linestep + c length(isqsor)
- isqsfig + 1 - linestep + c], ...
                                                      'FaceColor', 'none',
'EdgeColor', 'none', ...
                                                      'HitTest', 'on',
'PickableParts', 'all', ...
                                                      'ButtonDownFcn',@(src,
evt) lineClicked(src, evt, [WP__ sq{1}{1}(2 : length(sq{1}{1}))
'_ss.png'], tt, isqsfig, hse.Position([1, 2])));
                      region = [region; [pos-c length(isqsor)-isqsfig+1-
linestep-c (pos+len-1+c)-(pos-c) 2*c]];
                      if TEST | BOOTSTRAP
                            if h_or_p_
%                                 text(pos+len+5, length(isqsor) -
isqsfig + 1 - linestep, ['sc='
num2str(round(sc_{isqs}(sci_{isqs}(j))*10)/10) ' (p='
num2str(round(psc_(isqs)*1000)/1000) '), TED='
num2str(round(c_{isqs}(sci_{isqs}(j))*10)/10) ], 'FontSize', 8 );
                                  text(pos+len+5, length(isqsor) -
isqsfig + 1 - linestep, ['sc='
num2str(round(sc_{isqs}(sci_{isqs}(j))*10)/10) ', p='
num2str(round(psc_(isqs)*1000)/1000) ], 'FontSize', 8 );
```

```
                            else
%                                 text(pos+len+5, length(isqsor) -
isqsfig + 1 - linestep, ['sc='
num2str(round(sc_{isqs}(sci_{isqs}(j))*10)/10) ' (' textinfig '='
num2str(hsc_(isqs)) '), TED='
num2str(round(c_{isqs}(sci_{isqs}(j))*10)/10) ], 'FontSize', 8 );
                                  text(pos+len+5, length(isqsor) -
isqsfig + 1 - linestep, ['sc='
num2str(round(sc_{isqs}(sci_{isqs}(j))*10)/10) ', ' textinfig '='
num2str(hsc_(isqs)) ], 'FontSize', 8 );
                            end
                      else
%                           text(pos+len+5, length(isqsor) - isqsfig + 1
- linestep, ['sc=' num2str(round(sc_{isqs}(sci_{isqs}(j))*10)/10) ',
TED=' num2str(round(c_{isqs}(sci_{isqs}(j))*10)/10) ], 'FontSize', 8 );
                            text(pos+len+5, length(isqsor) - isqsfig + 1
- linestep, ['sc=' num2str(round(sc_{isqs}(sci_{isqs}(j))*10)/10) ],
'FontSize', 8 );
                      end
% draw dots for Us in 3' tail
                      if DO_3_END_U
                            for i = len - 10 : len
                                  if sq{1}{2}(i) == 'U' | sq{1}{2}(i) ==
'T'
                                        plot(pos + i - 1, length(isqsor)
- isqsfig + 1 - linestep - .1, 'k.');
                                  end
                            end
                      end
                      linestep = linestep + .15;
                      break
                end
          end
      end
      if DO_ALTERNATIVE_CANDIDATES
% draw a line for best-at-all sopt followed with score and #
          [ii ii] = sort(c_{isqs}, 'ascend');
          look_for_overlap_flag = 1;
          for iii = 1 : DO_ALTERNATIVE_CANDIDATES
                if exist([WP__ fn_best_sopts_{isqs}{ii(iii)}], 'file')
                      id = fopen([WP__ fn_best_sopts_{isqs}{ii(iii)}]);
sq = textscan(id,'%s','delimiter',char(10)); fclose(id);
                      s = strsplit(fn_best_sopts_{isqs}{ii(iii)}, '.');
                      s = strsplit(s{length(s) - 1}, '_');
                      pos = str2num(s{4}(~isletter(s{4})));
                      ss = strsplit(s{3}, '=');
                      len = str2num(ss{2});

                      if TOO_OLD_RNA_VIENNA
                            disp(['No regular PS figure for ' WP__
fn_best_sopts_{isqs}{ii(iii)} '. Most likely too old version of RNA
Vienna package installed.'])
                      else
                            unix(['RNAplot -t1 < ' WP__
fn_best_sopts_{isqs}{ii(iii)}]);
                            unix(['gs -sDEVICE=pngalpha -dEPSCrop -o '
WP__ sq{1}{1}(2 : length(sq{1}{1})) '_ss.png ' sq{1}{1}(2 :
length(sq{1}{1})) '_ss.ps >> /dev/null 2>&1']);
```

```
                    unix(['rm ' sq{1}{1}(2 : length(sq{1}{1}))
'_ss.ps']);
                end

                lineHandle = line([pos pos+len-1],[length(isqsor)
- isqsfig + 1 - linestep length(isqsor) - isqsfig + 1 -
linestep],'color', [1 192/255 203/255]);
                % text(pos+len, length(isqsor) - isqsfig + 1 -
linestep, ['TED=' num2str(c_{isqs}(j)) ', z='
num2str(round((mean(c_{isqs}) - c_{isqs}(j)) ./ std(c_{isqs}) .* 10) /
10)], 'FontSize', 8, 'VerticalAlignment', 'middle');

                if iii > 1
                    s = num2str(iii);
                    tt = [name ': ' s
SUFFIX{str2num(s(length(s))) + 1} ' best single' newline 'structure
candidate, TED=' num2str(c_{isqs}(ii(iii))) ];
                else
                    tt = [name ': best single' newline 'structure
candidate, TED=' num2str(c_{isqs}(ii(iii))) ];
                end
                if TEST | BOOTSTRAP
                    if h_or_p_
                        tt = [tt ', p='
num2str(round(pc_(isqs)*1000)/1000)];
                        text(pos+len+5, length(isqsor) -
isqsfig + 1 - linestep, ['TED=' num2str(c_{isqs}(ii(iii))) ', p='
num2str(round(pc_(isqs)*1000)/1000) ], 'FontSize', 8);
                    else
                        tt = [tt ', ' textinfig '='
num2str(hc_(isqs)) ];
                        text(pos+len+5, length(isqsor) -
isqsfig + 1 - linestep, ['TED=' num2str(c_{isqs}(ii(iii))) ', ' textinfig
'=' num2str(hc_(isqs)) ], 'FontSize', 8);
                    end
                else
                    tt = [tt '.'];
                    text(pos+len+5, length(isqsor) - isqsfig + 1
- linestep, ['TED=' num2str(c_{isqs}(ii(iii)))], 'FontSize', 8);
                end
                if DO_OVERLAP_WITH_SEQUENCE_SIMILARITY_REGIONS &
exist('cnsd_rgn_') & ~isempty(cnsd_rgn_i) &
~isempty(cnsd_rgn_{cnsd_rgn_i, 2})
      % here, find out whether the best set overlaps sequence similarity
region because of the str. window title
      % here you have the sequence similarity region within the predicted
structure
                    if cnsd_rgn_{cnsd_rgn_i, 2} >= pos &
cnsd_rgn_{cnsd_rgn_i, 3} <= pos + len
      % here you have overlap that must be positive and longer tha 25% of
the structure sequence length
      %             left = pos; if cnsd_rgn_{cnsd_rgn_i, 2} >=
pos, left = cnsd_rgn_{cnsd_rgn_i, 2}; end
      %             right = pos + len - 1; if
cnsd_rgn_{cnsd_rgn_i, 3} <= pos + len - 1, right = cnsd_rgn_{cnsd_rgn_i,
3}; end
      %             if right - left > len / 100 * 25
```

```
                                tt = [tt ' Also overlaps sequence
similarity region. ' char(10)];
                                look_for_overlap_flag = 0;
                           end
                      end
                      hPatch = patch('XData', [pos-c pos+len-1+c
pos+len-1-c pos+c], 'YData', [length(isqsor) - isqsfig + 1 - linestep - c
length(isqsor) - isqsfig + 1 - linestep - c length(isqsor) - isqsfig + 1
- linestep + c length(isqsor) - isqsfig + 1 - linestep + c], ...
                                'FaceColor', 'none', 'EdgeColor',
'none', ...
                                'HitTest', 'on', 'PickableParts',
'all', ...
                                'ButtonDownFcn', @(src, evt)
lineClicked(src, evt, [WP__ sq{1}{1}(2 : length(sq{1}{1})) '_ss.png'],
tt, isqsfig + .5, hse.Position([1 2])));
                      region = [region; [pos-c length(isqsor)-isqsfig+1-
linestep-c (pos+len-1+c)-(pos-c) 2*c]];

                      if DO_3_END_U
                           for i = len - 10 : len
                                if sq{1}{2}(i) == 'U' | sq{1}{2}(i) ==
'T'
                                     plot(pos + i - 1, length(isqsor)
- isqsfig + 1 - linestep - .05, 'k.');
                                end
                           end
                      end
                      linestep = linestep + .15;
                 else
                      disp([WP__ fn_best_sopts_{isqs}{ii(iii)} ' not
exist. No picture.'])
                 end
            end
% find an individual predicted str that overlap the sequence similarity
region and draw it
            if look_for_overlap_flag &
DO_OVERLAP_WITH_SEQUENCE_SIMILARITY_REGIONS & ~isempty(cnsd_rgn_i) &
exist('cnsd_rgn_') & ~isempty(cnsd_rgn_{cnsd_rgn_i, 2})
                 for iii = DO_ALTERNATIVE_CANDIDATES + 1 : length(ii)
                      s = strsplit(fn_best_sopts_{isqs}{ii(iii)}, '.');
                      s = strsplit(s{length(s) - 1}, '_');
                      pos = str2num(s{4}(~isletter(s{4})));
                      ss = strsplit(s{3}, '=');
                      len = str2num(ss{2});
% here you have the sequence similarity region within the predicted
structure
                      if cnsd_rgn_{cnsd_rgn_i, 2} >= pos &
cnsd_rgn_{cnsd_rgn_i, 3} <= pos + len
% % here you test for overlap that must be positive and longer tha 25% of
the structure sequence length
%                     left = pos; if cnsd_rgn_{cnsd_rgn_i, 2} >= pos,
left = cnsd_rgn_{cnsd_rgn_i, 2}; end
%                     right = pos + len - 1; if cnsd_rgn_{cnsd_rgn_i, 3}
<= pos + len - 1, right = cnsd_rgn_{cnsd_rgn_i, 3}; end
%                     if right - left > len / 100 * 25
```

```
                              id = fopen([WP__
fn_best_sopts_{isqs}{ii(iii)}]); sq =
textscan(id,'%s','delimiter',char(10)); fclose(id);
                              if TOO_OLD_RNA_VIENNA
                                    disp(['No regular PS figure for ' WP__
fn_best_sopts_{isqs}{ii(iii)} '. Most likely too old version of RNA
Vienna package installed.'])
                              else
                                    unix(['RNAplot -t1 < ' WP__
fn_best_sopts_{isqs}{ii(iii)}]);
                                    unix(['gs -sDEVICE=pngalpha -dEPSCrop -
o ' WP__ sq{1}{1}(2 : length(sq{1}{1})) '_ss.png ' sq{1}{1}(2 :
length(sq{1}{1})) '_ss.ps >> /dev/null 2>&1']);
                                    unix(['rm ' sq{1}{1}(2 :
length(sq{1}{1})) '_ss.ps']);
                              end
                              lineHandle = line([pos pos+len-
1],[length(isqsor) - isqsfig + 1 - linestep length(isqsor) - isqsfig + 1
- linestep],'color', [1 192/255 203/255],'LineStyle','--');
                              s = num2str(iii) ;
                              if TEST | BOOTSTRAP
                                    if h_or_p_
                                          tt = [name ': best single str.
sequence similarity' newline 'region-overlapping str. (' s
SUFFIX{str2num(s(length(s))) + 1} ' best in total, TED='
num2str(c_{isqs}(ii(iii))) ', p=' num2str(round(pc_(isqs)*1000)/1000)
')'];
                                          text(pos+len+5, length(isqsor) -
isqsfig + 1 - linestep, ['TED=' num2str(c_{isqs}(ii(iii))) ', p='
num2str(round(pc_(isqs)*1000)/1000) ], 'FontSize', 8);
                                    else
%                                         tt = [name ': best single str.
sequence similarity' newline 'region-overlapping str. (' s
SUFFIX{str2num(s(length(s))) + 1} ' best in total, TED='
num2str(c_{isqs}(ii(iii))) ', ' textinfig '=' num2str(hc_(isqs)) ')'];
                                          tt = [name ': best single str.
sequence similarity' newline 'region-overlapping str. (' s
SUFFIX{str2num(s(length(s))) + 1} ' best in total, TED='
num2str(c_{isqs}(ii(iii))) ')'];
                                          text(pos+len+5, length(isqsor) -
isqsfig + 1 - linestep, ['TED=' num2str(c_{isqs}(ii(iii))) ', ' textinfig
'=' num2str(hc_(isqs)) ], 'FontSize', 8);
                                    end
                              else
                                    tt = [name ': best single str. sequence
similarity' newline 'region-overlapping candidate (' s
SUFFIX{str2num(s(length(s))) + 1} ' best in total, TED='
num2str(c_{isqs}(ii(iii))) ')'];
                                    text(pos+len+5, length(isqsor) -
isqsfig + 1 - linestep, ['TED=' num2str(c_{isqs}(ii(iii)))], 'FontSize',
8);
                              end
                              hPatch = patch('XData', [pos-c pos+len-1+c
pos+len-1-c pos+c], 'YData', [length(isqsor) - isqsfig + 1 - linestep - c
length(isqsor) - isqsfig + 1 - linestep - c length(isqsor) - isqsfig + 1
- linestep + c length(isqsor) - isqsfig + 1 - linestep + c], ...
                                                'FaceColor', 'none',
'EdgeColor', 'none', ...
```

```
                                        'HitTest', 'on',
'PickableParts', 'all', ...
                                        'ButtonDownFcn', @(src, evt)
lineClicked(src, evt, [WP__ sq{1}{1}(2 : length(sq{1}{1})) '_ss.png'],
tt, isqsfig + .5, hse.Position([1 2])));
                            region = [region; [pos-c length(isqsor)-
isqsfig+1-linestep-c (pos+len-1+c)-(pos-c) 2*c]];
                            if DO_3_END_U
                                 for i = len - 10 : len
                                      if sq{1}{2}(i) == 'U' |
sq{1}{2}(i) == 'T'
                                           plot(pos + i - 1,
length(isqsor) - isqsfig + 1 - linestep - .1, 'k.');
                                      end
                                 end
                            end
                            linestep = linestep + .15;
                            break
                       end
                  end
             end
        end
        if SHOW_SEQUENCE_SIMILARITY_REGIONS & exist('cnsd_rgn_') &
~isempty(cnsd_rgn_i) & ~isempty(cnsd_rgn_{cnsd_rgn_i, 2})
             linestep = linestep - .1;
% you can have more BLAST hits of template in a single input sq. Show
them all.
             i = find(contains(cnsd_rgn_(:, 1), cnsd_rgn_(cnsd_rgn_i)));
             for ii = 1 : length(i)
                  if cnsd_rgn_{i(ii), 4} == '+'
                       line([cnsd_rgn_{i(ii), 2} cnsd_rgn_{i(ii), 3}],
[length(isqsor) - isqsfig + 1 - linestep length(isqsor) - isqsfig + 1 -
linestep],'color', [0 .2 200/255]);
                       line(cnsd_rgn_{i(ii), 3}, length(isqsor) - isqsfig
+ 1 - linestep,'color', [0 .2 200/255],'marker','>','MarkerSize',3);
                  else
                       line([cnsd_rgn_{i(ii), 2} cnsd_rgn_{i(ii), 3}],
[length(isqsor) - isqsfig + 1 - linestep length(isqsor) - isqsfig + 1 -
linestep],'color', [0 .2 200/255]);
                       line(cnsd_rgn_{i(ii), 2}, length(isqsor) - isqsfig
+ 1 - linestep,'color', [0 .2 200/255],'marker','<','MarkerSize',3);
                  end
             end
             linestep = linestep + .05;
        end
        if exist('tt5_') & ~isempty(tt5_)
             for i = 1 : length(tt5_{isqs})
                  if ttor_{isqs}(i) == '+'
                       line(tt3_{isqs}(i), length(isqsor) - isqsfig + 1 -
linestep,'color', 'k','marker','>','MarkerSize',3);
                       line([tt5_{isqs}(i) tt3_{isqs}(i)],
[length(isqsor) - isqsfig + 1 - linestep length(isqsor) - isqsfig + 1 -
linestep],'color', 'k');
                  % else
                  %     line(tt5_{isqs}(i), length(isqsor) - isqsfig + 1 -
linestep,'color', 'k','marker','<','MarkerSize',3);
                  end
                  linestep = linestep + .05;
```

```
            end
      end
end
xls = ['Input sequence length (in nucleotides). Legend: blue lines: query
sequences, +/-: genomic orienta-' newline ...
         'tion relative to template, red lines (full/dashed): best/best
sequence similarity region-overlapping candidates,' newline];
if DO_ALTERNATIVE_CANDIDATES
      xls = [xls  'pink lines (full/dashed): best/best sequence
similarity region-overlapping alternative candidates using single'
newline 'structure, ' ];
end
if DO_3_END_U
      xls = [xls 'black dots: 3'' Us, '];
end
xls = [xls 'sc: TED-based score, TED: tree edit distance of predicted
structure' newline 'to template, '];
if DO_OVERLAP_WITH_SEQUENCE_SIMILARITY_REGIONS & exist('cnsd_rgn_')
      xls = [xls 'blue arrows: sequence similarity regions oriented
relatively to template, '];
end
if h_or_p_
      xls = [xls 'p: t-test p-value, '];
else
      xls = [xls 't: t-test decision (0-unsignif., 1-signif.), '];
end
if exist('tt5_') & ~isempty(tt5_)
      xls = [xls 'black arrows: terminators.'];
else
      xls(length(xls)) = [];
      xls(length(xls)) = '.';
end
xlabel(xls,'FontSize', 8);
a=axis;
if h_or_p_
      axis([a(1) max(ll) + max(ll) / 3 a(3) length(isqsor)]);
else
%     axis([a(1) max(ll) + max(ll) / 20 a(3) length(isqsor)]);
      axis([a(1) max(ll) + max(ll) / 20 a(3)+(length(isqsor)-1)*.01
length(isqsor)+length(isqsor)*.04])
end
if BOOTSTRAP
      title([fnsqs__ ',' newline fntm__ ', SAMPLING LENGTH='
num2str(SAMPLINGLENGTH) ', Alpha=' num2str(ALPHA__) ', BOOTSTRAPS='
num2str(size(sc_rand_set_, 2))], 'interpreter','none','FontSize', 8);
else
      title([fnsqs__ ',' newline fntm__ ', SAMPLING LENGTH='
num2str(SAMPLINGLENGTH) ', Alpha=' num2str(ALPHA__)],
'interpreter','none','FontSize', 8);
end
yticks({});
yticks(1:length(isqsor));
yticklabels(isqsor(length(isqsor) : -1 : 1));
ylabel('Input sequence #');
if SAVE_FIGURES
      % s = cut_string(fnsqs__, '/'); saveas(gcf,[s{length(s)}
'.eps'],'epsc');
      saveas(gcf, [fnload_ '.eps'],'epsc');
```

```
end

set(hse, 'WindowButtonMotionFcn', @(src, event) cursorChangeCallback(hse,
event, region));

clear xls i id isqsor isqs isqsfig sqs n s ss pos ll hse len id a fn
lineHandle sq t c data dcm_obj hPatch p psc pc h ii j region tt xls
linestep REMOVEFROMHEADERS SUFFIX
clear left right iii overlap_flag name cnsd_rgn_i
NUMBER_OF_BEST_SETS_TO_SHOW NUMBER_OF_BEST_SOPTS_TO_SHOW lua MAXNOOFSTRS
textinfig

function cursorChangeCallback(hse, ~, region)
% Get the current point
    currPoint = get(gca, 'CurrentPoint');
    x = currPoint(1, 1);
    y = currPoint(1, 2);
% Check if the cursor is within the region
       for i = 1 : size(region, 1)
      if x >= region(i, 1) && x <= region(i, 1) + region(i, 3) && y >=
region(i, 2) && y <= region(i, 2) + region(i, 4)
           set(hse, 'Pointer', 'hand'); % Change cursor to hand
                 % disp('I am inside.')
                 break
           else
           set(hse, 'Pointer', 'arrow'); % Default cursor
                 % disp('I am outside.')
           end
       end
end

function output_txt = customDatatipCallback(~, event_obj)
    if event_obj.Button == 1
      pos = get(event_obj, 'Position');
      output_txt = {['X: ', num2str(pos(1))], ['Y: ', num2str(pos(2))]};
    end
end

function lineClicked(src, event, fn, t, isqs, start)
      if strcmp(event.EventName, 'Hit')
           f = ancestor(src, 'figure');
           data = guidata(f);
           lineHandle = gco;
           if event.Button == 3
                 if exist(fn, 'file')
                       % if ~any(data.strs == lineHandle, "all")
                             fig=figure;
                             fig.MenuBar='none';
fig.ToolBar='none';fig.NumberTitle='off';
                             imshow(fn);
                             fig.Position = [start(1) - 300 - isqs * 50
start(2) - isqs * 150 400 500];
                             title(t, 'FontSize', 8,
'Interpreter','none');
%                            x = version;
%                            x = strsplit(x, '(');
%                            if (str2num(x{2}(4)) >= 2 & str2num(x{2}(5))
> 0) | (str2num(x{2}(4)) == 2 & str2num(x{2}(5)) == 0 & x{2}(6) == 'b')
```

```
%                                   ax = gca;
%                                   ax.TitleHorizontalAlignment = 'left';
%                             end
                              data.strs = [data.strs; lineHandle];
                        % end
                  end
            end
            if event.Button == 1
                  if ~any(data.value(:, 3) == lineHandle, "all")
                  xdata = get(lineHandle, 'XData');
                        ydata = get(lineHandle, 'YData');
                        if strfind(t, 'best str.')
                              hText1 = text(xdata(1), ydata(1) - .05,
num2str(round(xdata(1))), 'Horizontalalignment', 'center' );
                              hText2 = text(xdata(2), ydata(1) - .05,
num2str(round(xdata(2))), 'Horizontalalignment', 'center');
                        else
                              hText1 = text(xdata(1), ydata(1) + .15,
num2str(round(xdata(1))), 'Horizontalalignment', 'center');
                              hText2 = text(xdata(2), ydata(1) + .15,
num2str(round(xdata(2))), 'Horizontalalignment', 'center');
                        end
                        data.value = [data.value; [hText1 hText2
lineHandle]];
                        guidata(f, data);
                  else
                        i = find(data.value(:, 3) == lineHandle);
                        delete(data.value(i, 1)); delete(data.value(i,
2));
                        data.value(i, :) = [];
                  end
            end
            guidata(f, data);
      end
end
```

Evaluation of structure search

indx_complete.m: f_un_gr_ve, run_rsol_IGRs_complete.m.

Scripts

indx_complete.m

```
if SET == 1, indx_ =
{{'/mnt/new/pepik/pac_compare/SRP_bact_small_esccol3d-97-11-17-
stretched.pdb.br'; 'igrs_myco_avi/';'Mycoavi/'; '171';0}
{'/mnt/new/pepik/pac_compare/SRP_bact_small_esccol3d-97-11-17-
stretched.pdb.br'; 'igrs_therm_therm/';'Thermtherm_Dh/';'321';0}
{'/mnt/new/pepik/pac_compare/SRP_bact_small_esccol3d-97-11-17-
stretched.pdb.br';'igrs_Therm_prof/'; 'Thermprof';'81';0}
{'/mnt/new/pepik/pac_compare/SRP_bact_small_esccol3d-97-11-17-
stretched.pdb.br'; 'igrs_therm_cald/';'Thermcald/';'205';0}
{'~/rsol/SRP_bact_small_Eco_RF00169.fa.rf';
'igrs_therm_therm_RF/';'Thermtherm_RF/';'399';0}
{'/mnt/new/pepik/pactool/6S_Ec.br'; 'igrs_bac_sub/';'Bacsub/'; '997'; 0}
```

```
{'/mnt/new/pepik/pactool/6S_Ec.br';    'igrs_burk_sem/';
'Burksem/';'556';0};
{'/mnt/new/pepik/pactool/6S_Ec.br';    'igrs_rhiz_et_165nts/';
'Rhizet_165nts/';'2129';165};
{'/mnt/new/pepik/pac_compare/5S_Ec_1C2X.br';
'igrs_pyr_fur/';'Pyrfur_Dh/';'721';0}
{'/mnt/new/pepik/pac_compare/5S_Ec_1C2X.br'; 'igrs_syne_elo';
'Syneelo';'345';0}
{'RsmZ_Pae_7YR7.br';'igrs_azoto_vin'; 'Azotovin/';'1155';0}
{'RsmZ_Pae_7YR7.br';'igrs_azomo_agi/';'Azomoagi/';'1137';0}
{'RsmZ_Pae_7YR7.br.fa.rf';'igrs_azomo_agi/';'Azomoagi_rf/';'1138';0}
{'RsmZ_Pae_7YR7.br';'igrs_halops_pach/';'Halopspach/';'1855';0}
{'/mnt/new/pepik/pac_sopts/oxys_Ec.br';
'igrs_trab_odon/';'Trabodon_Dh/';'21';0};
{'/mnt/new/pepik/pac_sopts/oxys_Ec.br';
'igrs_yok_reg/';'Yokreg/';'608';0}
{'/mnt/new/pepik/pac_sopts/oxys_Ec.br'; 'igrs_lel_nimi/';
'Lelnim';'2951';0};
{'~/hs/spot42_Eco.rf';'igrs_Arsu_per/';'Arsuper_Dh/';'1365';0}
{'~/hs/spot42_Eco.rf';'igrs_Arsu_tuo/';'Arsutuo/';'939';0}
{'~/hs/spot42_Eco.rf';'igrs_vib_ishi/'; 'Vibish';'57';0};
{'~/hs/spot42_Eco.rf';'igrs_photo_aqua/';'Photoaqua';'2973';0};
{'/mnt/new/pepik/pactool/Msmest_n_7.br';'igrs_rhod_gann';'Rhodgann';'327'
;0}
{'/mnt/new/pepik/pactool/Msmest_n_7.br';
'igrs_sa_coe_230/';'Sacoe_230/';'1657';230}
{'/mnt/new/pepik/pactool/Msmest_n_7.br';'igrs_strp_viola_230/';
'Strpviola_230';'867';230}
{'/mnt/new/pepik/pactool/Msmest_n_7.br';'igrs_gor_polyi/';
'Gorpolyi_Dh/'; '163'; 0}
{'/mnt/new/pepik/pactool/Msmest_n_7.br';'igrs_gor_eff/';'Goreff_Dh/';
'411';0}
{'/mnt/new/pepik/pactool/Msmest_n_7.br';'igrs_gord_ara/';'Gordara_Dh/';
'307';0}
{'/mnt/new/pepik/pactool/Msmest_n_7.br';'igrs_actino_preti_230/';'Actinop
reti_230/';'91';230}
{'/mnt/new/pepik/pactool/Msmest_n_7.br';'igrs_alder_kun';'Alderkun';'271'
;0}
{'/mnt/new/pepik/pactool/Msmest_n_7.br';'igrs_Actinoallote_spi_230/';'Act
inoallotespi_230_Dh/';'1771';230}
{'/mnt/new/pepik/pactool/gcvb_St.br';
'igrs_Hae_infl/';'Haeinfl_Dh/';'347'; 0}
{'/mnt/new/pepik/pactool/gcvb_St.br';
'igrs_Photo_aq/';'Photoaq/';'2031';0}
{'gcvb_St.br.fa.rf'; 'igrs_Hae_infl/';'Haeinfl_rf/';'347';0}
{'gcvb_St.br.fa.rf'; 'igrs_Photo_aq/';'Photoaq_rf/';'2031';0}
{'DsrA_Eco_paper.fa.br';'igrs_Ced_net/';'Cednet';'1409';0}
{'DsrA_Eco_paper.fa.br'; 'igrs_Ced_dav/'; 'Ceddav/';'1339';0}
{'DsrA_Eco_paper.fa.br'; 'igrs_citr_gill/'; 'Citrgill/';'1185';0}
{'~/rsol/tRNA_Thr_Eco_1QF6.br';
'igrs_therm_mari_NC_023151/';'Thermmari_NC_023151/'; '135';0}
{'~/rsol/tRNA_Thr_Eco_1QF6.br';'igrs_stre_halo/'; 'Strehalo/'; '4951';0}
{'~/rsol/RNaseP_bact_a_Ec_7UO1.fa.rf';'igrs_thermotog_caldi/';'Thermotogc
aldi_Dh/'; '48';328}
{'~/rsol/RNaseP_bact_a_Ec_7UO1.fa.rf';'igrs_thermotog_prof/';'Thermotogpr
of_Dh/'; '38';333}
{'~/rsol/RNaseP_bact_a_Ec_7UO1.fa.rf';'igrs_pseudofrank_sapro/';'Pseudofr
anksapro_Dh/'; '1148';0}
```

```
{'~/rsol/RNaseP_bact_a_Ec_7UO1.fa.rf';'igrs_Amedibac_hom/';'Amedibachom_D
h/'; '385';0}
{'~/rsol/16S_Ec.fa.rf';'igrs_therm_mar_16S/';'Thermmar16S_Dh'; '5';0}
{'~/rsol/micF_Ec.br';'igrs_salm_bongo/';'Salmbongo_Dh';'1944';0}
{'~/rsol/micF_Ec.br';'igrs_salm_enter_typhi/';'Salmentertyphi_Dh';'1798';
0}
{'~/rsol/ryhb_Ec.br';'igrs_vibr_chol_eltor/';'Vibrcholeltor_Dh';'85';0}
{'~/rsol/ryhb_Ec.br';'igrs_vibr_chol_RFB16/';'VibrcholRFB16_Dh';'47';0}
};
d___ = '/mnt/spin/data_and_results_for_rsol.odt_1/';
end
if ~SET
indx_ = {
{'/mnt/new/pepik/pactool/Msmest_n_7.br';
'igrs_sa_coe/';'Sacoe/';'1045';0}
{'/mnt/new/pepik/pactool/Msmest_n_7.br';'igrs_strp_viola/';
'Strpviola/';'481'; 0}
{'/mnt/new/pepik/pactool/Msmest_n_7.br'; 'igrs_strp_viola_240';
'Strpviola_240_Dh/'; '805'; 240}
{'/mnt/new/pepik/pactool/Msmest_n_7.br';'igrs_strp_viola_220/';
'Strpviola_220/';'933'; 220}
{'/mnt/new/pepik/pactool/Msmest_n_7.br';'igrs_strp_viola_210/';
'Strpviola_210/'; '1012'; 210}
{'/mnt/new/pepik/pactool/Msmest_n_7.br';'igrs_gor_polyi_210/';
'Gorpolyi_210/'; '261'; 210}
{'/mnt/new/pepik/pactool/Msmest_n_7.br';'igrs_gor_polyi_200/';
'Gorpolyi_200/'; '277'; 200}
{'/mnt/new/pepik/pactool/Msmest_n_7.br';'igrs_actino_preti/';'Actinopreti
/';'63';0}
{'/mnt/new/pepik/pactool/Msmest_n_7.br';'igrs_actino_preti_210/';'Actinop
reti_210/';'99';210}
};
d___ = '/mnt/spin/data_and_results_for_rsol.odt_1/';
end
if SET == 2
indx_= {
{'~/hs/spot42_Eco.rf';'igrs_glac_siph/';'Glacsiph_Dh/';'178';0}
{'~/hs/spot42_Eco.rf';'igrs_glac_KUL10/';'GlacKUL10_Dh/';'953';0}
{'~/hs/spot42_Eco.rf';'igrs_glac_HTCC2999/';'GlacHTCC2999_Dh/';'489';0}
{'~/hs/spot42_Eco.rf';'igrs_glac_33A/';'Glac33A_Dh/';'2350';0}
{'~/hs/spot42_Eco.rf';'igrs_glac_MH2013/';'GlacMH2013_Dh/';'2065';0}
{'~/hs/spot42_Eco.rf';'igrs_glac_2405UD65/';'Glac2405UD65_Dh/';'1593';0}
{'~/hs/spot42_Eco.rf';'igrs_glac_MF2/';'GlacMF2_Dh/';'367';0}
};
d___ = '/mnt/spin/data_and_results_for_rsol.odt_1/';
end
if SET == 6
indx_= {
{'~/hs/spot42_Eco.rf';'igrs_pseudoalterom_halopl/';'Pseudoalteromhalopl_D
h/';'446';0}
{'~/hs/spot42_Eco.rf';'igrs_pseudoalterom_maricalo/';'Pseudoalterommarica
lo_Dh/';'183';0}
{'~/hs/spot42_Eco.rf';'igrs_pseudoalterom_atl/';'Pseudoalteromatl_Dh/';'2
01';0}
{'~/hs/spot42_Eco.rf';'igrs_pseudoalterom_pir/';'Pseudoalterompir_Dh/';'2
49';0}
{'~/hs/spot42_Eco.rf';'igrs_pseudoalterom_mar/';'Pseudoalterommar_Dh/';'1
967';0}
```

```
{'~/hs/spot42_Eco.rf';'igrs_pseudoalterom_sim/';'Pseudoalteromsim_Dh/';'1
493';0}
};
d___ = '/mnt/spin/data_and_results_for_rsol.odt_1/';
end

if SET==3
% usage examples
indx_= {
{'~/hs/spot42_Eco.rf';'igrs_glac_siph/';'Glacsiph_Dh/';'178';0}
{'~/hs/spot42_Eco.rf';'igrs_glac_KUL10/';'GlacKUL10_Dh/';'953';0}
{'~/hs/spot42_Eco.rf';'igrs_glac_HTCC2999/';'GlacHTCC2999_Dh/';'489';0}
{'~/hs/spot42_Eco.rf';'igrs_glac_33A/';'Glac33A_Dh/';'2350';0}
{'~/hs/spot42_Eco.rf';'igrs_glac_MH2013/';'GlacMH2013_Dh/';'2065';0}
% 6.
{'~/hs/spot42_Eco.rf';'igrs_glac_2405UD65/';'Glac2405UD65_Dh/';'1593';0}
{'~/hs/spot42_Eco.rf';'igrs_glac_MF2/';'GlacMF2_Dh/';'367';0}
{'~/hs/spot42_Eco.rf';'igrs_pseudoalterom_halopl/';'Pseudoalteromhalopl_D
h/';'445';0}
{'~/hs/spot42_Eco.rf';'igrs_pseudoalterom_maricalo/';'Pseudoalterommarica
lo_Dh/';'183';0}
{'~/hs/spot42_Eco.rf';'igrs_pseudoalterom_atl/';'Pseudoalteromatl_Dh/';'2
01';0}
% 11
{'~/hs/spot42_Eco.rf';'igrs_pseudoalterom_pir/';'Pseudoalterompir_Dh/';'2
49';0}
{'~/hs/spot42_Eco.rf';'igrs_pseudoalterom_mar/';'Pseudoalterommar_Dh/';'1
967';0}
{'~/hs/spot42_Eco.rf';'igrs_pseudoalterom_sim/';'Pseudoalteromsim_Dh/';'1
';0}
{'~/hs/spot42_Eco.rf';'igrs_burk_cecocep_J2315/';'BurkcenocepJ2315_Dh/';'
249';0}
{'~/hs/spot42_Eco.rf';'igrs_burkh_cep_BC16/';'BurkhcepBC16_Dh/';'285';0}
% 16
{'~/hs/spot42_Eco.rf';'igrs_burk_cenocep/';'Burkcenocep_Dh/';'289';0}
};
d___ = '';
end
if SET == 4
      indx_= {
      {'/mnt/new/pepik/pactool/Msmest_n_7.br';'igrs_frank_aln/';'Frankaln
_Dh/';'';0}
      {'/mnt/new/pepik/pactool/Msmest_n_7.br';'igrs_frank_casu/';'Frankca
su_Dh/';'';0}
      {'/mnt/new/pepik/pactool/Msmest_n_7.br';'igrs_frank_Allo2/';'FrankA
llo2_Dh/';'';0}
      {'/mnt/new/pepik/pactool/Msmest_n_7.br';'igrs_frank_B2/';'FrankB2_D
h/';'';0}
      {'/mnt/new/pepik/pactool/Msmest_n_7.br';'igrs_frank_ACN1ag/';'Frank
ACN1ag_Dh/';'';0}

      {'/mnt/new/pepik/pactool/Msmest_n_7.br';'igrs_frank_QA3/';'FrankQA3
_Dh/';'';0}
      {'/mnt/new/pepik/pactool/Msmest_n_7.br';'igrs_frank_CcI6/';'FrankCc
I6_Dh/';'';0}
      {'/mnt/new/pepik/pactool/Msmest_n_7.br';'igrs_frank_cana/';'Frankca
na_Dh/';'';0}
```

```
      {'/mnt/new/pepik/pactool/Msmest_n_7.br';'igrs_frank_BMG5/';'FrankBM
G5_Dh/';'';0}
      {'/mnt/new/pepik/pactool/Msmest_n_7.br';'igrs_frank_ArI3/';'FrankAr
I3_Dh/';'';0}

      {'/mnt/new/pepik/pactool/Msmest_n_7.br';'igrs_frank_CeD/';'FrankCeD
_Dh/';'';0}
      {'/mnt/new/pepik/pactool/Msmest_n_7.br';'igrs_frank_AvcI1/';'FrankA
vcI1_Dh/';'';0}
      {'/mnt/new/pepik/pactool/Msmest_n_7.br';'igrs_frank_R43/';'FrankR43
_Dh/';'';0}
      {'/mnt/new/pepik/pactool/Msmest_n_7.br';'igrs_frank_nepa/';'Frankne
pa_Dh/';'';0}
      {'/mnt/new/pepik/pactool/Msmest_n_7.br';'igrs_frank_torr/';'Frankto
rr_Dh/';'';0}

      {'/mnt/new/pepik/pactool/Msmest_n_7.br';'igrs_frank_umida/';'Franku
mida_Dh/';'';0}
      {'/mnt/new/pepik/pactool/Msmest_n_7.br';'igrs_frank_nodulisporulans
AgUm/';'FranknodulisporulansAgUm_Dh/';'';0}
% % split IGRs
      {'/mnt/new/pepik/pactool/Msmest_n_7.br';'igrs_frank_gta/';'Frankgta
_Dh/';'';0}
      {'/mnt/new/pepik/pactool/Msmest_n_7.br';'igrs_frank_tis/';'Franktis
_Dh/';'';0}
      {'/mnt/new/pepik/pactool/Msmest_n_7.br';'igrs_frank_EI5c/';'FrankEI
5c_Dh/';'';0}

      {'/mnt/new/pepik/pactool/Msmest_n_7.br';'igrs_frank_KB5/';'FrankKB5
_Dh/';'';0}
      {'/mnt/new/pepik/pactool/Msmest_n_7.br';'igrs_frank_alp/';'Frankalp
_Dh/';'';0}
% % uncertain contexts
      {'/mnt/new/pepik/pactool/Msmest_n_7.br';'igrs_frank_RB7/';'FrankRB7
_Dh/';'';0}
      {'/mnt/new/pepik/pactool/Msmest_n_7.br';'igrs_frank_Hr75/';'FrankHr
75_Dh/';'';0}
% % not long enough IGRs
      {'/mnt/new/pepik/pactool/Msmest_n_7.br';'igrs_frank_Cj5/';'FrankCj5
_Dh/';'';0}



      %
{'/mnt/new/pepik/pactool/Msmest_n_7.br';'igrs_frank_umida230/';'Frankumid
a_Dh_230/';'';230}
      %
{'/mnt/new/pepik/pactool/Msmest_n_7.br';'igrs_frank_casuarinae_230/';'Fra
nkcasuarinae_Dh_230/';'';230}
      %
{'/mnt/new/pepik/pactool/Msmest_n_7.br';'igrs_frank_canadensis230/';'Fran
kcanadensis_Dh_230/';'';230}
      %
{'/mnt/new/pepik/pactool/Msmest_n_7.br';'igrs_frank_tisae230/';'Franktisa
e_Dh_230/';'';230}
      %
{'/mnt/new/pepik/pactool/Msmest_n_7.br';'igrs_frank_gtarii_230/';'Frankgt
arii_Dh_230/';'';230}
      %
```

```
      %
{'/mnt/new/pepik/pactool/Msmest_n_7.br';'igrs_frank_torreyi230/';'Frankto
rreyi_Dh_230/';'';230}
      %
{'/mnt/new/pepik/pactool/Msmest_n_7.br';'igrs_frank_QA3230/';'FrankQA3_Dh
_230/';'';230}
      %
{'/mnt/new/pepik/pactool/Msmest_n_7.br';'igrs_frank_nepalensis_230/';'Fra
nknepalensis_Dh_230/';'';230}

      %
{'/home/pepik/hs/Ms1_Sco_published.fa.br';'igrs_frank_casuarinae_230/';'F
rankcasuarinae_Df_Sco/';'';0}
      %
{'/home/pepik/hs/Ms1_Sco_published.fa.br';'igrs_frank_nepalensis_230/';'F
ranknepalensis_Df_Sco/';'';0}
      %
{'/home/pepik/hs/Ms1_Sco_published.fa.br';'igrs_frank_gtarii_230/';'Frank
gtarii_Df_Sco/';'';0}
      };
      d___ = '/mnt/spin/data_and_results_for_rsol.odt_1/';
end
if SET == 44
      indx_= {
      {'/home/pepik/hs/Ms1_Sco_published.fa.br';'igrs_frank_casuarinae_23
0/';'Frankcasuarinae_Dh_Sco/';'';0}
      {'/home/pepik/hs/Ms1_Sco_published.fa.br';'igrs_frank_nepalensis_23
0/';'Franknepalensis_Dh_Sco/';'';0}
      {'/home/pepik/hs/Ms1_Sco_published.fa.br';'igrs_frank_torreyi230/';
'Franktorreyi_Dh_Sco/';'';0}
      {'/home/pepik/hs/Ms1_Sco_published.fa.br';'igrs_frank_umida230/';'F
rankumida_Dh_Sco/';'';0}
      {'/home/pepik/hs/Ms1_Sco_published.fa.br';'igrs_frank_canadensis230
/';'Frankcanadensis_Dh_Sco/';'';0}

      {'/home/pepik/hs/Ms1_Sco_published.fa.br';'igrs_frank_QA3230/';'Fra
nkQA3_Dh_Sco/';'';0}
      {'/home/pepik/hs/Ms1_Sco_published.fa.br';'igrs_frank_R43230/';'Fra
nkR43_Dh_Sco/';'';0}
      {'/home/pepik/hs/Ms1_Sco_published.fa.br';'igrs_frank_alni230/';'Fr
ankalni_Dh_Sco/';'';0}
      {'/home/pepik/hs/Ms1_Sco_published.fa.br';'igrs_frank_ArI3230/';'Fr
ankArI3_Dh_Sco/';'';0}
      {'/home/pepik/hs/Ms1_Sco_published.fa.br';'igrs_frank_CcI6230/';'Fr
ankCcI6_Dh_Sco/';'';0}

      {'/home/pepik/hs/Ms1_Sco_published.fa.br';'igrs_frank_AvcI1230/';'F
rankAvcI1_Dh_Sco/';'';0}
      {'/home/pepik/hs/Ms1_Sco_published.fa.br';'igrs_frank_BMG5230/';'Fr
ankBMG5_Dh_Sco/';'';0}
      {'/home/pepik/hs/Ms1_Sco_published.fa.br';'igrs_frank_Allo2230/';'F
rankAllo2_Dh_Sco/';'';0}
      {'/home/pepik/hs/Ms1_Sco_published.fa.br';'igrs_frank_CeD230/';'Fra
nkCeD_Dh_Sco/';'';0}
      {'/home/pepik/hs/Ms1_Sco_published.fa.br';'igrs_frank_nodulisporula
nsAgUm230/';'FranknodulisporulansAgUm_Dh_Sco/';'';0}
```

```
      {'/home/pepik/hs/Ms1_Sco_published.fa.br';'igrs_frank_ACN1ag230/';'
FrankACN1ag_Dh_Sco/';'';0}
      {'/home/pepik/hs/Ms1_Sco_published.fa.br';'igrs_frank_B2230/';'Fran
kB2_Dh_Sco/';'';0}
% split IGRs
      {'/home/pepik/hs/Ms1_Sco_published.fa.br';'igrs_frank_gtarii_230/';
'Frankgtarii_Dh_Sco/';'';0}
      {'/home/pepik/hs/Ms1_Sco_published.fa.br';'igrs_frank_tisae230/';'F
ranktisae_Dh_Sco/';'';0}
      {'/home/pepik/hs/Ms1_Sco_published.fa.br';'igrs_frank_EI5c230/';'Fr
ankEI5c_Dh_Sco//';'';0}
% not long enough IGRs
      {'/home/pepik/hs/Ms1_Sco_published.fa.br';'igrs_frank_Cj5230/';'Fra
nkCj5_Dh_Sco//';'';0}
      };
      d___ = '/mnt/spin/data_and_results_for_rsol.odt_1/';
end
if SET == 9
      indx_= {
      {'/mnt/new/pepik/pactool/Msmest_n_7.br';'igrs_Sali_arenicola/';'Sal
iarenicola_Dh/';'';0}
      {'/mnt/new/pepik/pactool/Msmest_n_7.br';'igrs_Sali_vitiensis/';'Sal
ivitiensis_Dh/';'';0}
      {'/mnt/new/pepik/pactool/Msmest_n_7.br';'igrs_Sali_mooreana/';'Sali
mooreana_Dh/';'';0}
      {'/mnt/new/pepik/pactool/Msmest_n_7.br';'igrs_Sali_cortesiana/';'Sa
licortesiana_Dh/';'';0}
      {'/mnt/new/pepik/pactool/Msmest_n_7.br';'igrs_Sali_oceanensis/';'Sa
lioceanensis_Dh/';'';0}

      {'/mnt/new/pepik/pactool/Msmest_n_7.br';'igrs_Sali_fenicalii/';'Sal
ifenicalii_Dh/';'';0}
      {'/mnt/new/pepik/pactool/Msmest_n_7.br';'igrs_Sali_tropica/';'Salit
ropica_Dh/';'';0}
      {'/mnt/new/pepik/pactool/Msmest_n_7.br';'igrs_Sali_pacifica/';'Sali
pacifica_Dh/';'';0}

%
      {'/mnt/new/pepik/pactool/Msmest_n_7.br';'igrs_Sali_arenicola230/';'
Saliarenicola_Dh_230/';'';230}
%
      {'/mnt/new/pepik/pactool/Msmest_n_7.br';'igrs_Sali_vitiensis230/';'
Salivitiensis_Dh_230/';'';230}
%
      {'/mnt/new/pepik/pactool/Msmest_n_7.br';'igrs_Sali_mooreana230/';'S
alimooreana_Dh_230/';'';230}
%
      {'/mnt/new/pepik/pactool/Msmest_n_7.br';'igrs_Sali_cortesiana230/';
'Salicortesiana_Dh_230/';'';230}
%
      {'/mnt/new/pepik/pactool/Msmest_n_7.br';'igrs_Sali_oceanensis230/';
'Salioceanensis_Dh_230/';'';230}
%
%
      {'/mnt/new/pepik/pactool/Msmest_n_7.br';'igrs_Sali_fenicalii230/';'
Salifenicalii_Dh_230/';'';230}
```

```
%
      {'/mnt/new/pepik/pactool/Msmest_n_7.br';'igrs_Sali_tropica230/';'Sa
litropica_Dh_230/';'';230}
%
      {'/mnt/new/pepik/pactool/Msmest_n_7.br';'igrs_Sali_pacifica230/';'S
alipacifica_Dh_230/';'';230}
      };
      d___ = '/mnt/spin/data_and_results_for_rsol.odt_1/';
      d___ = '';
end
if SET == 10
      indx_= {
%
      {'/mnt/new/pepik/pactool/Msmest_n_7.br';'igrs_Glyco_arizonensis/';'
Glycoarizonensis_Dh/';'';0}
%
      {'/mnt/new/pepik/pactool/Msmest_n_7.br';'igrs_Glyco_dulcitolivorans
/';'Glycodulcitolivorans_Dh/';'';0}
%
      {'/mnt/new/pepik/pactool/Msmest_n_7.br';'igrs_Glyco_artemisiae/';'G
lycoartemisiae_Dh/';'';0}
%
      {'/mnt/new/pepik/pactool/Msmest_n_7.br';'igrs_Glyco_terrestris/';'G
lycoterrestris_Dh/';'';0}
%
      {'/mnt/new/pepik/pactool/Msmest_n_7.br';'igrs_Glyco_buryatensis/';'
Glycoburyatensis_Dh/';'';0}
%
%
      {'/mnt/new/pepik/pactool/Msmest_n_7.br';'igrs_Glyco_paridis/';'Glyc
oparidis_Dh/';'';0}
%
      {'/mnt/new/pepik/pactool/Msmest_n_7.br';'igrs_Glyco_tenuis/';'Glyco
tenuis_Dh/';'';0}
%
      {'/mnt/new/pepik/pactool/Msmest_n_7.br';'igrs_Glyco_xiaoerkulensis/
';'Glycoxiaoerkulensis_Dh/';'';0}

%
      {'/mnt/new/pepik/pactool/Msmest_n_7.br';'igrs_Glyco_arizonensis230/
';'Glycoarizonensis_Dh_230/';'';230}
%
      {'/mnt/new/pepik/pactool/Msmest_n_7.br';'igrs_Glyco_dulcitolivorans
230/';'Glycodulcitolivorans_Dh_230/';'';230}
%
      {'/mnt/new/pepik/pactool/Msmest_n_7.br';'igrs_Glyco_artemisiae230/'
;'Glycoartemisiae_Dh_230/';'';230}
%
      {'/mnt/new/pepik/pactool/Msmest_n_7.br';'igrs_Glyco_terrestris230/'
;'Glycoterrestris_Dh_230/';'';230}
%
      {'/mnt/new/pepik/pactool/Msmest_n_7.br';'igrs_Glyco_buryatensis230/
';'Glycoburyatensis_Dh_230/';'';230}
%
%
      {'/mnt/new/pepik/pactool/Msmest_n_7.br';'igrs_Glyco_paridis230/';'G
lycoparidis_Dh_230/';'';230}
```

```
%
      {'/mnt/new/pepik/pactool/Msmest_n_7.br';'igrs_Glyco_tenuis230/';'Gl
ycotenuis_Dh_230/';'';230}
%
      {'/mnt/new/pepik/pactool/Msmest_n_7.br';'igrs_Glyco_xiaoerkulensis2
30/';'Glycoxiaoerkulensis_Dh_230/';'';230}

      {'/home/pepik/hs/Ms1_Sco_published.fa.br';'igrs_Glyco_arizonensis23
0/';'Glycoarizonensis_Dh_230_Sco/';'';0}
      {'/home/pepik/hs/Ms1_Sco_published.fa.br';'igrs_Glyco_dulcitolivora
ns230/';'Glycodulcitolivorans_Dh_230_Sco/';'';0}
      {'/home/pepik/hs/Ms1_Sco_published.fa.br';'igrs_Glyco_artemisiae230
/';'Glycoartemisiae_Dh_230_Sco/';'';0}
      {'/home/pepik/hs/Ms1_Sco_published.fa.br';'igrs_Glyco_terrestris230
/';'Glycoterrestris_Dh_230_Sco/';'';0}
      {'/home/pepik/hs/Ms1_Sco_published.fa.br';'igrs_Glyco_buryatensis23
0/';'Glycoburyatensis_Dh_230_Sco/';'';0}

      {'/home/pepik/hs/Ms1_Sco_published.fa.br';'igrs_Glyco_paridis230/';
'Glycoparidis_Dh_230_Sco/';'';0}
      {'/home/pepik/hs/Ms1_Sco_published.fa.br';'igrs_Glyco_tenuis230/';'
Glycotenuis_Dh_230_Sco/';'';0}
      {'/home/pepik/hs/Ms1_Sco_published.fa.br';'igrs_Glyco_xiaoerkulensi
s230/';'Glycoxiaoerkulensis_Dh_230_Sco/';'';0}
      };
%     d___ = '/mnt/spin/data_and_results_for_rsol.odt_1/';
      d___ = '';
end
if SET == 5
% % from /mnt/spin/anns_Bacteria_sa/Bifidobacterium
      indx_= {
      {'/mnt/new/pepik/pactool/Msmest_n_7.br';'igrs_Bifi_adolescentis/';'
Bifiadolescentis_Dh/';'';0}
      {'/mnt/new/pepik/pactool/Msmest_n_7.br';'igrs_Bifi_angulatum/';'Bif
iangulatum_Dh/';'';0}
      {'/mnt/new/pepik/pactool/Msmest_n_7.br';'igrs_Bifi_animalis/';'Bifi
animalis_Dh/';'';0}
      {'/mnt/new/pepik/pactool/Msmest_n_7.br';'igrs_Bifi_bifidum/';'Bifib
ifidum_Dh/';'';0}
      {'/mnt/new/pepik/pactool/Msmest_n_7.br';'igrs_Bifi_dentium/';'Bifid
entium_Dh/';'';0}

      {'/mnt/new/pepik/pactool/Msmest_n_7.br';'igrs_Bifi_gallicum/';'Bifi
gallicum_Dh/';'';0}
      {'/mnt/new/pepik/pactool/Msmest_n_7.br';'igrs_Bifi_longum/';'Bifilo
ngum_Dh/';'';0}
      {'/mnt/new/pepik/pactool/Msmest_n_7.br';'igrs_Bifi_magnum/';'Bifima
gnum_Dh/';'';0}
      {'/mnt/new/pepik/pactool/Msmest_n_7.br';'igrs_Bifi_minimum/';'Bifim
inimum_Dh/';'';0}
      {'/mnt/new/pepik/pactool/Msmest_n_7.br';'igrs_Bifi_pseudolongum/';'
Bifipseudolongum_Dh/';'';0}
      };
%     d___ = '/mnt/spin/data_and_results_for_rsol.odt_1/';
      d___ = '';
end
if SET == 55
% % from /mnt/spin/anns_Bacteria_sa/Bifidobacterium
```

```
      indx_= {
      {'/home/pepik/hs/Ms1_Sco_published.fa.br';'igrs_Bifi_adolescentis_S
co/';'Bifiadolescentis_Dh_Sco/';'';0}
      {'/home/pepik/hs/Ms1_Sco_published.fa.br';'igrs_Bifi_angulatum_Sco/
';'Bifiangulatum_Dh_Sco/';'';0}
      {'/home/pepik/hs/Ms1_Sco_published.fa.br';'igrs_Bifi_animalis_Sco/'
;'Bifianimalis_Dh_Sco/';'';0}
      {'/home/pepik/hs/Ms1_Sco_published.fa.br';'igrs_Bifi_bifidum_Sco/';
'Bifibifidum_Dh_Sco/';'';0}
      {'/home/pepik/hs/Ms1_Sco_published.fa.br';'igrs_Bifi_boum_Sco/';'Bi
fiboum_Dh_Sco/';'';0}

      {'/home/pepik/hs/Ms1_Sco_published.fa.br';'igrs_Bifi_choerinum_Sco/
';'Bifichoerinum_Dh_Sco/';'';0}
      {'/home/pepik/hs/Ms1_Sco_published.fa.br';'igrs_Bifi_dentium_Sco/';
'Bifidentium_Dh_Sco/';'';0}
      {'/home/pepik/hs/Ms1_Sco_published.fa.br';'igrs_Bifi_gallicum_Sco/'
;'Bifigallicum_Dh_Sco/';'';0}
      {'/home/pepik/hs/Ms1_Sco_published.fa.br';'igrs_Bifi_longum_Sco/';'
Bifilongum_Dh_Sco/';'';0}
      {'/home/pepik/hs/Ms1_Sco_published.fa.br';'igrs_Bifi_magnum_Sco/';'
Bifimagnum_Dh_Sco/';'';0}

%
      {'/home/pepik/hs/Ms1_Sco_published.fa.br';'igrs_Bifi_minimum_Sco/';
'Bifiminimum_Dh_Sco/';'';0}
      {'/home/pepik/hs/Ms1_Sco_published.fa.br';'igrs_Bifi_moukalabense_S
co/';'Bifimoukalabense_Dh_Sco/';'';0}
      {'/home/pepik/hs/Ms1_Sco_published.fa.br';'igrs_Bifi_pseudolongum_S
co/';'Bifipseudolongum_Dh_Sco/';'';0}
      {'/home/pepik/hs/Ms1_Sco_published.fa.br';'igrs_Bifi_subtile_Sco/';
'Bifisubtile_Dh_Sco/';'';0}
      {'/home/pepik/hs/Ms1_Sco_published.fa.br';'igrs_Bifi_thermacidophil
um_Sco/';'Bifithermacidophilum_Dh_Sco/';'';0}

      {'/home/pepik/hs/Ms1_Sco_published.fa.br';'igrs_Bifi_tsurumiense_Sc
o/';'Bifitsurumiense_Dh_Sco/';'';0}
      };
%     d___ = '/mnt/spin/data_and_results_for_rsol.odt_1/';
      d___ = '';
end
if SET == 555
% % from /mnt/spin/anns_Bacteria_sa_alt/Bifidobacterium
      indx_= {
      {'/mnt/new/pepik/pactool/Msmest_n_7.br';'igrs_Bifi_A11_alt/';'BifiA
11_alt_Dh/';'';0}
      {'/mnt/new/pepik/pactool/Msmest_n_7.br';'igrs_Bifi_adolescentis_alt
/';'Bifiadolescentis_alt_Dh/';'';0}
      {'/mnt/new/pepik/pactool/Msmest_n_7.br';'igrs_Bifi_angulatum_alt/';
'Bifiangulatum_alt_Dh/';'';0}
      {'/mnt/new/pepik/pactool/Msmest_n_7.br';'igrs_Bifi_animalis_alt/';'
Bifianimalis_alt_Dh/';'';0}
      {'/mnt/new/pepik/pactool/Msmest_n_7.br';'igrs_Bifi_bifidum_alt/';'B
ifibifidum_alt_Dh/';'';0}

      {'/mnt/new/pepik/pactool/Msmest_n_7.br';'igrs_Bifi_bombi_alt/';'Bif
ibombi_alt_Dh/';'';0}
```

```
      {'/mnt/new/pepik/pactool/Msmest_n_7.br';'igrs_Bifi_boum_alt/';'Bifi
boum_alt_Dh/';'';0}
      {'/mnt/new/pepik/pactool/Msmest_n_7.br';'igrs_Bifi_catenulatum_alt/
';'Bificatenulatum_alt_Dh/';'';0}
      {'/mnt/new/pepik/pactool/Msmest_n_7.br';'igrs_Bifi_choerinum_alt/';
'Bifichoerinum_alt_Dh/';'';0}
      {'/mnt/new/pepik/pactool/Msmest_n_7.br';'igrs_Bifi_dentium_alt/';'B
ifidentium_alt_Dh/';'';0}

      {'/mnt/new/pepik/pactool/Msmest_n_7.br';'igrs_Bifi_gallicum_alt/';'
Bifigallicum_alt_Dh/';'';0}
% No IGR for ms1, see rsol_Bifi.odt
      {'/mnt/new/pepik/pactool/Msmest_n_7.br';'igrs_Bifi_longum_alt/';'Bi
filongum_alt_Dh/';'';0}
% No IGR for ms1, see rsol_Bifi.odt
      {'/mnt/new/pepik/pactool/Msmest_n_7.br';'igrs_Bifi_magnum_alt/';'Bi
fimagnum_alt_Dh/';'';0}
      {'/mnt/new/pepik/pactool/Msmest_n_7.br';'igrs_Bifi_minimum_alt/';'B
ifiminimum_alt_Dh/';'';0}
      {'/mnt/new/pepik/pactool/Msmest_n_7.br';'igrs_Bifi_moukalabense_alt
/';'Bifimoukalabense_alt_Dh/';'';0}

      {'/mnt/new/pepik/pactool/Msmest_n_7.br';'igrs_Bifi_pseudocatenulatu
m_alt/';'Bifipseudocatenulatum_alt_Dh';'';0}
      {'/mnt/new/pepik/pactool/Msmest_n_7.br';'igrs_Bifi_pseudolongum_alt
/';'Bifipseudolongum_alt_Dh/';'';0}
      {'/mnt/new/pepik/pactool/Msmest_n_7.br';'igrs_Bifi_ruminantium_alt/
';'Bifiruminantium_alt_Dh';'';0}
% No IGR for ms1, see rsol_Bifi.odt
      {'/mnt/new/pepik/pactool/Msmest_n_7.br';'igrs_Bifi_subtile_alt/';'B
ifisubtile_alt_Dh/';'';0}
      {'/mnt/new/pepik/pactool/Msmest_n_7.br';'igrs_Bifi_thermacidophilum
_alt/';'Bifithermacidophilum_alt_Dh/';'';0}

      {'/mnt/new/pepik/pactool/Msmest_n_7.br';'igrs_Bifi_thermophilum_alt
/';'Bifithermophilum_alt_Dh/';'';0}
      {'/mnt/new/pepik/pactool/Msmest_n_7.br';'igrs_Bifi_tsurumiense_alt/
';'Bifitsurumiense_alt_Dh/';'';0}
      };
%     d___ = '/mnt/spin/data_and_results_for_rsol.odt_1/';
      d___ = '';
end
if SET == 5555
% % from /mnt/spin/anns_Bacteria_sa_alt/Bifidobacterium
      indx_= {
      {'/home/pepik/hs/Ms1_Sco_published.fa.br';'igrs_Bifi_A11_alt_230/';
'BifiA11_alt_230_Dh/';'';0}
      {'/home/pepik/hs/Ms1_Sco_published.fa.br';'igrs_Bifi_adolescentis_a
lt_230/';'Bifiadolescentis_alt_230_Dh/';'';0}
      {'/home/pepik/hs/Ms1_Sco_published.fa.br';'igrs_Bifi_angulatum_alt_
230/';'Bifiangulatum_alt_230_Dh/';'';0}
      {'/home/pepik/hs/Ms1_Sco_published.fa.br';'igrs_Bifi_animalis_alt_2
30/';'Bifianimalis_alt_230_Dh/';'';0}
      {'/home/pepik/hs/Ms1_Sco_published.fa.br';'igrs_Bifi_bifidum_alt_23
0/';'Bifibifidum_alt_230_Dh/';'';0}

      {'/home/pepik/hs/Ms1_Sco_published.fa.br';'igrs_Bifi_bombi_alt_230/
';'Bifibombi_alt_230_Dh/';'';0}
```

```
      {'/home/pepik/hs/Ms1_Sco_published.fa.br';'igrs_Bifi_boum_alt_230/'
;'Bifiboum_alt_230_Dh/';'';0}
      {'/home/pepik/hs/Ms1_Sco_published.fa.br';'igrs_Bifi_choerinum_alt_
230/';'Bifichoerinum_alt_230_Dh/';'';0}
      {'/home/pepik/hs/Ms1_Sco_published.fa.br';'igrs_Bifi_catenulatum_al
t_230/';'Bificatenulatum_alt_230_Dh/';'';0}
      {'/home/pepik/hs/Ms1_Sco_published.fa.br';'igrs_Bifi_dentium_alt_23
0/';'Bifidentium_alt_230_Dh/';'';0}

      {'/home/pepik/hs/Ms1_Sco_published.fa.br';'igrs_Bifi_gallicum_alt_2
30/';'Bifigallicum_alt_230_Dh/';'';0}
      {'/home/pepik/hs/Ms1_Sco_published.fa.br';'igrs_Bifi_longum_alt_230
/';'Bifilongum_alt_230_Dh/';'';0}
      {'/home/pepik/hs/Ms1_Sco_published.fa.br';'igrs_Bifi_magnum_alt_230
/';'Bifimagnum_alt_230_Dh/';'';0}
      {'/home/pepik/hs/Ms1_Sco_published.fa.br';'igrs_Bifi_minimum_alt_23
0/';'Bifiminimum_alt_230_Dh/';'';0}
      {'/home/pepik/hs/Ms1_Sco_published.fa.br';'igrs_Bifi_moukalabense_a
lt_230/';'Bifimoukalabense_alt_230_Dh/';'';0}

      {'/home/pepik/hs/Ms1_Sco_published.fa.br';'igrs_Bifi_pseudocatenula
tum_alt_230/';'Bifipseudocatenulatum_alt_230_Dh/';'';0}
      {'/home/pepik/hs/Ms1_Sco_published.fa.br';'igrs_Bifi_pseudolongum_a
lt_230/';'Bifipseudolongum_alt_230_Dh/';'';0}
      {'/home/pepik/hs/Ms1_Sco_published.fa.br';'igrs_Bifi_ruminantium_al
t_230/';'Bifiruminantium_alt_230_Dh';'';0}
      {'/home/pepik/hs/Ms1_Sco_published.fa.br';'igrs_Bifi_subtile_alt_23
0/';'Bifisubtile_alt_230_Dh/';'';0}
      {'/home/pepik/hs/Ms1_Sco_published.fa.br';'igrs_Bifi_thermacidophil
um_alt_230/';'Bifithermacidophilum_alt_230_Dh/';'';0}

      {'/home/pepik/hs/Ms1_Sco_published.fa.br';'igrs_Bifi_thermophilum_a
lt_230/';'Bifithermophilum_alt_230_Dh';'';0}
      {'/home/pepik/hs/Ms1_Sco_published.fa.br';'igrs_Bifi_tsurumiense_al
t_230/';'Bifitsurumiense_alt_230_Dh/';'';0}
      };
%     d___ = '/mnt/spin/data_and_results_for_rsol.odt_1/';
      d___ = '';
end
if SET == 55555
% % from /mnt/spin/anns_Bacteria_sa_alt/Bifidobacterium
      indx_= {
      {'/home/pepik/hs/Ms1_Sco_published.fa.br';'igrs_Bifi_angulatum_alt_
230/';'Bifiangulatum_alt_230_Df/';'';0}
      {'/home/pepik/hs/Ms1_Sco_published.fa.br';'igrs_Bifi_bombi_alt_230/
';'Bifibombi_alt_230_Df/';'';0}
      {'/home/pepik/hs/Ms1_Sco_published.fa.br';'igrs_Bifi_boum_alt_230/'
;'Bifiboum_alt_230_Df/';'';0}
      {'/home/pepik/hs/Ms1_Sco_published.fa.br';'igrs_Bifi_choerinum_alt_
230/';'Bifichoerinum_alt_230_Df/';'';0}
      {'/home/pepik/hs/Ms1_Sco_published.fa.br';'igrs_Bifi_catenulatum_al
t_230/';'Bificatenulatum_alt_230_Df/';'';0}

      {'/home/pepik/hs/Ms1_Sco_published.fa.br';'igrs_Bifi_minimum_alt_23
0/';'Bifiminimum_alt_230_Df/';'';0}
      {'/home/pepik/hs/Ms1_Sco_published.fa.br';'igrs_Bifi_moukalabense_a
lt_230/';'Bifimoukalabense_alt_230_Df/';'';0}
```

```
      {'/home/pepik/hs/Ms1_Sco_published.fa.br';'igrs_Bifi_thermacidophil
um_alt_230/';'Bifithermacidophilum_alt_230_Df/';'';0}
      {'/home/pepik/hs/Ms1_Sco_published.fa.br';'igrs_Bifi_thermophilum_a
lt_230/';'Bifithermophilum_alt_230_Df';'';0}
      {'/home/pepik/hs/Ms1_Sco_published.fa.br';'igrs_Bifi_tsurumiense_al
t_230/';'Bifitsurumiense_alt_230_Df/';'';0}
      };
%     d___ = '/mnt/spin/data_and_results_for_rsol.odt_1/';
      d___ = '';
end
if SET == 7
indx_= {
{'~/hs/spot42_Eco.rf';'igrs_king_bona/';'Kingbona_Dh/';'';0}
{'~/hs/spot42_Eco.rf';'igrs_king_deni/';'Kingdeni_Dh/';'';0}
{'~/hs/spot42_Eco.rf';'igrs_king_king/';'Kingking_Dh/';'';0}
% strain SW7208426
{'~/hs/spot42_Eco.rf';'igrs_king_neg/';'Kingneg_Dh/';'';0}
% oralis_ATCC_51147
{'~/hs/spot42_Eco.rf';'igrs_king_or/';'Kingor_Dh/';'';0}
{'~/hs/spot42_Eco.rf';'igrs_king_pum/';'Kingpum_Dh/';'';0}
{'~/hs/spot42_Eco.rf';'igrs_king_oralisDSM/';'KingoralisDSM_Dh/';'';0}
{'~/hs/spot42_Eco.rf';'igrs_king_SNUBH/';'KingSNUBH_Dh/';'';0}
{'~/hs/spot42_Eco.rf';'igrs_king_potus/';'Kingpotus_Dh/';'';0}
{'~/hs/spot42_Eco.rf';'igrs_king_kingATCC23330/';'KingkingATCC23330_Dh/';
'';0}
{'~/hs/spot42_Eco.rf';'igrs_king_kingNCTC10529/';'KingkingNCTC10529_Dh/';
'';0}
{'~/hs/spot42_Eco.rf';'igrs_king_negD2292/';'KingnegD2292_Dh/';'';0}
{'~/hs/spot42_Eco.rf';'igrs_king_kingPYKK081/';'KingkingPYKK081_Dh/';'';0
}
{'~/hs/spot42_Eco.rf';'igrs_king_negCC443/';'KingnegCC443_Dh/';'';0}
{'~/hs/spot42_Eco.rf';'igrs_king_neg_BB331/';'KingnegBB331_Dh/';'';0}
{'~/hs/spot42_Eco.rf';'igrs_king_neg_AA267/';'KingnegAA267_Dh/';'';0}
{'~/hs/spot42_Eco.rf';'igrs_king_neg_EPA014/';'KingnegEPA014_Dh/';'';0}
{'~/hs/spot42_Eco.rf';'igrs_king_neg_PVC1712/';'KingnegPVC1712_Dh/';'';0}
};
d___ = '/mnt/spin/data_and_results_for_rsol.odt_1/';
end
if SET == 8
indx_= {
{'~/hs/spot42_Eco.rf';'igrs_neis_gonoATCC49226/';'NeisgonoATCC49226_Dh/';
'';0}
{'~/hs/spot42_Eco.rf';'igrs_neis_meninATCC13091/';'NeismeninATCC13091_Dh/
';'';0}
{'~/hs/spot42_Eco.rf';'igrs_neis_zoo/';'Neiszoo_Dh/';'';0}
{'~/hs/spot42_Eco.rf';'igrs_neis_muco/';'Neismuco_Dh/';'';0}
{'~/hs/spot42_Eco.rf';'igrs_neis_cine/';'Neiscine_Dh/';'';0}
{'~/hs/spot42_Eco.rf';'igrs_neis_subfATCC49275/';'NeissubfATCC49275_Dh/';
'';0}
{'~/hs/spot42_Eco.rf';'igrs_neis_lact/';'Neislact_Dh/';'';0}
{'~/hs/spot42_Eco.rf';'igrs_neis_baciDSM23338/';'NeisbaciDSM23338_Dh/';''
;0}
{'~/hs/spot42_Eco.rf';'igrs_neis_flav/';'Neisflav_Dh/';'';0}
{'~/hs/spot42_Eco.rf';'igrs_neis_macaATCC33926/';'Neis_macaATCC33926_Dh/'
;'';0}
{'~/hs/spot42_Eco.rf';'igrs_neis_siccDSM17713/';'NeissiccDSM17713_Dh/';''
;0}
{'~/hs/spot42_Eco.rf';'igrs_neis_elo/';'Neiselo_Dh/';'';0}
```

```
{'~/hs/spot42_Eco.rf';'igrs_neis_wea/';'Neiswea_Dh/';'';0}
{'~/hs/spot42_Eco.rf';'igrs_neis_shay/';'Neisshay_Dh/';'';0}
{'~/hs/spot42_Eco.rf';'igrs_neis_dent/';'Neisdent_Dh/';'';0}
{'~/hs/spot42_Eco.rf';'igrs_neis_poly/';'Neispoly_Dh/';'';0}
{'~/hs/spot42_Eco.rf';'igrs_neis_berg/';'Neisberg_Dh/';'';0}
{'~/hs/spot42_Eco.rf';'igrs_neis_can/';'Neiscan_Dh/';'';0}
{'~/hs/spot42_Eco.rf';'igrs_neis_bla/';'Neisbla_Dh/';'';0}
{'~/hs/spot42_Eco.rf';'igrs_neis_bene/';'Neisbene_Dh/';'';0}
{'~/hs/spot42_Eco.rf';'igrs_neis_uirg/';'Neisuirg_Dh/';'';0}
{'~/hs/spot42_Eco.rf';'igrs_neis_ora/';'Neisora_Dh/';'';0}
{'~/hs/spot42_Eco.rf';'igrs_neis_bass/';'Neisbass_Dh/';'';0}
{'~/hs/spot42_Eco.rf';'igrs_neis_muscu/';'Neismuscu_Dh/';'';0}
{'~/hs/spot42_Eco.rf';'igrs_neis_maig/';'Neismaig_Dh/';'';0}
{'~/hs/spot42_Eco.rf';'igrs_neis_animalo/';'Neisanim_Dh/';'';0}
{'~/hs/spot42_Eco.rf';'igrs_neis_viri/';'Neisviri_Dh/';'';0}
{'~/hs/spot42_Eco.rf';'igrs_neis_leo/';'Neisleo_Dh/';'';0}
{'~/hs/spot42_Eco.rf';'igrs_neis_chenw/';'Neischenw_Dh/';'';0}
{'~/hs/spot42_Eco.rf';'igrs_neis_perf/';'Neisperf_Dh/';'';0}
{'~/hs/spot42_Eco.rf';'igrs_neis_wei/';'Neiswei_Dh/';'';0}
{'~/hs/spot42_Eco.rf';'igrs_neis_HMSC31F04/';'NeisHMSC31F04_Dh/';'';0}
{'~/hs/spot42_Eco.rf';'igrs_neis_P0008/';'NeisP0008_Dh/';'';0}

{'~/hs/spot42_Eco.rf';'igrs_neis_elo_C2010010207/';'NeiseloC2010010207_Dh
/';'';0}
{'~/hs/spot42_Eco.rf';'igrs_neis_per_CCUG17915/';'NeisperCCUG17915_Dh/';'
';0}
{'~/hs/spot42_Eco.rf';'igrs_neis_leo_CCUG45853/';'NeisleoCCUG45853_Dh/';'
';0}
{'~/hs/spot42_Eco.rf';'igrs_neis_gon_FF05/';'NeisgonFF05_Dh/';'';0}
};
d___ ='';
end
if SET == 11
      indx_= {
      {'~/hs/spot42_Eco.rf';'igrs_Idiomarina_aquatica/';'Idiomarinaaquati
ca_Dh/';'';0}
      {'~/hs/spot42_Eco.rf';'igrs_Idiomarina_abyssalis/';'Idiomarinaabyss
alis_Dh/';'';0}
      {'~/hs/spot42_Eco.rf';'igrs_Idiomarina_xiamenensis/';'Idiomarinaxia
menensis_Dh/';'';0}
      {'~/hs/spot42_Eco.rf';'igrs_Idiomarina_piscisalsi/';'Idiomarinapisc
isalsi_Dh/';'';0}
      {'~/hs/spot42_Eco.rf';'igrs_Idiomarina_baltica/';'Idiomarinabaltica
_Dh/';'';0}

      {'~/hs/spot42_Eco.rf';'igrs_Idiomarina_zobellii/';'Idiomarinazobell
ii_Dh/';'';0}
      {'~/hs/spot42_Eco.rf';'igrs_Idiomarina_tyrosinivorans/';'Idiomarina
tyrosinivorans_Dh/';'';0}
      {'~/hs/spot42_Eco.rf';'igrs_Idiomarina_fontislapidosi/';'Idiomarina
fontislapidosi_Dh/';'';0}
      {'~/hs/spot42_Eco.rf';'igrs_Idiomarina_ramblicola/';'Idiomarinaramb
licola_Dh/';'';0}
      {'~/hs/spot42_Eco.rf';'igrs_Idiomarina_A28L/';'IdiomarinaA28L_Dh/';
'';0}

      {'~/hs/spot42_Eco.rf';'igrs_Idiomarina_MD25a/';'IdiomarinaMD25a_Dh/
';'';0}
```

```
      {'~/hs/spot42_Eco.rf';'igrs_Idiomarina_28-8/';'Idiomarina28-
8_Dh/';'';0}
      {'~/hs/spot42_Eco.rf';'igrs_Idiomarina_loihiensis/';'Idiomarinaloih
iensis_Dh/';'';0}
      {'~/hs/spot42_Eco.rf';'igrs_Idiomarina_seosinensis/';'Idiomarinaseo
sinensis_Dh/';'';0}
      {'~/hs/spot42_Eco.rf';'igrs_Idiomarina_rhizosphaerae/';'Idiomarinar
hizosphaerae_Dh/';'';0}

      {'~/hs/spot42_Eco.rf';'igrs_Idiomarina_aminovorans/';'Idiomarinaami
novorans_Dh/';'';0}
      };
      d___ = '';
end

DE_NOVO = 0; STAT =1;BOOTSTRAP=10;MAIN_FIGURES=0; VERBOSE = 0;
reply = 'n';
if exist('INDEXES', 'var')
      if ~isempty(INDEXES)
            v=f_un_gr_ve(INDEXES);
            s=[];for ii=1:length(v),if length(v{ii}) > 1,s=[s
num2str(v{ii}(1)) ':' num2str(v{ii}(length(v{ii}))) ' '];else,s=[s
num2str(v{ii}) ' '];end,end,s(length(s)) = [];
      end
      reply = input(['INDEXES = ' s '. OK? [y]/n, or write space
separated species indexes.'], 's');
end
if ~exist('INDEXES', 'var') | reply == 'n'
      reply = input(['Type space separated species indexes and press
ENTER. Only ENTER for 1 : length(indx_) = 1:' num2str(length(indx_))
'.'], 's');
      if isempty(reply)
            INDEXES = 1 : length(indx_);
      else
            INDEXES = str2num(reply);
      end
else
      if ~isempty(reply) & reply ~= 'y'
            INDEXES = str2num(reply);
      end
end
v=f_un_gr_ve(INDEXES);
s=[];for ii=1:length(v),if length(v{ii}) > 1,s=[s num2str(v{ii}(1)) ':'
num2str(v{ii}(length(v{ii}))) ' '];else,s=[s num2str(v{ii}) '
'];end,end,s(length(s)) = [];
disp(['INDEXES = ' s '.'])

if ~exist('GOON'), GOON=0; end, disp(['GOON=' num2str(GOON) '.'])
if ~GOON
      if ~isempty(indx_{INDEXES(1)}{4})
            sigt_scores = zeros(3, length(INDEXES));
            counts_scores = sigt_scores;
            sigt_distances = zeros(11, length(INDEXES)); counts_distances
= sigt_distances;
      end
      igrs_nos = zeros(1, length(INDEXES));
      ps1_ = cell(1, length(INDEXES)); ps2_ = ps1_; ps3_ = ps1_; ps4_ =
ps1_; ps5_ = ps1_; ps6_ = ps1_; ps7_ = ps1_; ps8_ = ps1_;
```

```
        ps9_ = ps1_; ps10_ = ps1_;ps11_ = ps1_;jc_ = ps1_; jsc_ = ps1_; j3_
= ps1_;
        if READ_MATS
                d_best_sopt__all_means_2_best_mean_ = ps1_;
d_best_sopt__mean_of_bests_2_best_ = ps1_;d_best_sopt__best_means_ =
ps1_;
                d_best_sopt__bests_ = ps1_; d_best_sopt__3_bests_ = ps1_;
d_best_set__all_means_2_best_mean_ = ps1_; d_best_set__best_means_ =
ps1_;
                min_c_ = ps1_;   min_sc_ = ps1_; min_3_ = ps1_;
d_best_set__3_best_means_2_best_mean_ = ps1_;
                d_best_sopt__worst_2_best_= ps1_; d_best_set__worst_2_best_=
ps1_; d_best_sopt__3_best_means_2_best_mean_ = ps1_;
        end
end

for iI = 1 : length(INDEXES)
        disp(' ')
        if iI <= length(indx_)
        if ~isempty(indx_{INDEXES(iI)}{1}) &
~isempty(indx_{INDEXES(iI)}{2}) & ~isempty(indx_{INDEXES(iI)}{3})
                disp(['Doing ' indx_{INDEXES(iI)}{1} ', '
indx_{INDEXES(iI)}{2} ' in ' indx_{INDEXES(iI)}{3} '.'])
                input_fns{2} = [d___ indx_{INDEXES(iI)}{2}];
                input_fns{1} = indx_{INDEXES(iI)}{1};
                CUSTOM_SCANNING_LENGTH = indx_{INDEXES(iI)}{5};
                MATDIR = [d___ indx_{INDEXES(iI)}{3}];
                if ~READ_MATS
% this is for recomputing with different ALPHAA without the need to read
*.mat
                        igrsno = igrs_nos(INDEXES(iI));

        d_best_sopt__all_means_2_best_mean=d_best_sopt__all_means_2_best_me
an_{INDEXES(iI)};

        d_best_sopt__mean_of_bests_2_best=d_best_sopt__mean_of_bests_2_best
_{INDEXES(iI)};

        d_best_sopt__best_means=d_best_sopt__best_means_{INDEXES(iI)};
                        d_best_sopt__bests=d_best_sopt__bests_{INDEXES(iI)};
                        d_best_sopt__3_bests=d_best_sopt__3_bests_{INDEXES(iI)};

        d_best_set__all_means_2_best_mean=d_best_set__all_means_2_best_mean
_{INDEXES(iI)};

        d_best_set__best_means=d_best_set__best_means_{INDEXES(iI)};
                        d_best_set__3_best_means_2_best_mean =
d_best_set__3_best_means_2_best_mean_{INDEXES(iI)};
                        d_best_sopt__worst_2_best =
d_best_sopt__worst_2_best_{INDEXES(iI)};
                        d_best_set__worst_2_best =
d_best_set__worst_2_best_{INDEXES(iI)};
                        d_best_sopt__3_best_means_2_best_mean =
d_best_sopt__3_best_means_2_best_mean_{INDEXES(iI)};
                        min_c=min_c_{INDEXES(iI)};
                        min_sc=min_sc_{INDEXES(iI)};
                        min_3=min_3_{INDEXES(iI)};
                        run_rsol_IGRs_complete
```

```
        else
              disp(['Reading .mat of ' indx_{INDEXES(iI)}{2} ' and '
indx_{INDEXES(iI)}{3} ' in ' d___ ' .' ]);
              if exist([d___ indx_{INDEXES(iI)}{2}], 'file') &
exist([d___ indx_{INDEXES(iI)}{3}], 'file')
                    disp('Dirs OK.')
                    run_rsol_IGRs_complete
                    igrs_nos(INDEXES(iI)) = igrsno;
                    d_best_sopt__all_means_2_best_mean_{INDEXES(iI)} =
d_best_sopt__all_means_2_best_mean;
                    d_best_sopt__mean_of_bests_2_best_{INDEXES(iI)} =
d_best_sopt__mean_of_bests_2_best;
                    d_best_sopt__best_means_{INDEXES(iI)} =
d_best_sopt__best_means;
                    d_best_sopt__bests_{INDEXES(iI)} =
d_best_sopt__bests;
                    d_best_sopt__3_bests_{INDEXES(iI)} =
d_best_sopt__3_bests;
                    d_best_set__all_means_2_best_mean_{INDEXES(iI)} =
d_best_set__all_means_2_best_mean;
                    d_best_set__best_means_{INDEXES(iI)} =
d_best_set__best_means;
                    d_best_set__3_best_means_2_best_mean_{INDEXES(iI)}
= d_best_set__3_best_means_2_best_mean;
                    d_best_sopt__worst_2_best_{INDEXES(iI)} =
d_best_sopt__worst_2_best;
                    d_best_set__worst_2_best_{INDEXES(iI)} =
d_best_set__worst_2_best;

      d_best_sopt__3_best_means_2_best_mean_{INDEXES(iI)} =
d_best_sopt__3_best_means_2_best_mean;
                    min_c_{INDEXES(iI)} = min_c;
                    min_sc_{INDEXES(iI)} = min_sc;
                    min_3_{INDEXES(iI)} = min_3;
              else
                    disp('Dirs not OK. Stop.')
                    return
              end
        end

        jc_{INDEXES(iI)} = jc; jsc_{INDEXES(iI)} = jsc;
j3_{INDEXES(iI)} = j3;
        ps1_{INDEXES(iI)} = ps1; ps2_{INDEXES(iI)} = ps2;
ps3_{INDEXES(iI)} = ps3;ps4_{INDEXES(iI)} = ps4;ps5_{INDEXES(iI)} = ps5;
ps6_{INDEXES(iI)} = ps6;
        ps7_{INDEXES(iI)} = ps7; ps8_{INDEXES(iI)} = ps8;
ps9_{INDEXES(iI)} = ps9; ps10_{INDEXES(iI)} = ps10; ps11_{INDEXES(iI)} =
ps11;

        if ~isempty(indx_{INDEXES(iI)}{4})
              sigt_scores(:, INDEXES(iI)) = 0; counts_scores(:,
INDEXES(iI)) = 0; sigt_distances(:, INDEXES(iI)) = 0;counts_distances(:,
INDEXES(iI)) = 0;

              n = str2num(indx_{INDEXES(iI)}{4});
              if rem(n,2)
                    n1 = n + 1;
              else
```

```
                    n1 = n-1;
                end

                if ~isempty([find(jc == n) find(jc == n1)])
                    sigt_scores(1, INDEXES(iI)) = min([find(jc == n)
find(jc == n1)]);
                end
                if ~isempty([find(jsc == n) find(jsc == n1)])
                    sigt_scores(2, INDEXES(iI)) = min([find(jsc == n)
find(jsc == n1)]);
                end
                if ~isempty([find(j3 == n) find(j3 == n1)])
                    sigt_scores(3, INDEXES(iI)) = min([find(j3 == n)
find(j3 == n1)]);
                end
                counts_scores(:, INDEXES(iI)) = [length(jc) length(jsc)
length(j3)];

                if ~isempty([find(ps1 == n) find(ps1 == n1)] )
                    sigt_distances(1,  INDEXES(iI)) = min([find(ps1 ==
n) find(ps1 == n1)]);
                end
                if ~isempty([find(ps2 == n) find(ps2 == n1)] )
                    sigt_distances(2,  INDEXES(iI)) = min([find(ps2 ==
n) find(ps2 == n1)]);
                end
                if ~isempty([find(ps3 == n) find(ps3 == n1)] )
                    sigt_distances(3,  INDEXES(iI)) = min([find(ps3 ==
n) find(ps3 == n1)]);
                end
                if ~isempty([find(ps4 == n) find(ps4 == n1)])
                    sigt_distances(4,  INDEXES(iI)) = min([find(ps4 ==
n) find(ps4 == n1)]);
                end
                if ~isempty([find(ps5 == n) find(ps5 == n1)] )
                    sigt_distances(5,  INDEXES(iI)) = min([find(ps5 ==
n) find(ps5 == n1)]);
                end
                if ~isempty([find(ps6 == n) find(ps6 == n1)] )
                    sigt_distances(6,  INDEXES(iI)) = min([find(ps6 ==
n) find(ps6 == n1)]);
                end
                if ~isempty([find(ps7 == n) find(ps7 == n1)])
                    sigt_distances(7,  INDEXES(iI)) = min([find(ps7 ==
n) find(ps7 == n1)]);
                end
                if ~isempty([find(ps8 == n) find(ps8 == n1)])
                    sigt_distances(8,  INDEXES(iI)) = min([find(ps8 ==
n) find(ps8 == n1)]);
                end
                if ~isempty([find(ps9 == n) find(ps9 == n1)])
                    sigt_distances(9,  INDEXES(iI)) = min([find(ps9 ==
n) find(ps9 == n1)]);
                end
                if ~isempty([find(ps10 == n) find(ps10 == n1)])
                    sigt_distances(10,  INDEXES(iI)) = min([find(ps10
== n) find(ps10 == n1)]);
                end
```

```
                 if ~isempty([find(ps11 == n) find(ps11 == n1)])
                      sigt_distances(11,  INDEXES(iI)) = min([find(ps11
== n) find(ps11 == n1)]);
                 end
                 counts_distances(:, INDEXES(iI)) = [length(ps1)
length(ps2) length(ps3) length(ps4) length(ps5) length(ps6) length(ps7)
length(ps8) length(ps9) length(ps10) length(ps11)];
           end
      end
      end
end

clear ALPHA__                             d_best_set__best_means
m5       ps11 ps10                          ps1
clear BOOTSTRAP                           d_best_sopt__3_bests
m6                                  ps7
clear CUSTOM_SCANNING_LENGTH
d_best_sopt__all_means_2_best_mean  m7
ps2
clear DE_NOVO                             d_best_sopt__best_means
min_c                               ps3
clear DO_3_END_U                          d_best_sopt__bests
min_c_i                             psc_
clear DO_ALTERNATIVE_CANDIDATES a  x j i
d_best_sopt__mean_of_bests_2_best   min_sc                           r
clear GOON                                fn_best_sopts_
n n1                                   sc_
clear fnsqs__                             notexistingmatsno
sc_rand_set_
clear ISQS                                fntm__
notuniquematsno                     sci_
clear MAIN_FIGURES d_best_set__3_best_means_2_best_mean iI o1 scs_
clear MATDIR                              igrsno
o2                                  scs_rand_set_
clear SAMPLINGLENGTH                      iigrsno
o3                                  scs_rand_sopt_
clear STAT d_best_sopt__best_2_worst o4                  jcx jscx
clear VERBOSE                             input_fns
o5
clear c_                                  jc
o6                                  sl
clear c_17_20                             jcsc      jsc3
o7                                  t
clear c_2_5_8_11_14                       jsc j3
pc_                                 u
clear c_rand_sopt_                        m1
uniqueexistingmatsno   reply
clear  m2                                  ps4
v
clear m3                                  ps5        ps8 ps9 ax reply cx
scx scs__ min_3 d_best_sopt__worst_2_best
d_best_sopt__3_best_means_2_best_mean
clear d_best_set__all_means_2_best_mean   m4           s        ii
ps6         foldermnt d___ input_fns reply d_best_set__worst_2_best


f_un_gr_ve.m
```

```
function ves = f_un_gr_ve(ve)
% find uniterupted growing vectors (ves) in vector ve

i = 1;
ves = {};
while i <= length(ve)
  ist = i;
  while i < length(ve) & ve(i+1) == ve(i)+1, i = i + 1; end
  ves = [ves {ve(ist:i)}];
  i = i + 1;
end
```

run_rsol_IGRs_complete.m

```
% input_fns = {'~/hs/spot42_Eco.rf'; '~/rsol/igrs_eco';};
% input_fns = {'~/hs/spot42_Eco.rf'; '~/rsol/igrs_glacpuni';};

if ~exist('ALPHAA'), ALPHAA = .05; end
if ~exist('VERBOSE'), VERBOSE = 1; end
if ~exist('READ_MATS'), READ_MATS = 1; end
if ~exist('BOOTSTRAP'), BOOTSTRAP = 10; end
if input_fns{2}(length(input_fns{2})) == '/',
input_fns{2}(length(input_fns{2})) = []; end
if exist('CUSTOM_SCANNING_LENGTH')
      if ~CUSTOM_SCANNING_LENGTH
            if exist(input_fns{1})
                  [sl sl] = unix(['head -n2 ' input_fns{1} ' | tail -n1 |
wc -c']); sl = num2str(str2num(sl) - 1);
            else
                  disp([input_fns{1} ' not exist. Stop.']), return
            end
      else
            sl = num2str(CUSTOM_SCANNING_LENGTH);
      end
else
      disp('CUSTOM_SCANNING_LENGTH must exist  [0 for template length |
custom length]. Stop.'), return
end

if DE_NOVO
      for iIGRS = 1 : length(NO_DE_NOVO)
            fn = dir([input_fns{2} '/*_igr' num2str(NO_DE_NOVO(iIGRS))
'.fa']);
            if ~isempty(fn) | length(fn) == 1
                  unix(['rsol -sqs=' input_fns{2} '/' fn.name ' -tm='
input_fns{1} ' -DO_ALTERNATIVE_CANDIDATES=0 -
NUMBER_OF_BEST_SETS_TO_SHOW=0 -BOOTSTRAP=' ...
                           num2str(BOOTSTRAP) ' -HISTOGRAM=0 -
GRAPHICAL_OUTPUT=0 -KEEP_DBS=1 -SAMPLINGLENGTH=' sl ' -DIST_LETT=h'])
% -MAXSTRSPERSQ=100 -WINDOWSIZE=-5 -PROCENTFE=20': a lot of irgs was
refused by the Unafold, incl. the one with glac33A spot42
% -MAXSTRSPERSQ=100 -WINDOWSIZE=-10 -PROCENTFE=2000': spot42 glac33A was
not identified by jc, jsc, j3.
% -MAXSTRSPERSQ=60 -WINDOWSIZE=10 -PROCENTFE=2000': more FPs in jc, jsc,
j3. spot42 glac33A was not identified by all, HTCC was by all.
```

```
% -MAXSTRSPERSQ=100 -WINDOWSIZE=10 -PROCENTFE=20: spot42 glac33A was not
identified by all, but less FPs in jc, jsc, j3. More FPs for HTCC, spot42
found. In MH2013,
% found in all, less FPs. In siph and KUL10 not found.
% MAXSTRSPERSQ=100 -WINDOWSIZE=5 -PROCENTFE=20: More FPs for HTCC, spot42
found in all. In glac33A, found by jc, but with more FPs.
%                 unix(['rsol -sqs=' input_fns{2} '/' fn.name ' -tm='
input_fns{1} ...
%                         ' -DO_ALTERNATIVE_CANDIDATES=0 -
NUMBER_OF_BEST_SETS_TO_SHOW=0 -BOOTSTRAP=0 -HISTOGRAM=0 -
GRAPHICAL_OUTPUT=0 -KEEP_DBS=1 -SAMPLINGLENGTH=' sl ...
%                         ' -DIST_LETT=h -MAXSTRSPERSQ=100 -WINDOWSIZE=-5
-PROCENTFE=20'])
           else
                 disp(['Either not exist or more than 1 ' input_fns{2}
'/*_igr' num2str(NO_DE_NOVO(iIGRS)) '.fa. Ignoring. Go to next subject
sequence.'])
           end
      end
end
% evaluate ALREADY COMPUTED *.mat for randomly chosen genomic gene/IGR
sequences
if STAT
      if length(MATDIR) & MATDIR(length(MATDIR)) ~= '/', MATDIR = [MATDIR
'/']; end;
      if READ_MATS
           disp(['Taking *.mat from ' MATDIR '.'])
% ISQS is an index of subject sq if there are more than one subject sqs,
which I do not count on in rsol.
           ISQS = 1;
           tmnoslash = strsplit(input_fns{1}, '/');
           tmnoslash = tmnoslash{length(tmnoslash)};
           t = strsplit(tmnoslash, '_');
           disp([t{1} ': '])

           if ~exist(input_fns{2}, 'dir'), disp([input_fns{2} ' does not
exist. Stop.']), return, end
           [igrsno igrsno] = unix(['ls ' input_fns{2} '/|wc -l']); igrsno
= str2num(igrsno);
           disp(['There should be ' num2str(igrsno) ' mats in ' MATDIR
'.'])

           if BOOTSTRAP
% ssq chosen acc to best-at-all sopt
% distances between single sopts
                 d_best_sopt__bests = NaN(size(ones(1, igrsno)));
                 d_best_sopt__3_bests = d_best_sopt__bests;
                 d_best_sopt__mean_of_bests_2_best = d_best_sopt__bests;
% distance between the mean of the set with best-at-all sopt and the mean
of the randomized set with best mean
                 d_best_sopt__best_means = d_best_sopt__bests;
                 d_best_sopt__all_means_2_best_mean = d_best_sopt__bests;
% distance between the mean of the set with best-at-all sopt and mean of
means of three randomized sets with three best means
                 d_best_sopt__3_best_means_2_best_mean =
d_best_sopt__bests;

                 d_best_sopt__worst_2_best = d_best_sopt__bests;
```

```
% ssq chosen acc to best mean
% distance between the mean of the set with best mean and the mean of the
randomized set with best mean
                d_best_set__best_means = d_best_sopt__bests;
% distance between the mean of the set with best mean and the mean of
means of all randomized sets
                d_best_set__all_means_2_best_mean = d_best_sopt__bests;
                d_best_set__3_best_means_2_best_mean =
d_best_sopt__bests;

                d_best_set__worst_2_best = d_best_sopt__bests;
          end
          min_c = NaN(size(ones(1, igrsno))); min_sc = min_c; min_3 =
min_c;

          uniqueexistingmatsno = 0; notexistingmatsno = 0;
notuniquematsno = 0;
% these are for testing purposes against all cs (min raw TEDs) and scs
(evaluated raw TEDs acc. to SCORINGSCHEME in
f_sample_sq_by_sopts_to_tm__no_PDF.m, i.e. mean)
%         scx = [];cx=[];
% this is for analytic purposes in xFPsTP.m
          %scs__ = cell(1, igrsno);
          for iigrsno = 1 : igrsno
                disp(['Reading ' num2str(iigrsno) '/' num2str(igrsno)
'.'])
                [x x] = unix(['ls ' MATDIR 'rsol_*_igr' num2str(iigrsno)
'.fa__' tmnoslash '__' sl '.mat']);
                if isempty(strfind(x, 'No such file or directory'))
     % check if it is an unique .mat and that there are not 2 same .mats
from different processes
                      if sum(x == char(10)) > 1
                            disp(['More than 1 ' MATDIR 'rsol_*_igr'
num2str(iigrsno) '.fa__' tmnoslash '__' sl '.mat. Computed by different
processes? Check. Now taking first of them.'])
     %                      return
                            notuniquematsno = notuniquematsno + 1;
                            i = find(x == char(10));
                            x(i + 1 : length(x)) = [];
                      end
     %                disp(['Doing ' num2str(iigrsno) '/'
num2str(igrsno) ' .mat in ' input_fns{2} '/.'])
                      uniqueexistingmatsno = uniqueexistingmatsno + 1;
                      x(length(x)) = [];
                      load(x)

%                     for i=1:length(scs_{ISQS})
%                           cx = [cx scs_{ISQS}{i}];
%                           scx = [scx mean(scs_{ISQS}{i})];
%                     end
%                     scs__{iigrsno} = scs_{ISQS};

                      min_sc(iigrsno) = sc_{ISQS}(sci_{ISQS}(1));
                      min_c(iigrsno) = min(c_{ISQS});

                      a = zeros(1, length(scs_{ISQS}));
                      for i=1:length(scs_{ISQS})
```

```
                        if length(scs_{ISQS}{i}) < 3
                             a(i) = mean(scs_{ISQS}{i});
                        else
                             b = sort(scs_{ISQS}{i});
                             a(i) = mean(b(1:3));
                        end
                   end
                   min_3(iigrsno) = min(a);

                   if BOOTSTRAP
                        [min_c_i min_c_i] = min(c_{ISQS});
                        a = sort(scs_{ISQS}{min_c_i});
                        u = []; for i = 1 : BOOTSTRAP, u = [u
min(scs_rand_sopt_{ISQS}{i})]; end
                        u = sort(u);
                        d_best_sopt__bests(iigrsno) = u(1) - a(1);
                        d_best_sopt__3_bests(iigrsno) = mean(u(1:3))
- mean(a(1:3));
                        d_best_sopt__mean_of_bests_2_best(iigrsno) =
mean(u) - a(1);
                        d_best_sopt__worst_2_best(iigrsno) =
mean(scs_rand_sopt_{ISQS}{BOOTSTRAP}) - mean(a);
                        r = [];for j = 1 : BOOTSTRAP, r = [r
mean(scs_rand_sopt_{ISQS}{j})]; end, r = sort(r);
                        d_best_sopt__best_means(iigrsno) = r(1) -
mean(a);
                        d_best_sopt__all_means_2_best_mean(iigrsno) =
mean(r - mean(a));

      d_best_sopt__3_best_means_2_best_mean(iigrsno) = mean(r(1 : 3)) -
mean(a);

                        v = [];for i = 1 : BOOTSTRAP, v = [v
mean(scs_rand_set_{ISQS}{i})]; end
                        v = sort(v);
                        d_best_set__best_means(iigrsno) = v(1) -
mean(scs_{ISQS}{sci_{ISQS}(1)});
                        d_best_set__all_means_2_best_mean(iigrsno) =
mean(v - mean(scs_{ISQS}{sci_{ISQS}(1)}));
                        d_best_set__3_best_means_2_best_mean(iigrsno)
= mean(v(1 : 3) - mean(scs_{ISQS}{sci_{ISQS}(1)}));

                        d_best_set__worst_2_best(iigrsno) =
v(BOOTSTRAP) - mean(scs_{ISQS}{sci_{ISQS}(1)});
                   end
              else
                   disp([MATDIR 'rsol_*_igr' num2str(iigrsno) '.fa__'
tmnoslash '__' sl '.mat not exist. Pausing. After key pressed, go to
next.']), pause
                   notexistingmatsno = notexistingmatsno + 1;
              end
         end

         if uniqueexistingmatsno ~= igrsno | notexistingmatsno |
notuniquematsno
              disp(['uniqueexistingmatsno ('
num2str(uniqueexistingmatsno) ')~= igrsno (' num2str(igrsno) ').'])
```

```
                disp(['notexistingmatsno = ' num2str(notexistingmatsno)
'.'])
                disp(['notuniquematsno = ' num2str(notuniquematsno)
'.'])
      %               return
      %         uniqueexistingmatsno = 0; notexistingmatsno = 0;
notuniquematsno = 0;
            end
      end
% HERE YOU COMPARE DISTANCES BETWEEN BEST SCORES OF REGULAR AND
RANDOMIZED SEQUENCES FOR TP AND TNS (= RANDOMLY CHOSEN IGR SQS IN THE
GENOME)
      ps_ = ones(length(igrsno), 14);
      if BOOTSTRAP
            for ia = 1 : length(d_best_sopt__all_means_2_best_mean)
                  [h pr] = ttest2(d_best_sopt__all_means_2_best_mean(ia),
d_best_sopt__all_means_2_best_mean([1 : ia - 1 ia + 1 :
length(d_best_sopt__all_means_2_best_mean)]), 'Alpha',ALPHAA,
'Tail','right');
                  ps_(ia, 5) = round(pr*1000)/1000;
            end
            for ia = 1 : length(d_best_sopt__mean_of_bests_2_best)
                  [h pr] = ttest2(d_best_sopt__mean_of_bests_2_best(ia),
d_best_sopt__mean_of_bests_2_best([1 : ia - 1 ia + 1 :
length(d_best_sopt__mean_of_bests_2_best)]), 'Alpha',ALPHAA,
'Tail','right');
                  ps_(ia, 3) = round(pr*1000)/1000;
            end
            for ia = 1 : length(d_best_sopt__best_means)
                  [h pr] = ttest2(d_best_sopt__best_means(ia),
d_best_sopt__best_means([1 : ia - 1 ia + 1 :
length(d_best_sopt__best_means)]), 'Alpha',ALPHAA, 'Tail','right');
                  ps_(ia, 4) = round(pr*1000)/1000 ;
            end
            for ia = 1 : length(d_best_sopt__bests)
                  [h pr] = ttest2(d_best_sopt__bests(ia),
d_best_sopt__bests([1 : ia - 1 ia + 1 : length(d_best_sopt__bests)]),
'Alpha',ALPHAA, 'Tail','right');
                  ps_(ia, 1) = round(pr*1000)/1000;
            end
            for ia = 1 : length(d_best_sopt__3_bests)
                  [h pr] = ttest2(d_best_sopt__3_bests(ia),
d_best_sopt__3_bests([1 : ia - 1 ia + 1 : length(d_best_sopt__3_bests)]),
'Alpha',ALPHAA, 'Tail','right');
                  ps_(ia, 2) = round(pr*1000)/1000;
            end
            for ia = 1 : length(d_best_sopt__worst_2_best)
                  [h pr] = ttest2(d_best_sopt__worst_2_best(ia),
d_best_sopt__worst_2_best([1 : ia - 1 ia + 1 :
length(d_best_sopt__worst_2_best)]), 'Alpha',ALPHAA, 'Tail','right');
                  ps_(ia, 7) = round(pr*1000)/1000;
            end
            for ia = 1 : length(d_best_sopt__3_best_means_2_best_mean)
                  [h pr] =
ttest2(d_best_sopt__3_best_means_2_best_mean(ia),
d_best_sopt__3_best_means_2_best_mean([1 : ia - 1 ia + 1 :
length(d_best_sopt__3_best_means_2_best_mean)]), 'Alpha',ALPHAA,
'Tail','right');
```

```
                ps_(ia, 6) = round(pr*1000)/1000;
            end
            ps1 = find(ps_(:, 1) <= ALPHAA); ps2 = find(ps_(:, 2) <=
ALPHAA); ps3 = find(ps_(:, 3) <= ALPHAA); ps4 = find(ps_(:, 4) <=
ALPHAA);
            ps5 = find(ps_(:, 5) <= ALPHAA); ps6 = find(ps_(:, 6) <=
ALPHAA); ps7 = find(ps_(:, 7) <= ALPHAA);

            for ia = 1 : length(d_best_set__all_means_2_best_mean)
                [h pr] = ttest2(d_best_set__all_means_2_best_mean(ia),
d_best_set__all_means_2_best_mean([1 : ia - 1 ia + 1 :
length(d_best_set__all_means_2_best_mean)]), 'Alpha',ALPHAA,
'Tail','right');
                ps_(ia, 9) = round(pr*1000)/1000;
            end
            for ia = 1 : length(d_best_set__best_means)
                [h pr] = ttest2(d_best_set__best_means(ia),
d_best_set__best_means([1 : ia - 1 ia + 1 :
length(d_best_set__best_means)]), 'Alpha',ALPHAA, 'Tail','right');
                ps_(ia, 8) = round(pr*1000)/1000;
            end
            for ia = 1 : length(d_best_set__3_best_means_2_best_mean)
                [h pr] =
ttest2(d_best_set__3_best_means_2_best_mean(ia),
d_best_set__3_best_means_2_best_mean([1 : ia - 1 ia + 1 :
length(d_best_set__3_best_means_2_best_mean)]), 'Alpha',ALPHAA,
'Tail','right');
                ps_(ia, 10) = round(pr*1000)/1000 ;
            end
            for ia = 1 : length(d_best_set__worst_2_best)
                [h pr] = ttest2(d_best_set__worst_2_best(ia),
d_best_set__worst_2_best([1 : ia - 1 ia + 1 :
length(d_best_set__worst_2_best)]), 'Alpha',ALPHAA, 'Tail','right');
                ps_(ia, 11) = round(pr*1000)/1000 ;
            end
            ps8 = find(ps_(:, 8) <= ALPHAA); ps9 = find(ps_(:, 9) <=
ALPHAA); ps10= find(ps_(:, 10) <= ALPHAA);ps11= find(ps_(:, 11) <=
ALPHAA);
        end
        for ia = 1 : length(min_c)
            [h pl] = ttest2(min_c(ia), min_c([1 : ia - 1 ia + 1 :
length(min_c)]), 'Alpha',ALPHAA, 'Tail','left');
            ps_(ia, 12) = round(pl*1000)/1000 ;
        end
        for ia = 1 : length(min_sc)
            [h pl] = ttest2(min_sc(ia), min_sc([1 : ia - 1 ia + 1 :
length(min_sc)]), 'Alpha',ALPHAA, 'Tail','left');
            ps_(ia, 13) = round(pl*1000)/1000 ;
        end
        for ia = 1 : length(min_3)
            [h pl] = ttest2(min_3(ia), min_3([1 : ia - 1 ia + 1 :
length(min_3)]), 'Alpha',ALPHAA, 'Tail','left');
            ps_(ia, 14) = round(pl*1000)/1000;
        end

        jc = find(ps_(:, 12) <= ALPHAA);
        [ic ic] = sort(min_c(jc), 'asce');
        jc = jc(ic);
```

```
      if VERBOSE, disp([newline 'Min c were significant (i.e. p<=' num2str(ALPHAA) ') for these IGRs (ascending c): ' num2str(jc') '.']);end

      jsc = find(ps_(:, 13) <= ALPHAA);
      [isc isc] = sort(min_sc(jsc), 'asce');
      jsc = jsc(isc);
      if VERBOSE, disp(['Min sc were significant (i.e. p<=' num2str(ALPHAA) ') for these IGRs (ascending sc): ' num2str(jsc') '.']);
end

      j3 = find(ps_(:, 14) <= ALPHAA);
      [i3 i3] = sort(min_3(j3), 'asce');
      j3 = j3(i3);
end
clear ALPHA__     iIGRS     i3           pl           ia
clear pr                         DO_3_END_U
fn_best_sopts_               psc_
clear DO_ALTERNATIVE_CANDIDATES    fn_pdf_                      r
vv vvno
clear ISQS                         fn_pdf_c_
clear fnsqs__                      sc_rand_set_
clear fntm__                      fn sci_
clear iigrsno                          h SAMPLINGLENGTH
hs_dist_best_set
clear hs_dist_best_sopt a                          i
sl
clear ans     t
clear c_                                             tmnoslash
ins q
clear c_rand_sopt_                 j                           u
clear min_c_i                      v                    x
p                   pc_
clear sc_ scs_ scs_rand_set_                  scs_rand_sopt_ ic isc  b
```

Synteny filtering

Scripts

syntenic_cands_among_all_cands_from_indexes.m:
-> f_get_synteny_from_fasta_comb.m -> f_find_all_words_of_str_in_str.m, f_make_header.m, f_clean_RNA_sequence.m, f_read_genome.m, f_read_anns.m, f_reconstruct_blast_hit.m, f_clean_fasta_headers.m, f_filter_fasta_by_ID.m.
-> f_try_anns.m:
   -> f_get_assembly_ID_from_ID_edirect.m, f_store_anns_simple_no_ONLYFEATURETABLE.m.

Matlab Code

syntenic_cands_among_all_cands_from_indexes.m

```
clear SWITCH INDEXES
ANNS_SA = '/mnt/spin/anns_Bacteria_sa';
SUFFIX =  '_Dh';
d = '/mnt/spin/data_and_results_for_rsol.odt_1/';
```

```
% LEGEND:
% IF pat IS A CELL, THEN ANY STRING OF IT MUST BE FOUND IN ANY GENOMIC
CONTEXT SIDE.
% IF pat IS TWO CELLS, THEN:
% IF CONTEXT = 2, THEN ANY STRING OF ONE OF THE CELLS IN pat MUST BE
FOUND IN ONE OF THE TWO GENOMIC CONTEXT SIDES, AND THEN ANY STRING OF THE
% OTHER CELL IN THE OTHER GENOMIC CONTEXT SIDE.
% IF CONTEXT = 1, THEN ANY STRING OF THE BOTH CELLS IN pat MUST BE FOUND
IN ONE OF THE TWO GENOMIC CONTEXT SIDES.
% THERE ALSO WORDS OF THE STRINGS AND/OR SUBWORDS CAN BE SEARCHED FOR
DEPENDING ON requireall AND allowsubstring PARAMS.

% genus='pseudoalterom';
% SWITCH = 'for_Idiomarina_Frankia_Pseudoalterom';
% gs={'igrs_pseudoalterom_halopl'; 'igrs_pseudoalterom_maricalo';
'igrs_pseudoalterom_atl'; 'igrs_pseudoalterom_pir';
'igrs_pseudoalterom_mar'; 'igrs_pseudoalterom_sim'};
% % SCORES=13; ns={[2278  1992  1957  1487  2394  1380   900   797   949
861  2653   815  2027  1378  2108  1730   299   238  1245  1416  1831
1832   155   233  1561   340  1518  2486  1513   923  1422  2245  2042
2366  2258  2494   394  1440  1404   636   175  2271  2338  1379     8
2305  2612  1189  1584    47   293   892   206   889  1102  1260   406
227  1171   668  1959  2299   440  1064  1697   446   702  1338  1306
2661   396   573  1755   943   609   785  1626   486  2263   184  1629
1588  1657  2480   950  1355  2289   784   250  1935  1774   772  2094
335   932  1401   825  1242  1818  1354   284  1129   899  2118   165
1199  1851   764  2516   853  2482  1029  1087  1053   904  2337  2088
1916  1145   688  1215  1566  2490  2378];[869   141    58   968  1549
652   422  1468   130   129  1618  1123  1787  1131   212   700   420
1100  2233   233  2302    55  1674   924  1711  2361  2373  1229  1710
260   846   636  1368   226  1431   432   721  1273     3   817   217
949  1381  1827  1075  1325   354   627  1695   184   439  1063  2428
463  1678  2028   325  2064  1137    50   358  1861  2125  1823  1231
823  2378  1903  2176  1534    89  1852   902  1539  1880   934  1647
622  1704  1945   634  1613  1857  1981  1083  1760  1150  2095  2478
1714   449  1483   107   234  1280  1354   763  2489  2256  2325   929
1119  1651  1367];[643  2209   101  1002   602   133  1837  2349   177
74  2155  1488  2063  2191  1537  1364  2176  1757   142   891  1655
135   183  1656   604   382  1104  1983   587  2249  1769   186   507
1505  1155  1883  1396    45   592   180  1957   801  1940  1172  1162
182  2261  2273  2156   200  1069  1808    88   676  1471   486  1531
848  1599   943  1022  1600   404   653  2193   284   143   344  2210
2062  2311  2300  1197  1458   642   620   852  1766   367   202  1185
977  1381  1004  1277  1306  1548   850  1592  1038   157   707   810
2000   130   428  2274  1857  1111   717   561  1629   794  1677  1495
2279];[1517   846   701   139  2475   442  2638  1006  2282  1664   482
2076  1252  1579  1226  1788  2190  1632  2367  1489  1357  2195  1054
2434    10   133    30  2463   864  1144  1379   883   210  1857  2442
347  1622   151  1124    86   321   494   513  1525  1462  1319   233
462  2080   113   982  2315    57  1057   514  1269   357  1878  1019
670  2598  1056   860  1446   729  1940   230   410  2697  2114   740
1598   584  1142   518  2128  1683   700  1465  2476   661   674   502
441  1702   369  2398   485  1040  1417  1580  1287    81  1234   141
1952  1423   365    97   436   556];[2144  1532  1258   121  1505  1349
163  1305   268   153   545  1051  1915  1805   104  1059    22   480
1987  2137  1946  1924   615  1684   784   301  2148  1070  1783  1099
1233   193  1367    24  2120  1655  2085  1493   372   264   604  1753
```

```
440   987  1067   831   483  1497    20   170  1135   782  2077   590
2044  1693  1706   458  1803  2005  2021   478   316   699  1201  2073
443  2139  2130   513  1836  1162  1025   204  1989  1097   297  1206
452  1618  1307   454   933    58   780    47   949   210  1458  1037
548   850   439  1967  1308  1323  1052   710  2123   300];[1966  2190
204  1537  1755  1419   644   611  1772   484   239  2189   781  1763
692  1959   597  2303  2514  2081  1284   488  1903   547  1363  1047
918   920  1780  1118   651  1978  2521  2506  1666  1809  2374  1875
765  2187  1278  2437    13   664  1600  2117  2459  2537   154  1493
1707   199  1932   536  1581  2481  1836  1743   864  1960   846   880
1732  2136  2370   554   105  2307  1762   432   595  1804   656  1988
1664   462  1128   594   835   854   314  1209  1705  1615  1371   539
2153  1593  2228   384  2080   257   842  1680  1472   313  2425   173
1268   743   879    12  1358  1893  2181  2074  2453   758   968  1468
2235  1101  2043  2015   928   472];};
% % % %  simplex had only one side synteny, an extra synteny filtering
must have been done for it with pat.
% % SCORES=13; gs={'igrs_pseudoalterom_sim'}; ns={[1966  2190   204  1537
1755  1419   644   611  1772   484   239  2189   781  1763   692  1959
597  2303  2514  2081  1284   488  1903   547  1363  1047   918   920
1780  1118   651  1978  2521  2506  1666  1809  2374  1875   765  2187
1278  2437    13   664  1600  2117  2459  2537   154  1493  1707   199
1932   536  1581  2481  1836  1743   864  1960   846   880  1732  2136
2370   554   105  2307  1762   432   595  1804   656  1988  1664   462
1128   594   835   854   314  1209  1705  1615  1371   539  2153  1593
2228   384  2080   257   842  1680  1472   313  2425   173  1268   743
879    12  1358  1893  2181  2074  2453   758   968  1468  2235  1101
2043  2015   928   472];};
% % % SCORES=9; gs={'igrs_pseudoalterom_pir';};ns={[10    30    57    77
81    82    86   103   113   133   139   151   210   230   233   235
237   242   246   249   261   263   321   344   348   365   429   441
442   450   460   462   474   482   493   502   514   518   543   554
568   592   668   670   699   700   701   707   726   729   823   846
864   883   892   913   942   982   998  1006  1019  1020  1040  1057
1088  1113  1143  1144  1170  1226  1234  1247  1252  1269  1287  1304
1319  1357  1404  1423  1424  1433  1434  1446  1448  1456  1462  1466
1467  1489  1508  1517  1578  1579  1580  1598  1614  1622  1632  1664
1702  1709  1749  1787  1788  1851  1857  2024  2076  2080  2111  2128
2195  2215  2282  2314  2315  2352  2367  2398  2434  2475  2522  2576
2598  2599  2624  2638  2670  2697  2717  2738];};
% SCORES=[13 9];ns={[2278  1992  1957  1487  2394  1380   900   797   949
861  2653   815  2027  1378  2108  1730   299   238  1245  1416  1831
1832   155   233  1561   340  1518  2486  1513   923  1422  2245  2042
2366  2258  2494   394  1440  1404   636   175  2271  2338  1379     8
2305  2612  1189  1584    47   293   892   206   889  1102  1260   406
227  1171   668  1959  2299   440  1064  1697   446   702  1338  1306
2661   396   573  1755   943   609   785  1626   486  2263   184  1629
1588  1657  2480   950  1355  2289   784   250  1935  1774   772  2094
335   932  1401   825  1242  1818  1354   284  1129   899  2118   165
1199  1851   764  2516   853  2482  1029  1087  1053   904  2337  2088
1916  1145   688  1215  1566  2490  2378     7    30    53   109   121
152   223   228   348   350   387   427   500   531   536   550   594
604   635   844   939   945  1146  1147  1319  1367  1450  1597  1667
1680  1688  1720  1907  1922  1963  2097  2099  2103  2120  2142  2204
2328  2370  2414  2454  2668  2669];[869   141    58   968  1549   652
422  1468   130   129  1618  1123  1787  1131   212   700   420  1100
2233   233  2302    55  1674   924  1711  2361  2373  1229  1710   260
846   636  1368   226  1431   432   721  1273     3   817   217   949
```

```
1381  1827  1075  1325   354   627  1695   184   439  1063  2428   463
1678  2028   325  2064  1137    50   358  1861  2125  1823  1231   823
2378  1903  2176  1534    89  1852   902  1539  1880   934  1647   622
1704  1945   634  1613  1857  1981  1083  1760  1150  2095  2478  1714
449  1483   107   234  1280  1354   763  2489  2256  2325   929  1119
1651  1367    45   170   197   283   338   421   453   467   473   628
651   705   864   935  1045  1103  1113  1115  1185  1223  1294  1388
1458  1484  1536  1572  1593  1596  1601  1604  1657  1896  2021  2082
2230  2267  2404  2493  2497];[643  2209   101  1002   602   133  1837
2349   177    74  2155  1488  2063  2191  1537  1364  2176  1757   142
891  1655   135   183  1656   604   382  1104  1983   587  2249  1769
186   507  1505  1155  1883  1396    45   592   180  1957   801  1940
1172  1162   182  2261  2273  2156   200  1069  1808    88   676  1471
486  1531   848  1599   943  1022  1600   404   653  2193   284   143
344  2210  2062  2311  2300  1197  1458   642   620   852  1766   367
202  1185   977  1381  1004  1277  1306  1548   850  1592  1038   157
707   810  2000   130   428  2274  1857  1111   717   561  1629   794
1677  1495  2279    72   148   154   162   174   181   189   213   264
268   279   296   353   356   492   563   606   611   637   718   721
750   760   799   829  1065  1177  1210  1256  1282  1337  1370  1400
1460  1675  1676  1717  1731  1812  1838  1847  1908  2018  2040  2050
2064  2095  2102  2167  2197  2356];[1517   846   701   139  2475   442
2638  1006  2282  1664   482  2076  1252  1579  1226  1788  2190  1632
2367  1489  1357  2195  1054  2434    10   133    30  2463   864  1144
1379   883   210  1857  2442   347  1622   151  1124    86   321   494
513  1525  1462  1319   233   462  2080   113   982  2315    57  1057
514  1269   357  1878  1019   670  2598  1056   860  1446   729  1940
230   410  2697  2114   740  1598   584  1142   518  2128  1683   700
1465  2476   661   674   502   441  1702   369  2398   485  1040  1417
1580  1287    81  1234   141  1952  1423   365    97   436   556    77
82   103   235   237   242   246   249   261   263   344   348   429
450   460   474   493   543   554   568   592   668   699   707   726
823   892   913   942   998  1020  1088  1113  1143  1170  1247  1304
1404  1424  1433  1434  1448  1456  1466  1467  1508  1578  1614  1709
1749  1787  1851  2024  2111  2215  2314  2352  2522  2576  2599  2624
2670  2717  2738];[2144  1532  1258   121  1505  1349   163  1305   268
153   545  1051  1915  1805   104  1059    22   480  1987  2137  1946
1924   615  1684   784   301  2148  1070  1783  1099  1233   193  1367
24  2120  1655  2085  1493   372   264   604  1753   440   987  1067
831   483  1497    20   170  1135   782  2077   590  2044  1693  1706
458  1803  2005  2021   478   316   699  1201  2073   443  2139  2130
513  1836  1162  1025   204  1989  1097   297  1206   452  1618  1307
454   933    58   780    47   949   210  1458  1037   548   850   439
1967  1308  1323  1052   710  2123   300     9   150   152   227   459
595   729   731   744   751   753   757   763   766   767   775   778
779   781   783   785   789   791   794   798   800   802   806   809
828  1004  1103  1108  1274  1414  1512  1515  1521  1523  1529  1531
1534  1535  1536  1561  1562  1563  1574  1579  1580  1586  1597  1604
1606  1610  1612  1732  1779  1806  1898  1964];[1966  2190   204  1537
1755  1419   644   611  1772   484   239  2189   781  1763   692  1959
597  2303  2514  2081  1284   488  1903   547  1363  1047   918   920
1780  1118   651  1978  2521  2506  1666  1809  2374  1875   765  2187
1278  2437    13   664  1600  2117  2459  2537   154  1493  1707   199
1932   536  1581  2481  1836  1743   864  1960   846   880  1732  2136
2370   554   105  2307  1762   432   595  1804   656  1988  1664   462
1128   594   835   854   314  1209  1705  1615  1371   539  2153  1593
2228   384  2080   257   842  1680  1472   313  2425   173  1268   743
879    12  1358  1893  2181  2074  2453   758   968  1468  2235  1101
```

```
2043  2015   928   472   252   380   414   461   501   505   574   640
722   729   766   805   821   886   891   939   993  1188  1219  1221
1292  1387  1453  1486  1531  1609  1695  1719  1720  1790  1858  1876
1897  1992  2052  2159  2330  2358  2491  2541  2566];};
%
% % SIX=0;ADJACENTONLY = 0;requireAll = 1; allowSubstring = 0;
% % CONTEXT = 2; pat={{'ribosome biogenesis GTP'; 'ribosome GTPase';
'YihA'; 'YsxC';'engb';};{'DNA polymerase I';'pola';'yceI';};};
% % pat={'ribosome biogenesis GTP'; 'ribosome GTPase'; 'YihA';
'YsxC';'engb';'DNA polymerase I';'pola';};
% SIX=0;ADJACENTONLY = 0;requireAll = 1; allowSubstring = 1;
pat={'cytochrome c'; 'c-type cytochrome';};

% genus='glac';
% SWITCH = 'for_Glaciecola';
%
gs={'igrs_glac_siph';'igrs_glac_KUL10';'igrs_glac_HTCC2999';'igrs_glac_33
A';'igrs_glac_MH2013';'igrs_glac_2405UD65';'igrs_glac_MF2'};
% % SCORES=13;ns={[1330  1080  2188   524   767   166   508  2112   567
395  1496  2493  2585  1742   120  2586  2048   186  1476  1958   408
1576   713   469    90   425  2053  1852  2470  2452  1656  2564  2469
1038    96  2457  2494  1837  1773  2368  1569   792  2213   359  2230
433  1709  1454  2588   742   174    37  1642   542  2047   998   173
556  1783   454  2505  1747   755   526   975  1001  2395  1564  2214
38   656  1559   376  2483   455   894   447     9    43  2462  2351
2519  1134  2243  1876   298    95  2461   889  1182  1241  1984  2207
1741  1214   576   805   153  1563  2106   464  1994   968  1194];[325
881  1722   107   445  1751  1953   802  1365  1502  1850   408   279
1440   768   565  2051    56    77  2008  2137   516  1183   518   912
866   121  2269     1  2091   252  1906  2065  1767  2292  1472  1584
428  1814  2300  2382  1612   911  1554  2139  1596   843  2274  1817
626   683  1035  1087   958  1246   427  1316   800   355  1360    83
2107   713  1706  2495  1391  2216   767   952  2234  1629   120    55
382   610   356   105  1663   957  1390   916  1553  1806    81  2159
517  1579  1250     4   585   649  2062];[830   910   799   980   513
319   738   827   185   982  1061   489   570   769  1111   270  1106
105   282   530   758   613   518   682  1051   767   615  1018   148
837   427   656  1053  1112   861    55   245   996  1123   539   858
803   640   610   707    35   772     8   496];[1141  2453  1171  1962
2868  2607   253  1345   818  2260  1922   727  2564  2939   435  3053
1719  2428   922  2602   644   788  2724  2496   692  2799   366  3147
1464  1858  2250   425  2813  1099  2631   874  2078  1565  2849  2343
309   923   209  2839  1086   419   319  1884  2045  2733  3144   402
560  1020  3004  2227  1516   226  2031   475  1423  1040  1638  2145
3279  3125   842  2725   870   647  2456  3048  1090  1968  3067  2503
1160   786   427  2529  1673  3084  3009  2392  2134  2761  2683  2741
2234  1019   488   494   787  1990   243   702  2204  1372  3185  2915
682  2217   747    94  1050   843   245  2315  1587  1607  1356  1916
2280  1666  2201   931  2331   194   876    81    35  1649  2388  2319
3149  2807    43  1954  3186  1822  1417   188  2651  2178  2650  1845
999];[1138  1793  2205   211   804   255  1935  1007  1781   790  1047
1052  1017   568  1059    39   321  1575   487  1412  1494  2366   599
82    92  2259   227  1276  1486  1885  1671  1928  1537  1008  1215
557   436  2206  1592  1051  2258  1278   338  2241   661   693  2227
1582  1693  1854  1676  2376  1231  1363   622  1255   736  1329  1647
1654  2272  1131  1418  1419  2041  1835  1915  1259    55   638   727
569  2065   258   226  1050   188   409   445   901  1668  1565   965
1496    32   749   457  1158   541  1809  1978   835   203   676   979
```

```
1782  1758  1640  2358];[358  1614  1603  1606   247   356  2161   222
511  1590   666   539  2162  1801  1459  1762  1356   457  1438  1593
2155   908  1009  2101   630   971  1010   308   162   345  1631   294
783    28   665  1254   340  1570   753  2107   232  2085   941  1806
1197   661   297  1819   943  1283   374  2184   656  1823   439   727
2250  1090  1861  1873  1779  1497  1162   295  1496  1196  1125  2031
947  2047  2256   398  1569   725   803  1993   974];[1187  2826   336
710  2249  1392  1225   473  1749  1408  1724  2274   925  2647  2149
1641  2091   976   889  2277  2294  1935  2243   588  1152   263  1657
154  1180  1539  2360   499   822  2572   773  2129  1505  2302  2045
2639  1477  2111    13  2121  2610   110  1773   118  2190  2315  2396
2321   707   429  2773   563  1359   831   371    23  1142   367  1145
767  2594  1211  2515   447  1835  1859  2827   864   700  2481  2026
2482  2163  1968  2411   244   693  2267   189  1726   238  2098   249
1182  2704  2874  1630  2324  2207  1125  1257  1772  2012  2386  1327
728  1510  2492  1222  1489  1558  1742  1761   435  1301   890  2009
2126  2843   962  1877  2089   264  1743  2148  1640];};
% %
SCORES=14;gs={'igrs_glac_siph';'igrs_glac_KUL10';'igrs_glac_33A';};ns={[1
080   425   408   704   166  2213  1364   524  2483   642   957  2053
376  1330  1958  1996  2493    43    96   186   447   542   605  1194
174   454  1576  1974  2343   433   455  1920  2112  2297  2395   120
363   381   540   901   911  1747  1867  2188  2214   556   753  1114
2230   375  1001  1199  1271  1668  2452  2585   185   742   767  1034
1214  1707  1781  1784  2368  2470    92   153   229   283   395   469
492   808  1210  1852  1870  1917  2054  2494  2516  2561    89   149
204   615   728  1038  1500  1650  1985  2153  2449  2588];[408  1722
881  1850   445  2091  1087  1472   107  2269   952   121   325  1360
610   649   911  1729  1751   420  1164  1365   291   733   866  1416
1471  1630  1923  2008  2382    77   200   419  1159    56   495   519
626   802   870  1508  1596  1671  1934  2051  2107  2212   195   273
329   743   768   874   902   912  1541  2013  2039  2046  2070  2300
2363  2457   279   639   800   851   953  1316  1552  1647  1770  1870
2065    91   144   180   346   389   432   631   758   935  1168  1430
1769  1798  2137  2143  2322];[253  1962  2453  2607   362   435  1566
2496   425  1149  2085  2750   633  1020  1171  1515  2031  2309  3053
3279   870  2817   647   787  1545  2145  2171  2755  2767   788   957
1345  2250  2564   818  1374  1615  1882  2133  2956  1141  1821  2134
3084   234   644  2280  2939    43   209   366   427   807   967  1166
1828  1922  1954  2315  2683  2868   810  1327  1417  1697  1990  2260
2350  2631  2768  2799  3102  3274    31    34   141   204   309   319
910   923  1086  1480  1516  1607  2019  2217  2894  3125   101   191
645  1099  1160  1278  1666  1685  1746  2045  2228  2359  2420  2853
2921   127   330   367   560   682   938  1150  1365  1640  1884  2219
2319  2368  2390  2428  2792  2849];};
% % SCORES=9;gs={'igrs_glac_siph';};ns={[90    96   120   153   166   173
174   178   186   208   227   288   332   333   350   376   395   408
425   433   437   448   455   469   507   508   524   526   556   567
711   713   742   755   767   792   808   894   957   967   977  1001
1007  1025  1038  1080  1182  1241  1250  1283  1330  1342  1382  1391
1447  1454  1469  1470  1476  1496  1559  1569  1576  1593  1618  1641
1656  1679  1709  1742  1747  1761  1773  1830  1832  1852  1876  1917
1944  1958  1984  1994  2048  2053  2063  2102  2112  2114  2188  2200
2213  2226  2230  2236  2243  2246  2299  2326  2350  2351  2368  2435
2445  2449  2452  2469  2470  2493  2544  2564  2585  2586  2593];};
% SCORES=[13 14 9];
% ns={[1330  1080  2188   524   767   166   508  2112   567   395  1496
2493  2585  1742   120  2586  2048   186  1476  1958   408  1576   713
```

```
469    90   425  2053  1852  2470  2452  1656  2564  2469  1038    96
2457  2494  1837  1773  2368  1569   792  2213   359  2230   433  1709
1454  2588   742   174    37  1642   542  2047   998   173   556  1783
454  2505  1747   755   526   975  1001  2395  1564  2214    38   656
1559   376  2483   455   894   447     9    43  2462  2351  2519  1134
2243  1876   298    95  2461   889  1182  1241  1984  2207  1741  1214
576   805   153  1563  2106   464  1994   968  1194   704  1364   642
957  1996   605  1974  2343  1920  2297   363   381   540   901   911
1867   753  1114   375  1199  1271  1668   185  1034  1707  1781  1784
92   229   283   492   808  1210  1870  1917  2054  2516  2561    89
149   204   615   728  1500  1650  1985  2153  2449   178   208   227
288   332   333   350   437   448   507   711   967   977  1007  1025
1250  1283  1342  1382  1391  1447  1469  1470  1593  1618  1641  1679
1761  1830  1832  1944  2063  2102  2114  2200  2226  2236  2246  2299
2326  2350  2435  2445  2544  2593];[325   881  1722   107   445  1751
1953   802  1365  1502  1850   408   279  1440   768   565  2051    56
77  2008  2137   516  1183   518   912   866   121  2269     1  2091
252  1906  2065  1767  2292  1472  1584   428  1814  2300  2382  1612
911  1554  2139  1596   843  2274  1817   626   683  1035  1087   958
1246   427  1316   800   355  1360    83  2107   713  1706  2495  1391
2216   767   952  2234  1629   120    55   382   610   356   105  1663
957  1390   916  1553  1806    81  2159   517  1579  1250     4   585
649  2062  1729   420  1164   291   733  1416  1471  1630  1923   200
419  1159   495   519   870  1508  1671  1934  2212   195   273   329
743   874   902  1541  2013  2039  2046  2070  2363  2457   639   851
953  1552  1647  1770  1870    91   144   180   346   389   432   631
758   935  1168  1430  1769  1798  2143  2322    76   109   123   295
541   577   630   836   895   901   960   977  1052  1078  1167  1506
1555  1567  1632  1658  1721  1746  1784  1786  1822  1834  1836  1898
1938  1959  2105  2169  2293  2297  2303  2324  2333];[830   910   799
980   513   319   738   827   185   982  1061   489   570   769  1111
270  1106   105   282   530   758   613   518   682  1051   767   615
1018   148   837   427   656  1053  1112   861    55   245   996  1123
539   858   803   640   610   707    35   772     8   496   919   574
607  1125   687   757   236   490   512   670   960  1076    51   765
781   987  1046   304   614   992  1092    50    79   145   176   262
309   347   348   356   387   393   417   473   479   586   683   744
895  1066  1179];[1141  2453  1171  1962  2868  2607   253  1345   818
2260  1922   727  2564  2939   435  3053  1719  2428   922  2602   644
788  2724  2496   692  2799   366  3147  1464  1858  2250   425  2813
1099  2631   874  2078  1565  2849  2343   309   923   209  2839  1086
419   319  1884  2045  2733  3144   402   560  1020  3004  2227  1516
226  2031   475  1423  1040  1638  2145  3279  3125   842  2725   870
647  2456  3048  1090  1968  3067  2503  1160   786   427  2529  1673
3084  3009  2392  2134  2761  2683  2741  2234  1019   488   494   787
1990   243   702  2204  1372  3185  2915   682  2217   747    94  1050
843   245  2315  1587  1607  1356  1916  2280  1666  2201   931  2331
194   876    81    35  1649  2388  2319  3149  2807    43  1954  3186
1822  1417   188  2651  2178  2650  1845   999   362  1566  1149  2085
2750   633  1515  2309  2817  1545  2171  2755  2767   957  1374  1615
1882  2133  2956  1821   234   807   967  1166  1828   810  1327  1697
2350  2768  3102  3274    31    34   141   204   910  1480  2019  2894
101   191   645  1278  1685  1746  2228  2359  2420  2853  2921   127
330   367   938  1150  1365  1640  2219  2368  2390  2792    77   115
221   273   321   441   471   515   678   728   811   859  1018  1057
1172  1271  1403  1413  1521  1594  1645  1717  1758  1860  1906  1917
2114  2210  2230  2239  2304  2354  2395  2557  2581  2715  2890  3093
3132];[1138  1793  2205   211   804   255  1935  1007  1781   790  1047
```

```
1052  1017   568  1059    39   321  1575   487  1412  1494  2366   599
82    92  2259   227  1276  1486  1885  1671  1928  1537  1008  1215
557   436  2206  1592  1051  2258  1278   338  2241   661   693  2227
1582  1693  1854  1676  2376  1231  1363   622  1255   736  1329  1647
1654  2272  1131  1418  1419  2041  1835  1915  1259    55   638   727
569  2065   258   226  1050   188   409   445   901  1668  1565   965
1496    32   749   457  1158   541  1809  1978   835   203   676   979
1782  1758  1640  2358   905  2260     3   498   514  1702  2084  2151
228  1348   704   729   854  1584   647  1349  1982  2297    78   833
1048  1670  1977  2144  2202  2252   316   545   684   737  1450  1841
2242   198   651  1394  1743  1966  2031  2126    30   117   144   154
166   167   293   341   404   413   439   455   530   629   649   701
824   826   874   988  1191  1221  1251  1325  1523  1524  1580  1756
1803  1846  1879  1886  2061  2135  2140  2264];[358  1614  1603  1606
247   356  2161   222   511  1590   666   539  2162  1801  1459  1762
1356   457  1438  1593  2155   908  1009  2101   630   971  1010   308
162   345  1631   294   783    28   665  1254   340  1570   753  2107
232  2085   941  1806  1197   661   297  1819   943  1283   374  2184
656  1823   439   727  2250  1090  1861  1873  1779  1497  1162   295
1496  1196  1125  2031   947  2047  2256   398  1569   725   803  1993
974   151   157   515  1093  2048   221   329   635   964  1753   224
994   644   465  1537  2009  1077  1106  1236  1568  1597  1848  1865
2050    11    21   298   454   458  1136  1499  1508  1851  1945  2049
10   462   899   900  1104  1240  1323  1737  1994  2140  2238    98
103   143   154   275   276   316   333   361   406   418   423   451
561   837   877   962  1004  1172  1203  1229  1282  1410  1423  1428
1485  1513  1622  1746  1751  1791  1830  1868  1869  1942  1969  2159
2214  2219  2260];[1187  2826   336   710  2249  1392  1225   473  1749
1408  1724  2274   925  2647  2149  1641  2091   976   889  2277  2294
1935  2243   588  1152   263  1657   154  1180  1539  2360   499   822
2572   773  2129  1505  2302  2045  2639  1477  2111    13  2121  2610
110  1773   118  2190  2315  2396  2321   707   429  2773   563  1359
831   371    23  1142   367  1145   767  2594  1211  2515   447  1835
1859  2827   864   700  2481  2026  2482  2163  1968  2411   244   693
2267   189  1726   238  2098   249  1182  2704  2874  1630  2324  2207
1125  1257  1772  2012  2386  1327   728  1510  2492  1222  1489  1558
1742  1761   435  1301   890  2009  2126  2843   962  1877  2089   264
1743  2148  1640   436  1294  2376  2643   155  1861  1686  2025  2825
1214  2316  1993  2692  2275    95   766  1136  1555  1817  2663   886
1131  1721  2196  2735   551   867  1253  1358  2154  2569  2714  2816
352   362   423   567   689  1299  1376  1520  2070  2244   321   380
1037  1349  1437  1465  1600  2465  2636  2698  2808    54   125   182
242   320   361   372   387   389   393   403   416   470   474   535
565   575   645   649   675   682   907   915   991  1050  1246  1272
1284  1323  1343  1360  1514  1622  1803  1818  1822  1923  1941  2127
2138  2319  2403  2437  2486  2625  2693];};

% SIX=0;ADJACENTONLY = 0;requireAll = 1; allowSubstring = 0;
% % pat={'ribosome biogenesis GTP'; 'ribosome GTPase'; 'YihA';
'YsxC';'engb';'DNA polymerase I';'pola';'ElbB'};
% % pat={'ribosome biogenesis GTP'; 'ribosome GTPase'; 'YihA';
'YsxC';'engb';'DNA polymerase I';'pola';};
% % pat={'DMT'; 'VRR-NUC'; 'shikimate';};
% CONTEXT = 2;
% pat={{'ribosome biogenesis GTP'; 'ribosome GTPase'; 'YihA';
'YsxC';'engb';};{'DNA polymerase I';'pola';};};
```

```
% genus='Idiomarina';
% d = '';
%
gs={'igrs_Idiomarina_aquatica';'igrs_Idiomarina_abyssalis';'igrs_Idiomari
na_xiamenensis';'igrs_Idiomarina_piscisalsi';'igrs_Idiomarina_baltica';
%
      'igrs_Idiomarina_zobellii';'igrs_Idiomarina_tyrosinivorans';'igrs_I
diomarina_fontislapidosi';'igrs_Idiomarina_ramblicola';'igrs_Idiomarina_A
28L';
%     'igrs_Idiomarina_MD25a';'igrs_Idiomarina_28-
8';'igrs_Idiomarina_loihiensis';'igrs_Idiomarina_seosinensis';'igrs_Idiom
arina_rhizosphaerae';
%     'igrs_Idiomarina_aminovorans';};
%
% %   {'~/hs/spot42_Eco.rf';'igrs_Idiomarina_aminovorans};
% SCORES=13;
% ns={[248  164   49  141  249  334  322  215  213  160   60  139  230
159    6  212  327  309  145  264  154  218  171  162];[169  179  103
262  241    7  107  255  205   61  279  122   53  104];[423  338  214
192  116  225  283   72  382   99  403  124  438  134  232  166  276  467
110  219  255  367  162];[56   16    3  110   64  165  100   55   84  183
89];[57  169  166  224  141  249  234  213   37  226  263  137  232  144
250   70  163   50  196  188   39];[148   86  197   40  116  212   38
248  281  127  118  135   47  259   74  202   41];[102  220  218   19
284   79  210   21  107   88  229   78  109];[139   50  218   53   75
284   79  279   31  294  167  238  292  175   52  129  168  295  176
125];[88  125  181  153  185  226  230  227  148  190   64   73  145  261
5];[215   25  207  287  247  513   89  193  246  189  510  365   41  431
458  439  186  322  366  260  214  333  228  221  343  388];[177   35
176  181  127   98  193   85   99   32   82  110];[151  336  136  330
223  161  186  173   44   32  219  145   46  171   17  142  216];[308
151    7   70  159  115   97  219  257   34  252   60   16  174  288   83
286];[112  241   29  201  229   15  143  213  126  170  147  234  193
34];[57  264  241  200   78  172   66  196  293   31  147  218  303  290
185];[215   74    4  349  155  144  187   56  199  143  202  136  201
68  319  108  347  327  158  386  307   86  364];};
% SCORES = 14;
% ns={[230  264  145  160  218  248  162  116  136  323  164   60  139
199   49  154  213  324  180  333    4];[103  169   53  179   70   95
107  221  205   22  127];[338  134  382  423  225  192   30  214  400
467  328  403  291   24   50   84   99  166  190  466  309  116  217
231];[3  191   57   64   16  165   60   84  110   56  209];[37   26  163
237  249   20   46  188   70  200  255   83  130  141  142  263   57  131
144  169  228  250];[86   40   47  148  197   89  248   62  149   68  127
212   16   38  116   52  259  280  320  115];[220  263   19   39   79
130  185   53  188  218   57  102  109  111];[218  108   52   60  167
279  140   22  128  166   75  238   53   79];[88  125  181  208   80  215
227  153   64   73  260  261   51  116  190];[215  507  284  189    3
287  426  217  247   56  135  228  216  345  474  186  207   25  193  366
280  513];[181    4   99   64   30   72  128   61  160   46];[151  223
219  165  186   58   62   79  136  256   45  119  286  304];};
% SCORES = 12;
% ns={[145  264    4  136  230   60  139  151  160  162  333  180  213
218  245  248];[103  133   22   70  107  127  163  169  221  277   19
41   53   95  179  262   16];[134  338  467  382  423  116  225  227  291
466  468   30   84  104  192  231  270  313  328  373  400];[3   57   16
191  209   64   84   51   60  110  165];[37  249   20   26   93    5   46
70  188   57  131  141  142  163  200  231  235  237];[47   68   40   86
```

```
148  89  62  127  149  197  34  50  116  248  270  320];[220  39
79  111  176  213  23  53  109  130  188  263  18  19  57  185
223];[218  108  279  52  60  167  12  140  22  163  166  240  252
14  66  87  110  128  238  271];[80  88  125  181  208  211  215  227
64  129  131  153  9  147  229  260  261];[215  507  284  189  290  230
3  127  216  217  287  320  345  426  474  66  69  169  228  247  309
418  432  508];[181  4  99  64  128  156  30  46  61  160  164
192];[151  165  223  186  219  58  79  136  160  40  32  62  135
237  304];};
% SCORES = 9;
% ns={[49  138  139  141  142  160  161  164  195  203  212  220  230
248  249  309  324  335];[7  50  53  74  103  107  122  149  169  179
183  205  241  255  262  279];[99  116  124  134  175  192  214  225  231
235  283  324  338  370  382  403  409  423  438  467];[3  16  35  38
53  56  64  84  100  110  113  146  151  155  161];[37  50  52  57
70  87  137  141  149  166  169  188  224  247  249  255];[38  40  41
86  89  103  118  120  127  135  148  185  197  202  217  248  259  263
281  299];[1  19  21  79  90  102  109  114  142  210  218  220  260
284];[31  50  52  53  75  104  139  167  168  175  218  238  261  276
279  284  295];[60  64  73  88  116  118  121  122  125  145  148  153
181  185  226  227  268];[3  4  25  36  89  106  181  186  187  193
207  215  221  229  230  246  247  287  365  381  412  447  451  463
510];[46  72  85  98  109  127  157  176  177  181  193];[17  32  35
44  46  136  142  145  151  173  186  219  223  304  307  336  338];};
% SCORES=3;
% ns={[4  7  69  106  116  118  139  145  148  151  153  160  162  180
204  212  236  249  264  302  309  323];[22  51  53  70  103  127  133
163  179  197  205  221  233  265  274  277];[30  50  116  134  164  215
227  231  238  263  291  321  338  340  342  382  400  403  408  423  466
467  468];[3  31  51  57  69  100  110  163  173  191];[5  19  26
46  70  93  131  141  153  200  226  235  249  250  280];[1  8  40
47  68  86  89  127  148  149  197  212  248  270  320];[20  47  57
79  109  111  145  147  174  188  213  220  253];[22  36  52  60  108
123  140  167  218  226  265  275  279];[9  57  64  80  87  88  125
129  147  153  181  208  211  215  260];[3  16  86  135  168  182  189
215  216  217  228  258  284  287  298  320  346  364  384  474  488
507];[4  30  46  64  70  87  99  128  160  164  181  192  197];[58
60  61  64  99  151  165  173  175  186  188  195  215  216  223  260
293  304  307];};
% SCORES=[11 10 1 2 4 5 6 7 8];
% ns={[23  60  138  141  154  160  161  164  195  215  220  230  243
248  249  286  305  309  322  49  65  89  139  142  203  212  273  281
324  335  4  7  12  40  55  69  116  130  145  148  151  180  204
236  323  6  62  72  103  218  334  162  45  135  206  209];[5  7
50  74  91  94  103  149  158  169  179  241  255  262  53  113  183
205  206  274  277  279  22  51  70  127  133  233  235  265  63  95
119  170  181  77  180  155  256  54  61  67  104  191  220  122];[19
77  96  99  110  114  116  124  125  134  192  214  221  225  231  232
233  235  283  284  338  343  382  403  423  70  175  219  264  340  370
409  438  442  467  30  34  146  164  179  215  227  256  263  274  289
321  337  342  400  408  50  162  174  238  258  291  24  72  153  276
310  320  356  441  45  8  84  166  266  277  279  325  327  457  68
201  228  452];[3  4  16  29  30  35  42  64  100  146  156  165
7  56  84  110  113  151  155  14  31  51  57  60  69  163  167
173  191  209  77  169  186  193  89  55  38  46  129  63];[37  50
52  57  70  94  103  113  141  144  149  155  166  188  196  235  238
255  39  87  114  137  213  224  280  5  19  20  26  46  64  93
97  130  131  175  200  226  237  243  250  132  15  83  169  228  249
```

```
263   23  122  123  195  147];[40   41   47   52   86  103  116  117  118
127  135  148  153  174  197  202  205  248  260  263  280  282  314   38
39   89  183  217  259  281    1    8   16   68   74  134  149  212  270
44   67   85  107  189   42   90  128  170  206  299  320  115  190];[1
21   79   83  102  107  109  114  155  186  220  229  263   19   77   90
142  210  212  218  241  259  260  284   47   57   80   95  111  147  204
20  174  280   78  182  213  232   91  125  148  154  156    8   87   88
145  254  258  271];[1   31   50  104  125  139  166  167  175  218  227
232  239  261  273  276  279  294  295    3   52   53   75   94  168  176
238  284   22  108  112  123  140  226  275   60  262   16   38   66   70
211  224  265  292  113  135   79  221];[51   60   64   88  118  122  125
145  153  174  181  185  228  267  268    5   73   78  116  121  148  226
227   22   53   77   86   87  123  135  147  208  260   80  215  190  230
10  161  261  237   70  146  188  246  264   72];[3    9   22   25   36
89  106  117  130  164  207  215  219  246  247  287  297  353  363  370
381  388  412  431  447  451  455  458  463  471  510  513    4   78  181
186  189  214  221  229  230  365  426    1    7   16   71   86  135  174
182  216  217  231  257  284  298  320  324  346  350  420  454  480  488
507   39  168  474   41  103  260  322  333  338  410  228  343  364  384
193  258  309  323  330  387  391  411  419   67  336];[35   48   61   67
72   85   99  108  109  112  127  157  160  177  193   46   64   82   98
176    4   74  164  169  181   30  128  197   32  158  159    7   37
150];[44   46   62   63   72  108  136  142  145  151  161  165  173  186
223  274  276  336  338   17   32   35   70  137  158  210  219  269  304
307  329   53   60   61   64  175  188  215  216  293  268  286   45  109
159  168  195  229  330   96  196  204  254];};
% SCORES=[11 7];
% ns={[49   65   89  138  139  141  142  161  164  203  212  248  273
281  324  335    4    6   40   45  135  154  162  206  209  249  334];[7
50   53  103  113  149  169  179  183  205  206  241  262  274  277  279
54   61   67   74  104  191  220  235  256];[70   99  116  124  134  175
192  219  225  235  264  283  338  340  370  382  409  423  438  442  467
34   72   84  166  214  232  266  276  277  279  325  327  457];[3    7
16   35   56   64   84  100  110  113  151  155   38   46   60  129
191];[39   57   70   87  114  137  141  149  166  188  213  224  255  280
19   23   50   97  122  123  144  169  195  196  226  235  249  250];[38
39   40   41   86   89  103  118  148  183  197  202  217  259  263  281
42   67   90  107  128  135  170  174  206  248  299  320];[19   21   77
79   90  102  114  142  210  212  218  220  241  259  260  284    8   87
88   91   95  145  147  182  213  254  258  271];[3   50   52   53   75
94  139  167  168  175  176  238  261  279  284  294  295   38  113  125
135  224];[5   60   64   73   78   88  116  118  121  125  145  148  181
185  226  227  268   10   70  146  161  188  215  230  246  264];[3    4
36   78  181  186  189  207  214  215  221  229  230  246  247  287  365
412  426  447  463    9   25   41   89  103  117  193  216  258  284  309
322  323  330  343  384  387  388  391  411  419  431  451  510];[46   64
72   82   85   98  109  127  176  177    7  150  164  181  193];[17   32
35   44   46   70  136  137  142  151  158  173  186  210  219  223  269
304  307  329  336  338   45   96  108  109  145  159  161  165  196  204
229  254];};
% SCORES=[12 9 10];
% ns={[145  264    4  136  230   60  139  151  160  162  333  180  213
218  245  248   49  138  141  142  161  164  195  203  212  220  249  309
324  335   65   89  273  281];[103  133   22   70  107  127  163  169
221  277   19   41   53   95  179  262   16    7   50   74  122  149  183
205  241  255  279  113  206  274];[134  338  467  382  423  116  225
227  291  466  468   30   84  104  192  231  270  313  328  373  400   99
124  175  214  235  283  324  370  403  409  438   70  219  264  340
```

```
442];[3   57   16  191  209   64   84   51   60  110  165   35   38   53
56  100  113  146  151  155  161    7];[37  249   20   26   93    5   46
70  188   57  131  141  142  163  200  231  235  237   50   52   87  137
149  166  169  224  247  255   39  114  213  280];[47   68   40   86  148
89   62  127  149  197   34   50  116  248  270  320   38   41  103  118
120  135  185  202  217  259  263  281  299   39  183];[220   39   79
111  176  213   23   53  109  130  188  263   18   19   57  185  223    1
21   90  102  114  142  210  218  260  284   77  212  241  259];[218  108
279   52   60  167   12  140   22  163  166  240  252   14   66   87  110
128  238  271   31   50   53   75  104  139  168  175  261  276  284  295
3   94  176  294];[80   88  125  181  208  211  215  227   64  129  131
153    9  147  229  260  261   60   73  116  118  121  122  145  148  185
226  268    5   78];[215  507  284  189  290  230    3  127  216  217
287  320  345  426  474   66   69  169  228  247  309  418  432  508    4
25   36   89  106  181  186  187  193  207  221  229  246  365  381  412
447  451  463  510   78  214];[181    4   99   64  128  156   30   46
61  160  164  192   72   85   98  109  127  157  176  177  193   82];[151
165  223  186  219   58   79  136  160   40   32   62  135  237  304   17
35   44   46  142  145  173  307  336  338   70  137  158  210  269
329];};
% SCORES=[12 9 10 7];
% % ns={[145  264    4  136  230   60  139  151  160  162  333  180  213
218  245  248   49  138  141  142  161  164  195  203  212  220  249  309
324  335   65   89  273  281    6   40   45  135  154  206  209
334];[103  133   22   70  107  127  163  169  221  277   19   41   53
95  179  262   16    7   50   74  122  149  183  205  241  255  279  113
206  274   54   61   67  104  191  220  235  256];[134  338  467  382
423  116  225  227  291  466  468   30   84  104  192  231  270  313  328
373  400   99  124  175  214  235  283  324  370  403  409  438   70  219
264  340  442   34   72  166  232  266  276  277  279  325  327  457];[3
57   16  191  209   64   84   51   60  110  165   35   38   53   56  100
113  146  151  155  161    7   46  129];[37  249   20   26   93    5   46
70  188   57  131  141  142  163  200  231  235  237   50   52   87  137
149  166  169  224  247  255   39  114  213  280   19   23   97  122  123
144  195  196  226  250];[47   68   40   86  148   89   62  127  149  197
34   50  116  248  270  320   38   41  103  118  120  135  185  202  217
259  263  281  299   39  183   42   67   90  107  128  170  174
206];[220   39   79  111  176  213   23   53  109  130  188  263   18
19   57  185  223    1   21   90  102  114  142  210  218  260  284   77
212  241  259    8   87   88   91   95  145  147  182  254  258
271];[218  108  279   52   60  167   12  140   22  163  166  240  252
14   66   87  110  128  238  271   31   50   53   75  104  139  168  175
261  276  284  295    3   94  176  294   38  113  125  135  224];[80   88
125  181  208  211  215  227   64  129  131  153    9  147  229  260  261
60   73  116  118  121  122  145  148  185  226  268    5   78   10   70
146  161  188  230  246  264];[215  507  284  189  290  230    3  127
216  217  287  320  345  426  474   66   69  169  228  247  309  418  432
508    4   25   36   89  106  181  186  187  193  207  221  229  246  365
381  412  447  451  463  510   78  214    9   41  103  117  258  322  323
330  343  384  387  388  391  411  419  431];[181    4   99   64  128
156   30   46   61  160  164  192   72   85   98  109  127  157  176  177
193   82    7  150];[151  165  223  186  219   58   79  136  160   40
32   62  135  237  304   17   35   44   46  142  145  173  307  336  338
70  137  158  210  269  329   45   96  108  109  159  161  196  204  229
254];};
% SCORES=[14 12 9 3];
% ns={[230  264  145  160  218  248  162  116  136  323  164   60  139
199   49  154  213  324  180  333    4  151  245  138  141  142  161  195
```

```
203  212  220  249  309  335    7   69  106  118  148  153  204  236
302];[103  169   53  179   70   95  107  221  205   22  127  133  163
277   19   41  262   16    7   50   74  122  149  183  241  255  279   51
197  233  265  274];[338  134  382  423  225  192   30  214  400  467
328  403  291   24   50   84   99  166  190  466  309  116  217  231  227
468  104  270  313  373  124  175  235  283  324  370  409  438  164  215
238  263  321  340  342  408];[3  191   57   64   16  165   60   84  110
56  209   51   35   38   53  100  113  146  151  155  161   31   69  163
173];[37   26  163  237  249   20   46  188   70  200  255   83  130  141
142  263   57  131  144  169  228  250   93    5  231  235   50   52   87
137  149  166  224  247   19  153  226  280];[86   40   47  148  197   89
248   62  149   68  127  212   16   38  116   52  259  280  320  115   34
50  270   41  103  118  120  135  185  202  217  263  281  299    1
8];[220  263   19   39   79  130  185   53  188  218   57  102  109  111
176  213   23   18  223    1   21   90  114  142  210  260  284   20   47
145  147  174  253];[218  108   52   60  167  279  140   22  128  166
75  238   53   79   12  163  240  252   14   66   87  110  271   31   50
104  139  168  175  261  276  284  295   36  123  226  265  275];[88  125
181  208   80  215  227  153   64   73  260  261   51  116  190  211  129
131    9  147  229   60  118  121  122  145  148  185  226  268   57
87];[215  507  284  189    3  287  426  217  247   56  135  228  216  345
474  186  207   25  193  366  280  513  290  230  127  320   66   69  169
309  418  432  508    4   36   89  106  181  187  221  229  246  365  381
412  447  451  463  510   16   86  168  182  258  298  346  364  384
488];[181    4   99   64   30   72  128   61  160   46  156  164  192
85   98  109  127  157  176  177  193   70   87  197];[151  223  219  165
186   58   62   79  136  256   45  119  286  304  160   40   32  135  237
17   35   44   46  142  145  173  307  336  338   60   61   64   99  175
188  195  215  216  260  293];[159  287  299  308  115   63    7  245
93  244  229   33   60  252  274   71   69  173  247    4   11   34   57
70   97  151  196  219  257  288    8   16   38   87  150  221];[29  229
19   25  112   34   68  182   82   87  106   15  172  213  236   26   52
99   16  126  143  147  193  201  234  241  117  206  217];[57  241   46
125  185  108  134  264  263   86  172   31   88   92  104  200  293   45
192   12   41   67  105  147  168  196  255  268  289  290  303  305   66
85  101  256  282];[215  349  233    4  144  167  257  354  187  307  199
364   34   56   57   85  202  342  103  388   72   77  158  163  247   68
74   92  108  136  139  155  209  303  319  347  386   55  110  181  200
201  330  368];};
% SCORES=[14 12 9 3 7];
% ns={[230  264  145  160  218  248  162  116  136  323  164   60  139
199   49  154  213  324  180  333    4  151  245  138  141  142  161  195
203  212  220  249  309  335    7   69  106  118  148  153  204  236  302
6   40   45  135  206  209  334];[103  169   53  179   70   95  107  221
205   22  127  133  163  277   19   41  262   16    7   50   74  122  149
183  241  255  279   51  197  233  265  274   54   61   67  104  191  220
235  256];[338  134  382  423  225  192   30  214  400  467  328  403
291   24   50   84   99  166  190  466  309  116  217  231  227  468  104
270  313  373  124  175  235  283  324  370  409  438  164  215  238  263
321  340  342  408   34   72  219  232  266  276  277  279  325  327
457];[3  191   57   64   16  165   60   84  110   56  209   51   35   38
53  100  113  146  151  155  161   31   69  163  173   46  129];[37   26
163  237  249   20   46  188   70  200  255   83  130  141  142  263   57
131  144  169  228  250   93    5  231  235   50   52   87  137  149  166
224  247   19  153  226  280   23   97  122  123  195  196];[86   40   47
148  197   89  248   62  149   68  127  212   16   38  116   52  259  280
320  115   34   50  270   41  103  118  120  135  185  202  217  263  281
299    1    8   42   67   90  107  128  170  174  206];[220  263   19
```

```
39   79  130  185   53  188  218   57  102  109  111  176  213   23   18
223    1   21   90  114  142  210  260  284   20   47  145  147  174  253
8   87   88   91   95  182  254  258  271];[218  108   52   60  167  279
140   22  128  166   75  238   53   79   12  163  240  252   14   66   87
110  271   31   50  104  139  168  175  261  276  284  295   36  123  226
265  275   38  113  125  135  224];[88  125  181  208   80  215  227  153
64   73  260  261   51  116  190  211  129  131    9  147  229   60  118
121  122  145  148  185  226  268   57   87   10   70  146  161  188  230
246  264];[215  507  284  189    3  287  426  217  247   56  135  228
216  345  474  186  207   25  193  366  280  513  290  230  127  320   66
69  169  309  418  432  508    4   36   89  106  181  187  221  229  246
365  381  412  447  451  463  510   16   86  168  182  258  298  346  364
384  488    9   41  103  117  322  323  330  343  387  388  391  411  419
431];[181    4   99   64   30   72  128   61  160   46  156  164  192
85   98  109  127  157  176  177  193   70   87  197    7  150];[151  223
219  165  186   58   62   79  136  256   45  119  286  304  160   40   32
135  237   17   35   44   46  142  145  173  307  336  338   60   61   64
99  175  188  195  215  216  260  293   96  108  109  158  159  161  196
204  229  254];[159  287  299  308  115   63    7  245   93  244  229
33   60  252  274   71   69  173  247    4   11   34   57   70   97  151
196  219  257  288    8   16   38   87  150  221   58   80   85   88  106
180  215  251  279];[29  229   19   25  112   34   68  182   82   87  106
15  172  213  236   26   52   99   16  126  143  147  193  201  234  241
117  206  217  186  205  210];[57  241   46  125  185  108  134  264  263
86  172   31   88   92  104  200  293   45  192   12   41   67  105  147
168  196  255  268  289  290  303  305   66   85  101  256  282   22
193];[215  349  233    4  144  167  257  354  187  307  199  364   34
56   57   85  202  342  103  388   72   77  158  163  247   68   74   92
108  136  139  155  209  303  319  347  386   55  110  181  200  201  330
368    6   52   81  123  145  152  273  359  360];};
% SCORES=[1 2 3 4 5 6 7 8 9 10 11 12 14];
% ns={[4    7   12   40   49   55   60   69  116  130  145  148  151  160
180  249  273  309  324  204  236  323  106  118  139  153  162  212  264
302    6   62   72  103  141  154  218  248  334  195  220   45  135  161
206  209  335   65   89  138  142  164  203  281  230   23  215  243  286
305  322  136  333  213  245  199];[22   51   53   70  103  127  133  149
179  205  206  233  235  265  277   63   95  119  170  181  279  163  197
221  274   77  169  180  155  241  256    7   54   61   67   74  104  191
220   50  113  122  183  262  107  255    5   91   94  158   19   41
16];[30   34  116  134  146  164  179  215  227  231  256  263  274  289
321  337  338  342  400  403  408  409  423  467   50  162  174  192  214
238  258  291  340  382  466  468   24   72   96  153  175  219  276  310
320  356  441   45   99  225  232    8   84  166  266  277  279  325  327
457   68   70  124  201  228  235  264  283  370  438  442  452  324   19
77  110  114  125  221  233  284  343  104  270  313  328  373  190  309
217];[3   14   31   51   57   60   69  110  163  167  173  191  209   77
100  169  186  193   16   56   89  151   55   64   38   46   84  129    7
63  113  155   35   53  146  161    4   29   30   42  156  165];[5   19
20   26   39   46   50   64   70   93   94   97  130  131  155  175  200
226  235  237  243  250  280  132  141  153  249   15   57   83  103  137
166  224  144  169  228  263   23  122  123  195  196  114  147  149  213
255   37   52   87  188  247  113  238  142  163  231];[1    8   16   40
47   68   74   86   89  127  134  148  149  197  202  212  248  270   38
320   44   67   85  107  116  189  118   42   90  103  128  135  170  174
206  281  299   39   41  115  190  217  259  263  120  185  183   52  117
153  205  260  280  282  314   62   34   50];[47   57   79   80   95  111
147  204  220   19   20  174  218  280  109  145  188  213  253    1   21
77   78  102  182  232   91  125  148  154  156    8   87   88  254  258
```

```
271  114  142  210  212  241  259  284   90  260   83  107  155  186  229
263   39  176   23   53  130   18  185  223];[3   22   52   94  108  112
123  139  140  166  167  218  226  275  279  294   53   60  262   36  265
16   31   38   50   66   70  175  176  211  224  292   75  113  125  135
238  295   79  168  221  261  284  104  276    1  227  232  239  273   12
163  240  252   14   87  110  128  271];[22   53   60   64   77   86   87
88  123  125  135  147  181  208  227  260   51   73   80  153  215    9
57  129  211  145  148  185  190  226  230  267   10  161  261  237   70
146  188  246  264    5   72   78  116  118  121  268  122  174  228  131
229];[1    3    7   16   25   71   86  135  174  182  189  215  216  217
231  257  284  287  298  320  324  346  350  420  451  454  480  488  507
39  168  474  228  258  364  384   41  103  186  207  214  247  260  322
333  338  381  388  410  431   89  343    9   36  117  193  221  309  323
330  387  391  411  419  510    4   67   78  181  229  230  246  336  365
412  426  447  106  187  463   22  130  164  219  297  353  363  370  455
458  471  513  290  127  345   66   69  169  418  432  508   56  366
280];[4   61   64   74   99  164  169  181   30  128  197   46   70   87
160  192   32   35   85  158  159  193    7   37  177   98  127  150   72
82  176  109  157   48   67  108  112  156];[53   60   61   64  151  165
173  175  188  215  216  223  293  304  307   62  186  219  268  286   58
99  195  260   17   45   46  109  142  158  159  168  229  330  145  161
338   96  108  136  196  204  254   32   35   44   63   70  137  210  269
329  336   72  274  276   79  160   40  135  237  256  119];[38   70   93
115  138  150  159  221  254  274  299  308    8   16   87    7   71  229
287   48   52   58   88   97  106  182  219  257  270  285   60   80   85
180  215  251  279    4   11   63   83  151  171  206  246   34   57  196
288  289   91  128  165   33   69  173  244  245  247  252];[19   29   68
87  112  126  143  151  184  201  203  217  154  117  172  206  236   15
147  229  234  241  170   56  186  205  210   16   89  124  193  213   26
49   93   95  171  182   25   82   34   52   99  106];[47   57   66   73
86   87   92  101  104  108  118  134  139  200  277  282  293   67  241
264   85   88  192  196  256   12   22  199  266  290  303  305   39   78
109   41  147  193  105  173  181  218  255  258  268   31  168  185  289
213  129  137  180  190  260   46  125   45  263  172];[4   44   57   72
77  103  110  136  144  173  184  187  200  233  279  311  330  368   58
74  133  215  265  349  364   55  155  158  167  181  201  202  319   92
204  208  273  298  299   56  145    6   52   81  123  152  342  359  360
22  108  139  143  209  217  258  303  347   68  386   17  134  253  282
327  388  163  247  257  354  307  199   34   85];};


% ADJACENTONLY = 0;requireAll = 1; allowSubstring = 0;
% pat={'ribosome biogenesis GTP'; 'ribosome GTPase'; 'YihA';
'YsxC';'engb';'DNA polymerase I';'pola';'ElbB'};
% pat={'ribosome biogenesis GTP'; 'ribosome GTPase'; 'YihA';
'YsxC';'engb';'DNA polymerase I';'pola';};
% pat={'DMT'; 'VRR-NUC'; 'shikimate';};
% CONTEXT = 2; pat={{'ribosome biogenesis GTP'; 'ribosome GTPase';
'YihA'; 'YsxC';'engb';};{'DNA polymerase I';'pola';};};
% pat={{'ribosome biogenesis GTP'; 'ribosome GTPase'; 'YihA';
'YsxC';'engb';};{'yceI';};};


% genus='Frankia';
% SWITCH = 'for_Idiomarina_Frankia_Pseudoalterom';
%
gs={'igrs_frank_aln';'igrs_frank_casu';'igrs_frank_Allo2';'igrs_frank_B2'
;'igrs_frank_ACN1ag';
```

```
% 'igrs_frank_QA3';'igrs_frank_CcI6';
'igrs_frank_cana';'igrs_frank_BMG5';'igrs_frank_ArI3';
%
'igrs_frank_CeD';'igrs_frank_AvcI1';'igrs_frank_R43';'igrs_frank_nepa';'i
grs_frank_torr';
% 'igrs_frank_umida'; 'igrs_frank_nodulisporulansAgUm';
% % IGRs broken between contigs
% 'igrs_frank_gta';'igrs_frank_tis';'igrs_frank_EI5c';'igrs_frank_KB5';
'igrs_frank_alp';
% % uncertain contexts
% 'igrs_frank_RB7';'igrs_frank_Hr75';
% % not long enough IGRs
% 'igrs_frank_Cj5';
%     };
% SCORES=[12 13 14];
% ns={[1911  2249   284   786  1152  1804  1801   399     4  1609  1924
2174    68   451   624  1011  1324  1850  2019   309   316   391   690
992  1672  1754  1928   534   726   968  1785  1860  2041  2155  2290
2351   379   456   457   658  1059  1253  1323  1475  1526  2042  2048
2111  2142   407   546   623   736   840   930   931  1043  1507  1554
1688  1742  1982  2140  2175  2301   107   286   303   378   562   833
843   871   884   985  1098  1270  1649  1741  1894  1935  2240  2329
722  1686  2212   473   872  2176  1575   607  1903   651   688  1944
691  1468  1511  2087  1246    65   400   718   492  2091  1284  2105
1583  1435  1525  1550   791   179  1140  1333  2275  1154   991  1674
202  1927  1560  2026  1779  2017  1477   374  1426   211  2215  2227
1607   819   925  2138   950  1547    67   734   258   596   891   982
168  2093  2071   249   444   320  2196  2357  1540];[53   121   372
1510   460  1402   277   337   392   955   243   339   370   694   957
1136  1216  1251  1343  1741  1612  1800   179   296   730   810  1319
1474  1530  1761   289   462   589   672   761   841   925   941  1096
1113  1384  1793  1812   170   256   290   535   551   692   713   754
775  1204  1367  1401  1548  1683  1473  1214   714   919  1759  1777
173   580   585  1552   473  1760   767  1512   304  1082   617  1314
840   232   798   417  1485   428   549   225   478  1171  1219   588
29  1001  1685   706  1538   124  1722   220   809     2   348   626
142  1088   803   920  1551  1154   616  1582   266   122  1537  1796
408   837  1311   831  1493  1678   373   376  1101  1494];[1846   190
442   709  1010   274   559  1088   567   745  1708  1727   132   288
355   569   657  1083  1124  1752    71    75   256  1149  1547    31
110   327   397   421   467   513   593   738   747   788   879   951
964  1332  1633  1693  1737  1840    78   235   286   448   663   684
730   812   818   963   972  1028  1288  1342  1615   787  1544  1868
1744   907  1171  1166   923   239   112   398    94  1437  1057  1568
557  1227  1321   111  1813   722  1012   708  1460  1717   456   646
149  1801  1306  1220   447   987    77   798   817   880   641  1232
1350   858   291  1347   208  1507   967    24   178  1090  1565   537
1442   530   959  1225  1277  1697  1122  1165   514   674  1326  1845
1087     6  1337  1349  1790];[1510  1545  1561  1293   133   382   498
1103    51   178   496   504  1547  1627   298  1128  1686   148   280
380   480   985  1339  1649  1682  1698  1778   143   176   396   427
519   587   722   855  1311  1679  1736    54   144   333   503   617
865   880  1037  1073  1197  1294  1326  1412  1570  1584  1650   986
1527   240   155   601  1663   779  1131  1229   302  1466  1559   740
891   803   606   511   263  1575  1195  1233  1461   581  1289   594
462   896  1232   127   616  1005   752   445   288  1507   340  1182
487  1165  1063   269   643   301   284    35   741  1453  1509  1583
287   794   438   776  1718   234  1152];[2094  2805   424   886  2551
```

247 2334 2477 108 1985 1010 1645 166 421 1421 1484 2002
2453 418 1616 1643 1892 2065 2184 2288 2545 225 226 325
420 960 995 1097 1198 1878 1904 2187 2386 2458 190 286
450 626 847 1250 1378 1422 1498 1596 1865 2018 2020 2478
2528 2614 2752 112 158 314 331 348 497 807 1304 1369
1646 2263 2293 2325 251 255 755 933 1023 1211 1621 1667
1761 1762 1801 1931 2062 2253 2287 2383 2415 2420 2434 2454
2720 212 2193 2268 2520 453 2612 2552 1287 1276 2061 1137
1446 464 1781 2280 2565 1591 1387 870 1757 134 2725 1249
890 553 1881 1712 1336 795 1090 2119 1277 1127 1979 1737
851 962 1332 320 2764 2120 877 2022 2757 2006 2058 64
895 328 1366 2368 2146 2609 898 2242 1342 272 1489 738
662 2092 2267 1283 2771 417 768 805 1047 621 411 1599
2515 1174 2483 2656 685 1647 211 326 620 996 1668
1932];[1671 2295 1900 50 1980 385 488 591 1031 1521 1724
527 717 857 900 958 1428 1705 1857 2388 49 627 868
1822 2011 2661 348 440 511 782 1091 1399 1848 2385 357
543 743 869 973 1131 1501 1788 1998 2004 2100 2421 2491
2539 2546 24 262 304 656 676 710 957 1092 1384 1439
1584 2103 2194 2422 2601 3 196 229 512 567 751 801
977 1147 1224 1326 1371 1373 1445 1503 1542 1691 1838 1975
2109 2126 2135 2281 2298 2530 2542 2543 2735 2276 1526 1864
948 1827 570 1823 2658 555 274 1027 2275 1089 2606 1331
186 344 780 343 867 2358 1683 2099 1227 1543 2018 1007
1578 699 2342 51 1875 1294 2188 2081 729 2154 521 950
1320 2545 2519 2405 1538 2502 2555 330 1164 752 1831 237
158 2006 1293 1163 1427 1188 1804 2720 1723 1841 141 2230
2115 231 715 134 114 232 111 1950 1555 764 1132 1751
2001 1943 2184 2262 248 2612 393 2082 1721 2386 1275 1446
2219 2282 205 282 899 2525 2738];[1276 315 575 633 1257
1291 733 1662 489 741 1558 36 300 382 635 743 817
1027 1154 604 879 883 278 504 671 885 1144 1403 1425
1556 1685 1767 9 154 677 819 886 927 956 1115 1221
1407 1455 1572 1724 21 221 325 500 549 794 955 1095
1143 1152 1178 1335 1413 1495 1661 672 1629 1496 279 280
902 455 434 158 1093 103 1732 1255 1296 574 474 1361
47 358 834 1354 1238 547 928 52 108 707 344 814
251 96 327 155 1072 1273 1396 1404 857 752 1670 1039
829 1111 1394 398 1275 841 1378 320 613 1076 1346 1383
1332 1345];[1503 1731 329 1912 1408 1788 212 336 636 1922
303 603 1251 1706 1910 733 951 1306 1600 1828 2006 2278
114 596 693 721 792 892 969 1409 1456 1599 2236 44
465 952 1348 1441 1622 1751 118 135 926 970 1175 1278
1641 2083 2164 9 98 275 442 542 634 1623 1759 1822
1981 2060 2158 49 131 175 222 293 563 594 1014 1056
1063 1246 1254 1820 1940 2070 2132 2156 967 615 639 590
1533 2233 43 404 201 1565 130 1177 315 741 225 573
1929 1774 1787 1867 802 791 720 413 1758 679 750 2076
1948 8 971 1963 1532 1205 1834 1129 379 749 1819 2123
1464 450 593 1762 897 2170 1908 1941 2201 1286 997 401
709 2167 409 1911 75 2235 213 2161 1596 383 1653 53
1149 164 384 642 799 1362 1934 538 1726 614 1272];[217
1693 1205 981 108 244 583 876 1233 219 242 392 557
585 689 1395 1538 1460 178 193 255 316 517 572 769
861 984 1353 1421 1604 1632 1806 56 90 188 195 362
516 559 868 1070 1422 1453 1477 1546 1551 1691 31 46
71 107 116 265 387 403 515 571 605 867 1088 1505
1537 1557 1624 1772 770 1175 468 298 854 1437 262 1090

291 85 1203 488 1154 1737 18 626 1017 475 827 1555
690 463 490 1162 733 339 11 1274 664 1807 447 1153
1738 1078 1696 1454 1512 1635 1354 233 115 1074 410 1456
266 437 1694 1077 939 911 390 982 1026 1528 1612];[1082
350 10 1977 2171 428 965 349 1893 2466 605 746 1289
2456 356 1085 1391 1935 1985 2168 1851 2170 371 427 453
749 910 1252 1364 1411 1622 1762 1832 1884 269 337 826
996 1244 1706 1960 2289 2334 167 430 652 889 932 1316
1320 1635 1668 1684 1894 1930 1983 2040 2074 2431 9 155
243 250 287 532 547 987 1050 1367 1496 1882 2352 2430
2432 1312 2199 63 492 381 1112 1330 834 539 1873 773
589 135 1733 168 1412 1713 1569 141 2325 2418 549 28
227 113 992 1212 114 248 556 2184 451 1724 171 742
1708 901 1246 2026 465 2237 1719 964 2082 1011 733 2263
1114 2167 924 1290 296 558 1763 1508 117 548 1278 2480
2224 1108 425 1025 2028 533 902 379 2164 234 2002 2537
1284 983 422 1698 2477 1621 1799 1820 2161 185 2367 1299
1687 768 1051];[1497 1140 333 1551 1714 1282 1796 74 132
182 1004 1194 72 364 503 850 44 530 564 643 682
789 793 795 1174 1252 1544 115 184 218 230 597 620
648 741 826 903 1030 1123 1157 1168 1204 1681 1711 1804
19 79 122 231 426 573 652 674 720 796 1173 1524
1634 1742 363 1029 1543 47 1069 1344 63 828 1588 1167
726 598 812 1219 430 211 435 1062 1607 270 183 1139
1809 1773 649 1318 1761 416 1158 279 1309 976 140 1016
928 2 634 647 1562 587 690 400 487 881 372 340
1365 1715 1114 593 226 427 637 531 1112 913 208 520
1198 1498 1549 1713 11 403 790];[3827 3857 3832 3947 3825
3941 3808 3925 3747 3970 3858 3770];[404 95 517 129 159 187
349 420 33 203 268 148 367 562 215 298 340 346 401 607 15
58 355 408 351 304 354 85 158 93 283 84 184 293 514 147
57 73 267 438 405 83 94 337 516 53 97];[1516 642 787
233 1608 1735 3651 271 2155 2214 329 595 255 2438 3093
3230 3331 3453 1182 1990 2400 2422 3659 1305 2050 2641 2986
3243 3551 3737 3852 90 168 258 884 1198 1481 1721 3080
167 259 407 564 756 1029 1302 1541 1945 2026 2130 2373
2620 3125 3227 3303 3439 3654 3744 3776 3924 328 470 561
575 929 930 1004 1616 1711 2038 2279 2877 3001 3304 3742
102 136 331 619 641 767 1103 1464 1508 1782 1788 1988
2384 2412 2568 2680 2706 2794 2803 2887 3106 3176 3177 3440
3841 4027 203 295 510 676 905 959 1166 1370 1409 1425
1776 1933 2051 2171 2513 2878 3108 3265 3341 3616 3673 3820
4010 4141 1188 3083 429 3136 4081 1252 960 1027 1028 2976
3683 2348 3321 1987 323 2927 2808 3998 1210 3756 3889 1358
4040 949 1102 1606 2825 613 2631 3961 2235 1934 398 906
1204 1343 3322 3690 1264 2380 4112 666 3921 2045 3160 4022
1618 1000 3002 3536 2177 3363 3433 987 2705 3958 3686 2650
2342 256 3458 4177 1128 2679 1189 1382 630 2465 3922 220
2246 3589 1141 1415 607 749 1011 438 1798 434 633 1442
634 2738 250 112 4097 3342 3388 3667 1607 2778 3178 2995
3229 834 1439 3271 324 2694 795 122 2550 3017 3531 3986
512 1034 1726 1979 2059 2021 1778 75 934 2789 2491 1469
1935 1301 1423 739 1142 529 3126 1869 2914 733 3825 2161
2509 2952 3099 3305 1535 1140 2316 2364 2777 4001 212 355
620 3663 3702 4050 4103 1646 1736 1878 2270 2987 476 831
1751 2575 3004 3960 1743 2574 2967 3863 3908];[2621 998 2203
270 2449 696 946 995 1944 2204 169 2153 2383 624 940
1268 1569 2106 2256 2618 262 314 992 1075 1403 2197 478

554 908 994 1680 1746 2144 109 117 497 627 649 695
813 1408 1532 1570 1592 1894 1987 2384 2476 2537 16 287
415 531 598 967 987 1049 1076 1212 1243 1524 151 195
243 264 321 469 840 939 945 1324 1681 1856 1984 2033
2120 2489 2521 850 1588 33 1470 35 2357 657 1025 518
1983 506 919 1286 2012 1541 893 717 1893 873 1713 2088
2004 724 1214 496 849 207 2036 2435 2035 477 2220 487
2227 107 886 833 740 1980 1626 868 244 1526 1308 234
1061 879 38 1435 258 1916 2505 1792 823 2467 2591 442
991 1361 1804 123 563 485 116 2283 2030 1558 2524 1048
2595 693 1612 1360 1411 1310 1179 727 690 858 1350 2295
821 985 1117 2154 140];[1681 1445 1883 283 640 184 389
535 776 446 844 99 308 848 1301 1916 73 268 475
532 772 1294 1469 1755 25 211 517 592 1040 1270 113
143 445 490 499 721 1256 1283 1508 1717 1884 378 384
597 698 819 1018 1154 1254 1338 1873 21 51 342 348
352 368 797 820 833 1156 1268 1302 1332 1381 1407 1535
1783 1850 1938 641 549 1282 1725 1371 242 440 363 236
1665 360 292 298 1889 1842 609 742 1433 1753 314 1572
835 62 459 1211 86 659 157 110 570 1387 1488 1871
1897 102 1133 377 1099 294 1898 778 1471 932 1872 100
94 518 962 926 1130 6 1630 364 1522 271 1449 1679
1205 1869 1206 1940 1874 281 103 441 1039 1646 432 574
1732 863 1212 1382];[876 306 569 1135 1331 305 307 1314
351 534 835 931 1161 1274 874 1105 1315 5 337 615
663 759 826 845 915 1095 1128 1291 67 116 120 228
266 438 586 669 865 948 1272 685 395 552 536 761
72 559 800 965 331 712 287 1157 230 566 736 343
429 836 725 1247 135 1127 484 33 1098 344 171 1174
772 155 483 89 802 1132 1069 670 555 3 96 664
565 1159 294 38 71 9 35 940 1292 375 765
816];[211 1992 2277 1392 128 704 941 1004 1846 1900 82
155 1262 890 962 1003 1021 1327 1819 1861 2067 2318 2398
967 1811 2003 2038 2063 2187 2366 60 65 402 433 1125
1140 1170 1314 1374 1531 1632 1759 2071 2079 2094 2239 2308
2372 2414 121 293 372 718 750 820 877 1355 1665 1746
1781 1859 2035 2081 2166 2340 2393 1289 740 1050 1258 1230
2388 1332 843 1922 1458 2259 783 408 1460 2088 196 2031
1631 2128 1757 1348 1567 373 1257 1061 2093 1224 198 998
377 474 707 1178 2101 2228 2080 1454 294 2164 950 1860
619 661 1555 1177 163 1174 502 726 2233 1636 1385 1987
2198 666 1562 494 1161 1017 316 1565 1169 982 378 1481
193 1564 148 861 2103 1602 1347 2234 1401 2002 422 236
1324 247 309 804 1055 2261 2346 325 379 450 772 990
2385];[2398 83 1920 1436 1795 1320 1794 1935 2329 752 878
2025 2204 719 1115 1181 1253 1294 1319 1390 1564 1661 1983
2194 503 1430 1966 205 392 486 746 801 936 1199 1476
2255 36 417 519 903 1135 1344 1383 1432 1490 1590 1677
1718 1977 2017 2038 2112 2155 80 115 226 240 307 764
866 985 1027 1119 1527 1639 1748 1778 1855 1856 2201 2297
2338 2193 1060 19 424 672 607 825 1692 2073 1429 1483
1162 1159 935 2172 1921 1503 2298 336 172 2191 163 1507
1816 355 380 2365 1971 118 1720 442 584 2014 158 1763
1762 611 126 689 580 1266 180 1213 1454 104 530 20
1563 1468 2183 2303 1660 540 100 2268 2346 1986 117 1071
916 1914 533 420 1260 177 1882 1932 423 2034 429 1840
553 852 54 1758 418 583 691 79 622 1768 223 1371
1908 1929 2225 998 1662];[2226 1043 1315 2035 1629 840 1916

2362 800 906 1180 1683 2016 2461 167 216 1188 24 771
945 1082 1113 1173 1401 1915 19 722 838 1563 1689 1740
1890 2303 1 508 584 699 1397 1630 2238 2270 2360 2428
88 253 288 991 1167 1174 1377 1394 1433 1434 1589 1996
2251 2420 60 437 446 453 525 830 837 843 848 996
1109 1211 1250 1271 1308 1323 1368 1452 1503 1875 2076 2157
2169 2171 2311 2393 2395 1081 1845 254 2219 1781 582 1429
2425 992 1045 1523 1469 1876 2149 1136 907 2097 1711 2391
1272 2433 447 1690 1463 548 1591 1670 1965 891 727 685
439 1461 1317 1135 706 1865 2317 2106 2304 2133 1685 434
1107 1096 1153 273 747 1896 799 2293 1612 1004 1846 653
238 2213 237 708 1749 1759 2447 1710 1618 1206 1338 1420
2166 740 1632 654 1470 1528 1726 1912 2100];[103 183 486
1352 886 901 58 613 1123 1757 792 946 1442 1906 287
1117 1125 1837 97 413 728 775 1471 1686 1862 274 516
555 849 944 1150 1189 1313 1554 1568 1728 1920 57 185
295 313 440 570 709 790 816 904 1223 1546 1619 1707
1718 1841 1856 335 1434 1640 708 707 490 561 1145 666
1776 1458 735 747 231 924 545 656 1630 229 1115 26
400 1284 1169 1054 1381 487 1734 1215 784 368 568 525
1152 727 1843 1459 302 1025 98 1202 435 662 328 1073
292 1330 902 459 1204 1449 1118 992 598 184 802 738
1893 860 1175 136 515 564 442 461 890 1013 1573 805
1468];[716 2040 1121 1379 2045 455 885 2020 265 1170 1494
126 489 1181 1331 1488 1736 376 1178 2007 309 948 1442
1513 1789 1855 2031 57 526 828 1086 1300 1312 1336 1497
1674 1887 1927 128 180 354 547 668 710 954 969 1051
1118 1241 1266 1538 1581 1965 1981 1982 2043 178 1260 54
76 713 1564 403 1196 993 1257 575 2 757 1069 1653
1281 1916 2060 2039 542 52 795 237 1791 1928 454 168
502 653 89 1592 1443 1203 730 1333 1234 1311 421 1818
785 1853 947 729 1388 1403 601 1141 1754 1805 858 1652
1259 896 233 164 857 1199 1432 1792 1525 1809 179 1103
1299 886 205 528 796 1237];[654 221 976 494 987 20
188 384 925 57 710 717 321 505 606 645 1131 23
184 257 417 769 995 1065 25 605 788 803 887 1068
1248 14 58 75 138 176 607 797 894 924 1009 1123
1176 1184 26 70 137 226 310 918 955 1033 1240 1241
1179 1182 178 781 895 555 1049 183 110 689 375 258
564 720 385 24 47 1094 993 48 897 165 390 733
643 1084 383 318 120 923 418 187 306 1014 18 175
812 809 28 156 770];[508 27 247 60 368 1101 8
23 407 702 704 1185 1319 1713 1787 96 272 1550 1758
1762 2300 28 248 626 1200 1417 366 694 964 1100 1213
1247 1284 1619 1971 2028 209 322 345 651 678 979 1018
1462 1716 1717 1828 2224 2290 25 240 571 621 695 703
946 1065 1193 1246 1692 1702 1706 1761 1810 1938 1981 46
47 106 126 152 199 243 622 648 696 709 890 988
1040 1382 1384 1408 1609 1634 1836 2296 2309 987 745 1343
202 597 2089 2177 741 677 645 346 1478 304 458 7
2305 530 1088 38 888 574 1137 614 348 421 921 1092
184 2184 2158 1449 2299 1198 1700 644 178 449 33 989
2241 779 386 748 1882 1549 1283 1231 1916 285 2027 1799
1460 619 1727 2268 686 1688 1856 2295 1026 680 317 1616
259 529 780 1257 755 1418 261 642 899 945 900 26
146 1023 1453 2250 2263];[1723 790 1834 1873 1803 438 1134
1335 1348 105 799 896 1612 398 692 1288 1660 1819 77
327 376 670 1167 1179 1336 1346 1370 1389 1659 1679 1686

```
625   707   786   950  1642  1654   446  1147  1390  1697  1726  1731
107   330   561   633   676   934  1050  1250  1316  1408  1591  1627
1724  1737  1837  1839  1861  1228  1570   906  1537   572   363   737
628  1486  1019  1450  1025  1028   290   427    57   771  1328   573
1114   848  1353   136   495   306    75   935  1682  1634   920  1337
1725   738  1533   257   905   269  1487  1105  1363    54  1796  1747
912  1090  1227  1213   961  1180  1347   895  1380   840   701   999
1704];};
% SCORES=[5 6 8 9 10];
% ns={[4    35    68   105   126   168   202   258   309   316   364
379   389   390   400   403   407   451   456   457   473   492   519
546   562   590   592   595   607   624   651   687   691   698   722
726   786   791   818   833   872   887   891   914   925   929   930
931   985   992  1011  1024  1035  1043  1058  1059  1098  1108  1140
1216  1222  1245  1246  1253  1284  1299  1323  1333  1372  1383  1398
1435  1451  1468  1475  1477  1478  1511  1560  1575  1583  1590  1607
1609  1630  1643  1649  1669  1674  1686  1688  1717  1730  1742  1751
1754  1779  1801  1834  1858  1860  1903  1908  1924  1928  1935  1944
1982  2017  2026  2087  2093  2142  2150  2165  2175  2176  2203  2206
2227  2234  2249  2251  2275  2293  2350  2351  2354   288   697   742
753   829   883   919  1044  1047  1102  1426  1473  1593  1844  1998
2023  2134  2155  2255  2332    65    67   161   179   254   279   281
341   387   397   399   581   584   589   596   608   718   819   849
981  1154  1250  1307  1324  1348  1356  1399  1526  1539  1547  1554
1579  1580  1741  1778  1786  1797  1805  1927  2020  2091  2105  2111
2138  2355   249   734   950   991  1114  1525  1955  2019  2071  2130
2212   899];[2    29    53    56    61   114   124   142   169   173
186   228   231   277   278   289   296   304   339   356   370   411
424   425   426   428   473   478   547   549   580   585   588   616
617   626   667   694   706   714   723   744   798   803   804   809
816   840   844   871   919   920   925   944   957   983  1017  1023
1087  1154  1199  1204  1214  1219  1314  1319  1343  1370  1399  1453
1473  1474  1485  1512  1537  1551  1552  1559  1560  1566  1612  1624
1640  1653  1659  1670  1677  1683  1698  1714  1741  1759  1760  1761
1777  1796  1799  1800  1812    69   107   286   347   490   727   739
775   846   958  1075  1209  1402  1486  1654    59   111   212   218
225   375   388   462   467   501   546   629   703   704   721   754
761   822   848   854  1001  1006  1082  1088  1299  1367  1449  1646
1651    22   167   220   329   577   941  1171  1685  1792   417   713
767   934];[24    47    71    75    94   110   111   112   132   149
159   178   180   182   186   189   190   286   291   369   393   398
403   405   426   448   450   456   475   504   530   557   558   569
645   646   648   674   681   709   722   738   747   748   787   788
798   810   812   880   907   908   923   953   959   963   972   987
1012  1032  1057  1083  1090  1099  1105  1130  1166  1171  1220  1225
1227  1232  1296  1306  1314  1321  1330  1331  1332  1347  1349  1350
1367  1442  1461  1475  1480  1538  1541  1544  1564  1568  1604  1607
1633  1693  1705  1707  1708  1715  1717  1752  1757  1787  1795  1802
1806  1808  1813  1840  1843  1860  1868  1916    32    67   105   402
438   537   818   875  1124  1293  1342  1698     6    19    45    77
185   324   339   404   442   498   543   560   617   663   684   708
796   879   951   964  1035  1088  1138  1193  1204  1288  1290  1322
1437  1441  1547  1573  1727  1744  1825  1915   229   239   549   559
578   703   729   932   967  1150  1460  1565  1571  1801    79];[6    35
127   133   135   143   148   176   181   240   269   288   298   301
302   323   396   427   438   445   466   496   502   511   519   533
581   582   587   598   601   606   640   644   671   688   740   741
752   779   810   858   892   893   937   954   971   974   985   986
```

1005 1037 1063 1064 1109 1128 1137 1138 1165 1193 1195 1211
1229 1232 1233 1249 1295 1296 1320 1339 1389 1404 1449 1461
1466 1468 1491 1507 1527 1547 1559 1567 1575 1610 1649 1650
1663 1682 1686 1745 83 105 117 134 247 284 325 394
709 728 786 911 953 1060 1213 1419 1509 1541 118 136
155 169 212 248 263 287 333 380 401 421 462 487
504 534 594 597 681 777 795 891 896 1031 1073 1074
1107 1131 1147 1157 1182 1224 1253 1263 1294 1452 1493 1584
1609 1627 1678 1714 1761 1771 616 643 803 1258 1570 1698
69 722];[7 37 43 45 46 133 134 138 158 190 211
212 225 230 247 251 255 258 272 308 316 320 325
328 331 355 395 418 421 423 424 449 453 464 482
542 580 607 626 662 719 738 755 768 794 807 812
870 872 877 895 898 913 914 951 962 981 991 1023
1026 1037 1047 1058 1083 1091 1097 1129 1137 1193 1250 1276
1287 1288 1315 1320 1336 1366 1376 1387 1399 1402 1408 1421
1422 1446 1484 1537 1543 1550 1591 1615 1616 1622 1670 1757
1760 1781 1854 1866 1878 1881 1883 1887 1903 1907 1930 1960
1979 1985 1994 2000 2018 2022 2061 2093 2103 2120 2146 2174
2184 2188 2218 2242 2247 2260 2264 2268 2280 2296 2334 2393
2399 2416 2434 2446 2478 2480 2487 2517 2519 2520 2551 2552
2577 2600 2612 2614 2637 2661 2695 2773 2782 12 121 417
545 620 717 806 853 1071 1160 1177 1188 1206 1249 1435
1482 1737 1848 1976 2011 2070 2102 2119 2141 2183 2320 2565
2780 108 112 220 239 286 313 341 365 386 396 400
553 633 656 701 795 890 996 1041 1066 1127 1198 1352
1379 1453 1573 1599 1645 1648 1707 1712 1749 1872 1922 1943
2020 2058 2065 2092 2126 2181 2193 2270 2288 2339 2386 2433
2453 2477 2515 2564 2635 2674 2697 2752 2764 329 363 382
1090 1197 1498 1644 1753 2287 2289 2368 2757 2771 639 1652
2319 2347];[26 29 43 49 50 111 114 134 141 176
185 186 196 229 231 237 242 248 261 274 325 330
343 371 385 439 469 470 483 484 491 493 504 521
522 527 567 588 591 616 628 636 699 704 715 729
752 756 765 767 780 837 849 859 867 869 901 948
950 1007 1027 1032 1035 1038 1089 1092 1131 1147 1163 1164
1188 1207 1223 1225 1227 1316 1331 1337 1395 1399 1400 1427
1428 1444 1485 1501 1523 1526 1537 1538 1544 1545 1546 1574
1578 1604 1644 1673 1677 1683 1694 1699 1701 1702 1705 1706
1721 1724 1799 1804 1823 1827 1837 1847 1864 1875 1950 1961
1976 1998 2001 2004 2018 2063 2081 2082 2099 2103 2113 2115
2125 2135 2154 2184 2188 2212 2213 2230 2237 2262 2275 2276
2283 2295 2298 2305 2337 2342 2358 2460 2461 2495 2501 2519
2525 2531 2539 2543 2585 2596 2606 2612 2618 2658 2737 25
107 427 433 648 741 826 891 944 1021 1129 1160 1269
1277 1335 1358 1394 1525 1582 1698 1723 1892 1979 1982 2126
2478 2500 2513 2545 2581 14 24 105 232 278 304 344
386 395 453 476 543 551 570 586 625 673 692 743
797 807 810 833 900 973 1091 1098 1103 1125 1270 1293
1320 1351 1502 1521 1541 1543 1832 1841 1857 1943 2006 2009
2011 2026 2164 2190 2257 2372 2405 2546 2555 2590 2601 2605
2719 51 338 393 555 764 1294 1475 1831 2101 2278
868];[12 49 103 132 154 155 157 158 251 278 279
280 300 315 316 320 325 327 358 398 408 434 455
465 474 475 489 490 491 499 504 514 547 575 604
635 638 671 672 697 743 767 768 794 814 819 827
829 857 873 879 883 902 928 955 956 968 969 1027
1039 1041 1042 1062 1066 1072 1076 1093 1094 1111 1127 1144

1178 1228 1238 1246 1255 1260 1261 1262 1273 1291 1296 1354
1357 1359 1360 1361 1370 1376 1396 1407 1474 1480 1496 1555
1570 1572 1629 1662 1672 1715 1720 1732 48 82 99 101
406 428 456 901 1065 1081 1504 1556 1650 47 96 120
221 326 407 409 420 500 545 574 613 617 752 844
886 1153 1154 1195 1221 1275 1327 1375 1404 1499 1567 1569
1609 1667 1670 244 344 348 642 707 885 1403 630
1733];[8 18 43 57 75 130 135 176 225 314 315
327 329 336 379 380 383 399 401 404 413 438 441
443 450 467 470 482 522 542 573 590 593 594 596
636 679 720 733 736 741 744 749 750 802 828 892
897 932 951 952 960 967 968 985 1056 1129 1136 1177
1205 1246 1249 1251 1254 1286 1306 1359 1362 1374 1380 1406
1408 1409 1416 1456 1463 1470 1503 1504 1532 1538 1565 1567
1599 1600 1618 1641 1676 1689 1706 1731 1758 1787 1788 1804
1819 1828 1834 1867 1868 1908 1912 1922 1929 1933 1940 1942
2016 2034 2076 2077 2083 2104 2123 2128 2132 2152 2154 2156
2158 2161 2170 2201 2221 2233 2234 102 259 412 427 523
574 641 665 682 752 970 1261 1349 1578 1606 1733 1742
1879 2075 2155 2167 2199 2237 2252 4 248 274 389 541
568 615 721 809 971 979 1184 1336 1360 1368 1596 1622
1653 1675 1681 1700 1762 1774 1910 1911 1941 1953 2178 2235
2278 98 182 201 435 473 639 709 791 810 997 1110
1117 1156 1175 1272 1464 1533 1623 1803 2236 214 553];[11
14 17 53 74 85 105 108 116 130 188 193 195
219 226 242 262 291 298 333 410 468 470 516 517
521 554 559 585 586 605 626 632 640 664 689 718
729 769 770 861 868 876 911 912 934 942 950 978
984 1009 1017 1042 1046 1049 1074 1078 1089 1090 1094 1104
1131 1154 1175 1181 1203 1219 1221 1249 1252 1272 1307 1344
1390 1395 1421 1422 1437 1456 1460 1468 1512 1546 1551 1553
1554 1555 1604 1635 1642 1669 1686 1709 1738 1772 1797 1807
35 115 250 445 576 611 949 1077 1216 1274 1343 1354
1363 1682 1736 1743 18 166 233 268 287 305 354 356
447 463 488 490 523 526 572 636 690 733 824 842
905 909 936 958 981 1026 1055 1162 1163 1321 1328 1336
1348 1353 1381 1453 1459 1462 1495 1505 1537 1544 1584 1658
1696 71 339 362 475 571 827 854 1103 1737 1127];[5
10 28 57 63 65 113 117 131 140 141 159 168
179 224 225 227 234 243 250 296 349 350 368 371
379 381 382 403 425 427 428 451 484 492 497 539
558 563 589 647 660 733 742 744 749 773 782 826
847 889 902 952 986 987 988 992 996 1000 1025 1026
1028 1035 1057 1082 1085 1086 1124 1197 1246 1278 1284 1289
1290 1312 1329 1330 1344 1411 1412 1430 1490 1508 1575 1628
1635 1693 1706 1708 1724 1731 1733 1796 1799 1820 1845 1883
1929 1935 1939 1960 1966 1977 1983 2002 2012 2026 2038 2056
2074 2082 2084 2131 2151 2167 2168 2171 2199 2224 2239 2263
2304 2319 2334 2353 2359 2397 2403 2409 2416 2419 2432 2525
2542 21 89 105 130 498 700 714 741 761 820 834
850 1103 1186 1256 1441 1577 1832 1851 1930 1942 2081 2099
2289 2477 2528 13 114 139 171 273 282 313 465 467
605 652 684 731 753 799 802 812 874 924 971 1009
1063 1106 1112 1163 1170 1212 1218 1269 1297 1365 1556 1569
1619 1622 1636 1698 1719 1763 1840 1906 1936 2028 2066 2075
2184 2202 2312 2368 2369 2418 2431 2452 2454 135 247 549
825 901 963 1108 1391 2164 2325 2537 833 1131 1802];[2 14
19 27 47 72 79 115 121 122 130 132 181 182

211 231 243 250 270 278 329 339 348 363 364 372
416 426 427 429 430 435 436 453 482 487 519 573
587 593 618 634 637 640 647 649 682 690 720 726
730 732 753 754 789 793 812 825 828 897 903 928
955 970 976 1016 1029 1030 1035 1062 1069 1112 1113 1114
1115 1139 1140 1159 1162 1167 1181 1194 1198 1214 1270 1309
1318 1319 1344 1383 1387 1424 1426 1500 1517 1524 1562 1573
1588 1607 1608 1634 1681 1701 1711 1715 1735 1744 1761 1773
1794 1 3 8 176 274 275 437 447 719 725 796
811 877 995 1202 1507 1513 46 63 140 183 279 509
624 689 795 826 850 881 913 920 931 1042 1096 1141
1157 1220 1235 1259 1448 1477 1529 1543 1549 1564 1713 1714
1723 152 178 226 598 1158 1219 1251 1436 1579 1809 1656
1795];[3702 3742 3795 3802 3808 3822 3825 3826 3858 3895 3919
3925 3933 3941 3946 3959 3973 3977 3989 3704 3784 3832 3968
3969 3970 3976 3982 3747 3956];[11 31 33 58 73 85 88
93 95 158 159 203 268 283 285 297 298 303 304 337 349 351
366 399 403 408 438 514 515 555 578 610 84 273 284 378 458
470 540 609 57 187 196 293 333 354 360 372 562 597 71 147
265];[10 22 57 75 80 90 102 112 120 122 132
167 216 233 250 256 259 317 323 324 329 354 356
420 428 429 453 470 476 512 563 581 595 613 620
636 666 729 748 760 768 795 831 855 884 904 906
909 910 931 934 949 967 987 1000 1004 1011 1027 1028
1054 1058 1102 1128 1132 1142 1166 1182 1192 1204 1206 1210
1236 1252 1264 1302 1343 1344 1358 1368 1370 1415 1423 1439
1460 1483 1516 1541 1606 1607 1608 1618 1686 1703 1733 1735
1778 1798 1807 1862 1869 1892 1931 1932 1964 1980 1987 2000
2021 2022 2038 2045 2050 2059 2084 2092 2139 2140 2170 2177
2187 2235 2245 2246 2270 2308 2316 2347 2361 2368 2379 2380
2384 2398 2400 2438 2465 2491 2510 2550 2556 2566 2569 2577
2596 2631 2679 2680 2681 2694 2701 2705 2753 2778 2789 2808
2825 2877 2878 2893 2911 2914 2916 2927 2976 2987 2994 2995
3001 3013 3017 3033 3066 3083 3084 3102 3113 3125 3136 3139
3160 3185 3243 3252 3265 3305 3321 3322 3341 3342 3349 3360
3388 3433 3440 3453 3458 3496 3531 3536 3551 3589 3602 3616
3652 3654 3667 3682 3683 3690 3756 3841 3890 3920 3921 3924
3958 3961 3968 3986 3998 4034 4040 4049 4074 4081 4090 4097
4112 4141 4177 4180 283 300 355 494 508 524 602 933
1069 1134 1188 1198 1382 1389 1394 1508 1743 1864 1951 2514
2539 2588 2692 2771 2829 2913 2945 2996 3317 3405 3406 3593
3620 3908 4083 4155 152 165 190 198 244 255 261 290
303 398 434 520 533 607 634 739 749 756 800 883
960 998 1003 1029 1034 1053 1189 1215 1336 1339 1367 1425
1469 1636 1726 1788 1804 1845 1878 2182 2184 2283 2346 2362
2422 2475 2632 2647 2668 2671 2704 2800 2870 2924 3002 3003
3080 3091 3093 3154 3178 3193 3269 3271 3280 3353 3354 3363
3371 3400 3606 3647 3686 3906 3922 3964 4022 4136 4156 4179
220 238 286 308 407 519 558 641 834 1141 1934 2348
2513 2591 2641 2650 3105 3229 3331 3601 3820 3889 4003 1616
1645 2525 2779 3478 3888];[12 33 35 49 56 94 107
108 125 151 177 207 213 258 263 264 266 270 287
314 321 322 442 455 465 477 480 485 506 509 554
563 618 622 627 649 657 693 695 699 717 724 730
781 813 840 841 886 891 893 905 908 919 938 939
940 969 991 992 997 998 1025 1048 1117 1126 1229 1240
1286 1308 1322 1384 1403 1444 1449 1470 1526 1541 1544 1551
1558 1569 1570 1588 1592 1602 1609 1611 1612 1649 1680 1721

1746 1755 1789 1792 1844 1855 1856 1873 1894 1916 1918 1983
1987 1990 2004 2005 2016 2030 2036 2054 2098 2120 2203 2204
2220 2227 2256 2264 2283 2299 2358 2414 2435 2447 2461 2468
2474 2489 2505 2521 2524 2526 2538 2595 2596 2618 2621 144
150 178 224 305 370 415 478 629 697 808 995 1144
1401 1556 1565 1583 1626 1637 1997 2007 2067 2406 109 116
195 498 559 587 696 728 740 749 794 833 839 849
850 868 875 933 948 957 986 987 1061 1075 1106 1179
1214 1243 1360 1406 1435 1437 1652 1713 1944 1945 1980 2012
2035 2042 2088 2153 2260 2310 2385 2410 2547 2576 2591 38
187 244 371 422 471 496 518 823 873 1076 1095 1310
1361 1635 1893 2357 2377 2384 2409 2467 2476 2540 454 879
2449 2544];[9 25 47 48 73 87 100 102 103 157
177 186 199 227 236 242 257 271 283 292 308 310
314 348 352 360 363 376 378 384 389 439 440 445
490 518 529 532 535 548 549 597 609 611 641 659
676 688 698 742 778 819 835 836 837 895 901 926
927 1040 1050 1099 1102 1108 1130 1156 1182 1205 1206 1221
1254 1268 1283 1320 1336 1363 1371 1387 1413 1469 1471 1488
1497 1523 1528 1559 1572 1574 1593 1613 1665 1669 1679 1681
1701 1725 1732 1753 1783 1822 1825 1842 1850 1871 1873 1897
1898 1934 1935 1939 1940 66 135 324 330 373 797 1009
1018 1044 1064 1071 1092 1214 1310 1328 1522 1660 1686 1796
1799 1841 1878 1977 2008 6 13 30 32 86 94 99
109 110 113 298 344 446 459 570 697 721 756 767
850 865 943 978 992 1017 1029 1035 1133 1134 1211 1282
1302 1368 1433 1449 1520 1646 1666 1868 1869 1872 1883 1889
1916 154 268 441 660 962 1039 1100 1381 1487 1570 1778
607];[33 38 71 72 86 89 96 118 120 135 155
230 246 294 306 307 331 343 347 351 375 395 396
443 483 484 534 536 555 559 562 565 566 626 634
663 664 712 725 728 761 772 800 836 865 874 940
994 1010 1061 1069 1076 1095 1105 1131 1135 1157 1159 1161
1174 1195 1215 1219 1234 1247 1274 1291 1292 1314 1315 1330
1332 3 496 613 671 876 1064 1077 1146 1156 1173 1324
4 13 14 111 171 217 287 305 344 460 511 535 552
569 631 668 672 674 689 714 825 835 845 909 961
1096 1127 1132 1177 1207 1221 1240 1267 1331 60 378 685
711 736 801 910 931 965 1027 929];[4 26 128 148 163
179 196 248 345 368 373 377 379 423 448 449 458
483 502 504 586 596 619 634 666 704 707 718 726
740 746 767 770 783 818 832 843 871 890 950 963
982 998 1003 1017 1050 1061 1091 1117 1138 1161 1174 1177
1178 1193 1224 1230 1244 1258 1261 1262 1289 1319 1332 1348
1349 1374 1429 1447 1454 1458 1460 1481 1508 1531 1564 1567
1570 1603 1621 1631 1665 1673 1719 1759 1787 1811 1819 1846
1852 1861 1881 1900 1922 1987 1992 2026 2067 2079 2081 2088
2093 2101 2103 2163 2166 2175 2179 2186 2187 2198 2228 2236
2239 2256 2259 2269 2329 2334 2340 2388 2398 2413 152 180
371 401 477 717 803 841 1004 1055 1184 1241 1298 1347
1500 1591 1604 1675 1691 1764 1939 1988 2092 2294 32 33
195 198 391 422 435 474 661 669 678 691 808 842
851 859 908 919 964 1077 1169 1257 1452 1516 1579 1602
1757 1781 1817 1860 1950 2003 2031 2080 2105 2113 2128 2143
2144 2158 2234 2273 2277 2359 2385 2386 2404 25 159 294
340 403 408 642 748 861 990 1176 1317 1415 1416 1555
1565 1636 2041 2302 2393 362 383 2346 2394];[18 19 20
21 27 36 79 80 106 118 126 163 177 194 197

204 211 281 336 380 383 417 423 424 442 454 486
506 512 530 533 572 580 607 611 641 653 655 672
689 719 778 856 866 874 878 916 935 964 985 1026
1027 1085 1115 1119 1140 1159 1162 1199 1240 1243 1260 1263
1266 1279 1299 1388 1390 1422 1429 1436 1454 1468 1483 1491
1503 1507 1510 1530 1557 1558 1582 1654 1660 1689 1692 1718
1720 1762 1763 1788 1794 1801 1807 1816 1841 1852 1868 1882
1903 1914 1921 1929 1935 1963 1971 1974 1983 2006 2014 2017
2023 2034 2064 2073 2112 2122 2158 2170 2172 2183 2191 2193
2268 2297 2298 2346 2394 23 58 381 399 525 761 889
1183 1278 1320 1352 1392 1562 1584 1645 2021 2038 2094 2361
104 117 158 171 172 222 240 303 316 350 355 450
540 573 584 752 812 825 922 938 963 1060 1082 1106
1190 1201 1258 1430 1444 1490 1535 1537 1586 1740 1777 1885
1894 1932 1936 1986 2102 2195 2219 2243 2287 2294 2305 2358
2365 83 157 429 721 887 1294 1316 1338 1349 1371 1563
1778 1787 1920 132 216 391 1662];[24 27 78 96 237
238 254 263 273 324 340 362 383 400 453 462 493
494 500 502 508 548 572 585 601 653 654 685 708
727 787 799 830 837 838 840 869 877 891 896 907
908 945 992 1025 1045 1081 1091 1096 1099 1102 1107 1122
1135 1136 1153 1180 1183 1250 1269 1271 1272 1282 1306 1315
1317 1323 1334 1338 1377 1385 1397 1401 1433 1434 1461 1463
1469 1523 1524 1551 1564 1587 1591 1629 1630 1676 1689 1710
1711 1781 1807 1845 1846 1876 1896 1913 1916 1923 1931 1932
1965 1966 1974 1978 1994 2035 2097 2101 2106 2133 2149 2169
2219 2224 2238 2270 2275 2293 2304 2317 2331 2344 2360 2361
2391 2425 2433 2440 2447 2455 2461 83 151 521 699 761
764 1140 1173 1191 1199 1238 1432 1445 1618 1673 1753 2009
2032 2036 2165 2213 2281 2289 2426 44 86 88 91 188
217 236 265 345 406 439 447 452 525 621 747 749
792 801 836 848 981 1082 1121 1201 1240 1378 1381 1486
1578 1617 1670 1685 1690 1726 1749 1949 1963 2016 2076 2077
2171 2172 2180 2209 2251 2276 2316 2326 2362 2374 2398 2428
19 388 582 740 1429 1742 2210 2303 2437 733 2105];[24 26
89 98 103 182 229 259 274 292 294 302 305 328
335 368 381 400 406 435 454 459 477 478 486 487
490 508 514 555 561 598 626 656 666 667 683 700
707 708 709 728 735 738 743 747 784 816 834 870
875 877 884 912 924 935 944 949 956 982 992 1025
1073 1145 1152 1169 1175 1189 1204 1215 1303 1304 1313 1318
1352 1375 1376 1434 1442 1449 1452 1468 1471 1482 1513 1546
1617 1640 1665 1694 1728 1734 1757 1776 1808 1837 1841 1855
1862 1899 1901 1905 1906 1936 43 100 185 243 288 306
556 639 737 755 792 958 1009 1055 1181 1475 1579 1619
1736 1800 1847 58 126 167 200 231 265 321 329 370
473 545 568 779 889 1000 1125 1218 1330 1459 1536 1554
1568 1630 1695 1717 1843 1869 1922 295 585 843 917 1077
1115 1116 1202 1284 1381 1595 1918 1155 1209 1784];[178 265
403 455 527 528 530 534 535 541 542 548 551 554
557 567 575 576 584 586 588 590 592 598 599 601
618 634 636 637 643 650 651 653 659 660 666 668
669 670 678 679 681 690 697 699 710 711 713 716
717 722 732 740 743 749 750 753 757 769 772 786
794 795 796 799 801 803 804 811 812 819 821 823
827 828 830 831 837 841 849 857 858 859 865 866
870 875 876 882 886 893 896 899 904 914 931 932
933 947 948 954 955 959 962 967 969 987 990 993

```
994   996  1003  1005  1008  1023  1032  1033  1234  1331  1379  1564
1581  1674  2040  2043   531   545   569   675   723   774   781   824
867   871   874   897   903   909   910   945   966  1496  1592   126
526   564   572   596   630   631   632   663   665   687   700   714
724   726   729   730   731   736   742   746   751   759   766   782
785   791   792   809   814   817   834   864   877   881   887   888
894   946   956   971   981   988   991  1004  1006  1024  1034  1196
558   605   622   671   802   885   906   975   982   989  1257   529
546];[20    26    47    48    98   110   120   165   178   187   188
199   221   222   257   258   282   295   317   321   372   375   383
385   390   397   398   418   484   504   555   564   596   605   607
617   643   645   654   689   720   731   769   781   785   788   820
841   846   847   895   897   899   918   925   976   987   993   995
1033  1049  1064  1065  1087  1094  1102  1179  1180  1182  1192   162
340   456   523   686   691   717   842   848   924   935   997  1085
1133  1176    24    30    34   138   157   176   183   226   310   318
351   417   478   619   680   694   719   733   770   803   810   955
956  1098  1124  1184   218   519   608   732   306   339   562];[2     23
27    28    38    47    96    98   111   126   141   152   178   184
189   202   215   247   248   261   272   281   284   285   304   325
350   366   368   373   396   403   404   415   421   445   449   458
527   568   571   574   579   597   614   619   622   644   651   674
677   680   702   703   741   745   747   756   776   780   856   900
937   987   989  1017  1018  1052  1088  1091  1092  1100  1135  1137
1152  1172  1193  1197  1198  1200  1204  1233  1248  1257  1261  1277
1283  1284  1322  1343  1361  1384  1406  1407  1408  1454  1460  1462
1478  1549  1609  1616  1620  1634  1646  1673  1700  1705  1706  1716
1764  1787  1799  1843  1855  1856  1938  1949  1981  2000  2069  2089
2139  2144  2158  2177  2184  2214  2224  2290  2295  2296  2299  2300
2305    48   136   164   362   476   479   816   851  1101  1184  1339
1373  1466  1533  1539  1870  2268  2309     5     7    18    25    26
44    60   162   316   317   346   348   386   491   508   517   518
645   695   704   730   732   748   826   888   890   921   946   963
979  1182  1186  1319  1381  1382  1550  1663  1694  1727  1810  2027
2030  2035  2048  2141  2163  2194  2213  2242  2250  2263  2294    33
34   160   243   530   613   678   694   755  1293  1391  1619  1828
1867  1916  2282   433   939  1418  1730];[16    31    75    77    95
98   105   107   177   238   249   257   269   290   304   306   316
330   363   374   376   398   427   446   539   572   625   628   633
642   675   676   689   692   701   707   738   756   759   809   848
884   888   895   897   898   905   906   950   961   999  1019  1025
1066  1074  1105  1145  1147  1177  1228  1250  1288  1328  1335  1336
1353  1428  1440  1459  1486  1487  1533  1537  1552  1570  1588  1607
1634  1637  1642  1659  1660  1667  1668  1672  1682  1689  1692  1731
1733  1747  1796  1803  1818  1837  1839  1844  1852   153   189   214
418   435   544   647   677   752   868  1168  1292  1300  1332  1348
1374  1723  1737    57    66    76   243   296   470   540   547   552
772   785   789   883   887   891   912   928  1114  1287  1316  1337
1380  1450  1587  1592  1639  1697   136   573   737   771   799   890
920   935  1227  1324  1596  1686  1725  1726];};
% SCORES=[1 2 3 4];
% ns={[4     9    16    68    85   102   107   126   141   179   197
211   258   284   308   316   379   387   391   399   407   444   451
456   534   540   546   562   609   623   624   658   722   739   742
786   883   930   931   992  1119  1152  1233  1246  1253  1300  1469
1475  1478  1507  1526  1554  1609  1635  1668  1688  1724  1730  1754
1785  1801  1804  1860  1863  1891  1894  1897  1903  1911  1928  1935
1982  2019  2048  2065  2097  2142  2158  2165  2175  2203  2206  2249
```

2290 2355 3 15 35 202 249 312 320 457 492 509
702 726 896 1011 1044 1086 1163 1289 1323 1333 1397 1547
1649 1672 1695 1738 1742 1850 1924 1941 2000 2041 2042 2303
67 303 309 833 1059 1356 1593 1701 2140 2155 2174 2227
61 105 168 288 384 403 440 473 519 590 595 655
676 697 753 818 829 872 914 919 925 929 985 1004
1024 1035 1047 1098 1107 1108 1140 1216 1222 1279 1299 1398
1426 1435 1468 1473 1509 1511 1575 1583 1590 1607 1643 1686
1736 1837 1844 1858 1929 1944 1998 2023 2087 2093 2099 2134
2150 2219 2234 2251 2255 2275 2293 2332 2354];[16 53 56
69 121 170 179 220 266 277 337 339 370 372 373
408 409 486 490 510 535 551 554 589 616 651 672
713 714 775 803 804 841 873 925 944 955 957 958
1006 1096 1117 1136 1204 1216 1311 1343 1367 1368 1440 1474
1510 1512 1530 1537 1548 1565 1598 1602 1612 1640 1659 1680
1684 1713 1741 1749 1750 1758 1774 1796 1800 83 142 174
232 243 256 304 374 692 694 810 863 976 1001 1023
1391 1493 1516 1653 1677 269 296 392 693 761 1017 1058
1251 1402 1484 1812 2 29 85 114 124 173 228 231
233 286 289 424 425 428 473 478 487 549 585 727
739 744 798 844 846 848 871 920 999 1075 1154 1209
1214 1313 1314 1370 1399 1453 1473 1485 1486 1540 1551 1552
1559 1560 1566 1606 1624 1654 1670 1683 1698 1759 1760 1777
1799];[31 32 71 75 132 190 239 274 286 288 289
327 398 442 448 504 512 532 657 674 701 709 748
788 796 818 907 959 972 987 1010 1012 1028 1060 1068
1090 1094 1119 1122 1124 1130 1150 1166 1171 1176 1241 1337
1341 1342 1349 1363 1367 1442 1458 1461 1564 1603 1615 1633
1664 1683 1716 1717 1722 1744 1752 1813 1846 18 77 97
110 123 197 355 397 409 467 537 559 663 684 722
747 810 812 895 908 923 954 1037 1083 1163 1264 1277
1328 1493 1697 1708 1774 1868 37 513 567 569 951 1046
1193 1514 1547 1790 24 47 67 94 105 112 152 178
180 189 393 402 403 405 426 438 450 456 468 475
530 542 557 641 645 646 648 681 730 738 787 817
875 963 1032 1105 1220 1225 1227 1232 1293 1296 1314 1321
1330 1332 1347 1466 1475 1544 1568 1654 1693 1698 1707 1727
1757 1787 1795 1802 1806 1840 1843 1860 1916];[51 83 120
127 130 133 140 178 257 259 280 301 315 379 382
410 427 440 480 498 503 521 582 587 601 606 626
640 710 714 721 722 728 794 795 849 855 865 866
880 891 985 1011 1031 1037 1128 1137 1140 1157 1269 1311
1325 1339 1456 1473 1510 1545 1547 1561 1569 1594 1627 1650
1679 1682 1686 1718 1736 1740 1777 1778 176 234 240 263
396 438 496 504 526 688 739 741 758 1013 1152 1195
1211 1233 1249 1412 1430 1494 1509 1570 1636 143 298 1165
1180 1197 1293 1584 1649 1698 35 105 117 118 126 134
135 247 284 288 302 323 325 394 466 502 511 709
779 786 810 892 893 911 953 974 986 1005 1060 1063
1064 1109 1138 1213 1229 1232 1295 1320 1322 1389 1404 1419
1449 1461 1466 1491 1527 1536 1541 1559 1575 1663 1759];[5
21 22 97 108 133 158 165 247 251 321 325 348
411 420 421 424 466 497 526 620 621 718 755 886
890 910 933 935 960 982 991 1010 1071 1072 1083 1085
1137 1160 1169 1187 1250 1299 1320 1402 1421 1484 1514 1551
1569 1573 1616 1643 1695 1712 1737 1843 1865 1878 1892 1909
1931 1985 2002 2004 2018 2020 2022 2065 2070 2094 2097 2120
2141 2154 2187 2188 2242 2258 2263 2268 2293 2296 2325 2334

2350 2399 2416 2434 2453 2477 2519 2533 2545 2552 2580 2612
2614 2621 2698 2752 2805 2806 226 255 318 320 326 328
418 523 639 847 952 1023 1047 1188 1192 1312 1351 1422
1489 1505 1718 1866 1894 1932 1979 2167 2176 2184 2253 2266
2287 2339 2386 2420 2528 2638 2659 2771 2778 2782 190 331
355 953 995 996 1097 1198 1211 1369 1378 1399 1645 1647
1904 2061 2062 2132 2133 2260 2415 2551 2633 7 12 37
43 121 138 212 225 258 272 286 412 417 423 449
453 464 542 545 677 717 719 724 730 738 806 807
812 853 870 877 895 898 913 914 932 951 962 981
1091 1127 1173 1177 1206 1231 1249 1276 1287 1288 1336 1387
1408 1431 1446 1482 1495 1550 1561 1591 1622 1754 1766 1781
1848 1881 1883 1903 1907 1930 1960 1976 2000 2011 2021 2102
2103 2125 2174 2183 2218 2247 2264 2267 2280 2320 2338 2393
2446 2478 2487 2520 2565 2577 2600 2637 2651 2661 2695 2773
2780];[3 24 43 50 96 113 196 205 282 296 298
304 330 348 357 371 385 412 440 488 504 511 527
543 567 581 591 628 636 699 710 715 729 751 778
780 782 796 810 857 893 900 944 958 1021 1031 1092
1131 1147 1190 1227 1250 1272 1316 1317 1337 1399 1428 1439
1467 1476 1521 1542 1544 1546 1584 1671 1683 1691 1698 1706
1721 1724 1788 1803 1832 1880 1900 1918 1949 1998 2001 2011
2093 2103 2108 2109 2154 2172 2194 2207 2214 2298 2332 2378
2388 2510 2520 2525 2539 2542 2543 2545 2601 2657 2735 2738
49 262 362 627 656 777 859 867 869 1035 1089 1237
1326 1371 1384 1427 1468 1501 1604 1677 1702 1823 1827 1848
1875 1921 1980 2004 2019 2082 2100 2126 2276 2281 2295 2342
2596 2737 299 555 717 764 792 868 987 1838 1864 2275
2341 2421 2422 2546 2661 2678 25 26 29 107 111 114
134 141 176 185 231 237 242 274 343 427 433 493
521 616 648 703 741 752 767 824 826 833 837 891
901 950 1007 1032 1129 1163 1164 1188 1207 1223 1225 1269
1277 1331 1335 1346 1358 1444 1523 1525 1537 1582 1638 1644
1701 1723 1804 1837 1889 1976 1979 1982 2018 2063 2081 2099
2113 2125 2135 2184 2212 2230 2233 2237 2337 2358 2460 2461
2500 2519 2531 2581 2604 2606 2612 2618 2683];[21 36 100
101 238 257 259 278 281 300 315 320 367 414 416
456 547 575 604 633 635 671 711 733 741 743 817
819 879 883 927 956 971 1027 1066 1074 1143 1178 1262
1276 1346 1361 1375 1376 1383 1388 1403 1410 1425 1496 1511
1572 1612 1629 1662 1670 1722 47 117 154 221 291 358
382 504 650 672 768 873 902 1039 1076 1152 1167 1255
1275 1332 1345 1420 1558 1673 1732 1767 9 483 489 496
514 794 955 1095 1257 1291 1335 1555 1661 1724 12 48
49 82 85 99 103 155 158 231 251 279 280 285
325 398 406 428 434 465 490 491 499 587 637 678
694 697 827 829 857 880 890 901 969 1041 1065 1072
1081 1093 1094 1111 1144 1228 1238 1246 1260 1273 1296 1311
1343 1349 1354 1357 1370 1396 1480 1504 1556 1672 1715];[44
54 98 118 130 131 135 136 214 217 311 329 336
383 442 450 453 466 482 563 596 603 621 635 641
682 693 715 721 792 799 848 855 892 951 969 970
1048 1056 1063 1149 1235 1246 1251 1254 1278 1306 1348 1359
1384 1408 1412 1441 1503 1532 1533 1538 1578 1599 1600 1607
1676 1706 1731 1751 1786 1822 1828 1899 1910 1911 1912 1916
1922 1933 1940 1942 1981 2006 2027 2039 2059 2070 2083 2183
53 114 170 212 327 370 390 441 573 614 636 709
733 750 967 985 1175 1177 1370 1409 1589 1596 1641 1759

1819 1862 1867 1868 1873 1908 1939 2012 2158 2199 2278 101
201 222 225 303 400 465 592 630 634 660 926 1014
1028 1456 1788 2084 2236 8 18 43 57 75 102 186
259 315 379 399 401 404 412 413 427 438 470 523
542 574 590 593 594 613 665 720 741 749 752 802
884 932 952 960 968 979 1129 1205 1249 1261 1349 1362
1364 1470 1504 1565 1606 1618 1659 1689 1733 1742 1758 1787
1834 1879 1929 2034 2075 2077 2104 2123 2128 2154 2155 2156
2167 2193 2233 2234 2237 2252];[4 21 71 90 107 108
130 131 188 193 217 219 244 250 262 287 298 301
322 392 490 559 567 572 605 689 769 819 854 861
911 915 939 958 981 1070 1077 1088 1094 1104 1127 1154
1205 1233 1252 1314 1335 1378 1388 1395 1405 1406 1421 1422
1453 1456 1460 1477 1505 1528 1538 1557 1558 1592 1632 1693
31 56 97 115 195 213 242 316 437 470 478 557
585 868 912 984 1078 1131 1175 1201 1214 1234 1339 1434
1459 1546 1560 76 255 456 517 851 871 896 905 1221
1322 1353 1512 1537 1772 11 17 35 53 105 230 234
291 333 349 445 457 468 586 611 626 632 664 718
729 770 876 934 942 949 1009 1017 1042 1049 1074 1105
1163 1181 1203 1216 1219 1249 1274 1307 1343 1344 1354 1363
1381 1414 1437 1468 1553 1554 1591 1596 1604 1635 1677 1682
1709 1736 1738 1797];[9 10 65 155 224 275 287 337
349 350 371 381 427 428 430 453 516 532 602 652
656 683 745 746 822 826 848 910 965 979 983 986
987 996 1025 1026 1050 1051 1057 1082 1085 1093 1180 1244
1246 1314 1316 1364 1367 1391 1409 1411 1508 1559 1580 1635
1643 1762 1799 1820 1845 1861 1873 1882 1883 1930 1936 1966
2018 2023 2028 2056 2063 2139 2144 2145 2151 2161 2167 2170
2171 2183 2224 2265 2301 2319 2371 2375 2387 2397 2431 2432
2439 2456 2477 2521 2535 2537 14 117 243 250 296 368
403 422 547 563 605 749 812 847 1081 1112 1124 1194
1207 1255 1292 1299 1412 1621 1713 1796 1851 1893 1935 1960
2040 2074 2168 2289 2334 2353 248 269 305 378 834 1000
1252 1289 1490 1555 1622 1687 1723 1881 1884 1894 1977 1983
1985 2352 2466 21 57 63 89 105 113 130 140 159
168 227 291 319 379 382 451 452 492 497 498 539
647 654 660 684 700 714 733 742 751 773 820 850
902 922 952 992 995 1002 1028 1035 1086 1103 1139 1186
1256 1269 1284 1330 1430 1441 1575 1706 1724 1733 1832 1852
1939 1942 1984 2002 2038 2081 2082 2099 2131 2199 2304 2359
2419 2525 2528 2529 2542];[19 27 44 72 74 81 115
140 152 176 182 207 211 218 231 256 329 333 339
364 436 437 510 530 573 597 634 643 670 674 682
720 795 812 826 841 850 869 874 878 913 931 1016
1042 1064 1107 1129 1139 1140 1168 1193 1194 1198 1219 1240
1282 1426 1429 1524 1551 1556 1561 1587 1609 1634 1680 1713
1714 1737 1739 1761 1766 1807 11 132 274 426 503 520
564 599 678 789 793 897 995 1017 1094 1112 1115 1220
1222 1309 1383 1516 1549 1562 1681 1711 1744 487 519 647
652 1004 1110 1123 1174 1252 1387 1497 1618 1795 1796 1805
1 2 3 122 130 181 243 270 275 348 363 372 429
430 435 447 482 593 598 637 690 725 732 786 796
800 811 828 877 882 903 928 970 1030 1062 1069 1113
1175 1202 1206 1318 1344 1424 1454 1500 1507 1513 1517 1607
1735 1773 1809];[3704 3729 3798 3857 3858 3933 3947 3969 3985
3770 3808 3822 3832 3926 3941 3946 3827 3702 3825 3826 3959
3973];[36 46 49 56 58 85 94 129 148 159 203 215 283

298 337 340 346 400 404 420 484 517 562 31 173 303 349 359
513 514 528 268 11 33 73 84 93 95 158 252 273 284 285
297 334 351 378 399 408 438 458 470 506 540];[90 102 112
131 151 167 168 193 218 233 258 259 271 329 337
355 446 476 524 527 561 564 575 595 619 642 733
749 766 767 768 787 795 831 856 884 906 910 933
1000 1004 1029 1034 1188 1198 1204 1210 1305 1324 1370 1437
1439 1463 1464 1508 1516 1541 1608 1616 1646 1652 1660 1735
1747 1751 1933 1945 1980 1988 2026 2045 2059 2070 2092 2155
2161 2214 2238 2279 2316 2317 2342 2373 2377 2384 2437 2438
2509 2514 2537 2547 2550 2599 2619 2629 2631 2632 2641 2650
2680 2706 2803 2877 2887 2967 2968 2976 2986 3002 3020 3083
3099 3116 3125 3243 3265 3304 3331 3361 3388 3405 3406 3439
3440 3453 3528 3531 3551 3589 3601 3620 3651 3654 3659 3665
3683 3737 3742 3744 3776 3820 3841 3852 3890 3913 3924 3960
3972 3986 4019 4029 4033 4040 4059 4107 4142 4165 4177 57
295 328 420 470 740 760 796 930 1011 1027 1099 1137
1182 1302 1358 1415 1721 1725 1788 1869 1878 1931 2032 2130
2182 2312 2380 2400 2422 2429 2465 2510 2566 2569 2575 2705
2794 2829 2945 2952 3001 3087 3102 3108 3160 3177 3227 3230
3303 3305 3321 3341 3526 3602 3604 3611 3619 3663 3667 3825
3907 3908 3968 3974 4103 4141 255 563 641 756 1103 1252
1344 1380 1535 1606 1990 2017 2050 2171 2620 2994 3080 3093
3106 3387 3478 3616 4151 75 120 132 250 290 300 317
323 344 354 356 428 429 453 508 512 557 602 613
636 721 729 748 855 904 931 934 1028 1069 1102 1128
1134 1142 1199 1206 1215 1236 1264 1306 1343 1375 1389 1394
1423 1470 1483 1607 1703 1733 1743 1778 1798 1807 1808 1892
1932 1964 1987 2000 2021 2038 2170 2177 2187 2221 2235 2246
2308 2398 2539 2556 2588 2596 2692 2694 2701 2715 2738 2771
2779 2789 2865 2878 2893 2913 2924 2987 2996 3012 3017 3033
3057 3066 3084 3113 3136 3139 3175 3342 3349 3360 3433 3536
3593 3682 3690 3756 3795 3833 3919 3920 3961 3998 4034 4049
4074 4081 4083 4090 4097 4112 4138 4155];[12 14 26 109
110 150 169 193 195 243 262 264 266 270 287 314
430 454 465 497 513 550 554 563 571 624 627 695
727 776 800 807 813 823 839 908 919 946 977 994
998 1049 1075 1106 1126 1222 1249 1268 1286 1310 1319 1403
1408 1415 1532 1535 1542 1569 1570 1574 1579 1587 1592 1637
1681 1713 1746 1824 1853 1894 1944 1947 1987 1993 2005 2106
2120 2153 2181 2197 2203 2204 2285 2383 2394 2422 2435 2449
2476 2537 2605 2618 2621 2622 13 16 108 321 322 458
477 506 531 599 696 724 738 781 840 940 1048 1164
1309 1449 1470 1550 1678 1680 1762 1792 1805 1918 1997 1998
2054 2067 2220 2357 2410 2489 2521 2592 276 533 598 833
967 995 1061 1305 1411 1753 1755 1952 1961 2218 2255 2384
33 49 107 144 151 178 187 207 213 224 258 263
370 397 415 442 459 509 622 649 655 657 693 697
730 808 821 841 891 939 1025 1117 1144 1173 1229 1240
1308 1322 1401 1526 1541 1551 1556 1565 1588 1609 1626 1635
1721 1789 1791 1856 1873 1983 1990 1992 2004 2007 2016 2098
2227 2260 2264 2283 2299 2358 2406 2414 2447 2468 2505 2524
2526 2538 2579 2595 2596];[15 23 25 33 73 100 103
140 146 169 184 211 223 236 242 271 281 298 319
352 360 378 384 389 475 514 517 532 592 611 640
688 721 724 746 772 776 777 819 820 844 927 1018
1039 1040 1102 1154 1212 1222 1238 1268 1270 1283 1294 1301
1302 1310 1328 1338 1348 1363 1445 1446 1452 1469 1535 1593

1604 1645 1669 1681 1725 1755 1883 1915 1916 1939 1977 21
99 143 199 283 308 446 476 490 535 708 789 806
833 1050 1508 1613 1646 1719 1732 1787 1850 1884 31 113
309 342 368 431 445 549 698 848 1005 1131 1254 1282
1407 48 66 85 110 135 177 186 227 257 292 314
317 324 330 348 373 395 439 529 548 609 641 712
742 832 835 836 882 901 926 943 1044 1064 1071 1108
1205 1214 1221 1317 1371 1381 1413 1471 1488 1528 1559 1572
1589 1660 1665 1686 1701 1753 1783 1796 1825 1841 1842 1878
1898 1940 2008];[3 5 56 63 71 73 96 116 169
230 287 305 306 307 331 337 344 347 351 390 438
504 534 566 593 610 642 659 669 711 712 747 757
758 808 835 845 874 876 915 940 948 1064 1076 1097
1105 1128 1135 1146 1150 1161 1234 1249 1272 1274 1292 1294
1314 1315 1331 100 290 342 375 429 569 586 594 631
816 865 1110 1339 266 759 807 826 931 1221 1291 9
33 38 86 89 135 155 191 226 246 252 270 294
483 484 496 555 559 562 613 662 671 725 728 764
772 800 836 1010 1061 1077 1156 1195 1215 1219 1229 1324
1332];[4 60 67 82 95 129 155 163 196 198 211
316 345 372 379 387 392 422 435 450 474 517 519
634 637 699 717 772 889 941 963 1003 1004 1021 1061
1125 1140 1160 1161 1193 1228 1261 1262 1265 1271 1283 1314
1348 1417 1458 1508 1565 1570 1621 1758 1761 1795 1811 1819
1828 1846 1861 1900 1985 2003 2031 2038 2063 2079 2140 2143
2164 2175 2187 2252 2273 2277 2308 2318 2366 2398 2413 2414
128 293 433 447 705 707 718 840 908 1029 1055 1117
1170 1181 1258 1324 1392 1447 1631 1632 1634 1759 1939 1940
1992 2035 2067 2078 2110 2144 2261 2294 2340 2359 65 121
236 425 434 446 513 962 967 1665 1746 1859 1946 2080
2094 2372 36 148 152 180 248 323 368 371 377 401
423 448 451 458 477 502 504 586 619 692 704 726
730 740 767 770 783 818 841 843 871 890 950 982
998 1017 1050 1066 1091 1144 1174 1178 1184 1224 1238 1241
1244 1257 1289 1298 1332 1347 1349 1374 1429 1454 1460 1481
1500 1531 1543 1562 1564 1567 1603 1675 1678 1711 1764 1852
1922 1987 1988 2081 2088 2092 2093 2101 2103 2163 2166 2179
2198 2228 2236 2239 2259 2269 2291 2334 2393];[7 80 83
132 177 226 240 307 394 424 442 491 553 587 605
606 617 638 689 695 709 717 719 752 794 866 878
879 903 979 1115 1119 1135 1181 1199 1206 1229 1242 1253
1287 1294 1319 1320 1352 1383 1390 1430 1436 1454 1476 1491
1527 1562 1660 1688 1689 1695 1718 1739 1778 1794 1795 1801
1807 1853 1908 1914 1920 1929 1966 1990 1996 2017 2025 2112
2131 2157 2178 2201 2204 2225 2297 2329 2361 2398 18 19
21 98 106 169 172 194 281 486 519 600 607 726
764 778 902 935 1159 1236 1245 1278 1379 1429 1521 1645
1727 1921 1932 1935 1983 2023 2094 2193 2268 2281 231 392
411 417 429 651 736 746 936 989 1027 1186 1388 1525
1677 1792 1863 1977 2254 2353 20 27 36 118 126 171
197 211 336 380 381 388 399 423 454 525 530 586
611 621 630 672 760 761 791 874 889 916 952 964
1005 1162 1183 1213 1266 1279 1299 1392 1422 1468 1483 1507
1510 1558 1692 1720 1762 1763 1777 1788 1841 1852 1868 1882
1896 1903 1963 1971 1974 2021 2038 2064 2073 2122 2170 2172
2183 2191 2239 2298 2309 2394];[1 5 24 37 87 88
96 101 151 153 167 190 235 288 348 383 400 437
438 453 454 457 468 474 508 525 548 642 653 686

```
688   732   765   771   792   799   800   837   840   848   896   906
928   946  1043  1078  1081  1082  1121  1135  1167  1173  1180  1187
1206  1250  1271  1279  1315  1317  1331  1368  1416  1433  1434  1470
1563  1587  1620  1629  1630  1683  1684  1689  1710  1740  1783  1795
1845  1880  1890  1915  1916  1923  1932  1978  1996  2016  2035  2075
2076  2078  2166  2169  2226  2238  2270  2360  2362  2394  2420  2425
2428  2461    78   110   347   511   584   665   801   838   945  1211
1294  1338  1401  1443  1503  1618  1733  1772  1813  1889  1946  2008
2149  2165  2275  2289  2359  2440    19   255   731  1122  1174  1188
1394  1452  1632  1983  2034  2182    34    59    83   237   254   263
324   340   362   462   493   572   585   601   699   708   727   739
764   787   830   869   877   880   907   966  1045  1091  1096  1099
1107  1136  1140  1153  1191  1199  1264  1269  1272  1282  1306  1334
1385  1417  1432  1445  1463  1469  1523  1524  1673  1676  1711  1753
1807  1846  1876  1896  1953  1965  1991  1994  2009  2036  2062  2097
2101  2106  2133  2213  2219  2224  2281  2293  2344  2361  2391  2419
2426  2433];[57   102   103   171   183   186   274   287   295   308
368   381   427   442   479   486   508   576   597   605   613   644
662   666   667   747   756   775   792   805   816   849   886   901
946   961   992  1020  1049  1054  1070  1096  1116  1123  1175  1189
1219  1223  1352  1442  1459  1515  1546  1554  1567  1568  1573  1583
1621  1639  1686  1707  1721  1841  1901  1906    98   112   185   334
404   487   561   564   567   614   728   738   783   812   884   918
1115  1150  1181  1313  1433  1629  1734  1736  1856  1862  1905  1920
25    58   302   516  1117  1125  1262  1376  1468  1559  1595  1619
1645  1718  1737  1757  1837  1852  1893    43    89   100   229   240
243   259   288   305   335   406   435   454   459   478   490   514
556   570   598   626   639   656   683   707   735   737   743   755
779   784   826   875   877   910   944   949   956   958   982  1009
1025  1055  1073  1145  1169  1204  1275  1303  1304  1318  1357  1434
1452  1469  1471  1475  1482  1514  1579  1617  1728  1800  1808  1843
1847  1855  1921  1936  1941];[265   528   530   535   542   545   547
551   554   556   557   567   571   572   583   584   594   596   599
600   608   630   635   643   653   658   665   668   671   672   679
697   710   711   713   714   716   722   729   730   731   732   735
744   745   746   747   750   759   762   769   774   785   786   794
795   796   801   802   803   808   819   823   828   835   836   841
857   858   862   864   865   866   870   875   876   877   879   885
886   888   892   893   896   897   898   903   904   913   914   940
946   947   948   953   954   955   956   959   969   970   971   973
975   981   987   988   993   994   999  1003  1007  1029  1033  1034
1035  1331  1379  1488  2040  2045   126   403   526   532   575   576
590   659   660   749   753   809   905   932   962  1008  1023  1032
2031   455   534   558   592   595   624   717   799   827   837   880
899   911   931  1013  1121   178   531   548   586   598   606   618
622   634   637   650   651   669   670   675   678   681   690   699
700   723   740   743   752   757   772   811   821   824   830   831
839   859   867   871   874   882   909   910   919   933   958   966
982   996  1005  1051  1234  1496  1581  1592  2007];[14    18    25
57    58   110   138   173   184   188   221   257   260   310   321
384   390   456   494   501   533   562   564   604   606   654   692
710   717   743   769   788   797   813   819   887   918   925   976
987   995  1065  1068  1098  1131  1176  1184  1192  1247  1248    20
70   226   385   643   645   691   770   935   993  1175  1180    23
484   505   605   607   955   999  1033  1240    47    48   165   178
187   199   282   292   317   340   397   504   523   555   617   680
686   689   731   765   839   841   842   848   859   897   997  1040
1085  1087  1102  1114  1182];[8    27    28    60    96   146   152
```

```
184   209   247   248   304   366   368   396   407   410   508   559
571   597   626   642   644   651   680   686   703   704   745   755
805   862   893   899   900  1018  1023  1065  1100  1101  1133  1151
1184  1185  1193  1206  1231  1247  1293  1303  1319  1320  1343  1351
1408  1417  1513  1550  1616  1634  1660  1692  1702  1706  1715  1717
1758  1787  1828  1859  1893  1942  1981  2032  2133  2186  2215  2224
2290  2296  2299  2300  2309    23    50   261   263   420   447   498
622   695   702   748   756   774   856   945   946   964   987  1197
1204  1212  1261  1284  1373  1376  1462  1566  1619  1666  1673  1694
1827  1841  1843  1882  1938  2028  2047  2214  2242    25   125   345
347   404   412   648   709   906   988  1040  1088  1213  1384  1514
1713  1761  1762  1836  1958  1994    38    47    48    65   111   126
129   136   164   178   284   285   350   362   373   458   476   479
527   530   574   614   677   685   747   776   780   816   903   937
989  1017  1072  1085  1091  1092  1137  1172  1200  1248  1322  1339
1361  1407  1460  1466  1478  1539  1549  1609  1705  1716  1799  1810
1856  1870  2000  2016  2089  2158  2175  2177  2184  2189  2209  2268
2305];[53    77   105   107   141   241   257   258   300   327   398
495   561   572   614   625   626   633   663   670   674   675   686
692   701   707   720   790   799   864   888   950   956   999  1025
1035  1063  1088  1105  1134  1145  1147  1218  1288  1335  1336  1346
1348  1428  1487  1488  1543  1568  1612  1654  1660  1672  1674  1682
1686  1697  1723  1726  1731  1737  1747  1819  1834  1861  1873    21
189   233   330   469   612   676   771   784   840   906  1019  1046
1066  1167  1225  1317  1326  1380  1591  1634  1642  1659  1803  1837
1839  1888   243   285   376   438   446   465   624   808   949  1092
1176  1177  1179  1228  1250  1316  1370  1389  1446  1484  1575  1679
1724  1852    16    31    34    75   214   238   249   269   363   374
408   418   419   427   435   451   544   647   677   689   691   738
752   759   848   868   895   897   898   905   961  1045  1074  1077
1107  1168  1201  1213  1292  1328  1332  1349  1353  1361  1374  1431
1440  1450  1459  1537  1570  1588  1637  1675  1692  1818  1844];};

% genus='frankia';
% SWITCH = 'for_Frankia_Sco';
% SUFFIX = '_Dh_Sco';
% d = '';
% % % didn't do
'igrs_frank_RB7';'igrs_frank_Hr75';igrs_frank_alp';'igrs_frank_KB5'; as
they are either with split IGRs or uncertain contexts, see Frankia above.
%
gs={'igrs_frank_casuarinae230';'igrs_frank_nepalensis230';'igrs_frank_tor
reyi230';'igrs_frank_umida230';'igrs_frank_canadensis230';
%
'igrs_frank_QA3230';'igrs_frank_R43230';'igrs_frank_alni230';'igrs_frank_
ArI3230';'igrs_frank_CcI6230';
%
'igrs_frank_AvcI1230';'igrs_frank_BMG5230';'igrs_frank_Allo2230';'igrs_fr
ank_CeD230';'igrs_frank_nodulisporulansAgUm230';
%       'igrs_frank_ACN1ag230';'igrs_frank_B2230';
% % split IGRs
%
'igrs_frank_gtarii230';'igrs_frank_tisae230';'igrs_frank_EI5c230';
% % not long enough IGRs
%       'igrs_frank_Cj5230';
%     };
% SCORES=[12 13 14];
```

```
% ns={[1355   106    21  1177  1722  2232  2210  2358  1520   249   354
923  1329  2302  2530   131   262  1218  1311  1551  1600  1851  2313
2403  2528  2571    80   167   420   970  2127  2568  2575  2594    88
172   327   328   336   484   625   894  1237  1439  1493  1800  1951
1986  2158  2239   342   350   555   613   647   720   842  1103  1178
1251  1599  1609  1683  1762  1786  1885  1903  2060  2278  2511    16
35    77   216   410   667   698   713   716   969   985  1016  1039
1143  1219  1269  1274  1299  1450  1585  1730  1957  2027  2037  2567
2432  1340   792  1222  2090  1874   370  1038  1422  1307  1586  2599
1215  2413  2366  2365  2014  2311   149    22   329  1953  2225   768
353  1507    83  1166   608  1726  1555  1358  1565   477    32  2082
1645   573   665  1200  1619  1887  2089  1059  1768  1738  1213  2289
261  2204  1636  1041  2007  2527   791   712   765  1216  2129   552
82  1747  2531  2249  1701  1426  2140   287  1309  1873   996  1724
2008  1673   549   718    78  2469   505  1657  2156  2230  2557    40
645  1021  2066];[1128  5425  1277  2089   593  1937  2562  4583  4099
4325  3180  4320  4991  5207   866  1203  1931  2104  2545  3333  3525
3836  3838  5758   140   615   753  1372  1482  1633  2079  2337  4092
4217  4438  5721   186   324   439   949  1980  3086  3468  3893  4535
4729  5255  5336  5392  5701    63   514   913  1051  1617  1705  1838
2097  2142  2284  2383  2905  3128  3368  3601  3849  3956  4300  4332
4662  4681  4995  5395  5483  5638    82   174   237   244   298   616
867  1135  1136  1365  1421  1507  1519  1819  1863  1881  1983  2000
2086  2246  2250  2439  2474  2589  2769  2794  2854  3172  3206  3291
3425  3798  4038  4235  4484  4649  4718  4846  4919  4940  5333    31
187   200   231   270   323   524   607   672   789   802   822   945
1186  1501  1866  2005  2387  2409  2535  2546  2590  2773  3018  3063
3118  3181  3182  3302  3672  3965  4017  4184  4297  4606  4612  4631
4682  4741  4914  4939  5099  5107  5157  5278  5327  5390  5561  5571
5690  5703  5727  5751  2735  4684  2927  2330  1453  1726  4656  2998
608  4221  3437  2129  2806  5442  4071  2840   859  5112  2356  2367
1065    36  1620  4574  4061  1674  3844   296  3889  3636  4169  2196
3370  1249  1113  1393   740  4434   808  3254  3908  1991  1495  2282
1871  3157  1052  1622  4533  2863  4928   199  4862     1  4546  3897
4538  3161  4102  1064  5000  1409  1858  4537  5512  4788   245  1814
1205   764  2877   523  4258  1788   640  1685  5274   173  3659   569
2774  1314  5524  3002  4009  4598   894  1926  3174  3975  2511  5748
3244  1321    52  3085  4151  3411  4901   863  5217  3898  4069   540
1185  1664  3158  1948  5488  5415   436  3702  2280  1879   612  1652
5160  1565  5477   776   641  4999   338  3450  5052   417  4908   486
2674   675  5647  1908  4644   941  1049  4712    68  5464  5564  1736
3635  3405   513  4437  4494  1139  3381  1024  4542  2841  4760    42
846   648   430  5677  5498  4691  5678  1360  1537  4121  5349  4811
2858   796    34   872   951  4100   233   372  2949   663  1262  1670
3166  3192  4308  5398    89   862  2984  3243  4444  4792   535  2163
2344  2369  2738  3142  3224  3766  3878  4013  4175  4724  5397];[363
2793  3101   285   729   904  3082  1350   292   756  1426  1492  2249
2836  3199  3311  3324  3619    40   905  1289  1349  1476  2055  2138
2829  3068  3228  3491   675   876   888  1056  1260  1474  1759  1977
2613  2958  3172  3342  3481  3506  3532   254   355   559   590   647
682   937  1029  1084  1500  1720  2295  2373  2572  2576  2620  2708
2960  3191  3282  3312  3513  3597  3626    25   118   260   298   364
366   408   416   489   567   873   889   922   986  1019  1020  1037
1129  1262  1290  1299  1300  1370  1609  1645  1819  1864  2409  2574
2676  2696  2889  2933  3093  3252  3384  3388  3414  3550  3560  3630
261  1244  3271  2056  1148  3360  2780  2697  1640  1046   690  1627
792  2768  1491  2884  3031  3072  2820  3235   789   323   471  2921
507  1534  1693  2934  3577  1077  1805  1961  3216  3408  1047  1164
```

1182 1689 3547 676 160 1024 1318 2202 2756 344 2484 3502
3248 3472 1109 701 1193 193 1824 1287 1127 3210 450 2536
2910 1842 3230 1670 790 1599 63 3386 2212 2509 3310 2436
3215 3393 826 429 494 663 2435 2124 1354 157 648 1963
2567 325 2680 1460 2167 3457 770 2755 3453 1571 2288 1776
2346 2610 2031 970 1198 3494 1823 2141 915 1163 1482 1519
3035 375 705 89 2363 1366 3555 352 3119 902 943 1106
3116 238 859 1213 1678 1901 2735 3094 730 1307 2209 2929
3176 3335 38 214 404 785 989 2042 2193 2259];[1338 1849
1615 823 855 2086 290 1048 1699 2738 1172 1288 1371 1710
2755 861 1439 1684 1704 1731 1766 1888 2061 2387 2460 2919
570 673 738 1310 1383 1847 1949 2059 2471 158 163 694
973 989 1117 1390 1626 1798 2400 2509 2613 2806 205 216
689 731 910 942 1345 1667 1700 2097 2353 2685 2705 2756
2808 31 218 254 278 421 487 520 647 685 706 763
810 860 869 884 916 917 974 1041 1094 1157 1405 1574
1610 1625 1691 1714 1765 2096 2126 2177 2389 2462 2650 2697
160 2298 1265 1285 1657 436 962 2631 1611 2163 2041 541
2666 1581 17 21 1182 1279 2893 252 672 2235 736 2859
669 1444 1958 2233 2098 2570 1107 783 2583 2496 2894 2569
2017 968 606 259 1272 1894 2702 1255 784 1128 2240 1824
1973 2095 2304 1961 1566 1701 952 529 253 1635 2442 2229
2531 2292 1732 441 1779 2686 2441 1557 569 2660 2178 2526
2717 443 1137 2746 658 1924 1713 1055 832 350 358 1181
800 1458 317 568 553 1479 932 1011 1268 1825 465 1080
175 597 1492 2345];[1887 1302 3166 369 1336 2001 1627 2088
2635 2848 677 775 1065 1771 1940 2124 2949 819 880 1052
1073 1214 1837 1871 2829 3132 51 171 321 325 420 1493
1835 1967 2476 2804 3029 3067 3169 594 776 800 808 1021
1172 1569 2122 2243 2417 2670 2821 2921 3068 3125 3186 3198
65 180 229 330 384 457 541 543 544 1043 1102 1198
1341 1539 1961 1990 2255 2336 2442 2695 2827 3055 3230 3254
8 269 272 290 367 427 442 522 574 675 791 1041 1053
1257 1422 1473 1706 1732 1781 2008 2473 2516 2573 2681 2826
2997 2998 3041 3184 3289 3339 281 844 839 904 2544 1012
1644 201 1678 727 1138 570 2822 2468 1515 1071 2350 1072
2186 2343 783 3162 1939 419 299 2944 609 347 68 1301
1224 583 558 587 799 109 573 2340 2619 1665 341 186
417 1827 676 814 639 440 2535 1289 1009 2035 1048 3283
712 1340 581 366 1646 2019 1117 458 2789 1104 507 2536
1912 840 602 472 519 81 3113 943 3260 2361 2866 3073
2265 258 2364 1529 2432 489 2377 1740 503 1380 2094 2622
399 636 826 1868 3275 538 616 2012 2440 3060 75 342
1010 1364 1595 1757 1956 3263 3265];[2961 453 1254 1632 2907
192 3705 138 961 1208 2297 2939 3014 3606 3620 282 439
510 1117 1221 1501 1754 1878 2613 2615 2765 2853 331 507
585 1015 1867 2251 3583 139 292 1011 1467 1482 1693 2887
3006 3033 3240 3510 37 178 665 743 770 963 1099 1308
1338 1343 1836 2027 2079 2190 2421 2689 2763 3122 3333 3392
134 297 374 394 721 734 964 1299 1439 1797 1799 1831
1929 1940 1956 1965 1992 2006 2093 2252 2322 2337 2580 2585
2738 2804 2891 2956 2976 2999 3088 3093 3113 3302 3351 3449
3540 1385 2952 1728 3376 3121 2787 2435 1055 3127 1259 1053
1853 2721 2602 98 1945 1124 2126 191 1430 3529 1115 2980
135 3450 2791 3671 1541 962 267 3428 3462 2733 651 2651
3348 2472 2764 1250 3239 2982 2362 281 256 2193 27 480
3539 1734 275 63 1135 2174 2886 908 1476 1184 3448 186
1021 1213 52 414 2722 2497 456 1900 847 2036 3631 2792

3084 1220 3346 180 1972 3005 3452 1464 3041 346 755 1453
1994 2258 3243 2908 1216 1946 3718 122 1242 921 3387 2444
2111 508 3374 1466 3447 18 2294 1241 1886 2328 3714 365
1868 2616 2898 1079 2916 2821 14 812 2660 132 1163 1431
3728 347 2664 2950 3076 313 525 723 1012 1781 1851 3025
3268 65 234 268 638 1211 1751 2112 2387 2533];[4138 1628
2802 3320 1069 2390 105 423 1154 1756 1851 1904 4455 192
370 714 1199 1450 2285 2923 3079 3126 3314 106 152 609
663 1108 1152 1639 2096 2808 3709 3782 3848 4013 4102 4107
4129 252 283 300 1014 1150 1376 1380 1506 1769 1887 1985
2409 2518 2845 3920 4183 4212 147 249 796 826 970 1084
1369 1393 1438 1761 1873 1968 2084 2378 2420 2470 2538 2683
2796 3105 3220 3359 3364 3413 3481 3616 3713 3908 4032 4171
4225 4232 4234 4244 4260 71 151 569 614 777 891 1006
1127 1606 1668 1908 1973 2124 2196 2297 2310 2593 2824 2831
2960 2997 3017 3043 3044 3204 3213 3336 3353 3519 3539 3898
3914 4007 4008 4040 4209 4331 4334 4466 28 159 258 575
719 1198 1247 1508 1517 1608 1737 1746 1850 1990 2038 2156
2261 2504 2536 2550 2590 2684 2745 2772 2920 2949 3122 3210
3247 3391 3669 3743 3750 3775 3810 3968 4044 4139 4211 4237
4347 4350 4408 4415 4485 4515 4518 4587 1670 3249 2512 655
3147 1762 2432 548 629 2377 102 3767 2347 3863 4457 4527
2441 1269 1555 1701 1370 1099 1915 1313 3528 1924 274 2544
3033 2780 3313 3416 788 1321 3346 3919 1918 1994 2287 2407
3000 2869 570 3341 3408 2872 4078 3923 3557 2886 2520 4328
40 876 3533 3254 1031 791 968 1320 3464 1520 4177 982
3420 491 2876 3728 1799 3597 3479 4128 3522 2427 96 4108
3296 1992 4385 513 1589 3466 1852 4031 4146 2844 425 1617
244 1655 1661 3868 740 2723 4227 664 2570 4181 4188 743
543 967 433 1228 3732 3994 661 2790 3945 516 1592 2589
1950 3890 204 521 3165 532 3723 3345 1501 2609 2428 2721
2830 2919 4300 361 1687 2507 2991 4548 239 934 1177 1252
1626 1718 2070 2902 4510];[1184 445 2583 1140 1984 2569 731
1191 1236 1294 1499 2126 3326 138 515 579 820 1226 2079
2282 2609 2633 2717 3021 3063 1077 1372 1405 1481 1572 1811
2359 2380 2955 105 215 850 1128 1395 2041 2195 2289 2290
2387 2613 2761 137 266 398 438 441 477 478 903 1078
1118 1152 1298 1307 1374 1602 1697 1779 1800 1925 2043 2056
2263 2273 2423 2520 2652 2730 3008 19 216 409 787 834
925 952 1014 1020 1065 1448 1449 1535 1758 1926 1992 2142
2145 2414 2468 2607 2737 3034 3069 3080 3181 3299 2412 1473
581 1408 2375 1845 1262 3218 2003 401 2597 2220 1884 1740
487 567 1329 2129 1684 1064 436 2398 1115 45 2207 3111
3007 982 3337 521 1255 3233 2253 582 442 3033 681 2255
1719 2748 2670 1538 2596 924 856 2229 428 3128 93 2013
298 702 384 2040 2406 2953 525 365 420 362 550 1622
256 1190 54 2545 2374 516 3216 2230 440 2009 684 2312
2323 73 1456 766 2692 892 2589 1752 617 2778 1450 2590
2350 2326 661 1921 469 2146 1728 987 1056 3314 1203 2099
2961 1426 3060 397 648 978 1681 1187 2727];[2970 2744 190
1137 469 542 638 1421 1710 3283 2906 3423 151 1478 2038
3062 481 543 1146 1247 1374 2003 2944 3110 3284 3451 15
79 367 502 516 838 1326 1328 1567 1934 2069 2513 2752
3108 3476 3497 49 77 144 573 729 736 897 1180 1467
1471 1500 1504 1589 1927 2128 2133 2628 2701 2796 2901 3005
3302 80 123 277 358 418 499 519 627 677 878 903
937 979 1050 1142 1297 1533 1545 1579 1580 1637 1752 1979
2085 2167 2510 2639 2640 2695 2741 2743 2794 2805 2848 3006

```
3227  3288  3422  3518  3534   890   808   114  2096  2943  2884  3289
1851   708  2033   305   721   695  1031   219   975  3523   437   477
2563   165  1996  1034   923  1645  2466  2572  2021  1617  1344  1548
2539  2421  1584  1149  1774  2900  1069  3462  2600  1983  3136  2536
3274  1223   233  1127  2361  2247  2810  2512  2027  1446  3233  2826
2001  3340  1033   667   320  1786   876  1417  2022  1632  3425   459
746  1696  2605  2990  1325  2787  2079  3363   345  3094   441  1737
2294  3457  2298   819  3173  1937  1272   314  2366  3454  2809   553
1198  1849  3116  2684  2451  2732  1129  3156  3002  1751   480   234
491  1540  2755   392   395   579   909  1258  2276  2381  3348   933
2065  2604  2879   540   976  1038  1761  2334  2705];[1939   359   633
666  1126  1587  1112  2582    41    57   658   718   760   894  1337
2473  2576  2586   475   905  1235  1319  1914  2001  2090   419   441
826  1276  1349  1880    73    79    80   747   820  1057  1739  1875
2188  2238  2419  2448  2494  2569    17    61    69   251   292   440
551   942  1073  1208  1609  1650  1726  1814  2152  2209  2259  2324
2513   176   201   354   373   621   750   864   976  1160  1364  1406
1522  1674  1695  1962  2129  2191  2249  2355  2587  2066  1666  1213
120   336  1999  1023  1694  1315  1961  1252   483  1838   893  2413
642  1594    58   988  1137   629   873  2321  2016   360   813   370
1031  1744  1283   788  1082  1024  2211  1179   611  1816   881  2232
1136  2089   576  2334  1813  2524  2497  2467  2468  1743   333  1896
1125   945  1202   418  1572  1442  1985   147  2519  1317   585   631
1045   948   663   290   699  1363  1538   597  1101  1589   378  1469
1652  1677  1697  2158  2567];[4275  1361  2801  1555   231  1002  4516
583   928  1933  3782  4203  4466  4475   263   790  1413  1953  2392
3019  3272  3367   211   430  1218  1432  1989  2698  2902  4563  4879
15   216   218   697   731  1125  2363  2503  2863  2940  3134  3238
3342  4182  4389  4574  4581  4584   179   781   808   938  1183  1310
1424  1723  1724  1738  1789  1954  1957  2064  2134  2344  2583  2659
2731  2778  2870  3096  3197  3514  3917  3976  4043  4187  4246  4328
4503  4783    62   740  1131  1232  1431  1573  1736  1813  1842  2066
2121  2171  2353  2456  2570  2571  2941  3057  3416  3556  3560  3578
3596  3716  3768  3789  4035  4090  4181  4208  4231  4330  4339  4355
2929  2612  1074   645  3059  3588  4207  1373  1631  2996   667    70
261   188  4034   155  4348  3985  2011  3732  1160  4835  3537  4462
3336  1433   219   283  2847  4058  2100   935  4103  2450  4499   153
4802   236  2519   458   436  2463  4178  4069  3668   521  4709  1866
726  4678   673   496   262  3175  1847   264  1768  3787   365  2042
1934   886   647  2627  3646  1289  2719  2748  2508  2521  4483  3455
1694  3271   725   122  2476  2196  3311   863  3031  3450  2961   684
1524  2553  3796  4333    96   396  4229  3036  2138  2554   128   715
2321   324   235  1184  4577  1945  2412  4025  4568  3471   636  4662
1016  1851   641  4487  2114  1220  2839  2577  1085  1105  2023  2383
2572  1303   700   492  4562  1460  2611  2765  3103   106  1501  2084
3900  1173  2122  2334  2606  3325   105  2514  3513  3679  3689  4329
601   973   977  1189  1291  3182  3638  4368  4539   435   586   646
2013  2123  2199  2257  2402  2459  2888  3680  4084  4582  4837   728
1026  1043  1341  2580  3899  4079];[1303  2203   889   805  1540  1834
38    54   184   366   656  1632  1866  2340  2440    16   164   275
937  1196  1290  2365  2565   509   910  1117  1153  1357  1431  2075
2399     8    97   128   280   741  1049  1118  1161  1388  1493  1547
1699  2121  2404     7    22    34    74   288   588   739  1061  1402
1518  1626  1858  1890  2009  2202  2256   168   358   613   761   784
795   859   955  1237  1358  1725  1772  1820  1839  1850  1915  1949
2067  2110  2176  2187  2287  2503  1263   261   270  1593  1777  2267
996  1661  1907   933  2175  1533   915  1282   423  2480    93  1833
2007  2566    37   801  1887  2206  1512  1168  2076   386  1823   529
```

```
1756  1869  2218  1805  2005  1959   589  1344  2030   950   113   307
1908  1274  1348  1444   787   242  2434  1724   536  2217     6   150
149  1660   339  1096  1728  1433  1594  2068  2513   890   776  1557
1137   644   575  2326  1662   545   803   196  1583  2294   328   415
742  1305  1538  1628  1817  2308  2384  2532];[477   522  2667    54
552   654   777   153   816  1346  2439   908   924  1024  1154  1334
1483  1646  1859  2309  2436   266   463   834   945  1172  1237  1944
2134  2287  2640    85   106   173   638  1404  1531  1533  1789  2032
2334  2335  2403  2430  2596   357   470   597   617   899   940  1048
1202  1247  1691  2107  2341  2591   172   287   458   478   605   618
634   920   960  1047  1186  1245  1356  1450  1560  1752  1834  2017
2039  2070  2229  2539  2580  2684  1143  2534   857  1798  1235  1592
1074  1934  2653   547  1748  1159  1176  1659  2435   312   497  1534
82   258  1238  2691  1417   936  2614  1015   351   543   784   602
1742   923   726   665   556   995  1353   900   714  2348  2434  2421
53  1118  2443   342   653   631  1073  1641   151   341  2674   438
568  1836  1003  1912   664   833   608  1623   405   406  1886  1938
1117   627  1795  1972  2555   560  1285  1290  1478  1913    40  1750
607  1593  1702  2011  1561  1854  2447];[2180  1715   925  1100  2277
179  1634  2299  2424   309   740   756   832  1201  1549  1765  1882
2056  2068  2254   253   722   761   993  1817  2093  2415  2445   803
876  1156  1297  1583  1689  1803  2413  2439   113   714   982  1398
1495  1505  2023  2131  2136  2195  2270  2340  2364  2496    54   243
327   404   449   530   569   584   713   736   831  1099  1147  1203
1248  1319  1397  1513  1521  1551  1671  1672  1673  1717  1750  1845
1993  2115  2193  2239  2348  2381  2469  1318  1779  1061   957  1605
1   669   197  2033   594  1445   930  1947   762   531  2203  1155  1032
526   752  2463   880  1660  2446   981   715  2284  1889   909   739
1786  1111   529   921  1079  2523  1387   120   874   199  1446   955
1633  1056  1348    73   536   422   192  2483  1846   811  1822   956
1716  1303  2177   965  1887   915  2524  1830   367  1680  1916  2472
245  1725  1848  2245  1427];[1228   993   341   808    42   578   599
938    36   179   498   754  1389  1653   540   617   948  1574  1787
89   320   562  1297  1487  1556  1660  1738  1848  1875   101   143
412   683   880  1444  1578  1591  1613  1678  1790    29    37    78
180   196   297   473   694   872   895   901   937  1039  1065  1424
1432  1490  1595  1729  1035   539   385    75  1596   191  1293   963
1703  1775   859  1037   366  1321   232   816  1632  1806  1320   864
1447   397   707   685   265   625  1609  1867   132  1201  1884  1781
1307  1400   165  1329  1319  1624  1883  1584  1672    90  1715  1086
1905  1232    79  1439   218  1169  1906   994   997  1837  1478  1614
1066  1430   825  1652  1791   581   582   755   913   990];[1312   322
2573  2574   106  1640  3550  3365  3432  1149  1482  1896  2032  2727
2847  3182  3699    85   596   678  1877  2111  2892  3741    71   105
429   643   814   925  1385  1758  2181  3364  3416  3503  3536  3703
330   468   540   815   927  1197  1345  1400  1697  2180  2247  2248
2383  2482  2625  2645  2672  2944  3041  3059  3213  3381  3395   374
496  1014  1304  1375  1498  1704  1744  1783  1849  1850  1871  1899
2081  2182  2537  2549  3080  3135  3305  3332  3347  3478  3626  3694
3713  3719  3748  3750   180   226   319   321   372   406   434   619
739   770   812  1152  1191  1212  1509  1621  1805  1821  2091  2101
2208  2376  2526  2632  2654  2711  2957  3003  3130  3148  3188  3306
3354  3380  3398  3437  3471  3518  3558  3637  3638  3668  3678  3311
1644  1012   495  2949  2575  1735  2090  2697   194  3088   671  3112
3237   967  2816  3099   310   392  3577  3695  2584   700   433  2384
1296  2878   179   636  2246  2651  1833  3066   698   870  3403  2586
1886  1799    32  2631   404  3633  3422  2806  1972   504  1396  3302
3284  3226  3097   623   899  1202  3282  2688   663     5  3442   641
```

1456 413 3720 600 3140 3057 383 2524 3310 2854 724 443
1816 2255 2223 170 1776 1880 384 1987 369 1477 3050 1018
35 3273 941 1760 169 1929 3581 1564 3068 2571 2805 112
258 2276 3673 333 1464 1299 3326 3367 3535 403 421 1182
1747 1957 2956 3271 3560 972 1029 1084 1543 2667 3230
3602];[1596 1420 568 936 2408 424 1549 1565 2001 519 1060
1129 1261 1277 1634 105 221 815 961 1295 1663 2570 2572
2583 192 499 592 745 1000 1406 1417 1706 2450 174 312
431 751 1013 1089 1301 1462 1857 1929 1947 2141 2223 2338
2464 98 120 270 298 313 337 402 502 569 835 837
886 1168 1169 1239 1281 1289 1302 1695 1790 1844 1882 2186
2262 2389 2457 132 208 682 715 747 803 896 906 1202
1405 1517 1678 1936 1996 2131 2305 2386 2407 2449 844 1740
1164 1342 1030 1902 1711 811 1677 1709 621 2312 1664 417
584 2584 1278 1538 355 1561 565 2081 396 133 2012 1807
1614 2568 2241 1139 767 1461 758 1454 629 1413 178 244
1776 2493 721 2326 73 453 330 129 1739 1184 2084 1550
255 441 2022 368 319 930 1214 1901 64 1563 583 2129
671 52 114 422 698 1592 83 454 833 867 978 1502
1884 1954 2308];[3195 2905 3365 1250 3178 3179 1183 2135 3121
3314 71 1699 2929 282 337 1201 1588 1700 1703 2365 2843
2865 3007 3353 308 366 742 1159 1298 1774 2063 2411 2517
2792 2799 3191 156 236 504 606 810 862 872 1172 1203
1257 1272 1510 2144 2234 2260 2374 2583 2606 2884 2910 3181
41 114 428 451 508 532 572 602 622 739 1032 1137
1154 1255 1322 1399 1432 1512 1531 1532 1540 1578 1590 1628
1629 2270 2648 2689 2886 2908 3017 3047 3058 3081 3177 3299
6 184 212 216 227 514 542 579 639 694 1104 1462 1761
1791 1804 1822 1911 1975 2130 2349 2364 2443 2592 2675 3116
3391 168 8 50 1352 581 2602 2857 199 1591 2319 2189
3117 1803 2412 2819 1855 1034 1418 930 3044 217 2902 96
3344 1224 1696 1953 1223 2572 1396 874 2993 1431 798 1862
1138 2072 1190 2431 2968 1167 2022 3056 1982 2903 2362 3399
3320 3282 2086 1279 1468 762 2112 338 1687 3261 3398 2486
2879 122 458 3386 256 3005 452 1607 3390 2498 3375 2718
1686 2191 1827 2452 1301 521 822 2507 2654 973 3240 3094
2330 3108 314 2185 852 840 2891 787 3243 421 3408 499
935 976 1205 2539 2632 163 641 983 1049 1905 1999 2071
2548 2734 2769 2881 3441];[1919 2009 107 1093 2020 143 141
770 2572 1107 1674 2170 2351 2668 2828 2876 217 608 668
1971 2381 2521 2982 3072 3151 34 113 256 303 545 659
678 2522 2922 3111 3316 3402 195 289 384 573 768 850
1037 1041 1122 1151 1646 1724 1728 1792 1941 2605 2770 2803
3036 3153 264 291 463 530 534 574 728 731 819 840
1026 1034 1067 1145 1213 1222 1423 1510 1673 1832 1873 2002
2034 2120 2126 2156 2311 2461 2703 3289 3306 3359 27 178
181 186 295 451 457 568 599 744 885 924 1142 1266
1273 1275 1328 1342 1357 1375 1608 1697 1726 2159 2200 2223
2353 2386 2455 2640 2843 3073 3085 3211 3249 3422 454 745
2824 2790 422 12 565 1838 3404 1823 1297 452 1700 501
3170 1430 896 837 1769 2666 255 1866 1828 508 2863 346
1632 2188 1916 374 1135 1351 2238 2180 93 282 2551 2472
871 3314 2388 1365 2352 2100 693 3386 2643 2242 730 856
773 1271 1861 2642 2433 2150 520 319 1330 3319 2838 2392
877 1475 2087 2932 1983 1663 696 3286 1894 2179 1306 78
1397 2032 2719 1900 3265 2357 522 483 385 2000 127 430
32 1688 2387 20 985 1287 1300 2491 2820 1047 1265 1321
1447 1715 450 1016 2637 2848 3141];[846 3213 1911 2927 3047

```
414  1129  1148  1250  2031  3268  1942  1979  2247    55   294   870
987  1130  2797  3316  1201  1371  1800  1802  2055  2362  2803  3077
3317  3320   259   285   355   396   592   643   693  1489  1538  1630
1958  1985  2124  2159  2528  2918  3183  3363    31   169   260   452
705   893   958   997  1018  1035  1275  1472  1612  1665  1727  1730
1853  2237  2269  2718  2731  2775  2851  2869  3001  3093  3200  2530
1061  2909   331  1463  3245   773   691   349  2155   424  1815  2738
2656  2642   116   807   547  2058  2512  1073  1880  1348  2164  2897
333  1708  3124  3062  1147  2543  3127    59  3225  1115  1874   925
1637  2802  3002  1639  1579   940  1733  1499  1477    67   894   437
2545  2183   568  3215   838  3138  2474  2099   769  1944   999  2248
806  2323   265   715  1314  2077  2428  3301  3110   442  2367   770
1218   602   539  2270  1713  1895  2644  1761   533  1296  1987  1647
488   417  1340  2730    32   421   505  2967  3072    82  1672  2033
2199   831  1460  2120  2301  3123  3265];[1591  1614  2600  2023  2286
1570  2130  2218  2590   652  1184  1566  2283  2346   414   933  1154
1262  1516  1565  1567  1745  2217  2629    37   393   625   761   816
998  1362  2102  2141  2536  2650   201   288   358   373   413   431
499   846   887   917  1021  1226  1536  1612  1748  2192  2528  2578
2646  2647   145   577   591   759   785   934  1001  1006  1027  1067
1401  1487  1531  1853  2086  2112  2211  2244  2264  2347  2549  2652
773  1651    95   183  2083   170  1656  1683  1087  1980  1942  2408
2108   784  2514   855   812  2129  1520  1694   274   217   844  1321
2117  2465  2458   906  1361   384   249  1403  2481  1583   562  2173
2061  2371  1630  2581   953  1592   365  1994  2427    62  1886  1695
1123  2313  2596  1599  1317  1131   181   766  2066  1420   424  1682
2166  2220  2222   318  2295  2349  2309   331  1738  2523  2649  1346
1382   187   272  1097  1104  1255  1351  1491  2213  1515  2639   896
1270  1360  2338  2367  2634]; };
% SCORES=[8 9 10];
% ns={[14    15    21    22    74    82   120   149   167   171   216
287   290   327   328   329   336   357   370   418   420   467   502
550   552   573   608   610   665   698   712   716   724   766   768
792   803   918   969   979   992   996  1016  1038  1050  1059  1127
1143  1166  1200  1213  1215  1219  1222  1309  1324  1329  1333  1340
1358  1422  1426  1464  1493  1507  1520  1536  1551  1555  1586  1600
1609  1619  1645  1657  1708  1722  1726  1747  1811  1859  1874  1887
1895  1903  1953  1978  1990  2003  2037  2042  2090  2091  2129  2130
2139  2140  2176  2210  2225  2232  2314  2358  2365  2403  2413  2430
2432  2528  2571  2594    32    83   106   214   315   353   409   477
549   667   707   723   791   842   923   985  1177  1218  1311  1353
1419  1450  1565  1724  1768  1833  1873  1885  2089  2204  2289  2302
2311  2366  2472  2531  2544    16    77   242  1034  1184  1810  2158
2296];[8    31    36    42    43   108   173   186   199   237   245
287   289   296   298   338   341   371   430   486   523   524   535
563   579   590   608   615   672   740   776   789   796   802   866
941  1049  1051  1065  1075  1102  1111  1113  1161  1185  1205  1253
1274  1278  1311  1360  1365  1372  1421  1453  1481  1495  1507  1517
1521  1565  1567  1571  1578  1601  1610  1611  1622  1652  1683  1705
1759  1788  1814  1819  1860  1862  1876  1920  1926  1931  1977  1987
2079  2089  2096  2142  2246  2250  2274  2280  2284  2330  2337  2356
2431  2440  2456  2498  2543  2546  2674  2735  2769  2806  2813  2840
2858  2863  2909  2911  2919  2927  2960  2966  2973  2998  3002  3018
3063  3078  3118  3157  3158  3161  3179  3181  3182  3213  3215  3248
3254  3287  3291  3296  3302  3321  3335  3425  3468  3499  3549  3555
3565  3600  3626  3636  3659  3780  3838  3844  3878  3897  4013  4054
4061  4069  4071  4102  4161  4169  4185  4186  4217  4221  4297  4315
4320  4418  4423  4434  4455  4477  4494  4533  4537  4542  4546  4550
```

4558 4598 4604 4654 4656 4681 4684 4691 4712 4788 4799 4805
4811 4862 4901 4991 4999 5052 5068 5107 5110 5122 5134 5139
5157 5160 5217 5255 5318 5388 5397 5415 5444 5512 5524 5571
5608 5647 5677 5701 5751 140 174 288 337 428 436 513
593 612 616 641 648 734 753 764 808 846 859 863
894 950 1052 1064 1066 1139 1186 1249 1277 1321 1326 1409
1475 1559 1674 1726 1858 1871 1879 1942 1948 1950 1959 1991
2097 2129 2143 2196 2367 2590 2678 2774 2788 2841 2877 2986
3085 3174 3244 3309 3357 3370 3437 3497 3525 3546 3601 3702
3815 3869 3889 3898 3907 3908 3956 4009 4092 4121 4197 4258
4300 4375 4574 4649 4717 4764 4867 4887 4928 4980 5000 5112
5235 5390 5391 5395 5431 5442 5488 5678 5721 5727 5744 860
1385 2282 3013 3466 4098 4437 4995 5162 5278 5297 5335];[25
39 63 78 85 130 142 157 160 214 247 260 261
280 323 325 337 355 366 376 406 408 450 453 471
494 572 577 589 647 663 690 705 718 729 756 785
792 817 823 826 876 902 904 939 959 1020 1024 1037
1038 1046 1083 1084 1148 1152 1164 1198 1282 1287 1299 1300
1335 1354 1369 1426 1460 1474 1482 1492 1534 1611 1615 1631
1633 1640 1642 1645 1670 1689 1710 1755 1805 1823 1864 1897
1961 1963 2031 2042 2051 2056 2124 2141 2193 2201 2202 2211
2212 2232 2249 2251 2288 2295 2331 2428 2444 2491 2504 2567
2613 2620 2640 2755 2761 2768 2780 2786 2793 2836 2866 2884
2910 2921 2925 2933 2934 2958 2960 3000 3031 3035 3072 3090
3093 3101 3118 3199 3204 3215 3228 3311 3324 3342 3360 3384
3393 3404 3408 3419 3425 3454 3457 3469 3472 3481 3497 3502
3506 3598 3630 3632 89 181 254 292 315 375 400 405
770 840 888 937 1056 1077 1109 1175 1244 1318 1350 1500
1627 1693 1759 1824 1842 1977 2393 2435 2436 2457 2509 2536
2610 2680 2697 2756 3086 3235 3271 3512 3547 3577 193 215
790 1700 2181 2484 2633 2676 2735 2820 3422 3494];[17 19
28 80 138 142 144 158 160 181 251 252 333 384
430 436 437 461 465 467 524 529 541 569 570 606
656 670 672 673 689 702 717 742 760 783 823 832
858 875 962 1055 1066 1094 1140 1172 1181 1182 1265 1272
1285 1288 1377 1386 1437 1458 1557 1611 1625 1630 1701 1710
1732 1766 1801 1894 1949 1958 1973 2017 2041 2059 2085 2086
2097 2111 2172 2229 2234 2246 2253 2264 2298 2389 2428 2429
2442 2496 2527 2528 2531 2569 2570 2583 2584 2631 2645 2650
2666 2701 2702 2715 2746 2755 2805 2859 15 21 22 253
736 738 763 860 916 968 1071 1128 1137 1157 1163 1279
1309 1310 1348 1444 1581 1655 1657 1684 1691 1713 1731 1779
1798 1815 1878 2010 2095 2100 2137 2163 2178 2233 2235 2284
2292 2345 2460 2624 2676 2717 2756 2806 2893 2894 637 676
771 1612 2046 2353 2441 2660 2791];[4 47 65 68 80 109
154 172 180 200 201 224 239 264 272 274 281 321
330 347 366 367 369 374 380 419 445 458 472 507
521 544 573 581 583 594 609 639 677 722 727 776
799 800 817 839 840 844 856 880 886 904 968 977
991 1009 1012 1052 1054 1065 1090 1101 1147 1175 1301 1302
1336 1353 1422 1429 1456 1480 1513 1515 1627 1639 1644 1665
1666 1669 1678 1687 1726 1805 1827 1837 1850 1865 1880 1887
1912 1940 1967 2001 2002 2019 2035 2124 2141 2219 2243 2334
2340 2350 2468 2473 2476 2494 2544 2561 2572 2573 2619 2635
2666 2667 2674 2755 2789 2804 2822 2848 2850 2866 2980 3055
3067 3073 3078 3107 3113 3125 3162 3170 3179 3184 3235 3254
40 142 243 299 341 417 440 487 508 509 519 558
570 587 602 675 763 814 838 914 943 1038 1048 1071

1073 1104 1117 1138 1163 1204 1265 1340 1408 1646 1706 1793
1901 2024 2094 2186 2217 2336 2343 2432 2501 2536 2593 3029
3068 3132 3153 3181 3244 258 678 2416 2943];[18 30 31
37 52 63 65 126 127 130 134 135 136 154 177
186 191 282 292 332 343 374 508 526 636 651 743
847 905 960 961 963 990 993 1009 1011 1015 1021 1032
1037 1041 1048 1053 1079 1115 1134 1160 1176 1177 1183 1186
1216 1218 1242 1250 1259 1299 1430 1464 1521 1567 1644 1728
1742 1754 1798 1799 1853 1874 1883 1886 1896 1900 1925 1940
1950 2006 2062 2066 2079 2083 2125 2126 2145 2170 2174 2252
2258 2284 2294 2319 2321 2333 2357 2365 2404 2435 2472 2530
2615 2651 2695 2721 2722 2886 2898 2899 2939 2952 2973 2978
2980 2989 3022 3033 3074 3084 3112 3121 3127 3137 3158 3240
3243 3335 3376 3378 3386 3391 3432 3441 3510 3529 3629 180
192 275 281 365 453 455 507 585 770 908 962 997
1012 1047 1055 1077 1222 1385 1476 1498 1541 1734 1836 1945
1956 2050 2140 2291 2328 2421 2444 2465 2559 2600 2602 2613
2643 2669 2733 2764 2787 2791 2792 2854 2891 2902 2916 2982
3041 3071 3093 3094 3346 3348 3448 3450 3452 3539 2481 2750
3387 3521];[12 28 40 72 105 147 192 203 212 227
249 251 263 279 296 340 370 392 423 425 426 434
491 521 532 548 562 609 629 655 663 714 740 743
759 791 852 876 896 924 963 966 969 978 1010 1019
1031 1099 1108 1199 1247 1256 1269 1313 1315 1321 1323 1370
1421 1506 1519 1520 1555 1565 1628 1644 1661 1670 1698 1749
1756 1758 1761 1762 1852 1860 1873 1888 1909 1950 1968 1973
2105 2118 2193 2209 2249 2347 2377 2390 2418 2432 2441 2446
2482 2495 2519 2520 2543 2544 2570 2607 2609 2644 2646 2647
2774 2780 2786 2802 2845 2869 2872 2949 2988 3000 3017 3033
3038 3079 3094 3105 3147 3249 3254 3287 3300 3313 3326 3345
3347 3383 3384 3408 3416 3420 3437 3449 3458 3464 3484 3489
3522 3597 3709 3728 3743 3759 3762 3787 3863 3868 3919 3920
3945 4023 4069 4078 4104 4107 4108 4138 4177 4211 4227 4233
4234 4242 4260 4320 4328 4340 4408 4415 4455 4457 4464 4485
4511 4527 96 102 106 161 204 244 253 258 274 334
374 614 788 929 968 982 1049 1069 1106 1320 1391 1465
1501 1655 1701 1799 1847 1904 1915 1924 1992 1994 2287 2325
2378 2480 2508 2512 2589 2683 2723 2769 2796 2833 2870 2876
2886 2920 3204 3320 3341 3346 3393 3413 3466 3528 3533 3539
3557 3713 3723 3732 3820 3914 4011 4013 4032 4102 4128 4146
4237 4385 4466 186 433 512 744 1182 2384 2427 2602 4187
4492];[19 53 54 93 174 215 249 297 298 322 338
362 365 398 399 401 429 441 467 487 515 516 519
521 567 581 582 681 778 820 850 856 862 903 919
922 923 925 943 948 952 977 980 982 984 1014 1029
1051 1064 1077 1142 1149 1151 1184 1190 1191 1226 1236 1255
1262 1329 1372 1408 1416 1426 1448 1450 1456 1473 1499 1572
1611 1623 1677 1684 1685 1697 1719 1725 1801 1811 1832 1844
1845 1926 1928 1964 1988 1991 2003 2009 2041 2079 2090 2142
2195 2205 2207 2229 2265 2273 2294 2312 2323 2326 2350 2374
2375 2412 2433 2554 2583 2589 2590 2596 2597 2609 2612 2613
2670 2681 2951 2952 2953 3007 3022 3033 3034 3069 3081 3111
3128 3181 3190 3217 3218 3233 3299 3326 3337 3338 3347 45
73 99 105 256 415 420 428 440 478 517 525 684
702 717 787 826 834 985 1027 1045 1115 1128 1131 1176
1247 1386 1449 1481 1538 1740 1884 1963 1992 2001 2040 2126
2129 2145 2204 2220 2253 2398 2406 2693 2778 3174 3309 14
987 1024 1781 2821 3328];[15 17 29 79 141 151 165

190 233 298 305 306 314 342 367 437 459 477 486
496 500 502 516 542 553 566 573 574 632 701 708
721 808 819 838 890 897 907 923 975 976 981 1010
1019 1034 1050 1061 1069 1142 1146 1172 1180 1247 1318 1343
1344 1374 1392 1415 1548 1567 1580 1584 1614 1617 1627 1632
1645 1737 1753 1849 1851 1863 1893 1927 1937 1981 2001 2021
2027 2033 2038 2079 2096 2147 2167 2230 2244 2247 2248 2255
2275 2294 2324 2361 2381 2400 2421 2484 2539 2545 2563 2600
2660 2684 2696 2697 2700 2704 2741 2769 2810 2848 2879 2884
2891 2906 2943 2970 2973 2982 3002 3056 3062 3108 3116 3127
3136 3143 3173 3185 3227 3284 3289 3338 3340 3363 3372 3457
3497 3507 3523 3524 7 114 157 170 219 418 491 596
667 695 786 796 876 1006 1033 1114 1127 1185 1198 1255
1326 1417 1446 1465 1657 1751 1760 1767 1774 1907 2152 2343
2366 2371 2466 2511 2536 2572 2628 2639 2640 2731 2826 2900
2948 3223 3233 3298 3347 3425 3462 569 1000 2202 2422 2606
2687 3482];[15 41 58 73 79 80 92 120 147 176
228 251 276 286 312 333 336 354 357 360 418 532
559 576 611 621 632 633 642 647 665 666 700 723
729 746 750 760 788 803 813 826 873 881 893 894
896 940 976 988 1023 1045 1057 1058 1082 1098 1124 1125
1179 1191 1201 1202 1213 1235 1238 1246 1286 1319 1332 1336
1337 1342 1353 1442 1545 1551 1587 1590 1609 1666 1694 1722
1743 1816 1925 1939 1961 1999 2049 2066 2094 2129 2191 2211
2272 2321 2373 2401 2413 2497 2524 2537 2582 2587 56 98
201 295 419 483 551 585 629 718 749 841 1031 1088
1126 1137 1252 1283 1315 1349 1364 1489 1572 1593 1594 1726
1833 1838 1841 1876 1880 1986 2001 2016 2152 2173 2188 2232
2339 2408 2452 2468 2586 2569];[15 62 70 128 153 155
206 216 219 245 261 346 365 396 492 496 521 583
586 616 636 645 664 700 715 725 740 769 831 886
937 938 949 1002 1014 1016 1048 1074 1082 1085 1105 1160
1197 1238 1245 1302 1303 1310 1336 1361 1373 1376 1433 1435
1460 1521 1552 1573 1596 1624 1631 1681 1693 1694 1724 1766
1768 1783 1813 1842 1934 1935 1945 1953 1976 1989 2011 2042
2063 2068 2100 2105 2121 2285 2321 2344 2363 2392 2407 2412
2426 2450 2456 2463 2469 2476 2487 2519 2544 2560 2570 2576
2583 2612 2627 2659 2662 2698 2714 2719 2732 2748 2857 2929
2949 2996 3019 3057 3058 3059 3069 3087 3096 3134 3143 3167
3238 3290 3294 3336 3343 3368 3374 3411 3434 3450 3537 3555
3560 3561 3588 3589 3596 3615 3644 3646 3668 3677 3732 3899
3985 4025 4034 4035 4036 4103 4140 4182 4207 4229 4231 4268
4275 4282 4348 4371 4389 4461 4462 4483 4499 4577 4581 4636
4662 4688 4801 4802 4835 4858 96 188 220 222 236 239
263 264 283 319 430 436 458 523 578 647 667 673
777 781 790 935 1086 1111 1125 1289 1432 1459 1555 1745
1836 1847 1866 1901 1951 2005 2023 2383 2508 2562 2571 2620
2839 2940 3039 3106 3175 3271 3272 3455 3787 3817 4058 4069
4107 4178 4368 4466 4568 649 1990 3769 4356 4852];[7 8
16 37 54 93 120 150 169 270 278 293 331 339
358 386 401 423 427 459 505 509 518 529 575 589
644 676 743 747 776 801 803 805 878 889 910 915
933 937 950 969 996 1083 1153 1178 1205 1226 1263 1274
1328 1405 1512 1518 1533 1539 1540 1593 1617 1656 1659 1661
1699 1707 1759 1777 1805 1823 1834 1837 1850 1851 1858 1866
1887 1907 1908 1959 1980 2005 2007 2030 2044 2139 2154 2170
2175 2187 2204 2255 2267 2302 2340 2369 2383 2413 2480 2503
2513 2573 2574 72 97 113 149 164 220 261 279 294

```
545   787  1024  1096  1105  1117  1137  1168  1282  1344  1433  1444
1594  1783  1833  1869  1933  2202  2206  2256  2399  2434  2566   517
930  1679  2111  2221];[40    53    68    82    85   122   137   153
258   287   291   312   350   357   366   406   446   474   477   478
486   487   497   530   547   552   554   602   608   617   618   638
665   714   726   753   771   777   786   833   857   884   887   899
908   940  1015  1016  1018  1024  1026  1028  1047  1074  1092  1106
1117  1172  1237  1275  1322  1334  1349  1357  1383  1404  1422  1450
1476  1623  1659  1676  1702  1742  1756  1836  1854  1856  1857  1858
1887  1934  1944  1951  2032  2039  2056  2062  2066  2079  2089  2164
2253  2310  2335  2348  2389  2398  2403  2422  2434  2435  2436  2439
2442  2443  2444  2451  2475  2534  2538  2596  2614  2674   151   207
342   405   413   438   522   556   597   654   664   784   901   920
923   936   995  1003  1143  1154  1159  1176  1214  1235  1238  1417
1437  1521  1562  1592  1748  1774  1795  1798  1859  1886  1912  1938
2047  2229  2309  2414  2421  2424  2478  2483  2653  2667  2691    39
545  1201  2288  2481];[1    40    59    95   120   179   187   192   197
199   253   359   367   406   443   506   526   536   584   669   713
715   722   739   740   751   756   762   832   915   921   939   956
957   981   982   994  1018  1056  1061  1079  1099  1111  1160  1163
1201  1248  1317  1318  1338  1360  1398  1445  1446  1450  1551  1592
1605  1606  1633  1637  1660  1689  1715  1765  1779  1791  1803  1824
1869  1882  1914  1928  1947  1968  1983  1994  2033  2068  2115  2131
2167  2175  2186  2189  2200  2277  2278  2284  2299  2343  2364  2386
2397  2414  2416  2446  2448  2456  2483  2500    57   161   240   434
529   531   594   752   811   874   879   880   909   925   934   955
965  1100  1155  1283  1303  1319  1397  1703  1707  1717  1723  1786
1817  1958  1964  2203  2381  2439  2463  2524   152   404   929  1451
1716  2124  2198  2357];[2     3     5    36    74    75    78    79
90   134   191   218   265   276   298   341   373   379   385   479
517   545   551   562   614   617   622   624   694   704   736   762
777   802   808   836   859   864   872   880   895   901   942   963
994  1035  1037  1065  1086  1293  1319  1329  1411  1429  1439  1476
1517  1521  1565  1574  1584  1609  1614  1624  1632  1640  1660  1672
1704  1707  1775  1781  1806  1835  1837  1845  1848  1851  1852  1883
1906   117   165   220   228   232   364   366   387   539   801   885
929   992   997  1031  1066  1169  1201  1320  1389  1465  1487  1509
1596  1703  1716  1802  1862  1867  1875  1917  1148  1843];[1    35
71    85   120   176   179   180   194   199   246   310   322   364
383   384   392   433   434   443   457   468   475   495   517   536
558   585   589   596   623   641   658   671   678   724   870   899
902   919   927   957   967  1014  1020  1202  1296  1329  1345  1375
1391  1396  1430  1454  1456  1477  1530  1590  1640  1644  1694  1697
1704  1735  1758  1816  1849  1899  1929  1940  1964  1971  1972  2032
2111  2180  2184  2206  2246  2248  2313  2321  2359  2376  2385  2391
2394  2430  2511  2524  2537  2570  2575  2586  2632  2645  2651  2678
2688  2697  2716  2739  2805  2816  2840  2847  2853  2892  2899  2901
2903  2925  2937  2944  2945  2949  2958  2973  3021  3057  3059  3066
3070  3088  3135  3188  3226  3274  3282  3302  3305  3306  3310  3311
3329  3332  3348  3365  3381  3395  3419  3478  3515  3550  3558  3560
3577  3633  3637  3659  3694  3713  3714     5    20    55   275   319
471   590   698   834   941  1012  1018  1191  1270  1312  1344  1400
1724  1833  1880  2181  2182  2247  2384  2404  2549  2584  2631  2806
2854  3068  3099  3112  3140  3237  3284  3354  3422  3437  3516  3581
3720   497   531  1061  1126  1301  1315  1388  2463  2968];[52    61
98   110   129   136   143   178   181   192   288   313   319   339
344   355   368   396   402   417   424   441   499   519   567   569
577   583   584   629   682   715   745   758   767   811   815   828
```

837 844 851 886 896 906 930 936 1019 1060 1163 1164
1214 1239 1255 1261 1278 1281 1295 1301 1302 1320 1342 1363
1420 1442 1454 1461 1465 1549 1561 1563 1565 1566 1592 1614
1634 1636 1645 1663 1664 1677 1706 1709 1711 1740 1757 1796
1804 1813 1844 1889 1890 1901 1903 1907 1969 1970 2002 2012
2034 2081 2131 2178 2212 2308 2312 2326 2338 2408 2540 2568
2584 2588 330 337 422 454 621 638 747 965 977 1013
1030 1139 1202 1314 1405 1538 1739 1776 1790 1807 1902 2022
2105 2171 2325 2412 2450 2453 802 816 1184 2170 2345];[5
6 17 50 71 100 150 156 168 199 200 218 282 298
308 317 334 366 411 428 433 438 504 514 521 577
581 590 602 606 691 713 753 798 810 823 842 852
930 936 943 973 980 1066 1128 1137 1167 1172 1183 1187
1223 1224 1272 1279 1301 1319 1322 1352 1396 1418 1426 1431
1498 1514 1540 1591 1603 1607 1687 1696 1703 1761 1803 1805
1855 1890 1899 2022 2028 2063 2071 2072 2086 2112 2123 2135
2185 2191 2228 2254 2260 2330 2410 2412 2431 2435 2452 2498
2512 2572 2577 2602 2649 2681 2689 2718 2792 2819 2823 2830
2857 2875 2876 2881 2891 2903 2905 2929 2962 3044 3048 3056
3081 3108 3124 3179 3212 3240 3259 3261 3313 3320 3331 3344
3348 3365 3375 3385 3390 3398 3399 3414 8 96 256 371
458 496 579 622 742 840 874 970 1034 1104 1138 1190
1250 1432 1468 1553 1677 1700 1827 1862 1953 1982 2189 2265
2319 2362 2407 2507 2583 2675 2799 2911 2968 3005 3041 3058
3094 3121 3195 3299 3407 787 1363 1560 1823 2307 2360 2408
2445 2959 3103];[1 12 15 28 32 34 36 68 78
93 107 113 186 195 217 230 239 255 282 292 319
374 384 385 422 434 452 454 463 477 501 534 545
573 662 770 773 837 840 850 856 864 865 882 894
947 1033 1093 1107 1121 1140 1141 1145 1194 1215 1230 1273
1282 1297 1300 1315 1326 1328 1330 1351 1365 1439 1484 1490
1587 1608 1643 1646 1663 1673 1700 1707 1724 1728 1750 1769
1777 1805 1823 1828 1832 1835 1866 1873 1894 1911 1916 1956
1971 1987 2000 2003 2009 2020 2034 2039 2103 2170 2180 2188
2238 2351 2352 2357 2381 2433 2454 2461 2572 2596 2604 2617
2642 2650 2657 2665 2666 2690 2719 2735 2736 2762 2770 2786
2790 2812 2824 2838 2876 2897 2903 2924 2953 3030 3068 3111
3141 3151 3152 3243 3314 3316 3330 3359 3386 3407 3422 143
271 295 346 508 565 745 871 877 896 913 919 1134
1342 1430 1514 1629 1838 1839 1861 1941 1980 1983 2032 2100
2223 2224 2227 2311 2386 2388 2392 2472 2522 2703 2740 2863
2872 2946 3014 3072 3082 3085 3089 3153 3170 3262 3319 3404
1234 1891 1944 2387 2843 3215];[59 115 116 217 252 265
331 349 361 396 422 424 442 465 509 517 533 539
547 549 563 569 614 615 652 691 693 715 743 769
773 799 831 832 846 893 894 925 953 964 987 997
1018 1057 1059 1061 1073 1115 1127 1148 1157 1181 1224 1227
1292 1314 1338 1348 1371 1397 1398 1402 1420 1463 1499 1514
1530 1551 1626 1708 1711 1724 1738 1799 1802 1806 1813 1815
1818 1841 1854 1874 1880 1911 1920 1924 1944 1951 2055 2058
2077 2103 2121 2124 2153 2160 2172 2173 2276 2315 2323 2362
2426 2448 2512 2530 2543 2551 2577 2600 2642 2644 2656 2706
2715 2727 2738 2798 2803 2823 2825 2851 2853 2869 2909 3001
3002 3017 3068 3089 3110 3114 3123 3127 3138 3148 3213 3215
3225 3280 3301 3315 3320 67 285 414 417 753 806 807
822 870 940 981 984 1269 1296 1477 1579 1637 1639 1713
2099 2155 2156 2164 2183 2210 2248 2275 2420 2468 2678 2700
2737 2777 2897 2999 3124 3200 3245 1223 1466 1761 1958 2247

```
2439  3187  3224];[47   130   170   195   217   274   347   350   362
365   373   399   413   414   431   499   609   612   635   697   707
711   712   724   749   759   760   766   773   784   786   887   896
899   905   933   953   994  1006  1008  1009  1027  1087  1092  1098
1134  1136  1205  1269  1270  1320  1362  1364  1381  1396  1410  1427
1462  1479  1502  1525  1531  1583  1591  1599  1612  1614  1630  1656
1683  1694  1748  1836  1886  1929  1937  1954  1980  2008  2066  2083
2086  2105  2108  2117  2128  2141  2217  2218  2264  2346  2356  2391
2407  2408  2416  2458  2460  2465  2471  2568  2581  2590  2596    95
183   201   384   393   768   779   812   844   906   998  1001  1007
1131  1140  1403  1420  1516  1520  1557  1561  1605  1651  1679  1745
1746  1942  1985  2129  2166  2194  2274  2371  2427  2481  2514  2536
2556  2578  2629   419   596  1536  1818  1932  2023  2450];};
% SCORES=[5 6];
% ns={[15    22    32    35    77    83   106   120   167   181   261
272   315   327   329   330   336   350   355   358   365   370   388
409   412   420   467   477   494   499   502   549   555   557   573
612   625   662   698   716   731   734   792   803   809   893   895
923   996  1021  1032  1059  1079  1094  1104  1120  1153  1166  1171
1177  1218  1222  1251  1257  1269  1307  1311  1324  1329  1340  1347
1358  1369  1376  1422  1426  1469  1520  1551  1555  1566  1580  1586
1595  1600  1609  1619  1628  1645  1657  1660  1673  1726  1730  1747
1748  1768  1778  1875  1881  1887  1888  1953  2007  2014  2027  2037
2042  2082  2089  2090  2091  2173  2204  2210  2231  2232  2296  2302
2311  2352  2365  2366  2376  2413  2430  2432  2472  2528  2530  2531
2544  2571  2594  2599    97   353   423   552   667   707   733   835
1127  1143  1200  1239  1309  1410  1494  1633  1725  1776  1789  1828
1926  2176  2239  2301  2324  2372  2374];[1    36    43    45    52
68    89   108   140   148   173   186   199   244   298   313   323
324   367   399   428   430   486   514   523   524   535   563   605
608   612   615   616   621   638   648   675   685   702   740   753
764   776   789   796   802   822   846   859   860   871   916   950
1004  1021  1057  1064  1065  1113  1139  1161  1170  1186  1205  1249
1262  1274  1277  1278  1298  1311  1313  1314  1321  1360  1365  1372
1385  1472  1482  1491  1495  1501  1507  1509  1519  1559  1565  1578
1619  1620  1622  1652  1664  1670  1674  1678  1685  1726  1788  1790
1809  1814  1819  1860  1863  1869  1871  1872  1876  1879  1899  1915
1919  1920  1929  1941  1942  1948  1977  1991  2037  2079  2089  2097
2143  2163  2244  2246  2250  2258  2266  2280  2284  2330  2356  2363
2367  2416  2439  2490  2498  2523  2545  2546  2554  2562  2570  2590
2625  2678  2691  2706  2735  2754  2769  2806  2813  2840  2845  2878
2927  2937  2952  2982  2986  2998  3002  3008  3013  3014  3036  3083
3085  3086  3099  3128  3142  3143  3157  3161  3174  3179  3183  3187
3203  3213  3224  3242  3248  3250  3295  3309  3350  3370  3386  3402
3405  3425  3437  3450  3465  3468  3474  3513  3525  3546  3565  3600
3601  3626  3636  3683  3702  3714  3838  3889  3898  3907  3908  3919
3956  3975  4009  4051  4061  4069  4071  4092  4098  4100  4102  4110
4151  4155  4182  4186  4191  4217  4221  4251  4258  4297  4300  4320
4404  4418  4423  4444  4455  4462  4472  4484  4494  4513  4533  4542
4565  4566  4574  4631  4644  4648  4649  4656  4684  4691  4760  4788
4799  4811  4813  4825  4844  4862  4908  4928  4944  4947  4991  4995
4997  4998  4999  5000  5074  5093  5107  5110  5112  5150  5157  5160
5161  5175  5206  5232  5284  5327  5335  5359  5390  5395  5431  5442
5498  5512  5524  5529  5563  5564  5569  5571  5624  5653  5654  5701
5721  5727  5744  5748  5797   251   287   296   387   445   573   607
745   840   879   913   917  1038  1076  1171  1322  1410  1414  1504
1601  1689  1705  1737  1792  1817  1908  1931  1971  1979  3178  3184
3336  3740  3989  4331  4487  4598  4850  4875  4899  4901  5001  5217
```

5308 5504 5628 5647 5678];[28 63 123 157 160 235 254
260 261 268 291 292 298 315 323 325 331 344 355
375 376 405 471 507 544 577 660 663 675 676 690
705 712 717 729 750 756 770 790 792 823 826 857
876 888 904 915 940 1024 1037 1046 1047 1056 1083 1106
1113 1148 1152 1164 1175 1182 1193 1244 1273 1287 1299 1318
1331 1354 1378 1390 1420 1426 1460 1474 1482 1491 1517 1519
1534 1599 1615 1627 1631 1640 1642 1645 1662 1670 1689 1693
1697 1719 1776 1823 1824 1842 1848 1857 1858 1864 1900 1910
1927 1961 1963 1967 1995 2031 2047 2049 2051 2056 2124 2126
2178 2193 2233 2249 2257 2288 2295 2311 2321 2394 2397 2444
2457 2484 2504 2521 2536 2549 2567 2609 2613 2620 2640 2697
2755 2768 2780 2786 2793 2815 2820 2829 2836 2866 2889 2910
2921 2934 2958 3020 3044 3069 3072 3086 3093 3101 3159 3162
3199 3216 3228 3230 3235 3239 3271 3311 3312 3324 3334 3342
3349 3360 3384 3386 3393 3408 3419 3453 3457 3472 3482 3494
3502 3506 3513 3547 3555 3572 3582 3615 3620 3632 64 117
453 454 816 1307 1338 1359 1500 1590 1622 1633 1691 1692
1977 2196 2251 2269 2330 2428 2674 3060 3192 3223];[14 17
21 43 62 79 160 251 253 256 259 296 350 358
412 436 463 467 491 517 529 541 556 569 570 597
669 672 673 685 689 691 711 717 736 742 763 783
784 788 823 855 860 909 910 940 952 953 962 968
1020 1032 1043 1053 1064 1071 1079 1094 1125 1128 1139 1157
1163 1172 1182 1255 1265 1272 1285 1288 1289 1310 1337 1344
1383 1396 1405 1464 1516 1537 1542 1553 1558 1566 1591 1612
1635 1655 1657 1659 1668 1713 1714 1731 1733 1743 1750 1822
1824 1847 1878 1894 1973 2010 2017 2041 2059 2086 2093 2097
2163 2177 2178 2203 2219 2229 2233 2235 2292 2295 2298 2357
2379 2388 2389 2430 2438 2442 2460 2464 2496 2509 2527 2528
2547 2569 2574 2584 2629 2645 2650 2660 2686 2702 2703 2755
2806 2813 2830 2859 2893 2894 465 610 658 683 705 756
1031 1077 1129 1192 1339 1461 1492 1557 1568 1576 1581 1751
1830 2043 2077 2216 2472 2588 2791 2870];[52 76 81 83
120 165 171 186 194 201 224 243 270 274 276 330
366 367 374 417 440 458 479 489 503 507 508 509
519 523 543 544 570 573 577 581 583 594 609 644
665 675 676 677 712 727 775 776 800 814 824 835
839 844 880 904 983 1009 1012 1048 1071 1072 1073 1090
1094 1104 1138 1162 1198 1278 1289 1299 1301 1302 1336 1342
1353 1364 1368 1415 1429 1456 1497 1513 1515 1521 1576 1627
1644 1665 1706 1736 1781 1802 1811 1827 1828 1837 1880 1887
1901 1912 1939 1940 1956 1967 2008 2012 2019 2035 2068 2081
2087 2186 2190 2199 2217 2219 2241 2255 2291 2343 2348 2350
2364 2377 2416 2417 2468 2473 2476 2501 2516 2529 2535 2544
2561 2587 2619 2635 2637 2673 2687 2697 2704 2755 2789 2804
2822 2848 2932 2944 2948 2965 2974 2997 2998 3026 3055 3067
3068 3132 3151 3153 3162 3174 3184 3203 3209 3212 3254 3255
3260 3283 15 118 178 436 541 783 879 929 994 1079
1099 1176 1181 1231 1518 1563 1758 1784 1799 2086 2123 2124
2243 2475 2609 2957 3005 3048 3265 3285];[9 18 30 51
52 53 55 63 66 111 147 156 160 180 191 192
255 268 275 289 292 374 387 414 439 452 454 455
456 477 522 585 625 642 651 665 708 723 743 810
862 905 908 921 955 959 961 963 978 984 1010 1015
1032 1033 1037 1044 1047 1053 1055 1066 1067 1077 1079 1087
1124 1135 1161 1163 1174 1205 1211 1213 1216 1218 1220 1238
1241 1242 1250 1259 1263 1278 1285 1333 1385 1397 1411 1430

```
1453  1464  1468  1476  1521  1524  1541  1591  1691  1754  1828  1836
1839  1853  1868  1874  1886  1900  1905  1931  1940  1945  1946  1955
1976  1994  2003  2036  2062  2126  2140  2174  2181  2193  2243  2252
2281  2294  2297  2362  2363  2404  2421  2435  2436  2444  2471  2472
2506  2541  2599  2602  2605  2615  2632  2669  2733  2787  2791  2792
2818  2821  2857  2891  2898  2908  2939  2940  2952  2955  2961  2973
2980  2982  2999  3005  3032  3040  3041  3071  3084  3088  3121  3137
3141  3152  3155  3161  3166  3229  3239  3240  3333  3348  3376  3391
3441  3449  3450  3462  3515  3527  3529  3580  3629  3661  3671  3675
3714  3718    27   134   136   138   177   288   425   523   565   789
873   990  1003  1012  1123  1184  1225  1248  1253  1302  1382  1498
1751  1797  1915  1918  2228  2304  2346  2384  2533  2559  2582  2763
2835  2850  2854  2888  3033  3093  3114  3156  3346  3383  3414  3440
3521];[28     40    72    78    96   102   138   147   186   192   196
218   244   249   256   274   299   317   331   334   340   361   370
400   423   425   472   498   516   521   526   532   541   548   562
570   588   593   609   663   664   714   743   754   777   788   837
843   852   875   876   891   966   967   968   969   982  1031  1049
1106  1125  1152  1199  1256  1269  1279  1296  1313  1316  1323  1339
1364  1376  1395  1450  1464  1500  1501  1506  1520  1555  1617  1652
1655  1661  1670  1701  1747  1756  1758  1762  1799  1832  1852  1867
1873  1904  1907  1909  1915  1924  1941  1954  1968  1972  1985  1990
1992  1994  2038  2118  2193  2209  2287  2295  2325  2347  2360  2383
2390  2402  2404  2407  2432  2441  2496  2504  2512  2520  2536  2539
2544  2549  2568  2570  2589  2646  2669  2723  2757  2761  2766  2774
2790  2796  2802  2805  2830  2844  2872  2886  2920  2949  2953  3000
3017  3033  3043  3105  3147  3214  3236  3247  3249  3254  3256  3258
3296  3320  3341  3346  3347  3408  3413  3416  3437  3447  3449  3466
3484  3512  3514  3515  3518  3522  3528  3533  3539  3606  3670  3709
3713  3728  3732  3733  3743  3767  3782  3787  3790  3820  3829  3839
3850  3879  3890  3914  3920  3923  3945  3994  4013  4023  4032  4040
4069  4070  4078  4102  4108  4120  4128  4138  4146  4177  4181  4187
4188  4211  4234  4237  4240  4242  4260  4288  4308  4328  4351  4385
4408  4415  4455  4457  4463  4466  4511  4527  4528   203   224   227
248   545   586   694   861  1178  1189  1248  1414  1472  1546  1579
1592  1709  1710  1854  2503  2519  2535  2645  2865  2968  3050  3417
3464  3592  3616  3618  3750  3863  4011  4054  4227  4233  4346
4473];[45     47    73    93   105   106   129   174   249   256   290
294   297   298   354   365   384   398   401   428   436   438   441
442   467   478   482   487   515   521   525   550   567   581   582
630   641   667   681   698   702   717   739   766   820   826   850
856   903   924  1014  1064  1077  1107  1115  1128  1151  1152  1176
1184  1191  1226  1236  1247  1262  1300  1310  1329  1372  1408  1448
1449  1450  1456  1473  1481  1499  1538  1569  1607  1608  1622  1634
1684  1740  1752  1801  1813  1832  1845  1884  1897  1918  1935  1988
1991  1992  2003  2009  2013  2033  2040  2041  2079  2085  2126  2129
2146  2177  2204  2214  2220  2229  2230  2253  2255  2273  2304  2312
2323  2326  2350  2375  2398  2406  2412  2448  2552  2554  2583  2589
2597  2609  2613  2670  2690  2693  2698  2741  2748  2778  2821  2878
2895  2924  2953  3007  3034  3063  3069  3111  3116  3131  3139  3181
3190  3218  3222  3233  3280  3299  3326  3337  3338  3357     4    53
215   399   440   617   697   742   763   840   919   977  1015  1288
1301  1330  1363  1462  1789  1862  1958  2027  2105  2881  2963
3060];[17     76    81    86   114   128   129   151   157   162   165
170   190   197   211   219   257   272   314   320   332   355   367
418   459   477   480   486   489   491   502   516   542   553   568
573   667   683   695   708   721   728   736   746   754   762   805
808   876   890   897   906   923   937   976  1031  1033  1102  1136
```

1142 1149 1180 1218 1221 1223 1233 1247 1272 1306 1326 1344
1374 1414 1446 1480 1482 1492 1504 1515 1523 1540 1584 1613
1614 1617 1632 1645 1696 1698 1738 1748 1751 1755 1761 1798
1811 1863 1907 1927 1934 1937 1996 2001 2003 2021 2038 2079
2096 2122 2152 2155 2205 2246 2251 2276 2278 2294 2298 2324
2343 2361 2377 2400 2421 2466 2487 2532 2536 2539 2563 2600
2629 2640 2660 2704 2731 2732 2741 2784 2787 2810 2826 2832
2842 2848 2884 2900 2901 2906 2909 2919 2943 2948 2949 2952
2970 2979 2982 2990 2995 3026 3051 3056 3062 3094 3108 3116
3127 3136 3144 3154 3173 3212 3233 3263 3281 3283 3289 3338
3340 3363 3367 3425 3454 3457 3462 3497 3523 79 85 254
501 575 608 905 912 918 1050 1061 1095 1129 1300 1426
1488 1852 1898 1945 1979 2033 2243 2291 2340 2398 2506 2605
2709 2756 2809 2818 2864 3110 3115 3146 3257];[7 41 53
56 58 61 65 73 80 92 120 147 172 176 201
251 252 315 334 336 360 370 373 419 448 462 470
483 500 514 540 543 551 576 585 629 642 643 666
718 748 749 752 760 788 803 813 826 841 845 873
874 888 894 945 948 1023 1024 1027 1031 1052 1057 1058
1088 1126 1132 1137 1171 1179 1201 1202 1209 1213 1235 1237
1252 1315 1317 1319 1336 1337 1341 1364 1537 1545 1587 1609
1624 1652 1653 1674 1711 1714 1722 1726 1734 1744 1780 1800
1813 1815 1816 1864 1876 1880 1909 1910 1939 1961 1999 2001
2003 2016 2066 2081 2087 2089 2115 2129 2134 2173 2232 2272
2305 2321 2401 2413 2467 2472 2473 2497 2519 2537 2586 2587
57 295 326 357 372 531 600 611 621 691 815 1124
1139 1236 1332 1534 1541 1584 1623 1677 1981];[62 69 74
122 128 153 155 188 206 216 219 220 222 236 239
244 245 261 263 264 283 336 353 365 388 430 436
452 458 464 476 496 521 540 555 576 578 583 616
631 636 645 647 673 684 700 725 726 739 740 744
750 765 769 776 777 886 889 910 935 938 940 949
1002 1011 1016 1018 1024 1074 1086 1105 1111 1125 1131 1133
1141 1160 1198 1207 1287 1303 1336 1376 1431 1432 1433 1435
1446 1498 1513 1524 1555 1564 1573 1620 1624 1631 1641 1653
1730 1745 1746 1768 1769 1771 1794 1836 1842 1847 1848 1866
1873 1901 1918 1945 1953 1989 2011 2042 2051 2068 2081 2100
2114 2121 2130 2138 2153 2196 2214 2242 2321 2354 2363 2390
2392 2407 2410 2433 2450 2452 2456 2474 2507 2514 2519 2521
2544 2553 2570 2571 2576 2577 2583 2602 2612 2620 2627 2659
2697 2698 2714 2753 2765 2810 2839 2847 2867 2888 2924 2954
2996 2997 3036 3051 3057 3058 3059 3096 3124 3133 3134 3166
3220 3238 3311 3336 3343 3353 3379 3416 3450 3482 3508 3525
3537 3560 3565 3570 3584 3589 3596 3615 3659 3668 3699 3732
3749 3780 3787 3791 3796 3812 3817 3818 3853 3903 3963 3985
4002 4020 4034 4042 4058 4066 4069 4103 4112 4178 4207 4221
4229 4230 4332 4337 4342 4348 4353 4389 4462 4466 4488 4499
4515 4575 4579 4581 4604 4624 4641 4653 4654 4662 4705 4709
4766 4802 4835 4845 246 451 457 652 775 863 1211 1232
1282 1378 1481 1614 1625 1814 2202 2218 2240 2332 2458 2469
2502 2525 2533 2555 2581 2597 2599 2761 2811 3143 3385 3404
3424 3667 3738 3900 3955 4094 4119 4154 4181 4368 4541 4573
4739 4760 4870];[16 34 39 54 72 93 97 113 132
150 164 168 184 223 242 261 270 294 307 331 339
341 358 386 401 438 451 463 486 509 529 545 546
575 589 613 656 667 676 743 747 750 765 775 776
795 801 815 889 910 915 930 933 937 950 954 955
996 997 1018 1071 1096 1117 1153 1168 1205 1238 1263 1274

1282 1319 1344 1348 1351 1396 1408 1413 1443 1444 1445 1476
1505 1512 1518 1539 1540 1542 1593 1660 1666 1724 1728 1736
1742 1777 1820 1823 1851 1859 1907 1908 1911 1915 1951 1959
2005 2007 2030 2031 2139 2175 2187 2202 2206 2223 2256 2258
2264 2267 2294 2320 2333 2340 2399 2413 2480 2503 2553 2559
26 32 37 41 137 149 201 278 288 547 558 1051
1083 1246 1533 1538 1655 1750 1783 1980 2067 2154 2181 2197
2250 2308 2326 2416 2434 2468];[40 41 43 53 82 85
137 142 151 153 154 180 202 258 307 312 341 351
401 407 413 419 474 477 478 488 497 522 534 541
543 556 560 602 617 654 665 753 777 778 784 785
792 833 857 887 899 900 908 920 923 940 958 991
995 1003 1024 1047 1073 1074 1075 1106 1118 1143 1150 1154
1155 1159 1172 1174 1176 1201 1202 1235 1238 1275 1301 1342
1353 1363 1421 1422 1450 1500 1534 1562 1599 1603 1633 1650
1659 1715 1717 1742 1748 1796 1798 1836 1854 1859 1885 1886
1912 1936 1938 1944 1965 1989 2032 2039 2080 2081 2123 2129
2134 2147 2164 2229 2301 2335 2348 2358 2375 2403 2414 2421
2422 2434 2435 2436 2440 2442 2475 2478 2534 2582 2592 2596
2667 2691 39 54 60 150 326 552 667 691 713 842
925 1027 1098 1475 1506 1608 1720 1751 1759 1788 1847 2037
2110 2182 2215 2451 2483];[1 54 60 95 109 120 152
161 179 192 197 240 253 254 272 341 342 406 421
434 444 459 461 506 526 531 536 544 584 594 669
713 715 722 756 774 797 832 874 876 880 909 915
921 934 939 955 956 957 965 972 982 983 989 1008
1032 1041 1056 1061 1075 1079 1100 1111 1155 1181 1201 1234
1248 1259 1303 1306 1318 1319 1338 1348 1387 1397 1398 1428
1446 1450 1466 1478 1480 1551 1558 1572 1583 1592 1605 1636
1651 1660 1680 1715 1717 1765 1778 1779 1786 1800 1803 1822
1823 1868 1869 1882 1887 1889 1947 1976 1983 2033 2068 2101
2131 2177 2203 2245 2270 2277 2299 2301 2341 2364 2416 2439
2446 2457 2463 2483 2500 2523 2524 404 496 621 651 718
741 932 941 1174 1203 1283 1298 1301 1575 1716 1994 2189
2200 2229 2351 2394 2434];[36 55 62 75 78 92 103
160 165 191 197 232 265 298 310 334 352 366 372
379 385 397 428 471 475 479 539 559 562 575 583
594 660 685 697 707 724 755 791 803 825 831 843
872 875 954 963 964 992 994 1031 1035 1037 1066 1086
1193 1201 1247 1252 1293 1303 1307 1320 1321 1389 1392 1430
1447 1478 1487 1521 1524 1537 1545 1574 1584 1606 1609 1614
1615 1660 1703 1715 1738 1747 1758 1771 1772 1775 1781 1806
1837 1845 1848 1851 1867 1883 1884 1906 196 210 218 297
551 662 694 723 864 888 965 1141 1290 1506 1640 1697
1704 1875 1913 1917];[35 71 74 124 133 170 179 194
258 269 281 284 285 310 319 330 364 392 413 432
433 434 438 457 471 495 504 518 558 589 596 636
641 658 671 698 700 703 706 724 771 814 830 834
899 919 927 941 958 967 991 997 1012 1061 1062 1094
1152 1197 1202 1224 1296 1300 1301 1312 1315 1344 1345 1388
1391 1396 1400 1456 1477 1503 1564 1640 1644 1645 1656 1697
1704 1735 1746 1758 1760 1776 1793 1799 1816 1833 1863 1865
1880 1882 1919 1925 1940 1945 1957 1961 1972 1987 2011 2032
2055 2090 2111 2174 2180 2248 2255 2322 2345 2347 2370 2384
2391 2490 2524 2537 2549 2575 2583 2586 2618 2631 2632 2645
2647 2651 2688 2697 2698 2772 2805 2816 2847 2854 2860 2878
2881 2892 2925 2932 2937 2944 2949 2952 2956 3021 3057 3059
3066 3068 3088 3099 3112 3124 3136 3140 3160 3181 3182 3188

3226 3266 3273 3284 3292 3302 3306 3311 3322 3329 3332 3365
3370 3381 3395 3404 3408 3411 3419 3422 3433 3451 3454 3471
3478 3496 3512 3515 3550 3558 3633 3673 3690 3694 3713 3727
3731 3737 3752 204 253 537 538 540 643 738 886 947
976 1209 1230 1270 1485 1510 1694 1889 1949 2009 2184 2245
2379 2548 2637 2716 2872 2973 3070 3214 3237 3310 3392 3518
3738];[64 81 89 98 102 143 181 192 197 252 270
284 288 304 319 330 337 339 342 344 345 355 356
357 396 402 423 424 431 485 491 499 539 565 569
584 621 629 671 674 682 715 721 745 747 757 758
811 813 815 816 828 844 896 910 930 936 950 965
997 999 1060 1139 1164 1178 1209 1214 1241 1261 1266 1273
1276 1277 1281 1295 1314 1342 1363 1412 1413 1417 1420 1442
1453 1454 1462 1489 1538 1541 1549 1561 1573 1601 1634 1663
1664 1677 1697 1706 1709 1711 1739 1740 1751 1776 1790 1807
1814 1840 1844 1883 1885 1903 1936 1970 1999 2012 2036 2083
2102 2124 2131 2212 2224 2241 2262 2265 2283 2288 2312 2326
2337 2408 2449 2461 2500 2563 2579 34 114 281 351 418
594 788 851 981 1090 1129 1517 1527 1902 1940 2004 2123
2141 2166 2292 2416 2428 2531];[6 50 60 71 122 156
168 199 213 217 227 236 256 282 308 314 323 335
366 376 419 421 428 452 458 496 504 551 581 598
602 609 680 683 742 798 810 822 840 842 852 874
875 930 935 943 973 981 1025 1028 1062 1073 1127 1128
1138 1172 1187 1223 1224 1250 1272 1286 1297 1301 1319 1322
1330 1348 1352 1362 1363 1396 1418 1426 1437 1464 1468 1526
1537 1540 1588 1591 1603 1641 1677 1685 1696 1700 1703 1761
1791 1798 1803 1804 1805 1813 1855 1862 1899 1953 1982 2022
2028 2036 2072 2086 2102 2111 2112 2135 2214 2215 2227 2254
2265 2295 2307 2310 2330 2360 2361 2362 2377 2386 2390 2431
2486 2507 2572 2602 2654 2675 2676 2689 2704 2752 2764 2769
2777 2778 2792 2799 2802 2857 2879 2903 2905 2911 2932 2959
2978 2993 3005 3021 3041 3053 3056 3058 3081 3094 3108 3121
3124 3159 3167 3179 3212 3240 3243 3299 3313 3320 3337 3344
3348 3365 3385 3390 3399 287 298 334 462 579 713 758
846 936 986 1076 1082 1169 1461 1535 1686 1766 1789 1796
1861 1911 1926 2079 2145 2213 2407 2434 2444 2470 2523 2822
2947 3043 3386];[12 33 34 36 68 78 99 107 113
117 120 127 136 143 178 186 195 282 307 319 346
374 385 422 433 434 452 454 508 522 534 538 545
566 583 639 678 688 693 707 730 731 738 744 745
770 819 832 837 840 842 856 864 871 896 907 919
959 985 986 993 997 1018 1033 1084 1086 1087 1093 1122
1134 1135 1215 1221 1230 1234 1271 1273 1306 1315 1328 1330
1338 1342 1351 1365 1397 1424 1430 1468 1475 1499 1608 1629
1632 1646 1663 1673 1674 1700 1704 1716 1728 1745 1750 1769
1805 1823 1828 1838 1839 1866 1870 1891 1900 1916 1932 1941
1944 1971 1983 2001 2003 2009 2014 2055 2089 2095 2100 2104
2118 2170 2180 2188 2200 2211 2223 2227 2242 2289 2312 2340
2351 2381 2388 2461 2472 2497 2522 2572 2632 2637 2642 2643
2657 2665 2666 2703 2719 2750 2762 2770 2786 2790 2803 2824
2838 2863 2876 2887 2943 2960 3036 3072 3073 3085 3089 3111
3113 3151 3163 3170 3260 3289 3316 3319 3386 3398 3404 3407
3422 35 48 93 265 295 316 379 384 527 530 948
1032 1141 1168 1276 1455 1490 1508 1551 1741 1810 1911 1980
2020 2121 2212 2287 2615 2617 2731 2953 3028 3152 3298];[54
59 67 82 115 116 209 248 250 285 331 333 348
349 355 362 381 396 417 424 502 517 533 539 547

```
549   568   592   601   615   629   643   652   691   702   715   753
806   834   846   862   870   892   902   913   925   940   964   974
979   989   997   999  1015  1018  1061  1107  1115  1123  1128  1147
1148  1151  1153  1181  1218  1227  1299  1314  1325  1340  1348  1362
1371  1402  1460  1463  1476  1499  1503  1509  1546  1607  1617  1642
1657  1686  1708  1711  1713  1733  1761  1802  1815  1817  1835  1875
1895  1903  1911  1920  1944  1951  1969  1975  2013  2018  2033  2055
2058  2077  2126  2153  2155  2158  2164  2172  2183  2205  2222  2247
2323  2362  2367  2376  2420  2426  2427  2428  2439  2450  2474  2483
2500  2512  2530  2543  2545  2577  2591  2633  2642  2644  2656  2676
2678  2706  2715  2731  2738  2764  2802  2834  2851  2869  2897  2917
2974  3017  3044  3048  3068  3108  3124  3127  3138  3141  3146  3148
3159  3200  3211  3213  3215  3225  3245  3301  3307  3312  3320  3328
361   453   515   706   721   807   893   984  1032  1042  1130  1157
1223  1342  1441  1702  1722  1724  1727  1765  1776  1855  1949  2118
2210  2269  2352  2473  2528  2843  3061  3070  3224  3298  3363];[5
47    62    95   130   170   183   201   215   217   259   272   274
318   330   331   356   367   373   393   413   414   418   431   434
445   499   500   562   588   591   612   652   719   749   759   760
772   773   784   794   820   844   855   871   896   899   933   934
953   998  1001  1087  1092  1113  1124  1140  1146  1154  1164  1172
1185  1208  1238  1262  1270  1317  1321  1362  1382  1402  1403  1428
1487  1516  1520  1525  1531  1570  1583  1599  1614  1630  1651  1655
1656  1683  1710  1715  1737  1745  1748  1857  1886  1901  1903  1937
1942  1954  2002  2017  2018  2038  2083  2086  2108  2117  2130  2141
2166  2173  2191  2192  2207  2211  2212  2218  2220  2226  2264  2274
2287  2288  2295  2313  2316  2317  2371  2447  2458  2481  2514  2562
2578  2581  2590  2596  2629  2650  2651    10    11    21   195   267
288   308   397   407   424   506   626   635   707   711   717   758
768   775   872   906   931  1138  1206  1269  1392  1420  1589  1695
1850  2217  2376  2576  2649  2672];};
% SCORES=[3 4];
% ns={[21    77    78    80    88   106   131   167   172   194   249
262   336   350   420   434   484   613   625   647   661   662   698
713   716   718   835   853   894   923   936   969   970   995  1018
1021  1103  1143  1153  1157  1177  1178  1218  1233  1237  1251  1307
1324  1329  1340  1355  1376  1389  1409  1422  1439  1453  1469  1490
1493  1520  1585  1599  1600  1609  1640  1683  1701  1722  1786  1800
1851  1926  1951  1986  2027  2037  2060  2066  2127  2139  2158  2173
2174  2210  2232  2239  2302  2313  2314  2358  2422  2435  2436  2468
2469  2511  2528  2530  2567  2568  2571  2575  2594    15    22    34
35    83    97   120   181   329   330   355   358   370   388   409
412   423   433   460   467   477   494   499   507   543   552   557
573   667   712   731   733   734   792   807   868   996  1059  1079
1094  1127  1166  1171  1200  1239  1311  1347  1358  1410  1426  1494
1555  1580  1595  1619  1628  1633  1645  1657  1673  1725  1738  1747
1748  1776  1826  1828  1874  1875  1876  1881  1887  1888  2000  2007
2014  2042  2082  2089  2090  2091  2176  2204  2231  2235  2296  2311
2337  2352  2365  2366  2372  2374  2413  2430  2432  2472  2531  2544
2593  2599];[48    63    81    82    89    97   140   174   187   231
232   237   244   298   323   324   374   439   514   524   535   593
607   615   616   628   672   725   753   789   796   802   822   866
868   872   876   913   949   991   992  1051  1052  1056  1128  1135
1136  1176  1185  1186  1203  1277  1278  1365  1472  1482  1501  1509
1516  1519  1537  1547  1610  1670  1685  1705  1734  1736  1792  1819
1838  1860  1863  1871  1872  1881  1937  1940  1980  1983  2005  2064
2079  2089  2097  2104  2125  2246  2270  2284  2337  2356  2383  2409
2455  2456  2490  2512  2522  2545  2546  2562  2625  2840  2846  2854
```

```
2877  2917  2949  2984  2990  2998  3013  3063  3086  3128  3142  3164
3180  3181  3183  3192  3193  3243  3250  3333  3368  3370  3405  3425
3429  3468  3525  3545  3600  3601  3627  3672  3766  3780  3798  3836
3838  3898  3953  3956  4038  4092  4099  4182  4186  4217  4300  4320
4325  4332  4438  4444  4484  4535  4583  4603  4631  4649  4662  4681
4682  4718  4792  4825  4914  4919  4921  4940  4991  4995  5035  5049
5070  5207  5232  5278  5333  5336  5350  5390  5392  5395  5398  5425
5483  5624  5654  5655  5690  5701  5721  5727  5736  5758  5789  5798
1    11    43    52   108   146   185   186   199   240   251   287   296
313   387   399   563   608   638   642   647   648   685   740   745
829   840   859   860  1004  1021  1038  1065  1076  1113  1139  1161
1205  1251  1274  1314  1321  1322  1360  1385  1386  1410  1414  1425
1491  1495  1504  1561  1565  1578  1601  1620  1622  1652  1674  1678
1726  1730  1737  1768  1788  1790  1809  1814  1817  1837  1869  1899
1919  1920  1929  1931  1942  1977  1991  2031  2037  2143  2244  2250
2258  2280  2330  2416  2439  2442  2466  2474  2554  2570  2590  2674
2735  2736  2754  2760  2769  2806  2845  2878  2884  2927  2937  2952
2953  2982  3002  3008  3014  3020  3083  3085  3099  3115  3143  3157
3161  3174  3178  3179  3184  3187  3200  3203  3213  3215  3224  3248
3295  3309  3336  3350  3351  3386  3402  3411  3437  3450  3465  3497
3513  3565  3626  3714  3740  3889  3907  3908  3919  3989  4009  4023
4069  4071  4091  4098  4102  4221  4258  4297  4331  4356  4404  4418
4423  4455  4494  4513  4533  4542  4556  4566  4598  4644  4656  4684
4760  4788  4811  4813  4844  4850  4875  4899  4901  4944  4947  4997
4998  4999  5000  5001  5072  5110  5112  5150  5157  5160  5176  5206
5217  5284  5308  5335  5431  5498  5504  5512  5527  5529  5563  5569
5628  5647  5653  5678  5688  5744  5748  5797];[40   118   141   254
261   285   292   298   352   363   395   471   489   542   544   567
675   729   756   789   792   800   873   876   888   902   904   905
915   922   986   989   995  1020  1029  1056  1129  1143  1171  1260
1280  1283  1289  1290  1300  1307  1318  1320  1349  1350  1426  1459
1474  1476  1492  1500  1524  1533  1693  1797  1824  1857  1864  1896
1945  1961  2055  2138  2249  2259  2295  2330  2373  2482  2572  2574
2576  2620  2793  2794  2815  2828  2829  2836  2889  2921  2929  2933
2958  2960  2976  3000  3004  3068  3082  3086  3101  3116  3172  3176
3199  3228  3248  3282  3311  3312  3324  3325  3334  3342  3384  3388
3481  3491  3506  3513  3532  3555  3560  3619  3626  3630    22    28
63    64    83   108   117   123   157   160   235   260   268   291
315   323   331   344   375   376   405   426   454   507   660   676
690   705   712   750   770   790   799   816   819   823   826   857
1024  1037  1046  1083  1113  1123  1148  1152  1164  1244  1273  1299
1331  1359  1390  1415  1482  1491  1517  1519  1534  1590  1599  1615
1616  1622  1627  1633  1640  1642  1670  1689  1691  1719  1776  1823
1842  1900  1967  1977  2031  2049  2051  2056  2124  2131  2178  2193
2196  2233  2251  2257  2269  2280  2288  2296  2311  2394  2397  2404
2428  2444  2457  2484  2534  2536  2567  2609  2640  2674  2697  2755
2768  2780  2786  2820  2866  2910  3020  3060  3069  3072  3093  3162
3230  3235  3239  3271  3349  3360  3386  3393  3408  3414  3419  3453
3457  3472  3494  3502  3547  3572  3598  3615  3620  3632];[31   105
157   158   159   160   162   163   205   254   278   290   460   487
529   646   647   658   685   689   691   694   738   763   783   800
823   855   861   884   917   926   942   952   953   973   989  1041
1048  1078  1081  1117  1172  1258  1285  1288  1310  1338  1345  1371
1383  1405  1439  1558  1574  1615  1626  1656  1667  1684  1691  1699
1704  1710  1714  1731  1738  1751  1766  1798  1847  1849  1888  1938
2059  2061  2086  2094  2096  2178  2219  2304  2312  2387  2388  2389
2460  2462  2509  2528  2574  2613  2650  2666  2702  2705  2738  2755
2806  2919    17    21    33    43    62    79   253   259   303   342
```

```
350   358   412   436   463   465   491   517   556   569   570   610
669   673   683   705   711   717   742   756   784   788   826   860
910   939   968   985  1020  1031  1032  1043  1077  1079  1100  1108
1157  1163  1192  1255  1265  1272  1279  1344  1348  1351  1461  1464
1492  1537  1557  1566  1568  1576  1581  1601  1612  1655  1657  1659
1668  1743  1750  1822  1830  1854  1878  1973  2041  2077  2097  2099
2163  2203  2216  2229  2233  2235  2292  2295  2298  2347  2357  2379
2430  2438  2463  2472  2496  2527  2547  2569  2584  2588  2629  2645
2791  2813  2830  2859  2870  2894  2911];[8    15    51    53    83
171   186   229   230   269   281   321   325   330   369   436   457
503   507   542   543   557   573   594   616   636   677   733   751
775   776   800   808   819   844   880   973   974  1041  1051  1052
1061  1065  1072  1073  1090  1102  1162  1172  1198  1230  1257  1302
1336  1341  1352  1368  1418  1422  1473  1493  1531  1539  1627  1706
1729  1732  1771  1781  1835  1837  1871  1887  1940  1956  1967  2001
2020  2088  2105  2122  2124  2193  2243  2255  2350  2442  2460  2476
2573  2587  2635  2675  2681  2687  2693  2804  2822  2825  2827  2848
2854  2921  2949  2955  2979  2997  2998  3029  3055  3067  3068  3075
3113  3125  3132  3151  3162  3166  3169  3186  3198  3230  3254  3263
3265  3325     6    52    76    81   105   118   124   165   194   201
224   341   366   417   440   489   519   523   525   541   544   570
577   581   583   609   644   665   675   676   727   783   814   824
839   879   911   929   983   994  1009  1012  1048  1055  1071  1079
1099  1104  1117  1138  1181  1231  1289  1301  1353  1415  1429  1497
1501  1513  1515  1518  1521  1644  1665  1678  1736  1743  1784  1799
1802  1827  1828  1861  1880  1901  1912  1939  2008  2012  2019  2035
2051  2067  2068  2081  2086  2101  2186  2190  2217  2219  2235  2241
2343  2348  2361  2364  2417  2468  2475  2516  2521  2529  2535  2544
2569  2619  2637  2658  2673  2697  2704  2755  2944  2948  2957  2965
2974  3005  3011  3048  3101  3184  3212  3255  3260  3283];[9    63
66   110   132   136   138   139   175   187   191   192   204   233
275   282   292   297   331   388   414   439   453   479   507   508
510   573   585   637   665   687   697   734   743   770   812   907
961   964   990  1010  1011  1015  1032  1033  1043  1079  1099  1117
1163  1208  1221  1237  1254  1261  1308  1370  1385  1397  1482  1501
1612  1632  1693  1733  1742  1754  1781  1799  1839  1867  1874  1878
1938  1956  1981  2027  2028  2079  2111  2141  2182  2193  2297  2450
2599  2613  2615  2664  2689  2730  2738  2763  2765  2769  2787  2804
2818  2821  2891  2898  2907  2913  2939  2955  2961  2976  2999  3014
3033  3041  3075  3081  3097  3113  3117  3160  3226  3240  3266  3302
3346  3353  3449  3507  3510  3516  3518  3554  3583  3606  3620  3705
3724    18    27    46    51    53    55    99   111   134   147   177
180   255   288   374   387   425   452   454   455   456   522   523
565   625   651   723   789   810   862   905   908   921   959   963
978   998  1003  1012  1044  1053  1055  1066  1067  1077  1087  1094
1123  1124  1161  1165  1184  1211  1213  1216  1220  1225  1238  1242
1248  1250  1253  1259  1278  1302  1382  1411  1430  1464  1468  1498
1521  1524  1541  1591  1691  1751  1797  1836  1853  1886  1894  1900
1905  1915  1918  1931  1940  1945  1946  1976  2003  2036  2126  2140
2143  2181  2243  2252  2257  2281  2294  2384  2421  2435  2436  2497
2533  2559  2582  2602  2605  2632  2669  2733  2791  2792  2811  2835
2850  2854  2857  2888  2908  2940  2952  2973  2978  2980  2982  3032
3040  3088  3093  3114  3121  3127  3137  3155  3166  3239  3348  3383
3391  3414  3440  3441  3450  3462  3521  3527  3675  3704  3714
3718];[28    41    71   102   105   106   138   143   159   163   192
195   223   224   249   250   252   256   258   260   279   283   292
300   370   412   420   423   498   516   521   535   548   562   575
580   586   600   609   623   714   759   777   796   817   826   834
```

852 888 891 894 968 970 986 996 1014 1039 1045 1069
1084 1105 1108 1124 1127 1147 1150 1152 1154 1198 1199 1251
1269 1316 1321 1376 1380 1393 1395 1438 1450 1500 1594 1606
1626 1628 1639 1643 1652 1668 1746 1756 1850 1851 1904 1905
1973 1990 2038 2245 2285 2310 2390 2404 2409 2420 2461 2470
2504 2512 2568 2590 2593 2613 2665 2683 2802 2824 2845 2923
2949 2960 2973 3079 3105 3126 3150 3213 3220 3244 3314 3320
3336 3359 3364 3416 3481 3499 3519 3539 3616 3653 3669 3682
3691 3709 3713 3742 3782 3848 3920 4007 4008 4013 4040 4044
4102 4129 4138 4181 4183 4187 4209 4212 4232 4234 4237 4244
4283 4408 4455 4612 18 40 78 96 147 196 203 218
227 248 274 340 400 425 545 570 583 588 593 663
664 728 754 779 788 843 861 966 967 969 982 1020
1031 1049 1125 1130 1178 1189 1248 1313 1323 1339 1364 1375
1381 1383 1414 1421 1472 1501 1520 1546 1555 1579 1592 1617
1655 1661 1670 1710 1747 1762 1799 1852 1854 1865 1875 1907
1968 1972 1994 2110 2118 2193 2287 2295 2325 2332 2347 2383
2392 2407 2441 2496 2503 2517 2519 2520 2535 2536 2544 2589
2615 2645 2646 2669 2723 2766 2774 2790 2830 2844 2865 2872
2886 2920 2953 2968 3033 3043 3147 3214 3249 3254 3256 3258
3274 3296 3321 3341 3346 3408 3413 3417 3447 3449 3464 3514
3515 3518 3533 3606 3618 3728 3733 3743 3748 3750 3767 3829
3863 3890 3914 3945 3957 3980 3994 4011 4023 4032 4054 4069
4070 4078 4108 4128 4146 4177 4188 4206 4227 4233 4242 4260
4288 4308 4320 4328 4346 4351 4415 4443 4457 4463 4466 4473
4528 4592];[56 93 105 115 138 204 215 216 219 266
293 316 398 409 436 438 445 515 531 539 561 565
579 641 702 731 734 787 820 850 854 903 904 925
952 977 1015 1065 1077 1078 1140 1184 1191 1226 1236 1247
1294 1297 1298 1307 1329 1330 1372 1374 1395 1405 1481 1499
1572 1602 1652 1698 1752 1779 1800 1925 1926 1984 2041 2043
2073 2079 2126 2145 2195 2263 2281 2282 2289 2290 2359 2380
2387 2412 2468 2520 2529 2552 2569 2583 2613 2643 2652 2670
2701 2717 2730 2737 2761 2856 2900 2955 3003 3021 3034 3063
3069 3080 3111 3233 3311 3315 3326 4 53 73 106 129
249 256 290 294 297 298 311 354 365 397 399 401
428 440 442 467 478 482 521 550 564 567 581 582
617 630 667 681 697 698 699 739 742 763 826 840
842 856 859 922 924 1064 1096 1107 1115 1128 1151 1152
1262 1288 1300 1301 1363 1408 1448 1449 1450 1473 1556 1608
1634 1684 1740 1832 1845 1862 1884 1918 1928 1935 1958 1964
1988 1991 1992 2003 2009 2013 2027 2040 2085 2120 2142 2146
2177 2220 2229 2230 2253 2255 2273 2304 2312 2323 2326 2350
2373 2375 2398 2406 2448 2554 2589 2590 2597 2609 2653 2662
2690 2727 2741 2780 2797 2821 2832 2878 2881 2895 2922 2924
2953 2963 3007 3060 3131 3161 3181 3190 3218 3222 3280 3299
3321 3337 3338 3357];[17 77 144 151 189 190 294 305
392 469 478 481 499 516 519 535 542 543 638 677
695 729 736 754 759 765 976 979 981 1031 1050 1051
1137 1146 1180 1247 1317 1325 1326 1328 1368 1374 1414 1421
1467 1478 1493 1496 1510 1515 1529 1533 1545 1579 1631 1696
1709 1710 1761 1785 1842 1850 1927 1934 1979 2003 2038 2069
2085 2128 2133 2155 2167 2217 2276 2298 2324 2357 2473 2503
2513 2564 2572 2600 2605 2639 2640 2663 2683 2695 2701 2726
2741 2744 2752 2755 2784 2798 2805 2875 2901 2906 2944 2970
2979 2981 3062 3108 3110 3144 3263 3283 3284 3286 3422 3423
3425 3451 3476 3518 3523 3528 8 60 81 85 114 128
157 162 165 197 211 219 257 298 314 320 332 355

418 459 465 480 491 501 502 553 568 573 575 608
627 662 667 683 721 746 762 805 808 890 901 905
906 912 918 937 975 1033 1061 1102 1129 1142 1218 1221
1223 1233 1306 1344 1387 1426 1477 1488 1492 1504 1523 1540
1584 1617 1632 1645 1698 1748 1751 1755 1779 1798 1811 1812
1852 1863 1865 1898 1907 1937 1945 1996 2001 2014 2033 2079
2096 2122 2186 2205 2230 2243 2246 2251 2273 2291 2294 2343
2361 2398 2400 2421 2466 2487 2506 2532 2536 2539 2563 2660
2704 2709 2718 2731 2732 2756 2809 2810 2818 2831 2843 2848
2858 2864 2884 2900 2909 2919 2943 2948 3026 3056 3094 3115
3116 3136 3147 3173 3233 3257 3285 3289 3338 3454 3457 3462
3497 3529];[17 35 41 56 57 69 80 290 292 313
359 372 373 419 441 475 495 532 576 633 658 666
718 747 750 760 788 826 858 864 874 894 900 905
988 1003 1052 1057 1112 1126 1208 1213 1235 1315 1319 1337
1349 1379 1399 1442 1506 1525 1538 1587 1652 1657 1666 1674
1695 1726 1739 1781 1875 1880 1914 1937 1939 1945 1962 2001
2090 2110 2152 2158 2173 2238 2249 2305 2419 2473 2513 2569
2576 2582 2586 2587 7 73 74 120 147 172 189 201
227 251 252 334 336 357 360 450 483 500 531 551
611 621 629 642 748 752 813 815 841 888 945 1023
1027 1031 1064 1065 1088 1098 1124 1132 1137 1139 1179 1186
1201 1202 1209 1252 1313 1317 1332 1336 1534 1541 1559 1584
1588 1609 1623 1677 1688 1711 1714 1722 1723 1744 1780 1813
1815 1816 1909 1961 1999 2003 2023 2066 2087 2089 2115 2129
2134 2157 2232 2270 2272 2413 2467 2497 2519 2537 2578];[15
62 128 153 179 211 216 218 220 225 231 235 263
397 430 576 583 622 647 649 653 667 697 700 716
726 731 733 740 741 744 790 808 928 938 949 966
1002 1024 1044 1125 1131 1141 1207 1218 1232 1242 1310 1337
1358 1361 1413 1424 1431 1432 1434 1501 1555 1603 1628 1723
1724 1725 1730 1738 1766 1794 1802 1808 1813 1842 1844 1933
1953 1954 1989 2055 2066 2084 2085 2134 2196 2199 2240 2344
2363 2392 2485 2486 2503 2514 2539 2570 2583 2589 2602 2606
2611 2659 2698 2731 2758 2765 2778 2801 2839 2863 2888 2902
2921 2940 2941 2949 3019 3057 3096 3105 3134 3197 3238 3272
3319 3342 3367 3399 3404 3411 3455 3514 3560 3596 3680 3687
3689 3700 3744 3782 3800 3863 4035 4090 4093 4106 4181 4182
4203 4207 4231 4267 4275 4389 4462 4463 4466 4475 4502 4503
4516 4563 4574 4581 4583 4584 4705 4879 74 122 127 155
222 236 244 245 246 261 283 336 365 388 436 451
452 457 458 464 496 521 540 555 645 652 725 734
750 769 775 863 910 935 1016 1074 1086 1111 1160 1174
1211 1222 1250 1303 1336 1418 1433 1435 1446 1498 1524 1573
1596 1614 1620 1624 1625 1631 1641 1653 1701 1768 1836 1847
1848 1866 1901 2011 2036 2042 2051 2064 2081 2100 2138 2153
2202 2214 2218 2242 2321 2332 2354 2390 2407 2456 2469 2502
2507 2519 2521 2525 2533 2544 2545 2553 2555 2571 2576 2581
2597 2599 2612 2620 2627 2697 2761 2810 2811 2847 2867 2924
2954 2996 2997 3036 3059 3124 3133 3143 3311 3336 3343 3379
3385 3443 3448 3449 3467 3486 3508 3537 3565 3584 3659 3663
3732 3738 3749 3780 3791 3796 3808 3812 3817 3853 3900 3903
3908 3939 3961 3985 3996 4020 4034 4047 4066 4084 4094 4103
4112 4119 4154 4229 4337 4346 4348 4368 4439 4488 4515 4573
4575 4579 4604 4624 4641 4653 4654 4662 4709 4739 4766 4802
4835];[8 16 22 34 38 54 97 128 164 168 184
220 269 280 358 366 401 415 438 440 509 545 550
588 656 743 761 784 787 805 851 889 910 937 955

1049 1061 1079 1118 1153 1196 1237 1282 1290 1303 1305 1345
1357 1372 1388 1402 1408 1431 1493 1518 1540 1593 1626 1632
1660 1696 1699 1725 1728 1771 1772 1817 1820 1833 1834 1850
1858 1866 1907 1915 1962 1997 2009 2051 2067 2068 2076 2110
2142 2176 2202 2203 2340 2365 2399 2428 2432 2440 2484 2565
32 37 72 93 113 137 150 242 261 270 278 294
331 339 386 451 486 529 535 536 546 547 558 575
589 676 747 801 815 915 930 933 950 996 997 1002
1018 1051 1083 1096 1168 1205 1238 1246 1263 1274 1319 1344
1348 1413 1443 1444 1445 1476 1488 1533 1538 1542 1655 1661
1724 1742 1750 1777 1783 1823 1851 1911 1959 1980 2005 2007
2030 2044 2139 2154 2175 2187 2197 2223 2250 2264 2267 2294
2308 2320 2383 2413 2416 2468 2503 2573 2586];[40 41 54
85 151 153 173 182 195 258 266 272 331 341 355
383 431 463 470 477 478 522 552 556 560 597 607
627 634 654 667 726 745 754 777 784 816 834 875
900 908 912 923 924 936 940 945 1018 1022 1024 1047
1048 1051 1098 1154 1172 1176 1199 1238 1247 1333 1334 1346
1356 1404 1413 1483 1526 1599 1646 1742 1752 1774 1775 1789
1798 1834 1840 1854 1859 1885 2017 2032 2065 2107 2134 2206
2220 2269 2287 2334 2335 2393 2403 2424 2430 2436 2439 2447
2534 2539 2591 2596 2640 2667 2684 14 39 53 82 87
137 150 160 180 219 291 307 312 326 342 401 405
407 413 414 419 474 488 534 541 543 602 617 683
691 713 785 792 833 842 857 887 902 920 958 991
1003 1027 1053 1073 1074 1075 1106 1143 1155 1174 1202 1235
1237 1257 1258 1298 1301 1342 1353 1421 1450 1475 1506 1534
1603 1608 1633 1650 1659 1715 1717 1720 1751 1759 1788 1796
1836 1847 1886 1938 1944 1965 1989 2080 2110 2147 2164 2182
2229 2235 2301 2414 2421 2422 2434 2435 2440 2451 2475 2478
2483 2538 2582 2592 2691];[179 243 253 309 327 346 404
529 530 584 651 706 713 722 756 761 777 803 831
832 876 925 967 982 993 1100 1147 1156 1201 1221 1248
1297 1310 1312 1350 1397 1398 1445 1466 1505 1549 1551 1558
1583 1592 1603 1634 1672 1673 1689 1715 1717 1725 1765 1803
1817 1882 1889 1975 1993 2068 2093 2101 2131 2134 2164 2180
2193 2196 2211 2239 2254 2270 2277 2299 2341 2364 2381 2415
2424 2451 2472 2496 1 54 60 95 109 120 161 166
197 254 272 302 406 461 496 531 594 668 669 715
718 741 880 909 921 934 939 941 942 956 957 972
1008 1032 1041 1056 1061 1075 1091 1155 1174 1203 1283 1298
1301 1303 1306 1318 1338 1348 1427 1428 1429 1446 1450 1480
1481 1552 1572 1575 1605 1657 1660 1680 1716 1778 1779 1786
1800 1822 1823 1868 1869 1887 1947 1983 1994 2033 2189 2200
2203 2229 2351 2394 2403 2416 2434 2439 2446 2457 2483 2523
2524 2526];[36 42 77 179 196 217 320 341 428 540
543 544 559 562 575 578 581 599 617 660 683 707
754 755 795 808 820 825 864 872 895 905 938 948
993 1039 1066 1115 1228 1297 1389 1394 1444 1487 1502 1578
1591 1613 1653 1660 1667 1678 1679 1738 1787 1791 1801 1848
1875 1883 1909 55 62 75 78 92 103 160 165 191
197 210 218 220 232 265 334 379 385 392 475 479
539 594 662 685 694 697 724 766 791 803 816 875
888 965 968 1031 1033 1035 1037 1086 1136 1141 1201 1252
1287 1290 1293 1303 1321 1392 1430 1433 1447 1478 1506 1521
1524 1545 1609 1615 1697 1703 1704 1715 1716 1736 1737 1758
1771 1772 1775 1781 1806 1845 1851 1867 1884 1906 1913
1917];[28 71 78 85 91 105 106 180 319 322 330

```
333   374   406   414   429   434   438   468   496   502   518   540
596   619   643   678   770   778   794   814   815   925   927   942
967  1149  1152  1191  1300  1304  1312  1345  1385  1400  1482  1498
1543  1640  1697  1704  1735  1747  1758  1776  1805  1849  1877  1894
1896  1957  2032  2111  2133  2180  2181  2203  2208  2224  2248  2310
2364  2383  2384  2387  2430  2444  2537  2545  2573  2574  2625  2632
2645  2654  2672  2673  2727  2799  2812  2847  2881  2892  2925  2949
2956  2957  3059  3068  3130  3135  3148  3182  3188  3213  3231  3302
3332  3364  3365  3367  3381  3392  3395  3404  3432  3454  3471  3503
3518  3535  3536  3550  3558  3577  3617  3637  3668  3673  3678  3694
3695  3699  3713  3741  3750    74   179   204   253   281   284   285
310   364   392   413   432   433   457   471   495   504   537   538
558   584   589   636   641   700   703   706   771   830   834   886
899   941   947   958   991   997  1061  1062  1094  1128  1197  1202
1209  1270  1296  1301  1344  1388  1391  1454  1456  1485  1503  1561
1581  1615  1644  1744  1760  1793  1799  1816  1865  1880  1889  1919
1925  1930  1940  1945  1949  1961  1972  2003  2009  2011  2055  2090
2091  2174  2184  2245  2255  2347  2370  2548  2575  2586  2618  2631
2647  2651  2697  2698  2716  2772  2805  2816  2854  2860  2872  2878
2887  2899  2906  2913  2926  2932  2944  2945  2952  2973  2981  3021
3066  3088  3112  3124  3136  3140  3214  3226  3237  3292  3306  3311
3322  3329  3337  3370  3372  3408  3411  3422  3451  3478  3496  3515
3597  3633  3690  3738];[83   105   120   132   174   192   208   221
270   284   312   319   339   424   431   499   502   519   568   569
592   611   671   682   745   747   815   835   867   886   936   961
1000  1013  1060  1067  1089  1106  1129  1169  1239  1261  1276  1277
1281  1289  1295  1370  1406  1420  1429  1445  1462  1489  1490  1509
1517  1527  1549  1565  1596  1634  1636  1663  1678  1706  1711  1751
1790  1799  1807  1844  1857  1882  1890  1902  1929  2001  2141  2186
2241  2257  2262  2305  2308  2338  2386  2408  2449  2450  2457  2458
2461  2464  2568  2570  2572  2583    34    62    64    89    98   102
114   129   143   181   197   281   303   304   330   337   342   345
351   355   356   357   396   402   539   545   565   584   599   621
629   638   674   757   758   788   792   813   816   840   844   851
896   910   925   965   981   997   999  1049  1139  1164  1214  1278
1314  1342  1363  1413  1442  1538  1541  1561  1601  1614  1650  1677
1722  1740  1814  1883  1885  1891  1936  1970  1982  1999  2004  2012
2082  2102  2123  2124  2131  2224  2265  2292  2303  2312  2326  2337
2416  2428  2500  2531  2534  2538  2563  2579];[50    71   114   156
163   168   199   227   282   298   308   366   421   451   452   455
499   504   532   542   568   572   577   590   605   609   622   739
754   810   822   852   862   872   935  1032  1082  1090  1127  1159
1162  1172  1183  1201  1250  1255  1257  1298  1322  1363  1384  1402
1420  1459  1468  1510  1536  1540  1578  1588  1699  1700  1703  1761
1774  1781  1791  1796  1803  1822  1883  1953  2022  2057  2135  2192
2214  2234  2260  2270  2295  2365  2374  2377  2422  2434  2441  2517
2606  2608  2648  2689  2769  2778  2792  2799  2843  2865  2880  2886
2905  2910  2929  3007  3017  3037  3047  3058  3081  3091  3108  3116
3121  3128  3142  3178  3179  3181  3191  3194  3195  3299  3314  3348
3365  3390  3391     6   213   236   287   314   334   376   396   419
458   508   579   581   598   648   680   683   694   700   703   726
758   842   846   850   874   875   886   936   973   986   995  1028
1062  1073  1076  1128  1138  1187  1224  1272  1286  1301  1319  1330
1352  1362  1396  1418  1437  1461  1535  1537  1591  1603  1635  1641
1685  1686  1744  1789  1798  1804  1805  1855  1861  1862  1911  1926
1936  1958  1982  2072  2079  2086  2111  2112  2145  2213  2215  2266
2307  2310  2330  2360  2362  2386  2407  2431  2470  2486  2523  2602
2654  2675  2676  2704  2752  2764  2802  2857  2879  2903  2911  2959
```

2962 2978 2993 3024 3027 3041 3043 3053 3056 3094 3124 3159
3161 3167 3240 3243 3313 3320 3337 3344 3380 3398];[27 34
35 76 78 107 113 117 135 141 143 178 179 186
195 217 256 264 292 417 418 430 451 530 545 565
574 608 659 668 688 694 697 698 728 770 819 880
885 930 985 994 1034 1037 1061 1093 1133 1135 1142 1145
1213 1275 1351 1357 1423 1468 1510 1629 1673 1674 1724 1728
1739 1824 1891 1911 1919 1971 2009 2020 2083 2126 2156 2159
2170 2194 2200 2206 2212 2311 2346 2351 2353 2381 2386 2411
2491 2521 2522 2572 2605 2640 2668 2762 2770 2803 2828 2876
2900 2920 2936 2982 3018 3024 3036 3070 3072 3085 3111 3151
3163 3249 3266 3289 3306 3316 3347 3402 3422 12 16 32
33 36 48 68 93 99 120 136 265 282 307 316
319 346 374 379 385 422 434 452 487 508 522 527
534 538 626 687 693 707 730 738 744 745 832 856
864 871 896 903 907 919 948 959 977 983 993 1032
1033 1055 1084 1086 1134 1141 1151 1157 1168 1215 1221 1230
1234 1271 1273 1276 1315 1328 1330 1342 1397 1415 1416 1425
1430 1447 1455 1475 1508 1522 1608 1700 1716 1741 1750 1769
1810 1813 1823 1828 1839 1866 1900 1916 1932 1941 1944 1980
1983 2001 2003 2014 2035 2055 2089 2095 2104 2121 2180 2188
2189 2211 2242 2287 2289 2312 2340 2367 2472 2604 2615 2617
2637 2642 2643 2657 2665 2666 2703 2719 2731 2735 2750 2786
2790 2824 2863 2937 2952 2960 3073 3113 3141 3152 3298 3386
3398 3404];[32 141 169 181 237 285 294 355 361 396
405 414 421 442 452 547 574 592 609 652 693 753
846 870 892 946 1018 1019 1058 1069 1077 1128 1129 1130
1148 1199 1201 1204 1250 1269 1275 1371 1460 1472 1489 1503
1575 1596 1630 1647 1672 1721 1727 1759 1800 1875 1911 1920
1942 1958 1969 1979 1985 2031 2055 2056 2120 2124 2144 2159
2172 2237 2247 2251 2269 2473 2474 2512 2528 2577 2700 2718
2731 2803 2846 2851 2901 2918 2927 3022 3047 3052 3062 3093
3102 3138 3200 3213 3229 3268 3316 3320 3361 3363 54 59
82 115 116 209 223 265 331 333 348 362 424 453
502 503 517 537 539 549 551 567 568 629 660 702
706 715 721 806 807 834 862 893 902 913 914 925
940 964 974 984 989 997 999 1032 1061 1107 1115 1126
1147 1151 1153 1157 1223 1227 1325 1340 1342 1346 1348 1362
1402 1441 1463 1476 1509 1513 1546 1585 1642 1702 1708 1711
1713 1722 1724 1730 1733 1765 1776 1817 1855 1903 1951 2013
2018 2033 2077 2099 2118 2126 2138 2140 2155 2158 2164 2183
2186 2205 2210 2222 2323 2352 2362 2367 2376 2420 2425 2426
2427 2428 2450 2483 2530 2543 2554 2591 2633 2644 2656 2678
2706 2715 2728 2738 2764 2834 2843 2869 2974 2998 3017 3044
3048 3058 3061 3070 3108 3127 3141 3142 3146 3148 3211 3215
3225 3245 3298 3301 3307 3312];[5 95 135 145 184 201
217 278 288 358 373 393 413 414 500 577 591 606
625 652 759 773 785 843 844 846 887 896 917 933
934 998 1001 1021 1027 1067 1086 1092 1154 1184 1202 1208
1262 1362 1374 1382 1401 1409 1458 1487 1516 1531 1536 1560
1561 1565 1566 1567 1570 1584 1591 1610 1612 1614 1656 1679
1705 1748 1871 2013 2023 2038 2083 2086 2102 2127 2130 2141
2142 2173 2192 2211 2218 2221 2264 2274 2283 2286 2309 2371
2465 2528 2536 2578 2590 2596 2600 2625 2629 2647 2661 2666
10 11 21 47 62 170 183 195 215 267 274 308
318 330 365 367 397 407 418 424 434 445 499 506
507 562 588 609 626 635 707 711 717 719 749 758
760 768 775 802 820 835 855 861 866 871 899 906

```
931   953  1087  1113  1124  1146  1157  1206  1269  1317  1320  1321
1365  1384  1392  1402  1420  1480  1525  1583  1589  1599  1630  1651
1683  1695  1745  1757  1850  1856  1857  1903  1911  1954  2002  2017
2035  2088  2108  2117  2156  2191  2207  2212  2226  2287  2288  2313
2316  2367  2376  2447  2458  2514  2562  2576  2649  2650  2651];};
% SCORES=[1 2];
% ns={[21    22    34    62    78   106   140   167   172   194   198
249   262   330   336   350   354   355   390   420   434   484   613
625   647   661   662   670   676   707   713   716   718   734   969
970   983  1039  1044  1153  1166  1177  1217  1218  1220  1233  1249
1263  1269  1307  1329  1340  1347  1355  1366  1409  1422  1469  1493
1494  1520  1599  1600  1628  1640  1683  1701  1722  1724  1730  1738
1786  1799  1803  1851  1881  1926  1951  1986  2007  2037  2065  2066
2090  2096  2127  2173  2210  2232  2304  2313  2314  2342  2358  2403
2422  2432  2436  2437  2469  2511  2528  2530  2575  2594    35    77
181   342   410   432   555   675   698   835   894   923  1057  1075
1103  1117  1143  1157  1311  1389  1426  1609  1792  1828  2014  2027
2060  2139  2302  2365  2529  2571];[18    48    63    81    82    97
134   140   168   174   187   204   231   232   237   240   244   290
298   323   324   366   371   374   422   438   439   524   535   593
607   608   615   616   628   672   725   753   822   826   860   866
868   872   876   896   991  1128  1136  1227  1251  1277  1278  1300
1472  1495  1501  1504  1515  1519  1520  1534  1542  1547  1570  1587
1610  1617  1620  1670  1674  1678  1705  1734  1736  1745  1792  1819
1838  1860  1881  1899  1900  1937  1940  1979  2024  2079  2097  2104
2112  2143  2194  2221  2246  2272  2284  2356  2383  2409  2435  2446
2455  2545  2551  2562  2570  2625  2684  2700  2744  2877  2905  2917
2945  2949  2982  2984  2990  2998  3008  3013  3017  3018  3063  3067
3086  3115  3123  3128  3140  3141  3142  3147  3157  3163  3164  3181
3183  3192  3243  3250  3267  3303  3312  3333  3370  3386  3452  3468
3471  3525  3545  3798  3838  3859  3898  3919  3956  4038  4099  4186
4217  4297  4320  4325  4332  4418  4438  4443  4444  4445  4451  4515
4535  4563  4576  4583  4596  4606  4662  4681  4718  4724  4742  4753
4760  4763  4792  4825  4919  4991  5035  5070  5157  5207  5232  5254
5293  5336  5390  5395  5406  5419  5425  5655  5701  5731  5735  5789
5798    89   287   513   514   730   789   796   802   913   992  1052
1056  1057  1186  1205  1211  1314  1365  1482  1622  1643  1652  1685
1817  1863  1866  1871  1872  1942  1983  2089  2102  2141  2490  2522
2546  2840  2846  2854  2878  3179  3180  3193  3203  3437  3513  3565
3600  3601  3719  3836  3878  4092  4191  4221  4285  4300  4404  4484
4514  4603  4631  4813  4841  5018  5049  5068  5195  5350  5359  5387
5398  5477  5690  5721  5736];[28    36    40    86   106   118   202
254   259   285   292   322   363   376   471   489   493   494   544
567   573   606   675   729   756   776   792   799   800   816   826
876   888   904   905   922   937   966   975   982   986   995  1019
1020  1029  1042  1079  1129  1135  1143  1246  1260  1261  1280  1289
1290  1300  1307  1318  1320  1349  1350  1426  1448  1474  1476  1492
1533  1572  1580  1609  1670  1678  1693  1776  1797  1823  1864  1896
1897  1907  1961  2031  2131  2202  2249  2256  2288  2295  2330  2521
2533  2572  2574  2613  2620  2640  2674  2749  2764  2780  2815  2828
2829  2836  2859  2921  2929  2958  2960  3000  3082  3086  3101  3116
3199  3215  3228  3248  3252  3311  3312  3324  3325  3384  3409  3414
3481  3506  3532  3579  3597  3619  3626  3630  3633    59   123   261
298   352   664   750   802   823   859   902   989  1056  1148  1152
1203  1283  1299  1455  1491  1500  1524  1819  1824  1842  1857  1945
2042  2056  2178  2193  2209  2233  2259  2312  2703  2735  2884  3172
3176  3216  3222  3278  3282  3318  3335  3360  3388  3394  3453  3491
3509  3513  3555  3635];[11    21    32    76   105   158   160   218
```

274 328 421 487 606 610 632 647 658 669 685 689
694 706 717 731 738 763 783 823 855 861 884 916
917 926 939 973 989 1041 1048 1117 1140 1172 1186 1194
1260 1288 1310 1338 1344 1457 1494 1540 1558 1574 1610 1615
1626 1651 1656 1684 1691 1704 1710 1714 1731 1751 1766 1798
1847 1849 1888 1949 2059 2061 2086 2099 2178 2208 2216 2219
2227 2240 2246 2345 2387 2389 2391 2430 2460 2479 2509 2528
2554 2574 2588 2613 2666 2702 2738 2755 2806 2919 43 51
216 261 278 290 646 734 788 932 942 952 953 958
1011 1080 1105 1255 1258 1351 1437 1451 1481 1492 1655 1825
2010 2096 2097 2126 2131 2248 2251 2304 2388 2462 2645
2650];[8 15 51 53 68 141 144 171 186 269 289
300 321 325 330 369 427 436 457 507 541 542 543
557 574 594 616 677 733 751 776 789 791 808 838
844 879 973 974 1041 1051 1052 1065 1066 1072 1073 1090
1097 1102 1106 1117 1198 1214 1230 1274 1302 1331 1336 1353
1364 1368 1380 1412 1473 1493 1539 1569 1627 1706 1727 1729
1736 1743 1771 1781 1835 1837 1855 1871 1887 1940 1961 1967
2020 2068 2122 2124 2193 2243 2255 2336 2343 2442 2460 2468
2476 2503 2573 2587 2607 2619 2635 2636 2687 2695 2804 2821
2822 2827 2848 2854 2883 2921 2943 2949 2955 2965 2998 3055
3067 3068 3073 3113 3132 3166 3198 3265 3289 3325 338 440
609 636 674 676 727 775 792 800 835 1162 1177 1231
1278 1352 1422 1515 1644 1748 1773 1811 1901 1956 2012 2035
2051 2094 2350 2569 2622 2693 2829 2862 2979 2997 3151 3199
3254 3260 3263 3339];[9 14 27 63 110 132 134 136
138 139 156 175 177 180 187 188 191 204 233 238
275 279 282 292 336 365 388 414 439 453 456 479
480 507 508 510 564 565 585 637 687 743 752 770
812 957 961 964 990 1011 1025 1032 1033 1043 1044 1067
1079 1099 1111 1117 1125 1163 1208 1230 1254 1308 1370 1385
1397 1431 1468 1482 1501 1675 1799 1836 1867 1874 1878 1886
1929 1937 1938 1965 1981 2011 2027 2028 2053 2065 2079 2095
2126 2141 2182 2193 2297 2311 2450 2613 2615 2638 2641 2730
2757 2787 2811 2850 2853 2854 2887 2898 2905 2907 2908 2939
2951 2955 2961 2964 2979 2999 3014 3019 3032 3033 3081 3097
3173 3216 3219 3226 3240 3264 3302 3346 3387 3447 3516 3529
3583 3606 3705 3724 66 78 107 111 193 347 573 723
789 908 921 956 962 984 1010 1087 1162 1205 1211 1221
1237 1238 1250 1430 1541 1632 1754 1839 1883 2015 2111 2142
2169 2190 2363 2444 2495 2599 2611 2632 2876 2916 3006 3041
3071 3334 3353 3382 3450 3620];[28 40 41 102 106 138
159 192 195 196 217 218 223 224 249 250 252 256
260 263 279 283 291 292 300 370 376 423 498 516
521 535 548 562 576 593 600 605 609 623 655 714
759 777 791 792 796 803 817 826 837 853 856 870
891 894 898 956 986 996 1031 1039 1083 1084 1088 1108
1124 1147 1150 1152 1154 1199 1248 1251 1252 1293 1313 1376
1380 1383 1393 1395 1438 1450 1500 1501 1514 1626 1628 1639
1670 1746 1756 1904 1905 1973 2038 2070 2390 2461 2504 2590
2593 2613 2665 2683 2802 2845 2960 2997 3126 3150 3220 3281
3314 3318 3320 3336 3364 3413 3449 3499 3533 3616 3669 3691
3713 3742 3782 3848 3914 3920 3956 4013 4040 4044 4102 4129
4138 4177 4183 4234 4237 4260 4408 4443 4455 4515 4518 64
151 186 203 287 303 372 420 545 575 586 614 852
970 1033 1105 1141 1160 1177 1198 1239 1269 1279 1316 1381
1495 1520 1592 1643 1701 1710 1925 1990 2285 2470 2508 2512
2568 2790 2844 3011 3079 3214 3247 3274 3416 3519 3539 3682

3709 3743 3760 3767 4007 4032 4078 4171 4187 4209 4232 4233
4240 4244 4328 4346 4466 4541];[56 92 93 105 115 116
129 138 204 215 293 311 312 315 316 409 445 477
515 531 539 647 658 671 702 731 734 787 820 854
903 904 922 952 965 977 1015 1064 1077 1078 1080 1114
1118 1128 1140 1184 1191 1208 1226 1236 1262 1280 1294 1297
1298 1307 1329 1330 1371 1372 1374 1437 1462 1481 1499 1535
1602 1652 1684 1686 1775 1779 1800 1844 1926 1984 2041 2043
2073 2079 2125 2142 2253 2273 2282 2290 2300 2520 2552 2583
2607 2652 2670 2687 2692 2701 2730 2797 2881 2918 2924 2955
2977 3021 3022 3034 3039 3048 3060 3063 3069 3111 3121 3161
3280 3311 3326 3337 48 137 398 436 438 442 561 579
582 630 651 681 763 1203 1247 1291 1459 1538 1572 1689
1752 1760 1884 1925 2003 2009 2099 2126 2195 2255 2263 2375
2387 2468 2597 2613 2643 2704 2727 2737 2761 2853 3003 3166
3190 3218 3299 3314];[12 27 81 101 123 144 151 166
189 190 211 264 305 392 395 459 469 478 481 499
516 533 535 542 543 568 606 627 638 677 688 695
736 754 759 765 810 894 900 906 935 937 943 979
981 992 1005 1013 1031 1050 1051 1119 1137 1146 1247 1262
1263 1297 1325 1326 1328 1364 1368 1374 1414 1421 1478 1493
1496 1510 1524 1529 1579 1645 1709 1710 1776 1785 1798 1807
1842 1852 1863 1893 1927 1934 1979 2003 2030 2033 2048 2128
2133 2167 2244 2247 2276 2298 2357 2362 2487 2605 2639 2645
2660 2663 2696 2703 2714 2717 2726 2731 2741 2744 2755 2784
2804 2805 2875 2906 2944 2970 2981 2989 3006 3062 3144 3183
3208 3248 3257 3263 3281 3283 3284 3288 3423 3476 3518 60
136 165 257 300 347 383 575 603 696 805 893 976
1105 1147 1180 1223 1272 1540 1561 1567 1580 1631 1637 1696
1751 1761 1850 1907 2001 2069 2421 2503 2539 2572 2600 2628
2640 2683 2699 2704 2798 2810 2901 2919 2943 2979 3026 3045
3051 3098 3110 3185 3286 3422 3454 3523];[17 19 35 39
41 56 57 73 258 290 305 359 372 441 449 450
483 492 498 531 552 576 597 611 621 631 633 658
666 689 747 750 760 820 826 841 844 866 894 904
957 1026 1038 1057 1101 1112 1126 1137 1139 1163 1171 1208
1209 1235 1276 1315 1319 1337 1349 1379 1399 1406 1469 1506
1515 1587 1589 1623 1649 1652 1657 1674 1698 1701 1739 1799
1849 1880 1900 1905 1939 1961 1968 2003 2049 2062 2152 2157
2158 2211 2238 2257 2324 2355 2356 2395 2448 2473 2513 2529
2569 2576 2586 2587 2596 58 69 80 147 201 252 292
373 378 419 462 475 518 532 604 663 718 748 858
905 942 988 1003 1052 1054 1213 1236 1341 1363 1609 1666
1726 1813 1875 1999 2001 2090 2419 2519 2537 2582];[15 78
155 211 216 217 218 231 235 242 262 263 324 354
395 404 435 537 583 587 622 649 653 667 700 716
726 731 733 740 741 744 746 771 777 790 808 857
910 916 928 949 973 1002 1005 1086 1131 1142 1211 1232
1242 1337 1361 1413 1424 1431 1432 1434 1435 1446 1501 1514
1532 1555 1704 1723 1724 1737 1738 1766 1794 1808 1810 1813
1841 1842 1874 1884 1903 1933 1954 1989 2000 2003 2055 2064
2084 2085 2100 2130 2196 2199 2234 2344 2392 2450 2459 2479
2480 2488 2503 2508 2544 2571 2580 2583 2586 2599 2606 2627
2698 2754 2765 2778 2801 2802 2850 2863 2888 2929 2940 2996
3051 3057 3070 3096 3134 3182 3197 3199 3238 3272 3367 3368
3404 3411 3416 3441 3449 3514 3560 3571 3572 3596 3678 3680
3687 3717 3744 3800 3813 3863 3996 3998 4008 4035 4080 4106
4181 4182 4183 4203 4208 4222 4267 4275 4310 4326 4389 4448

4466 4475 4502 4503 4515 4516 4540 4563 4573 4574 4575 4579
4583 4584 4676 4709 4760 4791 4867 62 122 128 206 220
224 415 521 586 646 647 697 715 821 863 938 977
1024 1074 1141 1160 1198 1271 1287 1291 1358 1378 1614 1628
1641 1736 1768 1825 1901 1912 1953 2013 2042 2066 2121 2122
2160 2240 2363 2442 2539 2570 2611 2612 2659 2742 2758 2949
2956 3399 3537 3667 3679 3689 3716 3985 4034 4207 4221 4229
4231 4368 4433 4581 4641 4705 4784 4814 4840];[6 8 16
38 54 72 128 164 168 184 204 220 269 358 366
401 509 588 656 718 741 743 784 805 869 889 910
914 955 1118 1137 1153 1167 1196 1213 1215 1282 1303 1305
1343 1345 1388 1402 1404 1408 1431 1445 1476 1493 1540 1593
1626 1632 1660 1661 1696 1699 1725 1728 1771 1783 1818 1839
1850 1858 1866 1902 1915 1959 1962 1986 1987 1997 2009 2067
2076 2094 2203 2204 2284 2294 2340 2365 2402 2432 2440 2484
2532 22 34 97 261 288 371 399 451 470 486 667
671 727 742 933 937 996 1002 1028 1168 1237 1290 1352
1357 1372 1518 1700 1777 1817 1820 1834 1907 1980 2066 2117
2142 2223 2249 2267 2565];[14 36 40 41 43 54 85
115 182 195 247 258 331 341 355 383 421 431 474
477 478 522 544 552 554 556 560 587 597 607 629
654 680 714 716 726 754 777 784 785 834 851 912
924 936 940 1015 1027 1047 1048 1051 1082 1098 1176 1186
1238 1298 1311 1333 1356 1357 1413 1421 1483 1526 1529 1646
1720 1751 1752 1758 1759 1774 1775 1789 1804 1824 1834 1836
1840 1854 1859 1885 1989 2017 2039 2055 2129 2134 2220 2287
2331 2334 2335 2336 2367 2393 2403 2424 2427 2435 2436 2439
2447 2539 2596 2667 357 381 419 458 463 592 715 816
875 900 908 923 945 1018 1143 1172 1202 1216 1237 1275
1301 1305 1334 1346 1506 1603 1607 1796 1798 1913 2032 2070
2534 2558 2591 2665];[54 154 179 200 220 221 243 253
309 327 351 359 404 449 530 638 706 713 714 722
740 756 761 762 777 807 831 832 925 929 988 993
1010 1100 1156 1164 1201 1221 1248 1265 1303 1310 1312 1318
1348 1397 1428 1445 1466 1470 1551 1575 1583 1592 1603 1634
1657 1664 1715 1716 1722 1786 1803 1817 1820 1842 1889 1907
1957 1993 1994 2068 2093 2101 2108 2131 2134 2164 2180 2196
2198 2234 2239 2245 2254 2270 2277 2299 2300 2331 2348 2364
2381 2397 2415 2424 2430 2439 2446 2451 2472 2496 120 346
352 443 497 531 624 636 651 715 982 1056 1099 1147
1174 1222 1297 1306 1398 1505 1521 1558 1605 1680 1689 1765
2064 2116 2253 2341 2394];[29 36 42 60 74 89 179
191 196 217 218 220 320 341 428 498 533 540 562
578 581 599 617 660 662 683 697 707 754 795 808
816 820 825 864 872 874 895 905 907 937 938 946
948 993 1039 1086 1136 1196 1228 1293 1297 1389 1400 1444
1487 1502 1521 1556 1660 1671 1678 1679 1704 1738 1791 1801
1839 1883 1909 62 90 92 103 282 575 584 649 755
888 929 990 1033 1290 1311 1591 1716 1787 1812 1845 1884];[1
28 71 74 85 106 179 180 185 204 319 322 330
374 406 433 438 457 493 495 496 502 518 537 643
651 722 770 778 794 815 925 958 967 1014 1084 1093
1149 1191 1192 1304 1312 1422 1482 1498 1543 1640 1697 1704
1747 1750 1877 1894 1896 1925 1948 2083 2141 2181 2191 2208
2248 2310 2364 2384 2387 2430 2444 2466 2482 2537 2545 2573
2574 2625 2631 2632 2654 2673 2727 2750 2799 2805 2812 2873
2878 2881 2925 2926 2944 2956 3067 3068 3182 3213 3216 3237
3302 3332 3365 3380 3381 3395 3404 3415 3416 3432 3536 3637

```
3668  3673  3694  3699  3713     6    36    83   225   275   281   321
333   364   429   434   540   596   657   700   997  1029  1051  1112
1152  1182  1197  1301  1345  1405  1735  1758  1776  1805  1880  1908
1957  2032  2111  2180  2203  2321  2383  2490  2526  2549  2645  2697
2811  2892  2949  3059  3188  3202  3310  3326  3364  3367  3370  3392
3454  3486  3550  3558  3611  3617];[83   192   197   208   221   263
270   298   312   313   357   422   424   438   499   502   519   568
569   583   592   630   671   698   714   747   757   835   858   867
886   888   929   936   961  1000  1013  1067  1089  1129  1163  1169
1202  1261  1277  1295  1301  1317  1366  1367  1370  1406  1420  1429
1462  1478  1489  1509  1549  1565  1569  1596  1634  1646  1678  1751
1790  1857  1870  1884  1902  1928  1929  1936  1954  2001  2131  2141
2241  2295  2305  2308  2337  2338  2386  2389  2408  2412  2449  2450
2457  2458  2461  2570  2572  2583    98   105   119   132   288   319
513   682   745   815   844   930   970  1060  1239  1276  1289  1309
1342  1490  1517  1563  1663  1706  1740  2129  2262];[52    71   102
107   156   163   168   199   228   236   282   298   308   366   433
452   499   542   568   577   579   590   606   614   620   641   700
726   754   810   822   872   935  1073  1075  1077  1082  1104  1127
1162  1172  1183  1201  1250  1255  1257  1272  1284  1286  1322  1341
1396  1452  1468  1482  1540  1562  1578  1588  1636  1685  1700  1703
1744  1761  1774  1781  1791  1883  1911  1919  1953  2022  2085  2135
2234  2260  2360  2365  2422  2432  2443  2517  2606  2621  2632  2764
2769  2778  2799  2843  2865  2886  2905  2910  2929  2993  3091  3116
3121  3178  3179  3181  3191  3194  3195  3259  3260  3314  3348  3365
3390  3391  3413    17    96   425   429   556   605   694   739   757
837   852   995  1076  1128  1190  1223  1311  1363  1371  1459  1512
1553  1603  1699  1796  1803  1982  2079  2086  2102  2133  2151  2431
2444  2473  2572  2648  2689  2704  2718  2752  2777  2857  3047  3081
3108  3221  3243  3313];[3    12    27    32    35    37    78   107
135   143   178   186   217   256   264   280   289   295   316   384
402   430   451   530   545   565   574   576   668   694   697   728
739   770   788   831   864   871   880   885   928   930   994  1009
1061  1093  1133  1135  1141  1145  1152  1221  1222  1234  1271  1273
1275  1321  1325  1342  1351  1357  1423  1447  1455  1468  1510  1612
1629  1646  1649  1673  1674  1724  1726  1740  1750  1823  1824  1827
1839  1886  1891  1919  1941  1971  1977  1983  1992  2001  2009  2020
2038  2083  2087  2156  2159  2194  2200  2206  2209  2212  2311  2346
2351  2381  2386  2387  2411  2522  2572  2605  2606  2637  2640  2642
2668  2675  2703  2716  2719  2760  2762  2770  2828  2843  2876  2900
2908  2922  2982  2984  3018  3024  3036  3070  3153  3163  3247  3266
3289  3331  3345  3402    20    76    99   117   141   177   181   195
241   450   452   573   688   768   819   832   896   985  1230  1704
1739  1911  2170  2242  2491  2558  2617  2727  2748  2803  2824  2932
3072  3085  3229  3249];[32    58   141   236   237   285   355   361
387   396   405   414   421   449   452   609   629   652   693   705
753   806   846   856   870   893  1018  1019  1031  1058  1069  1077
1079  1107  1130  1148  1195  1199  1201  1204  1250  1275  1329  1420
1460  1478  1575  1596  1672  1727  1753  1759  1773  1875  1911  1920
1942  1958  1979  2031  2033  2057  2124  2172  2210  2247  2251  2269
2301  2473  2512  2528  2577  2656  2678  2700  2716  2718  2731  2803
2854  2869  2901  2914  2918  2927  2967  2999  3004  3047  3052  3072
3199  3200  3213  3229  3268  3317  3320  3361  3363   169   294   356
417   424   515   552   592   604   614   831   925   984   997  1061
1157  1227  1269  1371  1472  1538  1612  1630  1647  1706  1722  1903
1961  2013  2055  2056  2067  2140  2144  2181  2183  2184  2222  2362
2420  2425  2474  2730  2825  2851  2899  3017  3197  3316];[1     5
9    10    27    37   170   184   193   241   267   276   358   365   373
```

```
393   413   420   423   480   500   506   518   591   625   652   710
719   759   760   762   765   835   843   846   887   896   917   933
1021  1027  1032  1067  1086  1092  1154  1184  1202  1208  1226  1262
1362  1374  1382  1398  1409  1420  1458  1487  1496  1516  1566  1567
1570  1584  1591  1612  1614  1627  1656  1679  1717  1745  1748  1809
1876  1930  1939  2016  2017  2019  2036  2038  2086  2102  2141  2156
2181  2192  2213  2217  2218  2286  2301  2346  2367  2371  2417  2465
2536  2578  2590  2596  2600  2629  2646  2649  2666    62   183   201
217   288   331   338   414   773   855   899   934  1097  1113  1270
1515  1560  1563  1569  1589  1814  1836  1911  2013  2026  2083  2173
2191  2220  2221  2244  2274  2283  2309  2360  2514  2568  2661]};

% SCORES= 13;
% ns={[420  2232  2594  2432  1340   167   792  1600  1311  2210  1222
2037   106  2090  1520  2530  1329   327  2528  1874   370   336  1038
1422  2358   716  1218  1307  1586  1722  2599  1215  2413  2366  2365
2014  2311   149  1609   923    22   329  1953  2225   768   625   353
1885  1903  2302  1507    83  1166   608  1726  1555  1358  1565  1177
2571   477    32  2082   216  1551  1645   573  1493   665  1200   842
1619  1887  2089  2060  1059  1768  1738  1213  2289   261  2204  1636
77  1041   698  2007   350  2527   791   712   765  1216  2129   552
82  1747  2531  2249  1701  1426    35  2140   287  1309  1873   996
1724  2008  1673   549];[4320  5701  3425  2769   186  3838  3468  2735
4297   615  4684  2089   140  1819  2927  2330  1453  1726  1507  4656
2998   608   789  2079  3601  4221  2097  3437  2129  1931  2806  2337
5442  4071  2840  3181   859  5112  2356  2367  1065  1051    36  1620
4574  4061  1674  3844   296  3889  3525  3636  4169  2196  2439  3370
1249  2562  1113  1393  2250   740  4434  5255  5157  1277  1421   808
3254  3908  1991  1372  1495   616  2282  1871  3157  1052  1622  4533
439  2546  2863  4928   199  4862     1  4546  5107  3897  4538  4092
3161  4102  1064  5000  1409  5721  1858   298  4537  5512  4788  2246
245  1814   324   593  1205   802   764  2284  2877   523  4258  1788
640  3118  2590  1685  5690  5274   173  3659   866  3302   569  2774
4484  1314  5524   270   822  3002  4009  4598  4631   894  1926  4217
3174  3975  2511  5327  5748  2142  3244   524  1321    52  3085  4151
3182  3411  4901   863  4649  5217  3898  4069   540  1185  1664  3158
1948  5333  5488  5415   436  4991  5727  3702  2280  1879   612  1652
5160  1565  5477  2545   776   641  4999  1365   338  3450  5052   417
4908   486  2674   675  5647  1937  1908  4644   941  1049   753  1617
4712    68  5464  5564  1736  3635  3405  3956   513  4437   244  4494
1139  3381  5571  1024  4542  2841  4760  1519    42   846   648   430
2474  5677  5498  4691  1186  5678  1360  1537  4121  5349   514  4811
2858];[904   729  2793   261  3481  1244  3093  3271  2056  1299  3199
2836   647  1474  1148  1020  3360  1492  2780  2697  1640  3101   876
260   888  3384  1056  1864  2249  1046   254   690  1627   792  2768
1491  2884  2620  2295  3031  2958  3072  3311  1426  1037  2820  3506
3235   366   789  3342   323   471  2921   507  1534  1693  2934  3577
1077  1805  1961  3216  3408  1047  1164  3513  1182  1689  3547  1759
676   160  1300  1024  1318  2202  2756   344  2484  2613  3502  3248
3472  1109   701  1193   193  1824  1287  1127  1289  3210   450   937
2536  2910  1842  3230  1670   790  1599    63  3386  2212  2509  3310
2436  3215  3393   408   826   429   494   663  2435  2124  2676  1354
157   648  3630  1963  2567   325  2680  1977  1460  2167  3457  2933
770  2576  2755  3453  1571  2829  2288  1776  2346  2610  2031   970
1198  3494  1823  2141   915  1163  1482  1519  3035   375   705    89
2363  1366   292];[2097   570  2059   823   160  1157  2650  2298  1265
1285   860  1094  1657  2806   436   962  1691  2631  2389  1172  1611
2163   673  1731  2041   541  2755  2666  1581    17    21  1182  1279
```

```
2893   252   672  2756  2235   736  2859   669  1444  1958  2233  2098
2570   158   738  1107   783  2583  2496  2894  1288  2569  2017   968
606   259  1272  1894  2702  1255   784  1128  2240  1824  1973  2095
689  2304  1961  1566  1701   952  1847   529   253  1635  2442  2229
2531  2292  1732  2460   441  1779  2686  2441  1557   569  1338  2660
2178  2526  2717   443  1137  2746   658  1924  1713  1055   832   350
358  1181   800  1458];[2476  2243  1940  1302  3184   281  1837   776
844  1336   800   839   904  2544  1012  1644   201  1678   727  3067
1967  1138   570   367  2822   321   544  2468  1515  1071  2350  3125
1887  2516  2635  2848  1072  2186  2343   783  3162  1939   419  1627
299  3068  2944   442   609  2001   347  2473    68  1301   675  1224
880   583   558   587   799  2336   109   573  2008  2340  2619   677
1665   341   186   417  1827   676   814   639   440  2535  1073  1289
330  1009  2035  1048   272  3283   712  1340   581  3055   171  2804
366  1646  2019   594  3132  1117   458  2789  1104   507  2536  1912
840   602   472   519    81  3113   943  3260  2361  2866  2417  3073
2265  1961   258  2364  1529  2432   489  2377];[3510   292  1754   743
2421  1385  2952  3033  1728  1836  3376  3121   963   770  1011  2787
2435  1055  3127  1015  1259  1099  1053  1853   282   961  2721  2602
98   585  1945  1124  2126   191  1430  3529  1115  2980   135  3450
2791  3671  1541  2006   962   267   192  3428  3462  2252  2613  2733
651  1299  2651  3093  3348  2472  2961  2764  1250  3239  2982  2362
281   256  2193    27   480  3539  1940  1734   275  1338    63  1135
2174  2886   908  1476  1184   134  3351  3448   186  1021  1213    52
414  2722  2497   456  1900   847  2036  3631  2792  3333  3084  1220
3346   180  1972  3005  3452  1464  3041   346   755  1453  1994   374
2258  3243  2908  1216  1946  3718   122  1242   921  3387  2444  2111
508  3374  2079  1466  3447    18  2294   439  2956  1241  1886  2328
3714   365  1868  2616  2580];[4234  1756  1670   663   423   370   714
4455   609  1968  4408  4211  2796  3249   192  2512   655  3147  1762
2390  2432   548  4466  4138  3709   629  2377  3920   102  4260  2802
3767  2347  3863  4457  3105  4527  1873  4040  1904  2441  1269  1555
1701  1973  3713  4485  1370  2920    28  1099  1915  1313  4013  3528
4415  1924  1506   274  2544  2378  3033  2780  3743  3313  1199  3416
1069   788  3204  1321   106  3346  3919  1918  1994  2287  2407  1084
3000  2869   570  3413  4032   105  3341  3408  2872  4078   614  3923
3557  3539  2886  2520  4102  4328    40   876  3533  3254  1031  1985
791   968  1320  3464  1520  4177   982  3420  1108  3017   491  2876
3043  3728  1799  3597  3479  4128  3522  2427    96  4108  3296  1992
3782  1152  4107  4385   513  1589  3466  1852  4031  3320  1990  4146
2844   249   425  1617   147   244  1655  1661  2536  3868   740  2723
1450  4227   664  2570  4181  4188   743   543  2285   967   433  1228
3914  3732  3994   661  2790  3945   516  1592  2589  2745  1950  3890
204   521  3165   532  3723  3345  1501  2609  2845];[820  1184  2613
478  1191  1226  2609  2412  1473  1499  3299   581  2079  2041  1408
2375  1845  1262  1372   903  3218   515  2003  1014   401  2126  2273
1236  2583   834  2597  2220  1884  3069  1740   487   567  1329  1992
2129  3034  1684  1064   215   436  2398  1697  1481  1115    45  2207
441   105  3181  2195  3111  3007   982  3337   521  1255  3233  2253
582   442  3033   681  2255  1719  2748  1077  2670  1538  2596   924
1152  1395   856  2229   428  3128    93  2013   850   298   702   384
2040  2406  2953  1926  1449  3326   525    19   365   420   362   550
1622   256  1190    54  2545  2374   516  3216  2230   440  2009   684
2312  1811  2323    73   137  1456   766  2692   892  2589  1128  1752
2142   617  2778  1450  2590  2350  2326   661  1921   469  2146  1728
987];[2906   542   190   890   808   114   367   418  2096  2943  2640
2038   151  3108  2884  1580  1146  3289  1851   708  2970   502  2033
305  2848   897   721   695  1031  1374   219   975  3523  1927  1180
```

437 79 477 516 2563 3062 165 1996 1247 2741 1034 923
1645 2466 2572 2021 1617 2628 1344 1548 15 838 2539 2421
2639 1584 1149 1774 2900 1069 1934 3462 2600 1983 3136 2536
3274 3497 1223 233 1127 1142 2361 2247 2810 2512 2027 573
1446 3233 2826 2001 3340 1033 1467 667 320 903 1786 876
1417 2022 1632 3425 459 746 1696 2605 979 2990 1325 2701
3005 2787 2079 3363 1326 345 3094 441 1737 2294 3457 2796
2298 819 3173 1937 1272 314 2366 3454 2809 1504 553 1198
1849 3116 2684 2451 2732 1129 3156 3002 1751 480 3227 234
491];[1235 666 760 1587 1609 41 826 2066 251 1666 1880
1213 120 1319 336 1999 2129 80 1939 1023 1337 73 1694
2582 201 2001 1315 1961 633 1252 483 419 1838 551 718
893 2413 642 1594 58 894 988 1137 629 873 2586 2321
79 1726 1126 2016 976 360 813 2513 370 1031 2188 1744
1283 788 1082 1024 2152 2211 1179 611 1816 881 2232 1136
2089 576 1364 2334 1813 176 2524 1057 2497 2467 61 2468
2473 1743 333 1896 1125 945 1202 418 1572 1442 373 1985
147 2519 1317 585 631 2191 1045 2587 2419 948];[1002 2929
1953 2392 2612 1074 2456 740 3560 645 3059 2940 2344 3588
938 4207 1361 430 4389 1373 1631 3238 1555 216 2996 667
70 2363 261 4581 2583 1573 188 1432 4034 155 4348 3985
2011 3732 1160 3272 4835 3596 3537 4462 3336 1433 4466 1813
219 283 2847 4058 2100 1842 935 4103 2659 4275 2450 4499
2698 153 4802 236 2519 458 436 2463 4178 4069 790 3668
521 4709 1866 726 4678 673 496 262 3175 1847 264 1768
3787 365 2042 1934 62 886 1989 647 2627 3646 1289 2719
2748 2508 1431 2521 583 2571 4483 3455 1694 3271 725 122
2476 2196 3311 863 3031 3450 2961 684 1524 263 2553 3796
4333 96 396 4229 3036 2138 2554 128 715 2321 324 235
1184 4577 1945 2412 4025 4568 3471 636 4662 1016 1851 641
4487 2114 1220 2839 2577 3134 1085 1105 2023 2383 2572 1303
700 492 4562 1460];[1540 910 1518 54 1153 1263 261 2256
2399 2340 270 1593 164 1777 2503 937 2267 996 7 1661
16 1907 358 933 2187 2175 805 1533 915 1282 423 2480
93 509 656 1833 8 2007 1915 2566 37 1402 801 1887
2206 1512 1168 2076 386 1823 889 529 1756 1117 1869 2218
1699 2202 1805 2005 1959 589 1344 2030 950 113 307 1908
1274 1348 1444 787 242 2434 1724 536 2217 613 6 150
1118 149 34 1660 339 1096 1728 1433 1594 2068 2513 890
168 776 1557 1137 1388 1820 644 575 2326 1662 545
803];[85 2403 777 477 1944 1450 2335 153 940 1143 2229
2596 2534 857 1172 617 1798 908 1235 899 1592 2667 1074
1154 1934 2653 547 2439 1748 597 1159 1176 1659 2435 312
552 497 1237 1534 2309 82 258 2032 1238 2691 1417 936
2614 1015 351 478 543 784 602 1742 923 726 665 556
995 1353 900 2039 2430 654 714 522 2348 2434 2436 2421
53 1118 2443 342 618 1024 653 631 1073 287 638 1641
151 341 1533 2674 438 568 1836 1003 1912 664 833 608
1623 1202 405 920 406 1859 1886 1938 1117 627 1795];[179
2277 1318 756 1779 1061 1551 2068 957 1605 1803 1 669
2299 197 2439 832 2033 594 2131 1445 722 2115 930 713
1201 1947 762 982 925 531 2203 1319 1155 1398 1032 526
752 2463 880 1882 1660 2446 981 715 1397 2284 1889 2469
909 739 2381 1786 1111 529 921 1079 1765 2523 584 1100
1387 120 1717 874 199 1446 955 1633 1056 1348 73 536
422 1715 192 253 2483 1846 811 1822 956 1716 803 1303
2177 1750 965 1887 915 2524];[36 872 1660 1035 539 385
1574 75 694 1596 191 1293 963 1389 1487 562 1703 1775

```
859  1037  1738   366  1321   232   816  1632  1806  1320   864  1447
397   707   685    89   265   625  1609  1867   132  1201  1884  1781
1848  1307  1875  1400    78   165  1329  1319   341  1624  1883  1584
1672    90  1715  1086  1905  1232    79  1439   218  1169  1906   994
997  1837  1478  1614    37  1066  1430   825];[3365   927  2032  3311
1640  2645  3558  1644  2180  3306  1012   495   678  2892  2949  2575
3637  3305  3135  1375  1735  2248   434  2090  3550  2697  3395   194
2537  3088  1899   671  3112  3237   967    71  2816   468  3099  3478
310   392  2111  1704  2247  2632  3577  3695  2847  2584  1498   700
433   596  2384  1296  2878   179  1400   636  2246  2651  1758  1345
1833  3066  3059   698   870  3403  2586  1886  1799    32  2549  2181
2631  2482  1152   404  3633  3422  2806  1972   504  1697  1396  3302
3284  3226  3097   623   899  3381  1202  3282  2688   663  3332     5
3442   641  1456   413  3720  1197   600    85  3140  3354  3057   383
2524  3310  2854   724  1850   443  1816   374   372  3694  2255  2944
2223   170  1776  1880  1014   384  1987   369  1477  3050  1018    35
3273   941  1760   169  1929  3581  1564  3068  2571  2805  3437   112
258   322  2276];[192   424  1420  1706  2408   402  1261   745   844
499  1740  1663  1164   815  1342  1030  1844   896   936  1302  1902
1295  1711   837  2131   811   715  1677  1565  1709   621  2312  1664
337    98  1060  1301   417   584  2584  1278   313  1634  1538   355
1561   565  2081  1549   396  1202   133  2012  1807  1417  1013  1614
2338  2568   569  2241  1139   767  1461  1790   758  1454   629  1413
178   244   682  1776  2493   721  2326    73   453   330  1281   747
129  1739  1184  2084   886  1550   255   441  2022   368   319   930
1214  1901    64  1563   583];[2905   168   366  3179   308  3365     6
504     8    50  1352  3081  1250   581  2602  2689  2857   199    71
1591   742  1703   428  2319  2189  1700  3117  1803  2412  3121  2819
1855  1034  1418   282  2135   930  3299  3044   217  2902    96   156
3344  1224  2675  1696  1953  1223  2572  1396   602   874  2929  2993
1431   798  2792  1862  1138  2072  1190  3195  1540  1804  1183   622
2431  1272  1322  2968  1167  2022  3056  1104  1982  2903  2362  3399
3320  1137  3282  2086  1279  1468   762  2112   338  1687  3261  3398
2486  2879  1172   514   122   458  3386   256  3005  1761   452  1607
1298  3390  2498  3375  2718  1686   579  2191  1827  2452  1301   521
822  2507  2654  1432   973  3240   606  3094  2330  3108   314  2185
852   810   840  2891   787];[2009   545   454  2351   745  2824  2703
1608  3316  2790   422    12  2170  2572  1673  3151  2034  2876   113
565  1728  1838  3404  1823  1297   452  1700  1093  1328   501  3170
1430   896   837   186  2311  1769  2666   840   255   195  1866  3153
107  1828   508  2863  2770   346   143  1632  2188  1916   374  1135
2461  1351  3289  2238  2180  2223  1873    93   282  1832  2551  2472
871    34   573   303  3314  2388  1365  2352  1342  1107  1122  2100
693  3386  2643  2242   730   856  1273   773  1271  1861  1971  3072
1646  2642  2433  2150   520   319  1330  2200  3319  2838  2392   877
1475  3036  2087  2932  1983  1663   696  3286   678   295  1894  2522
2179  1306    78  1397  2032  1941  3085  2719  1900   463  3265  2357
522   483   385  2000];[2851  1148   870  2530  3001  3213  1061  1802
1911  2909   331  1463  2362  3245   693   773   691   349  2155   424
2124   285  1815  2738  2656  2642   116  2055   997   807   547  2058
2512  1073  1880  1348  2164  2897   333  2803  1708  3124  3062  1147
2543  3127    59  3225  1115  1874   925  1637  2802  3002  1639  1579
1489   940  1733  1499  1477    67  1018   894  2247   437  2545  2183
568  3215   838  3138  2474  2099  3200   769  1944   999  2248   806
2323   265   715  1314    31   592  2077  2428  3301  3110   442  2367
770  1218   602   643   539  1730  2270  1713  1895  2644  1761   533
1296  1987];[1614   773  1651    95   759   431  2141   183  1001   201
2083   373   933  1531   170  1362  1656  1683  1748   414  1087  1980
```

```
1942  2408  2217  2108   784   413  1745  1516  2514   855  2264   812
887  2629  2086  2129  1520  1694   274   217   844  1321  2117   499
2465   577  2458   906  1361   384   249  1403  2481  1583   562  2173
2061  2130  2371  1630  2581   953  1592   365  1994  2427    62  1886
1695  1123  1006  2313  2596  1599  1317  1131  2650   181   766  2066
1420   424  1682  2166  2220  2222   318  2295  2349];};
% SCORES=14;
% ns={[106    21  2232  1177  1355   354  1218  1520  1311  2528   262
420  1600  1551  2210  2403  1722   625   894  2302  2530  1329  1439
2313  2358  2575  2594   923   970   167    80   342   555   249   336
484   969  1885  1903    77   327   985  1609  2027   172   613   716
720  1269  1274  1730    35    88   718   842  1222  1340  2037  2060
2571    78   370   573   713  1039  1166  1178  1219  1786  1957  1103
2469   505   698  1143  1450  1657  1701  2156  2230  2557    16    40
350   410   645   792  1021  1251  1307  1636  1683  2014  2066  2432
2511];[2089  1937  1277  4320  4099  5425   593  3180  4991   866  3838
5255  1931  2104  1372  4583  5701  4325   140   186   439  1482  3836
4217  2284  3468  4092  2079  2545  5758  1128  2337  5336   514   616
2250  2769  3849  1705  1983  2142  3601  4297  4940  5751   615   867
913  1365  1819  2000  2439  2562  3893  4332  5207  5721   298   324
4535  5107   789   802  1186  1866  2854  3291  4017  4438  4681  5392
5571  5690   796  1205  1863  1980  3370  4484  4939  5099  5333  1633
2246  3086  3118  3157  3425  3898  5390    34   672   872  1726  1838
2905  3128  4300  4995    68    82   607   949   951  1617  1674  1685
2086  2097  2409  2474  3956  4100   231   233   244   372   753  1136
2590  2949  2998  4662  5395  5477   663   945  1262  1507  1670  1858
1871  2546  3166  3192  4121  4184  4308  4649  5327  5398    89   524
862  1052  1135  1519  2005  2984  3161  3243  3798  4444  4684  4741
4792  4846  5157   174   323   535   608   648   822  1421  2163  2344
2367  2369  2735  2738  2840  3142  3182  3224  3333  3405  3525  3766
3878  4013  4071  4175  4606  4631  4724  5397  5488];[904  3101   729
292  2793  1350  2836  3199   756  3324   905  1426  1056   254  2572
2829  2620  3311  3342   876  3082  3513   285   647  3491  3532   298
1864  2933  3172  3282   366   675  1020  1349  1474  1819  2055  2295
2613  2958  3093  3388   261   364   489  1019  1299  1492  1977  3481
3555   352   889   986  1262  1300  2249  3228  3360  3453  3506  3619
363   559  2884  3119   408   888   902   943  1106  2056  2373  2576
2921  3116   238   859  1029  1037  1129  1213  1678  1901  2409  2735
2780  2889  3094   590   730   792   922  1307  2138  2209  2929  2934
3176  3335    38   214   355   404   785   989  1077  1084  1148  1842
2042  2193  2259  2574  3216];[1338  1849   290  1615  1710  2061   861
1172  2755  2919  1310  1371  2059  2086   855   570   823  1383  2460
989  1117  1288  2097  2353  2738   216  1626  1699  2126  1390  1691
2806   317   689   860  1439  1888  2389   568   647   673   685   884
942  1847  2235   553   706   763   973  1766  2509  2702   218   869
1157  1479  1731  2650  2859   487   738   910   932  1011  1094  1268
1825  2304  2756   253   465   520  1080  1405  1657  1704  2387  2462
158   175   597   669   952  1265  1345  1492  1765  1798  2345
2685];[3166  1052  1302  1887  2848  2949   775  1627  1065  1336  1835
2476  3067  1940  2829  3132   420  2124  2804   369  3068   800  1214
1961  1967   776   880  1569  3055   677   819  2001  2088  2635  3254
321   541   543  1198  1871  2442   171   269   442   594  1837  2417
2997   675  1422  1740  2243  2336  3198    51   839  2255  2670   384
457   503   727  1043  1380  2695  2998  3184  3230   281  2094  2473
2535  2544  2622  2826  3260    65   180   399   636   826  1868  2516
3029  3186  3275   330   367   538   544   616   791  1012  1073  1529
2012  2440  2822  2827  3060  3125    75   272   325   342   609   844
1010  1341  1364  1539  1595  1757  1956  2377  3263  3265  3339];[192
```

1632 2961 961 2907 3620 3705 510 1878 1208 2613 2939 453
2251 2297 3014 3510 1011 1867 3006 439 1015 1117 2765 3606
585 1501 2190 1221 1343 1482 2079 292 743 1099 1754 1836
2252 2887 282 2093 2421 3240 3333 138 37 1254 2615 2891
2898 3005 3093 1079 1467 2006 2916 3351 3583 1124 1430 1541
1940 2027 2804 2821 3088 14 394 812 964 1476 1728 2660
3529 132 134 1053 1135 1163 1431 1797 2322 2328 2435 2585
3540 3728 347 374 1184 1250 1259 1338 1799 2664 2763 2950
2952 3076 3239 3374 63 139 178 313 525 723 963 1012
1299 1781 1831 1851 1929 2337 2787 3025 3113 3268 65 191
234 268 480 638 921 1021 1211 1751 1886 1965 2112 2387
2533 2689 2853 2976 3376];[2802 3320 2390 4138 4455 423 1756
3079 663 370 1154 1069 1450 3782 192 1904 300 3709 4013
147 2285 3920 1014 1150 2845 4032 4171 4260 1506 1985 2808
3359 4234 4466 1084 1108 2096 2683 151 249 609 714 777
1199 1628 2378 2470 2538 4107 4244 3364 4102 4212 1517 2796
3204 4129 106 1908 2196 2297 3247 3519 3713 4225 548 2920
3043 3743 3908 4183 1152 2156 2550 2590 3017 3105 4232 4237
105 569 614 796 1508 1873 1990 2504 2844 2960 3213 3353
4211 4408 4485 252 1006 1269 1639 1670 1968 1992 2038 2124
2428 2441 2570 2721 3126 3210 3314 3391 3413 3669 4209 4415
159 891 1370 1438 1737 2287 2427 2790 2830 2919 3481 3898
4007 4044 4108 4300 4331 4587 283 361 1606 1608 1687 1769
2432 2507 2684 2991 3165 3914 4548 28 71 239 934 1177
1252 1626 1718 1746 1887 1950 2070 2420 2869 2902 3249 3416
3616 3863 3945 4177 4510];[1184 1236 3326 2126 1191 1499 2583
3063 515 2609 820 1226 105 138 1077 2079 2761 2613 3021
478 579 1078 1372 2289 137 398 2041 2282 1128 1140 2380
441 903 1118 1395 2195 2717 1481 2387 2569 3181 438 731
2142 2263 2290 3069 1152 1298 1535 2468 1697 1758 2412 3008
3299 19 477 582 702 850 1056 1065 1262 1572 1845 2359
3314 215 1014 1203 1449 1925 2099 2670 2961 3007 401 1294
1426 1926 1984 1992 2207 2220 2273 2633 3060 3233 397 487
648 978 1681 2009 2043 2056 216 409 925 1187 1448 1811
2003 2375 2692 2727 2955 3080];[2970 542 190 2906 151 3283
2038 543 1247 1421 3062 1137 897 1328 1927 367 469 502
2628 638 1567 3451 3497 627 937 1752 2069 2752 2848 3423
79 418 903 1180 1580 1637 1934 2741 2805 2944 3108 3284
144 878 890 1326 1540 1579 2640 2743 519 1050 1478 1504
2003 2298 2639 2755 2794 2539 2744 3422 3523 392 808 1146
395 579 923 1344 1374 2033 2133 2796 2943 3425 49 516
909 1142 1258 1589 2167 2276 2381 2510 2563 3005 3348 3518
123 736 933 1031 2065 2604 2701 2879 540 976 1038 1761
2021 2334 2572 2705 2826 2884];[1939 359 1587 666 1126 57
2001 1319 1235 760 2090 475 2586 633 2473 826 1112 41
894 905 1337 1739 2582 718 1880 69 747 1057 1609 942
1814 2513 73 419 658 820 2419 1522 551 1364 1666 2355
80 201 292 621 663 1208 1406 1875 2587 290 373 1276
1349 1999 2066 2129 2152 251 699 1137 1363 1538 1695 1726
2259 79 176 597 1101 1125 1160 1213 1442 1589 61 354
378 440 576 750 1073 1315 1469 1652 1677 1697 2158
2567];[4275 1361 1555 4203 928 2392 4516 1953 3367 1432 3782
4581 263 1002 1125 4182 1218 2363 3238 938 2583 2801 790
2698 3716 4389 216 697 1424 2659 2456 3096 4339 179 647
1183 1232 2066 2570 2611 4466 583 1954 2121 2765 3059 3103
3342 3514 4231 4879 106 726 740 1074 1501 1736 1789 2344
2940 3019 3560 4563 4584 15 62 231 1989 2084 2353 2571
2778 2941 3900 3985 4035 4246 1173 1738 2122 2138 2334 2606

2612 2731 2929 3325 4229 4330 105 1431 1842 2514 3272 3513
3679 3689 4043 4329 4475 218 430 601 935 973 977 1189
1291 1768 1933 2902 3057 3182 3197 3596 3638 4207 4368 4539
4783 128 435 586 646 715 1131 1934 2013 2123 2171 2199
2257 2402 2459 2839 2888 2996 3680 3768 3976 4084 4208 4462
4574 4582 4837 667 728 1026 1043 1341 2064 2134 2580 3537
3899 4079 4090 4178];[1834 1540 889 2203 38 937 2340 164
2565 1290 805 1153 54 280 16 509 910 1493 656 1357
2399 22 184 1518 288 97 366 741 1117 1161 1358 1402
1890 7 168 588 1303 1388 2440 358 795 1858 1949 2404
8 196 261 784 955 1583 1632 1820 2075 2503 1061 1756 2256
275 739 2202 2294 270 328 415 613 742 859 890 1137
1305 1538 1725 1866 34 536 933 1237 1263 1593 1628 1805
1817 1839 2009 2187 2308 2384 2480 2532];[2667 54 777 477
153 522 654 924 1237 2436 1172 1346 1944 463 834 908
2403 266 1646 2334 85 552 1334 1859 638 816 1789 2134
2335 945 1024 1154 1691 2439 1533 2430 618 2309 2596 478
1202 1450 2070 617 1834 357 1483 2341 940 1972 2039 605
608 1047 2032 2555 560 899 1245 1285 1290 1478 1748 1752
1913 2539 2591 40 106 287 312 597 1143 1235 1404 1742
1750 458 607 653 665 1593 1702 2011 2229 2435 1561 1854
2447];[179 1715 2277 1100 2180 740 1634 2056 2068 2445 832
1765 2254 2299 2415 925 1201 1549 756 761 1297 2439 722
1495 993 1882 1398 831 1397 1505 1689 54 2239 113 309
1845 2195 404 1803 1830 2131 2381 529 915 243 367 713
1680 1916 2136 2469 2472 245 253 530 569 594 1061 1156
1725 1889 2033 449 982 1099 1111 1147 1203 1248 1551 1671
1672 1848 2245 1427 1521 2115 2193];[341 1228 808 938 179
1389 1574 1660 36 540 562 599 1738 42 1487 694 993
1556 37 872 1065 1614 1848 89 880 1035 1729 1787 1790
412 754 1578 1591 1652 1781 1875 78 1490 1678 90 191
901 1297 1613 1653 1791 196 581 582 755 1624 101 180
825 913 990 1201 1905];[1640 3365 3550 2573 2574 2847 1312
3182 1149 2892 678 2032 2727 85 106 429 1482 596 1758
3364 1345 2180 2537 3059 3536 1704 3305 1877 2181 2549 2645
3395 3432 468 540 1385 2111 330 2247 3041 3188 319 927
1197 1697 1821 2944 3558 3637 226 321 1850 2526 2632 2711
3306 3311 3638 3673 105 180 333 643 1304 1400 1896 2625
3380 3713 3741 322 434 1464 2081 2248 3135 3237 1152 1375
1849 3068 71 374 443 1299 1776 2631 3066 3326 3367 3381
3437 3535 179 403 421 495 739 812 1182 1747 1805 1899
1957 2383 2956 3271 3310 3471 3560 3626 3694 3703 698 700
967 972 1012 1029 1084 1543 1735 1783 2667 3230 3602];[1420
936 424 2408 1549 568 1277 2001 815 192 1663 2583 1565
745 1089 1261 1634 1295 1706 1857 519 1060 1406 499 105
402 1289 1844 2262 2457 2570 751 1129 313 1405 1695 98
906 1302 1596 682 835 1169 1936 2131 2141 2223 2305 120
502 837 844 896 1301 1790 2129 174 208 298 337 1538
1614 569 671 886 2241 52 114 132 422 698 1517 1550
1592 2022 73 83 330 454 715 811 833 867 978 1164
1281 1462 1502 1711 1740 1884 1954 1996 2308 2389];[3195 3179
3365 2905 71 1250 1703 2843 308 1699 2135 3314 1172 1183
1700 3121 282 366 508 1588 2929 810 2063 2411 2689 2799
514 872 1512 1531 3058 236 504 742 1298 1532 3299 3353
6 602 1255 1540 1804 1975 428 1791 3081 41 337 451 694
1399 2144 2592 2602 2648 2792 3047 1761 2364 2443 2908 3178
3243 168 421 874 1104 1154 1322 1352 1432 2349 2583 3181
156 739 787 1431 1468 1628 2718 2884 2886 3240 3408 216

```
499   639   935   976  1205  2319  2539  2632  2675  2857   163   184
199   641   852   983  1049  1137  1138  1190  1201  1591  1774  1905
1953  1982  1999  2022  2071  2548  2734  2769  2881  3441];[107  1093
2351   143  1674  2572  2381   141  1971  2009  2020  3072   303  2170
2876  3316   768  1122  2803  1941   545   819  1107  2522   113   195
1724  1919  2982   384  2770  3151   463   573  2521  3111   291   668
744   770   840   850  1608  1646  1673  2034  2223  2455  2461  3289
186   289   534  2605  2668  2703  3402   217  1342  3085   127   430
454   568  1697  3036    12    32   451   530   678  1041  1151  1328
1430  1688  1726  2387  2642  2824  2843  3306    20   295   985  1271
1287  1300  1792  1832  1983  2242  2491  2820  3386    34   282   501
565   745  1047  1265  1321  1447  1715  3073   450   728  1016  1135
1145  1222  1275  1351  1375  1728  2200  2637  2848  3141  3170
3359];[846  1911  1148  2927  3213   414  1250  1800  2031   294   870
1371  3320  1802  2124  3316  1130  3363   693  1538    55   285   997
1018  1275  2055  2362  2718  2851   260   355  1129  1942  3317  1647
1727  2731   488   705   987  1730  2247  3001  3200   259   417  1612
2159  2474   806  1340  1630  1958  2644  2730  3183  3268    32   421
505   643  1761  2775  2869  2967  3072    82   592  1061  1472  1489
1672  2033  2058  2199  2797  2803  2918   424   568   831   999  1460
1708  2120  2237  2301  3123  3265];[1591  1614  2218  2023  2600  1570
2286  1745  2130   933  1184   652  2141  2346   431  1262  1362  2217
1516  2536   414  1748  2283   288   499  2647   201   393  1566  1612
2590   759   846   998  1154  1567  2108  2309  2408  2629   331   373
577   855   887  1001  2244  2578   358   413   816  1531  2086  2264
37    95   761   934  1006  1027  1226  1536  1738  2523  2649   773
1087  1346  1382  2192   187   272  1097  1104  1255  1351  1401  1420
1487  1491  1651  1942  2213  1361  1515  1694  1695  2639   625   896
1021  1270  1360  1656  2129  2338  2367  2596  2634];};
% SCORES=12;
% ns={[1355   106    21  1177  1722  2232  2210  2358  1520   249   354
923  1329  2302  2530   131   262  1218  1311  1551  1600  1851  2313
2403  2528  2571    80   167   420   970  2127  2568  2575  2594    88
172   327   328   336   484   625   894  1237  1439  1493  1800  1951
1986  2158  2239   342   350   555   613   647   720   842  1103  1178
1251  1599  1609  1683  1762  1786  1885  1903  2060  2278  2511    16
35    77   216   410   667   698   713   716   969   985  1016  1039
1143  1219  1269  1274  1299  1450  1585  1730  1957  2027  2037
2567];[1128  5425  1277  2089   593  1937  2562  4583  4099  4325  3180
4320  4991  5207   866  1203  1931  2104  2545  3333  3525  3836  3838
5758   140   615   753  1372  1482  1633  2079  2337  4092  4217  4438
5721   186   324   439   949  1980  3086  3468  3893  4535  4729  5255
5336  5392  5701    63   514   913  1051  1617  1705  1838  2097  2142
2284  2383  2905  3128  3368  3601  3849  3956  4300  4332  4662  4681
4995  5395  5483  5638    82   174   237   244   298   616   867  1135
1136  1365  1421  1507  1519  1819  1863  1881  1983  2000  2086  2246
2250  2439  2474  2589  2769  2794  2854  3172  3206  3291  3425  3798
4038  4235  4484  4649  4718  4846  4919  4940  5333    31   187   200
231   270   323   524   607   672   789   802   822   945  1186  1501
1866  2005  2387  2409  2535  2546  2590  2773  3018  3063  3118  3181
3182  3302  3672  3965  4017  4184  4297  4606  4612  4631  4682  4741
4914  4939  5099  5107  5157  5278  5327  5390  5561  5571  5690  5703
5727  5751];[363  2793  3101   285   729   904  3082  1350   292   756
1426  1492  2249  2836  3199  3311  3324  3619    40   905  1289  1349
1476  2055  2138  2829  3068  3228  3491   675   876   888  1056  1260
1474  1759  1977  2613  2958  3172  3342  3481  3506  3532   254   355
559   590   647   682   937  1029  1084  1500  1720  2295  2373  2572
2576  2620  2708  2960  3191  3282  3312  3513  3597  3626    25   118
```

260 298 364 366 408 416 489 567 873 889 922 986
1019 1020 1037 1129 1262 1290 1299 1300 1370 1609 1645 1819
1864 2409 2574 2676 2696 2889 2933 3093 3252 3384 3388 3414
3550 3560 3630];[1338 1849 1615 823 855 2086 290 1048 1699
2738 1172 1288 1371 1710 2755 861 1439 1684 1704 1731 1766
1888 2061 2387 2460 2919 570 673 738 1310 1383 1847 1949
2059 2471 158 163 694 973 989 1117 1390 1626 1798 2400
2509 2613 2806 205 216 689 731 910 942 1345 1667 1700
2097 2353 2685 2705 2756 2808 31 218 254 278 421 487
520 647 685 706 763 810 860 869 884 916 917 974
1041 1094 1157 1405 1574 1610 1625 1691 1714 1765 2096 2126
2177 2389 2462 2650 2697];[1887 1302 3166 369 1336 2001 1627
2088 2635 2848 677 775 1065 1771 1940 2124 2949 819 880
1052 1073 1214 1837 1871 2829 3132 51 171 321 325 420
1493 1835 1967 2476 2804 3029 3067 3169 594 776 800 808
1021 1172 1569 2122 2243 2417 2670 2821 2921 3068 3125 3186
3198 65 180 229 330 384 457 541 543 544 1043 1102
1198 1341 1539 1961 1990 2255 2336 2442 2695 2827 3055 3230
3254 8 269 272 290 367 427 442 522 574 675 791
1041 1053 1257 1422 1473 1706 1732 1781 2008 2473 2516 2573
2681 2826 2997 2998 3041 3184 3289 3339];[2961 453 1254 1632
2907 192 3705 138 961 1208 2297 2939 3014 3606 3620 282
439 510 1117 1221 1501 1754 1878 2613 2615 2765 2853 331
507 585 1015 1867 2251 3583 139 292 1011 1467 1482 1693
2887 3006 3033 3240 3510 37 178 665 743 770 963 1099
1308 1338 1343 1836 2027 2079 2190 2421 2689 2763 3122 3333
3392 134 297 374 394 721 734 964 1299 1439 1797 1799
1831 1929 1940 1956 1965 1992 2006 2093 2252 2322 2337 2580
2585 2738 2804 2891 2956 2976 2999 3088 3093 3113 3302 3351
3449 3540];[4138 1628 2802 3320 1069 2390 105 423 1154 1756
1851 1904 4455 192 370 714 1199 1450 2285 2923 3079 3126
3314 106 152 609 663 1108 1152 1639 2096 2808 3709 3782
3848 4013 4102 4107 4129 252 283 300 1014 1150 1376 1380
1506 1769 1887 1985 2409 2518 2845 3920 4183 4212 147 249
796 826 970 1084 1369 1393 1438 1761 1873 1968 2084 2378
2420 2470 2538 2683 2796 3105 3220 3359 3364 3413 3481 3616
3713 3908 4032 4171 4225 4232 4234 4244 4260 71 151 569
614 777 891 1006 1127 1606 1668 1908 1973 2124 2196 2297
2310 2593 2824 2831 2960 2997 3017 3043 3044 3204 3213 3336
3353 3519 3539 3898 3914 4007 4008 4040 4209 4331 4334 4466
28 159 258 575 719 1198 1247 1508 1517 1608 1737 1746
1850 1990 2038 2156 2261 2504 2536 2550 2590 2684 2745 2772
2920 2949 3122 3210 3247 3391 3669 3743 3750 3775 3810 3968
4044 4139 4211 4237 4347 4350 4408 4415 4485 4515 4518
4587];[1184 445 2583 1140 1984 2569 731 1191 1236 1294 1499
2126 3326 138 515 579 820 1226 2079 2282 2609 2633 2717
3021 3063 1077 1372 1405 1481 1572 1811 2359 2380 2955 105
215 850 1128 1395 2041 2195 2289 2290 2387 2613 2761 137
266 398 438 441 477 478 903 1078 1118 1152 1298 1307
1374 1602 1697 1779 1800 1925 2043 2056 2263 2273 2423 2520
2652 2730 3008 19 216 409 787 834 925 952 1014 1020
1065 1448 1449 1535 1758 1926 1992 2142 2145 2414 2468 2607
2737 3034 3069 3080 3181 3299];[2970 2744 190 1137 469 542
638 1421 1710 3283 2906 3423 151 1478 2038 3062 481 543
1146 1247 1374 2003 2944 3110 3284 3451 15 79 367 502
516 838 1326 1328 1567 1934 2069 2513 2752 3108 3476 3497
49 77 144 573 729 736 897 1180 1467 1471 1500 1504
1589 1927 2128 2133 2628 2701 2796 2901 3005 3302 80 123

277 358 418 499 519 627 677 878 903 937 979 1050
1142 1297 1533 1545 1579 1580 1637 1752 1979 2085 2167 2510
2639 2640 2695 2741 2743 2794 2805 2848 3006 3227 3288 3422
3518 3534];[1939 359 633 666 1126 1587 1112 2582 41 57
658 718 760 894 1337 2473 2576 2586 475 905 1235 1319
1914 2001 2090 419 441 826 1276 1349 1880 73 79 80
747 820 1057 1739 1875 2188 2238 2419 2448 2494 2569 17
61 69 251 292 440 551 942 1073 1208 1609 1650 1726
1814 2152 2209 2259 2324 2513 176 201 354 373 621 750
864 976 1160 1364 1406 1522 1674 1695 1962 2129 2191 2249
2355 2587];[4275 1361 2801 1555 231 1002 4516 583 928 1933
3782 4203 4466 4475 263 790 1413 1953 2392 3019 3272 3367
211 430 1218 1432 1989 2698 2902 4563 4879 15 216 218
697 731 1125 2363 2503 2863 2940 3134 3238 3342 4182 4389
4574 4581 4584 179 781 808 938 1183 1310 1424 1723 1724
1738 1789 1954 1957 2064 2134 2344 2583 2659 2731 2778 2870
3096 3197 3514 3917 3976 4043 4187 4246 4328 4503 4783 62
740 1131 1232 1431 1573 1736 1813 1842 2066 2121 2171 2353
2456 2570 2571 2941 3057 3416 3556 3560 3578 3596 3716 3768
3789 4035 4090 4181 4208 4231 4330 4339 4355];[1303 2203 889
805 1540 1834 38 54 184 366 656 1632 1866 2340 2440
16 164 275 937 1196 1290 2365 2565 509 910 1117 1153
1357 1431 2075 2399 8 97 128 280 741 1049 1118 1161
1388 1493 1547 1699 2121 2404 7 22 34 74 288 588
739 1061 1402 1518 1626 1858 1890 2009 2202 2256 168 358
613 761 784 795 859 955 1237 1358 1725 1772 1820 1839
1850 1915 1949 2067 2110 2176 2187 2287 2503];[477 522 2667
54 552 654 777 153 816 1346 2439 908 924 1024 1154
1334 1483 1646 1859 2309 2436 266 463 834 945 1172 1237
1944 2134 2287 2640 85 106 173 638 1404 1531 1533 1789
2032 2334 2335 2403 2430 2596 357 470 597 617 899 940
1048 1202 1247 1691 2107 2341 2591 172 287 458 478 605
618 634 920 960 1047 1186 1245 1356 1450 1560 1752 1834
2017 2039 2070 2229 2539 2580 2684];[2180 1715 925 1100 2277
179 1634 2299 2424 309 740 756 832 1201 1549 1765 1882
2056 2068 2254 253 722 761 993 1817 2093 2415 2445 803
876 1156 1297 1583 1689 1803 2413 2439 113 714 982 1398
1495 1505 2023 2131 2136 2195 2270 2340 2364 2496 54 243
327 404 449 530 569 584 713 736 831 1099 1147 1203
1248 1319 1397 1513 1521 1551 1671 1672 1673 1717 1750 1845
1993 2115 2193 2239 2348 2381 2469];[1228 993 341 808 42
578 599 938 36 179 498 754 1389 1653 540 617 948
1574 1787 89 320 562 1297 1487 1556 1660 1738 1848 1875
101 143 412 683 880 1444 1578 1591 1613 1678 1790 29
37 78 180 196 297 473 694 872 895 901 937 1039
1065 1424 1432 1490 1595 1729];[1312 322 2573 2574 106 1640
3550 3365 3432 1149 1482 1896 2032 2727 2847 3182 3699 85
596 678 1877 2111 2892 3741 71 105 429 643 814 925
1385 1758 2181 3364 3416 3503 3536 3703 330 468 540 815
927 1197 1345 1400 1697 2180 2247 2248 2383 2482 2625 2645
2672 2944 3041 3059 3213 3381 3395 374 496 1014 1304 1375
1498 1704 1744 1783 1849 1850 1871 1899 2081 2182 2537 2549
3080 3135 3305 3332 3347 3478 3626 3694 3713 3719 3748 3750
180 226 319 321 372 406 434 619 739 770 812 1152
1191 1212 1509 1621 1805 1821 2091 2101 2208 2376 2526 2632
2654 2711 2957 3003 3130 3148 3188 3306 3354 3380 3398 3437
3471 3518 3558 3637 3638 3668 3678];[1596 1420 568 936 2408
424 1549 1565 2001 519 1060 1129 1261 1277 1634 105 221

```
815   961  1295  1663  2570  2572  2583   192   499   592   745  1000
1406  1417  1706  2450   174   312   431   751  1013  1089  1301  1462
1857  1929  1947  2141  2223  2338  2464    98   120   270   298   313
337   402   502   569   835   837   886  1168  1169  1239  1281  1289
1302  1695  1790  1844  1882  2186  2262  2389  2457   132   208   682
715   747   803   896   906  1202  1405  1517  1678  1936  1996  2131
2305  2386  2407  2449];[3195  2905  3365  1250  3178  3179  1183  2135
3121  3314    71  1699  2929   282   337  1201  1588  1700  1703  2365
2843  2865  3007  3353   308   366   742  1159  1298  1774  2063  2411
2517  2792  2799  3191   156   236   504   606   810   862   872  1172
1203  1257  1272  1510  2144  2234  2260  2374  2583  2606  2884  2910
3181    41   114   428   451   508   532   572   602   622   739  1032
1137  1154  1255  1322  1399  1432  1512  1531  1532  1540  1578  1590
1628  1629  2270  2648  2689  2886  2908  3017  3047  3058  3081  3177
3299     6   184   212   216   227   514   542   579   639   694  1104
1462  1761  1791  1804  1822  1911  1975  2130  2349  2364  2443  2592
2675  3116  3391];[1919  2009   107  1093  2020   143   141   770  2572
1107  1674  2170  2351  2668  2828  2876   217   608   668  1971  2381
2521  2982  3072  3151    34   113   256   303   545   659   678  2522
2922  3111  3316  3402   195   289   384   573   768   850  1037  1041
1122  1151  1646  1724  1728  1792  1941  2605  2770  2803  3036  3153
264   291   463   530   534   574   728   731   819   840  1026  1034
1067  1145  1213  1222  1423  1510  1673  1832  1873  2002  2034  2120
2126  2156  2311  2461  2703  3289  3306  3359    27   178   181   186
295   451   457   568   599   744   885   924  1142  1266  1273  1275
1328  1342  1357  1375  1608  1697  1726  2159  2200  2223  2353  2386
2455  2640  2843  3073  3085  3211  3249  3422];[846  3213  1911  2927
3047   414  1129  1148  1250  2031  3268  1942  1979  2247    55   294
870   987  1130  2797  3316  1201  1371  1800  1802  2055  2362  2803
3077  3317  3320   259   285   355   396   592   643   693  1489  1538
1630  1958  1985  2124  2159  2528  2918  3183  3363    31   169   260
452   705   893   958   997  1018  1035  1275  1472  1612  1665  1727
1730  1853  2237  2269  2718  2731  2775  2851  2869  3001  3093
3200];[1591  1614  2600  2023  2286  1570  2130  2218  2590   652  1184
1566  2283  2346   414   933  1154  1262  1516  1565  1567  1745  2217
2629    37   393   625   761   816   998  1362  2102  2141  2536  2650
201   288   358   373   413   431   499   846   887   917  1021  1226
1536  1612  1748  2192  2528  2578  2646  2647   145   577   591   759
785   934  1001  1006  1027  1067  1401  1487  1531  1853  2086  2112
2211  2244  2264  2347  2549  2652];};
% % SCORES=9;
% % ns={[14    22    32    82    83   106   149   167   171   214   216
287   315   327   329   336   353   370   409   420   467   477   502
549   550   552   573   608   667   698   707   712   716   723   768
791   792   842   918   923   985   996  1016  1038  1059  1166  1177
1200  1213  1215  1218  1222  1309  1311  1324  1329  1340  1353  1419
1422  1426  1450  1493  1507  1520  1555  1565  1586  1600  1609  1619
1645  1722  1724  1726  1747  1768  1833  1873  1874  1885  1887  1903
1953  1990  2037  2089  2090  2129  2140  2176  2204  2210  2225  2232
2289  2302  2311  2314  2365  2366  2403  2413  2432  2472  2528  2531
2544  2571  2594];[31    36    42   140   174   186   199   245   288
296   298   337   338   341   428   430   436   486   513   523   524
535   563   593   608   612   615   616   641   648   734   740   753
764   776   789   796   802   808   846   859   863   866   894   941
950  1051  1052  1064  1065  1066  1113  1139  1185  1186  1205  1249
1253  1277  1278  1321  1326  1360  1365  1372  1409  1421  1453  1475
1495  1507  1559  1622  1652  1674  1705  1726  1788  1814  1819  1858
1860  1862  1871  1876  1879  1926  1931  1942  1948  1950  1959  1991
```

2079 2089 2096 2097 2129 2143 2196 2246 2250 2280 2284 2330
2337 2356 2367 2546 2590 2674 2678 2735 2769 2774 2788 2806
2840 2841 2858 2863 2877 2927 2966 2986 2998 3002 3063 3078
3085 3118 3157 3158 3174 3179 3181 3213 3244 3291 3309 3357
3370 3425 3437 3468 3497 3525 3546 3555 3565 3600 3601 3626
3636 3659 3702 3815 3838 3844 3869 3889 3898 3907 3908 3956
4009 4061 4069 4071 4092 4102 4121 4161 4185 4197 4217 4221
4258 4297 4300 4315 4320 4375 4418 4434 4455 4494 4533 4542
4546 4574 4598 4649 4681 4684 4717 4764 4788 4862 4867 4887
4901 4928 4980 4991 4999 5000 5052 5107 5110 5112 5134 5139
5157 5217 5235 5255 5390 5391 5395 5415 5431 5442 5444 5488
5512 5524 5571 5647 5678 5701 5721 5727 5744 5751];[63 89
142 157 160 181 254 260 261 292 315 323 325 337
366 375 400 405 408 450 453 471 494 647 663 690
705 729 770 792 817 823 840 876 888 902 904 937
1020 1024 1037 1038 1046 1056 1077 1109 1148 1164 1175 1198
1244 1282 1299 1300 1318 1350 1354 1426 1460 1474 1492 1500
1534 1615 1627 1633 1640 1645 1670 1689 1693 1759 1805 1823
1824 1842 1864 1897 1961 1963 1977 2031 2042 2056 2124 2193
2202 2249 2251 2288 2295 2393 2435 2436 2444 2457 2504 2509
2536 2567 2610 2620 2680 2697 2755 2756 2768 2780 2793 2836
2866 2884 2910 2921 2933 2934 3031 3035 3072 3086 3090 3093
3101 3199 3235 3271 3311 3342 3360 3384 3393 3408 3419 3425
3457 3469 3472 3481 3502 3506 3512 3547 3577 3598 3630];[15
17 21 22 142 144 158 160 252 253 436 465 467
529 541 569 570 606 656 672 673 689 702 717 736
738 742 763 783 823 860 916 962 968 1055 1071 1094
1128 1137 1140 1157 1163 1172 1181 1182 1265 1272 1279 1285
1309 1310 1348 1386 1437 1444 1458 1557 1581 1611 1655 1657
1684 1691 1701 1713 1731 1732 1766 1779 1798 1815 1878 1894
1949 1958 1973 2010 2017 2041 2059 2085 2095 2097 2100 2137
2163 2178 2229 2233 2234 2235 2264 2284 2292 2298 2345 2389
2428 2429 2460 2496 2531 2570 2583 2584 2624 2631 2650 2666
2676 2701 2702 2715 2717 2746 2755 2756 2805 2806 2859 2893
2894];[40 47 68 109 142 180 201 243 272 274 281
299 321 330 341 347 366 367 369 417 419 440 445
472 487 507 508 509 519 544 558 570 573 581 583
587 594 602 609 639 675 722 727 763 776 799 800
814 838 839 840 844 880 904 914 943 1009 1012 1038
1048 1054 1071 1073 1090 1104 1117 1138 1163 1204 1265 1301
1302 1336 1340 1353 1408 1429 1456 1480 1513 1515 1627 1644
1646 1665 1678 1706 1793 1805 1827 1837 1887 1901 1912 1940
1967 2001 2019 2024 2035 2094 2124 2186 2217 2219 2243 2336
2340 2343 2350 2432 2468 2473 2476 2501 2536 2544 2561 2573
2593 2619 2667 2789 2804 2822 2848 2850 2866 2980 3029 3067
3068 3073 3078 3107 3113 3125 3132 3153 3162 3181 3184
3244];[18 30 37 52 63 65 127 134 135 180 186
191 192 275 281 282 292 343 365 374 453 455 507
508 585 651 743 770 847 905 908 960 961 962 963
993 997 1009 1011 1012 1015 1021 1032 1047 1048 1053 1055
1077 1079 1115 1176 1186 1222 1250 1259 1299 1385 1430 1476
1498 1521 1541 1728 1734 1742 1754 1836 1853 1874 1883 1886
1900 1940 1945 1950 1956 2006 2050 2066 2079 2126 2140 2170
2174 2252 2258 2284 2291 2294 2319 2328 2333 2421 2435 2444
2465 2472 2530 2559 2600 2602 2613 2615 2643 2651 2669 2695
2721 2722 2733 2764 2787 2791 2792 2854 2886 2891 2899 2902
2916 2952 2973 2978 2982 2989 3033 3041 3071 3084 3093 3094
3112 3121 3127 3158 3240 3243 3335 3346 3348 3376 3378 3441

3448 3450 3452 3510 3529 3539 3629];[12 28 96 102 106
147 161 192 204 212 244 249 253 258 263 274 334
340 370 374 423 425 426 434 491 521 548 562 609
614 629 655 663 714 743 788 924 929 968 969 982
1031 1049 1069 1099 1106 1108 1199 1256 1269 1313 1320 1321
1323 1370 1391 1465 1501 1506 1520 1555 1628 1655 1661 1670
1701 1756 1758 1762 1799 1847 1852 1873 1888 1904 1915 1924
1950 1968 1973 1992 1994 2105 2287 2325 2347 2377 2378 2390
2432 2441 2480 2508 2512 2519 2520 2543 2544 2570 2589 2607
2609 2646 2647 2683 2723 2769 2774 2780 2796 2802 2833 2845
2869 2870 2876 2886 2920 3000 3017 3033 3038 3079 3094 3105
3147 3204 3249 3254 3313 3320 3341 3345 3346 3383 3393 3413
3416 3420 3458 3464 3466 3484 3522 3528 3533 3539 3557 3597
3709 3713 3723 3728 3732 3743 3820 3863 3868 3914 3919 3920
3945 4011 4013 4032 4078 4102 4107 4108 4128 4138 4146 4211
4234 4237 4242 4260 4328 4340 4385 4408 4415 4455 4457 4464
4466 4485 4527];[19 45 54 73 93 99 105 215 249
256 298 362 365 401 415 420 428 440 441 478 487
515 516 517 521 525 567 581 582 681 684 702 717
787 820 826 834 856 903 923 943 952 982 985 1014
1027 1029 1045 1064 1077 1115 1128 1131 1142 1149 1151 1176
1184 1190 1191 1226 1236 1247 1255 1262 1329 1372 1386 1408
1449 1456 1473 1481 1499 1538 1623 1684 1697 1719 1725 1740
1811 1832 1844 1845 1884 1926 1963 1964 1992 2001 2003 2009
2040 2041 2079 2090 2126 2129 2142 2145 2204 2205 2207 2220
2229 2253 2273 2312 2323 2326 2350 2374 2375 2398 2406 2412
2554 2583 2589 2590 2596 2597 2609 2612 2613 2681 2693 2778
2951 2953 3007 3022 3033 3034 3069 3081 3111 3128 3174 3181
3217 3218 3233 3299 3309 3337 3338];[7 15 79 114 151
157 165 170 190 219 233 305 314 342 367 418 437
459 477 486 491 500 502 516 542 553 573 574 596
667 695 708 721 786 796 808 819 838 876 890 897
923 975 976 1006 1010 1019 1033 1034 1050 1069 1114 1127
1142 1146 1172 1180 1185 1198 1255 1326 1343 1374 1392 1417
1446 1465 1548 1580 1584 1617 1627 1632 1645 1657 1737 1751
1760 1767 1774 1849 1851 1907 1927 1937 2021 2033 2038 2079
2096 2147 2152 2167 2247 2294 2343 2361 2366 2371 2400 2466
2511 2536 2539 2563 2572 2600 2628 2639 2640 2684 2696 2731
2741 2769 2810 2826 2848 2879 2884 2900 2906 2943 2948 2970
3002 3056 3062 3108 3127 3136 3173 3185 3223 3227 3233 3289
3298 3338 3340 3347 3363 3425 3457 3462 3497 3507 3523];[41
56 58 73 79 80 92 98 120 147 176 201 228
251 295 312 333 336 360 419 483 551 585 611 629
633 642 666 718 746 749 750 760 788 803 813 826
841 873 881 893 894 896 940 976 988 1023 1031 1045
1057 1082 1088 1126 1137 1179 1201 1202 1213 1235 1252 1283
1286 1315 1319 1332 1337 1349 1364 1442 1489 1545 1551 1572
1587 1593 1594 1609 1666 1694 1722 1726 1743 1816 1833 1838
1841 1876 1880 1939 1961 1986 1999 2001 2016 2066 2094 2129
2152 2173 2188 2191 2211 2232 2272 2321 2339 2401 2408 2413
2452 2468 2497 2524 2582 2586];[70 96 128 153 155 188
216 219 220 222 236 239 245 261 263 264 283 319
365 396 430 436 458 492 496 521 523 578 583 586
616 636 645 647 664 667 673 700 715 725 740 777
781 790 886 935 938 1002 1016 1074 1085 1086 1105 1111
1125 1160 1289 1361 1373 1432 1433 1459 1460 1521 1555 1573
1631 1694 1724 1745 1813 1836 1842 1847 1866 1901 1934 1945
1951 1953 1989 2005 2011 2023 2042 2100 2121 2321 2344 2363

2383 2392 2407 2412 2450 2456 2463 2469 2476 2487 2508 2544
2562 2571 2583 2612 2620 2627 2659 2698 2719 2839 2929 2940
2949 2996 3039 3057 3059 3096 3106 3175 3238 3271 3272 3290
3336 3343 3411 3455 3537 3560 3561 3588 3596 3646 3668 3677
3732 3787 3817 4025 4034 4058 4069 4103 4107 4178 4207 4268
4275 4282 4348 4368 4389 4461 4462 4466 4499 4568 4581 4662
4802 4835];[7 8 16 37 54 72 93 97 113 149
150 164 220 261 270 279 294 339 358 401 423 427
509 518 529 545 575 589 644 676 743 747 776 787
801 803 805 889 910 915 933 937 996 1024 1096 1105
1117 1137 1153 1168 1178 1205 1263 1274 1282 1344 1433 1444
1512 1518 1533 1539 1540 1593 1594 1661 1699 1707 1759 1777
1783 1805 1823 1833 1834 1837 1869 1887 1907 1908 1933 1959
2005 2007 2030 2139 2175 2187 2202 2206 2255 2256 2267 2340
2369 2383 2399 2413 2434 2480 2503 2513 2566 2573];[40 53
68 82 85 122 137 151 153 207 258 287 312 342
357 405 406 413 438 474 477 478 497 522 547 552
554 556 597 602 608 617 618 638 654 664 665 714
726 777 784 833 857 899 901 908 920 923 936 940
995 1003 1015 1024 1028 1047 1074 1092 1117 1143 1154 1159
1172 1176 1214 1235 1237 1238 1275 1322 1334 1357 1404 1417
1422 1437 1450 1521 1562 1592 1623 1659 1676 1742 1748 1774
1795 1798 1836 1854 1856 1859 1886 1912 1934 1938 1944 2032
2039 2047 2062 2164 2229 2309 2335 2348 2398 2403 2414 2421
2424 2434 2435 2436 2439 2443 2475 2478 2483 2534 2596 2614
2653 2667 2674 2691];[1 57 95 161 179 197 199 240
367 406 434 526 529 531 536 594 669 713 715 722
739 751 752 756 762 811 832 874 879 880 909 915
921 925 934 955 956 957 965 981 982 1056 1061 1100
1111 1155 1160 1163 1201 1248 1283 1303 1318 1319 1397 1398
1445 1446 1551 1592 1605 1606 1633 1637 1660 1689 1703 1707
1715 1717 1723 1779 1786 1803 1817 1824 1882 1947 1958 1964
1994 2033 2068 2115 2131 2203 2277 2284 2299 2381 2439 2463
2483 2524];[2 36 74 75 78 79 90 117 134 165
191 218 220 228 232 265 276 298 341 364 366 385
387 479 539 562 617 694 704 801 859 864 872 885
929 942 963 992 997 1031 1035 1037 1065 1066 1086 1169
1201 1293 1319 1320 1329 1389 1439 1465 1487 1509 1565 1574
1584 1596 1609 1624 1632 1660 1672 1703 1716 1775 1781 1802
1806 1837 1845 1848 1851 1862 1867 1875 1883 1906 1917];[1
5 20 35 55 71 85 194 275 310 319 322 383 384
392 433 434 443 457 468 471 495 558 590 596 623
641 658 671 678 698 724 834 870 899 919 927 941
967 1012 1014 1018 1191 1202 1270 1296 1312 1329 1344 1345
1375 1396 1400 1430 1456 1477 1640 1644 1694 1704 1724 1735
1758 1816 1833 1849 1880 1899 1929 1940 1972 2032 2111 2180
2181 2182 2184 2206 2246 2247 2248 2384 2404 2524 2537 2549
2575 2584 2586 2631 2632 2645 2651 2688 2697 2716 2805 2806
2816 2847 2853 2854 2892 2903 2925 2944 2949 2973 3021 3057
3059 3066 3068 3070 3088 3099 3112 3135 3140 3226 3237 3274
3282 3284 3302 3305 3306 3310 3311 3332 3354 3365 3381 3395
3419 3422 3437 3478 3516 3550 3558 3577 3581 3633 3637 3714
3720];[61 98 110 129 178 181 192 288 313 319 330
337 344 355 368 396 402 417 422 424 454 499 519
567 569 583 584 621 629 638 682 715 745 747 758
767 811 815 837 844 851 886 896 906 930 936 965
977 1013 1030 1060 1139 1164 1202 1214 1239 1261 1278 1281
1295 1301 1302 1314 1342 1363 1405 1420 1454 1461 1538 1549

```
1561  1563  1565  1566  1614  1634  1663  1664  1677  1706  1709  1711
1739  1740  1776  1790  1807  1813  1844  1902  1907  2012  2022  2034
2081  2105  2131  2171  2212  2312  2325  2338  2408  2412  2450  2453
2540  2568  2584];[6     8     50    71    96   156   168   199   256
282   308   334   366   371   411   428   433   438   458   496   504
514   521   577   579   581   602   622   742   798   810   840   874
930   943   970   973   980  1034  1066  1104  1128  1137  1138  1167
1172  1190  1223  1224  1250  1319  1322  1352  1396  1418  1426  1431
1432  1468  1514  1553  1591  1603  1607  1677  1687  1696  1700  1703
1761  1803  1805  1827  1855  1862  1899  1953  1982  2022  2028  2072
2086  2112  2135  2189  2228  2260  2265  2319  2330  2362  2407  2410
2412  2431  2498  2507  2512  2572  2577  2583  2602  2675  2689  2718
2792  2799  2819  2857  2876  2881  2891  2903  2905  2911  2962  2968
3005  3041  3044  3048  3056  3058  3081  3094  3121  3179  3195  3212
3240  3299  3313  3320  3331  3344  3365  3375  3390  3398  3399
3407];[1    12    36   113   143   186   195   230   255   271   292
295   319   346   374   385   422   434   452   454   463   508   534
545   565   573   662   745   837   856   864   871   877   882   896
913   919   947  1093  1107  1134  1145  1215  1230  1297  1328  1330
1342  1351  1430  1439  1514  1587  1608  1629  1643  1646  1673  1700
1728  1769  1777  1823  1828  1838  1839  1861  1866  1873  1894  1911
1941  1971  1980  1983  2000  2003  2009  2032  2034  2039  2100  2170
2180  2188  2223  2224  2227  2238  2311  2351  2352  2381  2386  2388
2392  2433  2461  2472  2522  2572  2604  2617  2666  2703  2719  2740
2762  2770  2786  2790  2824  2838  2863  2872  2876  2903  2946  2953
3014  3030  3068  3072  3082  3085  3089  3151  3153  3170  3262  3314
3316  3319  3330  3386  3404];[59    67    116   217   265   285   331
349   361   396   414   417   424   442   517   533   539   547   615
652   691   693   715   753   769   773   806   807   822   831   846
870   925   940   953   964   981   984   997  1018  1059  1061  1073
1115  1148  1269  1292  1296  1348  1371  1397  1463  1477  1499  1514
1579  1637  1639  1708  1711  1713  1802  1813  1874  1880  1911  1944
1951  2055  2058  2099  2124  2155  2156  2164  2172  2183  2210  2248
2275  2276  2323  2362  2420  2468  2512  2530  2543  2600  2642  2644
2656  2678  2700  2706  2715  2737  2738  2777  2803  2851  2853  2897
2909  2999  3001  3110  3124  3138  3200  3213  3215  3225  3245  3301
3320];[47    95   170   183   195   201   217   274   350   362   365
373   384   393   413   414   431   499   609   612   635   707   712
724   749   759   768   773   779   784   812   844   887   906   933
953   994   998  1001  1007  1008  1027  1087  1092  1131  1136  1140
1270  1362  1403  1410  1420  1479  1516  1520  1525  1531  1557  1561
1583  1599  1605  1614  1630  1651  1656  1679  1683  1694  1745  1746
1748  1929  1942  1954  1980  1985  2066  2083  2086  2105  2108  2117
2129  2141  2166  2194  2217  2218  2264  2274  2346  2371  2408  2427
2458  2481  2514  2536  2556  2578  2581  2590  2596  2629];};
% % SCORES=3;
% % ns={[21    77    78    80    88   106   131   167   172   194   249
262   336   350   420   434   484   613   625   647   661   662   698
713   716   718   835   853   894   923   936   969   970   995  1018
1021  1103  1143  1153  1157  1177  1178  1218  1233  1237  1251  1307
1324  1329  1340  1355  1376  1389  1409  1422  1439  1453  1469  1490
1493  1520  1585  1599  1600  1609  1640  1683  1701  1722  1786  1800
1851  1926  1951  1986  2027  2037  2060  2066  2127  2139  2158  2173
2174  2210  2232  2239  2302  2313  2314  2358  2422  2435  2436  2468
2469  2511  2528  2530  2567  2568  2571  2575  2594];[48    63    81
82    89    97   140   174   187   231   232   237   244   298   323
324   374   439   514   524   535   593   607   615   616   628   672
725   753   789   796   802   822   866   868   872   876   913   949
```

991 992 1051 1052 1056 1128 1135 1136 1176 1185 1186 1203
1277 1278 1365 1472 1482 1501 1509 1516 1519 1537 1547 1610
1670 1685 1705 1734 1736 1792 1819 1838 1860 1863 1871 1872
1881 1937 1940 1980 1983 2005 2064 2079 2089 2097 2104 2125
2246 2270 2284 2337 2356 2383 2409 2455 2456 2490 2512 2522
2545 2546 2562 2625 2840 2846 2854 2877 2917 2949 2984 2990
2998 3013 3063 3086 3128 3142 3164 3180 3181 3183 3192 3193
3243 3250 3333 3368 3370 3405 3425 3429 3468 3525 3545 3600
3601 3627 3672 3766 3780 3798 3836 3838 3898 3953 3956 4038
4092 4099 4182 4186 4217 4300 4320 4325 4332 4438 4444 4484
4535 4583 4603 4631 4649 4662 4681 4682 4718 4792 4825 4914
4919 4921 4940 4991 4995 5035 5049 5070 5207 5232 5278 5333
5336 5350 5390 5392 5395 5398 5425 5483 5624 5654 5655 5690
5701 5721 5727 5736 5758 5789 5798];[40 118 141 254 261
285 292 298 352 363 395 471 489 542 544 567 675
729 756 789 792 800 873 876 888 902 904 905 915
922 986 989 995 1020 1029 1056 1129 1143 1171 1260 1280
1283 1289 1290 1300 1307 1318 1320 1349 1350 1426 1459 1474
1476 1492 1500 1524 1533 1693 1797 1824 1857 1864 1896 1945
1961 2055 2138 2249 2259 2295 2330 2373 2482 2572 2574 2576
2620 2793 2794 2815 2828 2829 2836 2889 2921 2929 2933 2958
2960 2976 3000 3004 3068 3082 3086 3101 3116 3172 3176 3199
3228 3248 3282 3311 3312 3324 3325 3334 3342 3384 3388 3481
3491 3506 3513 3532 3555 3560 3619 3626 3630];[31 105 157
158 159 160 162 163 205 254 278 290 460 487 529
646 647 658 685 689 691 694 738 763 783 800 823
855 861 884 917 926 942 952 953 973 989 1041 1048
1078 1081 1117 1172 1258 1285 1288 1310 1338 1345 1371 1383
1405 1439 1558 1574 1615 1626 1656 1667 1684 1691 1699 1704
1710 1714 1731 1738 1751 1766 1798 1847 1849 1888 1938 2059
2061 2086 2094 2096 2178 2219 2304 2312 2387 2388 2389 2460
2462 2509 2528 2574 2613 2650 2666 2702 2705 2738 2755 2806
2919];[8 15 51 53 83 171 186 229 230 269 281
321 325 330 369 436 457 503 507 542 543 557 573
594 616 636 677 733 751 775 776 800 808 819 844
880 973 974 1041 1051 1052 1061 1065 1072 1073 1090 1102
1162 1172 1198 1230 1257 1302 1336 1341 1352 1368 1418 1422
1473 1493 1531 1539 1627 1706 1729 1732 1771 1781 1835 1837
1871 1887 1940 1956 1967 2001 2020 2088 2105 2122 2124 2193
2243 2255 2350 2442 2460 2476 2573 2587 2635 2675 2681 2687
2693 2804 2822 2825 2827 2848 2854 2921 2949 2955 2979 2997
2998 3029 3055 3067 3068 3075 3113 3125 3132 3151 3162 3166
3169 3186 3198 3230 3254 3263 3265 3325];[9 63 66 110
132 136 138 139 175 187 191 192 204 233 275 282
292 297 331 388 414 439 453 479 507 508 510 573
585 637 665 687 697 734 743 770 812 907 961 964
990 1010 1011 1015 1032 1033 1043 1079 1099 1117 1163 1208
1221 1237 1254 1261 1308 1370 1385 1397 1482 1501 1612 1632
1693 1733 1742 1754 1781 1799 1839 1867 1874 1878 1938 1956
1981 2027 2028 2079 2111 2141 2182 2193 2297 2450 2599 2613
2615 2664 2689 2730 2738 2763 2765 2769 2787 2804 2818 2821
2891 2898 2907 2913 2939 2955 2961 2976 2999 3014 3033 3041
3075 3081 3097 3113 3117 3160 3226 3240 3266 3302 3346 3353
3449 3507 3510 3516 3518 3554 3583 3606 3620 3705 3724];[28
41 71 102 105 106 138 143 159 163 192 195 223
224 249 250 252 256 258 260 279 283 292 300 370
412 420 423 498 516 521 535 548 562 575 580 586
600 609 623 714 759 777 796 817 826 834 852 888

891 894 968 970 986 996 1014 1039 1045 1069 1084 1105
1108 1124 1127 1147 1150 1152 1154 1198 1199 1251 1269 1316
1321 1376 1380 1393 1395 1438 1450 1500 1594 1606 1626 1628
1639 1643 1652 1668 1746 1756 1850 1851 1904 1905 1973 1990
2038 2245 2285 2310 2390 2404 2409 2420 2461 2470 2504 2512
2568 2590 2593 2613 2665 2683 2802 2824 2845 2923 2949 2960
2973 3079 3105 3126 3150 3213 3220 3244 3314 3320 3336 3359
3364 3416 3481 3499 3519 3539 3616 3653 3669 3682 3691 3709
3713 3742 3782 3848 3920 4007 4008 4013 4040 4044 4102 4129
4138 4181 4183 4187 4209 4212 4232 4234 4237 4244 4283 4408
4455 4612];[56 93 105 115 138 204 215 216 219 266
293 316 398 409 436 438 445 515 531 539 561 565
579 641 702 731 734 787 820 850 854 903 904 925
952 977 1015 1065 1077 1078 1140 1184 1191 1226 1236 1247
1294 1297 1298 1307 1329 1330 1372 1374 1395 1405 1481 1499
1572 1602 1652 1698 1752 1779 1800 1925 1926 1984 2041 2043
2073 2079 2126 2145 2195 2263 2281 2282 2289 2290 2359 2380
2387 2412 2468 2520 2529 2552 2569 2583 2613 2643 2652 2670
2701 2717 2730 2737 2761 2856 2900 2955 3003 3021 3034 3063
3069 3080 3111 3233 3311 3315 3326];[17 77 144 151 189
190 294 305 392 469 478 481 499 516 519 535 542
543 638 677 695 729 736 754 759 765 976 979 981
1031 1050 1051 1137 1146 1180 1247 1317 1325 1326 1328 1368
1374 1414 1421 1467 1478 1493 1496 1510 1515 1529 1533 1545
1579 1631 1696 1709 1710 1761 1785 1842 1850 1927 1934 1979
2003 2038 2069 2085 2128 2133 2155 2167 2217 2276 2298 2324
2357 2473 2503 2513 2564 2572 2600 2605 2639 2640 2663 2683
2695 2701 2726 2741 2744 2752 2755 2784 2798 2805 2875 2901
2906 2944 2970 2979 2981 3062 3108 3110 3144 3263 3283 3284
3286 3422 3423 3425 3451 3476 3518 3523 3528];[17 35 41
56 57 69 80 290 292 313 359 372 373 419 441
475 495 532 576 633 658 666 718 747 750 760 788
826 858 864 874 894 900 905 988 1003 1052 1057 1112
1126 1208 1213 1235 1315 1319 1337 1349 1379 1399 1442 1506
1525 1538 1587 1652 1657 1666 1674 1695 1726 1739 1781 1875
1880 1914 1937 1939 1945 1962 2001 2090 2110 2152 2158 2173
2238 2249 2305 2419 2473 2513 2569 2576 2582 2586 2587];[15
62 128 153 179 211 216 218 220 225 231 235 263
397 430 576 583 622 647 649 653 667 697 700 716
726 731 733 740 741 744 790 808 928 938 949 966
1002 1024 1044 1125 1131 1141 1207 1218 1232 1242 1310 1337
1358 1361 1413 1424 1431 1432 1434 1501 1555 1603 1628 1723
1724 1725 1730 1738 1766 1794 1802 1808 1813 1842 1844 1933
1953 1954 1989 2055 2066 2084 2085 2134 2196 2199 2240 2344
2363 2392 2485 2486 2503 2514 2539 2570 2583 2589 2602 2606
2611 2659 2698 2731 2758 2765 2778 2801 2839 2863 2888 2902
2921 2940 2941 2949 3019 3057 3096 3105 3134 3197 3238 3272
3319 3342 3367 3399 3404 3411 3455 3514 3560 3596 3680 3687
3689 3700 3744 3782 3800 3863 4035 4090 4093 4106 4181 4182
4203 4207 4231 4267 4275 4389 4462 4463 4466 4475 4502 4503
4516 4563 4574 4581 4583 4584 4705 4879];[8 16 22 34
38 54 97 128 164 168 184 220 269 280 358 366
401 415 438 440 509 545 550 588 656 743 761 784
787 805 851 889 910 937 955 1049 1061 1079 1118 1153
1196 1237 1282 1290 1303 1305 1345 1357 1372 1388 1402 1408
1431 1493 1518 1540 1593 1626 1632 1660 1696 1699 1725 1728
1771 1772 1817 1820 1833 1834 1850 1858 1866 1907 1915 1962
1997 2009 2051 2067 2068 2076 2110 2142 2176 2202 2203 2340

```
2365  2399  2428  2432  2440  2484  2565];[40   41   54   85   151
153   173   182   195   258   266   272   331   341   355   383   431
463   470   477   478   522   552   556   560   597   607   627   634
654   667   726   745   754   777   784   816   834   875   900   908
912   923   924   936   940   945  1018  1022  1024  1047  1048  1051
1098  1154  1172  1176  1199  1238  1247  1333  1334  1346  1356  1404
1413  1483  1526  1599  1646  1742  1752  1774  1775  1789  1798  1834
1840  1854  1859  1885  2017  2032  2065  2107  2134  2206  2220  2269
2287  2334  2335  2393  2403  2424  2430  2436  2439  2447  2534  2539
2591  2596  2640  2667  2684];[179   243   253   309   327   346   404
529   530   584   651   706   713   722   756   761   777   803   831
832   876   925   967   982   993  1100  1147  1156  1201  1221  1248
1297  1310  1312  1350  1397  1398  1445  1466  1505  1549  1551  1558
1583  1592  1603  1634  1672  1673  1689  1715  1717  1725  1765  1803
1817  1882  1889  1975  1993  2068  2093  2101  2131  2134  2164  2180
2193  2196  2211  2239  2254  2270  2277  2299  2341  2364  2381  2415
2424  2451  2472  2496];[36   42   77   179   196   217   320   341
428   540   543   544   559   562   575   578   581   599   617   660
683   707   754   755   795   808   820   825   864   872   895   905
938   948   993  1039  1066  1115  1228  1297  1389  1394  1444  1487
1502  1578  1591  1613  1653  1660  1667  1678  1679  1738  1787  1791
1801  1848  1875  1883  1909];[28   71   78   85   91   105   106
180   319   322   330   333   374   406   414   429   434   438   468
496   502   518   540   596   619   643   678   770   778   794   814
815   925   927   942   967  1149  1152  1191  1300  1304  1312  1345
1385  1400  1482  1498  1543  1640  1697  1704  1735  1747  1758  1776
1805  1849  1877  1894  1896  1957  2032  2111  2133  2180  2181  2203
2208  2224  2248  2310  2364  2383  2384  2387  2430  2444  2537  2545
2573  2574  2625  2632  2645  2654  2672  2673  2727  2799  2812  2847
2881  2892  2925  2949  2956  2957  3059  3068  3130  3135  3148  3182
3188  3213  3231  3302  3332  3364  3365  3367  3381  3392  3395  3404
3432  3454  3471  3503  3518  3535  3536  3550  3558  3577  3617  3637
3668  3673  3678  3694  3695  3699  3713  3741  3750];[83   105   120
132   174   192   208   221   270   284   312   319   339   424   431
499   502   519   568   569   592   611   671   682   745   747   815
835   867   886   936   961  1000  1013  1060  1067  1089  1106  1129
1169  1239  1261  1276  1277  1281  1289  1295  1370  1406  1420  1429
1445  1462  1489  1490  1509  1517  1527  1549  1565  1596  1634  1636
1663  1678  1706  1711  1751  1790  1799  1807  1844  1857  1882  1890
1902  1929  2001  2141  2186  2241  2257  2262  2305  2308  2338  2386
2408  2449  2450  2457  2458  2461  2464  2568  2570  2572  2583];[50
71   114   156   163   168   199   227   282   298   308   366   421
451   452   455   499   504   532   542   568   572   577   590   605
609   622   739   754   810   822   852   862   872   935  1032  1082
1090  1127  1159  1162  1172  1183  1201  1250  1255  1257  1298  1322
1363  1384  1402  1420  1459  1468  1510  1536  1540  1578  1588  1699
1700  1703  1761  1774  1781  1791  1796  1803  1822  1883  1953  2022
2057  2135  2192  2214  2234  2260  2270  2295  2365  2374  2377  2422
2434  2441  2517  2606  2608  2648  2689  2769  2778  2792  2799  2843
2865  2880  2886  2905  2910  2929  3007  3017  3037  3047  3058  3081
3091  3108  3116  3121  3128  3142  3178  3179  3181  3191  3194  3195
3299  3314  3348  3365  3390  3391];[27   34   35   76   78   107
113   117   135   141   143   178   179   186   195   217   256   264
292   417   418   430   451   530   545   565   574   608   659   668
688   694   697   698   728   770   819   880   885   930   985   994
1034  1037  1061  1093  1133  1135  1142  1145  1213  1275  1351  1357
1423  1468  1510  1629  1673  1674  1724  1728  1739  1824  1891  1911
1919  1971  2009  2020  2083  2126  2156  2159  2170  2194  2200  2206
```

```
2212  2311  2346  2351  2353  2381  2386  2411  2491  2521  2522  2572
2605  2640  2668  2762  2770  2803  2828  2876  2900  2920  2936  2982
3018  3024  3036  3070  3072  3085  3111  3151  3163  3249  3266  3289
3306  3316  3347  3402  3422];[32   141   169   181   237   285   294
355   361   396   405   414   421   442   452   547   574   592   609
652   693   753   846   870   892   946  1018  1019  1058  1069  1077
1128  1129  1130  1148  1199  1201  1204  1250  1269  1275  1371  1460
1472  1489  1503  1575  1596  1630  1647  1672  1721  1727  1759  1800
1875  1911  1920  1942  1958  1969  1979  1985  2031  2055  2056  2120
2124  2144  2159  2172  2237  2247  2251  2269  2473  2474  2512  2528
2577  2700  2718  2731  2803  2846  2851  2901  2918  2927  3022  3047
3052  3062  3093  3102  3138  3200  3213  3229  3268  3316  3320  3361
3363];[5    95   135   145   184   201   217   278   288   358   373
393   413   414   500   577   591   606   625   652   759   773   785
843   844   846   887   896   917   933   934   998  1001  1021  1027
1067  1086  1092  1154  1184  1202  1208  1262  1362  1374  1382  1401
1409  1458  1487  1516  1531  1536  1560  1561  1565  1566  1567  1570
1584  1591  1610  1612  1614  1656  1679  1705  1748  1871  2013  2023
2038  2083  2086  2102  2127  2130  2141  2142  2173  2192  2211  2218
2221  2264  2274  2283  2286  2309  2371  2465  2528  2536  2578  2590
2596  2600  2625  2629  2647  2661  2666];};

% SCORES=[13 14 12 9 3];
% ns={[420  2232  2594  2432  1340   167   792  1600  1311  2210  1222
2037   106  2090  1520  2530  1329   327  2528  1874   370   336  1038
1422  2358   716  1218  1307  1586  1722  2599  1215  2413  2366  2365
2014  2311   149  1609   923    22   329  1953  2225   768   625   353
1885  1903  2302  1507    83  1166   608  1726  1555  1358  1565  1177
2571   477    32  2082   216  1551  1645   573  1493   665  1200   842
1619  1887  2089  2060  1059  1768  1738  1213  2289   261  2204  1636
77  1041   698  2007   350  2527   791   712   765  1216  2129   552
82  1747  2531  2249  1701  1426    35  2140   287  1309  1873   996
1724  2008  1673   549    21  1355   354   262  2403   894  1439  2313
2575   970    80   342   555   249   484   969   985  2027   172   613
720  1269  1274  1730    88   718    78   713  1039  1178  1219  1786
1957  1103  2469   505  1143  1450  1657  2156  2230  2557    16    40
410   645  1021  1251  1683  2066  2511   131  1851  2127  2568   328
1237  1800  1951  1986  2158  2239   647  1599  1762  2278   667  1016
1299  1585  2567    14   171   214   315   409   467   502   550   707
723   918  1324  1353  1419  1833  1990  2176  2314  2472  2544   194
434   661   662   835   853   936   995  1018  1153  1157  1233  1376
1389  1409  1453  1469  1490  1640  1926  2139  2173  2174  2422  2435
2436  2468];
% [4320  5701  3425  2769   186  3838  3468  2735  4297   615  4684  2089
140  1819  2927  2330  1453  1726  1507  4656  2998   608   789  2079
3601  4221  2097  3437  2129  1931  2806  2337  5442  4071  2840  3181
859  5112  2356  2367  1065  1051    36  1620  4574  4061  1674  3844
296  3889  3525  3636  4169  2196  2439  3370  1249  2562  1113  1393
2250   740  4434  5255  5157  1277  1421   808  3254  3908  1991  1372
1495   616  2282  1871  3157  1052  1622  4533   439  2546  2863  4928
199  4862     1  4546  5107  3897  4538  4092  3161  4102  1064  5000
1409  5721  1858   298  4537  5512  4788  2246   245  1814   324   593
1205   802   764  2284  2877   523  4258  1788   640  3118  2590  1685
5690  5274   173  3659   866  3302   569  2774  4484  1314  5524   270
822  3002  4009  4598  4631   894  1926  4217  3174  3975  2511  5327
5748  2142  3244   524  1321    52  3085  4151  3182  3411  4901   863
4649  5217  3898  4069   540  1185  1664  3158  1948  5333  5488  5415
436  4991  5727  3702  2280  1879   612  1652  5160  1565  5477  2545
```

776   641  4999  1365   338  3450  5052   417  4908   486  2674   675
5647  1937  1908  4644   941  1049   753  1617  4712    68  5464  5564
1736  3635  3405  3956   513  4437   244  4494  1139  3381  5571  1024
4542  2841  4760  1519    42   846   648   430  2474  5677  5498  4691
1186  5678  1360  1537  4121  5349   514  4811  2858  4099  5425  3180
2104  4583  4325  1482  3836  5758  1128  5336  3849  1705  1983  4940
5751   867   913  2000  3893  4332  5207  4535  1866  2854  3291  4017
4438  4681  5392   796  1863  1980  4939  5099  1633  3086  5390    34
672   872  1838  2905  3128  4300  4995    82   607   949   951  2086
2409  4100   231   233   372  1136  2949  4662  5395   663   945  1262
1670  3166  3192  4184  4308  5398    89   862  1135  2005  2984  3243
3798  4444  4741  4792  4846   174   323   535  2163  2344  2369  2738
3142  3224  3333  3766  3878  4013  4175  4606  4724  5397  1203  4729
63  2383  3368  5483  5638   237  1881  2589  2794  3172  3206  4038
4235  4718  4919    31   187   200  1501  2387  2535  2773  3018  3063
3672  3965  4612  4682  4914  5278  5561  5703   288   337   341   428
563   734   950  1066  1253  1278  1326  1475  1559  1860  1862  1876
1942  1950  1959  2096  2143  2678  2788  2966  2986  3078  3179  3213
3309  3357  3497  3546  3555  3565  3600  3626  3815  3869  3907  4161
4185  4197  4315  4375  4418  4455  4717  4764  4867  4887  4980  5110
5134  5139  5235  5391  5431  5444  5744    48    81    97   232   374
628   725   868   876   991   992  1056  1176  1472  1509  1516  1547
1610  1734  1792  1872  1940  2064  2125  2270  2455  2456  2490  2512
2522  2625  2846  2917  2990  3013  3164  3183  3193  3250  3429  3545
3627  3780  3953  4182  4186  4603  4825  4921  5035  5049  5070  5232
5350  5624  5654  5655  5736  5789  5798];
% [904   729  2793   261  3481  1244  3093  3271  2056  1299  3199  2836
647  1474  1148  1020  3360  1492  2780  2697  1640  3101   876   260
888  3384  1056  1864  2249  1046   254   690  1627   792  2768  1491
2884  2620  2295  3031  2958  3072  3311  1426  1037  2820  3506  3235
366   789  3342   323   471  2921   507  1534  1693  2934  3577  1077
1805  1961  3216  3408  1047  1164  3513  1182  1689  3547  1759   676
160  1300  1024  1318  2202  2756   344  2484  2613  3502  3248  3472
1109   701  1193   193  1824  1287  1127  1289  3210   450   937  2536
2910  1842  3230  1670   790  1599    63  3386  2212  2509  3310  2436
3215  3393   408   826   429   494   663  2435  2124  2676  1354   157
648  3630  1963  2567   325  2680  1977  1460  2167  3457  2933   770
2576  2755  3453  1571  2829  2288  1776  2346  2610  2031   970  1198
3494  1823  2141   915  1163  1482  1519  3035   375   705    89  2363
1366   292  1350   756  3324   905  2572  3082   285  3491  3532   298
3172  3282   675  1349  1819  2055  3388   364   489  1019  3555   352
889   986  1262  3228  3619   363   559  3119   902   943  1106  2373
3116   238   859  1029  1129  1213  1678  1901  2409  2735  2889  3094
590   730   922  1307  2138  2209  2929  3176  3335    38   214   355
404   785   989  1084  2042  2193  2259  2574    40  1476  3068  1260
682  1500  1720  2708  2960  3191  3312  3597  3626    25   118   416
567   873  1290  1370  1609  1645  2696  3252  3414  3550  3560   142
181   315   337   400   405   453   817   823   840  1038  1175  1282
1615  1633  1897  2251  2393  2444  2457  2504  2866  3086  3090  3419
3425  3469  3512  3598   141   395   542   544   800   995  1143  1171
1280  1283  1320  1459  1524  1533  1797  1857  1896  1945  2330  2482
2794  2815  2828  2976  3000  3004  3325  3334];
% [2097   570  2059   823   160  1157  2650  2298  1265  1285   860  1094
1657  2806   436   962  1691  2631  2389  1172  1611  2163   673  1731
2041   541  2755  2666  1581    17    21  1182  1279  2893   252   672
2756  2235   736  2859   669  1444  1958  2233  2098  2570   158   738
1107   783  2583  2496  2894  1288  2569  2017   968   606   259  1272
1894  2702  1255   784  1128  2240  1824  1973  2095   689  2304  1961

1566 1701 952 1847 529 253 1635 2442 2229 2531 2292 1732
2460 441 1779 2686 2441 1557 569 1338 2660 2178 2526 2717
443 1137 2746 658 1924 1713 1055 832 350 358 1181 800
1458 1849 290 1615 1710 2061 861 2919 1310 1371 2086 855
1383 989 1117 2353 2738 216 1626 1699 2126 1390 317 1439
1888 568 647 685 884 942 553 706 763 973 1766 2509
218 869 1479 487 910 932 1011 1268 1825 465 520 1080
1405 1704 2387 2462 175 597 1345 1492 1765 1798 2345 2685
1048 1684 1949 2471 163 694 2400 2613 205 731 1667 1700
2705 2808 31 254 278 421 810 916 917 974 1041 1574
1610 1625 1714 2096 2177 2697 15 22 142 144 467 656
702 717 742 1071 1140 1163 1309 1348 1386 1437 1655 1815
1878 2010 2085 2100 2137 2234 2264 2284 2428 2429 2584 2624
2676 2701 2715 2805 105 157 159 162 460 646 691 926
953 1078 1081 1258 1558 1656 1738 1751 1938 2094 2219 2312
2388 2528 2574];
% [2476 2243 1940 1302 3184 281 1837 776 844 1336 800 839
904 2544 1012 1644 201 1678 727 3067 1967 1138 570 367
2822 321 544 2468 1515 1071 2350 3125 1887 2516 2635 2848
1072 2186 2343 783 3162 1939 419 1627 299 3068 2944 442
609 2001 347 2473 68 1301 675 1224 880 583 558 587
799 2336 109 573 2008 2340 2619 677 1665 341 186 417
1827 676 814 639 440 2535 1073 1289 330 1009 2035 1048
272 3283 712 1340 581 3055 171 2804 366 1646 2019 594
3132 1117 458 2789 1104 507 2536 1912 840 602 472 519
81 3113 943 3260 2361 2866 2417 3073 2265 1961 258 2364
1529 2432 489 2377 3166 1052 2949 775 1065 1835 2829 420
2124 369 1214 1569 819 2088 3254 541 543 1198 1871 2442
269 2997 1422 1740 3198 51 2255 2670 384 457 503 1043
1380 2695 2998 3230 2094 2622 2826 65 180 399 636 826
1868 3029 3186 3275 538 616 791 2012 2440 2827 3060 75
325 342 1010 1341 1364 1539 1595 1757 1956 3263 3265 3339
1771 1493 3169 808 1021 1172 2122 2821 2921 229 1102 1990
8 290 427 522 574 1041 1053 1257 1473 1706 1732 1781 2573
2681 3041 3289 40 47 142 243 274 445 487 508 509
722 763 838 914 1038 1054 1090 1163 1204 1265 1353 1408
1429 1456 1480 1513 1793 1805 1901 2024 2217 2219 2501 2561
2593 2667 2850 2980 3078 3107 3153 3181 3244 15 53 83
230 436 542 557 733 751 973 974 1051 1061 1162 1230
1352 1368 1418 1531 1729 2020 2105 2193 2460 2587 2675 2687
2693 2825 2854 2955 2979 3075 3151 3325];
% [3510 292 1754 743 2421 1385 2952 3033 1728 1836 3376 3121
963 770 1011 2787 2435 1055 3127 1015 1259 1099 1053 1853
282 961 2721 2602 98 585 1945 1124 2126 191 1430 3529
1115 2980 135 3450 2791 3671 1541 2006 962 267 192 3428
3462 2252 2613 2733 651 1299 2651 3093 3348 2472 2961 2764
1250 3239 2982 2362 281 256 2193 27 480 3539 1940 1734
275 1338 63 1135 2174 2886 908 1476 1184 134 3351 3448
186 1021 1213 52 414 2722 2497 456 1900 847 2036 3631
2792 3333 3084 1220 3346 180 1972 3005 3452 1464 3041 346
755 1453 1994 374 2258 3243 2908 1216 1946 3718 122 1242
921 3387 2444 2111 508 3374 2079 1466 3447 18 2294 439
2956 1241 1886 2328 3714 365 1868 2616 2580 1632 2907 3620
3705 510 1878 1208 2939 453 2251 2297 3014 1867 3006 1117
2765 3606 1501 2190 1221 1343 1482 2887 2093 3240 138 37
1254 2615 2891 2898 1079 1467 2916 3583 2027 2804 2821 3088
14 394 812 964 2660 132 1163 1431 1797 2322 2585 3540
3728 347 1799 2664 2763 2950 3076 139 178 313 525 723

```
1012  1781  1831  1851  1929  2337  3025  3113  3268    65   234   268
638  1211  1751  1965  2112  2387  2533  2689  2853  2976   331   507
1693   665  1308  3122  3392   297   721   734  1439  1956  1992  2738
2999  3302  3449    30   127   343   455   905   960   993   997  1009
1032  1047  1048  1077  1176  1186  1222  1498  1521  1742  1874  1883
1950  2050  2066  2140  2170  2284  2291  2319  2333  2465  2530  2559
2600  2643  2669  2695  2854  2899  2902  2973  2978  2989  3071  3094
3112  3158  3335  3378  3441  3629     9    66   110   136   175   187
204   233   388   479   573   637   687   697   907   990  1010  1033
1043  1237  1261  1370  1397  1612  1733  1839  1938  1981  2028  2141
2182  2450  2599  2730  2769  2818  2913  2955  3075  3081  3097  3117
3160  3226  3266  3353  3507  3516  3518  3554  3724];
% [4234  1756  1670   663   423   370   714  4455   609  1968  4408  4211
2796  3249   192  2512   655  3147  1762  2390  2432   548  4466  4138
3709   629  2377  3920   102  4260  2802  3767  2347  3863  4457  3105
4527  1873  4040  1904  2441  1269  1555  1701  1973  3713  4485  1370
2920    28  1099  1915  1313  4013  3528  4415  1924  1506   274  2544
2378  3033  2780  3743  3313  1199  3416  1069   788  3204  1321   106
3346  3919  1918  1994  2287  2407  1084  3000  2869   570  3413  4032
105  3341  3408  2872  4078   614  3923  3557  3539  2886  2520  4102
4328    40   876  3533  3254  1031  1985   791   968  1320  3464  1520
4177   982  3420  1108  3017   491  2876  3043  3728  1799  3597  3479
4128  3522  2427    96  4108  3296  1992  3782  1152  4107  4385   513
1589  3466  1852  4031  3320  1990  4146  2844   249   425  1617   147
244  1655  1661  2536  3868   740  2723  1450  4227   664  2570  4181
4188   743   543  2285   967   433  1228  3914  3732  3994   661  2790
3945   516  1592  2589  2745  1950  3890   204   521  3165   532  3723
3345  1501  2609  2845  3079  1154   300  1014  1150  4171  2808  3359
2096  2683   151   777  1628  2470  2538  4244  3364  4212  1517  4129
1908  2196  2297  3247  3519  4225  3908  4183  2156  2550  2590  4232
4237   569   796  1508  2504  2960  3213  3353   252  1006  1639  2038
2124  2428  2721  3126  3210  3314  3391  3669  4209   159   891  1438
1737  2830  2919  3481  3898  4007  4044  4300  4331  4587   283   361
1606  1608  1687  1769  2507  2684  2991  4548    71   239   934  1177
1252  1626  1718  1746  1887  2070  2420  2902  3616  4510  1851  2923
152  3848  1376  1380  2409  2518   826   970  1369  1393  1761  2084
3220  1127  1668  2310  2593  2824  2831  2997  3044  3336  4008  4334
258   575   719  1198  1247  1850  2261  2772  2949  3122  3750  3775
3810  3968  4139  4347  4350  4515  4518    12   161   212   253   263
334   340   374   426   434   562   924   929   969  1049  1106  1256
1323  1391  1465  1758  1847  1888  2105  2325  2480  2508  2519  2543
2607  2646  2647  2769  2774  2833  2870  3038  3094  3383  3393  3458
3484  3820  4011  4242  4340  4464    41   138   143   163   195   223
224   250   256   260   279   292   412   420   498   535   580   586
600   623   759   817   834   852   888   894   986   996  1039  1045
1105  1124  1147  1251  1316  1395  1500  1594  1643  1652  1905  2245
2404  2461  2568  2613  2665  2973  3150  3244  3499  3653  3682  3691
3742  4187  4283  4612];
% [820  1184  2613   478  1191  1226  2609  2412  1473  1499  3299   581
2079  2041  1408  2375  1845  1262  1372   903  3218   515  2003  1014
401  2126  2273  1236  2583   834  2597  2220  1884  3069  1740   487
567  1329  1992  2129  3034  1684  1064   215   436  2398  1697  1481
1115    45  2207   441   105  3181  2195  3111  3007   982  3337   521
1255  3233  2253   582   442  3033   681  2255  1719  2748  1077  2670
1538  2596   924  1152  1395   856  2229   428  3128    93  2013   850
298   702   384  2040  2406  2953  1926  1449  3326   525    19   365
420   362   550  1622   256  1190    54  2545  2374   516  3216  2230
440  2009   684  2312  1811  2323    73   137  1456   766  2692   892
```

```
2589  1128  1752  2142   617  2778  1450  2590  2350  2326   661  1921
469  2146  1728   987  3063   138  2761  3021   579  1078  2289   398
2282  1140  2380  1118  2717  2387  2569   438   731  2263  2290  1298
1535  2468  1758  3008   477  1056  1065  1572  2359  3314  1203  1925
2099  2961  1294  1426  1984  2633  3060   397   648   978  1681  2043
2056   216   409   925  1187  1448  2727  2955  3080   445  1405   266
1307  1374  1602  1779  1800  2423  2520  2652  2730   787   952  1020
2145  2414  2607  2737    99   249   415   517   717   826   923   943
985  1027  1029  1045  1131  1142  1149  1151  1176  1247  1386  1623
1725  1832  1844  1963  1964  2001  2090  2204  2205  2554  2612  2681
2693  2951  3022  3081  3174  3217  3309  3338    56   115   204   219
293   316   531   539   561   565   641   734   854   904   977  1015
1297  1330  1652  1698  2073  2281  2529  2552  2643  2701  2856  2900
3003  3311  3315];
% [2906   542   190   890   808   114   367   418  2096  2943  2640  2038
151  3108  2884  1580  1146  3289  1851   708  2970   502  2033   305
2848   897   721   695  1031  1374   219   975  3523  1927  1180   437
79   477   516  2563  3062   165  1996  1247  2741  1034   923  1645
2466  2572  2021  1617  2628  1344  1548    15   838  2539  2421  2639
1584  1149  1774  2900  1069  1934  3462  2600  1983  3136  2536  3274
3497  1223   233  1127  1142  2361  2247  2810  2512  2027   573  1446
3233  2826  2001  3340  1033  1467   667   320   903  1786   876  1417
2022  1632  3425   459   746  1696  2605   979  2990  1325  2701  3005
2787  2079  3363  1326   345  3094   441  1737  2294  3457  2796  2298
819  3173  1937  1272   314  2366  3454  2809  1504   553  1198  1849
3116  2684  2451  2732  1129  3156  3002  1751   480  3227   234   491
3283   543  1421  1137  1328   469   638  1567  3451   627   937  1752
2069  2752  3423  1637  2805  2944  3284   144   878  1540  1579  2743
519  1050  1478  2003  2755  2794  2744  3422   392   395   579  2133
49   909  1258  1589  2167  2276  2381  2510  3348  3518   123   736
933  2065  2604  2879   540   976  1038  1761  2334  2705  1710   481
3110  2513  3476    77   729  1471  1500  2128  2901  3302    80   277
358   499   677  1297  1533  1545  1979  2085  2695  3006  3288  3534
7   157   170   342   486   500   574   596   786   796  1006  1010  1019
1114  1172  1185  1255  1343  1392  1465  1627  1657  1760  1767  1907
2147  2152  2343  2371  2400  2511  2696  2731  2769  2948  3056  3127
3185  3223  3298  3338  3347  3507    17   189   294   478   535   754
759   765   981  1051  1317  1368  1414  1493  1496  1510  1515  1529
1631  1709  1785  1842  1850  2155  2217  2324  2357  2473  2503  2564
2663  2683  2726  2784  2798  2875  2979  2981  3144  3263  3286  3528];
% [1235   666   760  1587  1609    41   826  2066   251  1666  1880  1213
120  1319   336  1999  2129    80  1939  1023  1337    73  1694  2582
201  2001  1315  1961   633  1252   483   419  1838   551   718   893
2413   642  1594    58   894   988  1137   629   873  2586  2321    79
1726  1126  2016   976   360   813  2513   370  1031  2188  1744  1283
788  1082  1024  2152  2211  1179   611  1816   881  2232  1136  2089
576  1364  2334  1813   176  2524  1057  2497  2467    61  2468  2473
1743   333  1896  1125   945  1202   418  1572  1442   373  1985   147
2519  1317   585   631  2191  1045  2587  2419   948   359    57  2090
475  1112   905  1739    69   747   942  1814   658   820  1522  2355
292   621   663  1208  1406  1875   290  1276  1349   699  1363  1538
1695  2259   597  1101  1160  1589   354   378   440   750  1073  1469
1652  1677  1697  2158  2567  2576  1914   441  2238  2448  2494  2569
17  1650  2209  2324   864  1674  1962  2249    56    92    98   228
295   312   746   749   803   841   896   940  1088  1201  1286  1332
1489  1545  1551  1593  1722  1833  1841  1876  1986  2094  2173  2272
2339  2401  2408  2452    35   313   372   495   532   858   874   900
1003  1052  1379  1399  1506  1525  1657  1781  1937  1945  2110  2305];
```

```
% [1002  2929  1953  2392  2612  1074  2456   740  3560   645  3059  2940
2344  3588   938  4207  1361   430  4389  1373  1631  3238  1555   216
2996   667    70  2363   261  4581  2583  1573   188  1432  4034   155
4348  3985  2011  3732  1160  3272  4835  3596  3537  4462  3336  1433
4466  1813   219   283  2847  4058  2100  1842   935  4103  2659  4275
2450  4499  2698   153  4802   236  2519   458   436  2463  4178  4069
790  3668   521  4709  1866   726  4678   673   496   262  3175  1847
264  1768  3787   365  2042  1934    62   886  1989   647  2627  3646
1289  2719  2748  2508  1431  2521   583  2571  4483  3455  1694  3271
725   122  2476  2196  3311   863  3031  3450  2961   684  1524   263
2553  3796  4333    96   396  4229  3036  2138  2554   128   715  2321
324   235  1184  4577  1945  2412  4025  4568  3471   636  4662  1016
1851   641  4487  2114  1220  2839  2577  3134  1085  1105  2023  2383
2572  1303   700   492  4562  1460  4203   928  4516  3367  3782  1125
4182  1218  2801  3716   697  1424  3096  4339   179  1183  1232  2066
2570  2611  1954  2121  2765  3103  3342  3514  4231  4879   106  1501
1736  1789  3019  4563  4584    15   231  2084  2353  2778  2941  3900
4035  4246  1173  1738  2122  2334  2606  2731  3325  4330   105  2514
3513  3679  3689  4043  4329  4475   218   601   973   977  1189  1291
1933  2902  3057  3182  3197  3638  4368  4539  4783   435   586   646
1131  2013  2123  2171  2199  2257  2402  2459  2888  3680  3768  3976
4084  4208  4574  4582  4837   728  1026  1043  1341  2064  2134  2580
3899  4079  4090  1413   211   731  2503  2863   781   808  1310  1723
1724  1957  2870  3917  4187  4328  4503  3416  3556  3578  3789  4181
4355   220   222   239   245   319   523   578   616   664   777  1086
1111  1459  1521  1745  1836  1901  1951  2005  2407  2469  2487  2544
2562  2620  2949  3039  3106  3290  3343  3411  3561  3677  3817  4107
4268  4282  4461   225   397   576   622   649   653   716   733   741
744   949   966  1024  1044  1141  1207  1242  1337  1358  1434  1603
1628  1725  1730  1766  1794  1802  1808  1844  2055  2085  2240  2485
2486  2539  2589  2602  2758  2921  3105  3319  3399  3404  3687  3700
3744  3800  3863  4093  4106  4267  4463  4502  4583  4705];
% [1540   910  1518    54  1153  1263   261  2256  2399  2340   270  1593
164  1777  2503   937  2267   996     7  1661    16  1907   358   933
2187  2175   805  1533   915  1282   423  2480    93   509   656  1833
8  2007  1915  2566    37  1402   801  1887  2206  1512  1168  2076   386
1823   889   529  1756  1117  1869  2218  1699  2202  1805  2005  1959
589  1344  2030   950   113   307  1908  1274  1348  1444   787   242
2434  1724   536  2217   613     6   150  1118   149    34  1660   339
1096  1728  1433  1594  2068  2513   890   168   776  1557  1137  1388
1820   644   575  2326  1662   545   803  1834  2203    38  2565  1290
280  1493  1357    22   184   288    97   366   741  1161  1358  1890
588  1303  2440   795  1858  1949  2404   196   784   955  1583  1632
2075  1061   275   739  2294   328   415   742   859  1305  1538  1725
1866  1237  1628  1817  1839  2009  2308  2384  2532  1196  2365  1431
128  1049  1547  2121    74  1626   761  1772  1850  2067  2110  2176
2287    72   220   279   294   401   427   518   676   743   747  1024
1105  1178  1205  1539  1707  1759  1783  1837  1933  2139  2255  2369
2383  2413  2573   269   438   440   550   851  1079  1345  1372  1408
1696  1771  1962  1997  2051  2142  2428  2432  2484];
% [85  2403   777   477  1944  1450  2335   153   940  1143  2229  2596
2534   857  1172   617  1798   908  1235   899  1592  2667  1074  1154
1934  2653   547  2439  1748   597  1159  1176  1659  2435   312   552
497  1237  1534  2309    82   258  2032  1238  2691  1417   936  2614
1015   351   478   543   784   602  1742   923   726   665   556   995
1353   900  2039  2430   654   714   522  2348  2434  2436  2421    53
1118  2443   342   618  1024   653   631  1073   287   638  1641   151
341  1533  2674   438   568  1836  1003  1912   664   833   608  1623
```

```
1202   405   920   406  1859  1886  1938  1117   627  1795    54   924
1346   463   834   266  1646  2334  1334   816  1789  2134   945  1691
2070  1834   357  1483  2341  1972   605  1047  2555   560  1245  1285
1290  1478  1752  1913  2539  2591    40   106  1404  1750   458   607
1593  1702  2011  1561  1854  2447  2287  2640   173  1531   470  1048
1247  2107   172   634   960  1186  1356  1560  2017  2580  2684    68
122   137   207   413   474   554   901  1028  1092  1214  1275  1322
1357  1422  1437  1521  1562  1676  1774  1856  2047  2062  2164  2398
2414  2424  2475  2478  2483    41   182   195   272   331   355   383
431   667   745   754   875   912  1018  1022  1051  1098  1199  1333
1413  1526  1599  1775  1840  1885  2065  2206  2220  2269  2393];
% [179  2277  1318   756  1779  1061  1551  2068   957  1605  1803     1
669  2299   197  2439   832  2033   594  2131  1445   722  2115   930
713  1201  1947   762   982   925   531  2203  1319  1155  1398  1032
526   752  2463   880  1882  1660  2446   981   715  1397  2284  1889
2469   909   739  2381  1786  1111   529   921  1079  1765  2523   584
1100  1387   120  1717   874   199  1446   955  1633  1056  1348    73
536   422  1715   192   253  2483  1846   811  1822   956  1716   803
1303  2177  1750   965  1887   915  2524  2180   740  1634  2056  2445
2254  2415  1549   761  1297  1495   993   831  1505  1689    54  2239
113   309  1845  2195   404  1830   243   367  1680  1916  2136  2472
245   530   569  1156  1725   449  1099  1147  1203  1248  1671  1672
1848  2245  1427  1521  2193  2424  1817  2093   876  1583  2413   714
2023  2270  2340  2364  2496   327   736  1513  1673  1993  2348    57
95   161   240   406   434   751   879   934  1160  1163  1283  1592
1606  1637  1703  1707  1723  1824  1958  1964  1994   346   651   706
777   967  1221  1310  1312  1350  1466  1558  1603  1975  2101  2134
2164  2196  2211  2341  2451];
% [36   872  1660  1035   539   385  1574    75   694  1596   191  1293
963  1389  1487   562  1703  1775   859  1037  1738   366  1321   232
816  1632  1806  1320   864  1447   397   707   685    89   265   625
1609  1867   132  1201  1884  1781  1848  1307  1875  1400    78   165
1329  1319   341  1624  1883  1584  1672    90  1715  1086  1905  1232
79  1439   218  1169  1906   994   997  1837  1478  1614    37  1066
1430   825  1228   808   938   179   540   599    42   993  1556  1065
880  1729  1787  1790   412   754  1578  1591  1652  1490  1678   901
1297  1613  1653  1791   196   581   582   755   101   180   913   990
578   498   617   948   320   143   683  1444    29   297   473   895
937  1039  1424  1432  1595     2    74   117   134   220   228   276
298   364   387   479   704   801   885   929   942   992  1031  1465
1509  1565  1716  1802  1845  1851  1862  1917    77   217   428   543
544   559   575   660   795   820   905  1115  1394  1502  1667  1679
1801  1909];
% [3365   927  2032  3311  1640  2645  3558  1644  2180  3306  1012   495
678  2892  2949  2575  3637  3305  3135  1375  1735  2248   434  2090
3550  2697  3395   194  2537  3088  1899   671  3112  3237   967    71
2816   468  3099  3478   310   392  2111  1704  2247  2632  3577  3695
2847  2584  1498   700   433   596  2384  1296  2878   179  1400   636
2246  2651  1758  1345  1833  3066  3059   698   870  3403  2586  1886
1799    32  2549  2181  2631  2482  1152   404  3633  3422  2806  1972
504  1697  1396  3302  3284  3226  3097   623   899  3381  1202  3282
2688   663  3332     5  3442   641  1456   413  3720  1197   600    85
3140  3354  3057   383  2524  3310  2854   724  1850   443  1816   374
372  3694  2255  2944  2223   170  1776  1880  1014   384  1987   369
1477  3050  1018    35  3273   941  1760   169  1929  3581  1564  3068
2571  2805  3437   112   258   322  2276  2573  2574  1312  3182  1149
2727   106   429  1482  3364  3536  1877  3432   540  1385   330  3041
3188   319  1821   226   321  2526  2711  3638  3673   105   180   333
```

```
643  1304  1896  2625  3380  3713  3741  1464  2081  1849  1299  3326
3367  3535   403   421   739   812  1182  1747  1805  1957  2383  2956
3271  3471  3560  3626  3703   972  1029  1084  1543  1783  2667  3230
3602  3699   814   925  3416  3503   815  2672  3213   496  1744  1871
2182  3080  3347  3719  3748  3750   406   619   770  1191  1212  1509
1621  2091  2101  2208  2376  2654  2957  3003  3130  3148  3398  3518
3668  3678     1    20    55   275   457   471   558   590   658   834
919  1270  1329  1344  1430  1694  1724  1940  2184  2206  2404  2716
2853  2903  2925  2973  3021  3070  3274  3419  3516  3714    28    78
91   414   438   502   518   778   794   942  1300  1894  2133  2203
2224  2310  2364  2387  2430  2444  2545  2673  2799  2812  2881  3231
3392  3404  3454  3617];
% [192   424  1420  1706  2408   402  1261   745   844   499  1740  1663
1164   815  1342  1030  1844   896   936  1302  1902  1295  1711   837
2131   811   715  1677  1565  1709   621  2312  1664   337    98  1060
1301   417   584  2584  1278   313  1634  1538   355  1561   565  2081
1549   396  1202   133  2012  1807  1417  1013  1614  2338  2568   569
2241  1139   767  1461  1790   758  1454   629  1413   178   244   682
1776  2493   721  2326    73   453   330  1281   747   129  1739  1184
2084   886  1550   255   441  2022   368   319   930  1214  1901    64
1563   583   568  1277  2001  2583  1089  1857   519  1406   105  1289
2262  2457  2570   751  1129  1405  1695   906  1596   835  1169  1936
2141  2223  2305   120   502  2129   174   208   298   671    52   114
132   422   698  1517  1592    83   454   833   867   978  1462  1502
1884  1954  1996  2308  2389   221   961  2572   592  1000  2450   312
431  1929  1947  2464   270  1168  1239  1882  2186   803  1678  2386
2407  2449    61   110   181   288   344   567   638   851   965   977
1314  1363  1566  1813  1907  2034  2105  2171  2212  2325  2412  2453
2540   284   339   611  1067  1106  1276  1370  1429  1445  1489  1490
1509  1527  1636  1751  1799  1890  2257  2458  2461];
% [2905   168   366  3179   308  3365     6   504     8    50  1352  3081
1250   581  2602  2689  2857   199    71  1591   742  1703   428  2319
2189  1700  3117  1803  2412  3121  2819  1855  1034  1418   282  2135
930  3299  3044   217  2902    96   156  3344  1224  2675  1696  1953
1223  2572  1396   602   874  2929  2993  1431   798  2792  1862  1138
2072  1190  3195  1540  1804  1183   622  2431  1272  1322  2968  1167
2022  3056  1104  1982  2903  2362  3399  3320  1137  3282  2086  1279
1468   762  2112   338  1687  3261  3398  2486  2879  1172   514   122
458  3386   256  3005  1761   452  1607  1298  3390  2498  3375  2718
1686   579  2191  1827  2452  1301   521   822  2507  2654  1432   973
3240   606  3094  2330  3108   314  2185   852   810   840  2891   787
2843  1699  3314   508  1588  2063  2411  2799   872  1512  1531  3058
236  1532  3353  1255  1975  1791    41   337   451   694  1399  2144
2592  2648  3047  2364  2443  2908  3178  3243   421  1154  2349  2583
3181   739  1628  2884  2886  3408   216   499   639   935   976  1205
2539  2632   163   184   641   983  1049  1201  1774  1905  1999  2071
2548  2734  2769  2881  3441  2365  2865  3007  1159  2517  3191   862
1203  1257  1510  2234  2260  2374  2606  2910   114   532   572  1032
1578  1590  1629  2270  3017  3177   212   227   542  1462  1822  1911
2130  3116  3391   334   371   411   433   438   496   577   943   970
980  1066  1128  1319  1426  1514  1553  1603  1677  1805  1899  2028
2228  2265  2407  2410  2512  2577  2876  2911  2962  3041  3048  3212
3313  3331  3407   298   455   568   590   605   609   754  1082  1090
1127  1162  1363  1384  1402  1420  1459  1536  1781  1796  1883  2057
2192  2214  2295  2377  2422  2434  2441  2608  2778  2880  3037  3091
3128  3142  3194  3348];
% [2009   545   454  2351   745  2824  2703  1608  3316  2790   422    12
2170  2572  1673  3151  2034  2876   113   565  1728  1838  3404  1823
```

```
1297   452  1700  1093  1328   501  3170  1430   896   837   186  2311
1769  2666   840   255   195  1866  3153   107  1828   508  2863  2770
346   143  1632  2188  1916   374  1135  2461  1351  3289  2238  2180
2223  1873    93   282  1832  2551  2472   871    34   573   303  3314
2388  1365  2352  1342  1107  1122  2100   693  3386  2643  2242   730
856  1273   773  1271  1861  1971  3072  1646  2642  2433  2150   520
319  1330  2200  3319  2838  2392   877  1475  3036  2087  2932  1983
1663   696  3286   678   295  1894  2522  2179  1306    78  1397  2032
1941  3085  2719  1900   463  3265  2357   522   483   385  2000  1674
2381   141  2020   768  2803   819  1724  1919  2982   384  2521  3111
291   668   744   770   850  2455   289   534  2605  2668  3402   217
127   430   568  1697    32   451   530  1041  1151  1688  1726  2387
2843  3306    20   985  1287  1300  1792  2491  2820  1047  1265  1321
1447  1715  3073   450   728  1016  1145  1222  1275  1375  2637  2848
3141  3359  2828   608   256   659  2922  1037   264   574   731  1026
1034  1067  1213  1423  1510  2002  2120  2126  2156    27   178   181
457   599   885   924  1142  1266  1357  2159  2353  2386  2640  3211
3249  3422     1    36   230   271   292   434   662   864   882   913
919   947  1134  1215  1230  1439  1514  1587  1629  1643  1777  1839
1911  1980  2003  2039  2224  2227  2604  2617  2740  2762  2786  2872
2903  2946  2953  3014  3030  3068  3082  3089  3262  3330    35    76
117   135   179   417   418   688   694   697   698   880   930   994
1061  1133  1468  1739  1824  1891  2083  2194  2206  2212  2346  2411
2900  2920  2936  3018  3024  3070  3163  3266  3347];
% [2851  1148   870  2530  3001  3213  1061  1802  1911  2909   331  1463
2362  3245   693   773   691   349  2155   424  2124   285  1815  2738
2656  2642   116  2055   997   807   547  2058  2512  1073  1880  1348
2164  2897   333  2803  1708  3124  3062  1147  2543  3127    59  3225
1115  1874   925  1637  2802  3002  1639  1579  1489   940  1733  1499
1477    67  1018   894  2247   437  2545  2183   568  3215   838  3138
2474  2099  3200   769  1944   999  2248   806  2323   265   715  1314
31   592  2077  2428  3301  3110   442  2367   770  1218   602   643
539  1730  2270  1713  1895  2644  1761   533  1296  1987   846  2927
414  1250  1800  2031   294  1371  3320  3316  1130  3363  1538    55
1275  2718   260   355  1129  1942  3317  1647  1727  2731   488   705
987   259   417  1612  2159  1340  1630  1958  2730  3183  3268    32
421   505  2775  2869  2967  3072    82  1472  1672  2033  2199  2797
2918   831  1460  2120  2237  2301  3123  3265  3047  1979  1201  3077
396  1985  2528   169   452   893   958  1035  1665  1853  2269  3093
217   361   517   615   652   753   822   953   964   981   984  1059
1269  1292  1397  1514  1711  1813  1951  2156  2172  2210  2275  2276
2420  2468  2600  2678  2700  2706  2715  2737  2777  2853  2999   141
181   237   405   574   609   892   946  1019  1058  1069  1077  1128
1199  1204  1503  1575  1596  1721  1759  1875  1920  1969  2056  2144
2251  2473  2577  2846  2901  3022  3052  3102  3229  3361];
% [1614   773  1651    95   759   431  2141   183  1001   201  2083   373
933  1531   170  1362  1656  1683  1748   414  1087  1980  1942  2408
2217  2108   784   413  1745  1516  2514   855  2264   812   887  2629
2086  2129  1520  1694   274   217   844  1321  2117   499  2465   577
2458   906  1361   384   249  1403  2481  1583   562  2173  2061  2130
2371  1630  2581   953  1592   365  1994  2427    62  1886  1695  1123
1006  2313  2596  1599  1317  1131  2650   181   766  2066  1420   424
1682  2166  2220  2222   318  2295  2349  1591  2218  2023  2600  1570
2286  1184   652  2346  1262  2536  2283   288  2647   393  1566  1612
2590   846   998  1154  1567  2309   331  2244  2578   358   816    37
761   934  1027  1226  1536  1738  2523  2649  1346  1382  2192   187
272  1097  1104  1255  1351  1401  1487  1491  2213  1515  2639   625
896  1021  1270  1360  2338  2367  2634  1565  2102   917  2528  2646
```

```
145   591   785  1067  1853  2112  2211  2347  2549  2652    47   195
350   362   609   612   635   707   712   724   749   768   779   994
1007  1008  1092  1136  1140  1410  1479  1525  1557  1561  1605  1679
1746  1929  1954  1985  2105  2194  2274  2556     5   135   184   278
500   606   843  1086  1202  1208  1374  1409  1458  1560  1584  1610
1705  1871  2013  2038  2127  2142  2221  2625  2661  2666];};
% SCORES=8;
% ns={[14    15    21    22    74    82   120   149   167   171   216
287   290   327   328   329   336   357   370   418   420   467   502
550   552   573   608   610   665   698   712   716   724   766   768
792   803   918   969   979   992   996  1016  1038  1050  1059  1127
1143  1166  1200  1213  1215  1219  1222  1309  1324  1329  1333  1340
1358  1422  1426  1464  1493  1507  1520  1536  1551  1555  1586  1600
1609  1619  1645  1657  1708  1722  1726  1747  1811  1859  1874  1887
1895  1903  1953  1978  1990  2003  2037  2042  2090  2091  2129  2130
2139  2140  2176  2210  2225  2232  2314  2358  2365  2403  2413  2430
2432  2528  2571  2594];[8    31    36    42    43   108   173   186
199   237   245   287   289   296   298   338   341   371   430   486
523   524   535   563   579   590   608   615   672   740   776   789
796   802   866   941  1049  1051  1065  1075  1102  1111  1113  1161
1185  1205  1253  1274  1278  1311  1360  1365  1372  1421  1453  1481
1495  1507  1517  1521  1565  1567  1571  1578  1601  1610  1611  1622
1652  1683  1705  1759  1788  1814  1819  1860  1862  1876  1920  1926
1931  1977  1987  2079  2089  2096  2142  2246  2250  2274  2280  2284
2330  2337  2356  2431  2440  2456  2498  2543  2546  2674  2735  2769
2806  2813  2840  2858  2863  2909  2911  2919  2927  2960  2966  2973
2998  3002  3018  3063  3078  3118  3157  3158  3161  3179  3181  3182
3213  3215  3248  3254  3287  3291  3296  3302  3321  3335  3425  3468
3499  3549  3555  3565  3600  3626  3636  3659  3780  3838  3844  3878
3897  4013  4054  4061  4069  4071  4102  4161  4169  4185  4186  4217
4221  4297  4315  4320  4418  4423  4434  4455  4477  4494  4533  4537
4542  4546  4550  4558  4598  4604  4654  4656  4681  4684  4691  4712
4788  4799  4805  4811  4862  4901  4991  4999  5052  5068  5107  5110
5122  5134  5139  5157  5160  5217  5255  5318  5388  5397  5415  5444
5512  5524  5571  5608  5647  5677  5701  5751];[25    39    63    78
85   130   142   157   160   214   247   260   261   280   323   325
337   355   366   376   406   408   450   453   471   494   572   577
589   647   663   690   705   718   729   756   785   792   817   823
826   876   902   904   939   959  1020  1024  1037  1038  1046  1083
1084  1148  1152  1164  1198  1282  1287  1299  1300  1335  1354  1369
1426  1460  1474  1482  1492  1534  1611  1615  1631  1633  1640  1642
1645  1670  1689  1710  1755  1805  1823  1864  1897  1961  1963  2031
2042  2051  2056  2124  2141  2193  2201  2202  2211  2212  2232  2249
2251  2288  2295  2331  2428  2444  2491  2504  2567  2613  2620  2640
2755  2761  2768  2780  2786  2793  2836  2866  2884  2910  2921  2925
2933  2934  2958  2960  3000  3031  3035  3072  3090  3093  3101  3118
3199  3204  3215  3228  3311  3324  3342  3360  3384  3393  3404  3408
3419  3425  3454  3457  3469  3472  3481  3497  3502  3506  3598  3630
3632];[17    19    28    80   138   142   144   158   160   181   251
252   333   384   430   436   437   461   465   467   524   529   541
569   570   606   656   670   672   673   689   702   717   742   760
783   823   832   858   875   962  1055  1066  1094  1140  1172  1181
1182  1265  1272  1285  1288  1377  1386  1437  1458  1557  1611  1625
1630  1701  1710  1732  1766  1801  1894  1949  1958  1973  2017  2041
2059  2085  2086  2097  2111  2172  2229  2234  2246  2253  2264  2298
2389  2428  2429  2442  2496  2527  2528  2531  2569  2570  2583  2584
2631  2645  2650  2666  2701  2702  2715  2746  2755  2805  2859];[4
47    65    68    80   109   154   172   180   200   201   224   239
```

```
264   272   274   281   321   330   347   366   367   369   374   380
419   445   458   472   507   521   544   573   581   583   594   609
639   677   722   727   776   799   800   817   839   840   844   856
880   886   904   968   977   991  1009  1012  1052  1054  1065  1090
1101  1147  1175  1301  1302  1336  1353  1422  1429  1456  1480  1513
1515  1627  1639  1644  1665  1666  1669  1678  1687  1726  1805  1827
1837  1850  1865  1880  1887  1912  1940  1967  2001  2002  2019  2035
2124  2141  2219  2243  2334  2340  2350  2468  2473  2476  2494  2544
2561  2572  2573  2619  2635  2666  2667  2674  2755  2789  2804  2822
2848  2850  2866  2980  3055  3067  3073  3078  3107  3113  3125  3162
3170  3179  3184  3235  3254];[18    30    31    37    52    63    65
126   127   130   134   135   136   154   177   186   191   282   292
332   343   374   508   526   636   651   743   847   905   960   961
963   990   993  1009  1011  1015  1021  1032  1037  1041  1048  1053
1079  1115  1134  1160  1176  1177  1183  1186  1216  1218  1242  1250
1259  1299  1430  1464  1521  1567  1644  1728  1742  1754  1798  1799
1853  1874  1883  1886  1896  1900  1925  1940  1950  2006  2062  2066
2079  2083  2125  2126  2145  2170  2174  2252  2258  2284  2294  2319
2321  2333  2357  2365  2404  2435  2472  2530  2615  2651  2695  2721
2722  2886  2898  2899  2939  2952  2973  2978  2980  2989  3022  3033
3074  3084  3112  3121  3127  3137  3158  3240  3243  3335  3376  3378
3386  3391  3432  3441  3510  3529  3629];[12    28    40    72   105
147   192   203   212   227   249   251   263   279   296   340   370
392   423   425   426   434   491   521   532   548   562   609   629
655   663   714   740   743   759   791   852   876   896   924   963
966   969   978  1010  1019  1031  1099  1108  1199  1247  1256  1269
1313  1315  1321  1323  1370  1421  1506  1519  1520  1555  1565  1628
1644  1661  1670  1698  1749  1756  1758  1761  1762  1852  1860  1873
1888  1909  1950  1968  1973  2105  2118  2193  2209  2249  2347  2377
2390  2418  2432  2441  2446  2482  2495  2519  2520  2543  2544  2570
2607  2609  2644  2646  2647  2774  2780  2786  2802  2845  2869  2872
2949  2988  3000  3017  3033  3038  3079  3094  3105  3147  3249  3254
3287  3300  3313  3326  3345  3347  3383  3384  3408  3416  3420  3437
3449  3458  3464  3484  3489  3522  3597  3709  3728  3743  3759  3762
3787  3863  3868  3919  3920  3945  4023  4069  4078  4104  4107  4108
4138  4177  4211  4227  4233  4234  4242  4260  4320  4328  4340  4408
4415  4455  4457  4464  4485  4511  4527];[19    53    54    93   174
215   249   297   298   322   338   362   365   398   399   401   429
441   467   487   515   516   519   521   567   581   582   681   778
820   850   856   862   903   919   922   923   925   943   948   952
977   980   982   984  1014  1029  1051  1064  1077  1142  1149  1151
1184  1190  1191  1226  1236  1255  1262  1329  1372  1408  1416  1426
1448  1450  1456  1473  1499  1572  1611  1623  1677  1684  1685  1697
1719  1725  1801  1811  1832  1844  1845  1926  1928  1964  1988  1991
2003  2009  2041  2079  2090  2142  2195  2205  2207  2229  2265  2273
2294  2312  2323  2326  2350  2374  2375  2412  2433  2554  2583  2589
2590  2596  2597  2609  2612  2613  2670  2681  2951  2952  2953  3007
3022  3033  3034  3069  3081  3111  3128  3181  3190  3217  3218  3233
3299  3326  3337  3338  3347];[15    17    29    79   141   151   165
190   233   298   305   306   314   342   367   437   459   477   486
496   500   502   516   542   553   566   573   574   632   701   708
721   808   819   838   890   897   907   923   975   976   981  1010
1019  1034  1050  1061  1069  1142  1146  1172  1180  1247  1318  1343
1344  1374  1392  1415  1548  1567  1580  1584  1614  1617  1627  1632
1645  1737  1753  1849  1851  1863  1893  1927  1937  1981  2001  2021
2027  2033  2038  2079  2096  2147  2167  2230  2244  2247  2248  2255
2275  2294  2324  2361  2381  2400  2421  2484  2539  2545  2563  2600
2660  2684  2696  2697  2700  2704  2741  2769  2810  2848  2879  2884
```

```
2891  2906  2943  2970  2973  2982  3002  3056  3062  3108  3116  3127
3136  3143  3173  3185  3227  3284  3289  3338  3340  3363  3372  3457
3497  3507  3523  3524];[15    41    58    73    79    80    92   120
147   176   228   251   276   286   312   333   336   354   357   360
418   532   559   576   611   621   632   633   642   647   665   666
700   723   729   746   750   760   788   803   813   826   873   881
893   894   896   940   976   988  1023  1045  1057  1058  1082  1098
1124  1125  1179  1191  1201  1202  1213  1235  1238  1246  1286  1319
1332  1336  1337  1342  1353  1442  1545  1551  1587  1590  1609  1666
1694  1722  1743  1816  1925  1939  1961  1999  2049  2066  2094  2129
2191  2211  2272  2321  2373  2401  2413  2497  2524  2537  2582
2587];[15    62    70   128   153   155   206   216   219   245   261
346   365   396   492   496   521   583   586   616   636   645   664
700   715   725   740   769   831   886   937   938   949  1002  1014
1016  1048  1074  1082  1085  1105  1160  1197  1238  1245  1302  1303
1310  1336  1361  1373  1376  1433  1435  1460  1521  1552  1573  1596
1624  1631  1681  1693  1694  1724  1766  1768  1783  1813  1842  1934
1935  1945  1953  1976  1989  2011  2042  2063  2068  2100  2105  2121
2285  2321  2344  2363  2392  2407  2412  2426  2450  2456  2463  2469
2476  2487  2519  2544  2560  2570  2576  2583  2612  2627  2659  2662
2698  2714  2719  2732  2748  2857  2929  2949  2996  3019  3057  3058
3059  3069  3087  3096  3134  3143  3167  3238  3290  3294  3336  3343
3368  3374  3411  3434  3450  3537  3555  3560  3561  3588  3589  3596
3615  3644  3646  3668  3677  3732  3899  3985  4025  4034  4035  4036
4103  4140  4182  4207  4229  4231  4268  4275  4282  4348  4371  4389
4461  4462  4483  4499  4577  4581  4636  4662  4688  4801  4802  4835
4858];[7     8    16    37    54    93   120   150   169   270   278
293   331   339   358   386   401   423   427   459   505   509   518
529   575   589   644   676   743   747   776   801   803   805   878
889   910   915   933   937   950   969   996  1083  1153  1178  1205
1226  1263  1274  1328  1405  1512  1518  1533  1539  1540  1593  1617
1656  1659  1661  1699  1707  1759  1777  1805  1823  1834  1837  1850
1851  1858  1866  1887  1907  1908  1959  1980  2005  2007  2030  2044
2139  2154  2170  2175  2187  2204  2255  2267  2302  2340  2369  2383
2413  2480  2503  2513  2573  2574];[40    53    68    82    85   122
137   153   258   287   291   312   350   357   366   406   446   474
477   478   486   487   497   530   547   552   554   602   608   617
618   638   665   714   726   753   771   777   786   833   857   884
887   899   908   940  1015  1016  1018  1024  1026  1028  1047  1074
1092  1106  1117  1172  1237  1275  1322  1334  1349  1357  1383  1404
1422  1450  1476  1623  1659  1676  1702  1742  1756  1836  1854  1856
1857  1858  1887  1934  1944  1951  2032  2039  2056  2062  2066  2079
2089  2164  2253  2310  2335  2348  2389  2398  2403  2422  2434  2435
2436  2439  2442  2443  2444  2451  2475  2534  2538  2596  2614
2674];[1    40    59    95   120   179   187   192   197   199   253
359   367   406   443   506   526   536   584   669   713   715   722
739   740   751   756   762   832   915   921   939   956   957   981
982   994  1018  1056  1061  1079  1099  1111  1160  1163  1201  1248
1317  1318  1338  1360  1398  1445  1446  1450  1551  1592  1605  1606
1633  1637  1660  1689  1715  1765  1779  1791  1803  1824  1869  1882
1914  1928  1947  1968  1983  1994  2033  2068  2115  2131  2167  2175
2186  2189  2200  2277  2278  2284  2299  2343  2364  2386  2397  2414
2416  2446  2448  2456  2483  2500];[2     3     5    36    74    75
78    79    90   134   191   218   265   276   298   341   373   379
385   479   517   545   551   562   614   617   622   624   694   704
736   762   777   802   808   836   859   864   872   880   895   901
942   963   994  1035  1037  1065  1086  1293  1319  1329  1411  1429
1439  1476  1517  1521  1565  1574  1584  1609  1614  1624  1632  1640
```

1660 1672 1704 1707 1775 1781 1806 1835 1837 1845 1848 1851
1852 1883 1906];[1 35 71 85 120 176 179 180 194
199 246 310 322 364 383 384 392 433 434 443 457
468 475 495 517 536 558 585 589 596 623 641 658
671 678 724 870 899 902 919 927 957 967 1014 1020
1202 1296 1329 1345 1375 1391 1396 1430 1454 1456 1477 1530
1590 1640 1644 1694 1697 1704 1735 1758 1816 1849 1899 1929
1940 1964 1971 1972 2032 2111 2180 2184 2206 2246 2248 2313
2321 2359 2376 2385 2391 2394 2430 2511 2524 2537 2570 2575
2586 2632 2645 2651 2678 2688 2697 2716 2739 2805 2816 2840
2847 2853 2892 2899 2901 2903 2925 2937 2944 2945 2949 2958
2973 3021 3057 3059 3066 3070 3088 3135 3188 3226 3274 3282
3302 3305 3306 3310 3311 3329 3332 3348 3365 3381 3395 3419
3478 3515 3550 3558 3560 3577 3633 3637 3659 3694 3713
3714];[52 61 98 110 129 136 143 178 181 192 288
313 319 339 344 355 368 396 402 417 424 441 499
519 567 569 577 583 584 629 682 715 745 758 767
811 815 828 837 844 851 886 896 906 930 936 1019
1060 1163 1164 1214 1239 1255 1261 1278 1281 1295 1301 1302
1320 1342 1363 1420 1442 1454 1461 1465 1549 1561 1563 1565
1566 1592 1614 1634 1636 1645 1663 1664 1677 1706 1709 1711
1740 1757 1796 1804 1813 1844 1889 1890 1901 1903 1907 1969
1970 2002 2012 2034 2081 2131 2178 2212 2308 2312 2326 2338
2408 2540 2568 2584 2588];[5 6 17 50 71 100 150
156 168 199 200 218 282 298 308 317 334 366 411
428 433 438 504 514 521 577 581 590 602 606 691
713 753 798 810 823 842 852 930 936 943 973 980
1066 1128 1137 1167 1172 1183 1187 1223 1224 1272 1279 1301
1319 1322 1352 1396 1418 1426 1431 1498 1514 1540 1591 1603
1607 1687 1696 1703 1761 1803 1805 1855 1890 1899 2022 2028
2063 2071 2072 2086 2112 2123 2135 2185 2191 2228 2254 2260
2330 2410 2412 2431 2435 2452 2498 2512 2572 2577 2602 2649
2681 2689 2718 2792 2819 2823 2830 2857 2875 2876 2881 2891
2903 2905 2929 2962 3044 3048 3056 3081 3108 3124 3179 3212
3240 3259 3261 3313 3320 3331 3344 3348 3365 3375 3385 3390
3398 3399 3414];[1 12 15 28 32 34 36 68 78
93 107 113 186 195 217 230 239 255 282 292 319
374 384 385 422 434 452 454 463 477 501 534 545
573 662 770 773 837 840 850 856 864 865 882 894
947 1033 1093 1107 1121 1140 1141 1145 1194 1215 1230 1273
1282 1297 1300 1315 1326 1328 1330 1351 1365 1439 1484 1490
1587 1608 1643 1646 1663 1673 1700 1707 1724 1728 1750 1769
1777 1805 1823 1828 1832 1835 1866 1873 1894 1911 1916 1956
1971 1987 2000 2003 2009 2020 2034 2039 2103 2170 2180 2188
2238 2351 2352 2357 2381 2433 2454 2461 2572 2596 2604 2617
2642 2650 2657 2665 2666 2690 2719 2735 2736 2762 2770 2786
2790 2812 2824 2838 2876 2897 2903 2924 2953 3030 3068 3111
3141 3151 3152 3243 3314 3316 3330 3359 3386 3407 3422];[59
115 116 217 252 265 331 349 361 396 422 424 442
465 509 517 533 539 547 549 563 569 614 615 652
691 693 715 743 769 773 799 831 832 846 893 894
925 953 964 987 997 1018 1057 1059 1061 1073 1115 1127
1148 1157 1181 1224 1227 1292 1314 1338 1348 1371 1397 1398
1402 1420 1463 1499 1514 1530 1551 1626 1708 1711 1724 1738
1799 1802 1806 1813 1815 1818 1841 1854 1874 1880 1911 1920
1924 1944 1951 2055 2058 2077 2103 2121 2124 2153 2160 2172
2173 2276 2315 2323 2362 2426 2448 2512 2530 2543 2551 2577
2600 2642 2644 2656 2706 2715 2727 2738 2798 2803 2823 2825

```
2851  2853  2869  2909  3001  3002  3017  3068  3089  3110  3114  3123
3127  3138  3148  3213  3215  3225  3280  3301  3315  3320];[47   130
170   195   217   274   347   350   362   365   373   399   413   414
431   499   609   612   635   697   707   711   712   724   749   759
760   766   773   784   786   887   896   899   905   933   953   994
1006  1008  1009  1027  1087  1092  1098  1134  1136  1205  1269  1270
1320  1362  1364  1381  1396  1410  1427  1462  1479  1502  1525  1531
1583  1591  1599  1612  1614  1630  1656  1683  1694  1748  1836  1886
1929  1937  1954  1980  2008  2066  2083  2086  2105  2108  2117  2128
2141  2217  2218  2264  2346  2356  2391  2407  2408  2416  2458  2460
2465  2471  2568  2581  2590  2596];};
% % SCORES=10;
% % ns={[14    15    16    21    22    32    74    77    82   106   120
149   167   214   216   242   287   290   315   327   328   329   336
357   370   418   420   467   502   550   552   573   608   665   698
712   716   766   768   792   803   918   969   985   996  1016  1034
1038  1059  1166  1184  1200  1213  1215  1218  1219  1222  1309  1324
1329  1333  1340  1353  1358  1422  1426  1464  1493  1507  1520  1551
1555  1586  1600  1609  1619  1645  1657  1722  1724  1726  1747  1768
1810  1811  1833  1873  1874  1885  1887  1895  1903  1953  1990  2037
2042  2090  2091  2129  2130  2140  2158  2176  2210  2225  2232  2296
2302  2311  2314  2358  2365  2366  2403  2413  2432  2472  2528  2531
2571  2594];[36    42   140   173   174   186   199   245   296   298
341   371   430   523   524   535   563   593   608   615   641   672
740   753   764   776   789   796   802   846   859   860   866   941
950  1051  1065  1075  1113  1161  1185  1205  1253  1277  1278  1321
1326  1360  1365  1372  1385  1421  1453  1475  1481  1495  1507  1517
1521  1565  1567  1611  1622  1652  1674  1705  1788  1814  1819  1860
1862  1926  1931  1948  1977  1987  2079  2089  2096  2246  2250  2280
2282  2284  2330  2337  2356  2367  2440  2456  2498  2546  2674  2735
2769  2774  2806  2813  2840  2841  2858  2863  2909  2919  2927  2966
2973  2986  2998  3002  3013  3063  3078  3118  3157  3158  3161  3179
3181  3213  3248  3287  3291  3302  3335  3425  3437  3466  3468  3497
3549  3555  3565  3600  3601  3626  3636  3659  3815  3838  3844  3869
3897  3956  4009  4013  4054  4061  4069  4071  4098  4102  4185  4186
4197  4217  4221  4297  4300  4315  4320  4418  4423  4434  4437  4455
4494  4533  4542  4546  4574  4598  4656  4681  4684  4691  4712  4717
4764  4788  4805  4862  4887  4901  4991  4995  4999  5052  5107  5110
5112  5134  5139  5157  5162  5217  5255  5278  5297  5335  5390  5415
5442  5444  5488  5512  5524  5571  5647  5678  5701  5721  5751];[63
78    85    89   130   142   157   160   181   193   215   254   260
261   280   292   323   325   337   355   366   375   406   408   453
471   494   572   577   589   647   663   690   729   770   790   792
817   823   826   876   902   904   937   959  1020  1024  1037  1038
1046  1056  1109  1148  1152  1164  1198  1244  1282  1287  1299  1300
1354  1369  1426  1460  1474  1482  1492  1534  1611  1615  1627  1633
1640  1642  1645  1670  1689  1693  1700  1755  1805  1823  1864  1897
1961  1963  2031  2051  2056  2124  2141  2181  2193  2201  2202  2212
2249  2251  2288  2295  2428  2444  2484  2504  2509  2567  2613  2620
2633  2640  2676  2697  2735  2755  2768  2780  2786  2793  2820  2836
2866  2884  2910  2921  2933  2934  2958  3000  3031  3035  3072  3086
3090  3093  3101  3118  3199  3215  3228  3235  3271  3311  3342  3360
3384  3393  3408  3419  3422  3425  3457  3469  3472  3481  3494  3502
3506  3598  3630  3632];[17    19    80   138   142   144   158   160
252   253   333   384   430   436   437   465   467   524   529   541
569   570   606   637   656   670   672   673   676   689   702   717
738   742   760   771   783   823   860   962  1055  1094  1137  1140
1157  1163  1172  1181  1182  1265  1272  1285  1310  1386  1437  1444
```

1458 1557 1611 1612 1625 1630 1655 1657 1701 1732 1766 1779
1798 1801 1878 1894 1958 1973 2017 2041 2046 2059 2085 2097
2111 2172 2178 2229 2234 2246 2264 2298 2353 2389 2428 2429
2441 2442 2496 2528 2531 2569 2570 2583 2584 2631 2645 2650
2660 2666 2676 2701 2702 2715 2717 2746 2755 2791 2859
2893];[47 65 68 80 109 154 180 200 201 224 239
258 264 272 274 281 321 330 341 347 366 367 369
380 419 445 458 472 487 507 508 544 570 573 581
583 587 594 602 609 639 677 678 722 727 776 799
800 817 839 840 844 880 904 914 943 968 977 991
1009 1012 1038 1048 1052 1054 1065 1073 1090 1104 1175 1204
1301 1302 1336 1353 1408 1422 1429 1456 1480 1513 1515 1644
1665 1666 1669 1678 1726 1805 1827 1837 1880 1887 1912 1940
1967 2001 2002 2019 2035 2094 2124 2141 2186 2217 2219 2243
2340 2350 2416 2468 2476 2494 2536 2544 2561 2573 2619 2635
2666 2667 2789 2804 2822 2848 2850 2866 2943 2980 3067 3068
3073 3078 3107 3113 3125 3132 3162 3170 3184 3254];[18 30
31 37 52 63 65 126 127 134 135 154 186 191
192 275 281 282 292 343 374 453 507 508 526 585
636 651 743 770 847 905 960 961 962 963 990 993
1009 1011 1015 1021 1032 1037 1041 1048 1053 1079 1115 1160
1176 1177 1183 1186 1218 1250 1259 1299 1430 1464 1476 1521
1567 1644 1728 1742 1754 1799 1853 1874 1883 1886 1900 1925
1940 1950 1956 2006 2050 2062 2066 2079 2126 2140 2170 2174
2252 2258 2284 2291 2294 2319 2328 2333 2357 2404 2421 2435
2444 2465 2472 2481 2530 2600 2602 2615 2651 2695 2721 2722
2733 2750 2764 2787 2886 2898 2899 2952 2973 2978 2980 2989
3033 3041 3084 3093 3094 3112 3121 3127 3137 3158 3240 3243
3335 3348 3376 3378 3386 3387 3391 3432 3448 3510 3521 3529
3539 3629];[12 28 40 105 186 192 203 212 244 249
258 274 279 334 340 370 423 425 426 433 434 491
512 521 548 562 609 614 629 655 663 714 740 743
744 759 896 924 929 963 969 1019 1031 1049 1108 1182
1199 1256 1313 1321 1323 1370 1506 1519 1520 1555 1628 1644
1655 1661 1670 1698 1756 1758 1762 1852 1860 1873 1888 1909
1915 1924 1950 1968 1973 1994 2105 2118 2249 2287 2347 2377
2378 2384 2390 2427 2432 2441 2480 2482 2512 2519 2520 2543
2544 2570 2602 2607 2609 2646 2647 2774 2780 2786 2802 2845
2869 2870 2872 2920 2949 3000 3033 3038 3079 3094 3105 3147
3204 3249 3254 3287 3313 3320 3326 3341 3345 3347 3383 3393
3408 3416 3420 3437 3458 3464 3466 3484 3489 3557 3597 3709
3728 3743 3759 3762 3787 3863 3868 3919 3920 3945 4011 4013
4032 4069 4078 4107 4108 4128 4138 4146 4177 4187 4211 4234
4237 4242 4260 4320 4328 4340 4408 4415 4455 4457 4464 4466
4485 4492 4511 4527];[14 19 45 54 93 99 105 215
297 298 362 365 401 429 440 441 467 478 487 515
516 519 521 567 581 582 681 684 702 717 778 820
834 850 856 862 903 919 922 925 952 977 982 987
1014 1024 1029 1064 1077 1115 1131 1142 1149 1151 1184 1190
1191 1226 1236 1255 1262 1329 1372 1386 1408 1448 1449 1450
1456 1473 1481 1499 1572 1623 1685 1697 1719 1725 1781 1801
1811 1832 1844 1845 1926 1964 1988 1991 2003 2009 2040 2041
2079 2090 2129 2142 2145 2195 2205 2207 2265 2273 2312 2323
2326 2350 2375 2412 2433 2554 2583 2589 2590 2596 2597 2609
2612 2613 2681 2821 2951 2952 2953 3007 3022 3033 3034 3069
3081 3111 3128 3181 3190 3217 3218 3233 3299 3309 3326 3328
3337 3338 3347];[7 15 17 29 79 151 165 170 190
233 298 305 314 342 367 418 437 459 486 496 500

```
502   516   542   553   569   573   574   701   708   721   796   808
819   890   897   923   975   976  1000  1006  1010  1019  1034  1050
1069  1114  1127  1146  1172  1180  1185  1198  1326  1343  1344  1374
1392  1415  1548  1567  1580  1584  1617  1627  1632  1645  1657  1774
1849  1851  1893  1927  1937  1981  2001  2021  2033  2038  2079  2096
2147  2167  2202  2247  2248  2255  2294  2324  2361  2366  2381  2400
2421  2422  2466  2511  2539  2563  2572  2600  2606  2628  2640  2684
2687  2696  2697  2700  2741  2769  2810  2848  2879  2884  2891  2906
2943  2970  2973  3002  3056  3062  3108  3116  3127  3136  3173  3185
3227  3284  3289  3298  3338  3340  3363  3372  3425  3457  3482  3497
3507  3523  3524];[15    41    56    58    73    79    80    92    98
120   147   176   251   312   333   336   354   360   418   419   483
532   576   585   611   629   633   642   647   666   700   746   750
760   788   803   813   826   841   873   881   893   894   896   940
976   988  1023  1031  1045  1057  1058  1082  1088  1098  1137  1179
1191  1201  1202  1213  1235  1252  1286  1319  1332  1337  1349  1442
1489  1545  1551  1587  1609  1666  1694  1722  1743  1816  1838  1841
1880  1925  1939  1961  1999  2001  2016  2049  2066  2094  2129  2191
2211  2272  2321  2339  2373  2401  2408  2497  2524  2537  2569  2582
2586];[15    62    70    96   128   153   216   219   236   245   261
263   264   346   365   396   436   496   521   583   586   616   645
647   649   667   673   700   715   725   740   777   886   935   937
938   949  1002  1014  1016  1048  1074  1085  1105  1111  1160  1289
1303  1336  1361  1373  1433  1435  1459  1460  1521  1552  1573  1624
1631  1693  1694  1724  1745  1768  1813  1836  1842  1866  1934  1945
1951  1953  1989  1990  2011  2042  2100  2121  2321  2344  2363  2392
2407  2412  2426  2450  2456  2463  2469  2487  2544  2560  2562  2570
2571  2576  2583  2612  2627  2659  2698  2714  2719  2748  2839  2929
2940  2949  2996  3057  3058  3059  3069  3087  3096  3106  3175  3238
3271  3272  3290  3336  3343  3368  3411  3450  3455  3537  3560  3561
3588  3589  3596  3615  3644  3646  3668  3677  3732  3769  3899  3985
4025  4034  4058  4103  4107  4140  4182  4207  4229  4268  4275  4282
4348  4356  4371  4389  4461  4483  4499  4577  4581  4636  4662  4688
4801  4835  4852];[7     8    16    37    54    93   113   120   150
270   293   294   339   358   386   401   423   427   459   505   509
517   518   529   575   589   644   676   743   747   776   801   803
805   878   889   910   915   930   933   937   950   996  1083  1105
1153  1178  1205  1274  1328  1444  1512  1518  1533  1539  1540  1593
1659  1661  1679  1699  1707  1759  1777  1783  1805  1823  1833  1834
1837  1850  1858  1887  1907  1908  1959  2005  2007  2030  2111  2139
2175  2187  2204  2221  2255  2256  2267  2340  2369  2383  2399  2413
2434  2480  2503  2513  2573];[39    40    53    68    82    85   122
137   153   207   258   287   312   357   366   406   438   446   474
477   478   497   545   547   552   554   602   608   617   618   638
654   665   714   726   753   777   786   833   857   887   899   901
908   940  1015  1016  1018  1026  1028  1047  1074  1092  1117  1172
1176  1201  1214  1237  1238  1275  1322  1334  1357  1404  1422  1437
1450  1592  1623  1659  1676  1702  1742  1756  1836  1854  1856  1858
1934  1944  1951  2032  2039  2056  2062  2066  2079  2164  2229  2253
2288  2335  2348  2389  2398  2403  2422  2434  2435  2439  2443  2481
2534  2596  2614  2653  2667  2674  2691];[1    57    59   120   152
179   187   197   199   240   359   367   404   406   443   526   529
531   536   584   669   713   715   722   739   740   751   756   762
811   832   880   915   921   929   939   955   956   957   965   981
982  1056  1061  1079  1100  1111  1160  1163  1201  1248  1303  1317
1318  1338  1360  1398  1445  1446  1451  1551  1592  1605  1606  1633
1660  1689  1707  1715  1716  1723  1765  1779  1791  1803  1817  1824
1882  1928  1947  1964  1968  1994  2033  2068  2115  2124  2131  2186
```

2189 2198 2200 2277 2284 2299 2357 2364 2381 2386 2397 2439
2446 2456 2463 2483 2500 2524];[2 5 36 74 75 78
79 90 134 191 218 265 276 298 341 385 479 517
539 545 562 617 694 704 801 808 836 859 864 872
942 963 994 997 1035 1037 1065 1086 1148 1201 1293 1319
1320 1329 1439 1465 1476 1487 1509 1521 1565 1574 1584 1609
1614 1624 1632 1640 1660 1672 1707 1775 1781 1802 1806 1835
1837 1843 1845 1848 1875 1883 1906];[1 35 55 71 85
120 176 179 180 194 246 310 364 384 392 433 434
443 457 468 475 495 497 517 531 536 585 590 596
623 641 658 671 678 724 834 870 899 919 927 967
1012 1014 1061 1126 1202 1270 1296 1301 1312 1315 1329 1344
1345 1375 1388 1391 1396 1430 1454 1456 1477 1530 1590 1640
1644 1694 1704 1735 1758 1816 1833 1849 1880 1899 1929 1940
1971 1972 2032 2111 2180 2184 2206 2246 2248 2313 2321 2376
2391 2394 2404 2430 2463 2511 2524 2537 2549 2575 2586 2632
2645 2651 2688 2697 2716 2739 2805 2816 2840 2847 2853 2892
2903 2925 2944 2945 2949 2958 2968 2973 3021 3057 3059 3066
3070 3088 3135 3188 3226 3237 3274 3282 3284 3302 3305 3306
3311 3332 3354 3365 3381 3395 3419 3422 3437 3478 3515 3550
3558 3560 3577 3633 3637 3659 3694 3713 3714];[52 98 110
129 143 178 181 192 288 313 319 330 344 355 368
396 402 417 424 441 499 519 567 569 583 584 629
638 682 715 745 747 758 767 802 811 815 816 837
844 851 886 896 906 930 936 977 1030 1060 1164 1184
1202 1214 1239 1261 1278 1281 1295 1301 1302 1314 1320 1342
1363 1420 1442 1454 1461 1549 1561 1563 1566 1614 1634 1663
1664 1677 1706 1709 1711 1740 1776 1813 1844 1890 1907 1969
2002 2012 2034 2081 2131 2170 2171 2178 2212 2312 2338 2345
2408 2412 2450 2540 2568 2588];[5 6 17 50 71 96
100 156 168 199 200 218 282 308 317 334 366 411
428 433 438 504 514 521 577 581 590 602 606 691
713 753 787 798 810 840 842 852 930 936 943 970
973 980 1066 1128 1137 1167 1172 1183 1187 1223 1224 1250
1272 1279 1319 1322 1352 1363 1396 1418 1426 1431 1514 1540
1553 1560 1591 1603 1607 1687 1696 1700 1703 1761 1803 1805
1823 1855 1899 1953 1982 2022 2028 2063 2072 2086 2112 2135
2185 2189 2191 2228 2254 2260 2307 2319 2330 2360 2408 2410
2412 2431 2435 2445 2452 2498 2512 2572 2577 2583 2602 2675
2681 2689 2718 2792 2823 2830 2857 2881 2891 2903 2905 2911
2929 2959 2962 3005 3041 3044 3048 3056 3081 3103 3108 3121
3124 3179 3195 3212 3240 3261 3313 3320 3331 3344 3365 3375
3398 3399 3407 3414];[1 12 32 34 36 78 93 107
113 186 195 217 230 239 255 271 292 319 374 384
385 422 434 452 454 463 501 534 545 565 573 662
745 773 837 856 864 882 896 947 1033 1093 1107 1140
1141 1145 1194 1215 1230 1234 1282 1297 1326 1328 1330 1342
1351 1365 1430 1439 1587 1608 1629 1643 1646 1673 1700 1728
1750 1769 1777 1805 1823 1828 1838 1866 1873 1891 1911 1944
1956 1971 1983 1987 2000 2003 2009 2032 2034 2039 2103 2170
2188 2223 2227 2238 2351 2352 2357 2381 2387 2388 2433 2454
2461 2472 2522 2572 2604 2617 2642 2650 2657 2665 2666 2690
2719 2735 2762 2770 2786 2790 2812 2824 2843 2876 2897 2903
2924 2953 3030 3068 3072 3111 3141 3151 3152 3153 3170 3215
3243 3262 3314 3316 3330 3359 3386 3404 3407 3422];[59 115
116 217 252 265 331 349 361 396 422 424 442 517
533 539 547 569 614 615 652 691 693 715 769 773
799 822 831 832 846 870 893 894 925 953 964 984

```
987  1018  1059  1061  1073  1115  1148  1157  1223  1269  1292  1296
1314  1348  1371  1397  1398  1402  1463  1466  1477  1499  1514  1551
1579  1637  1708  1711  1761  1799  1802  1813  1815  1818  1874  1880
1911  1944  1951  1958  2055  2058  2099  2103  2121  2124  2155  2156
2172  2183  2210  2247  2275  2276  2315  2323  2362  2420  2426  2439
2512  2530  2543  2642  2644  2656  2700  2706  2715  2737  2738  2798
2803  2823  2825  2851  2853  2909  3001  3002  3068  3110  3114  3123
3127  3138  3148  3187  3200  3213  3215  3224  3225  3245  3301  3315
3320];[47   130   170   183   195   201   217   274   347   350   362
365   393   413   414   419   431   499   596   609   612   635   697
707   711   712   724   749   759   760   766   768   773   784   786
896   899   905   933   953   994   998  1001  1007  1008  1009  1027
1087  1092  1098  1131  1136  1140  1269  1270  1362  1410  1420  1427
1479  1502  1520  1525  1531  1536  1557  1583  1591  1599  1612  1614
1630  1656  1683  1694  1745  1748  1818  1836  1886  1929  1932  1954
2023  2066  2083  2086  2105  2108  2117  2141  2166  2194  2217  2218
2264  2274  2346  2391  2408  2416  2450  2458  2460  2471  2536  2568
2581  2590  2596  2629];};
% SCORES=[13 14 12 9 8];
% ns={[420  2232  2594  2432  1340   167   792  1600  1311  2210  1222
2037   106  2090  1520  2530  1329   327  2528  1874   370   336  1038
1422  2358   716  1218  1307  1586  1722  2599  1215  2413  2366  2365
2014  2311   149  1609   923    22   329  1953  2225   768   625   353
1885  1903  2302  1507    83  1166   608  1726  1555  1358  1565  1177
2571   477    32  2082   216  1551  1645   573  1493   665  1200   842
1619  1887  2089  2060  1059  1768  1738  1213  2289   261  2204  1636
77  1041   698  2007   350  2527   791   712   765  1216  2129   552
82  1747  2531  2249  1701  1426    35  2140   287  1309  1873   996
1724  2008  1673   549    21  1355   354   262  2403   894  1439  2313
2575   970    80   342   555   249   484   969   985  2027   172   613
720  1269  1274  1730    88   718    78   713  1039  1178  1219  1786
1957  1103  2469   505  1143  1450  1657  2156  2230  2557    16    40
410   645  1021  1251  1683  2066  2511   131  1851  2127  2568   328
1237  1800  1951  1986  2158  2239   647  1599  1762  2278   667  1016
1299  1585  2567    14   171   214   315   409   467   502   550   707
723   918  1324  1353  1419  1833  1990  2176  2314  2472  2544    15
74   120   290   357   418   610   724   766   803   979   992  1050
1127  1333  1464  1536  1708  1811  1859  1895  1978  2003  2042  2091
2130  2139  2430];
% [4320  5701  3425  2769   186  3838  3468  2735  4297   615  4684  2089
140  1819  2927  2330  1453  1726  1507  4656  2998   608   789  2079
3601  4221  2097  3437  2129  1931  2806  2337  5442  4071  2840  3181
859  5112  2356  2367  1065  1051    36  1620  4574  4061  1674  3844
296  3889  3525  3636  4169  2196  2439  3370  1249  2562  1113  1393
2250   740  4434  5255  5157  1277  1421   808  3254  3908  1991  1372
1495   616  2282  1871  3157  1052  1622  4533   439  2546  2863  4928
199  4862     1  4546  5107  3897  4538  4092  3161  4102  1064  5000
1409  5721  1858   298  4537  5512  4788  2246   245  1814   324   593
1205   802   764  2284  2877   523  4258  1788   640  3118  2590  1685
5690  5274   173  3659   866  3302   569  2774  4484  1314  5524   270
822  3002  4009  4598  4631   894  1926  4217  3174  3975  2511  5327
5748  2142  3244   524  1321    52  3085  4151  3182  3411  4901   863
4649  5217  3898  4069   540  1185  1664  3158  1948  5333  5488  5415
436  4991  5727  3702  2280  1879   612  1652  5160  1565  5477  2545
776   641  4999  1365   338  3450  5052   417  4908   486  2674   675
5647  1937  1908  4644   941  1049   753  1617  4712    68  5464  5564
1736  3635  3405  3956   513  4437   244  4494  1139  3381  5571  1024
4542  2841  4760  1519    42   846   648   430  2474  5677  5498  4691
```

1186 5678 1360 1537 4121 5349 514 4811 2858 4099 5425 3180
2104 4583 4325 1482 3836 5758 1128 5336 3849 1705 1983 4940
5751 867 913 2000 3893 4332 5207 4535 1866 2854 3291 4017
4438 4681 5392 796 1863 1980 4939 5099 1633 3086 5390 34
672 872 1838 2905 3128 4300 4995 82 607 949 951 2086
2409 4100 231 233 372 1136 2949 4662 5395 663 945 1262
1670 3166 3192 4184 4308 5398 89 862 1135 2005 2984 3243
3798 4444 4741 4792 4846 174 323 535 2163 2344 2369 2738
3142 3224 3333 3766 3878 4013 4175 4606 4724 5397 1203 4729
63 2383 3368 5483 5638 237 1881 2589 2794 3172 3206 4038
4235 4718 4919 31 187 200 1501 2387 2535 2773 3018 3063
3672 3965 4612 4682 4914 5278 5561 5703 288 337 341 428
563 734 950 1066 1253 1278 1326 1475 1559 1860 1862 1876
1942 1950 1959 2096 2143 2678 2788 2966 2986 3078 3179 3213
3309 3357 3497 3546 3555 3565 3600 3626 3815 3869 3907 4161
4185 4197 4315 4375 4418 4455 4717 4764 4867 4887 4980 5110
5134 5139 5235 5391 5431 5444 5744 8 43 108 287 289
371 579 590 1075 1102 1111 1161 1274 1311 1481 1517 1521
1567 1571 1578 1601 1610 1611 1683 1759 1920 1977 1987 2274
2431 2440 2456 2498 2543 2813 2909 2911 2919 2960 2973 3215
3248 3287 3296 3321 3335 3499 3549 3780 4054 4186 4423 4477
4550 4558 4604 4654 4799 4805 5068 5122 5318 5388 5608];
% [904 729 2793 261 3481 1244 3093 3271 2056 1299 3199 2836
647 1474 1148 1020 3360 1492 2780 2697 1640 3101 876 260
888 3384 1056 1864 2249 1046 254 690 1627 792 2768 1491
2884 2620 2295 3031 2958 3072 3311 1426 1037 2820 3506 3235
366 789 3342 323 471 2921 507 1534 1693 2934 3577 1077
1805 1961 3216 3408 1047 1164 3513 1182 1689 3547 1759 676
160 1300 1024 1318 2202 2756 344 2484 2613 3502 3248 3472
1109 701 1193 193 1824 1287 1127 1289 3210 450 937 2536
2910 1842 3230 1670 790 1599 63 3386 2212 2509 3310 2436
3215 3393 408 826 429 494 663 2435 2124 2676 1354 157
648 3630 1963 2567 325 2680 1977 1460 2167 3457 2933 770
2576 2755 3453 1571 2829 2288 1776 2346 2610 2031 970 1198
3494 1823 2141 915 1163 1482 1519 3035 375 705 89 2363
1366 292 1350 756 3324 905 2572 3082 285 3491 3532 298
3172 3282 675 1349 1819 2055 3388 364 489 1019 3555 352
889 986 1262 3228 3619 363 559 3119 902 943 1106 2373
3116 238 859 1029 1129 1213 1678 1901 2409 2735 2889 3094
590 730 922 1307 2138 2209 2929 3176 3335 38 214 355
404 785 989 1084 2042 2193 2259 2574 40 1476 3068 1260
682 1500 1720 2708 2960 3191 3312 3597 3626 25 118 416
567 873 1290 1370 1609 1645 2696 3252 3414 3550 3560 142
181 315 337 400 405 453 817 823 840 1038 1175 1282
1615 1633 1897 2251 2393 2444 2457 2504 2866 3086 3090 3419
3425 3469 3512 3598 39 78 85 130 247 280 376 406
572 577 589 718 939 959 1083 1152 1335 1369 1611 1631
1642 1710 1755 2051 2201 2211 2232 2331 2428 2491 2640 2761
2786 2925 3000 3118 3204 3404 3454 3497 3632];
% [2097 570 2059 823 160 1157 2650 2298 1265 1285 860 1094
1657 2806 436 962 1691 2631 2389 1172 1611 2163 673 1731
2041 541 2755 2666 1581 17 21 1182 1279 2893 252 672
2756 2235 736 2859 669 1444 1958 2233 2098 2570 158 738
1107 783 2583 2496 2894 1288 2569 2017 968 606 259 1272
1894 2702 1255 784 1128 2240 1824 1973 2095 689 2304 1961
1566 1701 952 1847 529 253 1635 2442 2229 2531 2292 1732
2460 441 1779 2686 2441 1557 569 1338 2660 2178 2526 2717
443 1137 2746 658 1924 1713 1055 832 350 358 1181 800

```
1458  1849   290  1615  1710  2061   861  2919  1310  1371  2086   855
1383   989  1117  2353  2738   216  1626  1699  2126  1390   317  1439
1888   568   647   685   884   942   553   706   763   973  1766  2509
218   869  1479   487   910   932  1011  1268  1825   465   520  1080
1405  1704  2387  2462   175   597  1345  1492  1765  1798  2345  2685
1048  1684  1949  2471   163   694  2400  2613   205   731  1667  1700
2705  2808    31   254   278   421   810   916   917   974  1041  1574
1610  1625  1714  2096  2177  2697    15    22   142   144   467   656
702   717   742  1071  1140  1163  1309  1348  1386  1437  1655  1815
1878  2010  2085  2100  2137  2234  2264  2284  2428  2429  2584  2624
2676  2701  2715  2805    19    28    80   138   181   251   333   384
430   437   461   524   670   760   858   875  1066  1377  1630  1801
2111  2172  2246  2253  2527  2528  2645];
% [2476  2243  1940  1302  3184   281  1837   776   844  1336   800   839
904  2544  1012  1644   201  1678   727  3067  1967  1138   570   367
2822   321   544  2468  1515  1071  2350  3125  1887  2516  2635  2848
1072  2186  2343   783  3162  1939   419  1627   299  3068  2944   442
609  2001   347  2473    68  1301   675  1224   880   583   558   587
799  2336   109   573  2008  2340  2619   677  1665   341   186   417
1827   676   814   639   440  2535  1073  1289   330  1009  2035  1048
272  3283   712  1340   581  3055   171  2804   366  1646  2019   594
3132  1117   458  2789  1104   507  2536  1912   840   602   472   519
81  3113   943  3260  2361  2866  2417  3073  2265  1961   258  2364
1529  2432   489  2377  3166  1052  2949   775  1065  1835  2829   420
2124   369  1214  1569   819  2088  3254   541   543  1198  1871  2442
269  2997  1422  1740  3198    51  2255  2670   384   457   503  1043
1380  2695  2998  3230  2094  2622  2826    65   180   399   636   826
1868  3029  3186  3275   538   616   791  2012  2440  2827  3060    75
325   342  1010  1341  1364  1539  1595  1757  1956  3263  3265  3339
1771  1493  3169   808  1021  1172  2122  2821  2921   229  1102  1990
8   290   427   522   574  1041  1053  1257  1473  1706  1732  1781  2573
2681  3041  3289    40    47   142   243   274   445   487   508   509
722   763   838   914  1038  1054  1090  1163  1204  1265  1353  1408
1429  1456  1480  1513  1793  1805  1901  2024  2217  2219  2501  2561
2593  2667  2850  2980  3078  3107  3153  3181  3244     4    80   154
172   200   224   239   264   374   380   521   817   856   886   968
977   991  1101  1147  1175  1639  1666  1669  1687  1726  1850  1865
1880  2002  2141  2334  2494  2572  2666  2674  2755  3170  3179  3235];
% [3510   292  1754   743  2421  1385  2952  3033  1728  1836  3376  3121
963   770  1011  2787  2435  1055  3127  1015  1259  1099  1053  1853
282   961  2721  2602    98   585  1945  1124  2126   191  1430  3529
1115  2980   135  3450  2791  3671  1541  2006   962   267   192  3428
3462  2252  2613  2733   651  1299  2651  3093  3348  2472  2961  2764
1250  3239  2982  2362   281   256  2193    27   480  3539  1940  1734
275  1338    63  1135  2174  2886   908  1476  1184   134  3351  3448
186  1021  1213    52   414  2722  2497   456  1900   847  2036  3631
2792  3333  3084  1220  3346   180  1972  3005  3452  1464  3041   346
755  1453  1994   374  2258  3243  2908  1216  1946  3718   122  1242
921  3387  2444  2111   508  3374  2079  1466  3447    18  2294   439
2956  1241  1886  2328  3714   365  1868  2616  2580  1632  2907  3620
3705   510  1878  1208  2939   453  2251  2297  3014  1867  3006  1117
2765  3606  1501  2190  1221  1343  1482  2887  2093  3240   138    37
1254  2615  2891  2898  1079  1467  2916  3583  2027  2804  2821  3088
14   394   812   964  2660   132  1163  1431  1797  2322  2585  3540
3728   347  1799  2664  2763  2950  3076   139   178   313   525   723
1012  1781  1831  1851  1929  2337  3025  3113  3268    65   234   268
638  1211  1751  1965  2112  2387  2533  2689  2853  2976   331   507
1693   665  1308  3122  3392   297   721   734  1439  1956  1992  2738
```

2999 3302 3449 30 127 343 455 905 960 993 997 1009
1032 1047 1048 1077 1176 1186 1222 1498 1521 1742 1874 1883
1950 2050 2066 2140 2170 2284 2291 2319 2333 2465 2530 2559
2600 2643 2669 2695 2854 2899 2902 2973 2978 2989 3071 3094
3112 3158 3335 3378 3441 3629 31 126 130 136 154 177
332 526 636 990 1037 1041 1134 1160 1177 1183 1218 1567
1644 1798 1896 1925 2062 2083 2125 2145 2321 2357 2365 2404
3022 3074 3137 3386 3391 3432];
% [4234 1756 1670 663 423 370 714 4455 609 1968 4408 4211
2796 3249 192 2512 655 3147 1762 2390 2432 548 4466 4138
3709 629 2377 3920 102 4260 2802 3767 2347 3863 4457 3105
4527 1873 4040 1904 2441 1269 1555 1701 1973 3713 4485 1370
2920 28 1099 1915 1313 4013 3528 4415 1924 1506 274 2544
2378 3033 2780 3743 3313 1199 3416 1069 788 3204 1321 106
3346 3919 1918 1994 2287 2407 1084 3000 2869 570 3413 4032
105 3341 3408 2872 4078 614 3923 3557 3539 2886 2520 4102
4328 40 876 3533 3254 1031 1985 791 968 1320 3464 1520
4177 982 3420 1108 3017 491 2876 3043 3728 1799 3597 3479
4128 3522 2427 96 4108 3296 1992 3782 1152 4107 4385 513
1589 3466 1852 4031 3320 1990 4146 2844 249 425 1617 147
244 1655 1661 2536 3868 740 2723 1450 4227 664 2570 4181
4188 743 543 2285 967 433 1228 3914 3732 3994 661 2790
3945 516 1592 2589 2745 1950 3890 204 521 3165 532 3723
3345 1501 2609 2845 3079 1154 300 1014 1150 4171 2808 3359
2096 2683 151 777 1628 2470 2538 4244 3364 4212 1517 4129
1908 2196 2297 3247 3519 4225 3908 4183 2156 2550 2590 4232
4237 569 796 1508 2504 2960 3213 3353 252 1006 1639 2038
2124 2428 2721 3126 3210 3314 3391 3669 4209 159 891 1438
1737 2830 2919 3481 3898 4007 4044 4300 4331 4587 283 361
1606 1608 1687 1769 2507 2684 2991 4548 71 239 934 1177
1252 1626 1718 1746 1887 2070 2420 2902 3616 4510 1851 2923
152 3848 1376 1380 2409 2518 826 970 1369 1393 1761 2084
3220 1127 1668 2310 2593 2824 2831 2997 3044 3336 4008 4334
258 575 719 1198 1247 1850 2261 2772 2949 3122 3750 3775
3810 3968 4139 4347 4350 4515 4518 12 161 212 253 263
334 340 374 426 434 562 924 929 969 1049 1106 1256
1323 1391 1465 1758 1847 1888 2105 2325 2480 2508 2519 2543
2607 2646 2647 2769 2774 2833 2870 3038 3094 3383 3393 3458
3484 3820 4011 4242 4340 4464 72 203 227 251 279 296
392 759 852 896 963 966 978 1010 1019 1315 1421 1519
1565 1644 1698 1749 1860 1909 2118 2193 2209 2249 2418 2446
2482 2495 2644 2786 2988 3287 3300 3326 3347 3384 3437 3449
3489 3759 3762 3787 4023 4069 4104 4233 4320 4511];
% [820 1184 2613 478 1191 1226 2609 2412 1473 1499 3299 581
2079 2041 1408 2375 1845 1262 1372 903 3218 515 2003 1014
401 2126 2273 1236 2583 834 2597 2220 1884 3069 1740 487
567 1329 1992 2129 3034 1684 1064 215 436 2398 1697 1481
1115 45 2207 441 105 3181 2195 3111 3007 982 3337 521
1255 3233 2253 582 442 3033 681 2255 1719 2748 1077 2670
1538 2596 924 1152 1395 856 2229 428 3128 93 2013 850
298 702 384 2040 2406 2953 1926 1449 3326 525 19 365
420 362 550 1622 256 1190 54 2545 2374 516 3216 2230
440 2009 684 2312 1811 2323 73 137 1456 766 2692 892
2589 1128 1752 2142 617 2778 1450 2590 2350 2326 661 1921
469 2146 1728 987 3063 138 2761 3021 579 1078 2289 398
2282 1140 2380 1118 2717 2387 2569 438 731 2263 2290 1298
1535 2468 1758 3008 477 1056 1065 1572 2359 3314 1203 1925
2099 2961 1294 1426 1984 2633 3060 397 648 978 1681 2043

2056 216 409 925 1187 1448 2727 2955 3080 445 1405 266
1307 1374 1602 1779 1800 2423 2520 2652 2730 787 952 1020
2145 2414 2607 2737 99 249 415 517 717 826 923 943
985 1027 1029 1045 1131 1142 1149 1151 1176 1247 1386 1623
1725 1832 1844 1963 1964 2001 2090 2204 2205 2554 2612 2681
2693 2951 3022 3081 3174 3217 3309 3338 53 174 297 322
338 399 429 467 519 778 862 919 922 948 977 980
984 1051 1416 1611 1677 1685 1801 1928 1988 1991 2265 2294
2433 2952 3190 3347];
% [2906 542 190 890 808 114 367 418 2096 2943 2640 2038
151 3108 2884 1580 1146 3289 1851 708 2970 502 2033 305
2848 897 721 695 1031 1374 219 975 3523 1927 1180 437
79 477 516 2563 3062 165 1996 1247 2741 1034 923 1645
2466 2572 2021 1617 2628 1344 1548 15 838 2539 2421 2639
1584 1149 1774 2900 1069 1934 3462 2600 1983 3136 2536 3274
3497 1223 233 1127 1142 2361 2247 2810 2512 2027 573 1446
3233 2826 2001 3340 1033 1467 667 320 903 1786 876 1417
2022 1632 3425 459 746 1696 2605 979 2990 1325 2701 3005
2787 2079 3363 1326 345 3094 441 1737 2294 3457 2796 2298
819 3173 1937 1272 314 2366 3454 2809 1504 553 1198 1849
3116 2684 2451 2732 1129 3156 3002 1751 480 3227 234 491
3283 543 1421 1137 1328 469 638 1567 3451 627 937 1752
2069 2752 3423 1637 2805 2944 3284 144 878 1540 1579 2743
519 1050 1478 2003 2755 2794 2744 3422 392 395 579 2133
49 909 1258 1589 2167 2276 2381 2510 3348 3518 123 736
933 2065 2604 2879 540 976 1038 1761 2334 2705 1710 481
3110 2513 3476 77 729 1471 1500 2128 2901 3302 80 277
358 499 677 1297 1533 1545 1979 2085 2695 3006 3288 3534
7 157 170 342 486 500 574 596 786 796 1006 1010 1019
1114 1172 1185 1255 1343 1392 1465 1627 1657 1760 1767 1907
2147 2152 2343 2371 2400 2511 2696 2731 2769 2948 3056 3127
3185 3223 3298 3338 3347 3507 17 29 141 298 306 496
566 632 701 907 981 1061 1318 1415 1614 1753 1863 1893
1981 2230 2244 2248 2255 2275 2324 2484 2545 2660 2697 2700
2704 2891 2973 2982 3143 3372 3524];
% [1235 666 760 1587 1609 41 826 2066 251 1666 1880 1213
120 1319 336 1999 2129 80 1939 1023 1337 73 1694 2582
201 2001 1315 1961 633 1252 483 419 1838 551 718 893
2413 642 1594 58 894 988 1137 629 873 2586 2321 79
1726 1126 2016 976 360 813 2513 370 1031 2188 1744 1283
788 1082 1024 2152 2211 1179 611 1816 881 2232 1136 2089
576 1364 2334 1813 176 2524 1057 2497 2467 61 2468 2473
1743 333 1896 1125 945 1202 418 1572 1442 373 1985 147
2519 1317 585 631 2191 1045 2587 2419 948 359 57 2090
475 1112 905 1739 69 747 942 1814 658 820 1522 2355
292 621 663 1208 1406 1875 290 1276 1349 699 1363 1538
1695 2259 597 1101 1160 1589 354 378 440 750 1073 1469
1652 1677 1697 2158 2567 2576 1914 441 2238 2448 2494 2569
17 1650 2209 2324 864 1674 1962 2249 56 92 98 228
295 312 746 749 803 841 896 940 1088 1201 1286 1332
1489 1545 1551 1593 1722 1833 1841 1876 1986 2094 2173 2272
2339 2401 2408 2452 15 276 286 357 532 559 632 647
665 700 723 729 1058 1098 1124 1191 1238 1246 1336 1342
1353 1590 1925 2049 2373 2537];
% [1002 2929 1953 2392 2612 1074 2456 740 3560 645 3059 2940
2344 3588 938 4207 1361 430 4389 1373 1631 3238 1555 216
2996 667 70 2363 261 4581 2583 1573 188 1432 4034 155
4348 3985 2011 3732 1160 3272 4835 3596 3537 4462 3336 1433

```
4466  1813   219   283  2847  4058  2100  1842   935  4103  2659  4275
2450  4499  2698   153  4802   236  2519   458   436  2463  4178  4069
790  3668   521  4709  1866   726  4678   673   496   262  3175  1847
264  1768  3787   365  2042  1934    62   886  1989   647  2627  3646
1289  2719  2748  2508  1431  2521   583  2571  4483  3455  1694  3271
725   122  2476  2196  3311   863  3031  3450  2961   684  1524   263
2553  3796  4333    96   396  4229  3036  2138  2554   128   715  2321
324   235  1184  4577  1945  2412  4025  4568  3471   636  4662  1016
1851   641  4487  2114  1220  2839  2577  3134  1085  1105  2023  2383
2572  1303   700   492  4562  1460  4203   928  4516  3367  3782  1125
4182  1218  2801  3716   697  1424  3096  4339   179  1183  1232  2066
2570  2611  1954  2121  2765  3103  3342  3514  4231  4879   106  1501
1736  1789  3019  4563  4584    15   231  2084  2353  2778  2941  3900
4035  4246  1173  1738  2122  2334  2606  2731  3325  4330   105  2514
3513  3679  3689  4043  4329  4475   218   601   973   977  1189  1291
1933  2902  3057  3182  3197  3638  4368  4539  4783   435   586   646
1131  2013  2123  2171  2199  2257  2402  2459  2888  3680  3768  3976
4084  4208  4574  4582  4837   728  1026  1043  1341  2064  2134  2580
3899  4079  4090  1413   211   731  2503  2863   781   808  1310  1723
1724  1957  2870  3917  4187  4328  4503  3416  3556  3578  3789  4181
4355   220   222   239   245   319   523   578   616   664   777  1086
1111  1459  1521  1745  1836  1901  1951  2005  2407  2469  2487  2544
2562  2620  2949  3039  3106  3290  3343  3411  3561  3677  3817  4107
4268  4282  4461   206   346   769   831   937   949  1014  1048  1082
1197  1238  1245  1302  1336  1376  1435  1552  1596  1624  1681  1693
1766  1783  1935  1976  2063  2068  2105  2285  2426  2560  2576  2662
2714  2732  2857  3058  3069  3087  3143  3167  3294  3368  3374  3434
3555  3589  3615  3644  4036  4140  4371  4636  4688  4801  4858];
% [1540   910  1518    54  1153  1263   261  2256  2399  2340   270  1593
164  1777  2503   937  2267   996     7  1661    16  1907   358   933
2187  2175   805  1533   915  1282   423  2480    93   509   656  1833
8  2007  1915  2566    37  1402   801  1887  2206  1512  1168  2076   386
1823   889   529  1756  1117  1869  2218  1699  2202  1805  2005  1959
589  1344  2030   950   113   307  1908  1274  1348  1444   787   242
2434  1724   536  2217   613     6   150  1118   149    34  1660   339
1096  1728  1433  1594  2068  2513   890   168   776  1557  1137  1388
1820   644   575  2326  1662   545   803  1834  2203    38  2565  1290
280  1493  1357    22   184   288    97   366   741  1161  1358  1890
588  1303  2440   795  1858  1949  2404   196   784   955  1583  1632
2075  1061   275   739  2294   328   415   742   859  1305  1538  1725
1866  1237  1628  1817  1839  2009  2308  2384  2532  1196  2365  1431
128  1049  1547  2121    74  1626   761  1772  1850  2067  2110  2176
2287    72   220   279   294   401   427   518   676   743   747  1024
1105  1178  1205  1539  1707  1759  1783  1837  1933  2139  2255  2369
2383  2413  2573   120   169   278   293   331   459   505   878   969
1083  1226  1328  1405  1617  1656  1659  1851  1980  2044  2154  2170
2204  2302  2574];
% [85  2403   777   477  1944  1450  2335   153   940  1143  2229  2596
2534   857  1172   617  1798   908  1235   899  1592  2667  1074  1154
1934  2653   547  2439  1748   597  1159  1176  1659  2435   312   552
497  1237  1534  2309    82   258  2032  1238  2691  1417   936  2614
1015   351   478   543   784   602  1742   923   726   665   556   995
1353   900  2039  2430   654   714   522  2348  2434  2436  2421    53
1118  2443   342   618  1024   653   631  1073   287   638  1641   151
341  1533  2674   438   568  1836  1003  1912   664   833   608  1623
1202   405   920   406  1859  1886  1938  1117   627  1795    54   924
1346   463   834   266  1646  2334  1334   816  1789  2134   945  1691
2070  1834   357  1483  2341  1972   605  1047  2555   560  1245  1285
```

```
1290  1478  1752  1913  2539  2591    40   106  1404  1750   458   607
1593  1702  2011  1561  1854  2447  2287  2640   173  1531   470  1048
1247  2107   172   634   960  1186  1356  1560  2017  2580  2684    68
122   137   207   413   474   554   901  1028  1092  1214  1275  1322
1357  1422  1437  1521  1562  1676  1774  1856  2047  2062  2164  2398
2414  2424  2475  2478  2483   291   350   366   446   486   487   530
753   771   786   884   887  1016  1018  1026  1106  1349  1383  1476
1756  1857  1858  1887  1951  2056  2066  2079  2089  2253  2310  2389
2422  2442  2444  2451  2538];
% [179  2277  1318   756  1779  1061  1551  2068   957  1605  1803     1
669  2299   197  2439   832  2033   594  2131  1445   722  2115   930
713  1201  1947   762   982   925   531  2203  1319  1155  1398  1032
526   752  2463   880  1882  1660  2446   981   715  1397  2284  1889
2469   909   739  2381  1786  1111   529   921  1079  1765  2523   584
1100  1387   120  1717   874   199  1446   955  1633  1056  1348    73
536   422  1715   192   253  2483  1846   811  1822   956  1716   803
1303  2177  1750   965  1887   915  2524  2180   740  1634  2056  2445
2254  2415  1549   761  1297  1495   993   831  1505  1689    54  2239
113   309  1845  2195   404  1830   243   367  1680  1916  2136  2472
245   530   569  1156  1725   449  1099  1147  1203  1248  1671  1672
1848  2245  1427  1521  2193  2424  1817  2093   876  1583  2413   714
2023  2270  2340  2364  2496   327   736  1513  1673  1993  2348    57
95   161   240   406   434   751   879   934  1160  1163  1283  1592
1606  1637  1703  1707  1723  1824  1958  1964  1994    40    59   187
359   443   506   939   994  1018  1317  1338  1360  1450  1791  1869
1914  1928  1968  1983  2167  2175  2186  2189  2200  2278  2343  2386
2397  2414  2416  2448  2456  2500];
% [36   872  1660  1035   539   385  1574    75   694  1596   191  1293
963  1389  1487   562  1703  1775   859  1037  1738   366  1321   232
816  1632  1806  1320   864  1447   397   707   685    89   265   625
1609  1867   132  1201  1884  1781  1848  1307  1875  1400    78   165
1329  1319   341  1624  1883  1584  1672    90  1715  1086  1905  1232
79  1439   218  1169  1906   994   997  1837  1478  1614    37  1066
1430   825  1228   808   938   179   540   599    42   993  1556  1065
880  1729  1787  1790   412   754  1578  1591  1652  1490  1678   901
1297  1613  1653  1791   196   581   582   755   101   180   913   990
578   498   617   948   320   143   683  1444    29   297   473   895
937  1039  1424  1432  1595     2    74   117   134   220   228   276
298   364   387   479   704   801   885   929   942   992  1031  1465
1509  1565  1716  1802  1845  1851  1862  1917     3     5   373   379
517   545   551   614   622   624   736   762   777   802   836  1411
1429  1476  1517  1521  1640  1704  1707  1835  1852];
% [3365   927  2032  3311  1640  2645  3558  1644  2180  3306  1012   495
678  2892  2949  2575  3637  3305  3135  1375  1735  2248   434  2090
3550  2697  3395   194  2537  3088  1899   671  3112  3237   967    71
2816   468  3099  3478   310   392  2111  1704  2247  2632  3577  3695
2847  2584  1498   700   433   596  2384  1296  2878   179  1400   636
2246  2651  1758  1345  1833  3066  3059   698   870  3403  2586  1886
1799    32  2549  2181  2631  2482  1152   404  3633  3422  2806  1972
504  1697  1396  3302  3284  3226  3097   623   899  3381  1202  3282
2688   663  3332     5  3442   641  1456   413  3720  1197   600    85
3140  3354  3057   383  2524  3310  2854   724  1850   443  1816   374
372  3694  2255  2944  2223   170  1776  1880  1014   384  1987   369
1477  3050  1018    35  3273   941  1760   169  1929  3581  1564  3068
2571  2805  3437   112   258   322  2276  2573  2574  1312  3182  1149
2727   106   429  1482  3364  3536  1877  3432   540  1385   330  3041
3188   319  1821   226   321  2526  2711  3638  3673   105   180   333
643  1304  1896  2625  3380  3713  3741  1464  2081  1849  1299  3326
```

```
3367  3535   403   421   739   812  1182  1747  1805  1957  2383  2956
3271  3471  3560  3626  3703   972  1029  1084  1543  1783  2667  3230
3602  3699   814   925  3416  3503   815  2672  3213   496  1744  1871
2182  3080  3347  3719  3748  3750   406   619   770  1191  1212  1509
1621  2091  2101  2208  2376  2654  2957  3003  3130  3148  3398  3518
3668  3678     1    20    55   275   457   471   558   590   658   834
919  1270  1329  1344  1430  1694  1724  1940  2184  2206  2404  2716
2853  2903  2925  2973  3021  3070  3274  3419  3516  3714   120   176
199   246   364   475   517   536   585   589   902   957  1020  1391
1454  1530  1590  1964  1971  2313  2321  2359  2385  2391  2394  2430
2511  2570  2678  2739  2840  2899  2901  2937  2945  2958  3329  3348
3515  3659];
% [192   424  1420  1706  2408   402  1261   745   844   499  1740  1663
1164   815  1342  1030  1844   896   936  1302  1902  1295  1711   837
2131   811   715  1677  1565  1709   621  2312  1664   337    98  1060
1301   417   584  2584  1278   313  1634  1538   355  1561   565  2081
1549   396  1202   133  2012  1807  1417  1013  1614  2338  2568   569
2241  1139   767  1461  1790   758  1454   629  1413   178   244   682
1776  2493   721  2326    73   453   330  1281   747   129  1739  1184
2084   886  1550   255   441  2022   368   319   930  1214  1901    64
1563   583   568  1277  2001  2583  1089  1857   519  1406   105  1289
2262  2457  2570   751  1129  1405  1695   906  1596   835  1169  1936
2141  2223  2305   120   502  2129   174   208   298   671    52   114
132   422   698  1517  1592    83   454   833   867   978  1462  1502
1884  1954  1996  2308  2389   221   961  2572   592  1000  2450   312
431  1929  1947  2464   270  1168  1239  1882  2186   803  1678  2386
2407  2449    61   110   181   288   344   567   638   851   965   977
1314  1363  1566  1813  1907  2034  2105  2171  2212  2325  2412  2453
2540   136   143   339   577   828  1019  1163  1255  1320  1442  1465
1636  1645  1757  1796  1804  1889  1890  1903  1969  1970  2002  2178
2588];
% [2905   168   366  3179   308  3365     6   504     8    50  1352  3081
1250   581  2602  2689  2857   199    71  1591   742  1703   428  2319
2189  1700  3117  1803  2412  3121  2819  1855  1034  1418   282  2135
930  3299  3044   217  2902    96   156  3344  1224  2675  1696  1953
1223  2572  1396   602   874  2929  2993  1431   798  2792  1862  1138
2072  1190  3195  1540  1804  1183   622  2431  1272  1322  2968  1167
2022  3056  1104  1982  2903  2362  3399  3320  1137  3282  2086  1279
1468   762  2112   338  1687  3261  3398  2486  2879  1172   514   122
458  3386   256  3005  1761   452  1607  1298  3390  2498  3375  2718
1686   579  2191  1827  2452  1301   521   822  2507  2654  1432   973
3240   606  3094  2330  3108   314  2185   852   810   840  2891   787
2843  1699  3314   508  1588  2063  2411  2799   872  1512  1531  3058
236  1532  3353  1255  1975  1791    41   337   451   694  1399  2144
2592  2648  3047  2364  2443  2908  3178  3243   421  1154  2349  2583
3181   739  1628  2884  2886  3408   216   499   639   935   976  1205
2539  2632   163   184   641   983  1049  1201  1774  1905  1999  2071
2548  2734  2769  2881  3441  2365  2865  3007  1159  2517  3191   862
1203  1257  1510  2234  2260  2374  2606  2910   114   532   572  1032
1578  1590  1629  2270  3017  3177   212   227   542  1462  1822  1911
2130  3116  3391   334   371   411   433   438   496   577   943   970
980  1066  1128  1319  1426  1514  1553  1603  1677  1805  1899  2028
2228  2265  2407  2410  2512  2577  2876  2911  2962  3041  3048  3212
3313  3331  3407     5    17   100   150   200   218   298   317   590
691   713   753   823   842   936  1187  1498  1890  2123  2254  2435
2649  2681  2823  2830  2875  3124  3259  3348  3385  3414];
% [2009   545   454  2351   745  2824  2703  1608  3316  2790   422    12
2170  2572  1673  3151  2034  2876   113   565  1728  1838  3404  1823
```

```
1297   452  1700  1093  1328   501  3170  1430   896   837   186  2311
1769  2666   840   255   195  1866  3153   107  1828   508  2863  2770
346   143  1632  2188  1916   374  1135  2461  1351  3289  2238  2180
2223  1873    93   282  1832  2551  2472   871    34   573   303  3314
2388  1365  2352  1342  1107  1122  2100   693  3386  2643  2242   730
856  1273   773  1271  1861  1971  3072  1646  2642  2433  2150   520
319  1330  2200  3319  2838  2392   877  1475  3036  2087  2932  1983
1663   696  3286   678   295  1894  2522  2179  1306    78  1397  2032
1941  3085  2719  1900   463  3265  2357   522   483   385  2000  1674
2381   141  2020   768  2803   819  1724  1919  2982   384  2521  3111
291   668   744   770   850  2455   289   534  2605  2668  3402   217
127   430   568  1697    32   451   530  1041  1151  1688  1726  2387
2843  3306    20   985  1287  1300  1792  2491  2820  1047  1265  1321
1447  1715  3073   450   728  1016  1145  1222  1275  1375  2637  2848
3141  3359  2828   608   256   659  2922  1037   264   574   731  1026
1034  1067  1213  1423  1510  2002  2120  2126  2156    27   178   181
457   599   885   924  1142  1266  1357  2159  2353  2386  2640  3211
3249  3422     1    36   230   271   292   434   662   864   882   913
919   947  1134  1215  1230  1439  1514  1587  1629  1643  1777  1839
1911  1980  2003  2039  2224  2227  2604  2617  2740  2762  2786  2872
2903  2946  2953  3014  3030  3068  3082  3089  3262  3330    15    28
68   239   477   865   894  1033  1121  1140  1141  1194  1282  1315
1326  1484  1490  1707  1750  1805  1835  1956  1987  2103  2454  2596
2650  2657  2665  2690  2735  2736  2812  2897  2924  3152  3243  3407];
% [2851  1148   870  2530  3001  3213  1061  1802  1911  2909   331  1463
2362  3245   693   773   691   349  2155   424  2124   285  1815  2738
2656  2642   116  2055   997   807   547  2058  2512  1073  1880  1348
2164  2897   333  2803  1708  3124  3062  1147  2543  3127    59  3225
1115  1874   925  1637  2802  3002  1639  1579  1489   940  1733  1499
1477    67  1018   894  2247   437  2545  2183   568  3215   838  3138
2474  2099  3200   769  1944   999  2248   806  2323   265   715  1314
31   592  2077  2428  3301  3110   442  2367   770  1218   602   643
539  1730  2270  1713  1895  2644  1761   533  1296  1987   846  2927
414  1250  1800  2031   294  1371  3320  3316  1130  3363  1538    55
1275  2718   260   355  1129  1942  3317  1647  1727  2731   488   705
987   259   417  1612  2159  1340  1630  1958  2730  3183  3268    32
421   505  2775  2869  2967  3072    82  1472  1672  2033  2199  2797
2918   831  1460  2120  2237  2301  3123  3265  3047  1979  1201  3077
396  1985  2528   169   452   893   958  1035  1665  1853  2269  3093
217   361   517   615   652   753   822   953   964   981   984  1059
1269  1292  1397  1514  1711  1813  1951  2156  2172  2210  2275  2276
2420  2468  2600  2678  2700  2706  2715  2737  2777  2853  2999   115
252   422   465   509   549   563   569   614   743   799   832  1057
1127  1157  1181  1224  1227  1338  1398  1402  1420  1530  1551  1626
1724  1738  1799  1806  1818  1841  1854  1920  1924  2103  2121  2153
2160  2173  2315  2426  2448  2551  2577  2727  2798  2823  2825  3017
3068  3089  3114  3148  3280  3315];
% [1614   773  1651    95   759   431  2141   183  1001   201  2083   373
933  1531   170  1362  1656  1683  1748   414  1087  1980  1942  2408
2217  2108   784   413  1745  1516  2514   855  2264   812   887  2629
2086  2129  1520  1694   274   217   844  1321  2117   499  2465   577
2458   906  1361   384   249  1403  2481  1583   562  2173  2061  2130
2371  1630  2581   953  1592   365  1994  2427    62  1886  1695  1123
1006  2313  2596  1599  1317  1131  2650   181   766  2066  1420   424
1682  2166  2220  2222   318  2295  2349  1591  2218  2023  2600  1570
2286  1184   652  2346  1262  2536  2283   288  2647   393  1566  1612
2590   846   998  1154  1567  2309   331  2244  2578   358   816    37
761   934  1027  1226  1536  1738  2523  2649  1346  1382  2192   187
```

```
272  1097  1104  1255  1351  1401  1487  1491  2213  1515  2639   625
896  1021  1270  1360  2338  2367  2634  1565  2102   917  2528  2646
145   591   785  1067  1853  2112  2211  2347  2549  2652    47   195
350   362   609   612   635   707   712   724   749   768   779   994
1007  1008  1092  1136  1140  1410  1479  1525  1557  1561  1605  1679
1746  1929  1954  1985  2105  2194  2274  2556   130   347   399   697
711   760   786   899   905  1009  1098  1134  1205  1269  1320  1364
1381  1396  1427  1462  1502  1836  1937  2008  2128  2356  2391  2407
2416  2460  2471  2568];};
% SCORES=11;
% ns={[14    22    32    34    83   106   149   167   181   216   272
294   309   327   329   336   353   358   370   387   388   400   409
420   467   477   549   608   640   704   707   716   723   768   791
792   842   867   891   893   918   923   934  1057  1059  1094  1140
1153  1164  1166  1177  1200  1213  1215  1222  1311  1329  1353  1357
1419  1422  1426  1468  1493  1507  1520  1555  1560  1565  1586  1595
1600  1603  1609  1619  1722  1724  1768  1792  1833  1873  1874  1885
1895  1903  1953  1957  1990  2007  2027  2037  2075  2079  2090  2138
2140  2179  2188  2204  2210  2225  2232  2289  2302  2311  2314  2366
2413  2432  2485  2528  2534  2557  2571  2573  2594];[1     36    52
68   140   186   245   296   323   324   337   367   417   428   436
486   523   524   535   608   615   623   640   702   725   734   740
764   776   789   794   796   808   846   859   861   863   866   867
881   882   916   941   950   972  1024  1051  1052  1064  1065  1066
1094  1113  1139  1172  1185  1186  1202  1205  1249  1253  1277  1310
1313  1314  1321  1358  1365  1372  1409  1410  1421  1453  1491  1512
1652  1664  1670  1675  1685  1687  1726  1814  1819  1839  1871  1876
1879  1920  1926  1948  1950  1977  1991  2015  2037  2046  2054  2079
2089  2096  2097  2123  2129  2141  2143  2196  2250  2280  2330  2356
2363  2367  2431  2456  2474  2511  2523  2545  2546  2564  2570  2590
2678  2691  2706  2735  2736  2769  2774  2788  2806  2839  2840  2841
2863  2877  2890  2927  2947  2965  3002  3078  3085  3118  3157  3158
3161  3174  3181  3213  3254  3258  3309  3411  3425  3437  3446  3468
3497  3525  3535  3565  3601  3626  3636  3640  3659  3702  3801  3809
3815  3816  3838  3840  3844  3869  3876  3889  3907  3908  3975  4009
4017  4061  4069  4071  4077  4082  4092  4102  4110  4151  4185  4186
4197  4203  4258  4284  4297  4300  4315  4320  4434  4494  4533  4542
4546  4566  4574  4577  4598  4622  4649  4684  4725  4773  4788  4805
4862  4869  4889  4901  4908  4917  4928  4977  4991  5008  5017  5107
5112  5121  5125  5127  5139  5151  5162  5175  5210  5235  5255  5274
5280  5327  5333  5360  5398  5416  5442  5470  5488  5512  5570  5573
5574  5678  5690  5701  5721  5727  5732  5797];[117   181   260   261
292   323   325   343   366   405   450   453   471   494   613   647
663   675   690   705   717   729   750   756   770   792   876   888
904   939  1020  1024  1038  1046  1056  1077  1127  1148  1163  1164
1193  1209  1244  1299  1318  1350  1354  1369  1426  1460  1474  1491
1492  1506  1519  1615  1627  1668  1670  1677  1689  1693  1697  1719
1759  1805  1823  1824  1842  1858  1864  1910  1912  1961  1977  2030
2031  2049  2051  2056  2124  2202  2249  2251  2272  2321  2346  2397
2436  2444  2457  2484  2535  2620  2680  2697  2755  2780  2793  2815
2820  2829  2836  2859  2866  2879  2884  2910  3020  3031  3035  3075
3089  3093  3101  3162  3199  3223  3235  3252  3271  3311  3324  3349
3360  3384  3393  3405  3419  3425  3472  3481  3506  3513  3547  3577
3615  3630];[17    22    41    43   158   160   163   252   342   343
412   436   465   490   491   516   541   570   597   670   672   673
689   736   763   783   823   940   962   968   985   987  1020  1055
1079  1094  1128  1140  1157  1163  1181  1182  1262  1265  1272  1285
1289  1309  1320  1321  1337  1348  1374  1443  1444  1457  1458  1508
```

1537 1557 1566 1591 1611 1657 1659 1684 1691 1713 1718 1731
1732 1743 1812 1815 1824 1878 1888 1924 1958 1961 1971 2010
2017 2021 2041 2059 2095 2097 2098 2100 2106 2110 2137 2163
2178 2219 2229 2233 2264 2292 2298 2428 2429 2460 2503 2631
2639 2650 2666 2676 2701 2702 2715 2805 2806 2813 2859 2893
2894];[15 40 47 68 109 120 142 178 180 274 281
290 299 321 335 341 366 367 369 387 417 440 442
472 489 507 518 519 525 532 544 558 570 573 581
583 602 605 609 635 639 655 675 676 707 727 751
776 783 799 800 814 839 844 880 914 929 943 1012
1038 1054 1071 1073 1090 1104 1138 1163 1224 1231 1289 1301
1302 1336 1340 1342 1353 1408 1456 1513 1515 1529 1537 1575
1601 1615 1616 1627 1644 1646 1665 1678 1736 1793 1805 1827
1837 1887 1901 1912 1940 1950 1967 1978 2001 2008 2012 2019
2024 2068 2186 2235 2243 2266 2286 2335 2336 2343 2357 2364
2377 2416 2417 2432 2442 2468 2476 2501 2535 2536 2544 2587
2593 2596 2629 2789 2804 2822 2848 2927 2944 2965 2979 2988
3026 3029 3048 3068 3073 3113 3114 3125 3132 3153 3162 3166
3169 3184 3188 3203 3244];[52 61 98 111 134 135 180
186 267 275 278 292 317 452 453 455 507 522 525
585 638 651 665 667 708 725 743 770 772 778 789
807 847 861 905 908 960 961 962 963 993 1012 1015
1047 1053 1099 1115 1118 1222 1259 1285 1299 1338 1385 1411
1430 1440 1466 1476 1482 1500 1521 1527 1541 1596 1715 1728
1733 1737 1742 1750 1751 1754 1779 1819 1836 1853 1905 1943
1945 2050 2140 2166 2174 2190 2193 2252 2284 2319 2333 2421
2435 2463 2471 2472 2497 2559 2568 2580 2581 2589 2600 2602
2613 2631 2643 2651 2671 2704 2721 2722 2731 2733 2734 2763
2764 2765 2799 2800 2854 2886 2902 2903 2908 2914 2933 2939
2952 2961 2978 2982 3033 3041 3071 3087 3112 3121 3127 3168
3228 3333 3348 3374 3376 3381 3428 3441 3448 3449 3450 3462
3510 3539 3540 3631 3661 3724 3728];[28 96 99 102 105
106 147 192 212 253 256 334 340 370 374 423 490
491 548 562 570 572 588 593 609 614 629 636 655
663 694 714 743 745 788 791 885 886 929 967 968
969 971 1049 1069 1084 1099 1106 1110 1152 1175 1228 1256
1317 1320 1321 1370 1475 1482 1501 1506 1555 1579 1592 1607
1615 1638 1670 1701 1717 1742 1756 1762 1770 1799 1847 1852
1867 1868 1876 1911 1915 1924 1968 1973 1992 1994 2012 2038
2117 2287 2295 2347 2390 2407 2432 2441 2508 2512 2520 2536
2543 2544 2570 2589 2590 2607 2646 2647 2669 2683 2772 2780
2796 2802 2833 2876 2886 2898 2920 2956 2986 2988 3000 3017
3033 3043 3059 3094 3097 3105 3123 3147 3165 3249 3254 3341
3346 3383 3413 3447 3464 3466 3475 3520 3522 3528 3533 3539
3557 3608 3709 3713 3732 3733 3741 3743 3781 3834 3839 3863
3868 3919 3927 3941 3964 4011 4032 4070 4102 4108 4128 4146
4211 4230 4234 4240 4243 4260 4287 4308 4328 4408 4415 4416
4455 4457 4464 4466 4485 4488 4527 4589];[19 45 54 73
93 105 256 300 306 311 349 399 401 428 436 441
445 478 487 515 516 521 525 564 567 581 582 642
681 684 702 717 732 752 820 826 834 859 897 903
923 924 943 952 982 987 1014 1064 1077 1078 1101 1115
1139 1149 1151 1184 1191 1221 1226 1236 1255 1262 1286 1312
1329 1372 1408 1416 1449 1473 1481 1499 1527 1538 1569 1623
1684 1697 1719 1725 1740 1801 1811 1845 1921 1963 1992 2001
2003 2013 2031 2033 2041 2043 2056 2063 2079 2087 2126 2129
2145 2207 2214 2230 2253 2273 2281 2302 2312 2326 2374 2375
2398 2406 2412 2433 2441 2545 2554 2583 2589 2597 2609 2613

2658 2681 2685 2693 2728 2748 2770 2778 2797 2868 2956 2987
3003 3008 3034 3041 3063 3069 3070 3111 3128 3174 3181 3195
3216 3217 3218 3233 3280 3288 3299 3307 3309 3320 3337];[15
79 114 151 175 190 219 249 272 278 305 314 320
367 418 422 431 437 442 459 462 477 491 502 516
542 553 573 667 683 695 708 721 786 796 808 838
876 890 897 975 979 1010 1033 1034 1067 1069 1095 1129
1142 1145 1146 1163 1180 1185 1198 1259 1272 1297 1325 1326
1327 1328 1343 1374 1446 1450 1482 1515 1548 1567 1580 1584
1613 1614 1617 1632 1682 1698 1708 1732 1737 1738 1748 1751
1767 1774 1851 1863 1907 1927 1934 1996 2021 2033 2038 2096
2152 2209 2238 2247 2248 2251 2298 2343 2366 2371 2377 2400
2451 2494 2536 2539 2563 2572 2600 2631 2639 2640 2731 2732
2741 2848 2879 2884 2891 2900 2906 2943 2948 2970 2995 3001
3062 3098 3108 3173 3212 3227 3233 3281 3289 3338 3340 3347
3367 3372 3425 3462 3482 3497 3523];[17 41 58 79 80
92 120 135 228 251 258 277 281 303 326 327 336
370 372 373 419 422 449 483 551 611 642 666 691
746 748 749 750 760 771 803 826 841 853 881 888
893 894 917 940 976 988 1023 1027 1031 1057 1088 1126
1137 1202 1213 1235 1252 1286 1315 1319 1337 1349 1363 1364
1463 1481 1489 1537 1572 1587 1593 1594 1609 1663 1666 1694
1712 1726 1743 1826 1827 1838 1876 1880 1910 1939 1961 1986
1999 2001 2003 2016 2066 2090 2129 2152 2171 2173 2187 2255
2272 2321 2359 2383 2384 2413 2452 2460 2473 2497 2567
2586];[70 127 128 153 155 188 219 220 222 225 231
234 236 239 244 261 262 263 264 283 319 365 430
436 458 492 496 502 523 578 583 601 636 645 647
652 667 677 700 715 740 770 781 826 831 875 886
917 935 937 938 1002 1048 1074 1085 1086 1125 1160 1215
1310 1362 1373 1413 1423 1432 1433 1435 1513 1521 1555 1573
1604 1631 1813 1842 1847 1901 1902 1934 1953 2005 2011 2023
2114 2121 2196 2214 2262 2321 2344 2347 2361 2383 2392 2450
2463 2515 2544 2547 2553 2555 2571 2612 2619 2627 2659 2719
2809 2847 2929 2940 2974 2980 2995 3019 3039 3059 3106 3175
3195 3271 3272 3311 3336 3343 3347 3376 3385 3471 3482 3537
3560 3565 3588 3590 3596 3599 3646 3660 3677 3699 3734 3740
3768 3787 3917 3935 3961 3999 4025 4058 4069 4103 4104 4112
4119 4154 4178 4189 4204 4207 4348 4353 4368 4378 4389 4461
4462 4463 4466 4475 4515 4562 4568 4574 4581 4655 4662 4675
4678 4802 4812 4831 4835 4844];[7 16 37 54 93 113
149 150 152 164 220 261 270 275 279 291 294 331
358 427 451 509 518 529 536 547 548 575 589 613
633 688 755 776 801 805 878 889 910 915 933 937
962 993 996 1096 1105 1110 1117 1137 1153 1168 1178 1205
1214 1263 1274 1282 1336 1344 1396 1402 1405 1426 1433 1444
1470 1518 1533 1539 1540 1593 1594 1645 1658 1660 1661 1699
1707 1728 1736 1738 1819 1823 1833 1837 1887 1902 1907 1908
1911 1912 1933 1974 1999 2005 2007 2030 2111 2139 2175 2181
2203 2206 2217 2218 2256 2267 2294 2340 2383 2399 2413 2468
2475 2503 2573];[36 40 43 82 85 137 153 213 258
342 351 438 477 497 552 554 595 608 617 618 653
664 665 714 764 777 783 784 857 899 900 908 920
923 925 935 936 940 970 972 995 1003 1010 1047 1074
1118 1143 1154 1159 1172 1176 1202 1214 1235 1237 1238 1245
1275 1293 1357 1384 1404 1450 1521 1534 1562 1618 1632 1658
1659 1676 1715 1742 1748 1798 1836 1859 1878 1886 1893 1912
1934 1938 1944 2009 2032 2220 2229 2335 2337 2348 2403 2414

2421 2435 2436 2439 2443 2444 2475 2481 2514 2533 2534 2596
2614 2625 2653 2667];[1 52 54 57 108 127 161 179
197 199 232 258 316 341 399 443 506 526 529 531
594 669 713 722 741 752 756 811 832 852 874 876
879 921 925 926 934 939 955 956 957 961 972 981
982 1008 1032 1061 1072 1079 1100 1160 1163 1177 1200 1201
1222 1283 1318 1319 1338 1348 1397 1398 1445 1446 1480 1540
1551 1572 1605 1633 1637 1703 1716 1717 1723 1779 1786 1803
1822 1868 1882 1887 1889 1897 1947 1964 1994 2033 2037 2068
2131 2203 2216 2277 2283 2284 2299 2351 2364 2439 2463 2472
2483 2518 2524];[36 38 71 74 75 78 79 90 117
176 179 191 218 232 265 298 341 364 366 381 385
387 397 461 479 539 559 562 582 694 723 806 816
825 840 864 872 885 929 963 992 1012 1035 1037 1065
1066 1086 1168 1169 1170 1201 1247 1257 1262 1293 1307 1320
1329 1389 1434 1447 1448 1463 1465 1478 1487 1510 1565 1574
1584 1596 1610 1615 1632 1660 1703 1716 1738 1775 1781 1806
1845 1848 1862 1875 1883 1884 1911 1917];[5 20 35 170
179 180 194 200 275 310 319 322 358 369 372 383
384 392 413 432 433 434 435 443 468 471 495 558
596 636 658 671 678 698 700 724 771 870 899 902
919 927 941 967 976 1012 1018 1068 1149 1151 1174 1202
1270 1296 1312 1329 1344 1345 1375 1396 1400 1464 1503 1586
1630 1640 1644 1704 1726 1735 1741 1758 1805 1833 1880 1886
1899 1929 1972 2011 2032 2037 2090 2091 2122 2174 2180 2181
2182 2206 2246 2247 2248 2255 2338 2384 2404 2524 2537 2549
2575 2584 2586 2618 2631 2632 2637 2645 2647 2651 2688 2697
2805 2806 2847 2873 2892 2925 2949 2968 3021 3059 3066 3068
3070 3088 3099 3112 3135 3203 3216 3237 3274 3284 3305 3306
3311 3330 3354 3365 3395 3403 3432 3437 3478 3486 3516 3550
3558 3577 3581 3597 3633 3637 3695 3719 3720 3727];[64 98
102 129 136 178 192 288 313 330 337 355 368 396
402 417 423 424 431 438 454 499 529 569 577 583
584 621 629 638 652 670 682 715 721 745 758 767
792 811 815 844 851 896 936 991 999 1013 1030 1044
1060 1081 1164 1202 1214 1261 1278 1295 1301 1302 1314 1363
1405 1411 1420 1433 1489 1541 1546 1563 1601 1634 1663 1664
1677 1706 1711 1713 1739 1740 1776 1790 1804 1813 1844 1885
1902 2022 2034 2058 2081 2105 2129 2131 2145 2170 2209 2212
2224 2312 2325 2408 2429 2450 2453 2493 2517 2528 2540 2568
2575 2584];[2 6 8 50 71 156 168 199 256 282
299 308 335 366 381 421 428 458 496 504 514 521
579 581 622 636 639 721 742 798 835 874 930 936
943 973 1025 1034 1104 1128 1138 1167 1172 1187 1190 1223
1224 1250 1319 1348 1352 1418 1426 1468 1497 1553 1558 1578
1591 1637 1677 1686 1703 1761 1778 1792 1804 1805 1827 1842
1862 1869 1933 1936 1953 1975 1976 1982 2000 2008 2022 2086
2112 2135 2189 2319 2330 2407 2412 2486 2507 2516 2572 2595
2602 2675 2689 2717 2792 2819 2845 2857 2881 2886 2903 2905
2911 2959 3005 3007 3028 3044 3056 3058 3081 3092 3094 3103
3117 3167 3179 3240 3252 3282 3299 3313 3320 3344 3353 3365
3386 3398 3399 3407 3413 3426 3443];[4 12 36 78 113
143 181 186 195 213 255 289 292 295 306 319 346
374 379 422 452 454 459 483 501 508 522 534 540
545 565 662 688 697 719 730 744 745 749 764 768
837 851 856 864 896 1107 1112 1122 1134 1194 1215 1245
1297 1315 1342 1351 1430 1462 1539 1551 1579 1586 1608 1612
1643 1673 1674 1700 1707 1728 1769 1779 1823 1828 1830 1834

```
1838  1839  1854  1861  1866  1873  1911  1916  1941  1974  1983  2001
2009  2032  2034  2087  2150  2170  2180  2188  2223  2227  2286  2289
2305  2311  2340  2351  2381  2386  2392  2461  2472  2572  2641  2666
2703  2739  2740  2756  2786  2790  2824  2843  2863  2876  2890  2903
2909  2932  2946  2953  2958  3014  3043  3068  3072  3080  3089  3151
3163  3170  3214  3316  3319  3330  3386  3404];[72   107   120   246
252   256   260   265   285   331   333   349   414   417   424   517
539   547   602   611   615   628   643   651   652   660   693   715
753   769   773   806   807   822   870   925   940   981   997   999
1015  1018  1057  1061  1073  1147  1148  1193  1248  1320  1325  1348
1451  1463  1477  1499  1514  1546  1547  1579  1637  1639  1669  1674
1708  1711  1713  1761  1802  1817  1832  1874  1875  1880  1895  1903
1911  1920  1924  2055  2058  2099  2118  2124  2126  2144  2155  2156
2183  2205  2269  2362  2377  2428  2476  2512  2516  2530  2533  2535
2543  2642  2656  2678  2700  2728  2730  2764  2777  2799  2803  2851
2869  2897  2909  2927  3001  3017  3027  3124  3126  3127  3138  3200
3211  3213  3225  3245  3268  3311  3320  3363];[47    95   128   150
170   183   184   201   214   274   331   373   406   414   424   431
434   466   475   562   568   609   611   635   707   724   759   768
773   779   784   799   829   844   846   887   896   906   933   934
945  1001  1008  1087  1092  1107  1123  1193  1255  1280  1281  1321
1362  1403  1410  1420  1515  1525  1531  1569  1583  1614  1651  1656
1683  1694  1745  1746  1748  1832  1875  1937  1954  1962  1980  1983
1985  1994  2038  2083  2086  2108  2117  2129  2141  2166  2167  2173
2217  2222  2228  2264  2274  2309  2313  2321  2346  2371  2408  2458
2481  2499  2514  2536  2568  2578  2596  2629  2647];};
%
% SIX=0;requireAll = 1; allowSubstring = 1; ADJACENTONLY = 0;
% % % 'conjugal'
% CONTEXT = 2;
pat={{'conjugal';'SCO6880';'TrbL';'DUF8175';'DUF6112';'MinD';}; {'helix-
turn-helix domain';'NlpC';'FAD oxidoreductase'}};
% % % 'septum'
% CONTEXT = 1; pat={{'conjugal'};{'septum';};};
% % % 'HAD-IB'
% allowSubstring = 0; pat={'HAD-IB hydrolase';'HAD-IB phosphatase';};

% % these are old, messy anns in '/mnt/spin/anns_Bacteria_sa'
% ANNS_SA = '/mnt/spin/anns_Bacteria_sa';
% SWITCH='for_Bifi';
% genus='Bifi';
% d = '';
%
gs={'igrs_Bifi_adolescentis';'igrs_Bifi_angulatum';igrs_Bifi_animalis';'i
grs_Bifi_bifidum';'igrs_Bifi_dentium';
%
'igrs_Bifi_gallicum';'igrs_Bifi_longum';'igrs_Bifi_magnum';'igrs_Bifi_min
imum';'igrs_Bifi_pseudolongum';};
% % requireAll = 1; allowSubstring = 0;
% % % pat={'hydrolase'; 'dehalogenase'; 'phosphatase';'morphological
differentiation';'hexosaminidase';'glucosidase';
'glucosidase';'glycosidase';...
% % %
'oxidoreductase';'Fic';'oxidase';'dehydrogenase';'reductase';};
% % % pat={'hydrolase';'pullulanase'; 'dehalogenase';'galactosidase';
'phosphatase';'morphological
differentiation';'hexosaminidase';'glycosylase';'glucosidase';...
% % % 'glycosidase';'Fic';};
```

```
% % pat={'HAD hydrolase';'glycosyl hydrolase'; 'glycoside
hydrolase';'glycosylase';'glycosylhydrolase';'glycoside
hydrolase';'dehalogenase';'galactosidase';...
% %    'phosphatase';'hexosaminidase';};
% % % CONTEXT = 2;pat={{'oxidoreductase';'Fic';}; {'hydrolase'}};
% % CONTEXT = 2;
% % pat={{'HAD hydrolase';'glycosyl hydrolase'; 'glycoside
hydrolase';'glycosylase';'glycosylhydrolase';'glycoside
hydrolase';'dehalogenase';'galactosidase';...
% %      'phosphatase';'hexosaminidase';};
{'transporter';'transcriptional';'regulator';'oxidoreductase';'Fic';}};
% % SIX=0; ADJACENTONLY = 1;requireAll = 1; allowSubstring = 0; pat={'HAD
hydrolase';};
% SCORES=[1 2 3 4 5 6 7 8 9 10 11 12 13 14];
% ns={[8   41   72  113  145  156  189  203  216  269  297  316  353  394
414  454  464  492  505    2   73  109  112  257  478  488   63  139  142
294  331  354  432  499   42   65  104  105  128  185  196  228  251  277
293  389  453  500  517  531   98  166  179  356  364  430  437   37  449
29   59   68   89  130  255  298  338  346  347  366  446  458  511   56
233  274  289  450  452  474   38   84  388   82  199  510  273  537  403
424  295  509];[16   27   28   29   66   88   90  147  184  215  251  254
312  336  342  362  390  112  255  260  326  361  386  126  371    3   60
84   93  118  119  125  131  133  161  163  299  324  365   54  100  157
333  359   78  122  328  239  253  288  325  392  395  308  311  159  160
380  259  302  216  217  158];[5    6   20   33   98  107  144  160  181
193  220  230  238  245  255  269  332   47  262  322   44  119  176  232
257  280  366   32   57   65  113  204  309  314   93  120  213  219  223
374    7   67   75  215  233  274  276  310  224  231  246  252  324  337
340  270  189  235  221  379  256  249];[36   46   65  111  146  148  164
215  239  304  309  310  323  327  400  447  451  152  260  298  441  104
227  283  329  378  412   23   48   70   97  101  103  137  145  238  284
302  322  330  355  390  407  422  423   76  290  297  416  276  134  147
176  226  268  333  398  428   12   53   98  218  212  228  240  175  179
321  301  365  187  461  273];[25   26   52   63   89  100  104  106  153
213  238  239  249  278  281  325  326  358  387  390  404   68  103  252
388  205  383  139  209  264  284  348  376   97   98  227  251  309  374
412   79   81  260  274  407    8   80  151  210  258   32  384   20  105
161  389   12   46   67  386];[48   59   81  101  107  122  183  204  238
244  246  255  271  276   60  181  187  221  326  149  304  324   68   78
166  170  191  214  216  243  270  292   85  121  126    8   70   47  105
281  127  203  256  167];[2   53   91  101  128  130  133  137  166  212
217  240  249  290  294  303  304  328  331  381   37   76  124  160  291
306  362   33   52  125  181  183  192  228  355  383  384  385  390  414
78  200  250  327  332  376   79  121  148  203  346   23   93  114  276
279  356  218  204  397  280  266  409];[13   56   73   81   92  100  196
228  260  264  267  314  322  335  339  342  344  370  423  445  447  456
502  504  516  537  588  621  634  668   22  188  265  307  336  421  523
664  108  176  256  341  477  521  539   66   74  122  135  152  175  240
431  443  484  492  503  511  638  653  103  166  229  286  469  568   99
185  232  266  268  276  288  305  393  395  466  519  630  636  646   85
97  123  243  263  343  427  459  465  499  500  514  654  131  315  396
410  411  501  351   16   59  205  340  365  400  478  513    9   55  182
51  515  524  369  483];[19   33   84   85  123  196  200  258  353    3
62   68  100  213  365  368  370   72  217  278  315  339   14   15   30
187  218  282  330  359   42  103  264  267  269  340  361   58   81   97
214  228  318  345  373   48  238  319  369  227  120  316  367];[3    5
6   70   75   85  105  170  262  336   86   93  110  263  179    8   57
94  137  196  205  225  236  251  261  315  317  106  312   33   52   69
```

```
84   97  108  119  273   43   83  107  188  242  289  313  302  311  128
204  318  290   76  115  136];};
% SIX=0; requireAll = 1; ADJACENTONLY = 0;
% % allowSubstring = 1; CONTEXT = 2; pat={{'CpaF';}; {'RNA degradosome
polyphosphate kinase';}};
% % allowSubstring = 1; CONTEXT = 1; pat={{'CpaF';};{'AAA family
ATPase';'ATPase';'P-loop NTPase';'hypothetical protein';'septum';};};
% allowSubstring = 0; pat={'CpaF';};
%
% ANNS_SA = '/mnt/spin/anns_Bacteria_sa';
% genus='bifi';
% SWITCH='for_Bifi';
% SUFFIX = '_Dh_Sco';
% d='';
%
gs={'igrs_Bifi_adolescentis_Sco/';'igrs_Bifi_angulatum_Sco/';'igrs_Bifi_a
nimalis_Sco/';'igrs_Bifi_bifidum_Sco/';'igrs_Bifi_boum_Sco/';
%
'igrs_Bifi_choerinum_Sco/';'igrs_Bifi_dentium_Sco/';'igrs_Bifi_gallicum_S
co/'; 'igrs_Bifi_longum_Sco/'; 'igrs_Bifi_magnum_Sco/';
%
'igrs_Bifi_moukalabense_Sco';'igrs_Bifi_pseudolongum_Sco/';'igrs_Bifi_sub
tile_Sco/';'igrs_Bifi_thermacidophilum_Sco/';
'igrs_Bifi_tsurumiense_Sco/';};
% SCORES=[1 2 3 4 5 6 7 8 9 10 11 12 13 14];
% ns={[8   12   17   62   71   82  175  218  228  305  306  345  364  407
411  433  435  457  473  486  499  502  537  583  603  637    3   53  243
361  548  648  209  290  362  422  493   19   21  105  111  189  248  253
325  333  384  389  423  467  470  491  558  588  167  289  408  465  517
404  546   61  169  180  285  315  372  519  544  563  600  162  195  210
277  308  417  458  469  363  552  653  168  439  510  567  573  633  288
346  412  420  307  436   72   14  540  276];[37   93  102  127  164  166
210  234  262  266  272  276  290  335  336  339  350  352  383  483  527
529  530  549  558  637  640  684   98  209  340  359  410  528  551  567
625  638  657  658   32   38  323  431  532   10   30  112  130  168  203
204  244  258  312  369  395  440  497  509  512  548  554  568  593  594
634  205  214  380  435  563  570  376   80  118  126  156  163  592  604
614  619  672  699  702  706   97  111  200  219  311  399  412  450  505
575  606  639  667  192  327  539  185    5   45  160  191  375  404  454
461  620  673  400   22    7  155  496  705  446];[11   19   24   31   70
72   89   93  111  126  147  217  240  266  308  366  372  438  440  480
495  545  546  553  568  660  668  671   40  474  672   33   54  127  155
205  265  312  338  497  567   73   74  102  110  142  179  204  221  256
257  278  290  307  319  337  418  498  528  554  586  593  625  675    2
32   61   64  128  167  239  243  267  396  684  333  353  411  419  512
552  144  168  351  479  530  532  573  104  255   79   91  161  601  615
112   18  607  137  289];[57   62  102  122  129  144  156  187  211  230
246  279  280  295  342  357  412  415  442  494  496  567  658  660  689
694  725  730  825  840  862  873  121  203  206  237  249  257  366  389
393  507  623  669  319  586  670  743   16   64   92  119  162  168  208
286  299  454  503  550  580  630  682  724  745  768  813  819  858  864
869   90  142  216  424  457  477  626  654  754  885  739   10   70  109
124  174  269  416  425  447  459  611  666  690  770   39   59   88  248
392  564  826  101  110  123  399  678    3  117  152  185  236  255  323
353  402  413  495  633  645  284  585  282  228  372];[63   64  110  115
128  146  160  218  293  327  332  336  338  399  418  420  424  426  527
569  625  627  638  651  670  712  736  781  878  920  935  271  458  635
726  815  849  895  898   19  449  482  574  637  709  196  243  351  460
```

477 490 541 566 572 594 609 645 647 665 675 808 836 860 914
922 139 206 214 220 311 401 407 409 410 508 593 604 676 824
839 227 580 729 863 8 28 69 75 95 100 158 287 314 319
400 521 545 571 711 845 20 45 88 93 121 148 207 208 223
263 272 281 310 325 457 491 678 850 926 340 448 505 662 890
47 286 282 290 328 330 339 428 480 669 691 903 203 412 917
626 24 812];[6 12 14 47 48 72 88 107 213 240 249 285
292 308 326 347 408 409 419 511 556 716 723 760 765 801 39
83 257 342 344 377 457 526 679 751 410 541 589 724 22 86
122 160 187 230 232 281 333 337 464 497 504 573 598 634 668
678 684 698 720 757 758 780 26 173 234 258 291 356 531 695
803 37 51 63 210 251 324 444 528 547 666 700 753 775 96
112 224 255 264 364 405 406 451 545 559 567 272 334 516 280
290 538 566 597 748 784 523 33 399 49 335 641];[9 43 116
125 126 177 219 239 243 262 267 360 371 402 406 427 440 446
449 472 473 549 577 609 633 636 670 679 702 742 746 763 2
61 87 344 348 359 642 669 774 250 515 613 12 72 224 316
317 325 340 345 353 413 480 569 631 692 695 704 705 788 214
614 175 65 86 99 212 230 283 293 411 514 573 646 671 714
715 787 161 448 455 624 706 785 781 274 326 392 525 536 649
699 24 232 583 183 534 261 396 507 297];[80 88 91 118 133
162 204 242 251 258 260 262 332 379 422 459 474 525 534 538
572 577 643 649 664 666 691 705 734 735 742 754 767 816 879
896 926 933 963 154 392 470 471 556 564 663 668 702 711 745
762 769 820 849 117 456 526 585 929 2 61 112 116 238 442
494 547 548 549 595 629 690 787 789 892 23 347 397 475 672
677 706 714 871 913 448 567 1 101 102 151 184 270 351 360
499 512 613 661 709 717 759 796 817 821 857 64 90 203 224
257 343 382 500 529 760 779 853 858 945 962 105 696 194 208
352 416 522 578 604 671 882 329 472 180 514 751 216 466
314];[42 46 66 107 154 171 191 269 270 303 326 347 354 366
382 387 389 407 424 449 483 485 563 568 591 639 640 646 692
707 727 740 747 755 767 23 52 104 192 322 412 486 508 542
635 712 96 186 360 390 776 1 47 71 100 120 213 223 349
357 474 503 613 651 667 670 675 681 714 733 777 95 111 136
146 196 410 599 756 130 39 182 184 201 271 423 461 480 520
576 589 604 665 748 780 789 63 93 103 127 153 187 195 644
683 522 541 350 79 614 147 106 43 61 124 48 249 391 521
699 25 190];[25 45 73 147 160 163 192 235 282 293 310 313
345 390 406 409 410 432 435 528 541 17 76 98 144 152 157
214 418 545 4 380 427 440 6 80 85 132 136 142 143 154
392 434 466 496 524 205 222 253 329 417 495 478 48 72 161
209 216 278 342 378 419 451 458 473 510 514 540 64 66 82
189 254 271 536 5 472 527 337 397 135 243 316 389 391 433
469 470 78 480 319 172 374];[12 29 54 75 76 104 105 165
181 184 211 220 232 236 256 281 287 290 328 344 362 398 422
455 466 484 537 653 686 217 219 384 452 481 495 514 518 587
170 206 535 545 21 36 98 130 222 229 234 330 385 395 399
438 493 555 565 592 624 630 661 669 14 93 146 187 243 482
579 616 679 10 43 136 149 264 303 321 329 343 347 364 369
396 494 659 4 39 59 61 101 141 145 162 387 441 641 646
117 186 381 389 654 64 99 323 348 688 693 263 697 660 388
380 666 342 423 691 27 161];[17 19 84 187 202 206 218 259
281 318 329 331 332 347 350 385 389 391 397 425 439 507 516
21 71 75 204 400 414 492 521 5 228 254 377 407 466 506
517 83 133 169 197 253 260 279 287 299 372 406 441 442 511
522 16 274 508 436 532 15 57 111 230 284 379 398 403 429
2 97 148 154 387 461 465 531 263 286 289 324 524 174 323

```
393   59  344  288   61];[51    86   118   142   143   195   202   234
256   269   313   349   375   391   473   486   491   498   510   572
580   606   609   622   627   638   640   698   726   767   784   807
834   838   942   948   956  1038  1079  1086  1178  1187  1219  1232
1238  1243  1258  1333  1345  1376  1381  1513    54   105   343   394
446   460   469   524   678  1124  1139  1173  1275  1337  1439  1503
30    58   207   208   214   337   509   617   748   926   957  1174
1292  1342  1407     9    32    53   110   124   174   310   331   366
474   499   513   532   537   573   618   671   704   749   756   772
781   794   845   872   903   978   995  1004  1093  1255  1277  1382
1426   182   275   305   479   523   562   623   679   692   734   743
747   827   927  1063  1101  1341  1373  1421   131   371   570   851
42    82   210   355   392   449   624   650   732   945  1007  1058
1075  1161  1207  1314  1335  1340  1492   140   229   333   341   481
531   569   681   712   719   778   804   810   843  1272  1284  1336
1371  1391  1428    47   930  1254  1394  1480  1460    76   109   192
356   427   757  1171  1321  1502   361  1078  1490    73   453   817
1287];[98   106   117   122   141   174   181   259   286   299   307
323   357   394   407   414   429   468   484   521   574   591   623
671   770   783   838   860   864   915  1078     5   208   249   292
297   345   370   445   483   633   683   709   715   743   753   765
772   801   914   993  1100    59   400   420   463   766   817   865
908  1109   126   133   157   182   264   285   333   378   403   423
427   432   459   515   522   543   573   592   626   732   798   940
956   981   988  1019  1088    42   165   251   352   478   563   684
734   751   769   777   834  1107   410   797    72   170   262   301
311   354   555   648   649   721   755   780   856   868   913   928
931   994  1051   137   330   600   652   897  1015   218   287   641
702   763   916   160   175   227   235   271   458   476   696  1035
1069    91   178   461   467   785    94  1077   556   176];[6    44
63    69    97   100   138   204   217   224   240   258   274   295
336   409   448   473   490   492   541   572   598   599   634   640
641   663   664   672   675   794   848   883   902   912   998  1025
1029  1038  1041   207   296   411   468   565   602   775   776   783
790   963   971    60    90   173   187   328   463   535   714   784
810   831   908    61   111   315   330   348   434   449   460   461
484   485   504   524   553   582   617   676   695   696   734   765
800   816   832   899   931   938   961   992  1052     4    59    79
143   179   213   456   601   615   644   702   705   768   814   820
950   952  1023   326   970    42   225   251   307   361   568   584
595   740   756   890   911   924   929   949   978  1014   185   195
319   369   474   578   685   791   843   884   962  1019    71   480
865   793  1008   398   491   708   773   135   453   916   787   982
302   397   310   901   612];};
% SIX=0; requireAll = 1; ADJACENTONLY = 0;
% % allowSubstring = 1; CONTEXT = 2; pat={{'CpaF';}; {'RNA degradosome
polyphosphate kinase';}};
% % allowSubstring = 1; CONTEXT = 1; pat={{'CpaF';};{'AAA family
ATPase';'ATPase';'P-loop NTPase';'hypothetical protein';'septum';};};
% allowSubstring = 0; pat={'CpaF';};

%
% % these are new, more consistent, clear and standard anns in
'/mnt/spin/anns_Bacteria_sa_alt';
% ANNS_SA = '/mnt/spin/anns_Bacteria_sa_alt';
% genus='Bifi_alt';
% SWITCH='for_Bifi';
% SUFFIX = '_alt_Dh';
```

```
% d='';
%
gs={'igrs_Bifi_A11_alt';'igrs_Bifi_adolescentis_alt';'igrs_Bifi_angulatum
_alt';'igrs_Bifi_animalis_alt';'igrs_Bifi_bifidum_alt';
%
      'igrs_Bifi_bombi_alt';'igrs_Bifi_boum_alt';'igrs_Bifi_catenulatum_a
lt';'igrs_Bifi_choerinum_alt';'igrs_Bifi_dentium_alt';
%     'igrs_Bifi_gallicum_alt';
'igrs_Bifi_longum_alt';'igrs_Bifi_magnum_alt';'igrs_Bifi_minimum_alt';
'igrs_Bifi_moukalabense_alt';
%
'igrs_Bifi_pseudocatenulatum_alt';'igrs_Bifi_pseudolongum_alt';'igrs_Bifi
_ruminantium_alt';'igrs_Bifi_subtile_alt';
%
      'igrs_Bifi_thermacidophilum_alt';'igrs_Bifi_thermophilum_alt';'igrs
_Bifi_tsurumiense_alt';};
% SCORES=[1 2 3 4 5 6 7 8 9 10 11 12 13 14];
% ns={[3   18   68   76  120  139  186  192  197  206  212  213  218  237
282  297  306  328  337   14   67   78  119  180  123  172  203  211  357
10   25   29   56  104  124  128  131  141  302  331  354    2    9   44
62   89  209  160  196  249  283  321  151  273  327  345  103  225  246
214  290  332   85];[20  160  173  193  219  221  224  255  256  309  340
345  379    6  176  181  252  274  297   33   44  172   57   65   67  113
204  269  280  301  310  342  144  250  374   32  120  234  247  257  337
362  363  369   12  189  223  246  270   93  231  356   50  119  222  282
286  324  230  249];[19   25   26   46  100  103  126  213  249  271  281
303  325  339  383  389  390  404  412   32   79   81  104  105  106  177
194  326  387   52   89   97   98  145  151  204  238  251  278  284  321
358  370  374  384   62  139  209  210  239  282  309  348  376   80  120
157  227  264  274   61  357  229  237   12   15  258  386  409   67
20];[14   29   33   68   83   84  120  123  200  213  214  217  228  236
278  315  365  366  367    3   13  124  196  316  368   15   30   62   66
97  103  194  218  258  267  269  275  324  340   71  171  292  319   57
58  109  298  238  370  353  369  133  187];[2   41   69   71   91  259
271  297  317  333  348  352  355  418  441  453  468  471  509  514  558
73  191  501  515    8   89   97  139  296  436  458  503   37   68  128
211  235  244  327  435  504  533  539  548   65  104  105  145  187  230
253  300  318  457  519  358   31   90   94  130  153  366  373  433  456
500   42   74  218  274  295  368  391  494  522   82   98  275  390  434
478    9  291  513  113  496  407  428];[28   50  111  112  149  215  248
289  331  352  381  382  426  431  436  440  459  460  494  531  532  548
607  667  680  701  729  227  315  441  505  656   56   82  108  118  403
616  625  702   17   39   59   71  114  181  196  209  351  389  415  452
600  648  664  681  714  732  170  186  260  555  594  599   36   91  230
241  298  353  354  370  500  554  668  684  691  708   35   96  127  128
169  325  507  508  556  676  722  179  195  275  439  493    2   12  420
283  512  584  624  570   90];[67   85   95  113  134  218  241  282  294
380  388  402  404  420  446  534   79  216  222  403  421  431  437  481
485  555   87   96  221  288  302  136  140  239  242  303  317  363  383
410  424  428  438  466  467  514  533  562  618   57   88  135  138  160
231  335  397  451  462  617   40   97  203  214  215  268  301  413  492
519  557  580  586  592  594   24   64  177  189  210  390  435  601  611
613  173  213  394  422  546  569   80  126  262  318  347  355  450   73
499  559   29];[16   87  122  136  141  148  154  194  195  199  200  203
278  329   13  115  196  231  248   14  283  357   67   91  121  173  204
250  256  257  274  347  359  380  404  126  255  276  338  386   85   97
130  189  211  290  390   72   95  197  349  356  395   30  328  292   33
82  281];[5   18   40   54  107  138  153  182  193  197  221  228  239
```

243 270 284 299 311 318 378 384 395 462 486 498 507 79 254
297 309 333 417 418 446 499 4 29 145 307 437 162 207 226
279 382 415 422 489 494 16 100 126 129 130 178 185 216 261
380 43 224 230 280 305 319 340 429 468 485 63 69 86 259
369 516 102 97 426 168 191 342];[104 148 178 205 215 218
256 283 293 301 309 310 322 323 365 399 429 437 441 297 304
298 327 329 36 137 145 155 175 238 273 358 389 394 407 423
447 23 70 76 97 103 135 164 226 369 290 330 77 91 181
284 333 337 416 12 98 146 210 212 240 357 111 228 355 263
321 46 152 187 461];[56 73 89 92 100 129 150 174 258 317
333 337 340 341 342 419 421 443 454 496 497 500 511 512 514
521 535 608 619 662 22 334 409 410 453 501 586 186 203 265
312 408 502 74 99 101 106 120 164 170 179 238 344 363 364
463 491 503 566 570 632 634 644 675 212 429 498 565 651 670
16 51 52 68 144 173 204 227 266 274 313 368 393 466 488
533 636 41 55 350 482 537 571 652 14 226 254 367 66 445
467 481 596 666 9 339 305 519 513 522 425];[27 104 126 185
215 217 239 299 312 336 361 371 386 389 390 47 58 253 28
29 125 302 385 3 66 100 112 118 119 231 324 326 359 380
54 93 157 310 333 59 114 131 163 164 169 328 159 232 311
325 16 60 194 287 216 259 308 90 254 158 255];[21 23 74
119 138 140 200 255 265 267 293 307 5 97 114 316 112 192
257 322 335 8 98 110 173 183 208 209 132 50 116 142 187
266 294 319 38 80 88 229 240 246 321 24 174 233 249 315
6 89 79];[16 53 101 102 114 124 126 137 164 166 210 240
249 321 328 331 362 37 48 129 160 212 218 285 294 355 385
128 33 52 66 78 125 192 203 279 327 346 383 384 390 401
403 414 91 250 306 332 376 381 21 142 157 213 217 291 322
335 400 402 405 94 130 187 266 276 303 345 93 186 280 183
409];[9 22 38 48 53 59 97 108 111 115 136 163 176 179
186 193 245 246 346 405 409 15 34 74 106 343 12 81 82
83 98 122 142 178 204 237 272 275 313 380 415 416 99 203
238 273 20 114 116 242 399 50 102 180 207 330 374 391 413
2 214 383 414 406 84 94 243 342 167];[1 3 55 62 75
77 89 119 157 174 197 227 266 299 301 177 187 300 357 117
231 255 307 33 71 76 78 164 268 291 343 359 371 394 397
405 449 52 95 131 245 292 43 94 97 123 208 214 347 395
444 18 61 90 175 201 308 311 358 382 450 17 282 455 125
22 445 461 356];[8 48 60 78 124 168 169 223 248 258 306
326 328 101 206 287 151 68 123 185 222 235 250 272 273 172
216 245 294 160 298 143 193 196 43 137 189 183];[20 57 58
63 106 206 263 295 297 317 326 345 346 381 382 415 433 298
336 398 429 434 55 122 5 154 189 236 306 351 367 374 402
416 427 148 190 216 150 165 174 192 226 302 412 11 18 80
156 322 347 352 358 394 422 423 391 121 271 321 395 397 12
2 324 343 379 425 272 305];[57 72 175 239 253 262 269
270 308 318 344 358 367 388 405 408 413 471 475
478 523 554 562 598 656 713 727 737 739 794 801
811 827 847 896 912 918 932 971 24 214 240 254
357 396 404 489 552 582 685 812 831 924 927 954
962 968 439 500 502 643 1009 7 23 34 75 81
124 132 208 213 221 293 353 419 421 526 531 532
583 646 797 843 886 1002 59 109 116 251 263 305
382 533 650 816 988 174 220 231 245 279 350 509
541 596 611 644 756 790 36 88 153 200 380 728
729 758 870 895 936 945 35 154 499 757 10 30
207 219 513 530 621 626 837 842 1007 82 252 92
227 126 476 866 216 597];[43 58 61 62 107 170 203 217

```
224  229  241  305  307  401  414  452  549  567  585  599  629  639  667
677  209  213  278  437  522  538  168  239  287  392  744  751   12   27
54   72   76  114  161  175  183  189  236  257  284  294  355  375  383
469  579  623  627  676   25  160  174  234  264  378  480  547  590  741
20   41  156  190  218  262  323  406  425  523  524  592  609  657   56
112  186  289  339  390  429  601  703  750  648   60  171  320  426  501
502  758  142  718   92];[25   49   58   59   99  262  278  319  320  434
487  523  571  651  714  755  771   72  161  213  277  289  390  455  461
586  684  709  789  118  205  369  413  425  474  495  517  615  701   43
74  111  117  159  199  232  260  263  283  337  366  377  380  385  389
414  426  521  552  597  609  670  679  681  705  723  726  734   24   52
208  223  361  438  444  566  588  699  811  673    2   15  372  511  533
546  565  635  696  717  831  100  124  402  469  613  747  776  780  787
40  316  411  504  637  360  580  290  498  713  440   73  123  687  388
545];[17   28   72  105  131  142  193  200  201  202  207  233  254  267
338  355  384  417  483  502  528  538  540  574  604  605  627   49  112
119  126  214  228  237  245  278  361  469  500  527  594   30   92  394
18  165  208  248  265  307  348  377  390  413  505  511  539  551  111
362  382  543  567  262   71  166  232  302  335  519  577  593    4   25
135  194  206  268  426  432  468  474  481  516  517  124  161  252  301
127  358  430];};
% SIX=0; requireAll = 1; ADJACENTONLY = 0;
% % allowSubstring = 1; CONTEXT = 2; pat={{'CpaE';'TadA';'CpaF';'P-loop
NTPase';'AAA family ATPase';}; {'RNA degradosome polyphosphate
kinase';}};
% allowSubstring = 1; CONTEXT = 1; pat={{'CpaE';'P-loop NTPase';'AAA
family ATPase';};{'CpaF';'TadA';};};
% % allowSubstring = 1; CONTEXT = 1; pat={'RNA degradosome';};

genus='bifi_alt';
ANNS_SA = '/mnt/spin/anns_Bacteria_sa_alt';
SWITCH='for_Bifi';
SUFFIX = '_alt_230_Dh';
d='';
gs={'igrs_Bifi_A11_alt_230';'igrs_Bifi_adolescentis_alt_230';'igrs_Bifi_a
ngulatum_alt_230';'igrs_Bifi_animalis_alt_230';
'igrs_Bifi_bifidum_alt_230';
      'igrs_Bifi_bombi_alt_230';'igrs_Bifi_boum_alt_230';'igrs_Bifi_choer
inum_alt_230';'igrs_Bifi_catenulatum_alt_230';'igrs_Bifi_dentium_alt_230'
;
      'igrs_Bifi_gallicum_alt_230';'igrs_Bifi_longum_alt_230';'igrs_Bifi_
magnum_alt_230';'igrs_Bifi_minimum_alt_230';'igrs_Bifi_moukalabense_alt_2
30';

'igrs_Bifi_pseudocatenulatum_alt_230';'igrs_Bifi_pseudolongum_alt_230';'i
grs_Bifi_ruminantium_alt_230';'igrs_Bifi_subtile_alt_230';'igrs_Bifi_ther
macidophilum_alt_230';
      'igrs_Bifi_thermophilum_alt_230';'igrs_Bifi_tsurumiense_alt_230';};
% SCORES=[1 2 3 4 5 6 7 8 9 10 11 12 13 14];

% SCORES=[1 2 3 4];

% % SCORES=[5 6 10];

% SCORES=[8 9 12 13 14];

% SCORES=[8 9];
```

```
% SCORES=[12 13 14];

% SCORES=[12 ];

% SCORES=[13];

% SCORES=[14];

SCORES=[12 13];
ns={[260   59  276    4   13  108  274  275  414  478  217  242  283  344
374  538  161  168  536  515  314  590  212  472    3  355   48  106  354
471  315  352  541  583  413  569  231  144  341   23   18];[62   71   82
305  502  288  637  345  175  306  346  412  420  499  583    3  210  307
362  364  436  537  248  465  417  111   17  473   72   14  467  653  228
540  389  486  363  168  253];[38  408  638  270  547  398  527   98  338
506  556  704   22  127  348  573    7  155  166  260  309  525  591  510
112  552  495  111  438  217  494  592  459  655  503  274  448  703
561];[89   24  126  569  112  205  340  497  499  673   11   18  111  240
265  355  609  674  104  137  155  476  670  204  500  440  482  339  309
64  398  292  128   74  595  168   73  127  278  532  534  310   72
79];[122  121  281  344  829  250  216  355  569  321   57  230  282  497
509  637  102  206  649  662  693  729  110  286  368  505  587  747  142
552  743  615  864   16  248  101  582  444  401   10  459   64   88  821
772  686  297  394  456  187];[1076   152   465    77   787   234   965
102   284   812    66   122   132   369   567   653   654   814   848
884   902   998   146   351   429   446   773   864  1038  1068   724
70   100    80   912   874  1036     4   876   145   694   908   890
794   743   663   868  1022   845   877   557   377   106   822   440
559    12   133   605   317   697   840   978   956   188   266
538];[627  160  898  736  332  638  203  399  449  527  670  311  566
625  635  662   63  218  338  412  424  815  917  935  505  336  281  214
676  922   88  645  541  836  626  604  327  407  243  480  594];[347
511  556  408  523  240  541  751  801   33  249  409  679  419  516  678
497   22   86  333  780  160  187   88  224  724  122  758  634  337
281];[347   11  389  443   53  145   23  356  472  566  595  625  122
201  471  485  519  650   47   73  142  303  180   67   68  442  151  269
652  228  434   46  331  355  202   69  124  465  416  322  560  243  571
206  230  332];[577  126  473  239  427  449  344  371  116  243  406
609  642   24   61  232  360  455  515  702  345  636  250  325  692  348
224  514  569  359  624  583  695  413  183  274  742  219  614  715  534
446  261];[133  689  709  160  532  703  896  933  472  583  647  249
260  570  765  327  420  468  470  562  707  926  929  178  202  258  349
512  575  743  749  847  523  700   61  222  473  670  390  694  492  380
602  152  546  666  554  758  712  593  856  214  664  464  206  118
64];[640  424  767  191  360  407   93  192  269  303  366  390   66  147
347  568  740   42   96  106  483  485  646  727   43   61  124  154  270
520  541  755   48   95  153  249  391  449  521  699  712  508  412   46
733  589   39  104  127  196  665  522  120  349  613  667  100  486
52];[412  194   47    6   80  149  347  442  482  237  312  315  392   50
300  321   82  165  429   87   75  299  391   17  331  480  159  471
474];[226  337  389  159  323   27   42  115  161  186  375  666  294
407  418  555  561  670  173  178  180  182  231  347  390  422  634  430
603  189    6  434  512  567  486  300  500  177  232  563];[518  219
686  384  211  263  697   12  184  660   93  388   76  380  452  466  666
21  130  232  342  369  423  162  661  328   59  624  482  323  141  387
243  117  187  222  290  691];[257  206  310   53   73  226  330  337
342  587   16   41  169  322   96  225  272  746  805   15  166  211  219
555  694  797  435  123  529  353  180   66   50   76   83  158  535  126
```

```
543   56  509  627  472  449  610  780   85   25  179  204];[84  331  256
206  519  349  468  508  518   59  346  444  494    5   17  230  523  290
467  197  261  204  443  265  534  463  526  202  288  276  441  510  320
408];[549  610   19  366  490  567   20  383  420  501   96  181  192
231  426  444   13  111  309  390  626  687  482  583  358  642   97  129
570  229  683  691  230  294  262  199  618  204  107  110];[1376  1381
361   491   118   375   767   195   698    86   349   473   834   843
1086  1513   726   810  1174  1187  1342   269   486   509  1078  1219
1490     9    73   105   207   453   510   524   618   817   957  1287
1503  1255   562   978   827  1238  1207  1139   704   449   531   756
845   498  1421  1480   110   305   681   743  1063   394   623   747
1277   903   851  1004   719    32   214  1391   926];[864   685  1082
122   208   484   868     5   106   463   574   842  1113   174    91
286   292   299   400   483   673   755   785   178   307   357   461
467   787   799   772   838   800   182   141  1023   522   218   249
323   175   378   429   771   736   698  1104   227   654   594   445
918   765   165   821   573   779   686   287];[567    58   264   838
119   271   351   311  1133   286   399   510   755    30   135   239
693   765   877   961  1010  1029  1057    57   173   414   503   568
668  1019  1066   115   174   337   359   411   498   522   626   671
985  1192  1193  1137   180   349   880   533  1135    33   663   384
51    38   821   604  1107   817   403    92    28   383   587   602
140  1035  1160   638];[794  1025   258   204   224   773   912   187
473   490   714   883    97   135   138   453   468   641   783   908
911   916   173   448   572   775   787   884   902   982    90   302
397   409   535   601   640   672   685   695   240   992   207   348
524   474   411   434   950   800   634   617   484   696   843    44
504    59   449   676   179   463   568  1023   310   644   708   602
820   740];};

SIX=0; requireAll = 1; ADJACENTONLY = 0;
% allowSubstring = 1; CONTEXT = 2; pat={{'CpaE';'TadA';'CpaF';'P-loop
NTPase';'AAA family ATPase';}; {'RNA degradosome polyphosphate
kinase';}};
allowSubstring = 1; CONTEXT = 1; pat={{'CpaE';'P-loop NTPase';'AAA family
ATPase';};{'CpaF';'TadA';};};
% allowSubstring = 1; CONTEXT = 1; pat={'RNA degradosome';};
% ADJACENTONLY = 1; allowSubstring = 1; CONTEXT = 1; pat={'HAD
hydrolase';};

genus='bifi_alt_Df';
ANNS_SA = '/mnt/spin/anns_Bacteria_sa_alt';
SWITCH='for_Bifi';
SUFFIX = '_alt_230_Df';
d='';
gs={'igrs_Bifi_angulatum_alt_230';
      'igrs_Bifi_bombi_alt_230';'igrs_Bifi_boum_alt_230';'igrs_Bifi_choer
inum_alt_230';'igrs_Bifi_catenulatum_alt_230';
      'igrs_Bifi_minimum_alt_230';'igrs_Bifi_moukalabense_alt_230';
    'igrs_Bifi_thermacidophilum_alt_230';
      'igrs_Bifi_thermophilum_alt_230';'igrs_Bifi_tsurumiense_alt_230';};
SCORES=[1 2 3 4 5 6 7 8 9 10 11 12 13 14];
ns={[32   37  111  160  190  212  236  259  262  308  330  334  357  407
408  488  547  556  565  577  623  635  637  638  703    5  110  127  158
217  321  369  529  574  624  636  658   85  221  235  495  525  618   87
128  189  230  256  320  344  433  473  506  510  513  589  634  700   83
163  169  183  201  222  413  549  552  566  122   22   36   65   73  112
150  164  210  226  283  295  337  402  422  435  584  697  155  229  260
```

377 561 562 564 26 97 170 348 86 322 441 496 38 159 555
239 98 215];[13 24 119 146 152 211 243 298 305 322
429 440 446 478 561 583 613 617 627 673 693 694
774 842 845 848 868 883 884 896 1026 1036 1076 7
22 58 77 108 140 220 244 255 424 453 585 598
640 729 787 794 814 881 902 4 284 315 317 451
609 811 901 994 84 85 98 117 163 188 213 269
292 302 321 383 398 401 415 423 444 445 552 563
565 569 616 677 704 719 720 724 747 822 833 866
874 912 917 945 1047 19 66 70 83 100 306 329
394 538 653 660 682 877 885 947 241 8 10 42
75 126 247 314 325 418 434 469 520 712 802 821
954 989 45 59 76 93 175 369 524 550 557 559
792 815 864 890 959 1041 118 479 610 630 1068 145
12 133 283 586 695 801 830 914 1021 1057 965 38
132 351 377 399 465 773 880 956 106 499 1 823
998 94];[12 78 114 131 160 203 246 279 328 358 399 400
401 410 439 449 458 481 499 542 545 570 572 574 590 607 631
636 637 638 647 777 898 920 926 20 47 55 255 310 332 577
814 871 57 58 229 339 340 736 752 76 77 81 99 121 166
185 243 287 291 342 352 362 448 546 579 608 627 642 646 655
666 675 678 757 786 797 835 836 862 872 887 903 907 915 101
165 196 269 348 390 426 428 465 541 589 604 626 668 673 718
773 900 409 8 68 135 270 311 447 496 512 609 667 670 735
874 927 31 91 148 218 327 393 500 505 521 573 594 625 676
693 812 842 860 868 905 115 220 331 223 301 407 648 759 781
550 730 187 412];[15 37 38 84 89 106 114 120 198 211 215
221 232 249 291 347 358 399 409 410 500 516 561 595 608 633
666 678 680 724 726 751 792 21 31 86 204 209 238 345 370
523 553 560 667 799 192 288 290 499 22 45 68 96 162 170
171 226 344 363 392 393 439 476 545 567 606 631 672 713 111
187 224 408 508 533 556 616 684 699 718 88 276 722 7 14
35 71 95 112 122 166 203 335 405 498 557 564 594 669 747
756 772 39 159 213 234 272 312 337 377 378 406 497 565 679
504 175 282 321 334 464 531 541 566 676 807 411 663 168 491
333 295 803 710];[11 47 70 73 122 170 172 201 225 278 347
355 364 386 399 402 411 416 437 439 454 532 565 572 577 578
582 603 643 650 665 683 12 45 108 164 265 435 465 25 40
145 171 215 241 288 361 434 531 566 8 86 120 124 126 144
155 202 209 247 277 281 290 294 312 321 336 401 424 447 496
504 575 602 616 642 680 32 62 102 115 151 206 228 230 269
491 524 571 641 652 521 553 65 68 103 335 449 518 533 570
33 46 104 180 234 244 303 331 519 522 564 620 685 55 450
666 48 53 147 276 457 573 610 644 17 142 368 476 189 69
471];[30 41 42 117 163 166 168 180 181 186 187 204 216 226
231 238 241 256 294 338 375 389 396 413 416 417 540 555 561
602 605 634 666 58 116 159 228 337 347 412 421 422 546 576
668 675 53 298 415 496 511 4 72 179 232 259 430 499 512
522 539 563 564 586 610 633 635 662 676 227 362 565 567 601
606 646 27 45 52 67 113 182 190 203 208 305 335 390 418
471 485 526 629 659 46 140 155 189 219 282 319 451 554 603
626 57 177 604 139 145 215 374 648 472 105 464 545 641 523
656 611 348];[9 55 57 130 141 145 146 165 166 183 203 211
222 231 248 263 268 282 290 328 341 343 348 359 369 386 441
445 492 495 512 514 579 603 616 698 133 233 626 686 289 504
631 697 26 39 43 48 50 54 112 117 134 168 176 184 187
196 219 229 236 281 367 387 429 431 517 518 581 20 64 306
344 403 508 15 409 36 90 103 120 272 323 332 423 540 600

```
17   30  217  232  321  395  538  699  243  298  452  576  624   14   51
93  410  513  541  558  587  654  665  691   76  475  470  547  330  170
98];[5    24    34    59    84    92   107   116   122   132   141   171
173   176   180   218   259   296   356   400   407   413   494   509
521   552   555   579   593   627   628   673   683   685   734   755
757   786   794   816   838   842   864   870   897   924   992  1007
1074  1101   249   430   432   515   677   711   967  1113   378   487
704   799  1082  1114   110   126   133   175   262   288   292   329
345   403   429   492   522   561   573   574   580   623   641   643
668   726   729   753   766   779   785   800   880   919   985   986
1023  1094   105   106   182   193   217   236   375   389   445   484
602   865   976   989  1104  1112   654  1020     6    20    91   142
223   252   269   286   299   324   334   351   489   549   559   625
636   686   696   836   849   918   965  1004  1031  1043    73   160
165   264   352   387   410   474   497   736   744   768   784   859
944   959   960  1022  1086   323   698   772   621   272   478   543
601   724   735   998   357  1038   723   912   920   330   390   140
483];[7    30    33    58    88   108   120   161   162   193   227   326
341   349   359   370   384   394   412   434   439   470   496   499
503   602   637   644   655   711   765   781   870   884   906   908
1078  1119  1121  1166    19   116   145   179   215   314   500   563
599   667  1066  1107  1185  1193    28   119   135   147   218   316
567   680   755   795  1026  1028  1170    74   159   173   184   208
240   272   321   323   325   334   383   389   399   460   482   490
498   509   517   522   569   587   604   630   633   666   693   806
846   937   945   963   988  1017  1019  1034  1035  1070  1076  1102
1167  1169  1178    51   107   311   403   454   508   533   654   838
1050  1160  1183   469  1058    34    43   143   548   586   591   607
770   947   983  1015  1020  1041  1043    14    29   100   126   137
160   174   180   271   414   497   527   547   600   730   880   907
1122  1192    52   668   879   818   244   372   981  1184   411   554
1029   206   351  1021   782   229  1051   656   805   985   701     4
917   986];[14    26    64    89    97   127   128   171   194   204
217   295   302   318   353   404   442   463   490   504   518   534
535   553   578   632   639   640   675   689   714   723   775   779
799   836   883   912   957   969   971   980   989  1037    45    58
187   224   258   290   460   489   598   665   685   750   765   794
924    71   421   480   540   631   890    49    61    84   104   111
166   179   188   212   240   276   294   409   431   459   473   474
478   524   593   602   617   721   767   776   790   884   899   938
992    79    93   273   274   434   456   594   708   928   988  1002
1008   321   595     7   422   454   514   563   679   691   695   696
734   840   894   939   955   973  1014  1024  1048  1050     8    75
213   348   453   464   528   601   608   716   754   774   861  1023
207   753   947   972    66   259   310   484   713   800   898   911
952  1034   109   902   936   692   634   676   491   740];};
% SCORES=[1 2 3 4];

% % SCORES=[5 6 10];

% SCORES=[8 9 12 13 14];

% SCORES=[8 9];

% SCORES=[12 13 14];

% SCORES=[12 ];
```

```
% SCORES=[13];

% SCORES=[14];

% SCORES=[12 13];

SIX=0; requireAll = 1; ADJACENTONLY = 0;
% allowSubstring = 1; CONTEXT = 2; pat={{'CpaE';'TadA';'CpaF';'P-loop
NTPase';'AAA family ATPase';}; {'RNA degradosome polyphosphate
kinase';}};
allowSubstring = 1; CONTEXT = 1; pat={{'CpaE';'P-loop NTPase';'AAA family
ATPase';};{'CpaF';'TadA';};};
% allowSubstring = 1; CONTEXT = 1; pat={'RNA degradosome';};
% ADJACENTONLY = 1; allowSubstring = 1; CONTEXT = 1; pat={'HAD
hydrolase';};


% genus='glyco';
% d = '';
% SCORES=13;
% % % CUSTOM_SCANNING_LENGTH=230, /mnt/new/pepik/pactool/Msmest_n_7.br
%
gs={'igrs_Glyco_arizonensis230';'igrs_Glyco_dulcitolivorans230';'igrs_Gly
co_artemisiae230';'igrs_Glyco_terrestris230';'igrs_Glyco_buryatensis230';
'igrs_Glyco_paridis230';'igrs_Glyco_tenuis230';'igrs_Glyco_xiaoerkulensis
230';};
% ns={[1166  1171  1296  1066   461   709   489   194   258   239  1147
167  1249   284  1172   131   671   630    67   393  1094   409   376
1377   555   967  1140   836   639   888   980  1292   456   757   831
93   804   304   230   879   982  1021  1357   315   418   704  1482
692  1525  1526   568  1330  1070   960  1476  1069  1436   710  1329
231   108  1260   685   225   721   520];
%     [1252   264   868  1087   706  1059   978   969   371  1225    75
625   573   908   458  1229  1140  1254   566   578   760   609   995
180   121   102   591   515   503   317   933   599   439   500   577
1021   979   365   626   380   355   867   934   213   263   409   652
1208   902   785    63   149   443  1164   812   374   834   873    64];
%     [353    46  1170   691   386   585   102   856  1179   524   428
300   124   113   417   692   590   460   753  1217   213   547   137
354   849   318   459   909  1096   128   363   843   541   271   916
697   297   705   587   241   626  1116  1123   294   544  1051   293
34  1147  1020   421   783];
%     [68   412  1353    38  1023  1116  1013  1501   777  1111   904
1248   467   260   611  1004   825  1042   764  1110  1040    72   680
469   665  1150   838    20  1126   402    52   476   149    78    19
1105   724  1230  1143   387   148   245   174  1049   840  1322  1294
758   173  1415   320  1058  1321  1234   699    58   881   452   274
894   151   275   348   952  1237  1218  1465];[1640   593  1327   361
1356   625  1380   207  1674  1093  1187  1820   346  1243  1153   305
1441   190   164   618   265    62   432   960  1607  1797   554   929
798   961  1789  1040   241  1068    33   429  1735   316  1218  1746
433  1299   626  1548   479  1850   243   385   353  1406  1196   245
1411  1442  1057  1412   812   175   642  1545   345  1510   773  1273
902   926   395  1791];[1585   839  1536   425   474   482   305   965
203  1250   473  1478  1335   335  1197  1436   665   184   187   501
1009  1307    74  1065   402   642  1477  1293  1474  1562   734   106
1141   629  1410   254    36  1392   352   986   218   521   366   519
601   599   891  1308   417    69   730   952   133   162  1378  1193
```

```
1060   840   434  1029  1534   165  1446   709   363   959  1116   505
1080   204   835   157  1245   748   400  1112   645];[1273  1505   480
1656    91  1217  1023  1113  1175  1366   247   533   117   812   922
1164   996   734   123    93  1232   934  1623   171   884   200  1441
438   463  1670   202  1654     7   111   168   188  1236  1252  1218
135   319   352  1005  1394  1178   248   347   913   191   452   684
1037   630   820  1127  1244   796   387   272   586  1234   847   887
1389  1202   581   622];[820   231   980   757    32   489   617   406
721  1104   550   407   948  1075   659    59   412   816   283   693
1160   273   989   878    77   809   360   390    16  1167   404   381
302   298   974   815   141   301   551   399   139    76    24   593
366];};
% % % /home/pepik/hs/Ms1_Sco_published.fa.br (227)
% ns={[1023  1021  1296  1291   836  1166   247   738  1505   868   758
1405  1069  1093   461  1370   518   245   399  1537  1475  1140   226
1332  1133   506   686   667   814  1316   710   229  1470   102   696
235  1170    49   813   805   108  1137  1471  1526   304    27   196
142   112   748  1220   670   173   131   980     6   265  1493  1371
888   133   679   171  1054   771    17  1519   968  1398   159  1531
580  1349];[726   911   482   859  1113   706   684   669   308  1012
1047   785    17   610  1142   881    97  1266   722    64   371   149
763   380   126   812   368   500   846    72   800   657   933    91
344  1251   815  1031   517  1208   444   735   591   910  1091   594
484   599   719    21   503   977   364   158   917   174   575   833
973];[859   399   314   903   916   839   626   857    58  1015  1123
483  1147   585   692   361  1187   397   358   822   353  1165   685
293   137   301   970   473   300   318   147  1196   299   495   796
909   232   659    54   534   896  1158  1020   519  1067   871   542
520  1170];[1256  1370  1446   947    13    29   720    58   356   690
1492  1127   668   342  1404   536  1336  1301  1013   949    26   623
667  1144   952   644   443   825  1454  1248   865  1258   405  1218
1040  1111    83   772  1150  1468  1423  1116   482   805   764  1289
374   149   442   420   719  1143  1052   894   148   268   145  1051
1089   259  1545   481   469];[1267   627  1335   345   828   361   862
401  1290   364   363  1632   610  1441  1537   709   934   362  1114
744  1604   947   385  1793   257    89   589   294  1356  1541  1273
851   640  1304   733  1486  1237   683  1548    62  1063   641  1213
1534   524  1603   413   889  1850   859  1722   593  1442    44  1773
1291  1844  1819   994  1805   692  1564  1783   631  1573  1797  1078
265  1735   852  1132  1131];[894   473  1166  1308   124   548   458
836   417   254   781  1336  1107   799  1509   991   645   948  1333
1503  1385  1193  1070   239  1534  1369   985   162  1456   183     8
1317  1562   347   585  1159  1085   192   474   768  1076  1180  1478
1149  1112  1270   917   972   840  1335   115   901   123   810   204
22   111   946   839   534   586  1245   777  1425   453   571  1378
550   955  1290  1420   915  1356   920  1197];[906   815  1217  1654
1558  1555  1505   147   798   282   141  1293  1128  1448   374    11
441   198  1044   830   733  1656   741   387   191   986   272  1576
151   271   144  1382   222   118   698  1670   609   142   782    10
1176   197   657  1438   475  1168   476  1239  1621   359  1079   989
388   549  1461   219   511   796   232  1077  1191    28   139   653
1167   630  1415   582   143   706   281  1423];[589  1030   146   948
1009   408   877   450   474   273  1029   684    59   507   897   301
16   366   989   285   861   960   382    78   942   172   223  1171
238  1160   758   994   272    10   721  1085   977  1173   658  1159];};
% % % /mnt/new/pepik/pactool/Msmest_n_7.br (304)
% %
gs={'igrs_Glyco_arizonensis';'igrs_Glyco_dulcitolivorans';'igrs_Glyco_art
```

```
emisiae';'igrs_Glyco_terrestris';'igrs_Glyco_buryatensis';'igrs_Glyco_par
idis';'igrs_Glyco_tenuis';'igrs_Glyco_xiaoerkulensis';};
% % ns={[972   807   368   746   892   113   466   536   875  1042   916
874   906   171   472   710   894   263   641   371   311   722   886
279   925  1044   284   506   650     3   932   528   900   519   389
308   721   285   824   286  1030   485   517   778   726   727   211
340    74];[674  463  207  659   33  191  163  267  348  231  530  321
482  368  464  529  317  143  491  111  237  116  268  451  347  478  442
456  677  671  648   11  168  636  514];[208  305  342  206  327  304
401  485  397  415  615  332  223  376  521  290  152  500  163  330   60
400  487  650  649  197   93  221  365  642  406   19  220  180  585  627
654  546];[477  288  839  437  641  181  212  537  854  376  116   10
758  358  830   75  703  206  397  262  805  264  284  276  327  534   81
547   22  123  150  844  161  384  329  538  833  300  778  753  634  837
681  411  338  115  365  440  810  523  721  640  370  200  170];[674
898   656   551   972   245   970   436   152   314   690   770   196
249   122   120   328    91   774   180   227   859   889   322  1054
61   552   831   200   273   671   597   938   689   560   900   823
563   522   848   353   792   544   292   735   325];[393  521  438   32
487  696  834  631  719  947  286  153  627   12  968  586  410  323  344
502   94  900  832  519  628  569  328   67  575   33  566  244  855   73
909  140  939   40  270  675  578  145];[276   227   841   780   334
800   566   685    71   244   391   122  1008   823   452   243   712
793   449   692   333   745   658   631    42  1061   580   524   479
113  1023   166   407   448   341   169   102   527   548   572   888
376   850   721   711   339   923   784   710   318   425];[555   11  405
180  264  438  207  411  615  659  399   49   25  382  250  223  190  602
610  153  613  462  544  400    3  376  557  605  641  682  393  151  632
44];};

% genus='sali';
% d = '';
% SCORES=13;
% % CUSTOM_SCANNING_LENGTH=230, /mnt/new/pepik/pactool/Msmest_n_7.br
%
gs={'igrs_Sali_arenicola230';'igrs_Sali_vitiensis230';'igrs_Sali_mooreana
230';'igrs_Sali_cortesiana230';'igrs_Sali_oceanensis230';'igrs_Sali_fenic
alii230';'igrs_Sali_tropica230';'igrs_Sali_pacifica230';};
% ns={[1683   400    99  1897  1223   197  1293  1647   482  1740   331
2103  1412   715   203  1135  1320   343    72   319    55   948  1529
660  1333   545  1566   896  2039  1631   783  1593   898  1346   929
1224  1263  1550  1961  1565  2055   640  1197  1063   692  1668     2
1442   644  1828  1413   579   639   332  1301  1154   399  1726   713
1030   630   258  1045  1270  1935   823   405   963  1530  2036   221
410   648   556  1864  1684  1995  1389  1616  2053  1013  1425
1337];[870  1432   771   730  1191   444  1026   435   827   569  1016
467  1480  1342   508   350   896  1295   589  1491   777  1289   757
865   609   365  1101   463  1075  1299    78    55  1399   916  1300
1350    92   169  1320  1261   174   685    89   890    30   498  1478
1506   810    60  1306   160  1248  1112   783    62   190   785   445
846   560    61   791  1242   327   825   780   214   834  1368];[722
1402  1410   563  1332  1607   221  1266  1045  1072    31  1362   720
323   440   115   439   947   344  1414  1115  1046   594   941   413
889   671   810  1440  1544   713   945   856  1100   719  1344  1158
777   649  1645   553   554   567   759   886  1475    56  1465   979
17   576  1439  1076  1077  1391  1526   385   569  1466   644  1436
386    36   542  1022   427  1269  1136   857   311];[670  1409  1108
680   312  1490  1280  1302   304  1170   462   222   618   765   848
```

```
120  1295  1623   468   449   406   827   192   723   666  1086   360
4  1257  1617  1169  1273  1543  1352   547   486  1136   156  1630  1635
1626   551   984   140   586   663  1304   821   909   863   159  1460
282  1058   436   590  1459   543   754    67   362  1515   585
302];[1148  1056   141  1156   916  1055  1280   841   393   339    96
787   607   782  1665   420   910   477   679     8   920   192   852
1235   983   864   284    70   171  1462   439   818  1537  1287   944
1021  1267   778  1404  1392   443  1344  1332  1169  1288    43  1144
45     7   399   951  1255  1126   232  1099   830   710  1268];[1151
912   546   863   325   218   290   770  1355  1578   547   724  1048
1039   933   554   503   355   801   579  1518  1196   106  1182  1378
540     7   472   166  1381  1210   309   814   219  1521  1303   806
1476  1100  1588   326   323  1412  1669   697    20  1413   564   998
366   822   504   563  1249   528   625  1336  1446  1234   459   184
890  1315  1483  1611  1051   163  1254   452   740   805  1095   413
365  1047   808   530  1614];[157  1457   799   230   576    50   932
697  1061   662   616   229   706   808   260   392   536  1650   607
915   149  1512   916  1225  1106   689   930   239   573  1555  1265
1080   803  1085   790  1416  1039   538  1105   358  1223   272  1043
650   196   316  1116   693   346   102   896   144   940   477   314
340   140  1634   391  1129  1153    33   416   266  1649   913  1079
918];[515   555  1402     3  1563   828  1106   707  1523   523  1416
150   202  1146   644  1278   277   305   499  1100   259   909  1184
512   299  1642   160   436   322   344  1543  1241  1608  1643  1122
45  1410   247  1457   749   248  1644   537  1674  1597  1138   133
506  1451   231   142  1615  1406  1640  1075   122  1024  1209   340
35  1536  1555   710   421];};
% % /mnt/new/pepik/pactool/Msmest_n_7.br (304)
%
gs={'igrs_Sali_arenicola';'igrs_Sali_vitiensis';'igrs_Sali_mooreana';'igr
s_Sali_cortesiana';'igrs_Sali_oceanensis';'igrs_Sali_fenicalii';'igrs_Sal
i_tropica';'igrs_Sali_pacifica';};
% ns={[1152  1256   788   197   793  1133   666  1352  1313  1254   633
831   925  1114   380   411  1042   280  1316  1261  1328   845   807
334  1132   210  1296  1031   996   913  1182   915   707   673   318
838   146   819  1118  1126   906  1197   620  1017   692   236   737
239    56  1201   450   631  1157    91  1372  1160   209    78  1315
1409  1169  1113   877   700];[866  382  401  440  415  553  806  864
886  165   12  631  352  496    9  220   69  780  753  749  494  522  811
870  827  124  431  406  339   20  654  218  169   80  655  174  254  863
402  841   88  240  521  419  897];[387   355   607   275   928   644
854   444   259    73   158   417   858   276   358   403  1022   233
238   247   885   585   856   313   132   566   376   910    30   344
1035   978  1054   237   540   638   796   759   260   746   590   157
707];[809  1068   224   922   124    96    39   150   727   990   212
921  1073   635   680   136   343   404  1170   352   350   512   483
1183   726  1195   935   271   408  1114  1119  1205   335   226   768
300   464   275   435  1150   135   127  1008   826   168   348
360];[803   926   653   707   728   336   588   600    73   625  1045
139   722   231   554   173   287   810   870   303   724   822   283
671   626   334   873   815   317   883    67   735   909    40   772
191   825    93   440];[185   938   304   445   658   843   661   785
550   649   493   964   183   863   779   341   706   303   566   819
1012   781   866   113   295   667   799   428   306   478   441   963
125   193   237   787   358   174   184   419   224];[659   635   865
403   651   703    42  1035   704  1036   384   272   569   571    83
148   692    84   337   436   145   418   174   781   297  1027     1
130   127   705    14   363   228   362   340   718   438   667   505
```

```
492  1021   147   582];[956   195  1023   204   115   986   521   984
960   640    87   722    88   608     3  1015   157    73   129   380
302   723   378   359   329   459   945   789   494   131  1016   928
584    98    31    97   899   130   175   958   572   505   971  1034
252  1012   385];};

% genus='frank';
% d = '';
% SCORES=13;
% % CUSTOM_SCANNING_LENGTH=230, /mnt/new/pepik/pactool/Msmest_n_7.br
%
gs={'igrs_frank_umida230';'igrs_frank_casuarinae230';'igrs_frank_canadens
is230';'igrs_frank_tisae230';'igrs_frank_gtarii230';'igrs_frank_torreyi23
0';'igrs_frank_QA3230';'igrs_frank_nepalensis230';};
% ns={[756   683  1193  2766  2830   160   705   256   421  1766  1765
2910    21  2705   158  2249  1288  2098  2340  1253    45  1828  2685
259  2430  2209  1734  1310  2569  1192  2233  2527  2874  2510   690
341   368   516  1337  2164  1691  1683   963   170  1961  2795  2871
2779   560  1107  2434   736   123   891    17  2097  2155   396  2668
1089   539  1444   649  1811  1364  2452   105  1055  2583   253  2842
1675   942  2875  2879  1623  2061  2644  1533   606  1733   967   563
2224  2352  1286  1363  2806  2808  2739  1065  2429  2699  2100  1887
1973  1344    12   862  1824  1078   468   975  1157   149   651   809
2791   530  1142  2189];[1021  2110   393  1855  2432  1335  2329  2605
1580    39  1673   643  1603  1181  1632  2594  1081  1200  1261  1359
1104   181  1800   692  1433  1080  1077  1788   915  1222  1007  1067
2271  1027   550   983  2547   239  2530  2413   351  2595   612   850
1038  2497  1215   365    87  1453  1360  2301  1959   847  2014  2527
297  1726  2499  2486  1186  1059    68   510  1334  1748   336  1874
1068   676  2085   840  1809  1463  2374  1599  2469  2006   154  2559
668   476   322   486   332  1747  2358  2169    36  1434   815   288
1425   572  2226  1884];[1707  3154  2340  2149  1408  2416  1726  3055
520   504  2223  2609  1708  2032  2826  1961  1071  3092  3244   546
2741   485  2436   609   715   471  1296  3140   343   573  1639  3204
2829  2570   250    14  2335  1666  2587  1354  2850   525   846  1968
3094   213  2432  2838   442  1855  1989  1885   137  2366  1303  1485
2002  3047  1497   396  2336   726  2334  2067  3048   889   113   242
301  2825  1705   880  2244  2102  1301  1151   678  1498  1859  1407
2441  2747  1101  1789  3242  1530    10   434   924  2963  3169   669
1310  2473   775  1009  1515   886  2245  1606   458  3156   799  1947
2362  2822    80   912  2267    79   577   986  1063   819  1208
2997];[987   914   330  3422  2003  1191   864  2021   749  1491  2837
567  2732  1949  1250   729  1937  1945   104   565  1194  1786  1894
2641     4   687  2739  1503   693   514  2237  2180  3371  2452   871
1988   866  2690    98   840  1085   770    27  1761   522  2386  1439
2872  1785   568  2703  3005  1630   195  1728  2169   730   586  2179
3243   807  2082   849  1911  1048  3382  1152  2575  2875  3030  3359
2470  1331  1684  1093    33  1201   239   119  2311  2666  3404  2254
1917   409  2903     3   345  2104  3402  1381  1447  1916    16  3399
3352   344   832  2039  1365  2734  2896   241  3379  2740  1128  3413
1482  2561  3262  1824  2392  2227  1026  2110   229  2887  3353];[1363
1190  1396  2932   981  3048  1125  2699  3121   936  1191  1884   978
707  3084  2440  3001   500  2987   798  2959  2292   640  3167  2198
1104     6  1432  1696  2583    50  3400   232  2450   231  2639   442
1897   740   308   138   974   215   935  1060  1636   410  2905  1386
1298  3021   560  2655  1205  2072  3299   973  2362  3117  1422   718
366  3395  1034  3362  2524   238  3195  1421   189  1827    86  1172
2691  3375  1558  2571   873  2473  1204  1686  3065  3353  2152  1147
```

```
1761  3414  2689  3212   721  1455  1463   106  3091   154  1540  1206
2270  2470  1369  3057  1883  2386   907  2489   515  1898  2648   437
2534   381  2717  2227  1703  2967  3053  1657  3413  2082   577  2807
2802];[769  3118  1786  3035   779  3041  2302  2796  3310   785    38
1290  2792  3349    37  1164   393  1558  1689  1289  3072  1348   852
3071  1928  1350  3093   193    29  1548  1491  2916  3426   916  1492
2143  1366    66  1509  3425   933  1372   423  2974  1302   648   507
259  1194  1719   187  3324   150  2802  1873  3467   590   955   938
2461  3374   939  1678  2820  2988  3406  1569  2761   536  3565   160
449  3020  1299  2204   497  1020  2436  1629  1460   937   278   711
576  2345   803  1564  1349   851   753  3153  2939  3512  1151  2262
2248  1694  2605  2909  2484  3319   412  2530   676  1127    28   546
2836  2999  2925    40   566   860  3419   280  2808   123  2026  1897
758];...
%     [859  2493  2126   414   267  1183  2252    65  1521     9   934
129  3450  1022   653  2840   110  2333   191   968  2135  2087  1317
1435  2787  2989   252  2770   585  3494  1539  3156   168  2579  3088
2582  3728  1440  1013   413   955  3264  1749  3073  3141  3087  3334
3606   738  1889  1285  3468  2551  1617  3431  3224   499  3022  2799
1865  1181  1953  3068  2737  2387  1093  2285  3286  1361   862  1238
769  1545  1733   286  1117   682  2530  3665  3155  3201   728   756
3711  2736   160  1278  1380   967   650  3033  3113   847  1633   154
1804   918  1943  3632  2036  2904  1025  2284   741  2811  3283  1751
1224  3009   586   329   703  1520  1486  1971  3060  1965   878  1920
761   962  3448  2462];[1726  1620  1453  2735  3336  2379    36  5435
5446  3118  2788  1471  5442  3405  1652  1435  5744  2997  4786    41
760  3721  2284  4690  1621  4919  1509  1185  2806  2678  4537  4691
5112  5677  1064  3276  5599  4875  3844  4747   153  1031  3254  5647
3890  5107   296  1130  3702  1680   793  1129  4650   798  2303  4788
3636   566  4545   771  3143   901  5569  2224  2096    68  2253  4159
2439  4275  5261  2824  5157  2505  1314  3002  1274  2356  1365  4379
4654   992  3670   894  1977  2260   861  2736   918  4151  1486  2349
3549  4866  2535  2546  3204  1789  5789    52  5734  2526   414  1814
466   893  2390  3564  4811   418  2959  4869   337  4760  4061   140
3088  2031  4319  2079   244  2246  4530  1467  4750  2266  5494  4169
763  4014   477  1097  4081   173  4812  4656  5133  4162  4098  2739
4251  2272  5563  5678  4410  4901  3250  1948  5207  1864  5616  3304
2926  3679  1706  2603   270  1991  1508  1052  2474  2112  4631  2003
2945   379  1144  5000  3946  3025  1819   387  2570  5582  2741  1941
1481   137  1037  3476  1915  1600  5498   323  4046  4580  3398  2270
664  4197  1932   347  1059  4217  2599  4246  5192  1705  3977   825
808  3526  2642  4297   833  1656    15  5294   311   207  1065];};

%
% ADJACENTONLY = 1;
% requireAll = 0; allowSubstring = 0; origigrname
% CONTEXT = 2;
% % pat={{'HAD hydrolase';'glycosyl hydrolase'; 'glycoside
hydrolase';'glycosylase';'glycosylhydrolase';'glycosidehydrolase';'glycos
idase';...
% %
'dehalogenase';'galactosidase';'phosphatase';'hexosaminidase';};
{'transporter';'transcriptional';'regulator';'oxidoreductase';'Fic';}};
%
pat={{'hydrolase';'dehalogenase';'galactosidase';'phosphatase';'hexosamin
idase';'glycosylase';'glycosylhydrolase';
```

```
%     'glycosidase'; 'glycosidehydrolase';
'glucosylase';'glucosylhydrolase';};
{'oxidase';'reductase';'oxidoreductase';'Fic';}};
% % pat={{'serine hydrolase'; 'HAD
hydrolase';'dehalogenase';'galactosidase';'phosphatase';'hexosaminidase';
% %   'glycosyl hydrolase';'glycosylase';'glycosylhydrolase';
% %   'glycoside hydrolase';'glycosidase'; 'glycosidehydrolase';
% %   'glucosyl hydrolase';'glucosylase';'glucosylhydrolase';};
{'oxidase';'reductase';'oxidoreductase';'Fic';}};
%
% % CONTEXT = 1; pat={'HAD hydrolase';'glycosyl hydrolase'; 'glycoside
hydrolase';'glycosylase';'glycosylhydrolase';'glycosidehydrolase';'glycos
idase';...
% %
'dehalogenase';'galactosidase';'phosphatase';'hexosaminidase';};


 % if you have more scores, then you have to make a string for file names
from them
t = regexprep(num2str(SCORES), '\s+', ' '); t(find(t == ' ')) = '_';

id = fopen([genus '_' t '_cands.fa_syntenic.broader'], 'w');
id1 = fopen([genus '_' t '_cands.fa_syntenic'], 'w');
if ~exist('INDEXES','var'), INDEXES = 1:length(gs); end
id2 = fopen([genus '_' t '_cands.fa_cntxs'], 'w');
for iINDEXES = 1 : length(INDEXES)
      igs = INDEXES(iINDEXES);
      for ins = 1 : length(ns{igs})
% get IGRs with candidates
            [origigrname origigrname] = unix(['ls ' d gs{igs} '/*r'
num2str(ns{igs}(ins)) '.fa']);
            origigrname(length(origigrname)) = [];
            [origigr origigr] = unix(['head -n2 ' d gs{igs} '/*r'
num2str(ns{igs}(ins)) '.fa']);
            if ~contains(origigr, 'head: cannot open')
% you also mistakenly put >> instead of > in the IGR fasta headers and
rewrite it for the case it is not rewritten later beacuse of the bad ID
                  if origigr(2) == '>'
                        origigr(2) = [];
                        unix(['echo -e ''' origigr ''' > ' origigrname]);
                  end
                  s=strsplit(origigr, char(10));
                  s=strsplit(s{1}, ' ');
                  c2 = str2num(s{length(s)});
                  c1 = str2num(s{length(s) - 1});
                  if c1 > c2
                        a = c2;
                        c2=c1;
                        c1=a;
                  end
% you put IDs of the first contigs in the IGR fasta files even when they
were from other configs; so now you have to find IDs of the right contigs
and put them
% in the original IGR fastas to be able to identify the contigs later by
f_get_synteny_from_fasta_comb.m. You also named IGR fastas with forward
sequences
% *_rc_igr*.fa, and vice versa; the IGR coordinates witihin the fastas
are all right.
```

```
                [r r] = unix(['grep ' s{1}(2:length(s{1})) ' ' ANNS_SA
'/' s{2} '/*fna*']);
                if ~contains(r, 'No such file or directory')
                    r=strsplit(r,':');
                    r=strsplit(r{1}, '/');
                    r=strsplit(r{length(r)}, '_');
                    [g g] = unix(['grep -w "'  num2str(c1 - 1) '.*[-
+]" ' ANNS_SA '/' s{2} '/' r{1} '_' r{2} '*fea* -A1|grep -w "' num2str(c2
+ 1) '.*[-+]"']);
    % it must be a single hit
                    if ~isempty(g)
                        if length(find(g==char(10))) ==
1%findCommonWords(g, 'chromosome', 0, 0) == 1 %length(strfind(g,
'chromosome')) == 1
                            g = strsplit(g, char(9));
    % dig out the new correct ID from the grep
                            origigr = strsplit(origigr, char(10));
                            h1 = strsplit(origigr{1}, ' ');
                            v = find(strcmp(g, '+') | strcmp(g, '-
'))-3;
    % does the contig ID obtained by grep of IGR coors now match the
contig ID in the IGR fasta?
                            if ~strcmp(h1{1}(2:length(h1{1})),
g{v})
                                r = ['>' g{v}];
                                for ih = 2 : length(h1)
                                    r = [r ' ' h1{ih}];
                                end
                                ido1 = fopen(origigrname, 'w');
fwrite(ido1, [r char(10) origigr{2} char(10) ]); fclose(ido1);
                            else
                %                   disp('there was the right ID in
the IGR fasta.')
                            end
                        else
                            disp([g ': more than one hit for this
IGR in *fea*. Leave the original ID in the IGR fasta for synteny.'])
    %                       pause
                        end
                    else
                        disp([g ': IGR not found in *fea*, e.g. for
contigs starts/ends. Leave the original ID in the IGR fasta for synteny
and going on...']);
    %                   pause
                    end

                    [names synteny syntenyext syntenyextext
syntenyextextext syntenytable syntenytableplain gls glsn namesext] =
f_get_synteny_from_fasta_comb({origigrname}, ANNS_SA, 1) ;
                    if ~isempty(syntenytable)
                        if ADJACENTONLY
                            syntenytable = replace(synteny, ' - ',
' : '); syntenytable = {[names{1} '::' syntenytable{1}]};
                        end
                        if SIX
                            s = replace(syntenytable{1}, '::',
'%');
                            s = replace(s, ' : ', '%');
```

```
                            s= replace(s, ' - ', '%');
                            s = strsplit(s,'%',
'CollapseDelimiters',false);
                            syntenytable = {[s{1} '::' s{3} ' - '
s{4} ' - ' s{5} ' - ' s{6} ' - ' s{7} ' - ' s{8} ' : ' ...
                                                        s{9} ' - ' s{10}
' - ' s{11} ' - ' s{12} ' - ' s{13} ' - ' s{14}]};
                        end
                        hit=0;
                        fwrite(id2,[syntenytable{1} char(10)]);
                        if iscell(pat{1})
          % substrings in both sides - you compare all words in the
string no matter their order
          %                 syntenytable
                            s = strsplit(syntenytable{1}, ' : ');
                            pat1i=0;
                            pat2i=0;
                            for ipat = 1 : length(pat{1})
                                if findCommonWords(s{1},
pat{1}{ipat}, requireAll, allowSubstring)
                                    pat1i=1;
                                    break
                                end
                            end
                            if ~pat1i
                                for ipat = 1 : length(pat{1})
                                    if findCommonWords(s{2},
pat{1}{ipat}, requireAll, allowSubstring)
                                        pat1i=2;
                                        break
                                    end
                                end
                            end
                            for ipat = 1 : length(pat{2})
                                if findCommonWords(s{1},
pat{2}{ipat}, requireAll, allowSubstring)
                                    pat2i=1;
                                    break
                                end
                            end
                            if ~pat2i
                                for ipat = 1 : length(pat{2})
                                    if findCommonWords(s{2},
pat{2}{ipat}, requireAll, allowSubstring)
                                        pat2i=2;
                                        break
                                    end
                                end
                            end
                            if CONTEXT == 2
      % any string of one sets in any of sides, any string of the other
set in the other side
                                if (pat1i==1 & pat2i==2) ||
(pat1i==2 & pat2i==1)
                                    hit = 1;
                                end
                            end
                            if CONTEXT == 1
```

```
      % any string of both sets in one of the sides
                                    if (pat1i==1 & pat2i==1) ||
(pat1i==2 & pat2i==2)
                                          hit = 1;
                                    end
                              end
                        else
      % any string of the set in any side
                              for ipat = 1 : length(pat)
                                    if
findCommonWords(syntenytable{1}, pat{ipat}, requireAll, allowSubstring)
                                          hit=1;
                                          break
                                    end
                              end
                        end
                        if hit
      % find sqs of syntenic candidates in their IGRs
                              igr = fastaread(origigrname);
                              z = strsplit(gs{igs}, '_');
      % find the rsol dir
                              if strcmp(SWITCH, 'for_Glaciecola')
                                    x = strsplit(igr(1).Header, ' ');
                                    [d1 d1] = unix(['ls ' d x{2}(1:4)
z{3} SUFFIX '/ -d']);
                              end
                              if strcmp(SWITCH, 'for_Bifi')
                                    [d1 d1] = unix(['ls ' d z{2} z{3}
SUFFIX '/ -d']);
                              end
      % % for Sali
      %                       [d1 d1] = unix(['ls ' d upper(z{2}(1))
z{2}(2:length(z{2})) z{3}(isletter(z{3})) SUFFIX '_'
z{3}(~isletter(z{3})) '/ -d']);
                              if strcmp(SWITCH,
'for_Idiomarina_Frankia_Pseudoalterom')
                                    [d1 d1] = unix(['ls ' d
upper(z{2}(1)) z{2}(2:length(z{2})) z{3} SUFFIX '/ -d']);
                              end
                              if strcmp(SWITCH, 'for_Frankia_Sco')
                                    [d1 d1] = unix(['ls ' d
upper(z{2}(1)) z{2}(2:length(z{2})) z{3}(1 : length(z{3}) - 3) SUFFIX '/
-d']);
                              end
                              if isempty(strfind(d1, 'No such file or
directory'))
                                    d1(length(d1)) = [];
                                    [x x] = unix(['ls ' d1 '*_igr'
num2str(ns{igs}(ins)) '.fa_*.inf']);
                                    if isempty(strfind(x, 'No such
file or directory'))
                                          if sum(x==char(10)) > 1
                                                i = find(x==char(10));
                                                x = x(1 : i(1));
                                          end
                                          id0 = fopen(x(1 : length(x)
- 1)); x = textscan(id0, '%s','delimiter',char(10)); fclose(id0);
```

```
                                                line = x{1}{strmatch('Best
set #', x{1})};
                                                lines = strsplit(line, '#');
                                                scores = strsplit(line,
'=');
                                                scores2=strsplit(scores{2},
' ');
                                                scores3=strsplit(scores{3},
'.');
                                                scores4=strsplit(scores{4},
'.');
            % best str of a set
                                                lines2 = strsplit(lines{2},
' ');
                                                [x x] = unix(['ls ' d1
'*_igr' num2str(ns{igs}(ins)) '.fa_*_' num2str((str2num(lines2{1}) - 1) *
5 + 1) 'nt*.db']);
                                                x1 = strsplit(x, '_');
                                                x1 =
strsplit(x1{length(x1)}, 'th');
                                                [bssq bssq] = unix(['head -
n2 ' x(1 : length(x) - 1) ' | tail -n1']);
            % best str at all
                                                lines3 = strsplit(lines{3},
' ');
                                                [xx xx] = unix(['ls ' d1
'*_igr' num2str(ns{igs}(ins)) '.fa_*_' num2str((str2num(lines3{1}) - 1) *
5 + 1) 'nt*.db']);
                                                x2 = strsplit(xx, '_');
                                                x2 =
strsplit(x2{length(x2)}, 'th');
                                                [basq basq] = unix(['head -
n2 ' xx(1 : length(xx) - 1) ' | tail -n1']);
      %                                         if ~strcmp(bssq, basq)
                                                      fwrite(id1, ['>'
igr(1).Header ': best set #' num2str(lines2{1}) ' with sc='
num2str(scores2{1}) ...
                                                                      '
with best sopt #' num2str(x1{1}) ' with TED=' num2str(scores3{1}) ', ' x
bssq ...
                                                                      '>'
igr(1).Header ': best-at-all sopt #' num2str(x2{1}) ' with TED='
num2str(scores4{1}) ...
                                                                      '
from set #' num2str(lines3{1}) ', ' xx basq]);
      %                                         else
      %                                               fwrite(id1, ['>'
igr(1).Header ' best set #' num2str(lines2{1}) ' best sopt #'
num2str(x1{1}) char(10) bssq]);
      %                                         end
      s = replace(syntenytable{1}, '::', ': ');
      s = replace(s, ' : ', ' # ');
      s = replace(s, ' - ', ' | ');
      % fwrite(id, [s char(10)]);
      % % its for diagnostic purposes
      fwrite(id, [origigrname ' ' igr(1).Header ' best-at-all sopt #'
num2str(x2{1}) ' with TED=' num2str(scores4{1}) ': ' s char(10)]);
                                          else
```

```
                                        disp(['No such file or
directory for: ' d1 '*_igr' num2str(ns{igs}(ins)) '.fa_*.inf']); return
                                   end
                              else
                                   disp(['No such file or directory
for: ' d upper(z{2}(1)) z{2}(2:length(z{2})) z{3} SUFFIX '/.']);return
                              end
                         end
                    end
               else
                    disp(r), return
               end
          else
               disp(['head: cannot open ' origigr]), return
          end
     end
end
fclose(id);fclose(id1);fclose(id2);
% remove identical candidates contexts
% unix(['sort -u ' genus '_' t '_cands.fa_cntxs -o ' genus '_' t
'_cands.fa_cntxs']);
% remove identical candidates as for some there are both genomic
orientaions
% unix(['sort -u ' genus '_' t '_cands.fa_syntenic.broader -o ' genus '_'
t '_cands.fa_syntenic.broader']);
% visualize_broader_contexts([genus '_' t '_cands.fa_syntenic.broader'])
% sort_broader_context([genus '_' t
'_cands.fa_syntenic.broader']);visualize_broader_contexts([genus '_' t
'_cands.fa_syntenic.broader.sorted'])
clear f i fns ist s id si ipat id1 sqs o pat1i pat2i gs fn namesext gls
glsn sqs hit id0 basq x x2 lines lines3 line d1 d z origigrname
clear ANNS_SA            c2                 ih                 r
syntenytableplain
clear SUFFIX             g                  ins                synteny
t
clear a                  genus              lines2             syntenyext
v          requireAll       SIX
clear h1                 names              syntenyextext      x1
clear bssq               igr                ns
syntenyextextext   allowSubstring
clear c1                 igs                origigr
syntenytable      CONTEXT expr words ADJACENTONLY SWITCH INDEXES iINDEXES
id2


f_get_synteny_from_fasta_comb.m

function [names synteny syntenyext syntenyextext syntenyextextext
syntenytable syntenytableplain gls glsn namesext] = ...
            f_get_synteny_from_fasta_comb(fns, ANNS_SA,
DO_5th_6th_7th_FLANKING_GENES)

TRESHBLAST = .05;
% TRESHBLAST = 1e-20;

for ifns = 1 : length(fns)
     fn = fns{ifns};
```

```
      INIGRONLY = 1;
      NOTINNAMES =
["MAG","MAG:","DNA","chromosome",",","complete","genome","MAG:","TPA_asm"
,"TPA_asm:","candidatus","cf.","Candidatus",...
                "sp.", "strain","isolate",
"bacterium","UNVERIFIED_ORG:","Sample","type","genome","assembly",":"," I
",...
                   "whole", "shotgun", "sequence", "uncultured"];

      WRITEOUT = 0;
      TRYTOLOAD = 0;
      DO_FLANKING_GENES = 0;
      DO_IDS_OF_FLANKING_GENES = 0;
      INCLUDERRNAS=1;
      INCLUDENCRNAS=0;
      INCLUDETRNAS=1;
      REMOVEPSEUDOGENES = 0;
      DO_FLANKING_IGRS = 0;
      WRITETOFILE = 0;
      OVERLAP = 2;
% for reverse compatibility
      if ~exist('DO_5th_6th_7th_FLANKING_GENES'),
DO_5th_6th_7th_FLANKING_GENES = 0; end

% SOMETIMES SYNTENY AND SYNTENYEXT CAN CONTAIN EMPTY STRINGS AS 1ST AND
LAST ANNOTATION STRINGS ARE EMPTY COS THE INDEXING OF IGRS (SEE
F_READ_ANNS.M)
      gls = []; names = {}; namesext={}; synteny = {}; syntenyext = {};
syntenyextext = {}; syntenyextextext = {}; glsn = []; syntenytable = {};
syntenytableplain = {};
      if DO_FLANKING_GENES
            if ~exist('fgs', 'dir')
                  !mkdir  fgs
            end
            if WRITETOFILE
                  idd1 = fopen(['fgs/' fn '_d1.fa'],'w'); idd2 =
fopen(['fgs/' fn '_d2.fa'],'w'); idd3 = fopen(['fgs/' fn '_d3.fa'],'w');
idd4 = fopen(['fgs/' fn '_d4.fa'],'w');
                  idu1 = fopen(['fgs/' fn '_u1.fa'],'w'); idu2 =
fopen(['fgs/' fn '_u2.fa'],'w'); idu3 = fopen(['fgs/' fn '_u3.fa'],'w');
idu4 = fopen(['fgs/' fn '_u4.fa'],'w');
            end
            count = 0; count_special_u1 = 1; count_special_d1 = 1;
count_special_u2 = 1; count_special_d2 = 1; count_special_u3 = 1;
count_special_d3 = 1;
            count_special_u4 = 1; count_special_d4 = 1;
      end
      if DO_IDS_OF_FLANKING_GENES
            if WRITETOFILE,
                  idsd1 = fopen(['id/' fn '_d1.id'],'w'); idsd2 =
fopen(['id/' fn '_d2.id'],'w'); idsd3 = fopen(['id/' fn '_d3.id'],'w');
idsd4 = fopen(['id/' fn '_d4.id'],'w');
                  idsu1 = fopen(['id/' fn '_u1.id'],'w'); idsu2 =
fopen(['id/' fn '_u2.id'],'w'); idsu3 = fopen(['id/' fn '_u3.id'],'w');
idsu4 = fopen(['id/' fn '_u4.id'],'w');
            end
      end
      if DO_FLANKING_IGRS
```

```
            if ~exist('igrs', 'dir')
                  !mkdir igrs
            end
            if WRITETOFILE,
                  idid1 = fopen(['igrs/' fn '_id1.fa'],'w'); idid2 =
fopen(['igrs/' fn '_id2.fa'],'w'); idid3 = fopen(['igrs/' fn
'_id3.fa'],'w');
            idiu1 = fopen(['igrs/' fn '_iu1.fa'],'w'); idiu2 =
fopen(['igrs/' fn '_iu2.fa'],'w'); idiu3 = fopen(['igrs/' fn
'_iu3.fa'],'w');
            end
      end

      if ANNS_SA(length(ANNS_SA)) ~= '/', ANNS_SA = [ANNS_SA '/']; end
      % if ANNS(length(ANNS)) == '/', ANNS(length(ANNS)) = []; end

      f = fastaread(fn);
      if WRITETOFILE, id = fopen([fn '.igr'], 'w');end

      if WRITEOUT, ido = fopen('get_synteny_from_fasta_comb.m.out', 'w');
end

      for fi = 1 : length(f)
            if DO_FLANKING_GENES
                  count_flag_u1 = 0; count_flag_d1 = 0; count_flag_u2 = 0;
count_flag_d2 = 0;count_flag_u3 = 0; count_flag_d3 = 0;count_flag_u4 = 0;
count_flag_d4 = 0;
            end

            s = ['Doing ' num2str(fi) '/' num2str(length(f)) ' '
f(fi).Header '.']; disp(s), if WRITEOUT, fwrite(ido, [s char(10)]); end

words = strsplit(f(fi).Header);
words = words(~ismember(words, NOTINNAMES));
f(fi).Header = strjoin(words, ' ');


            header = cut_string(f(fi).Header, ' ');
            [fna fna] = unix(['grep -l ' header{1} ' ' ANNS_SA header{2}
'/*.fna*']);
            if ~isempty(strfind(fna, 'No such file or directory')) |
isempty(fna)
                  s = ['Either ' ANNS_SA header{2} ' not exist or '
header{1} ' not found by grep in ' ANNS_SA header{2} '.']; disp(s), if
WRITEOUT, fwrite(ido, [s char(10)]);end
                  [fna fea] = f_try_anns(header{1}, [ANNS_SA header{2}],
TRYTOLOAD, 1);
                  if isempty(fna)
                        s = ['Could not download annotations for '
header{2} ' ' header{3} ': ' fea '.']; disp(s), if WRITEOUT, fwrite(ido,
[s char(10)]); end
                  else
      % move newly downloaded anns into their genus folders
                        if ~exist([ANNS_SA header{2}], 'dir')
                              unix(['mkdir ' ANNS_SA header{2}]);
                        end
                  end
            end
```

```
            if ~isempty(fna)
                  if fna(length(fna)) == char(10), fna(length(fna)) = [];
end
                  i = find(fna == char(10));
                  if ~isempty(i)
                        fna(i : length(fna)) = [];
                  end
                  i = strfind(fna, 'genomic.fna');
                  fea = [fna(1 : i - 1) 'feature_table.txt' fna(i +
length('genomic.fna') : length(fna))];
                  if ~exist(fea, 'file')
                        disp([fea ' does not exist.'])
                        pause
                  else
                        [x x] = unix(['head -n1 ' fna]);
                        [j g] = f_read_genome(fna);
                        [crs ors anns ids] = f_read_anns(fea, length(g),
INCLUDERRNAS , INCLUDENCRNAS, INCLUDETRNAS, REMOVEPSEUDOGENES);
                        if ~isempty(crs)
                              if str2num(header{length(header) - 1}) >
str2num(header{length(header)})
                                    i = find(crs(:, 2) <=
str2num(header{length(header)}) + OVERLAP);
                                    j = find(crs(:, 1) >=
str2num(header{length(header) - 1}) - OVERLAP);
                              else
                                    i = find(crs(:, 2) <=
str2num(header{length(header) - 1}) + OVERLAP);
                                    j = find(crs(:, 1) >=
str2num(header{length(header)}) - OVERLAP);
                              end
                              if ~isempty(j) & ~isempty(i)
                                    if j(1) - i(length(i)) == 1 | INIGRONLY
== 0
                                          if DO_FLANKING_GENES
                                                count = count + 1;
                                          end
                  % IT IS EITHER IN IGR OR IT IS NOT BUT YOU TAKE
EVERYTHING.
                                          if j(1) - i(length(i)) ~= 1
                                                s = 'It is not in IGR but
you take everything.'; disp(s), if WRITEOUT, fwrite(ido, [s
char(10)]);end
                                          else
                                                s = 'It is in IGR.';
disp(s), if WRITEOUT, fwrite(ido, [s char(10)]);end
                                          end
                                          if isempty([strfind(lower(x),
'contig') strfind(lower(x), 'scaffold')])
                                                gls = [gls;
[str2num(header{length(header) - 1}) str2num(header{length(header)})]];
                                                glsn = [glsn
(str2num(header{length(header) - 1}) + str2num(header{length(header)}) /
2) / length(g)];
                                          else
                                                gls = [gls; [0 0]];
                                                glsn = [glsn 0];
```

```
                                                disp('Contig or scaffold: 
gls=[0 0], glsn = 0.')
                                          end
                                          name = [header{2} ' ' header{3}];
                                          if length(header) > 3
                                                name = [name ' ' header{4}];
                                                if length(header) > 4
                                                      name = [name ' ' 
header{5}];
                                                end
                                          end
                                          names = [names; name];
                                          namesext = [namesext; 
f(fi).Header];
                                          synteny = [synteny; 
[anns{i(length(i))} ' - ' anns{j(1)}]];
                                          b = [anns{i(length(i))} ' : ' 
anns{j(1)}];
                                          c = [anns{i(length(i))} ' - ' 
anns{j(1)}];
            % here the genomic orientation is given by BLAST hit
                                          if str2num(header{length(header) 
- 1}) > str2num(header{length(header)})
                                                if WRITETOFILE, fwrite(id, 
['>' f(fi).Header char(10) seqrcomplement(g(crs(i(length(i)), 2) + 1 : 
crs(j(1), 1) - 1)) char(10)]);end
                                          else
                                                if WRITETOFILE,fwrite(id, 
['>' f(fi).Header char(10) g(crs(i(length(i)), 2) + 1 : crs(j(1), 1) - 1) 
char(10)]);end
                                          end
                                          if DO_FLANKING_GENES
                                                if i(length(i)) > 1
                                                      count_flag_d1 = 1;
                                                      if 
WRITETOFILE,fwrite(idd1, ['>' num2str(count_special_d1) ' ' num2str(fi) ' 
' anns{i(length(i))} char(10) rna2dna(g(crs(i(length(i)), 1) : 
crs(i(length(i)), 2))) char(10)]);end
                                                      if 
DO_IDS_OF_FLANKING_GENES
                                                            if 
WRITETOFILE,fwrite(idsd1, [ids{i(length(i))} char(10)]);end
                                                      end
                                                end
                                                if j(1) < length(anns)
                                                      count_flag_u1 = 1;
                                                      if 
WRITETOFILE,fwrite(idu1, ['>' num2str(count_special_u1) ' ' num2str(fi) ' 
' anns{j(1)} char(10) rna2dna(g(crs(j(1), 1) : crs(j(1), 2))) 
char(10)]);end
                                                      if 
DO_IDS_OF_FLANKING_GENES
                                                            if 
WRITETOFILE,fwrite(idsu1, [ids{j(1)} char(10)]);end
                                                      end
                                                end
                                          end
```

```
                                        a = ' - ';
                                        if i(length(i)) - 1 > 0
                                             a = [anns{i(length(i)) - 1}
a];
                                             b = [anns{i(length(i)) - 1}
' - ' b];
                                             c = [anns{i(length(i)) - 1}
' ' c];
                                             if i(length(i)) - 1 > 1
                                                  if DO_FLANKING_GENES
                                                       count_flag_d2 =
1;
                                                       if
WRITETOFILE,fwrite(idd2, ['>' num2str(count_special_d2) ' ' num2str(fi) '
' anns{i(length(i)) - 1} char(10) rna2dna(g(crs(i(length(i)) - 1, 1) :
crs(i(length(i)) - 1, 2))) char(10)]);end
                                                  end
                                                  if
DO_IDS_OF_FLANKING_GENES
                                                       if
WRITETOFILE,fwrite(idsd2, [ids{i(length(i)) - 1} char(10)]);end
                                                  end
                                                  if DO_FLANKING_IGRS
           % genomic orientation is unknown for the other IGRs
                                                       if
WRITETOFILE,fwrite(idid1, ['>' header{1} ' ' header{2} ' ' header{3} ' '
header{4} ' ' header{5} ' ' num2str(crs(i(length(i)) - 1, 2)) ' '
num2str(crs(i(length(i)), 1)) char(10) ...

g(crs(i(length(i)) - 1, 2) : crs(i(length(i)), 1)) char(10)]);end
                                                  end
                                             end
                                        else
                                             b = ['0 - ' b];
                                             c = ['0 ' c];
                                        end
                                        if j(1) + 1 <= length(anns)
                                             a = [a anns{j(1) + 1}];
                                             b = [b ' - ' anns{j(1) +
1}];
                                             c = [c ' ' anns{j(1) + 1}];
                                             if j(1) + 1 < length(anns)
                                                  if DO_FLANKING_GENES
                                                       count_flag_u2 =
1;
                                                       if
WRITETOFILE,fwrite(idu2, ['>'  num2str(count_special_u2) ' ' num2str(fi)
' ' anns{j(1) + 1} char(10) rna2dna(g(crs(j(1) + 1, 1) : crs(j(1) + 1,
2))) char(10)]);end
                                                  end
                                                  if
DO_IDS_OF_FLANKING_GENES
                                                       if
WRITETOFILE,fwrite(idsu2, [ids{j(1) + 1} char(10)]);end
                                                  end
                                                  if DO_FLANKING_IGRS
           % genomic orientation is unknown for the other IGRs
```

```
                                                            if
WRITETOFILE,fwrite(idiu1, ['>' header{1} ' ' header{2} ' ' header{3} ' '
header{4} ' ' header{5} ' ' num2str(crs(j(1), 2)) ' ' num2str(crs(j(1) +
1, 1)) char(10) ...

g(crs(j(1), 2) : crs(j(1) + 1, 1)) char(10)]);end
                                                      end
                                                end
                                          else
                                                b = [b ' - 0'];
                                                c = [c ' 0'];
                                          end
                                          syntenyext = [syntenyext; a];

                                          a = ' - ';
                                          if i(length(i)) - 2 > 0
                                                a = [anns{i(length(i)) - 2}
a];

                                                b = [anns{i(length(i)) - 2}
' - ' b];

                                                c = [anns{i(length(i)) - 2}
' ' c];

                                                if i(length(i)) - 2 > 1
                                                      if DO_FLANKING_GENES
                                                            count_flag_d3 =
1;

                                                            if
WRITETOFILE,fwrite(idd3, ['>' num2str(count_special_d3) ' ' num2str(fi) '
' anns{i(length(i)) - 2} char(10) rna2dna(g(crs(i(length(i)) - 2, 1) :
crs(i(length(i)) - 2, 2))) char(10)]);end
                                                      end
                                                      if
DO_IDS_OF_FLANKING_GENES
                                                            if
WRITETOFILE,fwrite(idsd3, [ids{i(length(i)) - 2} char(10)]);end
                                                      end
                                                      if DO_FLANKING_IGRS
            % genomic orientation is unknown for the other IGRs
                                                            if
WRITETOFILE,fwrite(idid2, ['>' header{1} ' ' header{2} ' ' header{3} ' '
header{4} ' ' header{5} ' ' num2str(crs(i(length(i)) - 2, 2)) ' '
num2str(crs(i(length(i)) - 1, 1)) char(10) ...

g(crs(i(length(i)) - 2, 2) : crs(i(length(i)) - 1, 1)) char(10)]);end
                                                      end
                                                end
                                          else
                                                b = ['0 - ' b];
                                                c = ['0 ' c];
                                          end
                                          if j(1) + 2 <= length(anns)
                                                a = [a anns{j(1) + 2}];
                                                b = [b ' - ' anns{j(1) +
2}];

                                                c = [c ' ' anns{j(1) + 2}];
                                                if j(1) + 2 < length(anns)
                                                      if DO_FLANKING_GENES
```

```
                                                  count_flag_u3 =
1;
                                                  if
WRITETOFILE,fwrite(idu3, ['>' num2str(count_special_u3) ' ' num2str(fi) '
' anns{j(1) + 2} char(10) rna2dna(g(crs(j(1) + 2, 1) : crs(j(1) + 2, 2)))
char(10)]);end
                                             end
                                             if
DO_IDS_OF_FLANKING_GENES
                                                  if
WRITETOFILE,fwrite(idsu3, [ids{j(1) + 2} char(10)]);end
                                             end
                                             if DO_FLANKING_IGRS
          % genomic orientation is unknown for the other IGRs
                                                  if
WRITETOFILE,fwrite(idiu2, ['>' header{1} ' ' header{2} ' ' header{3} ' '
header{4} ' ' header{5} ' ' num2str(crs(j(1) + 1, 2)) ' '
num2str(crs(j(1) + 2, 1)) char(10) ...

g(crs(j(1) + 1, 2) : crs(j(1) + 2, 1)) char(10)]);end
                                             end
                                        end
                                   else
                                        b = [b ' - 0'];
                                        c = [c ' 0'];
                                   end
                                   syntenyextext = [syntenyextext;
a];

                                   a = ' - ';
                                   if i(length(i)) - 3 > 0
                                        a = [anns{i(length(i)) - 3}
a];
                                        b = [anns{i(length(i)) - 3}
' - ' b];
                                        c = [anns{i(length(i)) - 3}
' ' c];
                                        if i(length(i)) - 3 > 1
                                             if DO_FLANKING_GENES
                                                  count_flag_d4 =
1;
                                                  if
WRITETOFILE,fwrite(idd4, ['>' num2str(count_special_d4) ' ' num2str(fi) '
' anns{i(length(i)) - 3} char(10) rna2dna(g(crs(i(length(i)) - 3, 1) :
crs(i(length(i)) - 3, 2))) char(10)]);end
                                             end
                                             if
DO_IDS_OF_FLANKING_GENES
                                                  if
WRITETOFILE,fwrite(idsd4, [ids{i(length(i)) - 3} char(10)]);end
                                             end
                                             if DO_FLANKING_IGRS
          % genomic orientation is unknown for the other IGRs
                                                  if
WRITETOFILE,fwrite(idid3, ['>' header{1} ' ' header{2} ' ' header{3} ' '
header{4} ' ' header{5} ' ' num2str(crs(i(length(i)) - 3, 2)) ' '
num2str(crs(i(length(i)) - 2, 1)) char(10) ...
```

```
g(crs(i(length(i)) - 3, 2) : crs(i(length(i)) - 2, 1)) char(10)]);end
                                                  end
                                             end
                                        else
                                             b = ['0 - ' b];
                                             c = ['0 ' c];

                                        end
                                        if j(1) + 3 <= length(anns)
                                             a = [a anns{j(1) + 3}];
                                             b = [b ' - ' anns{j(1) +
3}];

                                             c = [c ' ' anns{j(1) + 3}];
                                             if j(1) + 3 < length(anns)
                                                  if DO_FLANKING_GENES
                                                       count_flag_u4 =
1;
                                                       if
WRITETOFILE,fwrite(idu4, ['>' num2str(count_special_u4) ' ' num2str(fi) '
' anns{j(1) + 3} char(10) rna2dna(g(crs(j(1) + 3, 1) : crs(j(1) + 3, 2)))
char(10)]);end
                                                  end
                                                  if
DO_IDS_OF_FLANKING_GENES
                                                       if
WRITETOFILE,fwrite(idsu4, [ids{j(1) + 3} char(10)]);end
                                                  end
                                                  if DO_FLANKING_IGRS
           % genomic orientation is unknown for the other IGRs
                                                       if
WRITETOFILE,fwrite(idiu3, ['>' header{1} ' ' header{2} ' ' header{3} ' '
header{4} ' ' header{5} ' ' num2str(crs(j(1) + 2, 2)) ' '
num2str(crs(j(1) + 3, 1)) char(10) ...

g(crs(j(1) + 2, 2) : crs(j(1) + 3, 1)) char(10)]);end
                                                  end
                                             end
                                        else
                                             b = [b ' - 0'];
                                             c = [c ' 0'];
                                        end
                                        syntenyextextext =
[syntenyextextext; a];

                                        if DO_5th_6th_7th_FLANKING_GENES
                                             a = ' - ';
                                             if i(length(i)) - 4 > 0
%                                                 a = [anns{i(length(i))
- 4} a];

                                                  b = [anns{i(length(i))
- 4} ' - ' b];

                                                  c = [anns{i(length(i))
- 4} ' ' c];
     %                                            if i(length(i)) - 3 >
1
     %                                                 if
DO_FLANKING_GENES
```

```
      %
      count_flag_d4 = 1;
      %                                                         if
WRITETOFILE,fwrite(idd4, ['>' num2str(count_special_d4) ' ' num2str(fi) '
' anns{i(length(i)) - 3} char(10) rna2dna(g(crs(i(length(i)) - 3, 1) :
crs(i(length(i)) - 3, 2))) char(10)]);end
      %                                                   end
      %                                                   if
DO_IDS_OF_FLANKING_GENES
      %                                                         if
WRITETOFILE,fwrite(idsd4, [ids{i(length(i)) - 3} char(10)]);end
      %                                                   end
      %                                                   if
DO_FLANKING_IGRS
      %          % genomic orientation is unknown for the other IGRs
      %                                                         if
WRITETOFILE,fwrite(idid3, ['>' header{1} ' ' header{2} ' ' header{3} ' '
header{4} ' ' header{5} ' ' num2str(crs(i(length(i)) - 3, 2)) ' '
num2str(crs(i(length(i)) - 2, 1)) char(10) ...
      %
         g(crs(i(length(i)) - 3, 2) : crs(i(length(i)) - 2, 1))
char(10)]);end
      %                                                   end
      %                                             end
                                              else
                                                    b = ['0 - ' b];
                                                    c = ['0 ' c];
                                              end
                                              if j(1) + 4 <= length(anns)
%                                                   a = [a anns{j(1) +
4}];

                                                    b = [b ' - ' anns{j(1)
+ 4}];
                                                    c = [c ' ' anns{j(1) +
4}];
      %                                             if j(1) + 3 <
length(anns)
      %                                                   if
DO_FLANKING_GENES
      %
      count_flag_u4 = 1;
      %                                                         if
WRITETOFILE,fwrite(idu4, ['>' num2str(count_special_u4) ' ' num2str(fi) '
' anns{j(1) + 3} char(10) rna2dna(g(crs(j(1) + 3, 1) : crs(j(1) + 3, 2)))
char(10)]);end
      %                                                   end
      %                                                   if
DO_IDS_OF_FLANKING_GENES
      %                                                         if
WRITETOFILE,fwrite(idsu4, [ids{j(1) + 3} char(10)]);end
      %                                                   end
      %                                                   if
DO_FLANKING_IGRS
      %          % genomic orientation is unknown for the other IGRs
      %                                                         if
WRITETOFILE,fwrite(idiu3, ['>' header{1} ' ' header{2} ' ' header{3} ' '
header{4} ' ' header{5} ' ' num2str(crs(j(1) + 2, 2)) ' '
num2str(crs(j(1) + 3, 1)) char(10) ...
```

```
      %
         g(crs(j(1) + 2, 2) : crs(j(1) + 3, 1)) char(10)]);end
      %                                                    end
      %                                              end
                                               else
                                                     b = [b ' - 0'];
                                                     c = [c ' 0'];
                                               end
%                                              syntenyextextext =
[syntenyextextext; a];

                                               a = ' - ';
                                               if i(length(i)) - 5 > 0
%                                                    a = [anns{i(length(i))
- 5} a];
                                                     b = [anns{i(length(i))
- 5} ' - ' b];
                                                     c = [anns{i(length(i))
- 5} ' ' c];
      %                                              if i(length(i)) - 3 >
1
      %                                                    if
DO_FLANKING_GENES
      %
      count_flag_d4 = 1;
      %                                                          if
WRITETOFILE,fwrite(idd4, ['>' num2str(count_special_d4) ' ' num2str(fi) '
' anns{i(length(i)) - 3} char(10) rna2dna(g(crs(i(length(i)) - 3, 1) :
crs(i(length(i)) - 3, 2))) char(10)]);end
      %                                                    end
      %                                                    if
DO_IDS_OF_FLANKING_GENES
      %                                                          if
WRITETOFILE,fwrite(idsd4, [ids{i(length(i)) - 3} char(10)]);end
      %                                                    end
      %                                                    if
DO_FLANKING_IGRS
      %          % genomic orientation is unknown for the other IGRs
      %                                                          if
WRITETOFILE,fwrite(idid3, ['>' header{1} ' ' header{2} ' ' header{3} ' '
header{4} ' ' header{5} ' ' num2str(crs(i(length(i)) - 3, 2)) ' '
num2str(crs(i(length(i)) - 2, 1)) char(10) ...
      %
         g(crs(i(length(i)) - 3, 2) : crs(i(length(i)) - 2, 1))
char(10)]);end
      %                                                    end
      %                                              end
                                               else
                                                     b = ['0 - ' b];
                                                     c = ['0 ' c];
                                               end
                                               if j(1) + 5 <= length(anns)
%                                                    a = [a anns{j(1) +
5}];
                                                     b = [b ' - ' anns{j(1)
+ 5}];
                                                     c = [c ' ' anns{j(1) +
5}];
```

```
      %                                            if j(1) + 3 <
length(anns)
      %                                                  if
DO_FLANKING_GENES
      %
      count_flag_u4 = 1;
      %                                                        if
WRITETOFILE,fwrite(idu4, ['>' num2str(count_special_u4) ' ' num2str(fi) '
' anns{j(1) + 3} char(10) rna2dna(g(crs(j(1) + 3, 1) : crs(j(1) + 3, 2)))
char(10)]);end
      %                                                  end
      %                                                  if
DO_IDS_OF_FLANKING_GENES
      %                                                        if
WRITETOFILE,fwrite(idsu4, [ids{j(1) + 3} char(10)]);end
      %                                                  end
      %                                                  if
DO_FLANKING_IGRS
      %          % genomic orientation is unknown for the other IGRs
      %                                                        if
WRITETOFILE,fwrite(idiu3, ['>' header{1} ' ' header{2} ' ' header{3} ' '
header{4} ' ' header{5} ' ' num2str(crs(j(1) + 2, 2)) ' '
num2str(crs(j(1) + 3, 1)) char(10) ...
      %
         g(crs(j(1) + 2, 2) : crs(j(1) + 3, 1)) char(10)]);end
      %                                                  end
      %                                            end
                                             else
                                                   b = [b ' - 0'];
                                                   c = [c ' 0'];
                                             end
%                                            syntenyextextext =
[syntenyextextext; a];

                                             a = ' - ';
                                             if i(length(i)) -6 > 0
%                                                  a = [anns{i(length(i))
- 6} a];
                                                   b = [anns{i(length(i))
- 6} ' - ' b];
                                                   c = [anns{i(length(i))
- 6} ' ' c];
      %                                            if i(length(i)) - 3 >
1
      %                                                  if
DO_FLANKING_GENES
      %
      count_flag_d4 = 1;
      %                                                        if
WRITETOFILE,fwrite(idd4, ['>' num2str(count_special_d4) ' ' num2str(fi) '
' anns{i(length(i)) - 3} char(10) rna2dna(g(crs(i(length(i)) - 3, 1) :
crs(i(length(i)) - 3, 2))) char(10)]);end
      %                                                  end
      %                                                  if
DO_IDS_OF_FLANKING_GENES
      %                                                        if
WRITETOFILE,fwrite(idsd4, [ids{i(length(i)) - 3} char(10)]);end
      %                                                  end
```

```
      %                                                   if
DO_FLANKING_IGRS
      %          % genomic orientation is unknown for the other IGRs
      %                                                        if
WRITETOFILE,fwrite(idid3, ['>' header{1} ' ' header{2} ' ' header{3} ' '
header{4} ' ' header{5} ' ' num2str(crs(i(length(i)) - 3, 2)) ' '
num2str(crs(i(length(i)) - 2, 1)) char(10) ...
      %
         g(crs(i(length(i)) - 3, 2) : crs(i(length(i)) - 2, 1))
char(10)]);end
      %                                                   end
      %                                              end
                                                else
                                                     b = ['0 - ' b];
                                                     c = ['0 ' c];
                                                end
                                                if j(1) + 6 <= length(anns)
%                                                    a = [a anns{j(1) +
6}];

                                                     b = [b ' - ' anns{j(1)
+ 6}];
                                                     c = [c ' ' anns{j(1) +
6}];
      %                                              if j(1) + 3 <
length(anns)
      %                                                   if
DO_FLANKING_GENES
      %
      count_flag_u4 = 1;
      %                                                        if
WRITETOFILE,fwrite(idu4, ['>' num2str(count_special_u4) ' ' num2str(fi) '
' anns{j(1) + 3} char(10) rna2dna(g(crs(j(1) + 3, 1) : crs(j(1) + 3, 2)))
char(10)]);end
      %                                                   end
      %                                                   if
DO_IDS_OF_FLANKING_GENES
      %                                                        if
WRITETOFILE,fwrite(idsu4, [ids{j(1) + 3} char(10)]);end
      %                                                   end
      %                                                   if
DO_FLANKING_IGRS
      %          % genomic orientation is unknown for the other IGRs
      %                                                        if
WRITETOFILE,fwrite(idiu3, ['>' header{1} ' ' header{2} ' ' header{3} ' '
header{4} ' ' header{5} ' ' num2str(crs(j(1) + 2, 2)) ' '
num2str(crs(j(1) + 3, 1)) char(10) ...
      %
         g(crs(j(1) + 2, 2) : crs(j(1) + 3, 1)) char(10)]);end
      %                                                   end
      %                                              end
                                                else
                                                     b = [b ' - 0'];
                                                     c = [c ' 0'];
                                                end
%                                               syntenyextextext =
[syntenyextextext; a];
                                           end
```

```
                                        syntenytable = [syntenytable;
[name '::' b]];
                                        syntenytableplain =
[syntenytableplain; c];
                                  else
                                        s = 'It''s not in IGR and you
don''t take it.'; disp(s), if WRITEOUT, fwrite(ido, [s char(10)]);end
                                  end
                            else
                                  s = 'Either i or j is empty (e.g. when
only 1 gene in annotations or when at the end of contig/scaffold).';
disp(s), if WRITEOUT, fwrite(ido, [s char(10)]);end
                            end
                      end
                end
          end
          if DO_FLANKING_GENES
                if count_flag_u1, count_special_u1 = count_special_u1 +
1; end
                if count_flag_d1, count_special_d1 = count_special_d1 +
1; end
                if count_flag_u2, count_special_u2 = count_special_u2 +
1; end
                if count_flag_d2, count_special_d2 = count_special_d2 +
1; end
                if count_flag_u3, count_special_u3 = count_special_u3 +
1; end
                if count_flag_d3, count_special_d3 = count_special_d3 +
1; end
                if count_flag_u4, count_special_u4 = count_special_u4 +
1; end
                if count_flag_d4, count_special_d4 = count_special_d4 +
1; end
          end
      end
      if WRITETOFILE,
          if DO_FLANKING_GENES,
fclose(id);fclose(idd1);fclose(idd2);fclose(idd3);fclose(idd4);fclose(idu
1);fclose(idu2);fclose(idu3);fclose(idu4); end
      if DO_FLANKING_IGRS,
fclose(idid1);fclose(idid2);fclose(idid3);fclose(idiu1);fclose(idiu2);fcl
ose(idiu3); end
      if DO_IDS_OF_FLANKING_GENES,
fclose(idsd1);fclose(idsd2);fclose(idsd3);fclose(idsd4);fclose(idsu1);fcl
ose(idsu2);fclose(idsu3);fclose(idsu4); end
      end
%     [names synteny syntenyext]
%     clear ifns f n fi i g INIGRONLY TRYTOLOAD REMOVEPSEUDOGENES j crs
ors anns header a b s header fna fea NOTINHEADERS x y k WRITEOUT c id*
count* DO_FLANKING_GENES WRITETOFILE
%     save([fn '.igr.mat'], 'fn',
'gls','glsn','names','synteny','syntenyext','syntenyextext','syntenyextex
text','syntenytable','syntenytableplain')
end
```

f_find_all_words_of_str_in_str.m

```
function count = f_find_all_words_of_str_in_str(s1, s, SEPARATOR, EXACT,
CASESENS)
% count contains numbers of words in s found in the str s1

if ~exist('EXACT'), EXACT = 0; end
if ~exist('CASESENS'), CASESENS = 0; end
if ~exist('SEPARATOR'), SEPARATOR = ' '; end

if ~CASESENS, s = lower(s); s1 = lower(s1); end
s = cut_string(s, SEPARATOR);
count = 0;

for is = 1 : length(s)
    p = strfind(s1, s{is});
    if EXACT
        for ip = 1 : length(p)
% MATCHED STRINGS MUST BE THE SAME LONG AND HAVE NON-LETTER BEFORE AND
AFTER THE MATCH. IF THE MATCH STARTS AT THE START OF
% THE STRING, THEN CHECK ONLY THE CHARACTER FOLLOWING THE MATCH FOR BEING
NON-LETTER. ANALOGOUSLY IF THE END OF THE MATCH
% IS AT THE END OF THE STRING, CHECK ONLY THE CHARACTER PRECEDING THE
MATCH FOR BEING NON-LETTER.
            if (((p(ip) > 1 & ~isletter(s1(p(ip) - 1))) | p(ip) == 1) &
...
                ((p(ip) + length(s{is}) - 1 < length(s1) &
~isletter(s1(p(ip) + length(s{is})))) | p(ip) + length(s{is}) - 1 ==
length(s1)))
                    count = count + 1;
                    break
            end
        end
    else
        if ~isempty(p)
            count = count + 1;
        end
      end
end


f_make_header.m

function header = f_make_header(header, TERMS_SEPARATOR,
WANTED_TERMS_POSITIONS, WORDS_SEPARATOR, WANTED_WORDS_POSITIONS,
REMOVE_CHARS, REMOVE_WORDS, REMOVE_SINGLE_CHARS, OUTPUT_HEADER_SEPARATOR)
% function header = f_make_header(header, TERMS_SEPARATOR,
WANTED_TERMS_POSITIONS, WORDS_SEPARATOR, WANTED_WORDS_POSITIONS,
REMOVE_CHARS, REMOVE_WORDS, REMOVE_SINGLE_CHARS)

% examples:
% FROM SILVA HEADERS TO SPECIES NAMES
% SEPARATOR_TERMS = ';'; WANTED_TERMS_POSITIONS = 'last'; SEPARATOR_WORDS
= ' '; WANTED_WORDS_POSITIONS = 'all'; REMOVE_CHARS = '()';
% FROM SILVA HEADERS FOR rPREDICTOR
% SEPARATOR_TERMS = ' '; WANTED_TERMS_POSITIONS = [1]; SEPARATOR_WORDS =
''; WANTED_WORDS_POSITIONS = 'all'; REMOVE_CHARS = '';
% FROM BLAST SEARCH HEADERS TO SPECIES NAMES OLD STYLE
% SEPARATOR_TERMS = '|'; WANTED_TERMS_POSITIONS = [3]; SEPARATOR_WORDS =
' '; WANTED_WORDS_POSITIONS = [1 2]; REMOVE_CHARS = '';
```

```
% FROM BLAST SEARCH HEADERS TO SPECIES NAMES NEW STYLE
% SEPARATOR_TERMS = ''; WANTED_TERMS_POSITIONS = []; SEPARATOR_WORDS = '
'; WANTED_WORDS_POSITIONS = [1 2 3 4]; REMOVE_CHARS = '';

% FIRST FIND A SELECTED TERM(S)
if ~isempty(WANTED_TERMS_POSITIONS) &  ~isempty(TERMS_SEPARATOR)
      header = cut_string(header, TERMS_SEPARATOR);
      if ~isempty(WANTED_TERMS_POSITIONS)
            if strcmp(WANTED_TERMS_POSITIONS, 'last')
                  header = header{length(header)};
            else
                  ss = header{WANTED_TERMS_POSITIONS(1)};
                  for i = 2 : length(WANTED_TERMS_POSITIONS)
                        ss = [ss '_' header{WANTED_TERMS_POSITIONS(i)}];
                  end
                  header = ss;
            end
      end
end

for i = 1 : length(REMOVE_WORDS)
      j = strfind(header,REMOVE_WORDS{i});
      header(j : j + length(REMOVE_WORDS{i}) - 1) = [];
end

if REMOVE_SINGLE_CHARS
%     i = sort([find(isletter(header)) find(header == '0'|header ==
'1'|header == '2'|header == '3'|header == '4'|header == '5'|header ==
'6'|header == '7'|header == '8'|header == '9')]);
%     if ~isempty(i)
%           idel = [];
%           v = f_un_gr_ve(i);
%           for iv = 1 : length(v)
%                 if length(v{iv}) == 1
%                       idel = [idel v{iv}];
%                 end
%           end
%           header(idel) = [];
%     end
    i = isspace(header);
    i = find(i == 0);
    idel = [];
    v = f_un_gr_ve(i);
    for iv = 1 : length(v)
        if length(v{iv}) == 1
             idel = [idel v{iv} v{iv} + 1];
        end
    end
    header(idel) = [];
end

for i = 1 : length(REMOVE_CHARS)
      header(find(header == REMOVE_CHARS(i))) = '';
end

% REMOVE DOUBLE UNDERSCORES AFTER REMOVING SINGLE LETTERS
i = find(header == '_');
if ~isempty(i)
```

```
      idel = [];
      v = f_un_gr_ve(i);
      for iv = 1 : length(v)
            if length(v{iv}) > 1
                  idel = [idel v{iv}(2) : v{iv}(length(v{iv}))];
            end
      end
      header(idel) = [];
end

% REMOVE WHITE SPACES AT THE BEGINNING AND THE END
i = find(header == ' ');
if ~isempty(i)
      v = f_un_gr_ve(i);
      if i(length(i)) == length(header)
            header(v{length(v)}) = [];
      end
      if i(1) == 1
            header(v{1}) = [];
      end
end

% REMOVE DOUBLE SPACES
i = find(header == ' ');
if ~isempty(i)
      idel = [];
      v = f_un_gr_ve(i);
      for iv = 1 : length(v)
            if length(v{iv}) > 1
                  idel = [idel v{iv}(2) : v{iv}(length(v{iv}))];
            end
      end
      header(idel) = [];
end

% GET WORDS FROM THE SELECTED TERM(S)
if strcmp(WANTED_WORDS_POSITIONS, 'all')
      if ~isempty(WORDS_SEPARATOR)
            header(find(header == WORDS_SEPARATOR)) =
OUTPUT_HEADER_SEPARATOR;
      end
else
      header = cut_string(header, WORDS_SEPARATOR);
      ss = header{WANTED_WORDS_POSITIONS(1)};
      for i = 2 : length(WANTED_WORDS_POSITIONS)
            if i <= length(header)
                  ss = [ss OUTPUT_HEADER_SEPARATOR
header{WANTED_WORDS_POSITIONS(i)}];
            end
      end
      header = ss;
end


f_clean_RNA_sequence.m

function [sq i] = f_clean_RNA_sequence(sq)
% function [sq i] = clean_RNA_sequence(sq)
```

```
sq(find(~isletter(sq))) = [];
sq = upper(sq);

sq(find(sq ~= 'G' & sq ~= 'A' & sq ~= 'T' & sq ~= 'C' & sq ~= 'R' & sq ~=
'Y' & sq ~= 'M' & sq ~= 'K' & sq ~= 'S' & sq ~= 'W' & sq ~= 'H' & sq ~=
'B' & sq ~= 'V' & sq ~= 'D' & sq ~= 'N' & sq ~= 'U')) = 'N';

a = randperm(4);
n = 'ACGU';
sq(find(sq == 'N')) = n(a(1));

i = find(sq ~= 'A' & sq ~= 'C' & sq ~= 'G' & sq ~= 'T' & sq ~= 'U');
% if length(i) >
%     sq = 0;
%     return
% end
for ii = 1 : length(i)
      s=baselookup('code',sq(i(ii)));
      s=cut_string(s,char(9));
      s=cut_string(s{1},'|');
      sq(i(ii)) = s{1};
end
sq(find(sq == 'T')) = 'U';
```

f_read_genome.m

```
function [i f] = f_read_genome(fn)

fid=fopen(fn); f=char(fread(fid)'); fclose(fid);
i = find(f == 10);
%throw out the next line characters (the genomic sequence is formatted)
f(i)=[];
%throw out the 1st line which is the head
f(1:i(1)-1) = [];
[f i] = f_clean_RNA_sequence(f);
```

f_read_anns.m

```
function [crs ors anns ids] = f_read_anns(fn, lg, INCLUDERRNAS ,
INCLUDENCRNAS, INCLUDETRNAS, REMOVEPSEUDOGENES)
% function [crs ors anns] = f_read_anns(fn, lg, includernas = [1 0])

if ~exist('REMOVEPSEUDOGENES'), REMOVEPSEUDOGENES = 1; end
if ~exist('INCLUDERRNAS'), INCLUDERRNAS = 1; end
if ~exist('INCLUDENCRNAS'), INCLUDENCRNAS = 1; end
if ~exist('INCLUDETRNAS'), INCLUDETRNAS = 1; end
crs = []; ors = []; anns = {}; ids = {};

id=fopen(fn);
t=textscan(id,'%s %s %s %s %s %s %s %s %s %s %s %s %s %s %s %s %s %s %s
%s','delimiter','\t');
fclose(id);

%REMOVE THE FIRST LINE WITH THE HEAD
```

```
for i=1:length(t), t{i}(1)=[]; end

lines = strmatch('gene', t{1});

% COLUMNS WANTED: COORS 8 9, ORIENTATION 10, ANNS 14

for i=1:length(lines)
      if (i <= length(lines) && contains(t{1}{lines(i) + 1}, 'CDS')) ||
...
         (INCLUDERRNAS && contains(t{2}{lines(i)}, 'rRNA')) || ...
         (INCLUDENCRNAS && (contains(t{2}{lines(i)}, 'misc_RNA') ||
contains(t{2}{lines(i)}, 'ncRNA'))) || ...
         (INCLUDETRNAS && contains(t{2}{lines(i)}, 'tRNA'))
            crs = [crs; [str2num(t{8}{lines(i)})
str2num(t{9}{lines(i)})]];
            ors = [ors t{10}{lines(i)}];
            if i + 1 > length(lines)
                  anns = [anns; t{2}{lines(i)}];
            else
                  anns = [anns; t{14}(lines(i) + 1)];
            end
            ids = [ids; t{11}(lines(i))];
      else
            if ~REMOVEPSEUDOGENES && contains(t{2}{lines(i)},
'pseudogene')
                  crs = [crs; [str2num(t{8}{lines(i)})
str2num(t{9}{lines(i)})]];
                  ors = [ors t{10}{lines(i)}];
                  anns = [anns;  t{20}(lines(i))];
                  ids = [ids; t{17}(lines(i))];
            end
      end

end

% THIS IS 'COS YOU COMPUTE WITH THE IGRs AND NEED THE BEGINNING OF THE
1st IGR AND THE END OF THE LAST IGR.
if ~isempty(crs) & crs(1, 1) >= crs(1, 2)
    crs(1, 1) = 1;
end
% if crs(size(crs, 1), 2) < lg
      crs = [[0 0]; crs; [lg lg]];
      anns = [{''}; anns; {''}];
      ors = [' ' ors ' '];
      ids = [{''}; ids; {''}];
% else
%     crs = [[0 0]; crs;];
%     anns = [{''}; anns;];
%     ors = [' ' ors];
%     ids = [{''}; ids;];
% end
```

f_reconstruct_blast_hit.m

```
function d = f_reconstruct_blast_hit(d, l)
if  d.SubjectIndices(1) < d.SubjectIndices(2)
      d.SubjectIndices(1) = d.SubjectIndices(1) - d.QueryIndices(1) + 1;
```

```
      d.SubjectIndices(2) = d.SubjectIndices(2) + (1 -
d.QueryIndices(2));
else
      d.SubjectIndices(2) = d.SubjectIndices(2) - d.QueryIndices(1) + 1;
      d.SubjectIndices(1) = d.SubjectIndices(1)+ (1 - d.QueryIndices(2));
end
```

f_clean_fasta_headers.m

```
function fn = f_clean_fasta_headers(fn, RENAME, BAD_CHARS,
OUTPUTSEPARATOR, USEDFIRSTWORDS, WORDSEPARATOR)
% function fn_ = f_clean_fasta_headers(fn, RENAME, OUTPUTSEPARATOR [=
'_'], USEDFIRSTWORDS [=0], WORDSEPARATOR [= '_'])

if ~exist('RENAME'), RENAME = 1; end

% A COMPLETE SET OF BAD CHARS
% ':/\!''()";=,|+ -.';
if ~exist('BAD_CHARS'), BAD_CHARS = ':/\!''()";=,|+ '; end

if ~exist('OUTPUTSEPARATOR'), OUTPUTSEPARATOR = '_'; end
if ~exist('USEDFIRSTWORDS'), USEDFIRSTWORDS = 0; end % = use all words
if ~exist('WORDSEPARATOR'), WORDSEPARATOR = '_'; end

id = fopen(fn); sqs = textscan(id, '%s', 'delimiter', char(10));
fclose(id);
if RENAME
      fn=[fn '.cleaned'];
end
id = fopen(fn,'w');

for isqs = 1 : 2 : length(sqs{1})
% CLEAN HEADER
      j = fvi(sqs{1}{isqs}, BAD_CHARS);
      if ~isempty(j)
%           disp(['Bad chars (' sqs{1}{isqs}(j) ') in ''' sqs{1}{isqs} '''
header found and cleaned.'])
            sqs{1}{isqs}(j) = OUTPUTSEPARATOR;
      end
% THROW OUT WHITE SPACES AT THE END OF A HEADER
      j = find(sqs{1}{isqs} == ' ');
      k = find(sqs{1}{isqs} ~= ' ');
      i = find(j >= k(length(k)));
      sqs{1}{isqs}(j(i)) = [];
% THROW OUT UNDERSCORES AT THE END OF A HEADER
      j = find(sqs{1}{isqs} == '_');
      k = find(sqs{1}{isqs} ~= '_');
      i = find(j >= k(length(k)));
      sqs{1}{isqs}(j(i)) = [];
% THROW OUT UNINTERUPTED STRETCHES OF UNDERSCORES
      j = find(sqs{1}{isqs} == '_');
      i = f_un_gr_ve(j);
      idel = [];
      for ii = 1 : length(i)
            if length(i{ii}) > 1
                  idel = [idel i{ii}(2 : length(i{ii}))];
            end
```

```
      end
      sqs{1}{isqs}(idel) = [];
% THROW OUT UNINTERUPTED STRETCHES OF WHITE SPACES
      j = find(sqs{1}{isqs} == ' ');
      i = f_un_gr_ve(j);
      idel = [];
      for ii = 1 : length(i)
            if length(i{ii}) > 1
                  idel = [idel i{ii}(2 : length(i{ii}))];
            end
      end
      sqs{1}{isqs}(idel) = [];
      if USEDFIRSTWORDS
            s = cut_string(sqs{1}{isqs}, WORDSEPARATOR);
            sqs{1}{isqs} = s{1};
            if USEDFIRSTWORDS > length(s), USEDFIRSTWORDS = length(s); end
            for i = 2 : USEDFIRSTWORDS
                  sqs{1}{isqs} = [sqs{1}{isqs} WORDSEPARATOR s{i}];
            end
      end
      fwrite(id, [sqs{1}{isqs} char(10) sqs{1}{isqs + 1} char(10)]);
end
fclose(id);
```

f_filter_fasta_by_ID.m

```
function [fn idel] = f_filter_fasta_by_ID(fn, RENAME, NOTINHEADERS)

if ~exist('NOTINHEADERS') | ~ischar(NOTINHEADERS), NOTINHEADERS = {'MAG
'; 'MAG: '; 'TPA_asm ' ; 'TPA_asm: '; 'candidatus '; 'sp. '; 'isolate ';
' bacterium '; 'UNVERIFIED_ORG: ';}; end

if f_file_size(fn)
    t = fastaread(fn);
    if RENAME
        fn=[fn '.fltd'];
    end

    idel = []; id = fopen(fn, 'w'); indxs = 1 : length(t);

    while ~isempty(t)
        s = strsplit(t(1).Header, ' ');
    % this is because some IDs have the NZ_ prefix. you have to look for
ID without prefix in IDs both with and w/o the prefix.
        i = [];
        for it = 1 : length(t)
            ss = strsplit(t(it).Header, ' ');
            if ~isempty(findstr(ss{1}, s{1}(2 : length(s{1}))))
                i = [i it];
            end
        end

    % use ID with longest sequence
        l = [];
        for ii = 1 : length(i)
            l = [l length(t(i(ii)).Sequence)];
        end
```

```
        j = find(l == max(l));

    % if more IDs with the same longest sequence, then use that that has
not got any of the NOTINHEADERS in its header
        if length(j) > 1
            jdel = [];
            for ij = 1 : length(j)
                for iNOTINHEADERS = 1 : length(NOTINHEADERS)
                    if ~isempty(strfind(t(i(ij)).Header,
NOTINHEADERS{iNOTINHEADERS}))
                        jdel = [jdel ij];
                    end
                end
            end
            j(jdel) = [];
            if isempty(j), [j j] = max(l); else, j = j(1); end
        end

        fwrite(id, ['>' t(i(j)).Header char(10) t(i(j)).Sequence
char(10)]);
        t(i) = [];

        x = i(j);
        i(j) = [];
        for ii = 1 : length(i)
            idel = [idel indxs(i(ii)) * 2 - 1 indxs(i(ii)) * 2];
        end
        indxs([i x]) = [];

    end
    fclose(id);
else
    idel = [];
    if RENAME
        eval(['!cp ' fn ' ' fn '.fltd']);
        fn=[fn '.fltd'];
    end
end
```

f_try_anns

```
function [ngenomic nfeature] = f_try_anns(ID, ANNSPATH, TRYTOLOAD,
VERBOSE)

if ~exist('VERBOSE'), VERBOSE = 0; end
if ANNSPATH(1) == '~'
      [x x]=unix('echo $HOME');
      ANNSPATH = [x(1 : length(x) - 1) ANNSPATH(2 : length(ANNSPATH))];
end
ngenomic = []; nfeature = [];

% FIND OUT HAVE THIS GENOMIC SEQUENCE AND ITS ANNOTATIONS IN ANNSPATH.
YOU NEED TO HAVE GENOMIC SEQUENCES AS THERE IS NO ID IN ANNOTATIONS.
if ~exist(ANNSPATH, 'dir')
      n = [];
      mkdir(ANNSPATH);
else
```

```
      [n n] = unix(['grep -liR ' ID ' ' ANNSPATH '/.']);
end
if TRYTOLOAD & (isempty(n) | sum(n == char(10)) < 2)
      if VERBOSE, disp(['Can''t find annotations in ' ANNSPATH '/; trying
to download...']), end
% GET ASSEMBLY PATH FOR BLAST ID
      [assemblypath assemblystatus] =
f_get_assembly_ID_from_ID_edirect(ID);
      if ~isempty(assemblypath)
            i = [strfind(assemblypath, '/GCA_') strfind(assemblypath,
'/GCF_')];

% FIND IF THERE ALREADY ARE OTHER CHROMOSOMES OF THIS ASSEMBLY NAME THAT
HAVE DIFFERENT IDS. YOU SHOULD NOT DELETE THEM.
            [y y] = unix(['ls ' ANNSPATH '/' assemblypath(i + 1 :
length(assemblypath)) '*']);
            y = strsplit(y, char(10));
            y(length(y)) = [];

            [chr_g_fns chr_a_fns plasmids_g_fns plasmids_a_fns] =
f_store_anns_simple_no_ONLYFEATURETABLE(assemblypath, assemblypath(i + 1
: length(assemblypath)), ANNSPATH);
% FIND WHICH CHROMOSOME HAVE YOUR ID FOR NOT TO DELETE IT
            [n n] = unix(['grep -l '  ID ' ' ANNSPATH '/' assemblypath(i +
1 : length(assemblypath)) '*']);
            if ~isempty(n) & sum(n == char(10)) > 1 & isempty(strfind(n,
'No such file or directory'))
                  n = strsplit(n, char(10));
                  n(length(n)) = [];
                  if strfind(n{1}, 'genomic.fna'), ngenomic = n{1};
nfeature = n{2}; else, ngenomic = n{2}; nfeature = n{1}; end
% IF IT IS CONTIGS, TRY TO IDENTIFY THE RIGHT ONE
                  if length(chr_g_fns) > 1
% FIND THE ASSEMBLY IN WHICH THE BLAST HIT IS FOR NOT TO DELETE IT.
                        chr_g_fns(strmatch(ngenomic, chr_g_fns, 'exact'))
= [];
                        chr_a_fns(strmatch(nfeature, chr_a_fns, 'exact'))
= [];
                        if ~isempty(y) & isempty(strfind(y{1}, 'No such
file or directory'))
% YOU DON'T WANT TO DELETE CHROMOSOMES WITH THE SAME ASSEMBLY NAME THAT
YOU HAVE ALREADY ON THE DISK.
                              for iy = 1 : length(y)
% this is because different grep output in wide and narrow matlab command
window.
                                    if sum(y{iy}==' ')
                                          m = strsplit(y{iy}, ' ');
                                          for im = 1 : length(m)
                                                chr_g_fns(strmatch(m{im},
chr_g_fns, 'exact')) = [];
                                          end
                                    else
                                          chr_g_fns(strmatch(y{iy},
chr_g_fns, 'exact')) = [];
                                    end
                              end
                              for iy = 1 : length(y)
```

```
% this is because different grep output in wide and narrow matlab command window.
                              if sum(y{iy}==' ')
                                   m = strsplit(y{iy}, ' ');
                                   for im = 1 : length(m)
                                        chr_a_fns(strmatch(m{im}, chr_a_fns, 'exact')) = [];
                                   end
                              else
                                   chr_a_fns(strmatch(y{iy}, chr_a_fns, 'exact')) = [];
                              end
                         end
                    end
% DELETE ALL OTHER CHROMOSOMES WITH THE SAME ASSEMBLY NAME.
                    for i = 1 : length(chr_a_fns)
                         eval(['!rm ' chr_a_fns{i}]);
                    end
                    for i = 1 : length(chr_g_fns)
                         eval(['!rm ' chr_g_fns{i}]);
                    end
               end
               if length(plasmids_g_fns) > 1
% FIND THE ASSEMBLY IN WHICH THE BLAST HIT IS
                    plasmids_g_fns(strmatch(ngenomic, plasmids_g_fns, 'exact')) = [];
                    plasmids_a_fns(strmatch(nfeature, plasmids_a_fns, 'exact')) = [];
                    for i = 1 : length(plasmids_a_fns)
                         eval(['!rm ' plasmids_a_fns{i} ' ' plasmids_g_fns{i}]);
                    end
               end
          else
               nfeature = 'Can''t download annotations ''cos wget. ';
          end
     else
          nfeature = ['Can''t download annotations ''cos edirect (''' assemblystatus '''). '];
     end
end

if ~iscell(n) & (isempty(n) | sum(n == char(10)) < 2 | ~isempty(strfind(n, 'No such file or directory')))
     nfeature = [nfeature 'No annotations in ' ANNSPATH '/.'];
else
     if ~iscell(n)
          n = strsplit(n, char(10));
          if strfind(n{1}, 'genomic.fna')
               ngenomic = n{1}; nfeature = n{2};
          else
               ngenomic = n{2}; nfeature = n{1};
          end
     end
end
```

f_get_assembly_ID_from_ID_edirect.m

```
function [assemblypath assemblystatus]=
f_get_assembly_ID_from_ID_edirect(ID)

assemblypath = ''; assemblystatus = 'Nothing.';

% | xtract -pattern DocumentSummary -element FtpPath_RefSeq

[t t] = unix(['esearch -db nuccore -query "' ID '" | elink -db assembly -
target assembly | esummary']);

if sum(ID == '_')
      assemblypath = t(strfind(t, '<FtpPath_RefSeq>') +
length('<FtpPath_RefSeq>') : strfind(t, '</FtpPath_RefSeq>') - 1);
% if ~isempty(assemblypathrs) & strfind(assemblypathrs,
'ftp://ftp.ncbi.nlm.nih.gov/genomes/all/GCF/') == 1
else
      assemblypath = t(strfind(t, '<FtpPath_GenBank>') +
length('<FtpPath_GenBank>') : strfind(t, '</FtpPath_GenBank>') - 1);
% if ~isempty(assemblypathgb) & strfind(assemblypathgb,
'ftp://ftp.ncbi.nlm.nih.gov/genomes/all/GCA/') == 1
end

assemblystatus = t(strfind(t, '<AssemblyStatus>') +
length('<AssemblyStatus>') : strfind(t, '</AssemblyStatus>') - 1);
```

f_store_anns_simple_no_ONLYFEATURETABLE.m

```
function [chr_g_fns chr_a_fns plasmids_g_fns plasmids_a_fns] =
f_store_anns_simple_no_ONLYFEATURETABLE(ftppath, assemblyname, ANNSPATH)

chr_g_fns = {}; chr_a_fns={}; plasmids_g_fns = {}; plasmids_a_fns = {};

[a a]=unix(['wget -P ' ANNSPATH ' --timeout 30 -t 2 ' ftppath '/'
assemblyname '_feature_table.txt.gz']);
if ~isempty(strfind(a, 'saved'))
      [g g]=unix(['wget -P ' ANNSPATH ' --timeout 30 -t 2 ' ftppath '/'
assemblyname '_genomic.fna.gz']);
      if ~isempty(strfind(g, 'saved'))
            eval(['!gunzip -f ' ANNSPATH '/' assemblyname
'_feature_table.txt.gz']);
            eval(['!gunzip -f ' ANNSPATH '/' assemblyname
'_genomic.fna.gz']);
      else
            chr_a_fns = [assemblyname '_genomic.fna.gz doesn''t exist in
ftp path.'];
            return
      end
else
      chr_a_fns = [assemblyname '_feature_table.txt.gz doesn''t exist in
ftp path.'];
      return
end

% NOW ONLY COMPLETE GEMONMES, CHROMOSOMES AND PLASMIDS CAN BE IN FNA
FILES. CONTIGS AND SCAFFOLDS WERE FILTERED OUT IN
```

```
f_load_genomic_data_both.m

[chr_g_fns chr_a_fns plasmids_g_fns plasmids_a_fns] =
f_parse_genomic_data([ANNSPATH '/' assemblyname]);

if exist([ANNSPATH '/' assemblyname '_feature_table.txt.gz'], 'file')
      eval(['!rm ' ANNSPATH '/' assemblyname '_feature_table.txt.gz*']);
end
if exist([ANNSPATH '/' assemblyname '_genomic.fna.gz'], 'file')
      eval(['!rm ' ANNSPATH '/' assemblyname '_genomic.fna.gz*']);
end
% f_parse_genomic_data_gbID CHANGES FILE NAMES BY THE ADDITION OF
CHROMOSOME AND PLASMID KEYWORDS, SO THE ORIGINAL FILES GOT BY GUNZIPPING
SHOULD BE DELETED AS THEY ARE NOT BE USED
if exist([ANNSPATH '/' assemblyname '_genomic.fna'], 'file')
      eval(['!rm ' ANNSPATH '/' assemblyname '_genomic.fna']);
end
if exist([ANNSPATH '/' assemblyname '_feature_table.txt'], 'file')
      eval(['!rm ' ANNSPATH '/' assemblyname '_feature_table.txt']);
end
```

Conservation filtering

Scripts

syntenic_homs_of_cands.m:
-> f_read_local_BLAST_output_to_fasta_edirect_comb.m

Matlab Code

```
syntenic_homs_of_cands.m

% % example of how to make input files to this script:
% % cat glac_13_cands.fa_syntenic.broader
glac_14_cands.fa_syntenic.broader glac_9_cands.fa_syntenic.broader >
glac_cands.fa_syntenic.broader
% % cat glac_13_syntenic_cands.fa glac_14_syntenic_cands.fa
glac_9_syntenic_cands.fa > glac_syntenic_cands.fa

% LEGEND:
% IF pat IS A CELL, THEN ANY STRING OF IT MUST BE FOUND IN ANY GENOMIC
CONTEXT SIDE.
% IF pat IS TWO CELLS, THEN ANY STRING OF ONE OF THE CELL MUST BE FOUND
IN ONE OF THE TWO GENOMIC CONTEXT SIDES, AND THEN ANY STRING OF THE OTHER
CELL
% IN THE OTHER GENOMIC CONTEXT SIDE.
% THERE ALSO WORDS AND/OR SUBWORDS IN THE STRINGS CAN BE SEARCHED FOR
DEPENDING ON requireAll AND allowSubstring PARAMS.

annsd = '/mnt/spin/anns_Bacteria_sa';

% genus='King';
% DB = 'ref_prok_rep_gen';
% DB = 'nt_prok';
```

```
% DB = 'ref_seq_gen';
% DB = 'core_nt';
%
% % genus='Neis';
% % DB = 'ref_gen';
% % DB = 'nt_prok';
% % DB = 'core_nt';

% genus='pseudoalterom';
% DB = 'refseq_genomes';
% DB = 'refseq_genomes_in_Alteromonadales_no_Pseudoalteromonas';
% DB =
'refseq_genomes__Flocculibacter_collagenilyticus_in_Alteromonadales';

% genus='glac';
% DB = 'refseq_genomes';
% no syntenic homologs found: DB = 'nt_prok';

% % pat={'ribosome biogenesis GTP'; 'ribosome GTPase'; 'YihA';
'YsxC';'engb';'DNA polymerase I';'pola';'ElbB'};
% pat={{'ribosome biogenesis GTP'; 'ribosome GTPase'; 'YihA';
'YsxC';'engb';};{'DNA polymerase I';'pola';'YceI';};};

genus='89_frank_.05_13.txt';
% genus= '89_frank_.05_9.txt';
DB = 'refseq_genomes_in_Frankiales';
%  pat={{'HAD hydrolase'; 'haloacid dehalogenase'; 'phosphoserine
phosphatase';'morphological differentiation';};...
%      {'oxidoreductase';'beta hexosaminidase';'Beta glucosidase';
'glycosyl hydrolase';'glycoside hydrolase';};};
% % requireAll = 1; allowSubstring = 1; cntx = 4; ->
89_frank_.05_13_9.txt_cands.fa_syntenic_homologs_refseq_genomes_in_Franki
ales
requireAll = 1; allowSubstring = 1; cntx = 8; pat={{'hydrolase';
'dehalogenase'; 'phosphatase';'morphological differentiation';};...
{'oxidoreductase';'oxidase';'dehydrogenase';'reductase';'hexosaminidase';
'glucosidase'; 'glycosidase';'hydrolase';'phosphatase';};};
% requireAll = 1; allowSubstring = 0; cntx = 8;
% % pat={'hydrolase'; 'dehalogenase';
'phosphatase';'hexosaminidase';'glucosidase';'glycosidase';'morphological
differentiation';};
% pat={'HAD hydrolase'; 'haloacid dehalogenase'; 'phosphoserine
phosphatase';'morphological differentiation';'beta hexosaminidase';'Beta
glucosidase';...
%      'glycosyl hydrolase';'glycoside
hydrolase';'acylphosphatase';'pyrophosphatase';'trehalohydrolase';'amidoh
ydrolase';};

% SEPARATOR = ' ';SPECIESPOS=3; GENUSPOS=2;
% annsd = '/mnt/spin/anns_Bacteria_sa';
% ido = fopen([genus '_cands.fa_syntenic_homologs_' DB], 'w');
% fa = fastaread([genus '_cands.fa_syntenic']);
% bosfnsuffix = [genus '_cands.fa_syntenic_'];

% fafn = 'Supplementary File S7 - frank_manually_chosen_cand_strs.br.fa';
% SEPARATOR = '_'; SPECIESPOS=6; GENUSPOS=5;
% DB = 'refseq';
% requireAll = 1; allowSubstring = 1; cntx = 7;
```

```
% pat={{'conjugal';'SCO6880';'TrbL';'DUF8175';'DUF6112';'MinD';};
{'helix-turn-helix domain';'NlpC';'FAD oxidoreductase'}};
% ido = fopen([fafn '_syntenic_homologs_' DB], 'w');
% bo_fns_suffix = 'Supplementary File S7 -
frank_manually_chosen_cand_strs.br_';

fafn = 'Supplementary File S8 -
bifi_alt_manually_chosen_representative_sopts.br.fa';
SEPARATOR = '_'; SPECIESPOS=6; GENUSPOS=5;
% DB = 'ntprok';
DB = 'refseq';
requireAll = 1; allowSubstring = 1; cntx = 7; pat={'CpaE';'P-loop
NTPase';'AAA family ATPase';'CpaF';'TadA';};
ido = fopen([fafn '_syntenic_homologs_' DB], 'w');
bo_fns_suffix = 'Supplementary File S8 -
bifi_alt_manually_chosen_representative_sopts.br_';


% % --


NOTINNAMES =
["MAG","DNA","chromosome",",","complete","genome","MAG:","TPA_asm","TPA_a
sm:","candidatus","cf.","Candidatus",...
                "subsp.", "sp.", "strain","isolate", "_bacterium_", "
bacterium ", " bacterium,",
"UNVERIFIED_ORG:","Sample","type","genome","assembly",":"," I ",...
                   "whole", "shotgun", "sequence", "uncultured",
"Uncultured"];
fa = fastaread(fafn);
for ifa = 1 : length(fa)
      disp([char(10) 'Doing ' num2str(ifa) '/' num2str(length(fa)) ' '
fa(ifa).Header '.'])
      if exist([bo_fns_suffix num2str(ifa) '.bo'])
            [fn_ names_ evals_ gls_ anns_ annsext_ glsn_ wenthroughHSPs
usedHSPs noannsHSPs notinigrHSPs annsleft annsright annsextleft ...
             annsextright anns1extleft anns1extright posinquery_
anns2extleft anns2extright anns3extleft anns3extright anns4extleft ...
             anns4extright anns5extleft anns5extright anns6extleft
anns6extright] = ...
            f_read_local_BLAST_output_to_fasta_edirect_comb([bo_fns_suffix
num2str(ifa) '.bo'],annsd,0,.05, 1,100,0, '','',1,0,1,1,' ',1,0,0,0,2);
% the candidate BLAST hit does not always need to be the first one in the
BLAST output. Find the name of syntenic candidate in names of candidate
BLAST .bo file
            fa(ifa).Header = strtrim(regexprep(erase(fa(ifa).Header,
NOTINNAMES), '\s+', ' '));
            f = strsplit(fa(ifa).Header, SEPARATOR);
            candname = [f{GENUSPOS} ' ' f{SPECIESPOS}];
            j = find(contains(names_, f{SPECIESPOS}, 'IgnoreCase', true) &
contains(names_, f{GENUSPOS}, 'IgnoreCase', true));
%           if isempty(j)
% % the hit of a candidate can have different species name but is 100% no
gaps...???
%                 d_ = blastreadlocal([bo_fns_suffix num2str(ifa) '.bo'],
0);
```

```
%                 if (d_.Hits(1).HSPs(1).Identities.Match ==
d_.Hits(1).HSPs(1).Identities.Possible) &&
d_.Hits(1).HSPs(1).Identities.Percent == 100
%                       j = 1;
%                 end
%           end
%           if isempty(j)
% %               disp([fa(ifa).Header ' not found in its BLAST output '
genus '_syntenic_cands.fa_' num2str(ifa) '.bo_' DB '. Ignoring it.'])
% % % if you limit BLAST search to the taxa in which the query species
does not belong, then there is no query in BLAST output.
%                 candname = 'Noquery';
%                 sbroader = '';
%           else
%                 candname = strtrim(regexprep(erase(names_{j(1)},
NOTINNAMES), '\s+', ' '));
%                 candname = candname(find(candname == ' ') + 1 :
length(candname));
                  if length(candname ) > 30, candname = candname(1:30);
end
                  if candname(length(candname)) == ' ',
candname(length(candname)) = []; end
                  disp(['Identifying syntenic homologs of ' candname '
candidate.'])

                  if ~isempty(j)
                  switch cntx
                        case 8
                  sbroader = ['Cand. ' candname ' : ' ...
                                    anns6extleft{j(1)} ' | '
anns5extleft{j(1)} ' | ' anns4extleft{j(1)} ' | ' anns3extleft{j(1)} ' |
' anns2extleft{j(1)} ' | ' ...
                                    anns1extleft{j(1)} ' | '
annsextleft{j(1)} ' | ' annsleft{j(1)} ' # ' annsright{j(1)} ' | '
annsextright{j(1)} ' | ' ...
                                    anns1extright{j(1)} ' | '
anns2extright{j(1)} ' | ' anns3extright{j(1)} ' | ' anns4extright{j(1)} '
| ' anns5extright{j(1)} ' | ' ...
                                    anns6extright{j(1)} char(10)];
                        case 7
                  sbroader = ['Cand. ' candname ' : ' ...
                                    anns5extleft{j(1)} ' | '
anns4extleft{j(1)} ' | ' anns3extleft{j(1)} ' | ' anns2extleft{j(1)} ' |
' ...
                                    anns1extleft{j(1)} ' | '
annsextleft{j(1)} ' | ' annsleft{j(1)} ' # ' annsright{j(1)} ' | '
annsextright{j(1)} ' | ' ...
                                    anns1extright{j(1)} ' | '
anns2extright{j(1)} ' | ' anns3extright{j(1)} ' | ' anns4extright{j(1)} '
| ' anns5extright{j(1)} char(10)];
                        case 4
                  sbroader = ['Cand. ' candname ' : ' ...
                                    anns2extleft{j(1)} ' | '
anns1extleft{j(1)} ' | ' annsextleft{j(1)} ' | ' annsleft{j(1)} ' # '
annsright{j(1)} ' | ' ...
                                    annsextright{j(1)} ' | '
anns1extright{j(1)} ' | ' anns2extright{j(1)} char(10)];
                        case 1
```

```
                            sbroader = ['Cand. ' candname ' : '
annsleft{j(1)} ' # ' annsright{j(1)} char(10)];
                 end
                 else
                       sbroader = ['Cand. ' candname ' : the candidate
was not in .bo as most likely the taxa of the query species was excluded
from BLAST search to avoid lots of (almost) identical hits.'];
                 end
%           end
% find out homologs with one of the putative spot42-type genomic contexts
                 ss = ['Cand. ' candname ' -> '];
                 s = [];
                 for inames = 1 : length(names_)
                       if isempty(j) || inames ~= j(1)
                            namehom =
strtrim(regexprep(erase(names_{inames}, NOTINNAMES), '\s+', ' '));
                            namehom = namehom(find(namehom == ' ') + 1 :
length(namehom));
                            namehom = strsplit(namehom, ' ');
                            if length(namehom) > 2
                                  namehom3 = strsplit(namehom{3}, '_');
                                  namehom = [namehom{1} ' ' namehom{2} '
' namehom3{1}];
                            else
                                  namehom = [namehom{1} ' ' namehom{2}];
                            end
                            if length(namehom) > 30, namehom =
namehom(1:30); end
                            switch cntx
                                  case 8
                            syntenytable = [anns6extleft{inames} ' | '
anns5extleft{inames} ' | ' anns4extleft{inames} ' | '
anns3extleft{inames} ' | ' ...
                                                  anns2extleft{inames} '
| ' ...
                                                  anns1extleft{inames} '
| ' annsextleft{inames} ' | ' annsleft{inames} ' # ' annsright{inames} '
| ' ...
                                                  annsextright{inames} '
| ' anns1extright{inames} ' | ' anns2extright{inames} ' | '
anns3extright{inames} ' | ' ...
                                                  anns4extright{inames}
' | ' anns5extright{inames} ' | ' anns6extright{inames}];
                                  case 7
                            syntenytable = [anns5extleft{inames} ' | '
anns4extleft{inames} ' | ' anns3extleft{inames} ' | ' ...
                                                  anns2extleft{inames} '
| ' ...
                                                  anns1extleft{inames} '
| ' annsextleft{inames} ' | ' annsleft{inames} ' # ' annsright{inames} '
| ' ...
                                                  annsextright{inames} '
| ' anns1extright{inames} ' | ' anns2extright{inames} ' | '
anns3extright{inames} ' | ' ...
                                                  anns4extright{inames}
' | ' anns5extright{inames}];
                                  case 4
```

```
                              syntenytable = [anns2extleft{inames} ' | '
anns1extleft{inames} ' | ' annsextleft{inames} ' | ' annsleft{inames} ' #
' ...
                                              annsright{inames} ' |
' annsextright{inames} ' | ' anns1extright{inames} ' | '
anns2extright{inames}];
                                    case 1
                              syntenytable = [annsleft{j(1)} ' # '
annsright{j(1)}];
                              end
                              if iscell(pat{1})
                                    x = strsplit(syntenytable, ' # ');
                                    pat1i=0;
                                    pat2i=0;
                                    for ipat = 1 : length(pat{1})
                                          if findCommonWords(x{1},
pat{1}{ipat}, requireAll, allowSubstring)
                                                pat1i=1;
                                                break
                                          end
                                    end
                                    if ~pat1i
                                          for ipat = 1 : length(pat{1})
                                                if findCommonWords(x{2},
pat{1}{ipat}, requireAll, allowSubstring)
                                                      pat1i=2;
                                                      break
                                                end
                                          end
                                    end
                                    for ipat = 1 : length(pat{2})
                                          if findCommonWords(x{1},
pat{2}{ipat}, requireAll, allowSubstring)
                                                pat2i=1;
                                                break
                                          end
                                    end
                                    if ~pat2i
                                          for ipat = 1 : length(pat{2})
                                                if findCommonWords(x{2},
pat{2}{ipat}, requireAll, allowSubstring)
                                                      pat2i=2;
                                                      break
                                                end
                                          end
                                    end
% any string of one sets in any of sides, any string of the other set in
the other side
                                    if (pat1i==1 & pat2i==2) || (pat1i==2 &
pat2i==1)
                                          s = [s namehom ', '];
                                          sbroader = [sbroader namehom ' :
' syntenytable char(10)];
                                    end
                              else
% any string of the set in any side with words in any order
                                    for ipat = 1 : length(pat)
```

```
                                        if findCommonWords(syntenytable, 
pat{ipat}, requireAll, allowSubstring)
                                              s = [s namehom ', '];
                                              sbroader = [sbroader namehom 
' : ' syntenytable char(10)];
                                              break
                                        end
                                  end
                            end
                      end
                end
                if ~isempty(s)
% this is a list of syntenic candidates with their syntenic homologs
                      fwrite(ido, [ss s(1:length(s)-2) '.' char(10)]);
% these are genomic contexts of syntenic homologs for each of syntenic 
candidates
                      ido1name = [candname '.homologs.broader_' DB];
                      ido1name(find(ido1name == '/')) = '_';
                      ido1 = fopen(ido1name,'w'); fwrite(ido1, 
sbroader); fclose(ido1);
                end
                disp(['Found ' num2str(sum(s == ',')) ' syntenic 
homolog(s).'])
%          end
      else
%          disp([genus '_cands.fa_syntenic_' num2str(ifa) '.bo_' DB ' not 
exist - most likely, no BLAST hits for this candidate.'])
           disp([bo_fns_suffix num2str(ifa) '.bo not exist - most likely, 
no BLAST hits for this candidate.'])

      end
end
fclose(ido);
% Genus=genus;Genus(1) = upper(genus(1));
% fa=dir([Genus '*homologs.*' DB '.broader']);for 
i=1:length(fa),sort_broader_context(fa(i).name);end

clear cntxfn pars ido1 ido sbroader namehom name isqs annsd ido BLAST 
inames x name fn_ ido1name synteniccandscntxs
clear names_ evals_ gls_ anns_ annsext_ glsn_ wenthroughHSPs usedHSPs 
noannsHSPs notinigrHSPs annsleft annsright annsextleft annsextright 
anns1extleft
clear anns1extright posinquery_ anns2extleft anns2extright
clear db xl2 xl3 j k1 k2 sqs v vv namehom f s ss i type itype flag ctype 
candname namehom3 NOTINNAMES DB
clear anns3extleft anns3extright anns4extleft anns4extright anns5extleft 
anns5extright anns6extleft anns6extright
clear bos           fas           ifa           ipat          pat1i         
syntenytable
clear ans           fa            genus         ifas          pat           
pat2i         v  d_ requireAll allowSubstring cntx bo_fns_suffix fafn


f_read_local_BLAST_output_to_fasta_edirect_comb.m
```

```
function [fn_ names_ evals_ gls_ anns_ annsext_ glsn_ wenthroughHSPs
usedHSPs noannsHSPs notinigrHSPs annsleft annsright annsextleft
annsextright anns1extleft ...
             anns1extright posinquery_ anns2extleft anns2extright
anns3extleft anns3extright anns4extleft anns4extright anns5extleft
anns5extright...
             anns6extleft anns6extright leftannscoor_3prime
rightannscoor_5prime fnrec]=
f_read_local_BLAST_output_to_fasta_edirect_comb(fn_, ANNS_SA, EVAL_BEST,
...
             EVAL_WORST, FROM, TO, ALM_MIN_LEN_RAT, NAME, NOTNAME,
INIGRONLY, CLEANHEADERS, VERBOSE, TRYTOLOAD, OUTPUTHEADERSEPARATOR,
DO_ONLY_FIRST_HSP, ...
             FILTER, BASIC_BO_READ, WRITE_HSP_SQS, OVERLAP,
RECONSTRUCTHITS)

if ~exist('FROM'), FROM = 1; end
if ~exist('TO'), TO = realmax; end
if ~exist('ALM_MIN_LEN_RAT'), ALM_MIN_LEN_RAT = 0; end
if ~exist('NAME'), NAME = ''; end
if ~exist('NOTNAME'), NOTNAME = ''; end
if ~exist('INIGRONLY'), INIGRONLY = 1; end
if ~exist('CLEANHEADERS'), CLEANHEADERS = 0; end
if ~exist('VERBOSE'), VERBOSE = 1; end
if ~exist('TRYTOLOAD'), TRYTOLOAD = 1; end
if ~exist('OUTPUTHEADERSEPARATOR'), OUTPUTHEADERSEPARATOR = ' '; end
if ~exist('DO_ONLY_FIRST_HSP'), DO_ONLY_FIRST_HSP = 1; end
if ~exist('FILTER'), FILTER = 0; end
if ~exist('BASIC_BO_READ'), BASIC_BO_READ = 0; end
if ~exist('WRITE_HSP_SQS'), WRITE_HSP_SQS = 1; end
if ~exist('OVERLAP'), OVERLAP = 0; end
if ~exist('RECONSTRUCTHITS'), RECONSTRUCTHITS = 0; fnrec=''; end

% if strcmp(ANNS_SA, ANNS), disp('ANNS_SA == ANNS. Stop.'); return, end
if ~exist('REMOVEPSEUDOGENES'), REMOVEPSEUDOGENES = 0; end
if ~exist('INCLUDERRNAS'), INCLUDERRNAS = 0; end
if ~exist('INCLUDENCRNAS'), INCLUDENCRNAS = 0; end
if ~exist('INCLUDETRNAS'), INCLUDETRNAS = 1; end

if ~isempty(ANNS_SA) & ~BASIC_BO_READ
      if ANNS_SA(length(ANNS_SA)) == '/', ANNS_SA(length(ANNS_SA)) = [];
end
%     if ANNS(length(ANNS)) == '/', ANNS(length(ANNS)) = []; end
%     f_remove_incomplete_anns(ANNS,0);
%     f_remove_incomplete_anns(ANNS_SA,0);
end

NOTINNAMES =
["MAG","DNA","chromosome",",","complete","genome","MAG:","TPA_asm","TPA_a
sm:","candidatus","cf.","Candidatus",...
                "subsp.", "sp.", "strain","isolate", " bacterium ",
"bacterium,", "UNVERIFIED_ORG:","Sample","type","genome","assembly",":","
I ",...
                   "whole", "shotgun", "sequence", "uncultured",
"Uncultured"];

wenthroughHSPs = 0; usedHSPs = 0; noannsHSPs = 0; notinigrHSPs = 0;
```

```
d_ = blastreadlocal(fn_, 0);

no = 0;
for di = 1 : length(d_.Hits)
      for ihit = 1 : length(d_.Hits(di).HSPs)
            if d_.Hits(di).HSPs(ihit).Expect <= EVAL_WORST
                  no = no + 1;
                  break
            end
      end
end
disp(['Gonna do ' num2str(no) ' hits (out of total '
num2str(length(d_.Hits)) ') with at least 1 HSP under EVAL_WORST = '
num2str(EVAL_WORST) '.'])

if RECONSTRUCTHITS
      fnrec = [fn_ '.fa_rec'];
      idrec = fopen(fnrec,'w');
end
if WRITE_HSP_SQS
      fn_ = [fn_ '.fa'];
      id = fopen(fn_,'w');
end
% SOMETIMES SYNTENY AND SYNTENYEXT CAN CONTAIN EMPTY STRINGS AS 1ST AND
LAST ANNOTATION STRINGS ARE EMPTY COS THE INDEXING OF IGRS (SEE
F_READ_ANNS.M)
anns_ = {}; annsext_ = {}; gls_ = []; names_ = {}; glsn_ = []; evals_ =
[]; annsleft = {}; annsright = {}; annsextleft = {}; annsextright = {};
anns1extleft = {};
anns1extright = {}; posinquery_ = []; anns2extleft = {}; anns2extright =
{}; anns3extleft = {}; anns3extright = {}; anns4extleft = {};
anns4extright = {};
anns5extleft = {}; anns5extright = {}; anns6extleft = {}; anns6extright =
{}; leftannscoor_3prime = []; rightannscoor_5prime = [];

if TO > length(d_.Hits), TO = length(d_.Hits); end
for di = FROM : TO
%     if (isempty(NAME) & isempty(NOTNAME)) | ...
%        (~isempty(NAME) & isempty(NOTNAME) &
~isempty(strfind(d_.Hits(di).Name, NAME))) | ...
%        (isempty(NAME) & ~isempty(NOTNAME) &
isempty(strfind(d_.Hits(di).Name, NOTNAME))) | ...
%        (~isempty(NOTNAME) & ~isempty(NAME) &
~isempty(strfind(d_.Hits(di).Name, NAME)) &
isempty(strfind(d_.Hits(di).Name, NOTNAME)))
      z = f_find_all_words_of_str_in_str(d_.Hits(di).Name, NOTNAME, ' ');
      if (isempty(NAME) & isempty(NOTNAME)) | (~isempty(NAME) &
~isempty(strfind(d_.Hits(di).Name, NAME))) | (~isempty(NOTNAME) & z <=
length(find(NOTNAME == ' ')))
%           disp(['Doing hit #' num2str(di) '/' num2str(no) '/'
num2str(length(d_.Hits)) ', ' d_.Hits(di).Name '.'])
            if DO_ONLY_FIRST_HSP, hspsno = 1; else, hspsno =
length(d_.Hits(di).HSPs); end
            for ihit = 1 : hspsno
                  wenthroughHSPs = wenthroughHSPs + 1;
                  if d_.Hits(di).HSPs(ihit).Expect >= EVAL_BEST & ...
                        d_.Hits(di).HSPs(ihit).Expect < EVAL_WORST & ...
```

```
                        abs(d_.Hits(di).HSPs(ihit).QueryIndices(1) -
d_.Hits(di).HSPs(ihit).QueryIndices(2)) >= d_.Length * ALM_MIN_LEN_RAT

                        usedHSPs = usedHSPs + 1;
                        if VERBOSE, disp(['Reading HSP #' num2str(ihit) '
of hit ' num2str(di) ' (of ' num2str(TO-FROM+1) ' selected hits (of '
num2str(length(d_.Hits)) ' total) hits with E-val <= EVAL_WORST ('
num2str(EVAL_WORST) ')) ' d_.Hits(di).Name '.']), end

% MAKE SEQUENCE READY FOR POTENTIAL WRITING.
                        header = f_make_header(d_.Hits(di).Name,'',[],'
','all','()[]{},:/\;', {'MAG'; 'MAG:'; 'TPA_asm'; 'TPA_asm:';
'candidatus';'complete';'isolate';'genome';'strain';'assembly';'str.';'wh
ole';'shotgun';'sequence';'uncultured';}, 0, OUTPUTHEADERSEPARATOR);
                        sq = d_.Hits(di).HSPs(ihit).Alignment(3, :);
                        [sq i] = f_clean_RNA_sequence(sq);

% FIND IF THE HSP IS IN IGR.
                        if BASIC_BO_READ
                              if WRITE_HSP_SQS
                                    fwrite(id, ['>' header
OUTPUTHEADERSEPARATOR num2str(d_.Hits(di).HSPs(ihit).Expect)
OUTPUTHEADERSEPARATOR num2str(d_.Hits(di).HSPs(ihit).SubjectIndices(1))
OUTPUTHEADERSEPARATOR num2str(d_.Hits(di).HSPs(ihit).SubjectIndices(2))
char(10) sq char(10)]);
                              end
                              names_ = [names_; d_.Hits(di).Name];
                              evals_ = [evals_
d_.Hits(di).HSPs(ihit).Expect];
                              posinquery_ = [posinquery_;
d_.Hits(di).HSPs(ihit).QueryIndices];
                        else
                              headerd =
strtrim(regexprep(erase(d_.Hits(di).Name, NOTINNAMES), '\s+', ' '));
                              headerd = cut_string(headerd, ' ');
                              [fna fna] = unix(['grep ' headerd{1} ' '
ANNS_SA '/' headerd{2} '/*.fna*']);
                              if isempty(strfind(fna, 'Argument list too
long'))
                                          if ~isempty(strfind(fna, 'No such
file or directory')) | isempty(fna)
                                                if VERBOSE, disp(['Either '
ANNS_SA '/' headerd{2} ' not exist or ' headerd{1} ' not found by grep in
' ANNS_SA '/' headerd{2} '.']); end
                                                [fna fea] =
f_try_anns(headerd{1}, [ANNS_SA '/' headerd{2}], TRYTOLOAD, VERBOSE);
                                                if isempty(fna)
                                                      if VERBOSE,
disp(['Could not download annotations for ' headerd{2} ' ' headerd{3}
'.']), end
                                                end
                                          else
% if you get here, fna comes from grep. it is either with ':>' or without
it (when one file in directory).
                                                if isempty(strfind(fna,
':>'))
                                                      [fna fna] = unix(['ls
' ANNS_SA '/' headerd{2} '/*.fna*']);
```

```
                                        else
                                             fna = strsplit(fna,
':>');
                                             fna = fna{1};
                                        end
                                   end
                                   if ~isempty(fna)
                                        if VERBOSE,
disp('Annotations OK; trying to analyze...'), end
                                        fna(find(fna == char(10))) =
[];
                                        i = strfind(fna,
'genomic.fna');
                                        fea = [fna(1 : i - 1)
'feature_table.txt' fna(i + length('genomic.fna') : length(fna))];
                                        [x x] = unix(['head -n1 '
fna]);
                                        [j g] = f_read_genome(fna);
                                        if ~exist(fea, 'file')
                                             disp([fea ' NOT
EXIST!!!'])%, pause
                                        else
                                             [crs ors anns] =
f_read_anns(fea, length(g), INCLUDERRNAS, INCLUDENCRNAS, INCLUDETRNAS,
REMOVEPSEUDOGENES);
                                             if ~isempty(crs)
                                                  if
d_.Hits(di).HSPs(ihit).SubjectIndices(1) >
d_.Hits(di).HSPs(ihit).SubjectIndices(2)
                                                       i =
find(crs(:, 2) <= d_.Hits(di).HSPs(ihit).SubjectIndices(2) + OVERLAP);
                                                       j =
find(crs(:, 1) >= d_.Hits(di).HSPs(ihit).SubjectIndices(1) - OVERLAP);
                                                  else
                                                       i =
find(crs(:, 2) <= d_.Hits(di).HSPs(ihit).SubjectIndices(1) + OVERLAP);
                                                       j =
find(crs(:, 1) >= d_.Hits(di).HSPs(ihit).SubjectIndices(2) - OVERLAP);
                                                  end
                                                  if ~isempty(j) &
~isempty(i)
                                                       if j(1) -
i(length(i)) == 1 | INIGRONLY == 0
                      % IT IS EITHER IN IGR OR IT IS NOT BUT YOU TAKE
EVERYTHING.
                                                            if
j(1) - i(length(i)) ~= 1
                      % IT IS NOT IN IGR.

     if VERBOSE, disp(['Writing a subject sequence that is not in
IGR.']), end

     notinigrHSPs = notinigrHSPs + 1;
                      %
     synteny_ = [synteny_; anns(i(length(i)) + 1)];
                                                            else

     if VERBOSE, disp(['Writing a subject sequence.']), end
```

```
                    %
     synteny_ = [synteny_; [anns{i(length(i))} ' - ' anns{j(1)}]];
                                                                   end
                                                                   if
WRITE_HSP_SQS

     fwrite(id, ['>' header OUTPUTHEADERSEPARATOR
num2str(d_.Hits(di).HSPs(ihit).Expect) OUTPUTHEADERSEPARATOR
num2str(d_.Hits(di).HSPs(ihit).SubjectIndices(1)) OUTPUTHEADERSEPARATOR
num2str(d_.Hits(di).HSPs(ihit).SubjectIndices(2)) char(10) sq char(10)]);
                                                                   end
                                                                   if
RECONSTRUCTHITS

     v = f_reconstruct_blast_hit(d_.Hits(di).HSPs(ihit),
RECONSTRUCTHITS);

     if v.SubjectIndices(1) < 1

     v.SubjectIndices(1) = 1;

     end

     if v.SubjectIndices(2) < 1

     v.SubjectIndices(2) = 1;

     end

     if v.SubjectIndices(2) > v.SubjectIndices(1)

     fwrite(idrec, ['>' header OUTPUTHEADERSEPARATOR
num2str(d_.Hits(di).HSPs(ihit).Expect) OUTPUTHEADERSEPARATOR ...

                    num2str(v.SubjectIndices(1)) OUTPUTHEADERSEPARATOR
num2str(v.SubjectIndices(2)) char(10) ...

                    g(v.SubjectIndices(1) : v.SubjectIndices(2))
char(10)]);

     else

     fwrite(idrec, ['>' header OUTPUTHEADERSEPARATOR
num2str(d_.Hits(di).HSPs(ihit).Expect) OUTPUTHEADERSEPARATOR ...

                    num2str(v.SubjectIndices(2)) OUTPUTHEADERSEPARATOR
num2str(v.SubjectIndices(1)) char(10) ...

                    g(v.SubjectIndices(2) : v.SubjectIndices(1))
char(10)]);

     end
                                                                   end

                                                                   if
isempty([strfind(lower(x), 'contig') strfind(lower(x), 'scaffold')])

     gls_ = [gls_; d_.Hits(di).HSPs(ihit).SubjectIndices];
```

```
      glsn_ = [glsn_ (((d_.Hits(di).HSPs(ihit).SubjectIndices(1) +
d_.Hits(di).HSPs(ihit).SubjectIndices(2))) / 2) / length(g)];
                                                                      else

      gls_ = [gls_; [-1 -1]];

      glsn_ = [glsn_ -1];

      if VERBOSE, disp(['Contig or scaffold: gls = [-1 -1], glsn = -
1.']), end
                                                                      end

      names_ = [names_; d_.Hits(di).Name];

      evals_ = [evals_ d_.Hits(di).HSPs(ihit).Expect];

      posinquery_ = [posinquery_; d_.Hits(di).HSPs(ihit).QueryIndices];

                                                                      anns_
= [anns_; [anns{i(length(i))} ' - ' anns{j(1)}]];
                                                                      if
isempty(anns{i(length(i))})

      annsleft = [annsleft; ' '];
                                                                      else

      annsleft = [annsleft; anns{i(length(i))}];
                                                                      end
                                                                      if
isempty(anns{j(1)})

      annsright = [annsright;  ' '];
                                                                      else

      annsright = [annsright; anns{j(1)}];
                                                                      end

      leftannscoor_3prime = [leftannscoor_3prime; crs(i(length(i)), 2)];

      rightannscoor_5prime = [rightannscoor_5prime; crs(j(1), 1)];

                                                                      l = '
'; r = ' ';
                                                                      if
i(length(i)) - 1 > 0

      if ~isempty(anns{i(length(i)) - 1}), l = anns{i(length(i)) - 1};
end
                                                                      end
                                                                      if
j(1) + 1 <= length(anns)

      if ~isempty(anns{j(1) + 1}), r = anns{j(1) + 1}; end
                                                                      end

      annsext_ = [annsext_; [l ' - ' r]]; annsextleft = [annsextleft; l];
annsextright = [annsextright; r];
```

```
                                                                          l = '
'; r = ' ';
                                                                          if
i(length(i)) - 2 > 0

     if ~isempty(anns{i(length(i)) - 2}), l = anns{i(length(i)) - 2};
end
                                                                          end
                                                                          if
j(1) + 2 <= length(anns)

     if ~isempty(anns{j(1) + 2}), r = anns{j(1) + 2}; end
                                                                          end

     anns1extleft = [anns1extleft; l]; anns1extright = [anns1extright;
r];

                                                                          l = '
'; r = ' ';
                                                                          if
i(length(i)) - 3 > 0

     if ~isempty(anns{i(length(i)) - 3}), l = anns{i(length(i)) - 3};
end
                                                                          end
                                                                          if
j(1) + 3 <= length(anns)

     if ~isempty(anns{j(1) + 3}), r = anns{j(1) + 3}; end
                                                                          end

     anns2extleft = [anns2extleft; l]; anns2extright = [anns2extright;
r];

                                                                          l = '
'; r = ' ';
                                                                          if
i(length(i)) - 4 > 0

     if ~isempty(anns{i(length(i)) - 4}), l = anns{i(length(i)) - 4};
end
                                                                          end
                                                                          if
j(1) + 4 <= length(anns)

     if ~isempty(anns{j(1) + 4}), r = anns{j(1) + 4}; end
                                                                          end

     anns3extleft = [anns3extleft; l]; anns3extright = [anns3extright;
r];

                                                                          l = '
'; r = ' ';
                                                                          if
i(length(i)) - 5 > 0
```

```
      if ~isempty(anns{i(length(i)) - 5}), l = anns{i(length(i)) - 5};
end
                                                                  end
                                                                  if
j(1) + 5 <= length(anns)

      if ~isempty(anns{j(1) + 5}), r = anns{j(1) + 5}; end
                                                                  end

      anns4extleft = [anns4extleft; l]; anns4extright = [anns4extright;
r];


                                                                  l = '
'; r = ' ';
                                                                  if
i(length(i)) - 6 > 0

      if ~isempty(anns{i(length(i)) - 6}), l = anns{i(length(i)) - 6};
end
                                                                  end
                                                                  if
j(1) + 6 <= length(anns)

      if ~isempty(anns{j(1) + 6}), r = anns{j(1) + 6}; end
                                                                  end

      anns5extleft = [anns5extleft; l]; anns5extright = [anns5extright;
r];

                                                                  l = '
'; r = ' ';
                                                                  if
i(length(i)) - 7 > 0

      if ~isempty(anns{i(length(i)) - 7}), l = anns{i(length(i)) - 7};
end
                                                                  end
                                                                  if
j(1) + 7 <= length(anns)

      if ~isempty(anns{j(1) + 7}), r = anns{j(1) + 7}; end
                                                                  end

      anns6extleft = [anns6extleft; l]; anns6extright = [anns6extright;
r];
                                                            else
                        % IT IS NOT IN IGR AND YOU DON'T TAKE IT.
                                                                  if
VERBOSE, disp('BLAST hit is not in IGR and you don''t take it (INIGRONLY
= 1).'), end

      notinigrHSPs = notinigrHSPs + 1;
                                                            end
                                                      else
```

```
                                                            if VERBOSE,
disp('Either i or j is empty (e.g. when only 1 gene in annotations or
when at the end of contig/scaffold).'), end
                                                      end
                                                end
                                          else
                                                      if VERBOSE,
disp('Empty anns.'), end
                                                end
                                          end
                                    end
                              else
                                    if VERBOSE, disp([newline fna(1 :
length(fna) - 1) '. Go to the next hit.' newline]), end
                              end
                        end
                  end
            end
%     else
%           disp(['Do nothing as ' d_.Hits(di).Name ' is NOTNAME.'])
      end
end
if WRITE_HSP_SQS, fclose(id); end
if RECONSTRUCTHITS, fclose(idrec); end

if ~isempty(names_)
      if WRITE_HSP_SQS
            if CLEANHEADERS
                  fn_ = f_clean_fasta_headers(fn_, 1, '_', 0, '_',
':/\!''()";=,|+',
{'isolate';'strain';'candidatus';'complete';'sequence';'whole';'genome';'
shotgun';'chromosome';});
            end
      end
      if FILTER
            if WRITE_HSP_SQS
            %     [fn_ idel] = f_filter_fasta_by_sequence(fn_, 1, 0);
names_(idel(2 : 2 : length(idel)) / 2) = []; evals_(idel(2 : 2 :
length(idel)) / 2) = []; anns_(idel(2 : 2 : length(idel)) / 2) = [];
annsext_(idel(2 : 2 : length(idel)) / 2) = []; gls_(idel(2 : 2 :
length(idel)) / 2, :) = []; glsn_(idel(2 : 2 : length(idel)) / 2) = [];
                  [fn_ idel] = f_filter_fasta_by_ID(fn_, 1);
            end
            idel = idel(2:2:length(idel))/2;
            names_(idel) = []; anns_(idel) = []; annsext_(idel) = [];
gls_(idel, :) = []; glsn_(idel) = []; evals_(idel) = [];
posinquery_(idel, :) = [];
      %     anns2ext_(idel) = []; anns1ext_(idel) = [];
            annsleft(idel) = []; annsright(idel) = []; annsextleft(idel) =
[]; annsextright(idel) = []; anns1extleft(idel) = []; anns1extright(idel)
= [];
      end
end

%     filter_fasta_by_header_by_species
%     fn_ = [fn_ '.filtered'];

%     filter_for_header_keywords
%     fn_ = [fn_ '.filtered'];
```

```
% if VERBOSE, if ~isempty(names_), [names_ anns_], end, end
```

Supplementary File S12. ChatGPT-assisted analysis of functional relations between TraC and TadA proteins.

Question to ChatGPT 5.5:

Tell me, whether conjugal transfer TraC proteins and TadA family conjugal transfer-associated ATPases are functionally related.

ChatGPT 5.5 answer:

Yes, but only at a broad systems level.

TraC conjugal transfer proteins and TadA-family conjugal transfer-associated ATPases are both associated with bacterial cell-envelope transport machines that use ATP-powered components for conjugation, secretion, pilus assembly, or related macromolecular transfer. For example, TraC can be annotated as a type IV secretion/conjugal pilus assembly protein with ATP-binding activity, and some TraC-like proteins map to VirB4-like T4SS ATPases UniProt TraC entry, NCBI VirB4-like TraC/CD1110 family annotation. TadA-family proteins are also secretion/pilus ATPases; classic phylogenetic work places TadA in the type II/type IV secretion NTPase superfamily, close to VirB11/TrbB-like ATPases rather than simple DNA-processing enzymes Planet et al., PNAS 2001.

But they are **not usually the same protein family or direct functional equivalents**. TraC, depending on the plasmid/system, often refers to a T4SS conjugation component such as a pilus assembly or VirB4-like ATPase. TadA/CpaF proteins are motors of the Tad, or tight-adherence, pilus system; recent work describes TadA/CpaF as driving pilus extension and retraction through ATP-dependent hexameric motor activity Hohl et al., 2024.

So the clean answer is:

| Relationship | Assessment |
|---|---|
| Same pathway neighborhood? | Often yes: conjugation/secretion/pilus systems |
| Same broad biochemical role? | Yes: ATP-powered transport/assembly motors |
| Same protein family? | Usually no |
| Functionally interchangeable? | No, not without strong sequence/domain/context evidence |
| Likely evolutionary relationship? | Distant, through the broader type II/type IV secretion NTPase machinery |

If you're annotating genomes, I'd treat them as **functionally related but distinct ATPase families** unless domain analysis or synteny shows that a specific "TraC" is actually being used as a local name for a TadA-like motor.

Supplementary File S13. ChatGPT-assisted analysis of functional relations between MinD/ParA and Ssd proteins.

Question to ChatGPT 5.5:

Tell me what is the relationship between the MinD/ParA family ATP-binding proteins and septum site-determining proteins Ssd?

ChatGPT 5.5 answer:

I'll ground this in current protein-family/domain references and the specific "Ssd" terminology, since names can be overloaded across bacteria.

The relationship is **homology plus functional analogy**, but with some important nuance.

**MinD/ParA family ATP-binding proteins** are bacterial/archaeal "positioning ATPases." They use ATP binding/hydrolysis to spatially organize things inside cells. The classic examples are **ParA**, which helps segregate chromosomes/plasmids, and **MinD**, which helps position the cell-division machinery so the FtsZ/Z-ring forms at the correct place. Reviews describe ParA/MinD ATPases as systems that position genetic and protein cargos; MinD specifically is tied to divisome positioning.

**Ssd proteins**—"septum site-determining" proteins—are **MinD-like / ParA-MinD-family-related proteins** involved in septum placement or septation control, especially discussed in **mycobacteria/actinobacteria**. In *Mycobacterium tuberculosis*, **Ssd/Rv3660c** was originally identified as an ortholog of septum-site-regulating proteins and was reported to affect cell division: overexpression promoted filamentation and a dormancy/stress-like response, consistent with inhibition or misregulation of septation.

The key nuance is that **not every protein annotated as Ssd is necessarily the true functional equivalent of E. coli MinD**. A 2023 mycobacterial study compared candidate *M. tuberculosis* and *M. smegmatis* MinD homologues and concluded that **Rv1708/MSMEG_3743**, not **Rv3660c/MSMEG_6171**, are the closer/true *E. coli* MinD homologues based on complementation, overexpression, ATPase activity, and structural comparisons.

So, in one sentence: **Ssd proteins are septum-site regulators that belong to, or are closely related to, the broader MinD/ParA ATPase superfamily, but "Ssd" is a functional/annotation label and can include MinD-like proteins that differ in how closely they match canonical MinD.**

A useful way to think about it:

| Term | What it means | Relationship |
|---|---|---|
| **ParA/MinD family ATPase** | Broad family of Walker-type ATPases used for spatial organization | Superfamily/category |
| **MinD** | A ParA/MinD-family ATPase that positions the division site, often via MinC/MinE systems | A specific subfamily/function |
| **Ssd** | "Septum site-determining" protein, often MinD-like in actinobacteria/mycobacteria | Related/overlapping annotation; may be MinD-like but not always canonical MinD |

In genome annotations, you may therefore see a protein called **"MinD/ParA family ATP-binding protein," "septum site-determining protein," "Ssd," or "MinD-like ATPase"** for overlapping reasons: sequence similarity to ParA/MinD ATPases, ATP-binding motifs, and predicted or tested roles in septum placement.

B

Question to ChatGPT 5.5:

Give me a source and reference of this information."Ssd proteins are MinD-like ATPases / members of the ParA–MinD ATPase superfamily, and some Ssd proteins are orthologous to septum-site-regulating proteins in related actinobacteria."

ChatGPT 5.5 answer:

Thought for a couple of seconds

Yes. The best source for that exact statement is:

**Primary reference**

England, K., Crew, R., & Slayden, R. A. (2011). **"Mycobacterium tuberculosis septum site determining protein, Ssd encoded by rv3660c, promotes filamentation and elicits an alternative metabolic and dormancy stress response."** *BMC Microbiology*, 11, 79. DOI: **10.1186/1471-2180-11-79**. PMID: **21504606**.

This paper states that **Ssd, encoded by *rv3660c*, was discovered to be an ortholog of septum-site-regulating proteins in actinobacteria**, and their bioinformatic analysis compared the *M. tuberculosis* Ssd protein with proteins annotated as **MinD** and **Ssd**. It also reports that increased *ssd* expression inhibited septum formation, producing elongated cells lacking septa.

A supporting review/source for the broader "MinD/ParA family" framing is:

A polished sentence you can cite would be:

Ssd/Rv3660c in *Mycobacterium tuberculosis* has been described as an ortholog of septum-site-regulating proteins in actinobacteria and is related to proteins annotated as MinD/Ssd; more broadly, MinD belongs to the ParA/MinD family of Walker-type ATPases. However, Ssd should be described as MinD-like or ParA/MinD-family-related rather than assumed to be the canonical *E. coli* MinD ortholog.

Supplementary File S14. ChatGPT-assisted analysis of functional properties of SCO6880 proteins (A), TrbL/VirB6 proteins (B), RAD23 proteins (C), and TrbC/VirB2 family proteins (D).

**A**

Question to ChatGPT 5.5:

What are the SCO6880 family proteins?

ChatGPT 5.5 answer:

The **SCO6880 family protein** is not a well-characterized, single-function protein like an enzyme you'd find in textbooks. Instead, it's a **protein family designation** used in genome annotations—especially in bacteria of the genus *Streptomyces* and related actinomycetes.

**What the name means**

- **"SCO6880"** refers to a specific gene locus (SCO6880) in the model organism *Streptomyces coelicolor*.
- Proteins in other organisms that are **sequence-similar (homologous)** to that gene product are grouped as the **SCO6880 family**.

**What is known about the protein family**

- These proteins are typically:
  - **Bacterial proteins**, common in Actinobacteria (e.g., *Streptomyces*)
  - Around **~480–500 amino acids long** in many species
- They are usually annotated as:
  - **"hypothetical protein"** or **"uncharacterized protein"**
  - Meaning **no experimentally confirmed function** is available yet

**Functional insight (what scientists suspect)**

- Because they are conserved across related bacteria, they are likely:
  - **Not random proteins** (they're maintained by evolution)
  - Possibly involved in **core cellular processes** (e.g., metabolism, regulation, or cell structure)
- However, there is **no definitive biochemical function assigned** so far.

**Bottom line**

A **SCO6880 family protein** is:

A **conserved but uncharacterized bacterial protein family**, originally defined from a gene in *Streptomyces coelicolor*, with **unknown precise function** but likely biological importance due to its conservation.

If you want, I can dig into predicted domains, structural models, or any specific organism's SCO6880 protein to give more clues about its possible function.

**B**

Question to ChatGPT 5.5:

What is the TrbL/VirB6 family protein?

ChatGPT 5.5 answer:

The **TrbL/VirB6 family proteins** are **membrane proteins that form part of bacterial secretion systems**, especially those used to transfer DNA or proteins between cells.

---

 **What they are**

- Integral **inner membrane proteins** (span the bacterial membrane multiple times)
- Core components of **Type IV secretion systems (T4SS)**
- Found in many bacteria, especially those that:
  - Conjugate (exchange DNA)
  - Infect host cells
- Named after:
  - **TrbL** → conjugation systems in plasmids
  - **VirB6** → virulence system protein in Agrobacterium tumefaciens

---

 **What they do (main function)**

6

They help build and operate a **transport channel** that moves molecules across the cell envelope.

**Key roles:**

- Form part of the **membrane channel** of the secretion system
- Help **transfer DNA or proteins**:
  - Between bacteria (conjugation)
  - From bacteria into host cells (infection)
- Likely contribute to:
  - **Substrate translocation**
  - **Assembly/stability** of the secretion machinery

---

□ **Where they act**

- Located in the **inner (cytoplasmic) membrane**
- Work together with other T4SS proteins (VirB1–VirB11, etc.)
- Connect cytoplasmic energy sources (ATPases) to the export channel

---

□ **Functional contexts**

**1. Bacterial conjugation**

- Transfer of plasmids between bacteria
- TrbL/VirB6 is part of the machinery that moves DNA through the membrane

**2. Pathogenesis**

- In pathogens, T4SS injects effector molecules into host cells
- Example: Agrobacterium tumefaciens transfers DNA into plants (causing crown gall disease)

---

□ **Key features**

- **Highly hydrophobic** (many transmembrane helices)
- **Conserved across diverse bacteria**
- Often essential for a functional secretion system
- No enzymatic activity → mainly **structural/transport role**

---

□ **Big picture**

If we place this with your earlier proteins:

- **MinD/ParA** → spatial organization inside the cell
- **NlpC/P60** → remodeling the cell wall
- **TrbL/VirB6 → moving molecules across the cell envelope**

---

□ **Bottom line**

The **TrbL/VirB6 family proteins** are **core inner-membrane components of Type IV secretion systems**, enabling bacteria to **transfer DNA or proteins to other cells**, which is crucial for **conjugation and virulence**.

---

If you want, I can map all these protein families onto a single bacterial cell diagram so you can see how they work together spatially.

**C**

Question to ChatGPT 5.5:

What are the RAD23 proteins?

ChatGPT 5.5 answer:

RAD23 refers to a family of proteins involved primarily in **DNA repair** and **protein degradation** pathways.

There are a few closely related versions depending on the organism:

- In yeast, the original **Rad23** protein was discovered as part of the **nucleotide excision repair (NER)** system.
- In humans, there are two homologs:
  - RAD23A
  - RAD23B

**Main functions of RAD23 proteins**

**1. DNA repair**

RAD23 proteins help repair DNA damage caused by:

- ultraviolet (UV) radiation
- chemical mutagens
- oxidative stress

They work in the **nucleotide excision repair** pathway, which removes damaged sections of DNA and replaces them with correct sequences.

A major interaction partner is:

- XPC

RAD23 stabilizes XPC and helps detect DNA lesions.

**2. Protein degradation (ubiquitin–proteasome system)**

RAD23 proteins also shuttle ubiquitinated proteins to the proteasome for destruction.

They contain:

- a **UBA domain** (binds ubiquitin)
- a **UBL domain** (interacts with the proteasome)

So RAD23 acts like a delivery adaptor between tagged proteins and the cellular degradation machinery.

**Why RAD23 matters**

RAD23 proteins are important in:

- genome stability
- cancer biology
- aging research
- neurodegenerative disease studies

Defects in the NER pathway involving XPC/RAD23 are associated with disorders such as:

- Xeroderma Pigmentosum

**Simple analogy**

You can think of RAD23 as:

- partly a **DNA damage inspector**
- partly a **waste-management courier** for damaged proteins

It helps cells maintain both:

1. clean DNA
2. clean protein quality control

If you want, I can also explain:

- the RAD23 protein structure,
- how it works mechanistically in NER,
- differences between RAD23A vs RAD23B,

- or its role in cancer and proteasome biology.

Question to ChatGPT 5.5:

**D**

Characterize functionally TrbC/VirB2 family proteins.

ChatGPT 5.5 answer:

I'll characterize this as a family-level annotation, since "TrbC/VirB2 family protein, membrane protein" usually points to a secretion/conjugation apparatus component rather than a standalone enzyme.

Thought for 4s

**Functional characterization**

| **Protein annotation** | **Likely function** | **Functional category** | **Notes** |
|---|---|---|---|
| **TrbC/VirB2 family protein, membrane protein** | Likely **major pilin / pilus subunit** of a **Type IV secretion system** or bacterial conjugation system | Type IV secretion / conjugative DNA transfer / virulence-associated cell-envelope machinery | VirB2 and TrbC-family proteins are small, hydrophobic pilin-like proteins that form or contribute to the extracellular/conjugative pilus of Type IV secretion systems. These systems can mediate plasmid conjugation, horizontal DNA transfer, and in some pathogens delivery of effector proteins or nucleoprotein complexes into host cells. |

In annotation language, I would write:

**TrbC/VirB2 family protein, membrane protein:** "Putative Type IV secretion system major pilin/conjugative pilus subunit; likely membrane-associated precursor that contributes to pilus assembly and may participate in conjugative DNA transfer or effector translocation, depending on genomic context."

A cautious version would be:

**"Predicted membrane-associated TrbC/VirB2-family pilin, likely part of a Type IV secretion or conjugative transfer apparatus; no independent enzymatic activity expected."**

Supplementary File S15. ChatGPT-assisted analysis of functional properties of type VII secretion protein EccE, PrgI family protein (A), excisionase family DNA binding proteins (B), helix-turn-helix domain-containing protein MAD1 (C), Cif family virulence factors (D), NlpC/P60 family proteins (E), C40 family peptidases (F) and YjjA family protein (G).

**A**

Question to ChatGPT 5.5:

What is the functional characterization of the following proteins: type VII secretion protein EccE, PrgI family protein

ChatGPT 5.5 answer:

I'll treat these as protein-family annotations and distinguish what is known experimentally from what is usually inferred by homology.

**Functional characterization**

| **Protein annotation** | **Likely function** | **Functional category** | **Notes** |
|---|---|---|---|
| **Type VII secretion protein EccE** | Core component of a **Type VII / ESX secretion system** | Protein secretion / virulence-associated envelope transport | EccE is part of the conserved ESX membrane machinery, together with EccB, EccC, EccD and MycP. In mycobacteria, ESX systems secrete proteins such as Esx, Esp, PE/PPE substrates and are linked to virulence, metal homeostasis, nutrient acquisition, and host immune modulation depending on the ESX subtype. |
| **PrgI family protein** | Usually associated with **Type III secretion system needle/injectisome structure**, but some database "PrgI family" entries are annotated as functionally uncharacterized | Secretion apparatus structural protein / possible virulence factor | Experimentally characterized **PrgI** from *Salmonella* forms the T3SS needle complex, which helps inject effector proteins into host cells. However, the Pfam/InterPro "PrgI family" entry itself is described as functionally uncharacterized, so the safest annotation is "putative T3SS needle-related protein" unless genomic context confirms nearby T3SS genes. |

In genome-annotation language, I would write:

**EccE:** "Membrane-associated core component of the Type VII/ESX secretion system; likely involved in export of ESX substrates and may contribute to virulence or metal/nutrient homeostasis depending on the ESX locus."

**PrgI family protein:** "Putative Type III secretion system needle/injectisome-associated protein; likely structural component involved in effector translocation, but family-level annotation may be uncharacterized without supporting genomic context."

**B**

Question to ChatGPT 5.5:

What do the excisionase family DNA binding proteins do?

ChatGPT 5.5 answer:

**Functional characterization**

| Protein annotation | Likely function | Functional category | Notes |
|---|---|---|---|
| **DNA-binding protein, excisionase family** | Likely **excisionase / recombination directionality factor** involved in site-specific DNA recombination | Mobile genetic element regulation / prophage or integrative element excision | Excisionase-family proteins are usually small DNA-binding proteins encoded by bacteriophages, prophages, integrative conjugative elements, or other mobile genetic elements. They assist integrases by promoting **excision** of an integrated DNA element from the host chromosome, often by bending DNA or altering the synaptic recombination complex. |

In annotation language, I would write:

**DNA-binding protein, excisionase family:** "Putative excisionase/recombination directionality factor; small DNA-binding protein likely involved in site-specific excision of a prophage, integrative conjugative element, or other mobile genetic element from the host genome."

A more cautious version:

**"Predicted excisionase-family DNA-binding protein associated with mobile genetic element recombination; likely modulates integrase-mediated excision, but specific target sites and element type require genomic-context confirmation."**

**C**

Question to ChatGPT 5.5:

Might it be somehow biologically functionally related to the following protein:methylation-associated defense system helix-turn-helix domain-containing protein MAD1

ChatGPT 5.5 answer:

I'll compare them at the system/function level rather than just the names, because both annotations involve DNA-binding proteins but can belong to different mobile-element or defense contexts.

Thought for 4s

Yes — **they might be functionally related**, but the relationship is more likely **contextual/system-level** than "same protein function."

| Protein | Core inferred function | How it could relate to MAD1 |
|---|---|---|
| **DNA-binding protein, excisionase family** | Recombination directionality factor, usually helping excise prophages, integrative conjugative elements, or other mobile elements from chromosomes | Could indicate a nearby **mobile genetic element**, prophage, or integrative element |
| **Methylation-** | Component of **MADS**, a recently | Could be part |

| Protein | Core inferred function | How it could relate to MAD1 |
|---|---|---|
| **associated defense system HTH domain-containing protein MAD1** | described bacterial anti-phage defense system; MAD1 is predicted to contain a winged helix-turn-helix DNA-binding domain and deletion of **mad1** reduces restriction but does not affect methylation | of a nearby **anti-phage defense island** or mobile defense module |

The key shared theme is **DNA-binding in phage/mobile-element biology**. Excisionases are typically small DNA-binding proteins that help integrases drive excision of integrated phage or mobile-element DNA; they act as architectural/recombination directionality factors. MAD1, by contrast, belongs to the **methylation-associated defense system**, or MADS, which was described as a widespread bacterial defense system that limits phage infection and can interact with CRISPR-Cas immunity.

So the cautious interpretation is:

**Possible biological relationship:**

They may both be part of a broader **phage-defense / mobile genetic element region**, especially if the genes are adjacent or occur in the same genomic island.

**Less likely direct relationship:**

The excisionase-family protein is probably not itself a MAD1 homolog and probably does not perform the same role as MAD1. Excisionase proteins usually regulate **site-specific recombination/excision**, while MAD1 appears to be a **DNA-binding component of a methylation-associated restriction/defense system**. In the MADS study, **mad1 deletion reduced restriction but did not affect DNA methylation**, suggesting MAD1 contributes to the defensive restriction arm rather than methylation itself.

A good annotation note would be:

**"The excisionase-family DNA-binding protein and MAD1 are not necessarily homologous, but they may be functionally linked by genomic context if co-localized: the excisionase suggests a mobile/prophage or integrative element, while MAD1 suggests an anti-phage methylation-associated defense system. Together, they could indicate a defense island or mobile genetic element carrying phage-defense genes."**

To assess this more confidently, check whether the genes are near each other and whether the neighborhood also contains **integrase, recombinase, methyltransferase, nuclease, ATPase, CRISPR-Cas, restriction-modification, or other defense-associated genes**.

**D**

Question to ChatGPT 5.5:

Characterize functionally or biologically these proteins: Cif family virulence factors.

ChatGPT 5.5 answer:

Thought for a few seconds

**Functional characterization**

| Protein annotation | Likely function | Functional category | Notes |
|---|---|---|---|
| **Cif family** | Likely bacterial **cycle-** | Virulence factor / host-cell | In many enteric pathogens, Cif proteins are |

| Protein annotation | Likely function | Functional category | Notes |
|---|---|---|---|
| **virulence factor** | **inhibiting factor** or Cif-like effector that manipulates host-cell processes | modulation / possible Type III secretion effector | **cyclomodulins**: injected effectors that disrupt the eukaryotic cell cycle, commonly causing host-cell arrest around the **G2/M transition**. Cif-family proteins are found in pathogens of mammals and invertebrates and are often associated with Type III secretion systems. |

In annotation language, I would write:

**Cif family virulence factor:** "Putative Cif-family bacterial virulence effector/cyclomodulin; likely manipulates host-cell cycle or ubiquitin-dependent host regulatory pathways during infection, possibly as a Type III secretion substrate."

A more cautious version:

**"Predicted Cif-family virulence-associated protein; may function as a secreted effector that alters host-cell cycle progression or epithelial-cell physiology, but specific activity depends on the Cif subtype and organismal context."**

One important caveat: **"Cif" can refer to more than one virulence-factor lineage.** In enteric bacteria such as pathogenic *E. coli*, Cif usually means **cycle-inhibiting factor**, a cyclomodulin effector. In *Pseudomonas aeruginosa*, Cif can mean **CFTR inhibitory factor**, an α/β-hydrolase-family virulence factor that alters epithelial transporter trafficking, including CFTR degradation. So the best biological interpretation depends on the source organism and neighboring genes.

**E**

Question to ChatGPT 5.5:

Now characterize functionality of the NlpC/P60 family proteins.

ChatGPT 5.5 answer:

The **NlpC/P60 family proteins** are a large group of **bacterial enzymes that remodel or break down peptidoglycan**, the rigid mesh that forms the bacterial cell wall.

---

#### □ What they do (core function)

7

- They act as **peptidases (hydrolases)** that cut specific bonds in peptidoglycan
- Typically target **amide or peptide bonds** in the cell wall
- This activity is essential for:
  - **Cell growth** (expanding the wall)
  - **Cell division** (separating daughter cells)
  - **Cell wall turnover and recycling**

---

#### □ Why the name "NlpC/P60"?

- **NlpC**: originally identified as a *lipoprotein* in bacteria
- **P60**: a major secreted protein in Listeria monocytogenes
- Both were found to share a **common catalytic domain**, so they were grouped

into one family

---

⚙ **Structural & biochemical features**

- Belong to the **cysteine peptidase superfamily**
- Have a conserved **catalytic triad** (usually cysteine–histidine–polar residue)
- Often modular:
  - May include **cell wall–binding domains** (e.g., LysM domains)
  - Can be secreted, membrane-anchored, or cytoplasmic

---

🔑 **Functional roles (varies by organism)**

Different NlpC/P60 proteins can have specialized roles:

- **Cell separation enzymes**

→ split newly formed daughter cells after division

- **Autolysins**

→ controlled self-degradation of the cell wall

- **Virulence factors** (in pathogens)

→ help bacteria invade or escape host cells

- **Protein turnover / signaling**

→ some have adapted roles beyond simple wall degradation

---

□ **Big picture**

If MinD/ParA proteins are about **spatial organization**, NlpC/P60 proteins are about **physical remodeling**:

They act like **molecular scissors** that carefully cut the bacterial cell wall so the cell can grow, divide, and sometimes interact with its environment.

---

□ **Bottom line**

The **NlpC/P60 family proteins** are **cysteine peptidases that cleave peptidoglycan**, playing essential roles in **bacterial cell wall remodeling, division, and sometimes virulence**.

---

If you want, I can connect this back to your earlier SCO6880 protein—sometimes uncharacterized proteins end up having hidden NlpC/P60-like domains, which can hint at their function.

**F**

Question to ChatGPT 5.5:

Characterize functionally C40 family peptidases. Couldn't be they functionally related to the NlpC/P60 family proteins?

ChatGPT 5.5 answer:

I’ll compare the two at the catalytic-family and pathway level, because both are “peptidase-like” but they usually act on very different substrates.

Thought for 5s

Yes — **they could be functionally related**, and in this case the link is stronger than for many “peptidase” comparisons.

**Functional characterization**

| Protein annotation | Likely function | Functional category | Notes |
|---|---|---|---|
| **C40 family peptidase** | Predicted **bacterial cell-wall modifying peptidase**, often a cysteine peptidase involved in peptidoglycan processing | Cell wall remodeling / peptidoglycan turnover / cell separation or sporulation | MEROPS describes peptidase family C40 as containing bacterial **cell-wall modifying enzymes**, and the C40 holotype is a bacterial dipeptidyl-peptidase VI with a probable role in peptidoglycan turnover during sporulation. |

**Relationship to NlpC/P60 family protein**

They are **not necessarily the same family**, but they can be **biologically related** because both are commonly linked to **bacterial peptidoglycan/cell-wall remodeling**.

| Comparison point | NlpC/P60 family protein | C40 family peptidase |
|---|---|---|
| Broad activity | Peptidoglycan hydrolase / endopeptidase | Cell-wall modifying peptidase |
| Catalytic type | Usually cysteine peptidase, papain-like CA clan | Cysteine peptidase, MEROPS C40 |
| Likely substrate | Peptide bonds within peptidoglycan stem peptides or crosslinks | Bacterial cell-wall / peptidoglycan-associated peptides |
| Biological role | Cell-wall turnover, daughter-cell separation, division, host interaction | Cell-wall expansion, cell separation, sporulation, peptidoglycan turnover |

NlpC/P60 proteins are well described as peptidoglycan hydrolases important for bacterial cell-wall turnover, division, and host interactions. InterPro's C40 entry similarly links this family to peptidoglycan binding, cell-wall expansion, and cell separation during vegetative growth.

So I would annotate the relationship as:

**"The C40 family peptidase and the NlpC/P60 family protein may be functionally related as bacterial cell-wall remodeling enzymes. Both are likely peptidoglycan-associated peptidases/hydrolases, potentially involved in cell separation, cell-wall turnover, sporulation, or envelope remodeling. They are not necessarily homologous, but they may act in the same biological pathway or cellular process."**

A concise cautious version:

**"Predicted C40 cysteine peptidase; likely bacterial cell-wall/peptidoglycan-modifying enzyme. It could be functionally related to an NlpC/P60 protein through shared roles in peptidoglycan remodeling, although the two belong to distinct peptidase families."**

**G**

Question to ChatGPT 5.5:

And now characterize this one: YjjA family protein

ChatGPT 5.5 answer:

I'll treat "YjjA family" cautiously because this label is often attached to small conserved bacterial proteins with limited direct experimental characterization.

Thought for a few seconds

**Functional characterization**

| Protein annotation | Likely function | Functional category | Notes |
|---|---|---|---|
| **YjjA family protein** | Poorly characterized bacterial protein; often annotated as **DUF2501/YjjA-family** and predicted to be **periplasmic or envelope-associated** | Protein of unknown function; possible cell-envelope/periplasmic role | YjjA is generally not a well-characterized enzymatic protein. In *E. coli* K-12, yjjA is annotated as a **DUF2501 domain-containing protein**, and STRING describes it as a putative DUF2501-family periplasmic protein, possibly a glycoprotein/receptor-like protein. |

In annotation language, I would write:

**YjjA family protein:** “Predicted YjjA/DUF2501-family protein of unknown function; likely bacterial envelope- or periplasm-associated protein, with no confidently assigned catalytic activity.”

A more cautious version:

**“Conserved bacterial YjjA-family protein; function uncharacterized. Possible periplasmic/cell-envelope-associated role inferred from database annotations and predicted localization, but biological activity remains unclear.”**

Biologically, I would **not** assign this directly to peptidoglycan remodeling, secretion, defense, or virulence without genomic context. If it is located near cell-envelope genes, transport genes, or stress-response genes, it may be part of an envelope-associated process, but the family name alone is too vague for a strong functional claim.

Supplementary File S16. ChatGPT-assisted analysis of functional relations between genomic contexts of IGRs with predicted structure candidates in *Frankia casuarinae* and ms1 RNA synteny.

Question to ChatGPT 5.5:

These are NCBI genomic annotations of a succession of bacterial genes: Fic family protein, HAD-IB hydrolase, septum site-determining protein Ssd, TadA family conjugal transfer-associated ATPase. The genes are adjacent to each other in bacterial genomes and represent a segment of bacterial genomes. Tell me, which bacterial cellular biological processes/functions are related to the segment with such genes and also tell me which of the genes, if not all, are functionally/biologically related.

ChatGPT 5.5 answer:

I'll treat this as a genome-neighborhood/operon inference problem: identify likely functions of each annotation, then ask whether the adjacency suggests a shared biological module rather than four unrelated housekeeping genes.

The segment most likely points to a **cell-envelope / surface-structure / cell-cycle-stress module**, not a single classical metabolic pathway.

**Likely biological processes represented**

**1. Type IV/Tad pilus assembly, adhesion, surface colonization, and possibly DNA transfer**
The strongest signal is **TadA family conjugal transfer-associated ATPase**. TadA-like ATPases are part of **Tad/Flp type IV pilus systems**, which assemble pili used for bacterial surface interaction. Tad/T4P systems are linked to **adherence, motility, biofilm formation, host interaction, DNA uptake, and horizontal gene transfer**, depending on the organism and the rest of the locus. Recent reviews describe Tad pili as conserved bacterial type IV pilus machineries used in many physiological lifestyles, with T4P broadly facilitating motility, adherence, and DNA uptake.

**2. Septum positioning, cell division control, filamentation, and dormancy/stress physiology**
**Septum site-determining protein Ssd** is the clearest cell-division gene. In *Mycobacterium tuberculosis*, Ssd/Rv3660c was shown to inhibit septum formation when overexpressed, producing elongated cells lacking septa, and to induce a dormancy/alternative metabolism transcriptional response.

This suggests the segment may couple **surface-structure biogenesis** with **cell-cycle state or stress adaptation**.

**3. Fic-mediated post-translational regulation, toxin-like activity, stress arrest, or defense**
**Fic family proteins** often catalyze AMPylation or related post-translational modifications. Many bacterial Fic proteins are toxin-like regulators; some inhibit DNA gyrase or topoisomerase IV, causing growth arrest, filamentation, SOS induction, and disruption of chromosome segregation/cell separation.

So the Fic gene may connect biologically to **stress-induced growth modulation**, **cell-cycle arrest**, **toxin–antitoxin-like behavior**, or **defense/mobile-element biology**.

**4. Generic phosphatase/hydrolase activity, possible regulatory or metabolic accessory role**
**HAD-IB hydrolase** is the least specific annotation. HAD superfamily proteins include phosphatases and other aspartate-nucleophile hydrolases; InterPro classifies HAD-IB as a haloacid dehalogenase-superfamily subgroup.

Without the sequence/domain details, I would not assign it confidently to the same process. It could be a **small-molecule phosphatase**, **regulatory phosphatase**, or simply an accessory enzyme carried with the locus.

#### Which genes are functionally related?

**Strongly related:**
**Ssd + TadA** are very likely related. In actinobacterial/mycobacterial contexts, Ssd/Rv3660c sits in or near a type IV/Tad/Flp pilus-associated region, and TadA is a core ATPase for that kind of system. Search results for *M. tuberculosis* note that Rv3657c–Rv3660c resemble type IV pili/type II-IV secretion/tad genes, and later work labels Rv3660c as **tadZ** in an Flp type IV pilus operon context.

So **Ssd may be a cell-cycle/septation regulator embedded in or evolutionarily related to a Tad pilus locus**, rather than a random neighboring gene.

**Possibly related:**
**Fic + Ssd** may be biologically related through their effects on **cell division, filamentation, stress arrest, and persistence-like states**. Fic proteins can inhibit topoisomerase IV and cause filamentation; Ssd also affects septum formation and dormancy-related transcription.

However, this is a functional convergence, not proof that the two proteins directly interact.

**Weakly or uncertainly related:**
**HAD-IB hydrolase** may be part of the same conserved neighborhood, but the annotation alone is too broad. It could be a regulatory enzyme associated with the module, but I would not call it functionally linked without synteny conservation, co-occurrence, operon prediction, or domain/substrate information.

#### Best interpretation

This gene block most plausibly represents a **Tad/type IV pilus-associated genomic segment with cell-cycle/stress-regulatory accessories**. The core biological theme is:

**surface attachment / pilus or secretion-system assembly + septation/cell-cycle control + stress-growth modulation.**

A more conservative summary would be:

| Gene | Likely role | Relationship t[illegible] |
|---|---|---|
| Fic family protein | toxin-like AMPylating/regulatory protein; stress arrest, chromosome/cell-division effects | Possibly accessory s[illegible] regulator |
| HAD-IB hydrolase | broad hydrolase/phosphatase-like enzyme | Uncertain; possible a[illegible] enzyme |
| Ssd | septum placement/cell division, filamentation, dormancy response | Strongly relevant |
| TadA ATPase | Tad/type IV pilus assembly/conjugation-associated ATPase | Strongly relevant |

So, **Ssd and TadA are the most clearly functionally related pair**, likely tied to a Tad/Flp/type IV pilus locus. **Fic is plausibly related at the level of stress, growth arrest, and cell-division modulation. HAD-IB is the least interpretable from annotation alone.**

Question to ChatGPT 5.5:

Now I will send you 165 genomic contexts, one context per a line, terminated by dots. The contexts consist of NCBI genomic annotations, separated by the '-' characters. They represent segments of bacterial genomes that contain the genes encoding the proteins whose bilogical/functional roles are characterized by the annotations. Tell me, which of the contexts are biologically/functionally related to the characterisctics that you gave me above. Indentify the contexts using their order numbers. Here are the contexts: 3-oxoacyl-[acyl-carrier-protein] synthase III C-terminal domain-containing protein - hypothetical protein - hypothetical protein - recombinase family protein - hypothetical protein - hypothetical protein - albusnodin family lasso peptide - albusnodin/ikarugamycin family macrolactam cyclase - lasso peptide biosynthesis B2 protein - GNAT family N-acetyltransferase - replication initiator - recombinase family protein - recombinase family protein - MFS transporter. 7-carboxy-7-deazaguanine synthase - 6-carboxytetrahydropterin synthase QueD - 16S rRNA methyltransferase - preQ(1) synthase - IS5/IS1182 family transposase - IS5 family transposase - transposase - site-specific integrase - hypothetical protein - tyrosine-type recombinase/integrase - hypothetical protein - helix-turn-helix domain-containing protein - DUF397 domain-containing protein - hypothetical protein. AAA family ATPase - hypothetical protein - CBASS oligonucleotide cyclase - helix-turn-helix domain-containing protein - ImmA/IrrE family metallo-endopeptidase - hypothetical protein - IS110 family transposase - NUDIX hydrolase - hypothetical protein - hypothetical protein - hypothetical protein - hypothetical protein - hypothetical protein - hypothetical protein. ABC transporter ATP-binding protein - alpha-hydroxy acid oxidase - chorismate mutase - tryptophan 7-halogenase - ParB/RepB/Spo0J family partition protein - ISAs1 family transposase - IS66 family transposase - carbon starvation CstA family protein - hypothetical protein - DUF3311 domain-containing protein - sodium:solute symporter family protein - sigma-70 family RNA polymerase sigma factor - hypothetical protein - lipid-transfer protein. ABC transporter ATP-binding protein - lasso peptide biosynthesis B2 protein - lasso peptide biosynthesis PqqD family chaperone - asparagine synthase-related protein - lasso RiPP family leader peptide-containing protein - AfsR/SARP family transcriptional regulator - peptidoglycan-binding protein - alpha/beta hydrolase - four-helix bundle copper-binding protein - amidohydrolase family protein - methionine adenosyltransferase - DUF309 domain-containing protein - toll/interleukin-1 receptor domain-containing protein - TAXI family TRAP transporter solute-binding subunit. ABC transporter permease - ABC transporter substrate-binding protein - helix-turn-helix transcriptional regulator - hypothetical protein - response regulator transcription factor - 4-hydroxyphenylacetate 3-hydroxylase family protein - hypothetical protein - helix-turn-helix domain-containing protein - ATP-dependent Clp protease proteolytic subunit - ClpP family protease - IS607 family element RNA-guided endonuclease TnpB - IS607 family transposase - MerR family transcriptional regulator - MerR family transcriptional regulator. ABC transporter permease subunit - ATP-binding cassette domain-containing protein - hypothetical protein - 2OG-Fe(II) oxygenase family protein - hypothetical protein - aromatic ring-hydroxylating oxygenase subunit alpha - transposase - hypothetical protein - hypothetical protein - type I restriction-modification system subunit M - restriction endonuclease subunit S - type I restriction endonuclease subunit R - biotin/lipoyl-binding protein - DUF5666 domain-containing protein. acetamidase/formamidase family protein - thiamine-phosphate kinase - multifunctional oxoglutarate decarboxylase/oxoglutarate dehydrogenase thiamine pyrophosphate-binding subunit/dihydrolipoyllysine-residue succinyltransferase subunit - DUF6104 family protein - zinc-binding dehydrogenase - hypothetical protein - NAD(P)-dependent malic enzyme - dTDP-4-dehydrorhamnose 3,5-epimerase family protein - GNAT family N-acetyltransferase - anti-sigma factor - SigB/SigF/SigG family RNA polymerase sigma factor - hypothetical protein - diacylglycerol/lipid kinase family protein - glycerophosphodiester phosphodiesterase. acetyl-CoA acetyltransferase - class I adenylate-forming enzyme family protein - LuxR C-terminal-related transcriptional regulator - acyl-CoA dehydrogenase family protein - TetR/AcrR family transcriptional regulator - acyltransferase family protein - LysR family transcriptional regulator - MBL fold metallo-hydrolase - ABC transporter substrate-binding protein - carbohydrate ABC transporter permease - carbohydrate ABC transporter permease - ABC transporter ATP-binding protein - HAD-IA family hydrolase - AAA family ATPase. acetyl/propionyl/methylcrotonyl-CoA carboxylase subunit alpha - hypothetical protein - transposase domain-containing protein - dienelactone hydrolase family protein - transposase - LacI family DNA-binding transcriptional regulator - isocitrate lyase/PEP mutase family protein - TM0106 family RecB-like putative nuclease - LuxR C-terminal-related transcriptional regulator - MMPL family transporter - ChaB family protein - N-acetylmuramoyl-L-alanine amidase - class I SAM-dependent methyltransferase - FadR/GntR family transcriptional regulator. acyl-CoA carboxylase subunit epsilon - acyl-CoA carboxylase subunit beta - biotin--[acetyl-CoA-carboxylase] ligase - PH domain-containing protein - hypothetical protein - hypothetical protein - hypothetical protein - hypothetical protein - TetR/AcrR family transcriptional regulator - GtrA family protein - beta-class carbonic anhydrase - UDP-glucose dehydrogenase family protein - TIGR03089 family protein - LCP family protein. acyl-CoA dehydrogenase family protein - helix-turn-helix transcriptional regulator - tetratricopeptide repeat protein - hypothetical protein - hypothetical protein - HNH endonuclease - hypothetical protein - helix-turn-helix domain-containing protein - bifunctional DNA primase/polymerase - excisionase family DNA-binding protein - tyrosine-type recombinase/integrase - tRNA-Gly - tRNA-Pro - trigger factor. acyltransferase family protein - 30S ribosomal protein S1 - Trm112 family protein - glutathione peroxidase - glycoside hydrolase family 26 protein - hypothetical protein - hypothetical protein - choline/carnitine O-acyltransferase - hypothetical protein - HalD/BesD family halogenase - hemerythrin domain-containing protein - hypothetical protein - hypothetical protein - hydrogenase expression/formation protein HypE. adenylate kinase - glutamate synthase subunit beta - glutamate synthase large subunit - prolipoprotein diacylglyceryl transferase - tryptophan synthase subunit alpha - tryptophan synthase subunit beta - indole-3-glycerol phosphate synthase TrpC - hypothetical protein - Trp biosynthesis-associated membrane protein - anthranilate synthase component I - phosphoribosyl-AMP cyclohydrolase - hypothetical protein - imidazole glycerol phosphate synthase subunit HisF - bifunctional 1-(5-phosphoribosyl)-5-((5-phosphoribosylamino)methylideneamino)imidazole-4-

carboxamide isomerase/phosphoribosylanthranilate isomerase PriA. aldo/keto reductase - HAD family hydrolase - 3-oxoacyl-[acyl-carrier-protein] synthase III C-terminal domain-containing protein - hypothetical protein - hypothetical protein - recombinase family protein - hypothetical protein - hypothetical protein - albusnodin family lasso peptide - albusnodin/ikarugamycin family macrolactam cyclase - lasso peptide biosynthesis B2 protein - GNAT family N-acetyltransferase - replication initiator - recombinase family protein. alpha/beta fold hydrolase - tRNA (adenosine(37)-N6)-threonylcarbamoyltransferase complex ATPase subunit type 1 TsaE - tRNA (adenosine(37)-N6)-threonylcarbamoyltransferase complex dimerization subunit type 1 TsaB - ribosomal protein S18-alanine N-acetyltransferase - FMN-dependent NADH-azoreductase - MarR family winged helix-turn-helix transcriptional regulator - hypothetical protein - co-chaperone GroES - chaperonin GroEL - WhiB family transcriptional regulator - DUF5319 domain-containing protein - IMP dehydrogenase - GuaB3 family IMP dehydrogenase-related protein - hypothetical protein. amidohydrolase family protein - methionine adenosyltransferase - DUF309 domain-containing protein - toll/interleukin-1 receptor domain-containing protein - TAXI family TRAP transporter solute-binding subunit - thiolase family protein - DUF4232 domain-containing protein - hemolysin family protein - hemolysin family protein - class I SAM-dependent methyltransferase - WD40 repeat domain-containing serine/threonine protein kinase - Hsp70 family protein - methylmalonyl Co-A mutase-associated GTPase MeaB - methylmalonyl-CoA mutase. amino acid ABC transporter permease - ribonuclease Y - recombination regulator RecX - recombinase RecA - DUF3046 domain-containing protein - ATP-dependent helicase - RNA polymerase sigma factor - lantibiotic dehydratase - thiopeptide-type bacteriocin biosynthesis protein - lanthionine synthetase LanC family protein - protein-L-isoaspartate O-methyltransferase - tyrosine-type recombinase/integrase - hypothetical protein - hypothetical protein. AMP-binding protein - TauD/TfdA family dioxygenase - dehydrogenase - NUDIX hydrolase - AAA family ATPase - hypothetical protein - radical SAM protein - 4a-hydroxytetrahydrobiopterin dehydratase - flavoprotein - hypothetical protein - methyltransferase, FxLD system - FxLD family lanthipeptide - lantibiotic dehydratase - lanthionine synthetase C family protein. asparagine synthase-related protein - lasso RiPP family leader peptide-containing protein - AfsR/SARP family transcriptional regulator - peptidoglycan-binding protein - alpha/beta hydrolase - four-helix bundle copper-binding protein - amidohydrolase family protein - methionine adenosyltransferase - DUF309 domain-containing protein - toll/interleukin-1 receptor domain-containing protein - TAXI family TRAP transporter solute-binding subunit - thiolase family protein - DUF4232 domain-containing protein - hemolysin family protein. ATP-dependent helicase - ATP-dependent helicase - SDR family NAD(P)-dependent oxidoreductase - MGMT family protein - adenylyltransferase/sulfurtransferase MoeZ - CsbD family protein - Ku protein - hypothetical protein - TetR/AcrR family transcriptional regulator - DUF3107 domain-containing protein - ferritin-like fold-containing protein - DEAD/DEAH box helicase - hypothetical protein - ParA family protein. ATP-grasp domain-containing protein - cation:proton antiporter - alpha/beta fold hydrolase - lipocalin-like domain-containing protein - crotonyl-CoA carboxylase/reductase - 3-hydroxyacyl-CoA dehydrogenase family protein - hypothetical protein - group II intron maturase-specific domain-containing protein - flavodoxin family protein - SAM-dependent methyltransferase - putative bifunctional diguanylate cyclase/phosphodiesterase - hypothetical protein - GvpL/GvpF family gas vesicle protein - gas vesicle structural protein GvpA. bacterial proteasome activator family protein - NAD(P)H-quinone oxidoreductase - SigE family RNA polymerase sigma factor - hypothetical protein - hypothetical protein - DUF3817 domain-containing protein - hypothetical protein - tRNA-Ser - ATP-binding protein - roadblock/LC7 domain-containing protein - DUF742 domain-containing protein - GTP-binding protein - cytochrome P450 - terpene synthase family protein. beta-class carbonic anhydrase - Tex family protein - sensor histidine kinase - SRPBCC family protein - DUF3618 domain-containing protein - phage holin family protein - alpha/beta fold hydrolase - hypothetical protein - DegT/DnrJ/EryC1/StrS family aminotransferase - NAD-dependent epimerase/dehydratase family protein - MGDG synthase family glycosyltransferase - DMT family transporter - hypothetical protein - DUF397 domain-containing protein. beta-ketoacyl-ACP synthase III - acyl carrier protein - beta-ketoacyl-[acyl-carrier-protein] synthase family protein - aldo/keto reductase - HAD family hydrolase - 3-oxoacyl-[acyl-carrier-protein] synthase III C-terminal domain-containing protein - hypothetical protein - hypothetical protein - recombinase family protein - hypothetical protein - hypothetical protein - albusnodin family lasso peptide - albusnodin/ikarugamycin family macrolactam cyclase - lasso peptide biosynthesis B2 protein. beta-ketoacyl-[acyl-carrier-protein] synthase family protein - aldo/keto reductase - HAD family hydrolase - 3-oxoacyl-[acyl-carrier-protein] synthase III C-terminal domain-containing protein - hypothetical protein - hypothetical protein - recombinase family protein - hypothetical protein - hypothetical protein - albusnodin family lasso peptide - albusnodin/ikarugamycin family macrolactam cyclase - lasso peptide biosynthesis B2 protein - GNAT family N-acetyltransferase - replication initiator. C40 family peptidase - metal-dependent transcriptional regulator - DUF3039 domain-containing protein - tRNA-Arg - SulP family inorganic anion transporter - ArsR/SmtB family transcription factor - hypothetical protein - type II 3-dehydroquinate dehydratase - DegT/DnrJ/EryC1/StrS family aminotransferase - Gfo/Idh/MocA family protein - HAD-IA family hydrolase - ROK family protein - shikimate dehydrogenase family protein - 3-dehydroquinate synthase family protein. CapA family protein - Fur family transcriptional regulator - hypothetical protein - hypothetical protein - hypothetical protein - pyridoxal phosphate-dependent aminotransferase - hypothetical protein - type II toxin-antitoxin system VapB family antitoxin - hypothetical protein - HNH endonuclease - hypothetical protein - S1C family serine protease - large conductance mechanosensitive channel protein MscL - hypothetical protein. carbohydrate ABC transporter permease - ABC transporter ATP-binding protein - HAD-IA family hydrolase - AAA family ATPase - response regulator - SAM-dependent methyltransferase - DEAD/DEAH box helicase - acyl-CoA desaturase - hypothetical protein - hypothetical protein - recombinase family protein - hypothetical protein - DUF397 domain-containing protein - DUF5753 domain-containing protein. cation diffusion facilitator family transporter - hypothetical protein - helix-turn-helix domain-containing protein - enoyl-CoA hydratase/isomerase family protein - ABC transporter ATP-binding protein - hypothetical protein - pyruvate kinase - hypothetical protein - hypothetical protein - response regulator transcription factor - type II toxin-antitoxin system VapC family toxin - antitoxin - aminotransferase class V-fold PLP-dependent enzyme - recombinase family protein. CBASS oligonucleotide cyclase - helix-turn-helix domain-containing protein - ImmA/IrrE family metallo-endopeptidase - hypothetical protein - IS110 family transposase - NUDIX hydrolase - hypothetical protein - hypothetical protein - hypothetical protein - hypothetical protein - hypothetical protein - hypothetical protein -

hypothetical protein - FtsK/SpoIIIE domain-containing protein. class I SAM-dependent DNA methyltransferase - pyridoxal phosphate-dependent decarboxylase family protein - ISAs1 family transposase - hypothetical protein - nucleoside deaminase - replication-relaxation family protein - hypothetical protein - ArsR/SmtB family transcription factor - cation transporter - ATP-binding protein - hypothetical protein - recombinase family protein - hypothetical protein - SAM-dependent methyltransferase. class I SAM-dependent methyltransferase - glycosyltransferase family 2 protein - class E sortase - putative bifunctional diguanylate cyclase/phosphodiesterase - Gfo/Idh/MocA family oxidoreductase - protein-tyrosine-phosphatase - TylF/MycF family methyltransferase - glycosyltransferase family 4 protein - UDP-glucuronic acid decarboxylase family protein - methyltransferase FkbM - hypothetical protein - FkbM family methyltransferase - glycosyltransferase - FkbM family methyltransferase. CRISPR-associated helicase Cas3' - type I-C CRISPR-associated protein Cas5c - type I-C CRISPR-associated protein Cas8c/Csd1 - type I-C CRISPR-associated protein Cas7/Csd2 - CRISPR-associated protein Cas4 - type I-C CRISPR-associated endonuclease Cas1c - CRISPR-associated endonuclease Cas2 - site-specific integrase - Uma2 family endonuclease - alcohol dehydrogenase catalytic domain-containing protein - uracil-DNA glycosylase - benzoate/H(+) symporter BenE family transporter - hypothetical protein - epoxide hydrolase family protein. cyclase family protein - gephyrin-like molybdotransferase receptor GlpR - GNAT family N-acetyltransferase - MogA/MoaB family molybdenum cofactor biosynthesis protein - cyclic pyranopterin monophosphate synthase MoaC - molybdotransferase-like divisome protein Glp - UTP--glucose-1-phosphate uridylyltransferase - 5-formyltetrahydrofolate cyclo-ligase - FmdB family zinc ribbon protein - hypothetical protein - hypothetical protein - hypothetical protein - metal-sensitive transcriptional regulator - alpha-ketoglutarate-dependent dioxygenase AlkB. cytochrome bc1 complex cytochrome b subunit - cytochrome bc1 complex Rieske iron-sulfur subunit - hypothetical protein - aa3-type cytochrome oxidase subunit IV - aa3-type cytochrome oxidase subunit I - glycoside hydrolase family 15 protein - thiamine pyrophosphate-requiring protein - hopanoid C-3 methylase HpnR - cytochrome c oxidase assembly protein - Dyp-type peroxidase - SCO family protein - MarR family winged helix-turn-helix transcriptional regulator - chloride channel protein - acetyl-CoA acetyltransferase. cytochrome P450 - RNA-guided endonuclease InsQ/TnpB family protein - IS630 family transposase - hypothetical protein - MurR/RpiR family transcriptional regulator - hypothetical protein - hypothetical protein - hypothetical protein - L,D-transpeptidase - DUF4386 domain-containing protein - TetR/AcrR family transcriptional regulator C-terminal domain-containing protein - non-ribosomal peptide synthetase - hypothetical protein - type I polyketide synthase. DapH/DapD/GlmU-related protein - alpha/beta hydrolase family protein - IS5/IS1182 family transposase - ATP-binding protein - SCO6880 family protein - hypothetical protein - hypothetical protein - BTAD domain-containing putative transcriptional regulator - pilus assembly protein TadG-related protein - TadE/TadG family type IV pilus assembly protein - hypothetical protein - hypothetical protein - type II secretion system F family protein - type II secretion system F family protein. DHH family phosphoesterase - 30S ribosome-binding factor RbfA - DUF503 domain-containing protein - translation initiation factor IF-2 - YlxR family protein - transcription termination factor NusA - ribosome maturation factor RimP - hypothetical protein - DUF4439 domain-containing protein - RrF2 family transcriptional regulator - ABC transporter permease - ABC transporter ATP-binding protein - ABC transporter substrate-binding protein - GNAT family N-acetyltransferase. diguanylate cyclase domain-containing protein - isovaleryl-CoA dehydrogenase - hypothetical protein - hypothetical protein - cystathionine gamma-synthase - cystathionine beta-synthase - YccF domain-containing protein - diguanylate cyclase domain-containing protein - ABC transporter ATP-binding protein - ABC transporter ATP-binding protein - ABC transporter substrate-binding protein - ABC transporter permease subunit - ATP-binding cassette domain-containing protein - hypothetical protein. DivIVA domain-containing protein - cysteine--1-D-myo-inosityl 2-amino-2-deoxy-alpha-D-glucopyranoside ligase - SCO1664 family protein - hypothetical protein - DUF3090 domain-containing protein - histidine phosphatase family protein - ATP-dependent DNA helicase - glycosyl transferase - NAD(P)/FAD-dependent oxidoreductase - glycosyltransferase family 39 protein - hypothetical protein - DUF5703 family protein - hypothetical protein - M20/M25/M40 family metallo-hydrolase. DNA-binding protein - hypothetical protein - recombinase family protein - ATP-binding protein - hypothetical protein - B12-binding domain-containing radical SAM protein - hypothetical protein - hypothetical protein - flavodoxin family protein - DMT family transporter - hypothetical protein - hypothetical protein - DMT family transporter - FAD-dependent oxidoreductase. Dps family protein - helix-turn-helix domain-containing protein - competence/damage-inducible protein A - CDP-alcohol phosphatidyltransferase family protein - 30S ribosomal protein S12 methylthiotransferase RimO - helix-turn-helix domain-containing protein - FtsK/SpoIIIE family DNA translocase - ribonuclease J - 4-hydroxy-tetrahydrodipicolinate synthase - acetate/propionate family kinase - phosphoketolase family protein - FAD-dependent thymidylate synthase - histidine--tRNA ligase - UDP-glucose dehydrogenase family protein. DUF1918 domain-containing protein - hypothetical protein - DoxX family protein - UvrD-helicase domain-containing protein - DUF6342 family protein - IS607 family transposase - zinc ribbon domain-containing protein - tRNA-Ala - ParA family protein - maleylpyruvate isomerase family mycothiol-dependent enzyme - hypothetical protein - DLW-39 family protein - nuclease-related domain-containing protein - ergothioneine biosynthesis glutamate--cysteine ligase EgtA. DUF2784 domain-containing protein - lipoyl(octanoyl) transferase LipB - serine/threonine-protein kinase - alpha-ketoglutarate-dependent dioxygenase AlkB - ATP-binding protein - sensor domain-containing phosphodiesterase - universal stress protein - hypothetical protein - hemerythrin domain-containing protein - DUF6328 family protein - hypothetical protein - deaminase - tyrosine-type recombinase/integrase - hypothetical protein. DUF3180 domain-containing protein - 2-amino-4-hydroxy-6-hydroxymethyldihydropteridine diphosphokinase - dihydroneopterin aldolase - dihydropteroate synthase - GNAT family N-acetyltransferase - hypothetical protein - aminoglycoside phosphotransferase family protein - MinD/ParA family ATP-binding protein - geranylgeranyl reductase family protein - DUF5926 family protein - NAD(P)/FAD-dependent oxidoreductase - ATP-binding protein - phosphoribosylformylglycinamidine synthase subunit PurQ - phosphoribosylformylglycinamidine synthase subunit PurL. DUF397 domain-containing protein - hypothetical protein - hypothetical protein - GNAT family N-acetyltransferase - tyrosine-type recombinase/integrase - tRNA-Val - peroxiredoxin - DUF4234 domain-containing protein - DUF3052 domain-containing protein - secondary thiamine-phosphate synthase enzyme YjbQ - pyruvate dehydrogenase (acetyl-transferring), homodimeric type - hypothetical protein - bifunctional phosphatase PAP2/diacylglycerol kinase family protein - DUF5999 family protein. DUF5318 family protein - PadR family transcriptional regulator - inositol-3-phosphate synthase - MFS transporter - CCA tRNA nucleotidyltransferase - hypothetical protein - DUF6049 family protein - murein biosynthesis integral membrane protein MurJ - protein kinase family protein - hypothetical protein - hypothetical protein - thioredoxin-disulfide reductase - thioredoxin - N-acetylmuramoyl-L-alanine amidase. DUF6049 family protein - murein biosynthesis integral membrane protein MurJ - protein kinase family protein - hypothetical protein - hypothetical protein - thioredoxin-disulfide reductase - thioredoxin - N-acetylmuramoyl-L-alanine amidase - GNAT family N-acetyltransferase - ParB/RepB/Spo0J family partition protein - ParA family protein - 16S rRNA (guanine(527)-N(7))-methyltransferase RsmG - hypothetical protein - protein jag. DUF881 domain-containing protein - small basic family protein - DUF881 domain-containing protein - glycine cleavage system protein GcvH - FHA domain-containing protein - transcriptional regulator FtsR - bifunctional nuclease family protein - MerR family transcriptional regulator - daunorubicin resistance protein DrrA family ABC transporter ATP-binding protein - ABC transporter permease - hypothetical protein - FtsK/SpoIIIE domain-containing protein - hypothetical protein - hypothetical protein. enoyl-CoA hydratase-related protein - acyl-CoA dehydrogenase family protein - helix-turn-helix transcriptional regulator - tetratricopeptide repeat protein - hypothetical protein - hypothetical protein - HNH endonuclease - hypothetical protein - helix-turn-helix domain-containing protein - bifunctional DNA primase/polymerase - excisionase family DNA-binding protein - tyrosine-type recombinase/integrase - tRNA-Gly - tRNA-Pro. excisionase family DNA-binding protein - tyrosine-type recombinase/integrase - tRNA-Gly - tRNA-Pro - trigger factor - ATP-dependent Clp protease proteolytic subunit - ATP-dependent Clp protease proteolytic subunit - ATP-dependent Clp protease ATP-binding subunit ClpX - bifunctional folylpolyglutamate synthase/dihydrofolate synthase - dihydrofolate synthase - nucleoside-diphosphate kinase - rod shape-determining protein - rod shape-determining protein MreC - rod shape-determining protein MreD. FAD-binding oxidoreductase - aromatic/alkene/methane monooxygenase hydroxylase/oxygenase subunit alpha - GAF domain-containing protein - IS66 family transposase - helix-turn-helix domain-containing protein - diguanylate cyclase - hypothetical protein - hypothetical protein - HhH-GPD-type base excision DNA repair protein - MarR family transcriptional regulator - hypothetical protein - NAD-dependent epimerase/dehydratase family protein - RNA polymerase sigma factor SigJ - Fe-S cluster assembly protein HesB. FkbM family methyltransferase - aminomethyl-transferring glycine dehydrogenase subunit GcvPB - hypothetical protein - ISAs1 family transposase - hypothetical protein - IS607 family transposase - PIG-L deacetylase family protein - hypothetical protein - AAA family ATPase - hypothetical protein - hypothetical protein - ISKra4 family transposase - FUSC family protein - hypothetical protein. fluoride efflux transporter FluC - arsenate reductase ArsC - aquaporin - ArsI/CadI family heavy metal resistance metalloenzyme - alpha/beta hydrolase - ISAs1 family transposase - alpha/beta fold hydrolase - hypothetical protein - hypothetical protein - hypothetical protein - type II toxin-antitoxin system PemK/MazF family toxin - hypothetical protein - acyltransferase family protein - AMP-binding protein. fumarylacetoacetate hydrolase family protein - hypothetical protein - citramalate synthase - Crp/Fnr family transcriptional regulator - CBS domain-containing protein - branched-chain-amino-acid transaminase - 3-isopropylmalate dehydrogenase - EAL domain-containing protein - DUF1059 domain-containing protein - 2-

isopropylmalate synthase - septum formation family protein - phosphoglycerate dehydrogenase - ketol-acid reductoisomerase - acetolactate synthase small subunit. GH25 family lysozyme - tRNA-Thr - tRNA-Met - 50S ribosomal protein L33 - FAS1-like dehydratase domain-containing protein - MaoC family dehydratase - pyridoxal phosphate-dependent aminotransferase - tRNA-Trp - preprotein translocase subunit SecE - transcription termination/antitermination protein NusG - 50S ribosomal protein L11 - 50S ribosomal protein L1 - 50S ribosomal protein L10 - 50S ribosomal protein L7/L12. GlsB/YeaQ/YmgE family stress response membrane protein - DUF2599 domain-containing protein - TIGR03086 family metal-binding protein - TetR/AcrR family transcriptional regulator - formylglycine-generating enzyme family protein - hypothetical protein - methylenetetrahydrofolate reductase - hypothetical protein - Tn3 family transposase - DinB family protein - IS5 family transposase - transposase - group II intron reverse transcriptase/maturase - protein-tyrosine phosphatase family protein. glutathione peroxidase - glycoside hydrolase family 26 protein - hypothetical protein - hypothetical protein - choline/carnitine O-acyltransferase - hypothetical protein - HalD/BesD family halogenase - hemerythrin domain-containing protein - hypothetical protein - hypothetical protein - hydrogenase expression/formation protein HypE - hydrogenase formation protein HypD - HypC/HybG/HupF family hydrogenase formation chaperone - carbamoyltransferase HypF. glycosyltransferase family 2 protein - class E sortase - putative bifunctional diguanylate cyclase/phosphodiesterase - Gfo/Idh/MocA family oxidoreductase - protein-tyrosine-phosphatase - TylF/MycF family methyltransferase - glycosyltransferase family 4 protein - UDP-glucuronic acid decarboxylase family protein - methyltransferase FkbM - hypothetical protein - FkbM family methyltransferase - glycosyltransferase - FkbM family methyltransferase - heparinase II/III family protein. GNAT family N-acetyltransferase - hypothetical protein - transposase domain-containing protein - type II toxin-antitoxin system HicA family toxin - type II toxin-antitoxin system HicB family antitoxin - recombinase family protein - ISAs1 family transposase - Fic family protein - hypothetical protein - SCO4848 family membrane protein - SDR family NAD(P)-dependent oxidoreductase - RecQ family ATP-dependent DNA helicase - succinate dehydrogenase iron-sulfur subunit - succinate dehydrogenase flavoprotein subunit. group II intron maturase-specific domain-containing protein - ISAs1 family transposase - transposase - transposase family protein - IS5 family transposase - IS701 family transposase - hemerythrin domain-containing protein - family 2B encapsulin nanocompartment shell protein - family 2 encapsulin nanocompartment cargo protein terpene cyclase - SagB/ThcOx family dehydrogenase - YcaO-like family protein - YcaO-like family protein - CocE/NonD family hydrolase - TOMM precursor leader peptide-binding protein. HAD-IIA family hydrolase - hypothetical protein - hypothetical protein - hypothetical protein - recombinase family protein - peptide-methionine (S)-S-oxide reductase MsrA - PhoH family protein - phosphoenolpyruvate carboxykinase (GTP) - PAQR family membrane homeostasis protein TrhA - hypothetical protein - thioredoxin domain-containing protein - leucine--tRNA ligase - hypothetical protein - mycothiol conjugate amidase Mca. helix-turn-helix domain-containing protein - bifunctional DNA primase/polymerase - excisionase family DNA-binding protein - tyrosine-type recombinase/integrase - tRNA-Gly - tRNA-Pro - trigger factor - ATP-dependent Clp protease proteolytic subunit - ATP-dependent Clp protease proteolytic subunit - ATP-dependent Clp protease ATP-binding subunit ClpX - bifunctional folylpolyglutamate synthase/dihydrofolate synthase - dihydrofolate synthase - nucleoside-diphosphate kinase - rod shape-determining protein. helix-turn-helix transcriptional regulator - FKBP-type peptidyl-prolyl cis-trans isomerase - Pup--protein ligase - hypothetical protein - dienelactone hydrolase family protein - cation-translocating P-type ATPase - TMEM165/GDT1 family protein - hypothetical protein - type II toxin-antitoxin system VapC family toxin - hypothetical protein - GNAT family N-acetyltransferase - hypothetical protein - ATP-binding protein - ATP-binding cassette domain-containing protein. helix-turn-helix transcriptional regulator - VOC family protein - amidohydrolase family protein - glutamine synthetase family protein - saccharopine dehydrogenase NADP-binding domain-containing protein - pyruvate carboxylase - bifunctional glycosyltransferase family 2/GtrA family protein - ArnT family glycosyltransferase - sensor histidine kinase - response regulator transcription factor - hypothetical protein - Hsp70 family protein - hypothetical protein - response regulator. HepT-like ribonuclease domain-containing protein - transposase - ISKra4 family transposase - hypothetical protein - group II intron reverse transcriptase/maturase - hypothetical protein - epoxide hydrolase family protein - isocitrate lyase/PEP mutase family protein - hypothetical protein - NADP-dependent isocitrate dehydrogenase - DUF2142 domain-containing protein - hypothetical protein - N,N-dimethylformamidase beta subunit family domain-containing protein - winged helix-turn-helix transcriptional regulator. HRDC domain-containing protein - DUF3000 domain-containing protein - hypothetical protein - DUF3817 domain-containing protein - uroporphyrinogen decarboxylase - protoporphyrinogen oxidase - hydrogen peroxide-dependent heme synthase - sugar transferase - adenylyl-sulfate kinase - 3'(2'),5'-bisphosphate nucleotidase CysQ - sulfate adenylyltransferase subunit CysD - sulfate adenylyltransferase subunit 1 - hypothetical protein - hypothetical protein. hydroxymethylbilane synthase - uroporphyrinogen-III synthase - porphobilinogen synthase - hypothetical protein - heavy-metal-associated domain-containing protein - heavy metal translocating P-type ATPase - methionine--tRNA ligase - TnsA-like heteromeric transposase endonuclease subunit - hypothetical protein - Mu transposase C-terminal domain-containing protein - ATP-binding protein - TniQ family protein - hypothetical protein - hypothetical protein. hypothetical protein - bifunctional adenosylcobinamide kinase/adenosylcobinamide-phosphate guanylyltransferase - hypothetical protein - nicotinate-nucleotide--dimethylbenzimidazole phosphoribosyltransferase - adenosylcobinamide-GDP ribazoletransferase - glycine cleavage system aminomethyltransferase GcvT - leucyl aminopeptidase - dihydrolipoyl dehydrogenase - 2-oxoglutarate dehydrogenase, E2 component, dihydrolipoamide succinyltransferase - protein kinase - peptidase E - lipoyl synthase - hypothetical protein - DUF4191 domain-containing protein. hypothetical protein - class I SAM-dependent methyltransferase - dTDP-4-dehydrorhamnose 3,5-epimerase family protein - NAD-dependent epimerase/dehydratase family protein - serine O-acetyltransferase - acyl carrier protein - HAD-IIIC family phosphatase - S8 family serine peptidase - tyrosine--tRNA ligase - glycosyltransferase - glycosyltransferase family 2 protein - glycosyltransferase - PIG-L deacetylase family protein - aminoacyl-tRNA hydrolase. hypothetical protein - DDE-type integrase/transposase/recombinase - DHA2 family efflux MFS transporter permease subunit - winged helix-turn-helix transcriptional regulator - putative quinol monooxygenase - FAD-binding and (Fe-S)-binding domain-containing protein - GntR family transcriptional regulator - aa3-type cytochrome oxidase subunit I - LGFP repeat-containing protein - IclR family transcriptional regulator - PEP/pyruvate-binding domain-containing protein - DHA2 family efflux MFS transporter permease subunit - MarR family winged helix-turn-helix transcriptional regulator - aminotransferase class I/II-fold pyridoxal phosphate-dependent enzyme. hypothetical protein - DUF4277 domain-containing protein - class II 3-deoxy-7-phosphoheptulonate synthase - 6-phosphofructokinase - polyadenylate-specific 3'-exoribonuclease AS - hypothetical protein - lysophospholipid acyltransferase family protein - alpha/beta hydrolase - hypothetical protein - hypothetical protein - ROK family glucokinase - hypothetical protein - ArsA family ATPase - SRPBCC family protein. hypothetical protein - DUF5130 family protein - aminopeptidase N - AI-2E family transporter - hypothetical protein - ribose-5-phosphate isomerase - hypothetical protein - dolichyl-phosphate beta-glucosyltransferase - DUF3367 domain-containing protein - RyR domain-containing protein - glycosyltransferase family 4 protein - hypothetical protein - hypothetical protein - GNAT family N-acetyltransferase. hypothetical protein - DUF6624 domain-containing protein - AMP-binding protein - hypothetical protein - AMP-binding protein - TauD/TfdA family dioxygenase - dehydrogenase - NUDIX hydrolase - AAA family ATPase - hypothetical protein - radical SAM protein - 4a-hydroxytetrahydrobiopterin dehydratase - flavoprotein - hypothetical protein. hypothetical protein - endonuclease/exonuclease/phosphatase family protein - enoyl-CoA hydratase/isomerase family protein - hypothetical protein - acetate--CoA ligase - alpha-glucan family phosphorylase - glycogen debranching protein GlgX - hypothetical protein - hypothetical protein - cysteine protease StiP family protein - phosphoribosyltransferase family protein - HpcH/HpaI aldolase/citrate lyase family protein - glycoside hydrolase family 26 protein - hypothetical protein. hypothetical protein - GvpL/GvpF family gas vesicle protein - gas vesicle structural protein GvpA - GvpL/GvpF family gas vesicle protein - cyclase - MOSC domain-containing protein - arsenate reductase ArsC - acyl-CoA dehydrogenase family protein - alkaline phosphatase D family protein - formylglycine-generating enzyme family protein - hypothetical protein - hypothetical protein - hypothetical protein - tyrosine-type recombinase/integrase. hypothetical protein - helix-turn-helix transcriptional regulator - MFS transporter - TetR/AcrR family transcriptional regulator - TM2 domain-containing protein - hypothetical protein - hypothetical protein - amidohydrolase family protein - helix-turn-helix domain-containing protein - MFS transporter - prolipoprotein diacylglyceryl transferase - MarR family winged helix-turn-helix transcriptional regulator - helix-turn-helix domain-containing protein - hypothetical protein. hypothetical protein - hypothetical protein - ATP-binding protein - GNAT family N-acetyltransferase - DUF4037 domain-containing protein - helix-turn-helix domain-containing protein - PIN domain-containing protein - hypothetical protein - hypothetical protein - cytochrome P450 - 2-dehydropantoate 2-reductase - histidinol-phosphate transaminase - RiPP maturation radical SAM C-methyltransferase - IS66 family transposase. hypothetical protein - hypothetical protein - DUF3180 domain-containing protein - 2-amino-4-hydroxy-6-hydroxymethyldihydropteridine diphosphokinase - dihydroneopterin aldolase - dihydropteroate synthase - GNAT family N-acetyltransferase - hypothetical protein - aminoglycoside phosphotransferase family protein - MinD/ParA family ATP-binding protein - geranylgeranyl reductase family protein - DUF5926 family protein - NAD(P)/FAD-dependent oxidoreductase - ATP-binding protein. hypothetical protein - hypothetical protein - hypothetical protein - cytochrome P450 - OsmC family protein - thiopeptide-type bacteriocin biosynthesis protein - methyltransferase, FxLD system - cold-shock protein - response regulator - carboxymuconolactone decarboxylase family protein - ferredoxin - 2-oxoacid:ferredoxin oxidoreductase subunit beta - 2-oxoacid:acceptor oxidoreductase subunit alpha - cysteine desulfurase family protein. hypothetical protein - hypothetical protein - hypothetical protein - GroES family chaperonin - DUF4193 family protein - LuxR family transcriptional regulator - hypothetical protein - helix-turn-helix transcriptional regulator - type I polyketide synthase - type I polyketide synthase - acyl carrier protein - alpha/beta fold hydrolase - glycosyltransferase - class I SAM-dependent methyltransferase. hypothetical protein - hypothetical protein - hypothetical protein - hypothetical protein - hypothetical protein - hypothetical protein - hypothetical protein - FtsK/SpoIIIE domain-containing protein - hypothetical protein - hypothetical protein - type II toxin-antitoxin system VapC family toxin - type II toxin-antitoxin system Phd/YefM family antitoxin - IS5/IS1182 family transposase - IS66 family transposase. hypothetical protein - hypothetical protein - hypothetical protein - hypothetical protein - transposase - IS3 family transposase - IS701 family transposase - winged helix-turn-helix transcriptional regulator - BTAD domain-containing putative transcriptional regulator - pilus assembly protein TadG-related protein - TadE/TadG family type IV pilus assembly protein - TadE family protein - hypothetical protein - type II secretion system F family protein. hypothetical protein - hypothetical protein - phosphoenolpyruvate phosphomutase - hypothetical protein - class I SAM-dependent methyltransferase - SDR family NAD(P)-dependent oxidoreductase - LacI family DNA-binding transcriptional regulator - AMP-binding protein - HpcH/HpaI aldolase/citrate lyase family protein - acyl-CoA dehydrogenase family protein - MaoC family dehydratase - protein meaA - methylmalonyl Co-A mutase-associated GTPase MeaB - DnaJ C-terminal domain-containing protein. hypothetical protein - hypothetical protein - tyrosine-type recombinase/integrase - IS110 family transposase - IS630 family transposase - hypothetical protein - phosphoribosylglycinamide formyltransferase - 2'-5' RNA ligase family protein - helix-turn-helix domain-containing protein - hypothetical protein - hypothetical protein - helix-turn-helix domain-containing protein - hypothetical protein - hypothetical protein. hypothetical protein - ISAs1 family transposase - hypothetical protein - IS607 family transposase - PIG-L deacetylase family protein - hypothetical protein - AAA family ATPase - hypothetical protein - hypothetical protein - ISKra4 family transposase - FUSC family protein - hypothetical protein - hypothetical protein - SAM-dependent methyltransferase. hypothetical protein - lipid-transfer protein - TetR/AcrR family transcriptional regulator - metal ABC transporter permease - metal ABC transporter permease - metal ABC transporter ATP-binding protein - metal ABC transporter solute-binding protein, Zn/Mn family - SCO5389 family protein - ISAs1 family transposase - hypothetical protein - CapA family protein - WS/DGAT domain-containing protein - prephenate dehydratase - dihydrolipoamide acetyltransferase family protein. hypothetical protein - molybdenum cofactor biosynthesis protein MoaE - zinc-dependent metalloprotease - ThiF family adenylyltransferase - ABC1 kinase family protein - alkaline phosphatase family protein - hypothetical protein - WhiB family transcriptional regulator - hypothetical protein - ATP-dependent DNA helicase UvrD2 - mycoredoxin - GAF and ANTAR domain-containing protein - M16 family metallopeptidase - M16 family metallopeptidase. hypothetical protein - pyridoxal phosphate-dependent aminotransferase - hypothetical protein - type II toxin-antitoxin system VapB family antitoxin - hypothetical protein - HNH endonuclease - hypothetical protein - S1C family serine protease - large conductance mechanosensitive channel protein MscL - hypothetical protein - PHB depolymerase family esterase - hypothetical protein - mechanosensitive ion channel family protein - UdgX family uracil-DNA binding protein. hypothetical protein - transposase - CHAT domain-containing protein - hypothetical protein - hypothetical protein - TIGR03619 family F420-dependent LLM class oxidoreductase - MerR family transcriptional regulator - YncE family protein - polysaccharide deacetylase family protein - polysaccharide deacetylase family protein - SGNH/GDSL hydrolase family protein - RNA polymerase sigma factor SigJ - NAD(P)-dependent oxidoreductase - Kelch repeat-containing protein. hypothetical protein - transposase family protein - ABC transporter ATP-binding protein - ABC transporter permease - ABC transporter substrate-binding protein - TauD/TfdA dioxygenase family protein - ROK family transcriptional regulator - M56 family metallopeptidase - BlaI/MecI/CopY family transcriptional regulator - cytochrome ubiquinol oxidase subunit I - cytochrome d ubiquinol oxidase subunit II - CPBP family intramembrane glutamic endopeptidase - CYTH and CHAD domain-containing protein - cold-shock protein. hypothetical protein - transposase family protein - tyrosine-type recombinase/integrase - hypothetical protein - DUF2127 domain-containing protein - DUF2231 domain-containing protein - LLM class F420-dependent oxidoreductase - putative bifunctional diguanylate cyclase/phosphodiesterase - leucyl/phenylalanyl-tRNA--protein transferase - aminotransferase class I/II-fold pyridoxal phosphate-dependent enzyme - class I SAM-dependent methyltransferase - UvrD-helicase domain-containing protein - AAA domain-containing protein - acetamidase/formamidase family protein. hypothetical protein - type II toxin-antitoxin system VapB family antitoxin - hypothetical protein - HNH endonuclease - hypothetical protein - S1C family serine protease - large conductance mechanosensitive channel protein MscL - hypothetical protein - PHB depolymerase family esterase - hypothetical protein - mechanosensitive ion channel family protein - UdgX family uracil-DNA binding protein - L,D-transpeptidase - hypothetical protein. hypothetical protein - zinc-binding dehydrogenase - DHA2 family efflux MFS transporter permease subunit - ISAs1 family transposase - NADPH-dependent FMN reductase - hypothetical protein - hypothetical protein - helix-turn-helix domain-containing protein - DUF397 domain-containing protein - hypothetical protein - hypothetical protein - hypothetical protein - hypothetical protein - replication initiator. IclR family transcriptional regulator - serine/threonine-protein kinase - sensor histidine kinase - hypothetical protein - tRNA-Glu - tRNA-Gln - fumarylacetoacetate hydrolase family protein - hypothetical protein - citramalate synthase - Crp/Fnr family transcriptional regulator - CBS domain-containing protein - branched-chain-amino-acid transaminase - 3-isopropylmalate dehydrogenase - EAL domain-containing protein. ImmA/IrrE family metallo-endopeptidase - hypothetical protein - IS110 family transposase - NUDIX hydrolase - hypothetical protein - hypothetical protein - hypothetical protein - hypothetical protein - hypothetical protein - hypothetical protein - hypothetical protein - FtsK/SpoIIIE domain-containing protein - hypothetical protein - hypothetical protein. iron ABC transporter substrate-binding protein - AMP-binding protein - DUF308 domain-containing protein - ABC transporter ATP-binding protein - hypothetical protein - hypothetical protein - hypothetical protein - cytochrome P450 - ferredoxin - fibronectin type III domain-containing protein - fatty acyl-AMP ligase - thioesterase II family protein - cytochrome P450 - class I SAM-dependent methyltransferase. IS607 family transposase - protease - TerB family tellurite resistance protein - SCP2 sterol-binding domain-containing protein - aminotransferase class IV family protein - IS110 family transposase - IS110 family transposase - hypothetical protein - toll/interleukin-1 receptor domain-containing protein - cation:proton antiporter - malate dehydrogenase - ABC transporter substrate-binding protein - GlcG/HbpS family heme-binding protein - hypothetical protein. ISAs1 family transposase - alpha/beta fold hydrolase - hypothetical protein - hypothetical protein - hypothetical protein - type II toxin-antitoxin system PemK/MazF family toxin - hypothetical protein - acyltransferase family protein - AMP-binding protein - tryptophan 7-halogenase - RBBP9/YdeN family alpha/beta hydrolase - hypothetical protein - transposase - NUDIX hydrolase. L-aspartate oxidase - carboxylating nicotinate-nucleotide diphosphorylase - type III pantothenate kinase - SGNH/GDSL hydrolase family protein - (2Fe-2S)-binding protein - ATP-dependent Clp protease ATP-binding subunit - purple acid phosphatase family protein - hypothetical protein - PP2C family protein-serine/threonine phosphatase - hypothetical protein - polyprenol phosphomannose-dependent alpha 1,6 mannosyltransferase MptB - glycosyltransferase 87 family protein - Glu/Leu/Phe/Val family dehydrogenase - RNHCP domain-containing protein. lasso peptide biosynthesis B2 protein - lasso peptide biosynthesis PqqD family chaperone - asparagine synthase-related protein - lasso RiPP family leader peptide-containing protein - AfsR/SARP family transcriptional regulator - peptidoglycan-binding protein - alpha/beta hydrolase - four-helix bundle copper-binding protein - amidohydrolase family protein - methionine adenosyltransferase - DUF309 domain-containing protein - toll/interleukin-1 receptor domain-containing protein - TAXI family TRAP transporter solute-binding subunit - thiolase family protein. L,D-transpeptidase - phosphatase PAP2 family protein - OmpA family protein - (2Fe-2S)-binding protein - hypothetical protein - hypothetical protein - FadR/GntR family transcriptional regulator - catalase/peroxidase HPI - hypothetical protein - LysR substrate-binding domain-containing protein - Hsp20/alpha crystallin family protein - sulfotransferase - IS66 family transposase - sulfotransferase family protein. LysR family transcriptional regulator - MBL fold metallo-hydrolase - ABC transporter substrate-binding protein - carbohydrate ABC transporter permease - carbohydrate ABC transporter permease - ABC transporter ATP-binding protein - HAD-IA family hydrolase - AAA family ATPase - response regulator - SAM-dependent methyltransferase - DEAD/DEAH box helicase - acyl-CoA desaturase - hypothetical protein - hypothetical protein. malate dehydrogenase - ABC transporter substrate-binding protein - GlcG/HbpS family heme-binding protein - hypothetical protein - DUF397 domain-containing protein - helix-turn-helix domain-containing protein - ATP-binding protein - ATP-dependent RNA helicase HrpA - tetratricopeptide repeat protein - hypothetical protein - galactokinase - UDP-glucose--hexose-1-phosphate uridylyltransferase - redox-sensitive transcriptional activator SoxR - NADPH-dependent FMN reductase. MarR family winged helix-turn-helix transcriptional regulator - aminotransferase class I/II-fold pyridoxal phosphate-dependent enzyme - sulfotransferase family protein - NAD(P)-dependent oxidoreductase - winged helix-turn-helix transcriptional regulator - daunorubicin resistance protein DrrA family ABC transporter ATP-binding protein - ABC transporter permease - BTAD domain-containing putative transcriptional regulator - DUF2786 domain-containing protein - IS5/IS1182 family transposase - IS5 family transposase - heme-binding beta-barrel domain-containing protein - pyruvate:ferredoxin (flavodoxin) oxidoreductase - ParA family protein. MarR family winged helix-turn-helix transcriptional regulator - hypothetical protein - hypothetical protein - sulfatase family protein - Rpn family recombination-promoting nuclease/putative transposase - flavodoxin - prolyl aminopeptidase - nucleotidyltransferase family protein - HepT-like ribonuclease domain-containing protein - transposase - ISKra4 family transposase - hypothetical protein - group II intron reverse transcriptase/maturase - hypothetical protein. MauE/DoxX family redox-associated membrane protein - SCO family protein - DUF3618 domain-containing protein - lysylphosphatidylglycerol synthase transmembrane domain-containing protein - ATP-binding protein - hypothetical protein - hypothetical protein - hypothetical protein - GroES family chaperonin - DUF4193 family protein - LuxR family transcriptional regulator - hypothetical protein - helix-turn-helix transcriptional regulator - type I polyketide synthase. McrB family protein - zeta toxin family protein - DEAD/DEAH box helicase - ATP-dependent 6-phosphofructokinase - hypothetical protein - AAA domain-containing protein - hypothetical protein - antibiotic biosynthesis monooxygenase family protein - antibiotic biosynthesis monooxygenase family protein - SDR family NAD(P)-dependent oxidoreductase - beta-ketoacyl synthase N-terminal-like domain-containing protein - beta-ketoacyl-[acyl-carrier-protein] synthase family protein - cupin domain-containing protein - SRPBCC family protein. methionine--tRNA ligase - TnsA-like heteromeric transposase endonuclease subunit - hypothetical protein - Mu transposase C-terminal domain-containing protein - ATP-binding protein - TniQ family protein - hypothetical protein - hypothetical protein - putative immunity protein - hypothetical protein - class I SAM-dependent DNA methyltransferase - MerR family transcriptional regulator - arginine--tRNA ligase - IS66 family transposase. MFS transporter - ABC transporter substrate-binding protein - ABC transporter ATP-binding protein - FecCD family ABC transporter permease - ABC transporter ATP-binding protein - lipase - peptide deformylase - transglycosylase family protein - 23S rRNA (guanosine(2251)-2'-O)-methyltransferase RlmB - hypothetical protein - PP2C family protein-serine/threonine phosphatase - hypothetical protein - FHA domain-containing protein - cysteine--tRNA ligase. MFS transporter - HpcH/HpaI aldolase/citrate lyase family protein - HpcH/HpaI aldolase/citrate lyase family protein - hypothetical protein - PLP-dependent aminotransferase family protein - SRPBCC family protein - ArsR/SmtB family transcription factor - siderophore-interacting protein - serine/threonine-protein kinase - hypothetical protein - hypothetical protein - glutamate ABC transporter substrate-binding protein - hypothetical protein - hypothetical protein. NAD-dependent protein deacetylase - transcriptional regulator - hypothetical protein - copper resistance CopC family protein - cytochrome c oxidase assembly protein - HAD-IC family P-type ATPase - undecaprenyl-diphosphate phosphatase - helix-turn-helix domain-containing protein - ISAs1 family transposase - hypothetical protein - 2,4'-dihydroxyacetophenone dioxygenase family protein - PucR family transcriptional regulator - MFS transporter - HpcH/HpaI aldolase/citrate lyase family protein. NADH-ubiquinone oxidoreductase-F iron-sulfur binding region domain-containing protein - ferredoxin - alpha/beta hydrolase domain-containing protein - purple acid phosphatase family protein - Dyp-type peroxidase - FAD-dependent monooxygenase - keywimysin-related RiPP - TetR family transcriptional regulator - adenosyl-hopene transferase HpnH - beta-ketoacyl-[acyl-carrier-protein] synthase family protein - SpoIIE family protein phosphatase - cation diffusion facilitator family transporter - ArsR/SmtB family transcription factor - ribose-phosphate diphosphokinase. NAD(P)/FAD-dependent oxidoreductase - serine/threonine-protein kinase - hypothetical protein - cyclic dehypoxanthinyl futalosine synthase - adenosine deaminase - demethylmenaquinone methyltransferase - dienelactone hydrolase family protein - geranylgeranyl reductase family protein - NADH-quinone oxidoreductase subunit A - NuoB/complex I 20 kDa subunit family protein - NADH-quinone oxidoreductase subunit C - NADH-quinone oxidoreductase subunit D - NADH-quinone oxidoreductase subunit NuoE - NADH-quinone oxidoreductase subunit NuoF. nitrogenase iron-molybdenum cofactor biosynthesis protein NifE - nitrogenase molybdenum-iron protein subunit beta - nitrogenase molybdenum-iron protein alpha chain - nitrogenase iron protein - homocitrate synthase - hypothetical protein - hypothetical protein - TOBE domain-containing protein - site-2 protease family protein - hydrogenase maturation protease - Ni/Fe hydrogenase subunit alpha - oxidoreductase - FAD/NAD(P)-binding protein - cyclic nucleotide-binding domain-containing protein. NlpC/P60 family protein - hypothetical protein - transcriptional regulator - hypothetical protein - hypothetical protein - hypothetical protein - hypothetical protein - hypothetical protein - hypothetical protein - hypothetical protein - hypothetical protein - hypothetical protein - hypothetical protein - hypothetical protein. NTP transferase domain-containing protein - GTP 3',8-cyclase MoaA - polyphosphate--glucose phosphotransferase - septation protein SepH - DUF7455 domain-containing protein - YihY/virulence factor BrkB family protein - LCP family protein - aminopeptidase P family protein - histone-like nucleoid-structuring protein Lsr2 - hypothetical protein - ABC transporter ATP-binding protein - SpoIIE family protein phosphatase - bifunctional GNAT family N-acetyltransferase/acetate--CoA ligase family protein - acetoin utilization protein AcuC. nucleoside deaminase - replication-relaxation family protein - hypothetical protein - ArsR/SmtB family transcription factor - cation transporter - ATP-binding protein - hypothetical protein - recombinase family protein - hypothetical protein - SAM-dependent methyltransferase - hypothetical protein - hypothetical protein - hypothetical protein - hypothetical protein. nucleoside triphosphate pyrophosphohydrolase - glycoside hydrolase family 26 protein - sugar phosphate nucleotidyltransferase - DUF4910 domain-containing protein - IspD/TarI family cytidylyltransferase - hypothetical protein - SDR family oxidoreductase - hypothetical protein - Wzz/FepE/Etk N-terminal domain-containing protein - hypothetical protein - class I SAM-dependent methyltransferase - dTDP-4-dehydrorhamnose 3,5-epimerase family protein - NAD-dependent epimerase/dehydratase family protein - serine O-acetyltransferase. oxidoreductase - FAD/NAD(P)-binding protein - cyclic nucleotide-binding domain-containing protein - sulfite reductase subunit A - NUDIX hydrolase - DUF1918 domain-containing protein - hypothetical protein - DoxX family protein - UvrD-helicase domain-containing protein - DUF6342 family protein - IS607 family transposase - zinc ribbon domain-containing protein - tRNA-Ala - ParA family protein. peptide chain release factor 2 - polysaccharide deacetylase family protein - uroporphyrinogen-III synthase - nitrite reductase large subunit NirB - nitrite reductase small subunit NirD - uroporphyrinogen-III C-methyltransferase - hypothetical protein - cell division ATP-binding protein FtsE - permease-like cell division protein FtsX - HNH endonuclease signature motif containing protein - SsrA-binding protein SmpB - THUMP-like domain-containing protein - NAD(P)-dependent oxidoreductase - YoaK family protein. phosphate ABC transporter permease PstA - phosphate ABC transporter permease subunit PstC - phosphate ABC transporter substrate-binding protein PstS - hypothetical protein - proline--tRNA ligase - UMP kinase - ABC transporter substrate-binding protein - GntR family transcriptional regulator - hypothetical protein - hypothetical protein - FtsK/SpoIIIE domain-containing protein - replication initiator - excisionase family DNA-binding protein - site-specific integrase. phosphatidylglycerol lysyltransferase domain-containing protein - ribonuclease BN - GAP family protein - SHOCT domain-containing protein - ParA family protein - hypothetical protein - hypothetical protein -

winged helix-turn-helix transcriptional regulator - MFS transporter - rhodanese-like domain-containing protein - L-tyrosine/L-tryptophan isonitrile synthase family protein - TauD/TfdA dioxygenase family protein - NAD(P)/FAD-dependent oxidoreductase - MFS transporter. phosphopyruvate hydratase - S49 family peptidase - MerR family transcriptional regulator - transcription-repair coupling factor - SurA N-terminal domain-containing protein - nucleoside triphosphate pyrophosphohydrolase - glycoside hydrolase family 26 protein - sugar phosphate nucleotidyltransferase - DUF4910 domain-containing protein - IspD/TarI family cytidylyltransferase - hypothetical protein - SDR family oxidoreductase - hypothetical protein - Wzz/FepE/Etk N-terminal domain-containing protein. phosphotransferase - MFS transporter - class I SAM-dependent methyltransferase - sulfotransferase family protein - B12-binding domain-containing radical SAM protein - ISAs1 family transposase - cobalamin-dependent protein - IS6 family transposase - ATP-binding protein - hypothetical protein - helix-turn-helix domain-containing protein - DUF397 domain-containing protein - hypothetical protein - IS5/IS1182 family transposase. proline dehydrogenase family protein - DUF742 domain-containing protein - roadblock/LC7 domain-containing protein - sensor histidine kinase - ABC transporter substrate-binding protein - hypothetical protein - HNH endonuclease signature motif containing protein - bifunctional o-acetylhomoserine/o-acetylserine sulfhydrylase - homoserine O-acetyltransferase MetX - MurR/RpiR family transcriptional regulator - ABC transporter ATP-binding protein - ABC transporter substrate-binding protein - tyrosine-protein phosphatase - metallophosphoesterase. pyridoxal phosphate-dependent decarboxylase family protein - PHP domain-containing protein - hypothetical protein - class I SAM-dependent methyltransferase - MFS transporter - aldehyde dehydrogenase family protein - L-lysine 6-transaminase - protein kinase domain-containing protein - serine/threonine-protein kinase - superoxide dismutase family protein - type II toxin-antitoxin system VapC family toxin - type II toxin-antitoxin system Phd/YefM family antitoxin - 5,6-dimethylbenzimidazole synthase - hypothetical protein. recombinase family protein - hypothetical protein - hypothetical protein - hypothetical protein - albusnodin family lasso peptide - albusnodin/ikarugamycin family macrolactam cyclase - lasso peptide biosynthesis B2 protein - GNAT family N-acetyltransferase - replication initiator - recombinase family protein - recombinase family protein - MFS transporter - FAD-dependent monooxygenase - PPOX class F420-dependent oxidoreductase - cysteine hydrolase family protein. response regulator - potassium-transporting ATPase subunit C - potassium-transporting ATPase subunit KdpB - potassium-transporting ATPase subunit KdpA - K(+)-transporting ATPase subunit F - response regulator - DUF72 domain-containing protein - AbrB/MazE/SpoVT family DNA-binding domain-containing protein - type II toxin-antitoxin system VapC family toxin - winged helix-turn-helix transcriptional regulator - NAD(P)-binding domain-containing protein - NAD(P)-dependent oxidoreductase - ATP-binding protein - hypothetical protein. ROK family transcriptional regulator - M56 family metallopeptidase - BlaI/MecI/CopY family transcriptional regulator - cytochrome ubiquinol oxidase subunit I - cytochrome d ubiquinol oxidase subunit II - CPBP family intramembrane glutamic endopeptidase - CYTH and CHAD domain-containing protein - cold-shock protein - aspartate-semialdehyde dehydrogenase - aspartate kinase - TerD family protein - zinc-dependent alcohol dehydrogenase family protein - hypothetical protein - hypothetical protein. SAM-dependent methyltransferase - hypothetical protein - hypothetical protein - hypothetical protein - hypothetical protein - Eco57I restriction-modification methylase domain-containing protein -

DISARM system SNF2-like helicase DrmD - NADP-dependent oxidoreductase - hypothetical protein - transposase - helix-turn-helix domain-containing protein - type II toxin-antitoxin system HipA family toxin - hypothetical protein - hypothetical protein. SAM-dependent methyltransferase - ribulose-phosphate 3-epimerase - hypothetical protein - endonuclease/exonuclease/phosphatase family protein - aldo/keto reductase - hypothetical protein - B12-binding domain-containing radical SAM protein - hypothetical protein - DUF6624 domain-containing protein - AMP-binding protein - hypothetical protein - AMP-binding protein - TauD/TfdA family dioxygenase - dehydrogenase. SAVED domain-containing protein - AAA family ATPase - hypothetical protein - CBASS oligonucleotide cyclase - helix-turn-helix domain-containing protein - ImmA/IrrE family metallo-endopeptidase - hypothetical protein - IS110 family transposase - NUDIX hydrolase - hypothetical protein - hypothetical protein - hypothetical protein - hypothetical protein - hypothetical protein. SCO family protein - DUF3618 domain-containing protein - lysylphosphatidylglycerol synthase transmembrane domain-containing protein - ATP-binding protein - hypothetical protein - hypothetical protein - hypothetical protein - GroES family chaperonin - DUF4193 family protein - LuxR family transcriptional regulator - hypothetical protein - helix-turn-helix transcriptional regulator - type I polyketide synthase - type I polyketide synthase. SCP2 sterol-binding domain-containing protein - aminotransferase class IV family protein - IS110 family transposase - IS110 family transposase - hypothetical protein - toll/interleukin-1 receptor domain-containing protein - cation:proton antiporter - malate dehydrogenase - ABC transporter substrate-binding protein - GlcG/HbpS family heme-binding protein - hypothetical protein - DUF397 domain-containing protein - helix-turn-helix domain-containing protein - ATP-binding protein. SDR family NAD(P)-dependent oxidoreductase - acyl carrier protein - beta-ketoacyl synthase N-terminal-like domain-containing protein - IS6 family transposase - nucleotide disphospho-sugar-binding domain-containing protein - aromatase/cyclase - hypothetical protein - zinc-binding dehydrogenase - DHA2 family efflux MFS transporter permease subunit - ISAs1 family transposase - NADPH-dependent FMN reductase - hypothetical protein - hypothetical protein - helix-turn-helix domain-containing protein. serine/threonine-protein kinase - response regulator - hybrid sensor histidine kinase/response regulator - SpoIIE family protein phosphatase - ATP-binding protein - hypothetical protein - diaminopimelate decarboxylase - diaminobutyrate--2-oxoglutarate transaminase family protein - pyridoxal phosphate-dependent decarboxylase family protein - IucA/IucC family protein - lysine N(6)-hydroxylase/L-ornithine N(5)-oxygenase family protein - IucA/IucC family protein - glutamine synthetase family protein - MMPL family transporter. sulfotransferase family protein - methyltransferase domain-containing protein - rubrerythrin family protein - IS701 family transposase - IS701 family transposase - sulfatase - DUF6027 family protein - LacI family DNA-binding transcriptional regulator - MFS transporter - cytochrome P450 - adenosine deaminase - phosphotransferase - MFS transporter - class I SAM-dependent methyltransferase. TadE/TadG family type IV pilus assembly protein - hypothetical protein - aminoglycoside phosphotransferase family protein - CocE/NonD family hydrolase - ABC transporter permease - pilus assembly protein TadG-related protein - hypothetical protein - peptide chain release factor 2 - polysaccharide deacetylase family protein - uroporphyrinogen-III synthase - nitrite reductase large subunit NirB - nitrite reductase small subunit NirD - uroporphyrinogen-III C-methyltransferase - hypothetical protein. TerD family protein - aldo/keto reductase -

hypothetical protein - sporulation protein - GNAT family N-acetyltransferase - S9 family peptidase - N-acetylglutaminylglutamine amidotransferase - carboxylate--amine ligase/circularly permuted type 2 ATP-grasp protein - hypothetical protein - FMN-binding glutamate synthase family protein - VOC family protein - polyprenol phosphomannose-dependent alpha 1,6 mannosyltransferase MptB - glycosyltransferase 87 family protein - hypothetical protein. terpene cyclase/mutase family protein - 4-hydroxy-3-methylbut-2-enyl diphosphate reductase - winged helix-turn-helix transcriptional regulator - PaaI family thioesterase - adenosyl-hopene transferase HpnH - TetR/AcrR family transcriptional regulator - glycosyltransferase - urease accessory protein UreD - urease accessory protein UreF - urease subunit alpha - urease subunit beta - urease subunit gamma - hypothetical protein - adenine deaminase C-terminal domain-containing protein. TetR/AcrR family transcriptional regulator - FAD-binding oxidoreductase - hypothetical protein - SDR family NAD(P)-dependent oxidoreductase - LmeA family phospholipid-binding protein - ATP-binding cassette domain-containing protein - protein kinase domain-containing protein - Acg family FMN-binding oxidoreductase - hypothetical protein - type 1 glutamine amidotransferase domain-containing protein - HelD family protein - IS630 family transposase - helix-turn-helix domain-containing protein - hypothetical protein. TetR/AcrR family transcriptional regulator - PaaI family thioesterase - HNH endonuclease signature motif containing protein - MFS transporter - hypothetical protein - MarR family winged helix-turn-helix transcriptional regulator - GNAT family N-acetyltransferase - SHOCT domain-containing protein - PspA/IM30 family protein - PspA-associated protein PspAA - zinc metalloprotease HtpX - PspA-associated protein PspAB - M56 family metallopeptidase - BlaI/MecI/CopY family transcriptional regulator. thiamine phosphate synthase - sulfur carrier protein ThiS - Rv2175c family DNA-binding protein - Gfo/Idh/MocA family protein - NAD(P)/FAD-dependent oxidoreductase - 2-iminoacetate synthase ThiH - thiazole synthase - Stk1 family PASTA domain-containing Ser/Thr kinase - DUF5667 domain-containing protein - IS5/IS1182 family transposase - IS4 family transposase - helix-turn-helix domain-containing protein - aminotransferase class IV family protein - hypothetical protein. ThiF family adenylyltransferase - hypothetical protein - multiubiquitin domain-containing protein - helix-turn-helix domain-containing protein - ImmA/IrrE family metallo-endopeptidase - hypothetical protein - hypothetical protein - type I glyceraldehyde-3-phosphate dehydrogenase - hypothetical protein - hypothetical protein - IS66 family transposase - IS66 family transposase - LuxR C-terminal-related transcriptional regulator - AMP-binding protein. threonine synthase - chaperonin GroEL - Uma2 family endonuclease - DNA repair helicase XPB - helicase-associated domain-containing protein - hypothetical protein - hypothetical protein - DUF3180 domain-containing protein - 2-amino-4-hydroxy-6-hydroxymethyldihydropteridine diphosphokinase - dihydroneopterin aldolase - dihydropteroate synthase - GNAT family N-acetyltransferase - hypothetical protein - aminoglycoside phosphotransferase family protein. TlyA family RNA methyltransferase - hypothetical protein - alkyl sulfatase C-terminal domain-containing protein - HAD-IIA family hydrolase - tetratricopeptide repeat protein - DUF1015 family protein - tetratricopeptide repeat protein - 5S ribosomal RNA - 23S ribosomal RNA - 16S ribosomal RNA - DNA-3-methyladenine glycosylase - argininosuccinate lyase - argininosuccinate synthase - arginine repressor. translation elongation factor Ts - 30S ribosomal protein S2 - M23 family metallopeptidase - tyrosine-type recombinase/integrase - DNA-processing protein DprA - YifB family Mg chelatase-like AAA ATPase - YraN family protein - DUF2469 domain-containing protein - ribonuclease HII - signal peptidase I - 50S ribosomal protein L19 - tRNA (guanosine(37)-N1)-methyltransferase TrmD - ribosome maturation factor RimM - RNA-binding protein. transposase - hypothetical protein - maleylpyruvate isomerase N-terminal domain-containing protein - hypothetical protein - IS630 family transposase - IS5/IS1182 family transposase - transposase - NAD(P)/FAD-dependent oxidoreductase - peptide ligase PGM1-related protein - aminotransferase class III-fold pyridoxal phosphate-dependent enzyme - FAD-dependent monooxygenase - 3-dehydroquinate synthase II family protein - 2-amino-3,7-dideoxy-D-threo-hept-6-ulosonate synthase - aspartate kinase. transposase - hypothetical protein - transposase family protein - tyrosine-type recombinase/integrase - hypothetical protein - DUF2127 domain-containing protein - DUF2231 domain-containing protein - LLM class F420-dependent oxidoreductase - putative bifunctional diguanylate cyclase/phosphodiesterase - leucyl/phenylalanyl-tRNA--protein transferase - aminotransferase class I/II-fold pyridoxal phosphate-dependent enzyme - class I SAM-dependent methyltransferase - UvrD-helicase domain-containing protein - AAA domain-containing protein. trigger factor - ATP-dependent Clp protease proteolytic subunit - ATP-dependent Clp protease proteolytic subunit - ATP-dependent Clp protease ATP-binding subunit ClpX - bifunctional folylpolyglutamate synthase/dihydrofolate synthase - dihydrofolate synthase - nucleoside-diphosphate kinase - rod shape-determining protein - rod shape-determining protein MreC - rod shape-determining protein MreD - penicillin-binding protein 2 - rod shape-determining protein RodA - TIGR03960 family B12-binding radical SAM protein - TIGR03936 family radical SAM-associated protein. tRNA 2-thiouridine(34) synthase MnmA - cysteine desulfurase family protein - MarR family winged helix-turn-helix transcriptional regulator - fused isobutyryl-CoA mutase/GTPase IcmF - 5S ribosomal RNA - 23S ribosomal RNA - 16S ribosomal RNA - ROK family protein - phosphotransferase - hypothetical protein - vWA domain-containing protein - hypothetical protein - electron transfer flavoprotein subunit alpha/FixB family protein - electron transfer flavoprotein subunit beta/FixA family protein. tRNA-Trp - preprotein translocase subunit SecE - transcription termination/antitermination protein NusG - 50S ribosomal protein L11 - 50S ribosomal protein L1 - 50S ribosomal protein L10 - 50S ribosomal protein L7/L12 - DNA-directed RNA polymerase subunit beta - DNA-directed RNA polymerase subunit beta' - RluA family pseudouridine synthase - G/U mismatch-specific DNA glycosylase - 30S ribosomal protein S12 - 30S ribosomal protein S7 - elongation factor G. tryptophan--tRNA ligase - 16S rRNA (adenine(1518)-N(6)/adenine(1519)-N(6))-dimethyltransferase RsmA - TatD family hydrolase - ABC transporter substrate-binding protein - YdcF family protein - 16S rRNA (cytidine(1402)-2'-O)-methyltransferase - dolichyl-phosphate-mannose--protein mannosyltransferase - hypothetical protein - hypothetical protein - glycoside hydrolase family 15 protein - FHA domain-containing protein - DUF305 domain-containing protein - SulP family inorganic anion transporter - family 43 glycosylhydrolase. type I-C CRISPR-associated protein Cas5c - type I-C CRISPR-associated protein Cas8c/Csd1 - type I-C CRISPR-associated protein Cas7/Csd2 - CRISPR-associated protein Cas4 - type I-C CRISPR-associated endonuclease Cas1c - CRISPR-associated endonuclease Cas2 - site-specific integrase - Uma2 family endonuclease - alcohol dehydrogenase catalytic domain-containing protein - uracil-DNA glycosylase - benzoate/H(+) symporter BenE family transporter - hypothetical protein - epoxide hydrolase family protein - radical SAM protein. type I-E CRISPR-associated protein Cse2/CasB - type I-E CRISPR-associated protein Cas7/Cse4/CasC - type I-E CRISPR-associated protein Cas5/CasD - type I-E CRISPR-associated protein Cas6/Cse3/CasE - type I-E CRISPR-associated endonuclease Cas1e - type I-E CRISPR-associated endoribonuclease Cas2e - organic hydroperoxide resistance protein - lipase family protein - DedA family protein - P-II family nitrogen regulator - YbhB/YbcL family Raf kinase inhibitor-like protein - conjugal transfer protein TraC - SCO6880 family protein - hypothetical protein. type II toxin-antitoxin system death-on-curing family toxin - DUF433 domain-containing protein - hypothetical protein - tyrosine-type recombinase/integrase - hypothetical protein - 5'-methylthioadenosine/S-adenosylhomocysteine nucleosidase - NUDIX hydrolase - hypothetical protein - type ISP restriction/modification enzyme - IS66 family transposase - NF041680 family putative transposase - transposase - ISAzo13-like element ISFsp13 family transposase - ISAs1 family transposase. type I polyketide synthase - SGNH/GDSL hydrolase family protein - NAD(P)/FAD-dependent oxidoreductase - PaaI family thioesterase - alpha/beta hydrolase family protein - TetR/AcrR family transcriptional regulator - DUF1772 domain-containing protein - NAD-binding protein - GmrSD restriction endonuclease domain-containing protein - ISAs1 family transposase - ADP-ribosylglycohydrolase family protein - AMIN-like domain-containing (lipo)protein - aldehyde dehydrogenase family protein - zinc-dependent alcohol dehydrogenase. tyrosine-type recombinase/integrase - type II toxin-antitoxin system RelE/ParE family toxin - helix-turn-helix domain-containing protein - IS701 family transposase - ISAs1 family transposase - hypothetical protein - recombinase family protein - JmjC domain-containing protein - aldo/keto reductase - hypothetical protein - DUF397 domain-containing protein - helix-turn-helix domain-containing protein - hypothetical protein - ATP-binding protein. UbiX family flavin prenyltransferase - hypothetical protein - aminofutalosine synthase MqnE - hypothetical protein - DUF4229 domain-containing protein - menaquinone biosynthetic enzyme MqnA/MqnD family protein - helix-turn-helix domain-containing protein - response regulator - putative leader peptide - nitrite/sulfite reductase - hypothetical protein - phosphoadenylyl-sulfate reductase - uroporphyrinogen-III C-methyltransferase - sirohydrochlorin chelatase. uracil-DNA glycosylase - Ppx/GppA family phosphatase - DUF501 domain-containing protein - FtsB family cell division protein - phosphopyruvate hydratase - S49 family peptidase - MerR family transcriptional regulator - transcription-repair coupling factor - SurA N-terminal domain-containing protein - nucleoside triphosphate pyrophosphohydrolase - glycoside hydrolase family 26 protein - sugar phosphate nucleotidyltransferase - DUF4910 domain-containing protein - IspD/TarI family cytidylyltransferase. winged helix-turn-helix transcriptional regulator - NUDIX hydrolase - hypothetical protein - adenylate kinase family protein - hypothetical protein - HAD-IIA family hydrolase - hypothetical protein - hypothetical protein - hypothetical protein - recombinase family protein - peptide-methionine (S)-S-oxide reductase MsrA - PhoH family protein - phosphoenolpyruvate carboxykinase (GTP) - PAQR family membrane homeostasis protein TrhA. YbhB/YbcL family Raf kinase inhibitor-like protein - conjugal transfer protein TraC - SCO6880 family protein - hypothetical protein - hypothetical protein - DUF6112 family protein - MinD/ParA family ATP-binding protein - helix-turn-helix domain-containing protein - hypothetical protein - NlpC/P60 family protein - hypothetical protein - FAD-binding oxidoreductase - protein kinase domain-containing protein - glucose-6-phosphate isomerase. zinc ribbon domain-containing protein - IS701 family transposase - ribbon-helix-helix protein,

CopG family - type II toxin-antitoxin system death-on-curing family toxin - DUF433 domain-containing protein - hypothetical protein - tyrosine-type recombinase/integrase - hypothetical protein - 5'-methylthioadenosine/S-adenosylhomocysteine nucleosidase - NUDIX hydrolase - hypothetical protein - type ISP restriction/modification enzyme - IS66 family transposase - NF041680 family putative transposase.

ChatGPT 5.5 answer:

I'll screen for the strongest markers first: Tad/TadE/TadG, type IV pilus assembly, type II secretion F/pilus ATPase, TraC/conjugal transfer, and then weaker accessory matches such as Fic/HAD/toxin-mobile island context.

The strongest hits are clear: three contexts contain explicit Tad/type-IV-pilus assembly genes, and two contain a conjugal transfer ATPase/TraC-like signal. I'm separating those from broader biofilm-regulatory hints so the answer doesn't over-call unrelated mobile or metabolic islands.

Using each line as one numbered context, the **best matches to the Tad/Flp type-IV-pilus / adhesion / colonization / conjugation-like segment** are:

**High-confidence related contexts**

**38, 84, 140, 157, 164**

These are the contexts I would treat as biologically/functionally related to the segment discussed above.

| Context # | Why it matches |
|---|---|
| 38 | Contains **pilus assembly protein TadG-related protein**, **TadE/TadG family type IV pilus assembly protein**, and **type II secretion system F family protein**. This is a strong Tad/type-IV-pilus assembly signal. |
| 84 | Contains **BTAD domain-containing putative transcriptional regulator**, **pilus assembly protein TadG-related**, **TadE/TadG family type IV pilus assembly protein**, **TadE family protein**, and **type II secretion system F family protein**. This is the strongest match in the list. |
| 140 | Starts with **TadE/TadG family type IV pilus assembly protein** and also contains **pilus assembly protein TadG-related protein**. Strong Tad/Flp/type-IV-pilus relationship. |
| 157 | Contains **conjugal transfer protein TraC**. This is not a TadE/TadG context, but it matches the **conjugal-transfer-associated ATPase / type-IV secretion-pilus-like ATPase** aspect of the original interpretation. |
| 164 | Contains **conjugal transfer protein TraC** plus nearby regulatory/cell-envelope proteins. This is related to the conjugation-like ATPase side of the original segment. |

**Possible but weaker related context**

**61**

Context **61** contains a **Fic family protein**:

GNAT family N-acetyltransferase - hypothetical protein - transposase domain-containing protein - type II toxin-antitoxin system HicA family toxin - type II toxin-antitoxin system HicB family antitoxin - recombinase family protein - ISAs1 family transposase - **Fic family protein** - ...

This is **not a Tad/Flp pilus context**, but it is biologically related to the **accessory stress/toxin/mobile-island part** of the interpretation I gave earlier. Fic proteins are often associated with toxin-like, stress, virulence, or mobile-element functions. So I would mark **61 as a weak/accessory match**, not as a direct pilus/adhesion module.

**Final classification**

**Direct Tad/type-IV-pilus/conjugation-related contexts:**
**38, 84, 140, 157, 164**

**Accessory/mobile-island Fic-related but not direct Tad/pilus:**
**61**

I would **not** include the many contexts with only integrases, transposases, restriction-modification, CRISPR, lasso peptides, HAD hydrolases, septation/cell-division proteins, or generic ATPases unless additional neighboring genes show Tad/Flp/type-IV-pilus components. Those features can occur in mobile or defense islands, but they do not by themselves identify the same biological module.

Supplementary File S17. ChatGPT-assisted analysis of functional relations between genomic contexts of IGRs with predicted structure candidates in *Frankia sp. ArI3* and ms1 RNA synteny.

Question to ChatGPT 5.5:

These are NCBI genomic annotations of a succession of bacterial genes: Fic family protein, HAD-IB hydrolase, septum site-determining protein Ssd, TadA family conjugal transfer-associated ATPase. The genes are adjacent to each other in bacterial genomes and represent a segment of bacterial genomes. Tell me, which bacterial cellular biological processes/functions are related to the segment with such genes and also tell me which of the genes, if not all, are functionally/biologically related.

ChatGPT 5.5 answer:

I'll treat this as a genome-neighborhood/operon inference problem: identify likely functions of each annotation, then ask whether the adjacency suggests a shared biological module rather than four unrelated housekeeping genes.

The segment most likely points to a **cell-envelope / surface-structure / cell-cycle-stress module**, not a single classical metabolic pathway.

### Likely biological processes represented

**1. Type IV/Tad pilus assembly, adhesion, surface colonization, and possibly DNA transfer**
The strongest signal is **TadA family conjugal transfer-associated ATPase**. TadA-like ATPases are part of **Tad/Flp type IV pilus systems**, which assemble pili used for bacterial surface interaction. Tad/T4P systems are linked to **adherence, motility, biofilm formation, host interaction, DNA uptake, and horizontal gene transfer**, depending on the organism and the rest of the locus. Recent reviews describe Tad pili as conserved bacterial type IV pilus machineries used in many physiological lifestyles, with T4P broadly facilitating motility, adherence, and DNA uptake.

**2. Septum positioning, cell division control, filamentation, and dormancy/stress physiology**
**Septum site-determining protein Ssd** is the clearest cell-division gene. In *Mycobacterium tuberculosis*, Ssd/Rv3660c was shown to inhibit septum formation when overexpressed, producing elongated cells lacking septa, and to induce a dormancy/alternative metabolism transcriptional response.

This suggests the segment may couple **surface-structure biogenesis** with **cell-cycle state or stress adaptation**.

**3. Fic-mediated post-translational regulation, toxin-like activity, stress arrest, or defense**
**Fic family proteins** often catalyze AMPylation or related post-translational modifications. Many bacterial Fic proteins are toxin-like regulators; some inhibit DNA gyrase or topoisomerase IV, causing growth arrest, filamentation, SOS induction, and disruption of chromosome segregation/cell separation.

So the Fic gene may connect biologically to **stress-induced growth modulation**, **cell-cycle arrest**, **toxin–antitoxin-like behavior**, or **defense/mobile-element biology**.

**4. Generic phosphatase/hydrolase activity, possible regulatory or metabolic accessory role**
**HAD-IB hydrolase** is the least specific annotation. HAD superfamily proteins include phosphatases and other aspartate-nucleophile hydrolases; InterPro classifies HAD-IB as a haloacid dehalogenase-superfamily subgroup.

Without the sequence/domain details, I would not assign it confidently to the same process. It could be a **small-molecule phosphatase**, **regulatory phosphatase**, or simply an accessory enzyme carried with the locus.

### Which genes are functionally related?

**Strongly related:**
**Ssd + TadA** are very likely related. In actinobacterial/mycobacterial contexts, Ssd/Rv3660c sits in or near a type IV/Tad/Flp pilus-associated region, and TadA is a core ATPase for that kind of system. Search results for *M. tuberculosis* note that Rv3657c–Rv3660c resemble type IV pili/type II-IV secretion/tad genes, and later work labels Rv3660c as **tadZ** in an Flp type IV pilus operon context.

So **Ssd may be a cell-cycle/septation regulator embedded in or evolutionarily related to a Tad pilus locus**, rather than a random neighboring gene.

**Possibly related:**
**Fic + Ssd** may be biologically related through their effects on **cell division, filamentation, stress arrest, and persistence-like states**. Fic proteins can inhibit topoisomerase IV and cause filamentation; Ssd also affects septum formation and dormancy-related transcription.

However, this is a functional convergence, not proof that the two proteins directly interact.

**Weakly or uncertainly related:**
**HAD-IB hydrolase** may be part of the same conserved neighborhood, but the annotation alone is too broad. It could be a regulatory enzyme associated with the module, but I would not call it functionally linked without synteny conservation, co-occurrence, operon prediction, or domain/substrate information.

### Best interpretation

This gene block most plausibly represents a **Tad/type IV pilus-associated genomic segment with cell-cycle/stress-regulatory accessories**. The core biological theme is:

**surface attachment / pilus or secretion-system assembly + septation/cell-cycle control + stress-growth modulation.**

A more conservative summary would be:

| Gene | Likely role | Relationship to s |
|---|---|---|
| Fic family protein | toxin-like AMPylating/regulatory protein; stress arrest, chromosome/cell-division effects | Possibly accessory stre regulator |
| HAD-IB hydrolase | broad hydrolase/phosphatase-like enzyme | Uncertain; possible acc enzyme |
| Ssd | septum placement/cell division, filamentation, dormancy response | Strongly relevant |
| TadA ATPase | Tad/type IV pilus assembly/conjugation-associated ATPase | Strongly relevant |

So, **Ssd and TadA are the most clearly functionally related pair**, likely tied to a Tad/Flp/type IV pilus locus. **Fic is plausibly related at the level of stress, growth arrest, and cell-division modulation. HAD-IB is the least interpretable from annotation alone.**

Question to ChatGPT 5.5:

Now I will send you genomic contexts, one context per a line, terminated by dots. The contexts consist of NCBI genomic annotations, separated by the '-' characters. They represent segments of bacterial genomes that contain the genes encoding the proteins whose bilogical/functional roles are characterized by the annotations. Tell me, which of the contexts are biologically/functionally related to the characterisctics that you gave me above. Indentify the contexts using their order numbers. First count the contexts and write me how many of them you counted. Here are the contexts: 0 - 0 -

- hypothetical protein - group II intron reverse transcriptase/maturase - IS4 family transposase - DinB family protein : transposase - transposase - DDE-type integrase/transposase/recombinase - IS6 family transposase - hypothetical protein - IS6 family transposase - potassium transporter Kup. 1-acyl-sn-glycerol-3-phosphate acyltransferase - MMPL family transporter - ChaB family protein - N-acetylmuramoyl-L-alanine amidase - trans-aconitate 2-methyltransferase - zinc-binding dehydrogenase - NADP-dependent oxidoreductase : IS701 family transposase - helix-turn-helix domain-containing protein - carbonic anhydrase - dienelactone hydrolase family protein - LLM class flavin-dependent oxidoreductase - NtaA/DmoA family FMN-dependent monooxygenase - phosphotransferase family protein. 2-hydroxyacid dehydrogenase - NAD-dependent epimerase/dehydratase family protein - hypothetical protein - PAC2 family protein - hypothetical protein - sensor histidine kinase - DHA2 family efflux MFS transporter permease subunit : TetR/AcrR family transcriptional regulator C-terminal domain-containing protein - pyridoxamine 5'-phosphate oxidase family protein - ammonium transporter - carbon-nitrogen family hydrolase - MBL fold metallo-hydrolase - helix-turn-helix domain-containing protein - PHB depolymerase family esterase. 30S ribosomal protein S15 - polyribonucleotide nucleotidyltransferase - pitrilysin family protein - 4-hydroxy-tetrahydrodipicolinate reductase - Hsp70 family protein - hypothetical protein - UDP-glucuronic acid decarboxylase family protein : UDP-glucose 4-epimerase GalE - UDP-glucose/GDP-mannose dehydrogenase family protein - histidine--tRNA ligase - FAD-dependent thymidylate synthase - hypothetical protein - acetate/propionate family kinase - hypothetical protein. 3-isopropylmalate dehydratase large subunit - 3-isopropylmalate dehydratase small subunit - HU family DNA-binding protein - RNA degradosome polyphosphate kinase - transglutaminase family protein - bifunctional diguanylate cyclase/phosphodiesterase - 2-phospho-L-lactate guanylyltransferase : NAD(P)H-dependent glycerol-3-phosphate dehydrogenase - UDP-N-acetylglucosamine 1-carboxyvinyltransferase - bifunctional hydroxymethylpyrimidine kinase/phosphomethylpyrimidine kinase - Lrp/AsnC family transcriptional regulator - DUF3515 domain-containing protein - D-alanine--D-alanine ligase family protein - HNH endonuclease signature motif containing protein. 4-hydroxy-3-methylbut-2-enyl diphosphate reductase - exodeoxyribonuclease VII large subunit - exodeoxyribonuclease VII small subunit - DUF4245 domain-containing protein - class II fructose-bisphosphatase - sterol desaturase family protein - polysaccharide deacetylase family protein : DUF6542 domain-containing protein - DNA recombination protein RmuC - redox-regulated ATPase YchF - DUF952 domain-containing protein - ribosylglycohydrolase - 4a-hydroxytetrahydrobiopterin dehydratase - (deoxy)nucleoside triphosphate pyrophosphohydrolase. 50S ribosomal protein L25/general stress protein Ctc - aminoacyl-tRNA hydrolase - glycosyltransferase - glycosyltransferase family 2 protein - glycosyltransferase - tyrosine--tRNA ligase - S8 family serine peptidase : HAD family hydrolase - acyl carrier protein - serine O-acetyltransferase - DUF4062 domain-containing protein - NAD(P)-dependent oxidoreductase - dTDP-4-dehydrorhamnose 3,5-epimerase family protein - class I SAM-dependent methyltransferase. 50S ribosomal protein L32 - phosphate starvation protein PhoH - ribonuclease III - bifunctional DNA-formamidopyrimidine glycosylase/DNA-(apurinic or apyrimidinic site) lyase - DUF2786 domain-containing protein - acylphosphatase - hypothetical protein : AAA family ATPase - signal recognition particle-docking protein FtsY - signal recognition particle protein - 30S ribosomal protein S16 - RNA-binding protein - ribosome maturation factor RimM - tRNA (guanosine(37)-N1)-methyltransferase TrmD. 5S ribosomal RNA - hypothetical protein - lipopolysaccharide assembly protein LapB - DUF1015 family protein - lipopolysaccharide assembly protein LapB - HAD-IIA family hydrolase - alkyl sulfatase C-terminal domain-containing protein : hypothetical protein - TlyA family RNA methyltransferase - NAD kinase - DNA repair protein RecN - aldo/keto reductase - CTP synthase - NUDIX domain-containing protein. AAA domain-containing protein - UvrD-helicase domain-containing protein - class I SAM-dependent methyltransferase - hypothetical protein - aminotransferase class I/II-fold pyridoxal phosphate-dependent enzyme - leucyl/phenylalanyl-tRNA--protein transferase - bifunctional diguanylate cyclase/phosphodiesterase : hypothetical protein - M56 family metallopeptidase - hypothetical protein - BlaI/MecI/CopY family transcriptional regulator - DUF2231 domain-containing protein - DUF2127 domain-containing protein - hypothetical protein. AAA family ATPase - hypothetical protein - hypothetical protein - hypothetical protein - hypothetical protein - hypothetical protein - hypothetical protein : hypothetical protein - hypothetical protein - hypothetical protein - YbaB/EbfC family nucleoid-associated protein - hypothetical protein - hypothetical protein - WXG100 family type VII secretion target. ABC transporter ATP-binding protein - ABC transporter permease - FUSC family protein - hypothetical protein - MFS transporter - MarR family winged helix-turn-helix transcriptional regulator - tRNA-Ser : nuclear transport factor 2 family protein - helix-turn-helix domain-containing protein - hypothetical protein - DUF3817 domain-containing protein - hypothetical protein - hypothetical protein - SigE family RNA polymerase sigma factor. ABC transporter permease - ABC transporter ATP-binding protein - hypothetical protein - BlaI/MecI/CopY family transcriptional regulator - MarR family winged helix-turn-helix transcriptional regulator - MFS transporter - cyclopropane-fatty-acyl-phospholipid synthase family protein : serine/threonine-protein kinase - helix-turn-helix domain-containing protein - NADPH-dependent F420 reductase - MFS transporter - hypothetical protein - alpha/beta fold hydrolase - cupin domain-containing protein. ABC transporter permease - ABC transporter substrate-binding protein - ABC transporter permease - hypothetical protein - M20/M25/M40 family metallo-hydrolase - hypothetical protein - hypothetical protein : ATP-binding protein - hypothetical protein - DUF5703 family protein - hypothetical protein - glycosyltransferase family 39 protein - NAD(P)/FAD-dependent oxidoreductase - glycosyltransferase. ABC transporter permease subunit - hypothetical protein - hypothetical protein - hypothetical protein - DUF1003 domain-containing protein - aldo/keto reductase - DUF2142 domain-containing protein : bifunctional SulP family inorganic anion transporter/carbonic anhydrase - ATP-binding domain-containing protein - ABC transporter ATP-binding protein - type I restriction-modification system subunit M N-terminal domain-containing protein - RloB family protein - ATP/GTP-binding protein - cytochrome P450. ABC transporter substrate-binding protein - acyl-CoA dehydrogenase family protein - acyl-CoA dehydrogenase family protein - ABC transporter substrate-binding protein - amidohydrolase family protein - acetoacetate decarboxylase family protein - CaiB/BaiF CoA transferase family protein : ATP-binding cassette domain-containing protein - ABC transporter ATP-binding protein - FadR/GntR family transcriptional regulator - hypothetical protein - LLM class F420-dependent oxidoreductase - iron-containing alcohol dehydrogenase - aldehyde dehydrogenase family protein. ABC transporter substrate-binding protein - hypothetical protein - hypothetical protein - DUF2752 domain-containing protein - RNA methyltransferase - 8-oxoguanine deaminase - bacterioferritin : aldolase/citrate lyase family protein - anion permease - hypothetical protein - DUF3349 domain-containing protein - hydroxyisourate hydrolase - allantoicase - allantoinase AllB. acyl carrier protein - DUF5615 family PIN-like protein - DUF433 domain-containing protein - HIT family protein - Dyp-type peroxidase - hypothetical protein - DNA-binding protein : MerR family transcriptional regulator - MFS transporter - uridine kinase - trans-aconitate 2-methyltransferase - TetR/AcrR family transcriptional regulator - alpha/beta fold hydrolase - hypothetical protein. acyl-CoA dehydrogenase family protein - helix-turn-helix domain-containing protein - RNA-guided endonuclease InsQ/TnpB family protein - nuclear transport factor 2 family protein - cytochrome P450 - TIGR01458 family HAD-type hydrolase - arsenate reductase ArsC : polyketide cyclase - hypothetical protein - MOSC domain-containing protein - enoyl-CoA hydratase - acyl-CoA synthetase - bifunctional diguanylate cyclase/phosphodiesterase - SAM-dependent methyltransferase. acyl-CoA dehydrogenase family protein - SfnB family sulfur acquisition oxidoreductase - LLM class flavin-dependent oxidoreductase - histidine phosphatase family protein - RNA-guided endonuclease InsQ/TnpB family protein - DUF2993 domain-containing protein - phosphatidylglycerol lysyltransferase domain-containing protein : zinc-binding dehydrogenase - helix-turn-helix domain-containing protein - CBS domain-containing protein - RNA polymerase sigma factor - IS4 family transposase - LysR family transcriptional regulator - MFS transporter. acyltransferase - oxidoreductase - LysR family transcriptional regulator - hypothetical protein - hypothetical protein - LysR substrate-binding domain-containing protein - Uma2 family endonuclease : phosphotransferase - FCD domain-containing protein - hypothetical protein - 5,6-dimethylbenzimidazole synthase - TetR/AcrR family transcriptional regulator - hypothetical protein - LLM class F420-dependent oxidoreductase. adenosine deaminase - glycerate kinase - N-acetylmuramoyl-L-alanine amidase - GNAT family N-acetyltransferase - hypothetical protein - adenosylhomocysteinase - RND family transporter : DUF1918 domain-containing protein - phosphatase PAP2 family protein - acetyl-CoA C-acetyltransferase - C2 family cysteine protease - universal stress protein - hypothetical protein - hypothetical protein. adenylosuccinate synthase - DUF3151 domain-containing protein - transglycosylase family protein - transglycosylase family protein - class II fructose-bisphosphate aldolase - DedA family protein - CAP domain-containing protein : NAD(P)-dependent oxidoreductase - hypothetical protein - hypothetical protein - PH domain-containing protein - PH domain-containing protein - MMPL family transporter - pseudouridine-5'-phosphate glycosidase. aldehyde dehydrogenase family protein - hypothetical protein - hypothetical protein - hypothetical protein - metallophosphoesterase - plastocyanin/azurin family copper-binding protein - hypothetical protein : VOC family protein - helix-turn-helix domain-containing protein - IS630 family transposase - helix-turn-helix domain-containing protein - IS630 family transposase - DUF1360 domain-containing protein - pyridoxamine 5'-phosphate oxidase family protein. aldo/keto reductase - CTP synthase - NUDIX domain-containing protein - PKD domain-containing protein - hypothetical protein - site-specific tyrosine recombinase XerD - hypothetical protein : NADPH-dependent F420 reductase - helical backbone metal receptor - 3-methyl-2-oxobutanoate hydroxymethyltransferase - TetR/AcrR family transcriptional regulator C-terminal ligand-binding domain-containing protein - threonine--tRNA ligase - HEPN domain-containing protein - KGGVGR-motif variant AAA ATPase. aldo/keto reductase family protein - DUF3043 domain-containing protein - quinolinate synthase NadA - iron-sulfur cluster insertion protein ErpA - carbohydrate kinase family protein - sulfurtransferase TusA family protein - cysteine desulfurase family protein : cytochrome c oxidase subunit II - cytochrome c oxidase subunit I - cytochrome c oxidase subunit 4 - IS1182 family transposase - anthranilate phosphoribosyltransferase - response regulator - Fur family transcriptional regulator. aldo/keto reductase - SDR family NAD(P)-dependent oxidoreductase - PaaX family transcriptional regulator C-terminal domain-containing protein - LLM class F420-dependent oxidoreductase - enoyl-CoA hydratase/isomerase family protein - FkbM family methyltransferase - LLM class F420-dependent oxidoreductase : trypsin-like peptidase domain-containing protein - S1C family serine protease - metallophosphoesterase - TIR domain-containing protein - hypothetical protein - DUF6461 domain-containing protein - hypothetical protein. alkaline phosphatase - MaoC family dehydratase - suppressor of fused domain protein - TerD family protein - NAD(P)-dependent oxidoreductase - acyl-CoA dehydrogenase family protein - CoA ester lyase : magnesium transporter MgtE N-terminal domain-containing protein - DUF1003 domain-containing protein - chorismate mutase - PD-(D/E)XK nuclease family protein - pyridoxal phosphate-dependent aminotransferase - hypothetical protein - hypothetical protein. AlkA N-terminal domain-containing protein - methylated-DNA--[protein]-cysteine S-methyltransferase - 2OG-Fe(II) oxygenase - UBP-type zinc finger domain-containing protein - multicopper oxidase family protein - RtcB family protein - hypothetical protein : thiamine pyrophosphate-dependent dehydrogenase E1 component subunit alpha - aldehyde dehydrogenase family protein - LppP/LprE family lipoprotein - NAD(P)-binding oxidoreductase - methylmalonyl-CoA epimerase - MerR family transcriptional regulator - MerR family transcriptional regulator. alkene reductase - VOC family protein - PPOX class F420-dependent oxidoreductase - stage II sporulation protein M - acyltransferase domain-containing protein - beta-ketoacyl synthase N-terminal-like domain-containing protein - response regulator : RDD family protein - adenosylhomocysteinase - MerR family transcriptional regulator - transglycosylase domain-containing protein - hypothetical protein - SIS domain-containing protein - Trm112 family protein. allophanate hydrolase - cysteine hydrolase family protein - GntR family transcriptional regulator - alpha/beta fold hydrolase - hypothetical protein - hypothetical protein - hypothetical protein : FmdB family zinc ribbon protein - 5-formyltetrahydrofolate cyclo-ligase - UTP--glucose-1-phosphate uridylyltransferase GalU - gephyrin-like molybdotransferase Glp - cyclic pyranopterin monophosphate synthase MoaC - molybdenum cofactor biosynthesis protein B - GNAT family N-acetyltransferase. alpha-amylase family protein - hypothetical protein - nuclear transport factor 2 family protein - amidohydrolase family protein - hypothetical protein - FadR/GntR family transcriptional regulator - serine hydrolase : helix-turn-helix transcriptional regulator - GNAT family N-acetyltransferase - TIR domain-containing protein - pyridoxal-dependent decarboxylase - NAD(P)-dependent oxidoreductase - hypothetical protein - tetratricopeptide repeat protein. alpha/beta fold hydrolase - phosphopantetheine-binding protein - beta-ketoacyl synthase N-terminal-like domain-containing protein - SDR family NAD(P)-dependent oxidoreductase - hypothetical protein - beta-ketoacyl synthase N-terminal-like domain-containing protein - LuxR C-terminal-related transcriptional regulator : hypothetical protein - LuxR family transcriptional regulator - DUF4193 family protein - co-chaperone GroES - hypothetical protein - hypothetical protein - ATP-binding protein. alpha/beta hydrolase - class I adenylate-forming enzyme family protein - enoyl-CoA hydratase/isomerase family protein - cytochrome P450 - NDMA-dependent alcohol dehydrogenase - aldehyde dehydrogenase - class I adenylate-forming enzyme family protein : hypothetical protein - phosphotransferase family protein - dienelactone hydrolase family protein - helix-turn-helix domain-containing protein - TIR domain-containing protein - tRNA-Asn - hypothetical protein. alpha/beta hydrolase domain-containing protein - alpha/beta hydrolase domain-containing protein - HNH endonuclease signature motif containing protein - mycofactocin-coupled SDR family oxidoreductase - aldehyde dehydrogenase family protein - hypothetical protein - TetR/AcrR family transcriptional regulator : cytochrome P450 - ferredoxin - helix-turn-helix domain-containing protein - alpha/beta hydrolase - glycosyltransferase family 2 protein - class I SAM-dependent methyltransferase - acyl-CoA dehydrogenase family protein. amidohydrolase family protein - hypothetical protein - FadR/GntR family transcriptional regulator - serine hydrolase - helix-turn-helix transcriptional regulator - GNAT family N-acetyltransferase - TIR domain-containing protein : pyridoxal-dependent decarboxylase - NAD(P)-dependent oxidoreductase - hypothetical protein - tetratricopeptide repeat protein - tRNA-Ser - tRNA adenosine(34) deaminase TadA - tRNA adenosine deaminase-associated protein. amino acid ABC transporter permease - transporter substrate-binding domain-containing protein - citrate/2-methylcitrate synthase - citrate/2-methylcitrate synthase - hypothetical protein - transposase - DNRLRE domain-containing protein : transposase - RHS repeat-associated core domain-containing protein - DDE-type integrase/transposase/recombinase - IS110 family transposase - FkbM family methyltransferase - serine/threonine-protein kinase - fibronectin type III domain-containing protein. aminopeptidase N - DUF6529 family protein - Rieske (2Fe-2S) protein - FHA domain-containing protein - hypothetical protein - type II toxin-antitoxin system RelE/ParE family toxin - type II toxin-antitoxin system Phd/YefM family antitoxin : glycosyltransferase family 2 protein - SCO family protein - hypothetical protein - PA14 domain-containing protein - glycosyltransferase family 2 protein - FAD-binding and (Fe-S)-binding domain-containing protein - ATP-binding domain-containing protein. AMP-binding protein - nuclear transport factor 2 family protein - hypothetical protein - hypothetical protein - hypothetical protein - alpha/beta hydrolase domain-containing protein - alpha/beta hydrolase domain-containing protein : HNH endonuclease signature motif containing protein - mycofactocin-coupled SDR family oxidoreductase - aldehyde dehydrogenase family protein - hypothetical protein - TetR/AcrR family transcriptional regulator - cytochrome P450 - ferredoxin. antibiotic biosynthesis monooxygenase - antibiotic biosynthesis monooxygenase - hypothetical protein - flavin reductase family protein - enoyl-CoA hydratase/isomerase family protein - AAA domain-containing protein - NB-ARC domain-containing protein : ATP-dependent 6-phosphofructokinase - FAD-dependent oxidoreductase - nuclear transport factor 2 family protein - GNAT family N-acetyltransferase - carbohydrate ABC transporter permease - carbohydrate ABC transporter permease - metallophosphoesterase. antibiotic biosynthesis monooxygenase - collagen-like protein - dihydroxy-acid dehydratase - L-histidine N(alpha)-methyltransferase - ergothioneine biosynthesis protein EgtC - ergothioneine biosynthesis protein EgtB - ergothioneine biosynthesis glutamate--cysteine ligase EgtA : nuclease-related domain-containing protein - DLW-39 family protein - alternative ribosome rescue aminoacyl-tRNA hydrolase ArfB - hypothetical protein - ABC transporter permease - ABC transporter permease - PotD/PotF family extracellular solute-binding protein. ATP-binding cassette domain-containing protein - hypothetical protein - hypothetical protein - PASTA domain-containing protein - hypothetical protein - WXG100 family type VII secretion target - FtsK/SpoIIIE domain-containing protein : serine/threonine-protein kinase - MoxR family ATPase - DUF58 domain-containing protein - transglutaminase domain-containing protein - SigE family RNA polymerase sigma factor - hypothetical protein - hypothetical protein. ATP-binding protein - DUF234 domain-containing protein - patatin-like phospholipase family protein - hypothetical protein - DUF4407 domain-containing protein - PP2C family serine/threonine-protein phosphatase - LLM class F420-dependent oxidoreductase : hypothetical protein - NADP-dependent isocitrate dehydrogenase - DUF6328 family protein - serine/threonine-protein kinase - flavin reductase family protein - DUF4397 domain-containing protein - hypothetical protein. ATP-binding protein - hypothetical protein - FxSxx-COOH system tetratricopeptide repeat protein - winged helix-turn-helix domain-containing protein - transposase - ATP-binding protein - hypothetical protein : effector-associated domain EAD1-containing protein - hypothetical protein - FAD-binding oxidoreductase - hypothetical protein - transposase - winged helix-turn-helix domain-containing protein - IS701 family transposase. ATP-binding protein - YqcI/YcgG family protein - HD family hydrolase - effector-associated domain EAD1-containing protein - trans-aconitate 2-methyltransferase - daunorubicin resistance protein DrrA family ABC transporter ATP-binding protein - ABC transporter permease : BTAD domain-containing putative transcriptional regulator - flavodoxin family protein - hypothetical protein - hypothetical protein - flavodoxin - type II toxin-antitoxin system PemK/MazF family toxin - type II toxin-antitoxin system HicA family toxin. ATP/GTP-binding protein - ABC transporter substrate-binding protein - hypothetical protein - hypothetical protein - GuaB3 family IMP dehydrogenase-related protein - IMP dehydrogenase - DUF5319 domain-containing protein : WhiB family transcriptional regulator - chaperonin GroEL - co-chaperone GroES - hypothetical protein - MarR family winged helix-turn-helix transcriptional regulator - FMN-dependent NADH-azoreductase - ribosomal protein S18-alanine N-acetyltransferase. beta-ketoacyl synthase N-terminal-like domain-containing protein - SDR family NAD(P)-dependent oxidoreductase - hypothetical protein - beta-ketoacyl synthase N-terminal-like domain-containing protein - LuxR C-terminal-related transcriptional regulator - hypothetical protein - LuxR family transcriptional regulator : DUF4193 family protein - co-chaperone GroES - hypothetical protein - hypothetical protein - ATP-binding protein - lysylphosphatidylglycerol synthase transmembrane domain-containing protein - histidine phosphatase family protein. bifunctional diguanylate cyclase/phosphodiesterase - class II glutamine amidotransferase - CoA transferase - phosphatase PAP2 family protein - alpha/beta fold hydrolase - HAD family phosphatase - lipase : ABC transporter substrate-binding protein - pre-mycofactocin synthase MftD - hypothetical protein - hypothetical protein - YceI family protein - helix-turn-helix domain-containing protein - mycofactocin radical SAM maturase. bifunctional GNAT family N-acetyltransferase/acetate--CoA ligase family protein - PAS domain-containing protein - SpoIIE family protein phosphatase - ABC transporter ATP-binding protein - hypothetical protein - Lsr2 family protein - aminopeptidase P family protein : LCP family protein - YihY/virulence factor BrkB family protein - hypothetical protein - septation protein SepH - polyphosphate--glucose phosphotransferase - GTP 3',8-cyclase MoaA - molybdenum cofactor guanylyltransferase. bifunctional nuclease family protein - MerR family transcriptional regulator - FHA domain-containing protein - glycine cleavage system protein GcvH - DUF881 domain-containing protein - DUF1290 domain-containing protein - DUF881 domain-containing protein : sugar phosphate nucleotidyltransferase - CDP-alcohol phosphatidyltransferase family protein - NlpC/P60 family protein - class F sortase - hypothetical protein - tRNA-Pro - orotidine-5'-phosphate decarboxylase. C2 family cysteine protease - universal stress protein - hypothetical protein - hypothetical protein - FAD-dependent oxidoreductase - hypothetical protein - peptidylprolyl isomerase : rhomboid family intramembrane serine protease - PH domain-containing protein - glycoside hydrolase family 15 protein - hypothetical protein - RidA family protein - cell division protein CrgA - DUF881 domain-containing protein. CaiB/BaiF CoA-transferase family protein - acyl-CoA dehydrogenase - FadR/GntR family transcriptional regulator - ABC transporter substrate-binding protein - ABC transporter ATP-binding protein - branched-chain amino acid ABC transporter permease/ATP-binding protein - SDR family oxidoreductase : ABC transporter substrate-binding protein - hypothetical protein - hypothetical protein - transcriptional regulator - amidohydrolase family protein - TetR/AcrR family transcriptional regulator - nuclear transport factor 2 family protein.

CapA family protein - DoxX family protein - alpha/beta fold hydrolase - NTP transferase domain-containing protein - GNAT family N-acetyltransferase - substrate-binding domain-containing protein - DUF6158 family protein : sensor histidine kinase - response regulator - hypothetical protein - Hsp70 family protein - dienelactone hydrolase family protein - hypothetical protein - hypothetical protein. carbohydrate ABC transporter permease - metallophosphoesterase - ABC transporter substrate-binding protein - ABC transporter ATP-binding protein - MurR/RpiR family transcriptional regulator - homoserine O-acetyltransferase - bifunctional o-acetylhomoserine/o-acetylserine sulfhydrylase : alkaline phosphatase D family protein - hypothetical protein - RNA polymerase sigma factor - amidohydrolase family protein - hypothetical protein - ABC transporter permease subunit - ABC transporter ATP-binding protein. carbohydrate kinase family protein - DUF3616 domain-containing protein - helix-turn-helix transcriptional regulator - UbiA family prenyltransferase - toll/interleukin-1 receptor domain-containing protein - biotin carboxylase N-terminal domain-containing protein - carboxyl transferase domain-containing protein : DUF5925 domain-containing protein - NADAR family protein - DUF3710 domain-containing protein - hypothetical protein - hypothetical protein - precorrin-4 C(11)-methyltransferase - cobalt-precorrin-6A reductase. cation diffusion facilitator family transporter - hypothetical protein - helix-turn-helix domain-containing protein - enoyl-CoA hydratase/isomerase family protein - ABC transporter ATP-binding protein - hypothetical protein - pyruvate kinase : hypothetical protein - hypothetical protein - response regulator transcription factor - DUF5685 family protein - aminotransferase class V-fold PLP-dependent enzyme - L-2-hydroxyglutarate oxidase - glycoside hydrolase family 15 protein. citrate/2-methylcitrate synthase - citrate/2-methylcitrate synthase - hypothetical protein - transposase - DNRLRE domain-containing protein - transposase - RHS repeat-associated core domain-containing protein : DDE-type integrase/transposase/recombinase - IS110 family transposase - FkbM family methyltransferase - serine/threonine-protein kinase - fibronectin type III domain-containing protein - bifunctional 2-polyprenyl-6-hydroxyphenol methylase/3-demethylubiquinol 3-O-methyltransferase UbiG - cupin domain-containing protein. class II glutamine amidotransferase - CoA transferase - phosphatase PAP2 family protein - alpha/beta fold hydrolase - HAD family phosphatase - lipase - ABC transporter substrate-binding protein : pre-mycofactocin synthase MftD - hypothetical protein - hypothetical protein - YceI family protein - helix-turn-helix domain-containing protein - mycofactocin radical SAM maturase - mycofactocin biosynthesis chaperone MftB. class II glutamine amidotransferase - glycoside hydrolase family 15 protein - MFS transporter - SHOCT domain-containing protein - ribonuclease BN - DUF2252 domain-containing protein - carboxymuconolactone decarboxylase family protein : hypothetical protein - diacylglycerol kinase family protein - hypothetical protein - MarR family winged helix-turn-helix transcriptional regulator - anthrone oxygenase family protein - helix-turn-helix domain-containing protein - IS630 family transposase. class II glutamine amidotransferase - LLM class F420-dependent oxidoreductase - hypothetical protein - STAS domain-containing protein - GAF and ANTAR domain-containing protein - ABC transporter ATP-binding protein - ABC transporter permease : FUSC family protein - hypothetical protein - MFS transporter - MarR family winged helix-turn-helix transcriptional regulator - tRNA-Ser - nuclear transport factor 2 family protein - helix-turn-helix domain-containing protein. class I SAM-dependent methyltransferase - choline/carnitine O-acyltransferase - DUF2461 domain-containing protein - thiamine pyrophosphate-requiring protein - phosphotransferase - LLM class F420-dependent oxidoreductase - ABC transporter ATP-binding protein/permease : hypothetical protein - hypothetical protein -

glycoside hydrolase family 26 protein - lysine--tRNA ligase - AraC family transcriptional regulator - DUF1772 domain-containing protein - helix-turn-helix domain-containing protein. class I SAM-dependent methyltransferase - hemolysin family protein - transporter associated domain-containing protein - hemolysin family protein - DUF4232 domain-containing protein - TetR/AcrR family transcriptional regulator - acetyl-CoA C-acyltransferase : DUF309 domain-containing protein - methionine adenosyltransferase - LLM class flavin-dependent oxidoreductase - amidohydrolase family protein - four-helix bundle copper-binding protein - alpha/beta hydrolase - peptidoglycan-binding protein. co-chaperone GroES - hypothetical protein - hypothetical protein - ATP-binding protein - lysylphosphatidylglycerol synthase transmembrane domain-containing protein - histidine phosphatase family protein - DUF3618 domain-containing protein : hypothetical protein - SCO family protein - DoxX family protein - thioredoxin domain-containing protein - thioredoxin-dependent thiol peroxidase - hypothetical protein - tRNA-Leu. conjugal transfer protein TraC - hypothetical protein - YbhB/YbcL family Raf kinase inhibitor-like protein - P-II family nitrogen regulator - DedA family protein - lipase family protein - IS1634 family transposase : TetR/AcrR family transcriptional regulator - organic hydroperoxide resistance protein - glycosyltransferase family 4 protein - class I SAM-dependent methyltransferase - hypothetical protein - class I SAM-dependent methyltransferase - peptide deformylase. copper resistance CopC/CopD family protein - YcnI family protein - response regulator transcription factor - HNH endonuclease signature motif containing protein - hypothetical protein - type II toxin-antitoxin system VapB family antitoxin - DUF2470 domain-containing protein : hypothetical protein - OmpA family protein - metal ABC transporter ATP-binding protein - metal ABC transporter permease - metal ABC transporter permease - pyridoxal phosphate-dependent aminotransferase - hypothetical protein. cystathionine beta-synthase - cystathionine gamma-synthase - hypothetical protein - carbohydrate kinase family protein - hypothetical protein - NADPH-dependent FMN reductase - hypothetical protein : hypothetical protein - isovaleryl-CoA dehydrogenase - GGDEF domain-containing protein - ribokinase - response regulator - histidine kinase - PAS domain S-box protein. cysteine desulfurase family protein - cytochrome c oxidase subunit II - cytochrome c oxidase subunit I - cytochrome c oxidase subunit 4 - IS1182 family transposase - anthranilate phosphoribosyltransferase - response regulator : Fur family transcriptional regulator - rubrerythrin family protein - DUF3501 family protein - toll/interleukin-1 receptor domain-containing protein - SseB family protein - hypothetical protein - heme-copper oxidase subunit III. cytochrome C - AIM24 family protein - CoA transferase - DUF4386 domain-containing protein - response regulator - bacterioferritin - superoxide dismutase : alpha/beta hydrolase - C4-dicarboxylate ABC transporter substrate-binding protein - HD domain-containing protein - hypothetical protein - glutamate ABC transporter substrate-binding protein - serine/threonine-protein kinase - GlsB/YeaQ/YmgE family stress response membrane protein. D-alanyl-D-alanine carboxypeptidase/D-alanyl-D-alanine-endopeptidase - inorganic diphosphatase - hypothetical protein - DUF6247 family protein - tRNA-Thr - response regulator transcription factor - stage 0 sporulation family protein : DNA polymerase III subunit delta' - dTMP kinase - type I DNA topoisomerase - sodium-translocating pyrophosphatase - ATP-binding protein - STAS domain-containing protein - DEAD/DEAH box helicase. diaminopimelate epimerase - VOC family protein - helix-turn-helix transcriptional regulator - GTPase HflX - transcriptional repressor LexA - LysM peptidoglycan-binding domain-containing protein - transcriptional regulator NrdR : vitamin B12-dependent ribonucleotide reductase - hypothetical protein - type II toxin-antitoxin system VapC family toxin - trehalase family glycosidase - RNA polymerase sigma factor -

acetoacetate--CoA ligase - acetoin utilization protein AcuC. dienelactone hydrolase family protein - CoA transferase - type II toxin-antitoxin system Phd/YefM family antitoxin - Txe/YoeB family addiction module toxin - DUF6326 family protein - response regulator transcription factor - sensor histidine kinase : tRNA-Leu - ANTAR domain-containing response regulator - branched-chain amino acid ABC transporter substrate-binding protein - branched-chain amino acid ABC transporter substrate-binding protein - PaaI family thioesterase - DNA polymerase I - putative bifunctional diguanylate cyclase/phosphodiesterase. diguanylate cyclase domain-containing protein - hypothetical protein - FtsX-like permease family protein - hypothetical protein - ABC transporter ATP-binding protein - PadR family transcriptional regulator - enoyl-CoA hydratase/isomerase family protein : class I SAM-dependent methyltransferase - hypothetical protein - HAF repeat-containing protein - hypothetical protein - hypothetical protein - hypothetical protein - WD40 repeat domain-containing serine/threonine protein kinase. DNA-3-methyladenine glycosylase - 16S ribosomal RNA - 23S ribosomal RNA - 5S ribosomal RNA - hypothetical protein - lipopolysaccharide assembly protein LapB - DUF1015 family protein : lipopolysaccharide assembly protein LapB - HAD-IIA family hydrolase - alkyl sulfatase C-terminal domain-containing protein - hypothetical protein - TlyA family RNA methyltransferase - NAD kinase - DNA repair protein RecN. DUF1003 domain-containing protein - D-alanyl-D-alanine carboxypeptidase family protein - YihY/virulence factor BrkB family protein - 2'-5' RNA ligase family protein - serine hydrolase - NAD(P)/FAD-dependent oxidoreductase - DUF3017 domain-containing protein : FHA domain-containing protein - hypothetical protein - bifunctional phosphoribosylaminoimidazolecarboxamide formyltransferase/IMP cyclohydrolase - phosphoribosylglycinamide formyltransferase - helix-turn-helix domain-containing protein - DUF6350 family protein - hypothetical protein. DUF2273 domain-containing protein - Asp23/Gls24 family envelope stress response protein - DUF6286 domain-containing protein - hypothetical protein - 50S ribosomal protein L25/general stress protein Ctc - aminoacyl-tRNA hydrolase - glycosyltransferase : glycosyltransferase family 2 protein - glycosyltransferase - tyrosine--tRNA ligase - S8 family serine peptidase - HAD family hydrolase - acyl carrier protein - serine O-acetyltransferase. DUF262 domain-containing protein - alkaline phosphatase PhoX - hypothetical protein - hypothetical protein - amidohydrolase family protein - hypothetical protein - CAP domain-containing protein : ATP-binding cassette domain-containing protein - ABC transporter substrate-binding protein - NADPH-dependent FMN reductase - redox-sensitive transcriptional activator SoxR - alkaline phosphatase PhoX - UDP-glucose--hexose-1-phosphate uridylyltransferase - galactokinase. DUF3090 family protein - SCO1664 family protein - cysteine--1-D-myo-inosityl 2-amino-2-deoxy-alpha-D-glucopyranoside ligase - DivIVA domain-containing protein - PAC2 family protein - HAD family hydrolase - type IV toxin-antitoxin system AbiEi family antitoxin domain-containing protein : ABC transporter permease - ABC transporter ATP-binding protein - PD-(D/E)XK nuclease family protein - site-2 protease family protein - tRNA (adenine-N1)-methyltransferase - ferredoxin - proteasome ATPase. DUF3117 domain-containing protein - M17 family metallopeptidase - hypothetical protein - hypothetical protein - O-methyltransferase - RNA polymerase sigma factor SigE - anti-sigma factor : hypothetical protein - S1C family serine protease - Sec-independent protein translocase protein TatB - nucleotidyltransferase family protein - Mrp/NBP35 family ATP-binding protein - MarR family winged helix-turn-helix transcriptional regulator - PHP domain-containing protein. DUF3159 domain-containing protein - OB-fold nucleic acid binding domain-containing protein - DUF3710 domain-containing protein - dUTP diphosphatase - DUF4235 domain-containing protein - gamma carbonic anhydrase family protein - sigma-70 family RNA polymerase sigma factor : NarK/NasA family nitrate transporter - valine--tRNA ligase - inositol monophosphatase family protein - LuxR family transcriptional regulator - hypothetical protein - non-hydrolyzing UDP-N-acetylglucosamine 2-epimerase - hypothetical protein. DUF4190 domain-containing protein - serine/threonine-protein kinase - carboxyl transferase domain-containing protein - triacylglycerol lipase - DNA helicase PcrA - threonine/serine dehydratase - glutamine-hydrolyzing GMP synthase : nitrate- and nitrite sensing domain-containing protein - roadblock/LC7 domain-containing protein - DUF742 domain-containing protein - ATP/GTP-binding protein - ABC transporter substrate-binding protein - hypothetical protein - hypothetical protein. DUF559 domain-containing protein - FHA domain-containing protein - DUF2993 domain-containing protein - SDR family oxidoreductase - FAD-binding oxidoreductase - TetR/AcrR family transcriptional regulator - glycerol-3-phosphate dehydrogenase/oxidase : sensor histidine kinase - acyl-CoA dehydrogenase family protein - acyl-CoA dehydrogenase family protein - glycerol-3-phosphate dehydrogenase/oxidase - GPGG-motif small membrane protein - NAD(P)/FAD-dependent oxidoreductase - phosphogluconate dehydrogenase (NAD(+)-dependent, decarboxylating). DUF6049 family protein - NUDIX hydrolase - CCA tRNA nucleotidyltransferase - MFS transporter - inositol-3-phosphate synthase - PadR family transcriptional regulator - DUF5318 family protein : transglycosylase domain-containing protein - glycosyltransferase 87 family protein - hypothetical protein - hypothetical protein - 30S ribosomal protein S6 - single-stranded DNA-binding protein - 30S ribosomal protein S18. DUF6113 family protein - flavin reductase family protein - transglutaminase-like domain-containing protein - ferredoxin - succinyldiaminopimelate transaminase - 2,3,4,5-tetrahydropyridine-2,6-dicarboxylate N-succinyltransferase - succinyl-diaminopimelate desuccinylase : sugar transferase - DUF4012 domain-containing protein - WecB/TagA/CpsF family glycosyltransferase - glycosyltransferase family 4 protein - glycosyltransferase - glycosyltransferase - glycosyltransferase family 1 protein. DUF86 domain-containing protein - nucleotidyltransferase family protein - hypothetical protein - protein kinase - NERD domain-containing protein - SAM-dependent methyltransferase - recombinase family protein : FtsK/SpoIIIE domain-containing protein - hypothetical protein - hypothetical protein - hypothetical protein - hypothetical protein - hypothetical protein - AlpA family transcriptional regulator. EAL domain-containing protein - VC0807 family protein - ABC transporter substrate-binding protein - ABC transporter ATP-binding protein - branched-chain amino acid ABC transporter permease/ATP-binding protein - ABC transporter substrate-binding protein - ABC transporter substrate-binding protein : GntR family transcriptional regulator - MFS transporter - hypothetical protein - PKD domain-containing protein - SDR family NAD(P)-dependent oxidoreductase - TauD/TfdA family dioxygenase - SDR family NAD(P)-dependent oxidoreductase. electron transfer flavoprotein subunit alpha/FixB family protein - ROK family protein - mycofactocin-coupled SDR family oxidoreductase - mycofactocin biosynthesis peptidyl-dipeptidase MftE - 16S ribosomal RNA - 23S ribosomal RNA - 5S ribosomal RNA : fused isobutyryl-CoA mutase/GTPase IcmF - MarR family winged helix-turn-helix transcriptional regulator - mycofactocin biosynthesis glycosyltransferase MftF - cysteine desulfurase family protein - tRNA 2-thiouridine(34) synthase MnmA - methionine synthase - NAD-dependent DNA ligase LigA. enoyl-CoA hydratase-related protein - SDR family NAD(P)-dependent oxidoreductase - enoyl-CoA hydratase-related protein - mycofactocin-coupled SDR family oxidoreductase - acyl-CoA thioesterase II - nitroreductase/quinone reductase family protein - oxidoreductase : N-6 DNA methylase - hypothetical protein - hypothetical protein - YdeI/OmpDassociated family protein - APC family permease - PadR family transcriptional regulator - cation transporter. excinuclease ABC subunit UvrC - RNase adapter RapZ - uridine diphosphate-N-acetylglucosamine-binding protein YvcK - DNA-binding protein WhiA - type I glyceraldehyde-3-phosphate dehydrogenase - phosphoglycerate kinase - triose-phosphate isomerase : preprotein translocase subunit SecG - ABC transporter substrate-binding protein - RNA polymerase-binding protein RbpA - serine/threonine-protein kinase - SHOCT domain-containing protein - 6-phosphogluconolactonase - glucose-6-phosphate dehydrogenase assembly protein OpcA. F0F1 ATP synthase subunit epsilon - type II toxin-antitoxin system PemK/MazF family toxin - hypothetical protein - RNA-guided endonuclease TnpB family protein - cob(I)yrinic acid a,c-diamide adenosyltransferase - protein kinase - DUF4158 domain-containing protein : LysR family transcriptional regulator - EthD family reductase - LytR C-terminal domain-containing protein - protein phosphatase 2C domain-containing protein - hypothetical protein - NUDIX hydrolase - UDP-N-acetylglucosamine 1-carboxyvinyltransferase. FAD-binding oxidoreductase - MFS transporter - thioesterase II family protein - LLM class flavin-dependent oxidoreductase - carboxyl transferase domain-containing protein - hypothetical protein - acyl-CoA carboxylase subunit beta : MMPL family transporter - LysR family transcriptional regulator - SDR family oxidoreductase - IS630 family transposase - helix-turn-helix domain-containing protein - hypothetical protein - AMP-binding protein. FAD-dependent monooxygenase - AfsR/SARP family transcriptional regulator - TetR/AcrR family transcriptional regulator - nuclear transport factor 2 family protein - PhnD/SsuA/transferrin family substrate-binding protein - ABC transporter ATP-binding protein - ABC transporter permease : LLM class flavin-dependent oxidoreductase - TetR/AcrR family transcriptional regulator - MFS transporter - AfsR/SARP family transcriptional regulator - 3-oxoacyl-ACP synthase III family protein - FAD-dependent monooxygenase - type I polyketide synthase. ferredoxin - helix-turn-helix domain-containing protein - alpha/beta hydrolase - glycosyltransferase family 2 protein - class I SAM-dependent methyltransferase - acyl-CoA dehydrogenase family protein - Fpg/Nei family DNA glycosylase : DUF1684 domain-containing protein - Rrf2 family transcriptional regulator - acyl-CoA dehydrogenase family protein - NtaA/DmoA family FMN-dependent monooxygenase - rhodanese-like domain-containing protein - LLM class flavin-dependent oxidoreductase - ABC transporter substrate-binding protein. Fic family protein - antitoxin VbhA family protein - Panacea domain-containing protein - type II toxin-antitoxin system RelE/ParE family toxin - CoA-binding protein - hypothetical protein - glycosyltransferase family 2 protein : DUF5999 family protein - bifunctional phosphatase PAP2/diacylglycerol kinase family protein - hypothetical protein - hypothetical protein - pyruvate dehydrogenase (acetyl-transferring), homodimeric type - secondary thiamine-phosphate synthase enzyme YjbQ - DUF3052 domain-containing protein. FxSxx-COOH system tetratricopeptide repeat protein - tRNA-Arg - IS3 family transposase - transposase - IS3 family transposase - IS5 family transposase - NADP-dependent oxidoreductase : hypothetical protein - acyltransferase family protein - acyltransferase family protein - hypothetical protein - hypothetical protein - hypothetical protein - hypothetical protein. FxSxx-COOH system tetratricopeptide repeat protein - tyrosine-protein kinase family protein - hypothetical protein - hypothetical protein - TIR domain-containing protein - toll/interleukin-1 receptor domain-containing protein - hypothetical protein : TerD family protein - DsbA family protein - hypothetical protein - WD40 repeat domain-containing protein - YkvA family protein - metallophosphoesterase - 4'-phosphopantetheinyl transferase. GAF domain-containing protein - mycoredoxin - ATP-dependent helicase - HRDC domain-containing protein -

glycerate kinase - WhiB family transcriptional regulator - alkaline phosphatase family protein : ABC1 kinase family protein - hypothetical protein - ThiF family adenylyltransferase - zinc-dependent metalloprotease - molybdenum cofactor biosynthesis protein MoaE - hypothetical protein - PDZ domain-containing protein. glucose-1-phosphate thymidylyltransferase - glycosyltransferase family 39 protein - LCP family protein - TIGR03089 family protein - UDP-glucose/GDP-mannose dehydrogenase family protein - carbonic anhydrase - GtrA family protein : TetR/AcrR family transcriptional regulator - hypothetical protein - hypothetical protein - hypothetical protein - hypothetical protein - PH domain-containing protein - biotin--[acetyl-CoA-carboxylase] ligase. glucose-6-phosphate isomerase - serine/threonine-protein kinase - FAD-binding oxidoreductase - hypothetical protein - NlpC/P60 family protein - hypothetical protein - helix-turn-helix domain-containing protein : hypothetical protein - DUF6112 family protein - hypothetical protein - hypothetical protein - SCO6880 family protein - conjugal transfer protein TraC - hypothetical protein. glutamate ABC transporter substrate-binding protein - serine/threonine-protein kinase - GlsB/YeaQ/YmgE family stress response membrane protein - ABC transporter ATP-binding protein - siderophore-interacting protein - dimethylargininase - PLP-dependent aminotransferase family protein : keywimysin-related RiPP - FAD-dependent monooxygenase - mycofactocin system FadH/OYE family oxidoreductase 2 - mycofactocin system transcriptional regulator - hypothetical protein - hypothetical protein - hypothetical protein. glutamine-hydrolyzing GMP synthase - nitrate- and nitrite sensing domain-containing protein - roadblock/LC7 domain-containing protein - DUF742 domain-containing protein - ATP/GTP-binding protein - ABC transporter substrate-binding protein - hypothetical protein : hypothetical protein - GuaB3 family IMP dehydrogenase-related protein - IMP dehydrogenase - DUF5319 domain-containing protein - WhiB family transcriptional regulator - chaperonin GroEL - co-chaperone GroES. glycoside hydrolase family 26 protein - uroporphyrinogen-III C-methyltransferase - nitrite reductase small subunit NirD - nitrite reductase large subunit NirB - uroporphyrinogen-III synthase - polysaccharide deacetylase family protein - peptide chain release factor 2 : hypothetical protein - pilus assembly protein TadG-related protein - SPFH domain-containing protein - IS4 family transposase - TetR/AcrR family transcriptional regulator - daunorubicin resistance protein DrrA family ABC transporter ATP-binding protein - ABC transporter permease. glycosyltransferase - chaperonin GroEL - hypothetical protein - cation diffusion facilitator family transporter - hypothetical protein - universal stress protein - SAM-dependent methyltransferase : hypothetical protein - TetR family transcriptional regulator - epoxide hydrolase family protein - phosphotransferase family protein - Nonspecific lipid-transfer protein - Zn-ribbon domain-containing OB-fold protein - hypothetical protein. glycosyltransferase family 2 protein - cellulase family glycosylhydrolase - glycosyltransferase - glycosyltransferase family 4 protein - hypothetical protein - hypothetical protein - lipopolysaccharide biosynthesis protein : glycosyltransferase family 2 protein - TIGR00730 family Rossman fold protein - TIGR00730 family Rossman fold protein - dihydropteroate synthase - DivIVA domain-containing protein - DUF3117 domain-containing protein - M17 family metallopeptidase. glycosyltransferase family 4 protein - condensation domain-containing protein - hypothetical protein - RyR domain-containing protein - hypothetical protein - DUF3367 domain-containing protein - dolichyl-phosphate beta-glucosyltransferase : ribose-5-phosphate isomerase - DsbA family protein - AI-2E family transporter - aminopeptidase N - DUF5130 family protein - hypothetical protein - 4-alpha-glucanotransferase. GNAT family N-acetyltransferase - dTDP-4-dehydrorhamnose 3,5-epimerase - amino acid ABC transporter ATP-binding protein - amino acid

ABC transporter permease - ABC transporter substrate-binding protein - NADP-dependent malic enzyme - hypothetical protein : Zn-dependent alcohol dehydrogenase - DUF6104 family protein - multifunctional oxoglutarate decarboxylase/oxoglutarate dehydrogenase thiamine pyrophosphate-binding subunit/dihydrolipoyllysine-residue succinyltransferase subunit - hypothetical protein - hypothetical protein - thiamine-phosphate kinase - acetamidase/formamidase family protein. GNAT family N-acetyltransferase - GNAT family N-acetyltransferase - histidine kinase - response regulator transcription factor - DUF4097 family beta strand repeat-containing protein - transposase - lytic transglycosylase domain-containing protein : hypothetical protein - DNRLRE domain-containing protein - hypothetical protein - hypothetical protein - FtsX-like permease family protein - ABC transporter ATP-binding protein - PadR family transcriptional regulator. GNAT family N-acetyltransferase - vanadium-dependent haloperoxidase - ABC transporter permease - ABC transporter permease - ATP-binding cassette domain-containing protein - SpoIIE family protein phosphatase - ATP-binding protein : WhiB family transcriptional regulator - NAD(P)/FAD-dependent oxidoreductase - YafY family protein - FxSxx-COOH system tetratricopeptide repeat protein - FHA domain-containing protein - VWA domain-containing protein - protein phosphatase 2C domain-containing protein. GntR family transcriptional regulator - class IV adenylate cyclase - ADP-ribosylglycohydrolase family protein - DarT ssDNA thymidine ADP-ribosyltransferase family protein - 3'-5' exonuclease - UvrD-helicase domain-containing protein - hypothetical protein : hypothetical protein - IS6 family transposase - hypothetical protein - recombinase family protein - IS6 family transposase - type I-E CRISPR-associated endoribonuclease Cas2e - type I-E CRISPR-associated endonuclease Cas1e. GntR family transcriptional regulator - MFS transporter - hypothetical protein - PKD domain-containing protein - SDR family NAD(P)-dependent oxidoreductase - TauD/TfdA family dioxygenase - SDR family NAD(P)-dependent oxidoreductase : class II aldolase/adducin family protein - hypothetical protein - phosphotransferase - hypothetical protein - SAM-dependent methyltransferase - ABC transporter substrate-binding protein - phosphotransferase family protein. HAD family hydrolase - type IV toxin-antitoxin system AbiEi family antitoxin domain-containing protein - ABC transporter permease - ABC transporter ATP-binding protein - PD-(D/E)XK nuclease family protein - site-2 protease family protein - tRNA (adenine-N1)-methyltransferase : ferredoxin - proteasome ATPase - depupylase/deamidase Dop - ubiquitin-like protein Pup - proteasome subunit beta - proteasome subunit alpha - dienelactone hydrolase family protein. HD domain-containing protein - DsbA family oxidoreductase - hypothetical protein - toll/interleukin-1 receptor domain-containing protein - Sec-independent protein translocase subunit TatA - GlsB/YeaQ/YmgE family stress response membrane protein - TIGR03620 family F420-dependent LLM class oxidoreductase : CrcB family protein - CrcB family protein - DUF2599 domain-containing protein - sigma-70 family RNA polymerase sigma factor - SigE family RNA polymerase sigma factor - hypothetical protein - TetR/AcrR family transcriptional regulator. HEAT repeat domain-containing protein - hypothetical protein - hypothetical protein - hypothetical protein - integrase core domain-containing protein - IS256 family transposase - integrase : RHS repeat-associated core domain-containing protein - DNRLRE domain-containing protein - lytic transglycosylase domain-containing protein - hypothetical protein - heme-binding beta-barrel domain-containing protein - pyruvate:ferredoxin (flavodoxin) oxidoreductase - NAD(+) synthase. helix-turn-helix domain-containing protein - amino acid ABC transporter ATP-binding protein - amino acid ABC transporter permease - transporter substrate-binding domain-containing protein - citrate/2-methylcitrate synthase - citrate/2-methylcitrate synthase - hypothetical protein :

transposase - DNRLRE domain-containing protein - transposase - RHS repeat-associated core domain-containing protein - DDE-type integrase/transposase/recombinase - IS110 family transposase - FkbM family methyltransferase. helix-turn-helix domain-containing protein - DUF6350 family protein - hypothetical protein - succinate--CoA ligase subunit alpha - ADP-forming succinate--CoA ligase subunit beta - DUF4190 domain-containing protein - serine/threonine-protein kinase : carboxyl transferase domain-containing protein - triacylglycerol lipase - DNA helicase PcrA - threonine/serine dehydratase - glutamine-hydrolyzing GMP synthase - nitrate- and nitrite sensing domain-containing protein - roadblock/LC7 domain-containing protein. Hsp70 family protein - DUF6461 domain-containing protein - AAA family ATPase - hypothetical protein - hypothetical protein - hypothetical protein - hypothetical protein : hypothetical protein - hypothetical protein - hypothetical protein - hypothetical protein - hypothetical protein - YbaB/EbfC family nucleoid-associated protein - hypothetical protein. hypothetical protein - aldehyde dehydrogenase family protein - PP2C family protein-serine/threonine phosphatase - DUF427 domain-containing protein - TetR/AcrR family transcriptional regulator - ABC transporter permease - GAF domain-containing protein : universal stress protein - MFS transporter - pyridoxamine 5'-phosphate oxidase family protein - TetR/AcrR family transcriptional regulator - hemerythrin domain-containing protein - MarR family transcriptional regulator - RNA-binding S4 domain-containing protein. hypothetical protein - alkane 1-monooxygenase - hypothetical protein - AEC family transporter - 3-hydroxybutyryl-CoA dehydrogenase - acyl-CoA dehydrogenase family protein - LCP family protein : 2-hydroxyacid dehydrogenase - NAD-dependent epimerase/dehydratase family protein - hypothetical protein - PAC2 family protein - hypothetical protein - sensor histidine kinase - DHA2 family efflux MFS transporter permease subunit. hypothetical protein - Asp-tRNA(Asn)/Glu-tRNA(Gln) amidotransferase subunit GatB - penicillin-binding transpeptidase domain-containing protein - acetolactate synthase large subunit - acetolactate synthase small subunit - ketol-acid reductoisomerase - phosphoglycerate dehydrogenase : septum formation family protein - 2-isopropylmalate synthase - DUF1059 domain-containing protein - EAL domain-containing protein - 3-isopropylmalate dehydrogenase - branched-chain amino acid transaminase - cyclic nucleotide-binding/CBS domain-containing protein. hypothetical protein - glycoside hydrolase family 26 protein - SDR family oxidoreductase - hypothetical protein - 2-C-methyl-D-erythritol 4-phosphate cytidylyltransferase - DUF4910 domain-containing protein - sugar phosphate nucleotidyltransferase : glycoside hydrolase family 26 protein - MerR family transcriptional regulator - transcription-repair coupling factor - SurA N-terminal domain-containing protein - nucleoside triphosphate pyrophosphohydrolase - S49 family peptidase - S49 family peptidase. hypothetical protein - HAD hydrolase-like protein - FKBP-type peptidyl-prolyl cis-trans isomerase - macrolide family glycosyltransferase - TIGR03620 family F420-dependent LLM class oxidoreductase - SDR family NAD(P)-dependent oxidoreductase - HAD family hydrolase : hypothetical protein - ABC transporter ATP-binding protein - nucleotide disphospho-sugar-binding domain-containing protein - alpha/beta fold hydrolase - acyl carrier protein - acyltransferase domain-containing protein - type I polyketide synthase. hypothetical protein - hypothetical protein - ABC transporter permease - ABC transporter ATP-binding protein - ABC transporter ATP-binding protein - ABC transporter substrate-binding protein - hypothetical protein : hypothetical protein - DUF2752 domain-containing protein - RNA methyltransferase - 8-oxoguanine deaminase - bacterioferritin - aldolase/citrate lyase family protein - anion permease. hypothetical protein - hypothetical protein - aminopeptidase N - DUF6529 family protein - Rieske (2Fe-2S) protein - FHA domain-containing protein - hypothetical protein : type II toxin-antitoxin system RelE/ParE family toxin - type II toxin-antitoxin system Phd/YefM family antitoxin - glycosyltransferase family 2 protein - SCO family protein - hypothetical protein - PA14 domain-containing protein - glycosyltransferase family 2 protein. hypothetical protein - hypothetical protein - CHAT domain-containing protein - hypothetical protein - hypothetical protein - VapC toxin family PIN domain ribonuclease - IS4 family transposase : group II intron reverse transcriptase/maturase - IS3 family transposase - group II intron maturase-specific domain-containing protein - hypothetical protein - hypothetical protein - toll/interleukin-1 receptor domain-containing protein - tetratricopeptide repeat-containing protein. hypothetical protein - hypothetical protein - DUF2795 domain-containing protein - Rieske (2Fe-2S) protein - metallophosphoesterase - BON domain-containing protein - hypothetical protein : MarR family winged helix-turn-helix transcriptional regulator - CapA family protein - DoxX family protein - alpha/beta fold hydrolase - NTP transferase domain-containing protein - GNAT family N-acetyltransferase - substrate-binding domain-containing protein. hypothetical protein - hypothetical protein - DUF86 domain-containing protein - nucleotidyltransferase family protein - helix-turn-helix domain-containing protein - SDR family oxidoreductase - hypothetical protein : hypothetical protein - VOC family protein - YoaK family protein - class I SAM-dependent methyltransferase - SsrA-binding protein SmpB - permease-like cell division protein FtsX - cell division ATP-binding protein FtsE. hypothetical protein - hypothetical protein - Hint domain-containing protein - hypothetical protein - Imm52 family immunity protein - polymorphic toxin-type HINT domain-containing protein - DNRLRE domain-containing protein : hypothetical protein - aldehyde dehydrogenase family protein - hypothetical protein - IS3 family transposase - toll/interleukin-1 receptor domain-containing protein - IS701 family transposase - hypothetical protein. hypothetical protein - hypothetical protein - Hsp70 family protein - DUF6461 domain-containing protein - AAA family ATPase - hypothetical protein - hypothetical protein : hypothetical protein - hypothetical protein - hypothetical protein - hypothetical protein - hypothetical protein - hypothetical protein - hypothetical protein. hypothetical protein - hypothetical protein - hypothetical protein - DUF6531 domain-containing protein - hypothetical protein - DUF6531 domain-containing protein - hypothetical protein : FtsK/SpoIIIE domain-containing protein - nuclear transport factor 2 family protein - NAD(P)/FAD-dependent oxidoreductase - hypothetical protein - YhjD/YihY/BrkB family envelope integrity protein - hypothetical protein - HypC/HybG/HupF family hydrogenase formation chaperone. hypothetical protein - hypothetical protein - hypothetical protein - hypothetical protein - hypothetical protein - hypothetical protein - hypothetical protein : DUF2255 family protein - pentapeptide repeat-containing protein - hypothetical protein - hypothetical protein - DUF305 domain-containing protein - GGDEF domain-containing protein - FtsK/SpoIIIE domain-containing protein. hypothetical protein - hypothetical protein - hypothetical protein - hypothetical protein - hypothetical protein - TIR domain-containing protein - helix-turn-helix domain-containing protein : alpha/beta fold hydrolase - hypothetical protein - holin - LuxR family transcriptional regulator - tetratricopeptide repeat protein - CHAT domain-containing protein - hypothetical protein. hypothetical protein - hypothetical protein - hypothetical protein - integrase core domain-containing protein - IS256 family transposase - integrase - RHS repeat-associated core domain-containing protein : DNRLRE domain-containing protein - lytic transglycosylase domain-containing protein - hypothetical protein - heme-binding beta-barrel domain-containing protein - pyruvate:ferredoxin (flavodoxin) oxidoreductase - NAD(+) synthase - TetR family transcriptional regulator. hypothetical protein - hypothetical protein - hypothetical protein - metallophosphoesterase family protein - sulfite exporter TauE/SafE family protein - ATP-dependent Clp protease ATP-binding subunit - (2Fe-2S)-binding protein : type II toxin-antitoxin system HicA family toxin - type II toxin-antitoxin system HicB family antitoxin - SGNH/GDSL hydrolase family protein - type III pantothenate kinase - carboxylating nicotinate-nucleotide diphosphorylase - L-aspartate oxidase - aspartate 1-decarboxylase. hypothetical protein - hypothetical protein - IS1380 family transposase - hypothetical protein - hypothetical protein - acyl-CoA dehydrogenase family protein - TetR/AcrR family transcriptional regulator : peptidoglycan-binding protein - hypothetical protein - DUF2889 domain-containing protein - urease accessory protein UreG - tRNA (adenosine(37)-N6)-threonylcarbamoyltransferase complex transferase subunit TsaD - Fur family transcriptional regulator - peptide deformylase. hypothetical protein - hypothetical protein - OB-fold nucleic acid binding domain-containing protein - RNA-guided endonuclease TnpB family protein - PHP domain-containing protein - SAV_6107 family HEPN domain-containing protein - hypothetical protein : BTAD domain-containing putative transcriptional regulator - spheroidene monooxygenase - NAD(P)-dependent oxidoreductase - FAD-binding oxidoreductase - GNAT family N-acetyltransferase - bisanhydrobacterioruberin hydratase CruF - glycosyltransferase family 2 protein. hypothetical protein - hypothetical protein - RDD family protein - hypothetical protein - hypothetical protein - IS701 family transposase - helix-turn-helix domain-containing protein : recombinase family protein - hypothetical protein - tRNA-Ile - DUF3566 domain-containing protein - DNA gyrase subunit A - DNA topoisomerase (ATP-hydrolyzing) subunit B - DciA family protein. hypothetical protein - hypothetical protein - thiamine-phosphate kinase - acetamidase/formamidase family protein - AAA domain-containing protein - UvrD-helicase domain-containing protein - class I SAM-dependent methyltransferase : hypothetical protein - aminotransferase class I/II-fold pyridoxal phosphate-dependent enzyme - leucyl/phenylalanyl-tRNA--protein transferase - bifunctional diguanylate cyclase/phosphodiesterase - hypothetical protein - M56 family metallopeptidase - hypothetical protein. hypothetical protein - IS1634 family transposase - methyltransferase domain-containing protein - hypothetical protein - excalibur calcium-binding domain-containing protein - ABC transporter substrate-binding protein - metallophosphoesterase : TetR family transcriptional regulator - acyl-CoA dehydrogenase - MBL fold metallo-hydrolase - SDR family NAD(P)-dependent oxidoreductase - thiolase family protein - aldo/keto reductase - SDR family NAD(P)-dependent oxidoreductase. hypothetical protein - LysM peptidoglycan-binding domain-containing protein - hypothetical protein - MFS transporter - metallophosphoesterase - dihydrofolate reductase family protein - DMT family transporter : enoyl-CoA hydratase-related protein - SDR family NAD(P)-dependent oxidoreductase - glycoside hydrolase family 3 protein - SAM-dependent methyltransferase - MarR family winged helix-turn-helix transcriptional regulator - LLM class flavin-dependent oxidoreductase - hypothetical protein. hypothetical protein - NADP-dependent oxidoreductase - hypothetical protein - ABC transporter ATP-binding protein - ABC transporter permease - YhjD/YihY/BrkB family envelope integrity protein - hypothetical protein : SRPBCC domain-containing protein - hypothetical protein - ATP-binding protein - SDR family NAD(P)-dependent oxidoreductase - type I polyketide synthase - PfaD family polyunsaturated fatty acid/polyketide biosynthesis protein - SDR family oxidoreductase. hypothetical protein - PHP domain-containing protein - hypothetical protein - hypothetical protein - SAM-dependent methyltransferase - hypothetical protein - hypothetical protein : helix-turn-helix transcriptional regulator - GNAT family N-acetyltransferase - 3'-5' exonuclease - hypothetical protein - nucleoside-diphosphate kinase - thymidylate synthase - hypothetical protein. hypothetical protein - RNA-guided endonuclease TnpB family protein - cob(I)yrinic acid a,c-diamide adenosyltransferase - protein kinase - DUF4158 domain-containing protein - LysR family transcriptional regulator - EthD family reductase : LytR C-terminal domain-containing protein - protein phosphatase 2C domain-containing protein - hypothetical protein - NUDIX hydrolase - UDP-N-acetylglucosamine 1-carboxyvinyltransferase - hypothetical protein - STAS domain-containing protein. hypothetical protein - RNA polymerase sigma factor - amidohydrolase family protein - hypothetical protein - ABC transporter permease subunit - ABC transporter ATP-binding protein - ABC transporter substrate-binding protein : nitrate- and nitrite sensing domain-containing protein - hypothetical protein - roadblock/LC7 domain-containing protein - DUF742 domain-containing protein - hypothetical protein - ABC transporter substrate-binding protein - iron ABC transporter permease. hypothetical protein - SCO2521 family protein - SCO2522 family protein - SCO2523 family variant P-loop protein - SCO2524 family protein - SCO2525 family SAM-dependent methyltransferase - N-acetyltransferase : mechanosensitive ion channel family protein - trans-aconitate 2-methyltransferase - GlsB/YeaQ/YmgE family stress response membrane protein - hypothetical protein - amidase - PP2C family protein-serine/threonine phosphatase - DUF4236 domain-containing protein. hypothetical protein - Sec-independent protein translocase subunit TatA - twin-arginine translocase subunit TatC - RNA helicase - cobalamin biosynthesis protein - cobyric acid synthase - hypothetical protein : SDR family NAD(P)-dependent oxidoreductase - phosphotransferase family protein - ferredoxin - cytochrome P450 - mycofactocin-coupled SDR family oxidoreductase - aldehyde dehydrogenase - TetR/AcrR family transcriptional regulator. hypothetical protein - TetR/AcrR family transcriptional regulator - SDR family oxidoreductase - class I SAM-dependent methyltransferase - hypothetical protein - MFS transporter - WS/DGAT domain-containing protein : SDR family NAD(P)-dependent oxidoreductase - dihydrolipoyl dehydrogenase - hypothetical protein - SDR family NAD(P)-dependent oxidoreductase - MarR family winged helix-turn-helix transcriptional regulator - MFS transporter - Zn-ribbon domain-containing OB-fold protein. hypothetical protein - transposase - winged helix-turn-helix domain-containing protein - IS701 family transposase - IS256 family transposase - hypothetical protein - IS6 family transposase : DUF4037 domain-containing protein - hypothetical protein - nucleotidyltransferase family protein - DUF86 domain-containing protein - hypothetical protein - hypothetical protein - toll/interleukin-1 receptor domain-containing protein. hypothetical protein - trehalose-6-phosphate synthase - hypothetical protein - hypothetical protein - FtsK/SpoIIIE domain-containing protein - hypothetical protein - MerR family transcriptional regulator : bifunctional nuclease family protein - MerR family transcriptional regulator - FHA domain-containing protein - glycine cleavage system protein GcvH - DUF881 domain-containing protein - DUF1290 domain-containing protein - DUF881 domain-containing protein. hypothetical protein - vWA domain-containing protein - hypothetical protein - hypothetical protein - hypothetical protein - DUF6531 domain-containing protein - hypothetical protein : DUF6531 domain-containing protein - hypothetical protein - FtsK/SpoIIIE domain-containing protein - nuclear transport factor 2 family protein - NAD(P)/FAD-dependent oxidoreductase - hypothetical protein - YhjD/YihY/BrkB family envelope integrity protein. hypothetical protein - WD40 repeat domain-containing serine/threonine protein kinase - substrate-binding domain-containing protein - class I SAM-dependent methyltransferase - hemolysin family protein - transporter associated domain-containing protein - hemolysin family protein : DUF4232 domain-containing protein - TetR/AcrR family transcriptional regulator - acetyl-CoA C-acyltransferase - DUF309 domain-containing protein - methionine adenosyltransferase - LLM class flavin-dependent oxidoreductase - amidohydrolase family protein. hypothetical protein - YceI family protein - helix-turn-helix domain-containing protein - mycofactocin radical SAM maturase - mycofactocin biosynthesis chaperone MftB - mycofactocin precursor MftA - zinc ribbon domain-containing protein : TetR/AcrR family transcriptional regulator - 3-keto-5-aminohexanoate cleavage protein - polyprenyl diphosphate synthase - TetR/AcrR family transcriptional regulator - TetR/AcrR family transcriptional regulator - acyl-CoA dehydrogenase family protein - SDR family NAD(P)-dependent oxidoreductase. hypothetical protein - YdcF family protein - GntR family transcriptional regulator - hypothetical protein - hypothetical protein - hypothetical protein - DUF222 domain-containing protein : acetyl-CoA hydrolase/transferase C-terminal domain-containing protein - thiolase family protein - Zn-ribbon domain-containing OB-fold protein - acyl-CoA dehydrogenase family protein - acyl-CoA dehydrogenase family protein - Rieske 2Fe-2S domain-containing protein - amidohydrolase family protein. IclR family transcriptional regulator - hypothetical protein - NAD(P)H-binding protein - LysR family transcriptional regulator - helicase-associated domain-containing protein - hypothetical protein - CocE/NonD family hydrolase : hypothetical protein - AAA family ATPase - hypothetical protein - EndoU domain-containing protein - NADPH-dependent FMN reductase - pentapeptide repeat-containing protein - TIR domain-containing protein. inorganic polyphosphate kinase - SPFH domain-containing protein - NUDIX domain-containing protein - DMT family transporter - glycosyl hydrolase - NAD(P)-dependent oxidoreductase - DegT/DnrJ/EryC1/StrS aminotransferase family protein : metallophosphoesterase - MSMEG_0572/Sll0783 family nitrogen starvation response protein - alpha/beta fold hydrolase - YihY/virulence factor BrkB family protein - phage holin family protein - DUF3618 domain-containing protein - DUF4407 domain-containing protein. integrase core domain-containing protein - IS256 family transposase - integrase - RHS repeat-associated core domain-containing protein - DNRLRE domain-containing protein - lytic transglycosylase domain-containing protein - hypothetical protein : heme-binding beta-barrel domain-containing protein - pyruvate:ferredoxin (flavodoxin) oxidoreductase - NAD(+) synthase - TetR family transcriptional regulator - MFS transporter - MFS transporter - DUF4062 domain-containing protein. iron ABC transporter permease - ABC transporter ATP-binding protein - proline dehydrogenase family protein - L-glutamate gamma-semialdehyde dehydrogenase - hypothetical protein - DUF5134 domain-containing protein - cytochrome C : AIM24 family protein - CoA transferase - DUF4386 domain-containing protein - response regulator - bacterioferritin - superoxide dismutase - alpha/beta hydrolase. iron uptake system protein EfeO - iron uptake transporter deferrochelatase/peroxidase subunit - hypothetical protein - HIT family protein - NUDIX hydrolase - hypothetical protein - hypothetical protein : LysR family transcriptional regulator - DMT family transporter - SAM-dependent methyltransferase - C4-dicarboxylate transporter DctA - transposase - response regulator - aldehyde dehydrogenase family protein. IS4 family transposase - ATP-binding cassette domain-containing protein - serine/threonine-protein kinase - family 2 encapsulin nanocompartment cargo protein terpene cyclase - family 2B encapsulin nanocompartment shell protein - isopentenyl-diphosphate Delta-isomerase - hemerythrin domain-containing protein : DUF3291 domain-containing protein - superoxide dismutase family protein - hypothetical protein - serine hydrolase - methyltransferase - iron-containing redox enzyme family protein - CYTH domain-containing protein. IS6 family transposase - ZIP family metal transporter - DUF86 domain-containing protein - TetR/AcrR family transcriptional regulator - hypothetical protein - ABC transporter substrate-binding protein - hypothetical protein : hypothetical protein - hypothetical protein - Hint domain-containing protein - hypothetical protein - Imm52 family immunity protein - polymorphic toxin-type HINT domain-containing protein - DNRLRE domain-containing protein. lipase - ABC transporter ATP-binding protein - IS4 family transposase - ATP-binding cassette domain-containing protein - serine/threonine-protein kinase - family 2 encapsulin nanocompartment cargo protein terpene cyclase - family 2B encapsulin nanocompartment shell protein : isopentenyl-diphosphate Delta-isomerase - hemerythrin domain-containing protein - DUF3291 domain-containing protein - superoxide dismutase family protein - hypothetical protein - serine hydrolase - methyltransferase. LLM class flavin-dependent oxidoreductase - FxSxx-COOH system tetratricopeptide repeat protein - serine/threonine-protein kinase - ice-binding family protein - helix-turn-helix domain-containing protein - NAD(P)-dependent oxidoreductase - Clp protease N-terminal domain-containing protein : hypothetical protein - CopG family transcriptional regulator - type II toxin-antitoxin system death-on-curing family toxin - hypothetical protein - acyltransferase - oxidoreductase - LysR family transcriptional regulator. LLM class flavin-dependent oxidoreductase - LLM class flavin-dependent oxidoreductase - ABC transporter ATP-binding protein - ABC transporter ATP-binding protein - ABC transporter permease - ABC transporter permease - ABC transporter substrate-binding protein : hypothetical protein - hypothetical protein - hypothetical protein - hypothetical protein - hypothetical protein - TIR domain-containing protein - helix-turn-helix domain-containing protein. LuxR C-terminal-related transcriptional regulator - 1-acyl-sn-glycerol-3-phosphate acyltransferase - MMPL family transporter - ChaB family protein - N-acetylmuramoyl-L-alanine amidase - trans-aconitate 2-methyltransferase - zinc-binding dehydrogenase : NADP-dependent oxidoreductase - IS701 family transposase - helix-turn-helix domain-containing protein - carbonic anhydrase - dienelactone hydrolase family protein - LLM class flavin-dependent oxidoreductase - NtaA/DmoA family FMN-dependent monooxygenase. LysM peptidoglycan-binding domain-containing protein - Rv3235 family protein - preprotein translocase subunit SecA - GNAT family N-acetyltransferase - response regulator transcription factor - ribosome hibernation-promoting factor, HPF/YfiA family - ComF family protein : hypothetical protein - DUF4129 domain-containing protein - DUF4350 domain-containing protein - AAA family ATPase - DUF58 domain-containing protein - DUF5990 family protein - nitroreductase family deazaflavin-dependent oxidoreductase. LysR family transcriptional regulator - FUSC family protein - DUF3696 domain-containing protein - hypothetical protein - DUF262 domain-containing protein - alkaline phosphatase PhoX - hypothetical protein : hypothetical protein - amidohydrolase family protein - hypothetical protein - CAP domain-containing protein - ATP-binding cassette domain-containing protein - ABC transporter substrate-binding protein - NADPH-dependent FMN reductase. LysR family transcriptional regulator - NAD(P)H-binding protein - Rieske 2Fe-2S domain-containing protein - SRPBCC family protein - antitoxin - EAL domain-containing protein - VC0807 family protein : ABC transporter substrate-binding protein - ABC transporter ATP-binding protein - branched-chain amino acid ABC transporter permease/ATP-binding protein - ABC transporter substrate-binding protein - ABC transporter substrate-binding protein - GntR family transcriptional regulator - MFS transporter. MBL fold metallo-hydrolase - excinuclease ABC subunit UvrA - SsgA family sporulation/cell division regulator - excinuclease ABC subunit UvrC - RNase adapter RapZ - uridine diphosphate-N-acetylglucosamine-binding protein YvcK - DNA-binding protein WhiA : type I glyceraldehyde-3-phosphate dehydrogenase - phosphoglycerate kinase - triose-phosphate isomerase - preprotein translocase subunit SecG - ABC transporter substrate-binding protein - RNA polymerase-binding protein RbpA - serine/threonine-protein kinase. MBL fold metallo-hydrolase - GlsB/YeaQ/YmgE family stress response membrane protein - serine/threonine-protein kinase - lipopolysaccharide assembly protein LapA domain-containing protein - hypothetical protein - protein kinase - hypothetical protein : hypothetical protein - AIPR family protein - IS607 family element RNA-guided endonuclease TnpB - IS607 family transposase - AIPR family protein - class I SAM-dependent DNA methyltransferase - restriction endonuclease subunit S. MBL fold metallo-hydrolase - ribonuclease PH - RdgB/HAM1 family non-canonical purine NTP pyrophosphatase - PLP-dependent cysteine synthase family protein - MoaD/ThiS family protein - Mov34/MPN/PAD-1 family protein - DUF2017 domain-containing protein : ATP-dependent Clp protease adapter ClpS - NUDIX domain-containing protein - nicotinate phosphoribosyltransferase - hypothetical protein - helix-turn-helix domain-containing protein - NAD-dependent epimerase/dehydratase family protein - hypothetical protein. MbtH family protein - ABC transporter ATP-binding protein - ABC transporter ATP-binding protein - iron chelate uptake ABC transporter family permease subunit - iron ABC transporter permease - lysine N(6)-hydroxylase/L-ornithine N(5)-oxygenase family protein - iron-siderophore ABC transporter substrate-binding protein : ABC transporter ATP-binding protein - hypothetical protein - ABC transporter ATP-binding protein - amino acid adenylation domain-containing protein - dihydrofolate reductase family protein - VOC family protein - rifampin monooxygenase. metallophosphoesterase - dihydrofolate reductase family protein - DMT family transporter - enoyl-CoA hydratase-related protein - SDR family NAD(P)-dependent oxidoreductase - glycoside hydrolase family 3 protein - SAM-dependent methyltransferase : MarR family winged helix-turn-helix transcriptional regulator - LLM class flavin-dependent oxidoreductase - hypothetical protein - HEAT repeat domain-containing protein - TetR/AcrR family transcriptional regulator - phosphotransferase family protein - effector-associated domain EAD1-containing protein. methionyl-tRNA formyltransferase - RsmB/NOP family class I SAM-dependent RNA methyltransferase - ribulose-phosphate 3-epimerase - dCMP deaminase - riboflavin synthase - bifunctional 3,4-dihydroxy-2-butanone-4-phosphate synthase/GTP cyclohydrolase II - 6,7-dimethyl-8-ribityllumazine synthase : hypothetical protein - SseB family protein - translation initiation factor IF-3 - 50S ribosomal protein L35 - 50S ribosomal protein L20 - TrmH family RNA methyltransferase - HAMP domain-containing sensor histidine kinase. MFS transporter - alpha/beta fold hydrolase - DUF2889 domain-containing protein - TetR/AcrR family transcriptional regulator - hypothetical protein - glycosyltransferase family 39 protein - dolichyl-phosphate beta-glucosyltransferase : IS256 family transposase - tetratricopeptide repeat protein - IS66 family transposase - threonine/serine dehydratase - molecular chaperone HtpG - hypothetical protein - IS1634 family transposase. MFS transporter - hypothetical protein - response regulator transcription factor - GAF domain-containing sensor histidine kinase - hypothetical protein - cysteine dioxygenase family protein - rhodanese-like domain-containing protein : WhiB family transcriptional regulator - histidine phosphatase family protein - hypothetical protein - trehalose-6-phosphate synthase - hypothetical protein - hypothetical protein - FtsK/SpoIIIE domain-containing protein. MFS transporter - SHOCT domain-containing protein - ribonuclease BN - DUF2252 domain-containing protein - carboxymuconolactone decarboxylase family protein - hypothetical protein - diacylglycerol kinase family protein : hypothetical protein - MarR family winged helix-turn-helix transcriptional regulator - anthrone oxygenase family protein - helix-turn-helix domain-containing protein - IS630 family transposase - helix-turn-helix domain-containing protein - hypothetical protein. MMPL family transporter -

phosphoribosylaminoimidazolesuccinocarboxamide synthase - hypothetical protein - hypothetical protein - adenylosuccinate lyase - 5-(carboxyamino)imidazole ribonucleotide mutase - phosphoribosylamine--glycine ligase : hypothetical protein - adenylosuccinate synthase - DUF3151 domain-containing protein - transglycosylase family protein - transglycosylase family protein - class II fructose-bisphosphate aldolase - DedA family protein. molecular chaperone DnaK - nuclease-related domain-containing protein - (Fe-S)-binding protein - dCTP deaminase - beta-phosphoglucomutase family hydrolase - glycoside hydrolase family 65 protein - tRNA-Gly : pentapeptide repeat-containing protein - carbohydrate kinase family protein - glycosyltransferase family 4 protein - class II glutamine amidotransferase - LLM class F420-dependent oxidoreductase - hypothetical protein - STAS domain-containing protein. molybdopterin-binding protein - acyl-CoA dehydrogenase family protein - helix-turn-helix domain-containing protein - RNA-guided endonuclease InsQ/TnpB family protein - nuclear transport factor 2 family protein - cytochrome P450 - TIGR01458 family HAD-type hydrolase : arsenate reductase ArsC - polyketide cyclase - hypothetical protein - MOSC domain-containing protein - enoyl-CoA hydratase - acyl-CoA synthetase - bifunctional diguanylate cyclase/phosphodiesterase. mycofactocin system FadH/OYE family oxidoreductase 2 - mycofactocin system transcriptional regulator - hypothetical protein - hypothetical protein - hypothetical protein - cation-translocating P-type ATPase - hypothetical protein : undecaprenyl-diphosphate phosphatase - cation-translocating P-type ATPase - Na+/H+ antiporter NhaA - cytochrome c oxidase assembly protein - hypothetical protein - vWA domain-containing protein - hypothetical protein. NAD-dependent protein deacetylase - GNAT family N-acetyltransferase - AlkA N-terminal domain-containing protein - methylated-DNA--[protein]-cysteine S-methyltransferase - 2OG-Fe(II) oxygenase - UBP-type zinc finger domain-containing protein - multicopper oxidase family protein : RtcB family protein - hypothetical protein - thiamine pyrophosphate-dependent dehydrogenase E1 component subunit alpha - aldehyde dehydrogenase family protein - LppP/LprE family lipoprotein - NAD(P)-binding oxidoreductase - methylmalonyl-CoA epimerase. Na+/H+ antiporter NhaA - hypothetical protein - hypothetical protein - hypothetical protein - tRNA-Ala - histidine phosphatase family protein - ribosome silencing factor : nicotinate-nucleotide adenylyltransferase - phosphoglucomutase (alpha-D-glucose-1,6-bisphosphate-dependent) - VWA domain-containing protein - MoxR family ATPase - glutamate-5-semialdehyde dehydrogenase - exonuclease SbcCD subunit D C-terminal domain-containing protein - AAA family ATPase. nitroreductase family protein - acyl-CoA dehydrogenase family protein - hypothetical protein - protein kinase - hypothetical protein - DUF559 domain-containing protein - FHA domain-containing protein : DUF2993 domain-containing protein - SDR family oxidoreductase - FAD-binding oxidoreductase - TetR/AcrR family transcriptional regulator - glycerol-3-phosphate dehydrogenase/oxidase - sensor histidine kinase - acyl-CoA dehydrogenase family protein. NTP transferase domain-containing protein - GNAT family N-acetyltransferase - substrate-binding domain-containing protein - DUF6158 family protein - sensor histidine kinase - response regulator - hypothetical protein : Hsp70 family protein - dienelactone hydrolase family protein - hypothetical protein - hypothetical protein - response regulator transcription factor - cell wall metabolism sensor histidine kinase WalK - glycosyltransferase family 39 protein. nuclear transport factor 2 family protein - helix-turn-helix domain-containing protein - hypothetical protein - DUF3817 domain-containing protein - hypothetical protein - hypothetical protein - SigE family RNA polymerase sigma factor : hypothetical protein - nitroreductase family protein - NAD(P)H-quinone oxidoreductase - bacterial proteasome activator family protein - metallopeptidase TldDrelated protein - TldD/PmbA family protein - hypothetical protein. ParA family protein - ParA family protein - helicase HerA domain-containing protein - hypothetical protein - TIR domain-containing protein - serine/threonine-protein kinase - flavoprotein : hypothetical protein - N,N-dimethylformamidase beta subunit family domain-containing protein - MFS transporter - ResA-like WAxxUGC motif-containing protein - hypothetical protein - phosphotransferase family protein - ResA-like WAxxUGC motif-containing protein. PAS domain S-box protein - bifunctional diguanylate cyclase/phosphodiesterase - hypothetical protein - class I adenylate-forming enzyme family protein - acyl-CoA thioesterase II - Na+/H+ antiporter NhaA - hypothetical protein : hypothetical protein - hypothetical protein - tRNA-Ala - histidine phosphatase family protein - ribosome silencing factor - nicotinate-nucleotide adenylyltransferase - phosphoglucomutase (alpha-D-glucose-1,6-bisphosphate-dependent). PASTA domain-containing protein - hypothetical protein - TadE/TadG family type IV pilus assembly protein - hypothetical protein - type II secretion system F family protein - type II secretion system F family protein - CpaF family protein : PAS domain-containing protein - hypothetical protein - hypothetical protein - patatin-like phospholipase family protein - transcriptional regulator - chromosome partitioning protein - SAF domain-containing protein. PfkB family carbohydrate kinase - serine hydrolase - Si-specific NAD(P)(+) transhydrogenase - class II fumarate hydratase - 4-hydroxy-3-methylbut-2-enyl diphosphate reductase - exodeoxyribonuclease VII large subunit - exodeoxyribonuclease VII small subunit : DUF4245 domain-containing protein - class II fructose-bisphosphatase - sterol desaturase family protein - polysaccharide deacetylase family protein - DUF6542 domain-containing protein - DNA recombination protein RmuC - redox-regulated ATPase YchF. phage holin family protein - alpha/beta fold hydrolase - MarP family serine protease - plastocyanin/azurin family copper-binding protein - hypothetical protein - endonuclease III - hypothetical protein : Crp/Fnr family transcriptional regulator - adenylate/guanylate cyclase domain-containing protein - MBL fold metallo-hydrolase - NUDIX hydrolase - RidA family protein - DUF4177 domain-containing protein - ArsA-related P-loop ATPase. PHB depolymerase family esterase - adenylate kinase - class I SAM-dependent methyltransferase - bifunctional methylenetetrahydrofolate dehydrogenase/methenyltetrahydrofolate cyclohydrolase FolD - nucleotidyltransferase domain-containing protein - class I SAM-dependent methyltransferase - GNAT family N-acetyltransferase : hypothetical protein - TauD/TfdA family dioxygenase - NADH:flavin oxidoreductase - YciI family protein - hypothetical protein - hypothetical protein - PP2C family serine/threonine-protein phosphatase. PHB depolymerase family esterase - C40 family peptidase - hypothetical protein - helix-turn-helix domain-containing protein - IS630 family transposase - phosphoenolpyruvate carboxylase - ROK family protein : TauD/TfdA family dioxygenase - PhnD/SsuA/transferrin family substrate-binding protein - ABC transporter permease - ABC transporter ATP-binding protein - hypothetical protein - BlaI/MecI/CopY family transcriptional regulator - MarR family winged helix-turn-helix transcriptional regulator. PH domain-containing protein - glycoside hydrolase family 15 protein - hypothetical protein - RidA family protein - cell division protein CrgA - DUF881 domain-containing protein - DUF222 domain-containing protein : aminodeoxychorismate/anthranilate synthase component II - Stk1 family PASTA domain-containing Ser/Thr kinase - serine/threonine-protein kinase - penicillin-binding protein 2 - FtsW/RodA/SpoVE family cell cycle protein - Stp1/IreP family PP2C-type Ser/Thr phosphatase - FHA domain-containing protein. P-loop ATPase, Sll1717 family - FadR/GntR family transcriptional regulator - carboxymuconolactone decarboxylase family protein - peroxiredoxin - LysR substrate-binding domain-containing protein -

glutamate--cysteine ligase - N-acetylglutaminylglutamine amidotransferase : alpha/beta hydrolase family protein - GNAT family N-acetyltransferase - GNAT family N-acetyltransferase - histidine kinase - response regulator transcription factor - DUF4097 family beta strand repeat-containing protein - transposase. polyketide cyclase - hypothetical protein - MOSC domain-containing protein - enoyl-CoA hydratase - acyl-CoA synthetase - bifunctional diguanylate cyclase/phosphodiesterase - SAM-dependent methyltransferase : hypothetical protein - NAD(P)-dependent alcohol dehydrogenase - DUF1918 domain-containing protein - histidine kinase - TetR/AcrR family transcriptional regulator - glycosyltransferase 87 family protein - molybdopterin-dependent oxidoreductase. polysaccharide deacetylase family protein - DUF6542 domain-containing protein - DNA recombination protein RmuC - redox-regulated ATPase YchF - DUF952 domain-containing protein - ribosylglycohydrolase - 4a-hydroxytetrahydrobiopterin dehydratase : (deoxy)nucleoside triphosphate pyrophosphohydrolase - translational GTPase TypA - N-acetyl-1-D-myo-inositol-2-amino-2-deoxy-alpha-D-glucopyranoside deacetylase - S1C family serine protease - hypothetical protein - DUF6113 family protein - flavin reductase family protein. PP2C family serine/threonine-protein phosphatase - LLM class F420-dependent oxidoreductase - hypothetical protein - NADP-dependent isocitrate dehydrogenase - DUF6328 family protein - serine/threonine-protein kinase - flavin reductase family protein : DUF4397 domain-containing protein - hypothetical protein - class F sortase - class I SAM-dependent methyltransferase - alpha/beta hydrolase family protein - SGNH/GDSL hydrolase family protein - TauD/TfdA family dioxygenase. prepilin peptidase - chorismate synthase - shikimate kinase - 3-dehydroquinate synthase - elongation factor P - transcription antitermination factor NusB - transcriptional regulator : bifunctional pyr operon transcriptional regulator/uracil phosphoribosyltransferase PyrR - aspartate carbamoyltransferase catalytic subunit - dihydroorotase - hypothetical protein - hypothetical protein - glutamine-hydrolyzing carbamoyl-phosphate synthase small subunit - carbamoyl-phosphate synthase large subunit. proteasome ATPase - depupylase/deamidase Dop - ubiquitin-like protein Pup - proteasome subunit beta - proteasome subunit alpha - dienelactone hydrolase family protein - hypothetical protein : Pup--protein ligase - FKBP-type peptidyl-prolyl cis-trans isomerase - YafY family protein - YafY family protein - hypothetical protein - Sec-independent protein translocase subunit TatA - twin-arginine translocase subunit TatC. pseudouridine-5'-phosphate glycosidase - TIGR03667 family PPOX class F420-dependent oxidoreductase - ATP-dependent chaperone ClpB - heat shock protein transcriptional repressor HspR - molecular chaperone DnaJ - nucleotide exchange factor GrpE - molecular chaperone DnaK : nuclease-related domain-containing protein - (Fe-S)-binding protein - dCTP deaminase - beta-phosphoglucomutase family hydrolase - glycoside hydrolase family 65 protein - tRNA-Gly - pentapeptide repeat-containing protein. pyridoxamine 5'-phosphate oxidase family protein - endo alpha-1,4 polygalactosaminidase - FMN-dependent NADH-azoreductase - ABC transporter substrate-binding protein - metallophosphoesterase family protein - nitroreductase family protein - hypothetical protein : amidohydrolase family protein - nitroreductase family protein - aldose 1-epimerase family protein - ABC transporter substrate-binding protein - LLM class flavin-dependent oxidoreductase - iron ABC transporter permease - ABC transporter ATP-binding protein. response regulator - ATP-binding protein - potassium-transporting ATPase subunit KdpA - potassium-transporting ATPase subunit C - potassium-transporting ATPase subunit KdpB - potassium-transporting ATPase subunit KdpA - K(+)-transporting ATPase subunit F : hypothetical protein - type I polyketide synthase - type I polyketide synthase - type I polyketide synthase - beta-ketoacyl synthase N-terminal-like domain-containing protein - methylaspartate mutase - cobalamin B12-binding domain-containing protein. response regulator - Fur family transcriptional regulator - rubrerythrin family protein - DUF3501 family protein - toll/interleukin-1 receptor domain-containing protein - SseB family protein - hypothetical protein : heme-copper oxidase subunit III - c-type cytochrome - ubiquinol-cytochrome c reductase iron-sulfur subunit - ubiquinol-cytochrome c reductase cytochrome b subunit - DEDD exonuclease domain-containing protein - VOC family protein - HIT family protein. response regulator transcription factor - DUF5685 family protein - aminotransferase class V-fold PLP-dependent enzyme - L-2-hydroxyglutarate oxidase - glycoside hydrolase family 15 protein - hypothetical protein - PGPGW domain-containing protein : cation:proton antiporter regulatory subunit - cation:proton antiporter - enoyl-ACP reductase FabI - pyruvate dehydrogenase - DUF167 family protein - 3-oxoacyl-ACP reductase FabG - hypothetical protein. response regulator transcription factor - sensor histidine kinase KdpD - NAD-dependent epimerase/dehydratase family protein - sulfite exporter TauE/SafE family protein - pentapeptide repeat-containing protein - NAD(P)/FAD-dependent oxidoreductase - MarR family winged helix-turn-helix transcriptional regulator : protein-tyrosine phosphatase family protein - PPOX class F420-dependent oxidoreductase - aldo/keto reductase - helix-turn-helix domain-containing protein - hypothetical protein - class I SAM-dependent methyltransferase - phosphotransferase family protein. rhodanese-like domain-containing protein - FAD-binding oxidoreductase - MFS transporter - thioesterase II family protein - LLM class flavin-dependent oxidoreductase - carboxyl transferase domain-containing protein - hypothetical protein : acyl-CoA carboxylase subunit beta - MMPL family transporter - LysR family transcriptional regulator - SDR family oxidoreductase - IS630 family transposase - helix-turn-helix domain-containing protein - hypothetical protein. RidA family protein - DUF4177 domain-containing protein - ArsA-related P-loop ATPase - ArsA-related P-loop ATPase - WhiB family transcriptional regulator - IS630 family transposase - helix-turn-helix domain-containing protein : transglycosylase domain-containing protein - GatB/YqeY domain-containing protein - metallophosphoesterase - tRNA-Pro - glyoxalase/bleomycin resistance/dioxygenase family protein - UMP kinase - hypothetical protein. RNA polymerase-binding protein RbpA - serine/threonine-protein kinase - SHOCT domain-containing protein - 6-phosphogluconolactonase - glucose-6-phosphate dehydrogenase assembly protein OpcA - glucose-6-phosphate dehydrogenase - transaldolase : heme o synthase - tetratricopeptide repeat protein - FxSxx-COOH system tetratricopeptide repeat protein - toll/interleukin-1 receptor domain-containing protein - PPOX class F420-dependent oxidoreductase - heme A synthase - ABC transporter permease. rod shape-determining protein - nucleoside-diphosphate kinase - dihydrofolate synthase - folylpolyglutamate synthase/dihydrofolate synthase family protein - ATP-dependent Clp protease ATP-binding subunit ClpX - ATP-dependent Clp protease proteolytic subunit - ATP-dependent Clp protease proteolytic subunit : trigger factor - tRNA-Pro - tRNA-Gly - class I SAM-dependent methyltransferase - peptidylprolyl isomerase - L,D-transpeptidase - DUF3291 domain-containing protein. SAM-dependent methyltransferase - hypothetical protein - AAA family ATPase - hypothetical protein - hypothetical protein - hypothetical protein - ATP-binding protein : hypothetical protein - FxSxx-COOH system tetratricopeptide repeat protein - winged helix-turn-helix domain-containing protein - transposase - ATP-binding protein - hypothetical protein - effector-associated domain EAD1-containing protein. SCO6880 family protein - conjugal transfer protein TraC - hypothetical protein - YbhB/YbcL family Raf kinase inhibitor-like protein - P-II family nitrogen regulator - DedA family protein - lipase family protein : IS1634 family transposase - TetR/AcrR family transcriptional regulator -

organic hydroperoxide resistance protein - glycosyltransferase family 4 protein - class I SAM-dependent methyltransferase - hypothetical protein - class I SAM-dependent methyltransferase. ScpA family protein - Zn-dependent membrane protease - ParA family protein - S-(hydroxymethyl)mycothiol dehydrogenase - MBL fold metallo-hydrolase - GlsB/YeaQ/YmgE family stress response membrane protein - serine/threonine-protein kinase : lipopolysaccharide assembly protein LapA domain-containing protein - hypothetical protein - protein kinase - hypothetical protein - hypothetical protein - AIPR family protein - IS607 family element RNA-guided endonuclease TnpB. SDR family NAD(P)-dependent oxidoreductase - TauD/TfdA family dioxygenase - SDR family NAD(P)-dependent oxidoreductase - class II aldolase/adducin family protein - hypothetical protein - phosphotransferase - hypothetical protein : SAM-dependent methyltransferase - ABC transporter substrate-binding protein - phosphotransferase family protein - hypothetical protein - MoxR family ATPase - DUF58 domain-containing protein - transglutaminase domain-containing protein. SDR family oxidoreductase - LLM class F420-dependent oxidoreductase - pyridoxamine 5'-phosphate oxidase family protein - hypothetical protein - hypothetical protein - amidohydrolase family protein - thioesterase family protein : DUF2752 domain-containing protein - TM2 domain-containing protein - hypothetical protein - AAA family ATPase - hypothetical protein - response regulator transcription factor - RNA helicase. SDR family oxidoreductase - tRNA-Tyr - YajQ family cyclic di-GMP-binding protein - zinc metalloprotease HtpX - Rossmann-like and DUF2520 domain-containing protein - 2-oxoacid:acceptor oxidoreductase subunit alpha - 2-oxoacid:ferredoxin oxidoreductase subunit beta : hypothetical protein - hypothetical protein - polyprenyl synthetase family protein - NADH-quinone oxidoreductase subunit NuoN - NADH-quinone oxidoreductase subunit M - NADH-quinone oxidoreductase subunit L - NADH-quinone oxidoreductase subunit NuoK. sensor histidine kinase - ABC transporter permease subunit - ABC transporter ATP-binding protein - TetR/AcrR family transcriptional regulator - alpha/beta hydrolase fold domain-containing protein - NtaA/DmoA family FMN-dependent monooxygenase - ABATE domain-containing protein : ABC transporter substrate-binding protein - ABC transporter permease - ABC transporter permease - ABC transporter ATP-binding protein - NtaA/DmoA family FMN-dependent monooxygenase - FAD-dependent monooxygenase - LLM class flavin-dependent oxidoreductase. sensor histidine kinase - tRNA-Leu - ANTAR domain-containing response regulator - branched-chain amino acid ABC transporter substrate-binding protein - branched-chain amino acid ABC transporter substrate-binding protein - PaaI family thioesterase - DNA polymerase I : putative bifunctional diguanylate cyclase/phosphodiesterase - hypothetical protein - M4 family metallopeptidase - hypothetical protein - hypothetical protein - dephospho-CoA kinase - excinuclease ABC subunit UvrB. serine O-acetyltransferase - cysteine synthase A - (Fe-S)-binding protein - lactate utilization protein B - LUD domain-containing protein - hypothetical protein - NAD(P)-dependent oxidoreductase : LLM class flavin-dependent oxidoreductase - LLM class flavin-dependent oxidoreductase - ABC transporter substrate-binding protein - ABC transporter permease - ABC transporter ATP-binding protein - IS1182 family transposase - TetR/AcrR family transcriptional regulator. serine/threonine-protein kinase - Rv3654c family TadE-like protein - TadE family type IV pilus minor pilin - DUF4244 domain-containing protein - type II secretion system F family protein - TadA family conjugal transfer-associated ATPase - septum site-determining protein Ssd : hypothetical protein - phage holin family protein - alpha/beta fold hydrolase - MarP family serine protease - plastocyanin/azurin family copper-binding protein - hypothetical protein - endonuclease III. succinate dehydrogenase flavoprotein subunit - succinate dehydrogenase iron-sulfur subunit - hypothetical protein - RecQ family ATP-dependent DNA helicase - SCO4848 family membrane protein - type II toxin-antitoxin system HicA family toxin - type II toxin-antitoxin system HicB family antitoxin : hypothetical protein - toll/interleukin-1 receptor domain-containing protein - hypothetical protein - GNAT family N-acetyltransferase - putative zinc-binding metallopeptidase - DUF1003 domain-containing protein - D-alanyl-D-alanine carboxypeptidase family protein. SulP family inorganic anion transporter - HAD-IC family P-type ATPase - CBS domain-containing protein - cation-transporting P-type ATPase - hypothetical protein - hypothetical protein - PP2C family protein-serine/threonine phosphatase : hypothetical protein - hypothetical protein - Hsp20/alpha crystallin family protein - hypothetical protein - phosphoenolpyruvate synthase - heavy metal translocating P-type ATPase - GAF domain-containing protein. TetR/AcrR family transcriptional regulator - ABC transporter substrate-binding protein - MaoC family dehydratase N-terminal domain-containing protein - branched-chain amino acid ABC transporter permease/ATP-binding protein - ABC transporter ATP-binding protein - ABC transporter substrate-binding protein - LuxR C-terminal-related transcriptional regulator : hypothetical protein - enoyl-CoA hydratase/isomerase family protein - FadR/GntR family transcriptional regulator - SpoIIE family protein phosphatase - pyridoxamine 5'-phosphate oxidase family protein - M20/M25/M40 family metallo-hydrolase - GNAT family N-acetyltransferase. TetR/AcrR family transcriptional regulator - CbtB-domain containing protein - CbtA family protein - MmgE/PrpD family protein - methylisocitrate lyase - VIT family protein - hypothetical protein : hypothetical protein - TetR/AcrR family transcriptional regulator - aldo/keto reductase - TIR domain-containing protein - globin domain-containing protein - SDR family NAD(P)-dependent oxidoreductase - carboxymuconolactone decarboxylase family protein. TetR/AcrR family transcriptional regulator - FAD-dependent oxidoreductase - EthD family reductase - DUF2252 domain-containing protein - class F sortase - hypothetical protein - DJ-1/PfpI family protein : SMI1/KNR4 family protein - PPOX class F420-dependent oxidoreductase - sulfurtransferase - P1 family peptidase - hypothetical protein - hypothetical protein - hypothetical protein. TetR/AcrR family transcriptional regulator - hypothetical protein - glycosyltransferase family 39 protein - dolichyl-phosphate beta-glucosyltransferase - IS256 family transposase - tetratricopeptide repeat protein - IS66 family transposase : threonine/serine dehydratase - molecular chaperone HtpG - hypothetical protein - IS1634 family transposase - methyltransferase domain-containing protein - hypothetical protein - excalibur calcium-binding domain-containing protein. TetR/AcrR family transcriptional regulator - hypothetical protein - hypothetical protein - hypothetical protein - hypothetical protein - PH domain-containing protein - biotin--[acetyl-CoA-carboxylase] ligase : acyl-CoA carboxylase subunit beta - acyl-CoA carboxylase subunit epsilon - biotin carboxylase N-terminal domain-containing protein - DUF5313 family protein - hypothetical protein - hypothetical protein - NAD(P)H-quinone dehydrogenase. TetR/AcrR family transcriptional regulator - LLM class flavin-dependent oxidoreductase - TetR/AcrR family transcriptional regulator - ABC-F family ATP-binding cassette domain-containing protein - hypothetical protein - recombinase family protein - hypothetical protein : YdcF family protein - GntR family transcriptional regulator - hypothetical protein - hypothetical protein - hypothetical protein - DUF222 domain-containing protein - acetyl-CoA hydrolase/transferase C-terminal domain-containing protein. thiaminase II - class I SAM-dependent methyltransferase - glycosyltransferase - alpha/beta fold hydrolase - phosphopantetheine-binding protein - beta-ketoacyl synthase N-terminal-like domain-containing protein - SDR family NAD(P)-dependent oxidoreductase : hypothetical protein - beta-ketoacyl synthase N-terminal-like domain-containing protein - LuxR C-terminal-related transcriptional regulator - hypothetical protein - LuxR family transcriptional regulator - DUF4193 family protein - co-chaperone GroES. threonine/serine dehydratase - molecular chaperone HtpG - hypothetical protein - IS1634 family transposase - methyltransferase domain-containing protein - hypothetical protein - excalibur calcium-binding domain-containing protein : ABC transporter substrate-binding protein - metallophosphoesterase - TetR family transcriptional regulator - acyl-CoA dehydrogenase - MBL fold metallo-hydrolase - SDR family NAD(P)-dependent oxidoreductase - thiolase family protein. TIGR03667 family PPOX class F420-dependent oxidoreductase - ATP-dependent chaperone ClpB - heat shock protein transcriptional repressor HspR - molecular chaperone DnaJ - nucleotide exchange factor GrpE - molecular chaperone DnaK - nuclease-related domain-containing protein : (Fe-S)-binding protein - dCTP deaminase - beta-phosphoglucomutase family hydrolase - glycoside hydrolase family 65 protein - tRNA-Gly - pentapeptide repeat-containing protein - carbohydrate kinase family protein. TIR domain-containing protein - TetR/AcrR family transcriptional regulator - bifunctional 5,10-methylenetetrahydrofolate dehydrogenase/5,10-methenyltetrahydrofolate cyclohydrolase - diguanylate cyclase - adenosylmethionine--8-amino-7-oxononanoate transaminase - rhomboid family intramembrane serine protease - integrase core domain-containing protein : IS4 family transposase - IS3 family transposase - IS3 family transposase - transposase - DUF222 domain-containing protein - MFS transporter - hypothetical protein. transcriptional repressor LexA - LysM peptidoglycan-binding domain-containing protein - transcriptional regulator NrdR - vitamin B12-dependent ribonucleotide reductase - hypothetical protein - type II toxin-antitoxin system VapC family toxin - trehalase family glycosidase : RNA polymerase sigma factor - acetoacetate--CoA ligase - acetoin utilization protein AcuC - bifunctional GNAT family N-acetyltransferase/acetate--CoA ligase family protein - PAS domain-containing protein - SpoIIE family protein phosphatase - ABC transporter ATP-binding protein. transglutaminase family protein - bifunctional diguanylate cyclase/phosphodiesterase - 2-phospho-L-lactate guanylyltransferase - NAD(P)H-dependent glycerol-3-phosphate dehydrogenase - UDP-N-acetylglucosamine 1-carboxyvinyltransferase - bifunctional hydroxymethylpyrimidine kinase/phosphomethylpyrimidine kinase - Lrp/AsnC family transcriptional regulator : DUF3515 domain-containing protein - D-alanine--D-alanine ligase family protein - HNH endonuclease signature motif containing protein - 50S ribosomal protein L28 - ATP-dependent DNA helicase RecG - 16S rRNA (guanine(966)-N(2))-methyltransferase RsmD - pantetheine-phosphate adenylyltransferase. transglutaminase-like domain-containing protein - hypothetical protein - PIG-L deacetylase family protein - aminomethyl-transferring glycine dehydrogenase - FkbM family methyltransferase - GGDEF domain-containing protein - HD domain-containing phosphohydrolase : hypothetical protein - TetR/AcrR family transcriptional regulator - MMPL family transporter - glutamine synthetase family protein - IucA/IucC family siderophore biosynthesis protein - lysine N(6)-hydroxylase/L-ornithine N(5)-oxygenase family protein - IucA/IucC family siderophore biosynthesis protein. tRNA-Glu - hypothetical protein - sensor histidine kinase KdpD - serine/threonine-protein kinase - IclR family transcriptional regulator - 3-isopropylmalate dehydratase large subunit - 3-isopropylmalate dehydratase small subunit : HU family DNA-binding protein - RNA degradosome polyphosphate kinase - transglutaminase family protein - bifunctional diguanylate cyclase/phosphodiesterase - 2-phospho-L-lactate guanylyltransferase - NAD(P)H-dependent glycerol-3-phosphate dehydrogenase - UDP-N-acetylglucosamine 1-carboxyvinyltransferase. tRNA (N6-isopentenyl adenosine(37)-C2)-

methylthiotransferase MiaB - alpha/beta fold hydrolase - tRNA (adenosine(37)-N6)-dimethylallyltransferase MiaA - EAL domain-containing protein - diaminopimelate epimerase - VOC family protein - helix-turn-helix transcriptional regulator : GTPase HflX - transcriptional repressor LexA - LysM peptidoglycan-binding domain-containing protein - transcriptional regulator NrdR - vitamin B12-dependent ribonucleotide reductase - hypothetical protein - type II toxin-antitoxin system VapC family toxin. tRNA-Thr - response regulator transcription factor - stage 0 sporulation family protein - DNA polymerase III subunit delta' - dTMP kinase - type I DNA topoisomerase - sodium-translocating pyrophosphatase : ATP-binding protein - STAS domain-containing protein - DEAD/DEAH box helicase - hypothetical protein - PASTA domain-containing protein - hypothetical protein - serine/threonine-protein kinase. type 1 glutamine amidotransferase - PHB depolymerase family esterase - C40 family peptidase - hypothetical protein - helix-turn-helix domain-containing protein - IS630 family transposase - phosphoenolpyruvate carboxylase : ROK family protein - TauD/TfdA family dioxygenase - PhnD/SsuA/transferrin family substrate-binding protein - ABC transporter permease - ABC transporter ATP-binding protein - hypothetical protein - BlaI/MecI/CopY family transcriptional regulator. type I-E CRISPR-associated endonuclease Cas1e - type I-E CRISPR-associated protein Cas6/Cse3/CasE - type I-E CRISPR-associated protein Cas5/CasD - type I-E CRISPR-associated protein Cas7/Cse4/CasC - type I-E CRISPR-associated protein Cse2/CasB - type I-E CRISPR-associated protein Cse1/CasA - CRISPR-associated helicase Cas3' : hypothetical protein - recombinase family protein - transposase - winged helix-turn-helix domain-containing protein - hypothetical protein - type II toxin-antitoxin system Phd/YefM family antitoxin - PIN domain-containing protein. type II secretion system F family protein - CpaF family protein - PAS domain-containing protein - hypothetical protein - hypothetical protein - patatin-like phospholipase family protein - transcriptional regulator : chromosome partitioning protein - SAF domain-containing protein - helix-turn-helix domain-containing protein - hypothetical protein - LysM peptidoglycan-binding domain-containing protein - Rv3235 family protein - preprotein translocase subunit SecA. type II toxin-antitoxin system PemK/MazF family toxin - cyclopropane-fatty-acyl-phospholipid synthase family protein - DNA polymerase III subunit alpha - aspartate aminotransferase family protein - alpha/beta fold hydrolase - hypothetical protein - DEAD/DEAH box helicase : DUF445 domain-containing protein - hypothetical protein - DUF6343 family protein - enoyl-CoA hydratase-related protein - acyl-CoA thioesterase II - ATP-binding protein - roadblock/LC7 domain-containing protein. type II toxin-antitoxin system Phd/YefM family antitoxin - Txe/YoeB family addiction module toxin - DUF6326 family protein - response regulator transcription factor - sensor histidine kinase - tRNA-Leu - ANTAR domain-containing response regulator : branched-chain amino acid ABC transporter substrate-binding protein - branched-chain amino acid ABC transporter substrate-binding protein - PaaI family thioesterase - DNA polymerase I - putative bifunctional diguanylate cyclase/phosphodiesterase - hypothetical protein - M4 family metallopeptidase. type I phosphomannose isomerase catalytic subunit - DUF5719 family protein - hypothetical protein - glycosyltransferase - WhiB family transcriptional regulator - bifunctional FO biosynthesis protein CofGH - hypothetical protein : TetR/AcrR family transcriptional regulator - 2-phospho-L-lactate transferase - coenzyme F420-0:L-glutamate ligase - DUF3105 domain-containing protein - DNA polymerase III subunit beta - sensor histidine kinase - sugar phosphate nucleotidyltransferase. type I polyketide synthase - SDR family oxidoreductase - nuclear transport factor 2 family protein - nuclear transport factor 2 family protein - TetR/AcrR family transcriptional regulator - GNAT family N-acetyltransferase - hypothetical protein : hypothetical protein -

alpha/beta fold hydrolase - cysteine hydrolase family protein - HNH endonuclease signature motif containing protein - MFS transporter - alpha/beta fold hydrolase - DUF2889 domain-containing protein. type I polyketide synthase - type I polyketide synthase - methylmalonyl-CoA epimerase - fatty acyl-AMP ligase - hypothetical protein - hypothetical protein - beta-propeller fold lactonase family protein : hypothetical protein - hypothetical protein - hypothetical protein - tRNA-Phe - tRNA-Asp - FadR/GntR family transcriptional regulator - MFS transporter. UbiA family prenyltransferase - SpoIIE family protein phosphatase - HAMP domain-containing protein - acyl-CoA desaturase - ferredoxin reductase - HdeD family acid-resistance protein - SDR family oxidoreductase : molybdopterin-binding protein - molecular chaperone DnaK - nucleotide exchange factor GrpE - DnaJ C-terminal domain-containing protein - chaperone modulator CbpM - UBP-type zinc finger domain-containing protein - SRPBCC family protein. undecaprenyldiphospho-muramoylpentapeptide beta-N-acetylglucosaminyltransferase - putative lipid II flippase FtsW - UDP-N-acetylmuramoyl-L-alanine--D-glutamate ligase - phospho-N-acetylmuramoyl-pentapeptide-transferase - UDP-N-acetylmuramoyl-tripeptide--D-alanyl-D-alanine ligase - UDP-N-acetylmuramoyl-L-alanyl-D-glutamate--2,6-diaminopimelate ligase - penicillin-binding protein 2 : hypothetical protein - 16S rRNA (cytosine(1402)-N(4))-methyltransferase RsmH - division/cell wall cluster transcriptional repressor MraZ - MoxR family ATPase - DUF58 domain-containing protein - DUF3488 and transglutaminase-like domain-containing protein - hypothetical protein. universal stress protein - chaperonin GroEL - DDE-type integrase/transposase/recombinase - hypothetical protein - transposase - GyrI-like domain-containing protein - cell division protein FtsK : effector-associated domain EAD1-containing protein - effector-associated domain EAD1-containing protein - hypothetical protein - hypothetical protein - hypothetical protein - IS5 family transposase - tRNA-Arg. UvrD-helicase domain-containing protein - hypothetical protein - hypothetical protein - IS6 family transposase - hypothetical protein - recombinase family protein - IS6 family transposase : type I-E CRISPR-associated endoribonuclease Cas2e - type I-E CRISPR-associated endonuclease Cas1e - type I-E CRISPR-associated protein Cas6/Cse3/CasE - type I-E CRISPR-associated protein Cas5/CasD - type I-E CRISPR-associated protein Cas7/Cse4/CasC - type I-E CRISPR-associated protein Cse2/CasB - type I-E CRISPR-associated protein Cse1/CasA. VOC family protein - TetR/AcrR family transcriptional regulator - class I SAM-dependent methyltransferase - hypothetical protein - DsrE family protein - TetR/AcrR family transcriptional regulator - class I SAM-dependent methyltransferase : MFS transporter - universal stress protein - chaperonin GroEL - DDE-type integrase/transposase/recombinase - hypothetical protein - transposase - GyrI-like domain-containing protein. WD40 repeat domain-containing protein - hypothetical protein - helix-turn-helix transcriptional regulator - MFS transporter - hypothetical protein - cytochrome P450 - hypothetical protein : cytochrome P450 - FAD-dependent monooxygenase - AfsR/SARP family transcriptional regulator - TetR/AcrR family transcriptional regulator - nuclear transport factor 2 family protein - PhnD/SsuA/transferrin family substrate-binding protein - ABC transporter ATP-binding protein. WD40 repeat domain-containing protein - hypothetical protein - site-specific integrase - hypothetical protein - LLM class flavin-dependent oxidoreductase - HD domain-containing protein - DsbA family oxidoreductase : hypothetical protein - toll/interleukin-1 receptor domain-containing protein - Sec-independent protein translocase subunit TatA - GlsB/YeaQ/YmgE family stress response membrane protein - TIGR03620 family F420-dependent LLM class oxidoreductase - CrcB family protein - CrcB family protein. WD40 repeat domain-containing serine/threonine protein kinase - universal stress protein - YihY/virulence factor BrkB family protein - lipopolysaccharide assembly LapA domain-containing protein - hypothetical protein - mechanosensitive ion channel family protein - DUF3710 domain-containing protein : SsgA family sporulation/cell division regulator - GNAT family N-acetyltransferase - hypothetical protein - DUF664 domain-containing protein - aldo/keto reductase - AMP-binding protein - IS200/IS605 family transposase. WhiB family transcriptional regulator - histidine phosphatase family protein - hypothetical protein - trehalose-6-phosphate synthase - hypothetical protein - hypothetical protein - FtsK/SpoIIIE domain-containing protein : hypothetical protein - MerR family transcriptional regulator - bifunctional nuclease family protein - MerR family transcriptional regulator - FHA domain-containing protein - glycine cleavage system protein GcvH - DUF881 domain-containing protein. YafY family protein - FxSxx-COOH system tetratricopeptide repeat protein - FHA domain-containing protein - VWA domain-containing protein - protein phosphatase 2C domain-containing protein - tetratricopeptide repeat protein - hypothetical protein : glutamate ABC transporter substrate-binding protein - hypothetical protein - hypothetical protein - molybdate ABC transporter substrate-binding protein - ABC transporter permease - ABC transporter ATP-binding protein - STAS domain-containing protein. YajQ family cyclic di-GMP-binding protein - zinc metalloprotease HtpX - Rossmann-like and DUF2520 domain-containing protein - 2-oxoacid:acceptor oxidoreductase subunit alpha - 2-oxoacid:ferredoxin oxidoreductase subunit beta - hypothetical protein - hypothetical protein : polyprenyl synthetase family protein - NADH-quinone oxidoreductase subunit NuoN - NADH-quinone oxidoreductase subunit M - NADH-quinone oxidoreductase subunit L - NADH-quinone oxidoreductase subunit NuoK - NADH-quinone oxidoreductase subunit J - NADH-quinone oxidoreductase subunit NuoI. YceI family protein - ferredoxin - HdeD family acid-resistance protein - tyrosine-protein phosphatase - SOS response-associated peptidase - M50 family metallopeptidase - 1-acyl-sn-glycerol-3-phosphate acyltransferase : hypothetical protein - hypothetical protein - hypothetical protein - alkane 1-monooxygenase - hypothetical protein - AEC family transporter - 3-hydroxybutyryl-CoA dehydrogenase. Zn-ribbon domain-containing OB-fold protein - hypothetical protein - hypothetical protein - CHAT domain-containing protein - hypothetical protein - NADP-dependent oxidoreductase - hypothetical protein : ABC transporter ATP-binding protein - ABC transporter permease - YhjD/YihY/BrkB family envelope integrity protein - hypothetical protein - SRPBCC domain-containing protein - hypothetical protein - ATP-binding protein.

ChatGPT 5.5 answer:

I’ll number the contexts by their order in your pasted list, starting with **1** for the first line. I’m screening for the same themes as before: Tad/type IV pilus or conjugation, septation/cell division, envelope/cell-wall stress, Fic/toxin-stress regulation, and closely adjacent HAD/phosphatase-type accessory genes.

I counted **256 contexts**.

I numbered them by order of appearance, starting with **1** for the first context. Using that numbering, the contexts biologically/functionally related to the previous segment are:

**Strongest / direct matches:**
**217**

This is the closest match because it contains:

> serine/threonine-protein kinase - Rv3654c family TadE-like protein - TadE family type IV pilus minor pilin - DUF4244 - type II secretion system F family protein - **TadA family conjugal transfer-associated ATPase - septum site-determining protein Ssd**

This is essentially the same biological neighborhood: **Tad/type IV pilus machinery + Ssd/septation/cell-cycle control**.

### Other clearly related contexts

These contain **Tad/type IV pilus, prepilin, type II secretion F/CpaF, TraC/conjugal transfer, or pilus-assembly genes**:

**64, 98, 101, 186, 196, 209, 217, 238**

Why:

| Context | Relevant signal |
|---|---|
| 64 | **conjugal transfer protein TraC** |
| 98 | **conjugal transfer protein TraC** |
| 101 | **pilus assembly protein TadG-related protein** |
| 186 | **TadE/TadG family type IV pilus assembly**, type II secretion F, CpaF |
| 196 | **prepilin peptidase**, often linked to type IV pilus/prepilin processing |
| 209 | **conjugal transfer protein TraC** |
| 217 | **TadE, TadA, Ssd** — direct match |
| 238 | type II secretion F, CpaF, adjacent cell-envelope/pilus-associated genes |

### Contexts related through septation, cell division, cell-wall growth, or envelope remodeling

These are related to the **Ssd / septation / cell-cycle / envelope-stress** side of the interpretation:

**9, 49, 51, 73, 75, 77, 84, 103, 106, 118, 125, 128, 129, 139, 147, 148, 153, 166, 167, 173, 182, 191, 207, 210, 245, 246, 250, 251, 252, 256**

Important examples:

| Context | Relevant signal |
|---|---|
| 49 | **septation protein SepH**, LCP family protein, BrkB-like virulence/envelope factor |
| 51 | **cell division protein CrgA** |
| 77 | **DivIVA domain-containing protein**, HAD hydrolase, AbiE antitoxin |
| 84 | **FtsK/SpoIIIE domain-containing protein** |
| 103 | **DivIVA domain-containing protein** |
| 106 | **FtsX-like permease**, ABC transporter ATP-binding protein |
| 118 | **septum formation family protein** |
| 125 | **FtsX / FtsE cell division system** |

| Context | Relevant signal |
|---|---|
| 128 | **FtsK/SpoIIIE**, BrkB/YhjD/YihY envelope-integrity protein |
| 166 | **SsgA family sporulation/cell division regulator**, WhiA, Ser/Thr kinase |
| 182 | **WalK cell wall metabolism sensor histidine kinase** |
| 191 | **CrgA, PASTA-domain kinase, PBP2, FtsW/RodA/SpoVE, PP2C phosphatase** |
| 245 | large **cell wall/division cluster**, including FtsW and PBP2 |
| 251 | **SsgA**, LapA, BrkB/GlsB-type stress/envelope proteins |
| 252 | **FtsK/SpoIIIE** |

These are not necessarily Tad/Ssd loci, but they are biologically connected to the same broader theme: **cell division, septum placement, envelope biogenesis, peptidoglycan remodeling, and stress-controlled growth**.

### Contexts related through toxin–antitoxin, Fic-like stress arrest, DNA damage, or defense modules

These are related to the **Fic/toxin-stress-growth arrest** part of the earlier interpretation:

**38, 70, 71, 89, 93, 108, 122, 123, 132, 158, 160, 165, 218, 230, 237, 239, 240**

Most relevant among these:

| Context | Relevant signal |
|---|---|
| 93 | **Fic family protein**, antitoxin VbhA, RelE/ParE toxin — very strong |
| 38 | RelE/ParE toxin + Phd/YefM antitoxin |
| 70 | LexA, VapC toxin, stress/DNA-damage-associated neighborhood |
| 71 | Phd/YefM antitoxin + Txe/YoeB toxin |
| 108 | DarT toxin-like ADP-ribosyltransferase + CRISPR-associated genes |
| 123 | VapC toxin family PIN ribonuclease |
| 132 | HicA/HicB toxin-antitoxin system |
| 160 | death-on-curing toxin |
| 218 | HicA/HicB toxin-antitoxin system |
| 230 | LexA + VapC toxin + SpoIIE phosphatase |
| 239 | PemK/MazF toxin |
| 240 | Phd/YefM antitoxin + YoeB/Txe toxin |

### Best overall answer

The contexts I would call related are:

**9, 38, 49, 51, 64, 70, 71, 73, 75, 77, 84, 89, 93, 98, 101, 103, 106, 108, 118, 122, 123, 125, 128, 129, 132, 139, 143, 147, 148, 153, 158, 160, 165, 166, 167, 173, 182, 186, 191, 196, 207, 209, 210, 217, 218, 230, 237, 238, 239, 240, 245, 246, 250, 251, 252, 256**

Among these, the **highest-confidence biologically related contexts** are:

**49, 51, 64, 77, 93, 98, 101, 118, 125, 166, 186, 191, 196, 209, 217, 238, 245, 251**

And the single best match is:

**217**

Supplementary File S18. ChatGPT-assisted analysis of functional relations of minD, TadZ-like proteins to ATPases (A) and P-loop NTPases (B).

**A**

Question to ChatGPT 5.5:

Are the septum site-determining protein minD, TadZ-like proteins biologically/functionally related to ATPases?

ChatGPT 5.5 answer:

I'll ground this in current protein-family evidence rather than relying on names alone, because "TadZ-like" can refer to several related PilM/ParA/MinD-family ATPase-like proteins.

Thought for 8s

Yes — **biologically/functionally, they are related to ATPases**, especially to the **ParA/MinD family of P-loop ATPases**.

**MinD** is directly an ATPase. It is a bacterial cell-division positioning protein whose ATP-bound state controls membrane association, MinC recruitment, and division-site placement. Structural work describes MinD as a membrane-associated ATPase, and reviews place MinD within the broader **ParA/MinD ATPase family** involved in spatial positioning inside bacterial cells.

**TadZ-like proteins** are also related to that same family, but with an important nuance: many TadZ proteins are **ATPase-like ParA/MinD-family proteins**, often involved in **polar localization or assembly of Tad/Flp pili**, rather than being strong motor ATPases themselves. A TadZ structure study found an atypical ATPase domain that binds ATP but has low ATPase activity, and a 2025 Tad pilus paper describes TadZ as belonging to the ParA/MinD family of ATPases implicated in polar localization of macromolecular assemblies.

So the relationship is:

**MinD** → bona fide ParA/MinD-family ATPase involved in septum/division-site positioning.

**TadZ-like proteins** → ParA/MinD-family ATPase-like proteins, commonly linked to pilus localization/assembly; may bind ATP and may have weak or regulated hydrolysis.

**TadA**, by contrast, is often the main cytoplasmic ATPase that powers Tad pilus extension/retraction, so TadZ should not always be interpreted as the principal energy-generating ATPase of the Tad system.

Bottom line: **yes, they are functionally and evolutionarily connected to ATPases through the ParA/MinD P-loop NTPase/ATPase superfamily, but TadZ-like proteins may be better described as ATPase-like spatial-positioning factors unless ATP hydrolysis has been demonstrated for the specific protein.**

**B**

Question to ChatGPT 5.5:

Are the septum site-determining protein minD, TadZ-like proteins biologically/functionally related to P-loop NTPases?

ChatGPT 5.5 answer:

Thought for 4s

Yes. **MinD and TadZ-like proteins are biologically related to P-loop NTPases**, specifically the **ParA/MinD branch of P-loop ATPases/NTPases**.

**MinD** is the clearest case: it is a septum site–determining protein and an ATPase required for correct cell-division site placement. UniProt classifies *E. coli* MinD in the **ParA family, MinD subfamily**, and structural studies describe nucleotide binding/hydrolysis features characteristic of P-loop proteins.

**TadZ-like proteins** are also related, but often in a more "ATPase-like" or specialized way. TadZ has been described as belonging to the **ParA/MinD family of ATPases**, involved in polar localization of Tad/Flp pilus systems; structural work notes that TadZ family proteins contain P-loop/Walker-A regions, though some have atypical Walker-A motifs and weak or unusual ATPase activity.

So, in annotation terms:

| Protein | Relationship to P-loop NTPases | Functional theme |
|---|---|---|
| **MinD** | Direct member of ParA/MinD P-loop ATPase family | Septum/division-site positioning |
| **TadZ-like** | ParA/MinD-family ATPase-like P-loop NTPase | Polar localization/assembly of Tad pili or related complexes |

Bottom line: **yes — both are connected to P-loop NTPases through the ParA/MinD P-loop ATPase lineage.** For TadZ-like proteins, it is safest to say **"P-loop NTPase/ATPase-like ParA-MinD family protein"** unless ATP hydrolysis has been experimentally shown for the exact protein.

Supplementary File S19. ChatGPT-assisted analysis of functional relations of CpaE/TadZ family proteins to septum site-determining proteins Ssd.

Question to ChatGPT 5.5:

Tell me if the CpaE/TadZ family proteins are biologically/functionally related to septum site-determining proteins Ssd?

ChatGPT 5.5 answer:

I'll check current primary/curated sources for both protein families, then separate strong evidence from weaker domain/annotation similarity.

Thought for 13s

Yes — **they are biologically related at the broad protein-family/mechanistic level**, but **not functionally equivalent in the narrow sense**.

**Best interpretation:** CpaE/TadZ proteins and Ssd septum site-determining proteins are both members or close relatives of the broader **ParA/MinD / Walker-type ATPase spatial-regulation superfamily**, which bacteria use to position cellular structures. But they are assigned to different cellular modules:

| Protein family | Main biological role | Shared relationship |
|---|---|---|
| **CpaE/TadZ** | Positions/assembles **Tad/Flp type IV pilus systems**, often at the cell pole | MinD/ParA-like ATPase; spatial localization protein |

| Protein family | Main biological role | Shared relationship |
| --- | --- | --- |
| **Ssd / MinD-like septum site-determining proteins** | Regulates **division-site placement / septation**, often through FtsZ/septum effects | MinD-like ATPase involved in spatial regulation |

The key link is that **TadZ/CpaE is described as a MinD/ParA-like ATPase** in the tad pilus locus, and recent reviews explicitly group TadZ/CpaE with ParA/MinD spatial-positioning systems. TadZ/CpaE localizes Tad pilus machinery to the cell pole and is required for secretion/pilus function.

Ssd, especially the *Mycobacterium tuberculosis* Rv3660c protein, was identified as an ortholog of actinobacterial septum-regulating proteins; overexpression inhibits septum formation and causes elongated, septum-deficient cells. The same work notes alignment/relationship to MinD-like septum site proteins.

So the clean answer is:

**Related?** Yes, likely through the **ParA/MinD ATPase spatial-positioning lineage**.
**Same function?** No. **CpaE/TadZ positions pilus/secretion machinery**, while **Ssd/MinD-like proteins regulate septum placement or cell division**.
**Functional analogy?** Yes: both help position cellular structures in bacteria using related ATPase-based spatial-regulation logic.
**Direct orthology?** Usually no, unless a specific annotation calls a particular protein "Ssd/MinD @ possible CpaE," which should be treated as a broad MinD/ParA-family annotation rather than proof of identical biological role. One MTB systems annotation does label Rv3660c as "Septum site-determining protein MinD @ possible CpaE," which supports family-level ambiguity but not interchangeable function.
